# Lack of adoption of computational gene-panel-selection tools in targeted spatial transcriptomics

**DOI:** 10.64898/2026.10.06.757034

**Authors:** Ceyhun Alar, Lorenzo Venturelli, Jelle Jacobs, Alejandro Sifrim

## Abstract

Targeted spatial transcriptomics requires selecting a gene panel before any experiment is run. Choosing well across tens of thousands of candidate genes under platform-specific probe, panel-size, and sensitivity constraints is difficult to do manually, and impossible to correct once a panel has been used. A growing number of computational tools to automate this selection have been designed, yet it remains unclear whether they are adopted in practice. Several factors could limit uptake: validation uncertainty about panel performance, workflow friction between a ranked gene list and a submission-ready panel, limited platform awareness in tools developed for dissociated single-cell data, and the cost advantage of vendor panels. We analyzed 18 computational tools for gene-panel selection from two angles: their citation landscape and their use across a targeted spatial transcriptomics corpus. Across 540 full-text-classified records, only 35 (6.5%) reported using the cited tool for panel design; most came from computational, benchmark, review, or non-spatial papers, or papers that cited but did not use it. In an independent corpus, only 34 of 479 (7.1%) involved a computational tool, alone or in a hybrid workflow; most used manually curated, premade vendor, or modified vendor panels. Both lines of evidence point the same way: computational selection is rarely part of the reported experimental workflow, and it more often supplemented manual curation than replaced it. Adoption is unlikely to turn on further benchmark gains alone; it will depend on reducing the validation uncertainty and workflow friction that keep users on established panel-design practice.

## Introduction

Targeted spatial transcriptomics now spans imaging- and sequencing-based platforms including MERFISH (1), seqFISH(2), HCR-FISH (3), Xenium (4), MERSCOPE, and CARTANA (5). The first design decision in each targeted experiment is which genes the panel will measure, and that choice constrains further downstream analysis: cell-type annotation, rare-cell-state detection, and the biological claims the experiment can support. Vendor-designed panels are commercially available, but purpose-built panels have been shown to outperform them: in a comparison of expert, vendor, and algorithmic strategies, computational panels performed well for major cell-type discrimination, while expert and computationally derived panels retained advantages for minor populations and feature diversity (6). Adding genes does not make panel composition matter less: broader-coverage panels trade per-gene sensitivity for breadth and still perform worse downstream than smaller targeted panels, so the choice of genes continues to drive measurement quality (7).

Computational methods have been developed to automate this design step, and they fall into two groups that behave differently in the analysis that follows. One group is spatially motivated: these tools were developed for, or explicitly applied to, targeted spatial transcriptomics panel design, and several also encode platform-aware constraints such as probe feasibility, panel size, and detection sensitivity: Spapros (8), gpsFISH (9), PERSIST (10), scGIST (11), ReconST (12), scGPD (13), geneBasis (14), panelScope(6), geneCover (15), and scMAGS (16). The other group was developed for marker or feature selection in dissociated single-cell RNA-seq: COSG (17), SCMER (18), scGeneFit (19), RankCorr (20), NS-Forest v2 (21) and v4.0 (22), scPNMF (23), and SMaSH (24). Together these make up the 18 included tools summarized in Table 1.

A larger collection of tools does not by itself establish that those tools currently shape experimental practice. Citation counts are difficult to interpret in this setting: a panel-selection tool can be cited by a new computational method, a benchmark, a review, a non-spatial single-cell study, or an experimental spatial paper that mentions the tool without using it. Hence citation alone are not very informative. Spatial transcriptomics is a commercialized assay with high per-experiment costs mainly constituted by the necessary panel-specific detection probe kits, and vendor-designed panels are priced well below custom panels of similar size, so the design choice carries a cost as well as an analytical consequence. Panel design is where computational optimization has the most room to change what an experiment can resolve. If adoption of the published tools is in fact low, that matters for the developers building them, the funders supporting them, and the experimentalists trying to choose among them.

This paper asks a narrow empirical question: are computational gene panel selection tools for targeted spatial transcriptomics being used in experimental studies? We approach this question from two sides. First, we assemble a corpus of citing papers for the 18 tools and classify how each citing paper engages with the cited tool, distinguishing actual experimental use from citation without reported use, computational benchmarking, other computational work, reviews, non-spatial work, and other categories. Second, we assemble a corpus of targeted spatial transcriptomics papers up until April 2026 and classify how each reported panel was designed, including computational tools, hybrid computational/manual workflows, manual curation, vendor panels, vendor panels with manual modification, and cases with insufficient evidence. Together, these two analyses estimate adoption from both the tool side and the field side and show that computational tools are reported for fewer than one in ten panel design records. The paper does not benchmark algorithm quality or rank panel selection methods; instead, it surveys the uptake of gene panel selection tools in this field as we see commercial maturation of spatial transcriptomics technologies.

## Methods

### Tool set

We defined a canonical set of computational gene-panel selection tools before running any citation or corpus analyses. A tool was eligible if it described an explicit computational procedure for selecting genes, markers, or panels, and if the method was plausibly relevant to targeted spatial transcriptomics panel design. We included tools built specifically for spatial panels and general marker-selection methods that have been applied to spatial or targeted-panel contexts. We excluded downstream spatial analysis methods, segmentation tools, probe and barcode design utilities with no gene-selection step, differential-expression workflows with no distinct panel-selection method, and papers that only described experimental platforms. The resulting set contains 18 tools: COSG, geneBasis, geneCover, gpsFISH, NS-Forest v2, NS-Forest v4.0, panelScope, PERSIST, RankCorr, ReconST, SCMER, SMaSH, Spapros, scGeneFit, scGIST, scGPD, scMAGS, and scPNMF. For each tool we recorded the published DOI where available, the preprint DOI, the preprint date, and a category label (spatially-motivated tool or general marker-selection tool). Full bibliographic details are reported in Supplementary Table S1.

#### Citation harvest

For each tool, we queried citing papers against both the published DOI and the preprint DOI where both existed. The full DOI manifest and the OpenAlex(25) and Semantic Scholar(26) query specifications are provided in Supplementary Methods S1. Citation records were harvested from OpenAlex and Semantic Scholar and deduplicated within each tool by normalized DOI, or by source-specific identifier for DOI-less records. The harvest was frozen on 17 April 2026 and then cleaned for same-tool preprint/published duplicates and known false-positive citation mappings. Three additional tool-paper records, corresponding to two unique papers, were added during cross-validation against the spatial corpus analysis: both papers explicitly named one of the 18 tools in their methods text but did not formally cite the tool’s published or preprint DOI, so neither OpenAlex nor Semantic Scholar had surfaced them as citing papers. We added these three records to the citation-screening flow before classification. A paper that cited multiple tools contributed one tool-paper record per cited tool, but only one record to the unique-paper count.

#### Citation classification

We assigned each tool-paper record one of seven mutually exclusive labels capturing how the citing paper engaged with the cited tool. **EXP-USED** is an experimental spatial transcriptomics paper that reported using the cited tool to select or design genes for its own panel. **EXP-CITED** is an experimental spatial transcriptomics paper that cited the tool without reporting that it used it for panel design. **COMP-BENCH** is a computational benchmarking or comparison paper that ran the tool as part of a benchmark. **COMP** covers computational methods or analysis papers that cited the tool without being an experimental deployment or benchmark. **REV** covers reviews, perspectives, protocol overviews, and other literature-synthesis articles. **NON-ST** covers papers outside spatial transcriptomics. **OTHER** is the residual category for corrections, non-research items, and otherwise ineligible records.

Classification ran in stages. A pre-filter resolved records that could be typed without full-text review (86/626): 77 by deterministic rules, labelled REV or OTHER when their OpenAlex work type or Semantic Scholar publication types identified them as reviews, book chapters, dissertations, or clinical trials, or when the title began with a correction/erratum/corrigendum/retraction or editorial/special-issue marker, and 9 by manual review (exact fields and patterns in Supplementary Methods S2.1). The remaining 540/626 entered a second stage: PDFs were retrieved where possible, parsed with PyMuPDF V1.27 (27), and passed to a local Gemma4 (28) model (31B, released 2 April 2026) running through Ollama v0.20.7 (29). The Gemma4 prompt supplied the target tool name, aliases, and known DOI variants, and requested document type, explicit tool-use evidence, paper profile, spatial-transcriptomics status, confidence, and supporting verbatim quotes. The full prompt template is provided in Supplementary Methods S2. Final citation labels were produced by an independent GPT-5.4 audit (release 5 March 2026) (30) that reviewed and reconciled the Gemma4 classifications. Record routing through the full pipeline is summarized in **Supplementary Fig. S1**. To quantify classifier reliability against human judgement, a human annotator validated every EXP-USED record, the positive class of primary interest, against the full text, and the remaining labels (EXP-CITED, COMP-BENCH, COMP, REV, NON-ST, OTHER) were checked on a stratified random sample of 20 records per label.

### Targeted spatial transcriptomics corpus

To assess panel-design strategy in the broader targeted spatial transcriptomics literature, independent of whether a paper cited any of the 18 tools, we harvested a topic corpus from Europe PMC (31), OpenAlex, Scopus (32), and Semantic Scholar. Search terms covered the major targeted transcriptomics platforms and method families: MERFISH, seqFISH, Xenium, MERSCOPE, HCR-FISH, CosMx (33), Molecular Cartography (34), STARmap (35), osmFISH (36), HybISS (37), BaristaSeq (38), SCRINSHOT (39), EASI-FISH (40), ExSeq (41), SABER-FISH (42), in situ sequencing, targeted spatial, and in situ transcriptomics. The full query strings for each database are provided in Supplementary Methods S3. The query window covered 2015 through a publication cutoff of 31 March 2026. Database records were then screened to remove non-primary research and off-topic material. Exclusions covered reviews, editorials, letters, errata, retractions, datasets, book chapters, protocols, conference abstracts, low-information supplement records, blocked non-paper DOI prefixes, non-English records, records published after the freeze date, and a curated list of manual exclusions. Papers that cite one of the 18 tools might not be retrieved by a topic search alone. To make sure none were missed, we added the citing papers with a resolvable DOI to the corpus during DOI deduplication, before panel-design classification. Deduplication ran in two stages. DOI-bearing records were collapsed first, with a canonical record retained per DOI by source priority: Europe PMC, OpenAlex, Scopus, Semantic Scholar. Source labels were carried through, so multi-source discovery was preserved. Remaining duplicates and preprint/published variants were resolved by fuzzy title matching. Candidate pairs were scored on normalized title similarity, author overlap, and record status. Pairs above the automatic-merge threshold were collapsed; ambiguous pairs were resolved manually. Stage-by-stage record counts are reported in **Supplementary Fig. S2**.

### Panel-design classification

We classified each corpus record by the reported source of its targeted gene panel, using six final labels. **TOOL** is a paper in which one of the 18 canonical tools explicitly selected the final panel. **HYBRID-TOOL** is a paper in which one of the 18 tools contributed to panel selection but manual, vendor, or other additions also shaped the final panel. **MANUAL-PANEL** covers panels curated from biological markers, differential expression, ranking output, pathway lists, literature knowledge, or similar non-canonical-tool logic. **VENDOR** is a fixed commercial or catalog panel. **VENDOR-MANUAL** is a vendor panel that was manually modified, extended, or adapted. **FILTERED** covers records that did not contain targeted panel-design evidence for this analysis.

Initial panel-design classification used the same PDF-first local Gemma4 workflow as citation screening, with the prompt adapted to request document type, panel-design approach, experimental/computational profile, spatial-transcriptomics status, targeted-panel status, evidence, and confidence. The full prompt template is provided in Supplementary Methods S**4**. To prevent inflated estimates of computational tool adoption, the classifier treated differential-expression lists, marker ranking output, mutual-information heuristics, pathway curation, and literature-based gene lists as MANUAL-PANEL unless one of the 18 canonical tools explicitly selected the final panel. Probe, barcode, oligo, primer, decoding, segmentation, registration, and downstream analysis tools were not counted as gene-panel selection tools. Final panel-design labels were produced by an independent GPT-5.4 that reviewed and reconciled the Gemma4 classifications. Audit rows from earlier harvest iterations that fall outside the deduplicated corpus after manual exclusions were not used in our analyses. For the Fig. 2 analytic set, records using untargeted spatial platforms (e.g., Visium, Slide-seq) or otherwise outside scope were excluded. Platform-stratified summaries use paper-platform records, so a paper reporting more than one platform can contribute to more than one platform count. As with citation classification, a human annotator validated every TOOL and HYBRID-TOOL call, the positive classes that drive our estimates of computational-tool adoption, against the source full text. The remaining panel-design labels (MANUAL-PANEL, VENDOR, VENDOR-MANUAL, FILTERED) were checked on a stratified random sample of 20 records per label.

## Results

### Citation composition across the tools

The 540 full-text-classified records correspond to 403 unique citing papers and cover 15 of the 18 included tools. Of the 540 only 35 (6.5%; Wilson 95% CI, 4.7–8.9%) were classified as EXP-USED, indicating that the citing paper reported using the cited tool to select or design genes for its own targeted spatial transcriptomics panel ***(Fig. 1a-b)***. The remaining records resolved as 52 (9.6%) EXP-CITED, 98 (18.1%) COMP-BENCH, 181 (33.5%) COMP, 36 (6.7%) REV, 108 (20.0%) NON-ST, and 30 (5.6%) OTHER. Citations from computational papers (COMP plus COMP-BENCH; 279 records) outnumbered those from experimental papers (EXP-USED plus EXP-CITED; 87 records) by approximately 3.2:1. Most citing papers cited a single tool (342 of 403; 84.9%). Of the 61 multi-tool citing papers, 37 (61%) were benchmarks or reviews (32 benchmarks, 5 reviews), and a further 12 (20%) were other computational papers.

**Figure 1.**
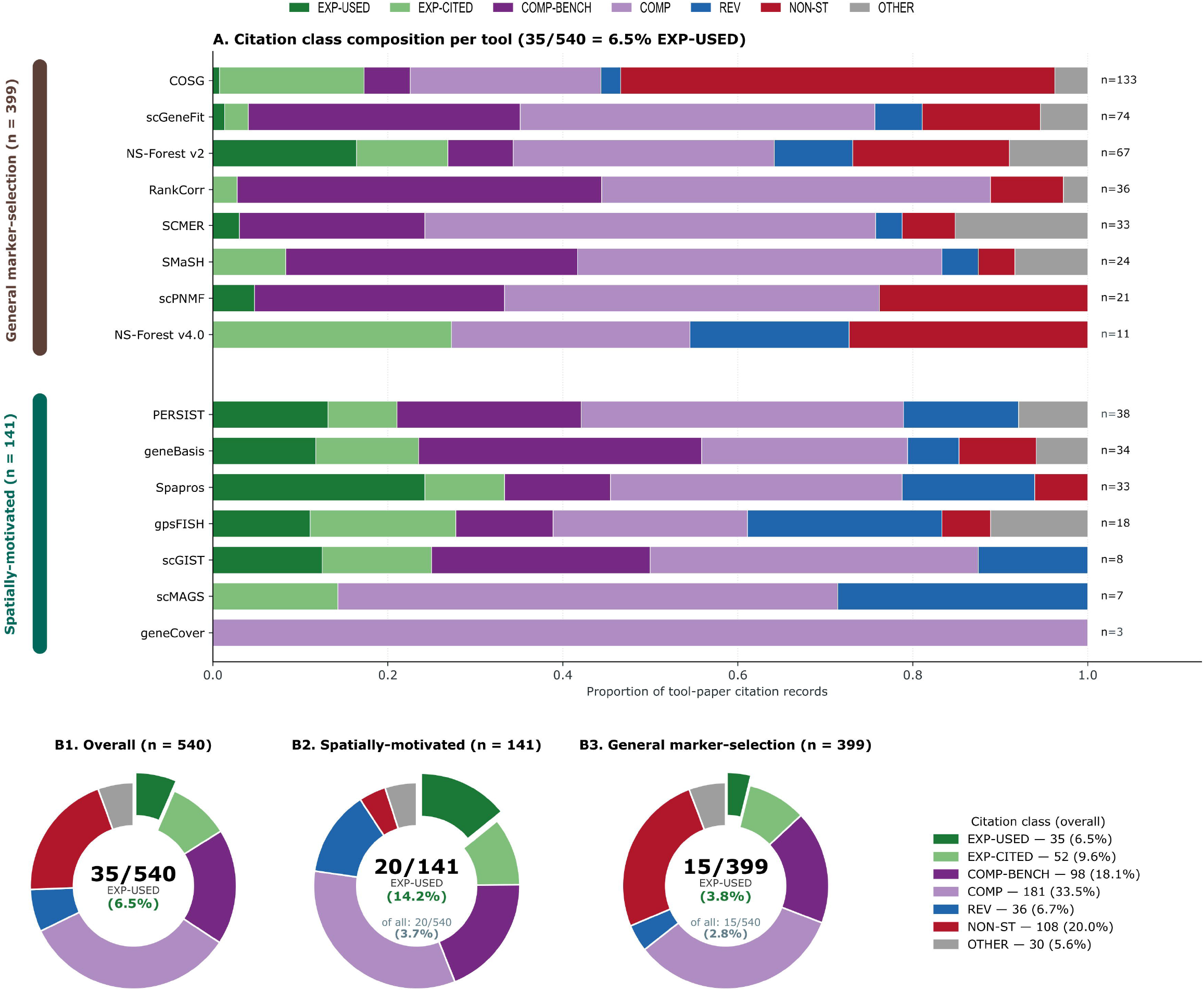
Citations to panel-selection tools rarely represent experimental use. (A) Per-tool composition of the 540 full-text-classified records, shown as 100% stacked bars for the 15 tools with at least one citing paper. Tools are arranged in two blocks (general marker-selection, top; spatially-motivated, bottom) and ordered by citation volume within each block. EXP-USED records (dark green) form the small left-most slice for most tools. (B) Donut charts comparing the EXP-USED fraction across all records (35/540, 6.5%), spatially-motivated tools (20/141, 14.2%; 3.7% of all records), and general marker-selection tools (15/399, 3.8%; 2.8% of all records). Citation classes: EXP-USED, the citing paper used the tool to design a panel; EXP-CITED, an experimental paper cited the tool without using it; COMP-BENCH, computational benchmark or comparison; COMP, other computational use; REV, review or perspective; NON-ST, non-spatial-transcriptomics paper; OTHER, residual ineligible records.

Spatially-motivated tools accounted for 141 of 540 (26.1%), of which 20 (14.2% within the group; 3.7% of all tool-paper records) were EXP-USED, a higher rate than the 15 of 399 (3.8%) for general marker-selection tools despite their lower citation volume (***Fig. 1b***). Within the spatially-motivated group, EXP-USED shares were highest for Spapros (8 of 33; 24.2%), PERSIST (5 of 38; 13.2%), scGIST (1 of 8; 12.5%), geneBasis (4 of 34; 11.8%), and gpsFISH (2 of 18; 11.1%) ***(Fig. 1a)***. Among the highest-volume general-marker tools the pattern was mixed: COSG accumulated 133 records but only 1 (0.8%) was EXP-USED and scGeneFit 74 records with 1 (1.4%), whereas NS-Forest v2 (67 records) was EXP-USED in 11 (16.4%), more EXP-USED records than any other tool in either group **(Fig. 1a)**. The two groups also differed in non-spatial citation drift: 6 of 141 (4.3%) were NON-ST, compared with 102 of 399 (25.6%).

### Computational tools are absent from most reported panels

The deduplicated targeted spatial transcriptomics corpus contained 1,646 records. Out of these 479 records contained a clear description of how the targeted panel was selected or sourced. Of these, 34 (7.1%; Wilson 95% CI, 5.1–9.8%) involved one of the 18 computational tools — 7 (1.5%) classified as TOOL and 27 (5.6%) as HYBRID-TOOL (**Fig. 2a)**. The remaining 445 in-scope records used non-computational strategies: 301 records (62.8%) were MANUAL-PANEL, 96 (20.0%) were VENDOR, and 48 (10.0%) were VENDOR-MANUAL **(Fig. 2a)**.

**Figure 2.**
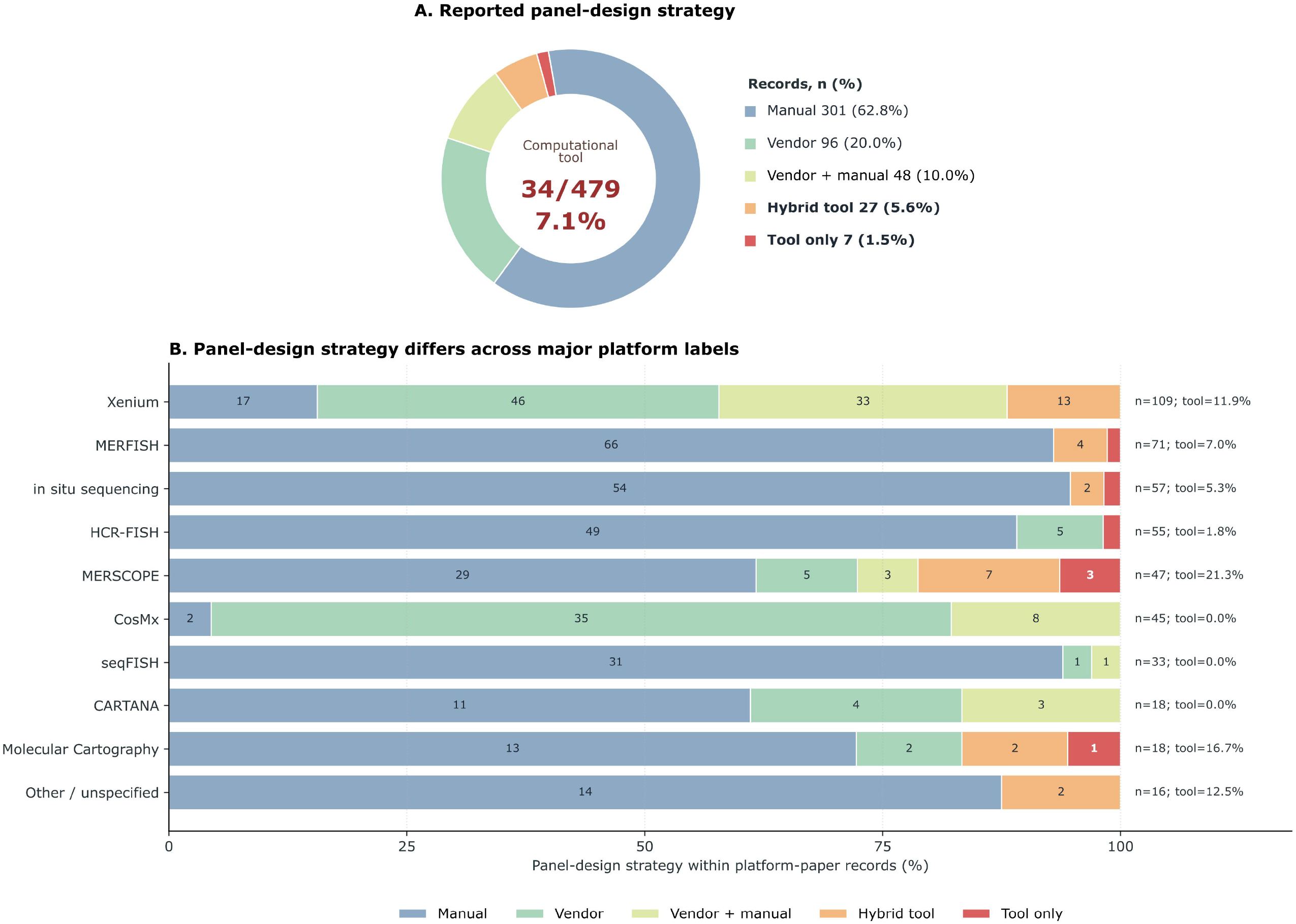
Computational tools are rarely reported in targeted spatial panel design. (A) Overall panel-design strategy across the 479 in-scope targeted spatial transcriptomics records. Donut wedges show MANUAL-PANEL (n = 301, 62.8%), VENDOR (n = 96, 20.0%), VENDOR-MANUAL (n = 48, 10.0%), HYBRID-TOOL (n = 27, 5.6%), and TOOL (n = 7, 1.5%); computational tool adoption (TOOL plus HYBRID-TOOL) was 34/479 (7.1%; Wilson 95% CI, 5.1–9.8%). (B) Within-platform panel-design strategy, shown as 100% stacked bars per paper-platform record using the same five categories as (A); a paper that reports multiple platforms contributes to each. Panel-design classes: TOOL, a computational tool selected the final panel; HYBRID-TOOL, a computational tool was combined with manual or vendor input; MANUAL-PANEL, manually curated panel; VENDOR, premade vendor panel; VENDOR-MANUAL, vendor panel with manual modification.

The reported design source of the chosen panel differed by platform **(Fig. 2b)**. Among Xenium paper-platform records, 79 of 109 (72.5%) were vendor-associated (46 VENDOR and 33 VENDOR-MANUAL). Among MERSCOPE records, 8 of 47 (17.0%) were vendor-associated (5 VENDOR and 3 VENDOR-MANUAL), while 29 of 47 (61.7%) were MANUAL-PANEL. CosMx records were almost entirely vendor-associated (43 of 45; 95.6%) with no TOOL or HYBRID-TOOL appearances, while seqFISH (33 records) and CARTANA (18 records) likewise showed no computational tool appearances. MERSCOPE had the highest tool fraction at 10 of 47 records (21.3%).

Tool appearances in the corpus-level analysis were concentrated. NS-Forest v2 appeared 11 times across TOOL and HYBRID-TOOL records, followed by Spapros (8), PERSIST (5), and geneBasis (4). The remaining seven appearances came from six other tools: gpsFISH (twice), and scGeneFit, COSG, SCMER, scGIST, and scPNMF (once each). Eight tools — RankCorr, scMAGS, NS-Forest v4.0, geneCover, SMaSH, panelScope, scGPD, and ReconST — had no TOOL or HYBRID-TOOL appearances in the current corpus, despite together accounting for 138 tool-paper citation records in the citation analysis.

### Citation-level and corpus-level adoption estimates

The two analyses count different things but converge on the same set of papers **(Fig. 1, Fig. 2a)**. The citation analysis counts tool-paper records: a paper contributes one record per tool it cites, so a benchmark citing three tools contributes three records, producing 540 records across 18 tools. The corpus analysis counts papers, identifying 479 with adjudicable panel-design evidence regardless of whether they cited any of the 18 tools. The 35 EXP-USED tool-paper records in the citation analysis correspond to 34 unique papers; one paper reported using and citing two canonical tools (NS-Forest v2 and Spapros), contributing two records. All 34 papers appear in the deduplicated corpus, and all 34 receive a TOOL or HYBRID-TOOL label in the panel-design audit: 7 TOOL and 27 HYBRID-TOOL. The same four tools dominate both rankings, in the same order: NS-Forest v2 (11 EXP-USED records, 11 corpus appearances), Spapros (8 and 8), PERSIST (5 and 5), and geneBasis (4 and 4). The two analyses approach the question from opposite directions. One starts with the tools and surveys their usage; the other starts with the ST literature and asks how panels are primarily designed. They return the same answer: computational panel-selection methods are rarely used as part of the experimental panel-design process.

## Discussion

Panel design is not a neutral upstream choice: different panels yield materially different biological readouts on the same tissue (7). The relevance of panel design is itself shifting as full-transcriptome spatial technologies narrow the regime in which targeted panels are strictly necessary (43). Even so, well-designed smaller panels remain attractive on cost and throughput grounds and stay the practical choice for many tissues and questions. Such panels may be premade by a vendor, curated manually leveraging biological expertise, optimized computationally, or built as a mosaic of these. The latter approaches theoretically provide panels tailored to capture variability in specific biological conditions in an unbiased, data-driven manner, reducing redundancy to increase experimental power.

In this study we meta-analyzed the adoption of such tools by the community using two alternative analysis strategies that returned similarly low estimates: fewer than one in ten panel-design records involved a computational tool **(Fig. 1, Fig. 2a)**. These trends seem to be platform-specific (**Fig. 2b)**. Xenium records in our analytic set were heavily vendor-associated. General adoption of the same panel by the community has its benefits and limitations. It allows for easy comparison between studies and enhances reproducibility. But if vendor panels have systematic biases in a tissue, disease state, or cell-state resolution task, that dependence could propagate the same design constraints across many studies. MERSCOPE showed the opposite pattern, dominated by manually curated panels. This marks a setting where computational panel-selection tools could be especially useful.

Several mechanisms could explain the low adoption rates. One mechanism is validation uncertainty. A computationally selected panel is useful only to the extent that someone can judge whether it will perform well in a particular tissue, disease setting, cell-type hierarchy, and biological question. Many tools report internal metrics such as classification accuracy, reconstruction error, marker specificity, or reference-data coverage, but these are computed on dissociated single-cell reference data. Strong performance there does not guarantee that the same genes will behave the same way as probes in tissue: the expression profiles, dynamic range differences, and cell-type separation seen in single-cell data may not transfer to a spatial assay, where platform-specific sensitivity, tissue autofluorescence, probe constraints, optical crowding, and segmentation errors all intervene between the selected gene list and the measured signal. Part of this gap stems from how these tools were built: many general marker-selection tools were built for dissociated single-cell RNA-seq, where they identify discriminative genes well but do not address probe design, detection sensitivity, limited dynamic range, tissue-specific background, or the preservation of spatially patterned cell states. The higher experimental-use fraction for spatially motivated tools than for general marker-selection tools (14.2% vs. 3.8%) is consistent with this account, though our data cannot establish causality. As more tools are built around these platform constraints, the translation from gene list to trustworthy ST panel should become easier, but at present the burden of judging assay-readiness still falls on the user.

Recent benchmarking efforts rank panels against one another, but neither computational selection nor manual curation has been validated against tissue-specific assay endpoints. Manual curation rests on expert intuition, prior literature, and informal heuristics that have not themselves been systematically tested against those endpoints. Expert curation may miss genes outside a curator’s familiar terrain just as algorithmic selection may miss well-described context-specific markers. Treating expert judgment as a more trustworthy standard than quantitative selection is field convention rather than evidence of differential validation. A symmetric view suggests that low computational adoption is partly a status-quo effect. Manual curation is the established practice, so its limitations are accepted as familiar, while the limitations of computational tools draw scrutiny precisely because they are new.

Computational tools rarely produce a finished, orderable panel. This workflow friction is a second mechanism limiting adoption, since bridging this gap adds work instead of saving it. Many computational tools produce ranked genes or candidate panels, while experimental workflows need a finished panel design that a vendor or core facility can manufacture, requiring probe-feasibility checks, the addition of positive and negative controls, housekeeping genes, and biology-driven additions. These steps require additional effort and time, after the generation of a gene list. A predesigned vendor panel alleviates that necessity, even when a computational tool could have provided improved designs.

A third mechanism is that visibility can be misleading. Several tools accumulated many citations from computational papers, benchmarks, reviews, or non-spatial applications. These citations are legitimate scholarly uses, but reflect citations of the computational tool development community, conflating apparent impact if treated as evidence of experimental adoption.

A fourth mechanism is cost. On both leading targeted-spatial platforms, vendor panels are priced approximately an order of magnitude below a fully customized panel of comparable size on a per-gene basis. Their fixed design and validation costs are spread across all the vendor’s customers rather than re-incurred for a single study. Both vendors also offer the possibility to add a small number of custom genes to a validated panel. The per-gene cost of custom content is highest when a panel is small and run on few samples, and drops only as that fixed cost is divided across more genes and reactions. These economics favour vendor-defined content and small, incremental customisation. In the current situation, optimizing an entire custom panel offers the least marginal benefit per unit cost.

Our analysis indicates that computational involvement was more often a hybrid approach, combining manual curation with data-driven approaches, compared to purely automated selection. Automation may not completely replace expert curation. A more realistic adoption pathway is for computational methods to be useful within expert-led workflows, where they can expose trade-offs, guarantee cell-state coverage, and identify candidate genes that manual curation might miss.

This study has several limitations. First, both classification tasks depend on what authors reported. Some authors may have used computational tools without naming them, relied on internal scripts not tied to a published method, or used vendor-facing design support that does not appear in the manuscript. The estimates therefore describe reported adoption and are best read as lower bounds: undisclosed tool use is counted as non-use, and the agreement between the two estimates shows they are consistent, not that they are free of a shared downward bias. If a panel was shaped by a computational method that goes undisclosed, readers cannot evaluate the design logic, reproduce the selection process, or compare the panel with other studies, so the adoption gap has consequences whether it reflects low actual use or low disclosure. Second, the corpus is a frozen snapshot through the March 2026 publication cutoff. Several of the tools are recent, and their citation and adoption patterns may change as they mature. In particular, ReconST, scGPD, and panelScope had no citing papers in the frozen window and should not be read as failed-adoption cases. They were retained in the tool set because they are part of the current methods ecosystem, but citation uptake cannot yet be meaningfully assessed for them. Third, we measured adoption rather than algorithm quality. A tool can be underused and still be scientifically strong; a manual or vendor panel can be widely used and still be suboptimal for a particular biological question. Our analysis does not rank tools, compare selected panels experimentally, or determine which panel-design strategy is best. It asks whether computational panel-selection tools are reported in the experimental workflows where they could plausibly matter. Fourth, citation and DOI tracking are imperfect. Preprint and published versions, duplicate records, incomplete metadata, and false-positive citation mappings can distort tool-level counts. Deduplication and audit steps reduce these errors, but residual errors are possible. Finally, the tool set is bound by design. We included 18 canonical computational gene-panel selection tools and excluded downstream spatial analysis methods, segmentation tools, probe and barcode design utilities with no gene-selection step, and generic differential-expression workflows. This boundary makes the classification reproducible but means the analysis does not capture every possible computational influence on panel design.

The lack of adoption of computationally optimized panel designs observed here is not a failure of individual tools or individual experimentalists. It reflects how computational biology often rewards method development on terms that do not match how experimental users adopt infrastructure(44–46). A method can be novel, peer-reviewed, benchmarked, and well-cited and still remain underused, because users default to established practice rather than change a functional workflow when the benefits are not easily predictable. For targeted spatial transcriptomics, panel design determines the outcome of an expensive physical assay, where missing genes cannot be recovered from the same experiment once a panel is ordered. If these tools are to be adopted, development will need to shift from improving benchmark performance toward easing the validation uncertainty and workflow friction that keep users on established practice.

## Supporting information

supplementary_data_s1_citation_classifications

supplementary_data_s2_panel_classifications

supplementary_evidence_dossier_citation

supplementary_evidence_dossier_panel

supplementary_methods_s1_doi_manifest_queries

supplementary_methods_s2_citation_prompts

supplementary_methods_s3_corpus_queries

supplementary_methods_s4_panel_prompts

## Supplementary figures

**Figure S1**. PRISMA flow for the citation harvest and citation classification. Source merge: 577 OpenAlex hits and 478 Semantic Scholar hits, minus 432 cross-source overlaps, gave 626 unique source-merge records. After filtering, 540 records were classified by Gemma4 full-text classification and an independent GPT-5.4 audit. Terminal labels were EXP-USED, 35; EXP-CITED, 52; COMP-BENCH, 98; COMP, 181; REV, 36; NON-ST, 108; OTHER, 30.

**Figure S2**. PRISMA flow for the targeted spatial transcriptomics corpus and panel-design classification. Identification yielded 5,820 records: 5,355 database records (Europe PMC, 1,039; OpenAlex, 2,020; Scopus, 626; Semantic Scholar, 1,670). Screening removed 1,531 database records, leaving 3,827; 460 unique citing papers with a DOI were then added at the deduplication stage, giving 4,287 records entering deduplication. Removing 2,641 duplicates produced a final deduplicated corpus of 1,646. Eligibility screening then classified 1,167 records as filtered and 479 as in-scope. The 479 in-scope records comprised MANUAL-PANEL, 301; VENDOR, 96; VENDOR-MANUAL, 48; HYBRID-TOOL, 27; and TOOL, 7. Computational tool adoption (TOOL plus HYBRID-TOOL) was 34/479 (7.1%).

## Tables

**Table 1**. Feature comparison of the 18 computational gene-panel-selection tools, one row per tool, grouped into spatially-motivated and general marker-selection categories and ordered by citation volume within each. Method columns cover each tool’s optimization objective, spatial-platform awareness, probe-design feasibility and panel-size guidance; adoption columns give citing records and the EXP-USED count and fraction over the full-text-classified set (n = 540), corpus appearances as sole (TOOL) or combined (HYBRID-TOOL) panel selector, and the primary-reference DOIs.

## Supplementary Methods

**Supplementary Methods S1**. Tool DOI manifest and citation-harvest query specifications. The canonical published and preprint DOIs for the 18 reviewed gene-panel-selection tools, the exact OpenAlex and Semantic Scholar queries used to harvest citing papers for each DOI, and the DOI-normalization and same-tool version-deduplication rules. Enables exact reproduction of the citation harvest (frozen 17 April 2026).

**Supplementary Methods S2**. Citation-classification prompts. The deterministic metadata/title pre-filter rules, the Gemma4 full-text classification prompts (document type, tool use, paper profile), and the GPT-5.4 audit instructions and structured-output schema used to assign each tool-paper citation record to one of seven classes. Of 626 records, 86 were resolved before full-text classification (77 by metadata, 9 by manual review) and the remaining 540 proceeded to Gemma4 classification.

**Supplementary Methods S3**. Topic-corpus query specifications. The full query strings and API-call parameters for the four databases (Europe PMC, OpenAlex, Scopus, Semantic Scholar) used to harvest the targeted spatial transcriptomics corpus, including per-backend field-scoping, pagination, and preprint handling.

**Supplementary Methods S4**. Panel-design classification prompts. The Gemma4 prompts (document type, panel-design approach, paper profile, panel scope) and the GPT-5.4 audit instructions and schema used to classify each corpus paper’s reported panel-design strategy as MANUAL-PANEL, VENDOR, VENDOR-MANUAL, HYBRID-TOOL, or TOOL.

## Supplementary Data

**Supplementary Data S1**. Citation-screening classifications (n = 540). One row per full-text-classified tool-paper citation record (the denominator for all citation-screening rates). Columns give the final class, cited tool, DOI, year, and title; the GPT-5.4 audit decision, evidence quote, and validation status; and traceability fields (Gemma4 and audit labels and confidences, reconciled class, error type, record keys, and PDF filename).

**Supplementary Data S2**. Panel-design classifications (n = 1,646). One row per in-corpus paper. Columns give the final panel-design class, DOI, year, and title; the audit decision and evidence quote; the derived imaging platform and canonical tool name(s); validation status; the Figure 2 exclusion flag (fig2_platform_excluded); and source/paper keys. The 479 in-scope records (non-FILTERED, fig2_platform_excluded = FALSE) underlie Figure 2.

## Key Points

Across 540 papers citing at least one of 18 computational gene-panel selection tools, only a small fraction reported using the cited tool for panel design.

In an independent corpus of targeted spatial transcriptomics studies, computational tools were used in only a small fraction of panels; most studies used manually curated or vendor panels.

The results show an adoption gap between the development and citation of computational panel-design tools and their use in real experiments.

Closing the gap will likely require clearer validation standards, easier workflow integration, and platform-aware probe constraints.

## Acknowledgements

We thank the developers of the gene-panel selection tools surveyed here for making their methods openly available; this analysis would not have been possible otherwise, and our intent is to support their continued development rather than diminish it. We also thank the maintainers of the open bibliographic resources this study drew on: OpenAlex, Europe PMC, and Semantic Scholar (Allen Institute for AI), and Scopus (Elsevier).

## Data Availability

All harvest, screening, classification, and audit code, together with the final tables and the scripts that generate the PRISMA figures, are available at https://github.com/sifrimlab/Alar-et-al. The two canonical machine-readable classification tables are provided as Supplementary Data S1 (citation-screening records, n = 540) and Supplementary Data S2 (panel-design audit rows, n = 1,646). Per-record evidence dossiers are provided as Additional File 1 (citation analysis) and Additional File 2 (panel-design analysis).

