## supplementary_evidence_dossier_citation for "Lack of adoption of computational gene-panel-selection tools in targeted spatial transcriptomics"

### Citation classification evidence dossier

*Per-record audit dossier for the 626 tool-paper citation records. For each record this document shows the final classification (EXP-USED / EXP-CITED / COMP-BENCH / COMP / REV / NON-ST / OTHER), the GPT-5.4 audit decision (AGREE / DISAGREE / NEEDS_MANUAL or MANUAL_DEMOTE / MANUAL_ADD where a post-audit edit applies), the verbatim PDF evidence quote used by the audit, the audit's reason, and the validation status. The dossier is the human-readable companion to Data S1 (GPT_citation_audit_FINAL.csv).*

### Summary

This dossier covers every tool-paper record in the frozen citation classification set after the 2026-05-21 reconciliation. Each row below is one tool-paper pair: a multi-tool paper appears once per cited tool. The underlying CSV is analysis/pipeline/results/tool_citations/citation_screening/gpt54_audit/GPT_citation_audit_FINAL.csv (Data S1 in the supplement).

**Total tool-paper records: 540**

**Breakdown by final classification:**

EXP-USED: 35 — Experimental spatial-transcriptomics paper that reported using the cited tool to select or design genes for its own panel.

EXP-CITED: 52 — Experimental spatial-transcriptomics paper that cited the tool without reporting that it used it for panel design.

COMP-BENCH: 98 — Computational benchmarking or comparison paper that ran the tool as part of a benchmark.

COMP: 181 — Computational methods or analysis paper that cited the tool without being an experimental deployment or benchmark.

REV: 36 — Review, perspective, protocol overview, or other literature-synthesis article.

NON-ST: 108 — Paper outside spatial transcriptomics.

OTHER: 30 — Residual category: corrections, non-research items, or otherwise ineligible records.

**Breakdown by tool:**

COSG: 133

scGeneFit: 74

NS-Forest v2: 67

PERSIST: 38

RankCorr: 36

geneBasis: 34

SCMER: 33

Spapros: 33

SMaSH: 24

scPNMF: 21

gpsFISH: 18

NS-Forest v4.0: 11

scGIST: 8

scMAGS: 7

geneCover: 3

### EXP-USED (n = 35)

*Experimental spatial-transcriptomics paper that reported using the cited tool to select or design genes for its own panel.*

#### NS-Forest v2 (11)

| DOI | Year | Title | Audit decision | Evidence (verbatim quote) | Audit reason / validation status |
|---|---|---|---|---|---|
| 10.1016/j.cell.2024.02.030 | 2024 | Cellular architecture of evolving neuroinflammatory lesions and multiple sclerosis pathology | DISAGREE | This selection was made manually, based on the literature and single-cell datasets ... in combination with different computational frameworks.97,98 [Ref. 97 = NS-Forest v2]. | Treat this as EXP-USED for consistency with the custom-panel ISS/Xenium cases where the panel was selected manually in combination with computational frameworks and the cited framework reference resolves to NS-Forest v2. [accepted_disagree] |
| 10.1038/s41586-024-07171-z | 2024 | Spatially organized cellular communities form the developing human heart | AGREE | "These 238 target genes ... Using MERFISH-encoding probes designed for these selected genes, we performed MERFISH studies ..." | The evidence explicitly links NS-Forest-selected genes to MERFISH probe design for this study. [accepted_agree] |
| 10.1038/s41586-024-08030-7 | 2024 | Temporally distinct 3D multi-omic dynamics in the developing human brain | AGREE | The MERFISH gene selection was carried out by first using ... NSForest v245 ... The target sequences for each gene were concatenated ... | The packet explicitly links NS-Forest v2 to MERFISH gene selection and probe construction. [accepted_agree] |
| 10.1038/s41588-024-01990-6 | 2024 | A temporal cortex cell atlas highlights gene expression dynamics during human brain maturation | AGREE | "NS-Forest ... revealed ... minimal marker genes ... MERFISH spatial transcriptomic analysis of a subset of minimal markers ..." | The evidence links NS-Forest minimal markers to a subset used in MERFISH spatial transcriptomic analysis, which is explicit targeted-panel use. [accepted_agree] |
| 10.1038/s42003-021-02517-z | 2021 | Comprehensive in situ mapping of human cortical transcriptomic cell types | AGREE | The genes targeted in the HybISS experiments ... The main computational algorithms used were NSforest23 | The packet explicitly links NSforest to computational selection of genes targeted in the HybISS experiments. [accepted_agree] |
| 10.1101/2023.06.29.547074 | 2023 | Single cell-resolution <i>in situ</i> sequencing elucidates spatial dynamics of multiple sclerosis lesion and disease evolution | DISAGREE | This selection was made manually in combination with different computational frameworks116,117. [Ref. 116 = NS-Forest] | Manual plus computational-framework selection is being treated as EXP-USED here because Ref. 116 is NS-Forest and the user requested this case be counted as tool use for the custom mouse panel. [accepted_disagree] |

| DOI | Year | Title | Audit decision | Evidence (verbatim quote) | Audit reason / validation status |
|---|---|---|---|---|---|
| 10.1101/2025.06.13.659535 | 2025 | Non-disruptive 3D profiling of combinations of epigenetic marks in single cells | AGREE | To aid gene selection, we used a custom algorithm as well as NS-Forest101. | This targeted in situ epigenetic profiling paper says NS-Forest was used to aid gene selection, which is a direct local link to target selection for the assay. [accepted_agree] |
| 10.3389/fncel.2023.1256619 | 2023 | Cochlear transcriptome analysis of an outbred mouse population (CFW) | AGREE | The NSForest cell-type specific marker genes ... were utilized as MERFISH probes | The packet explicitly states that NSForest marker genes were used as MERFISH probes. [accepted_agree] |
| 10.64898/2026.01.10.694705 | 2026 | The Caudate Nucleus Exhibits Distinct Pathology and Cell Type-Specific Responses Across Alzheimer's Disease | AGREE | "An additional set of 11 genes were added to improve discrimination of MSN subtypes ... using NSForest ..." | The quote says additional genes were added using NS-Forest to improve subtype discrimination, which supports tool-driven targeted-gene addition for the study. [accepted_agree] |
| 10.64898/2026.02.03.703645 | 2026 | Single-cell Multiome Analysis of Chromatin State and Transcriptome in the Human Basal Ganglia | AGREE | Two initial subsets of genes were selected based on the snRNA-seq DEGs using NS Forest124 and snm3C-seq DMGs using Spapros separately | The packet explicitly states that NS-Forest was used to select an initial gene subset for the final 920-gene MERFISH panel. [accepted_agree] |
| 10.64898/2026.02.12.705355 | 2026 | Multi-gene biomarkers reveal spatial organization and subpopulation-specific damage response in intrahepatic biliary epithelial cells | AGREE | A machine learning algorithm, NSForest, identified minimal, multi-gene signatures for BEC subpopulations. These metagenes ... were validated using hybridization chain reaction (HCR) FISH | The packet abstract explicitly states NSForest identified minimal multi-gene signatures, and those metagenes were validated by HCR FISH; this links the named tool to the in situ biomarker panel. [accepted_agree] |

#### Spapros (8)

| DOI | Year | Title | Audit decision | Evidence (verbatim quote) | Audit reason / validation status |
|---|---|---|---|---|---|
| 10.1016/j.immuni.2025.10.003 | 2025 | Single-cell spatial transcriptomic profiling defines a pathogenic inflammatory niche in chronic active multiple sclerosis lesions | AGREE | For the selection of 500 genes for MERFISH imaging ... the top 150 genes identified using Spapros ... were selected | The packet explicitly states that Spapros was used to select genes for a 500-gene MERFISH panel. [accepted_agree] |
| 10.1038/s41467-025-65704-0 | 2025 | Spatial single-cell atlas reveals regional variations in healthy and diseased human lung | AGREE | generated a probe panel for HybISS using Spapros ... We applied this panel | The packet explicitly states that the HybISS probe panel was generated using Spapros and then applied experimentally. [accepted_agree] |
| 10.1101/2024.09.17.613303 | 2024 | Endothelial regulatory circuits control cranial neural crest | AGREE | We used SPAPROS, a probe selection algorithm ... | The packet explicitly states that the MERSCOPE gene panel was |

| DOI | Year | Title | Audit decision | Evidence (verbatim quote) | Audit reason / validation status |
|---|---|---|---|---|---|
| | | migration and plasticity | | | designed using SPAPROS. [accepted_agree] |
| 10.1101/2025.05.13.653495 | 2025 | A spatiotemporal cancer cell trajectory underlies glioblastoma heterogeneity | AGREE | "The additional custom probes were largely selected by applying ... Spapros ... to our snRNA-seq data." | The evidence directly states that custom Xenium probes were largely selected by applying Spapros to the study's snRNA-seq data. [accepted_agree] |
| 10.1101/2025.08.29.673064 | 2025 | Spatial Analysis of T Cell Clonality in Autoimmune Kidney Disease Using TRV Probes | AGREE | "150 genes were selected using the Spapros algorithm ... for the 10x Xenium panel." | The evidence explicitly states that genes for the 10x Xenium panel were selected using the Spapros algorithm, which is direct panel-design use. [accepted_agree] |
| 10.1101/2025.10.29.684766 | 2025 | A dual genetic constraint underlies the conservation of early brains in vertebrates | AGREE | We identify this blueprint by running spapros74 on this integrated dataset ... Padlock probes against the selected genes were designed ... Direct RNA In situ sequencing (ISS). | The packet explicitly shows that Spapros was run to identify the 100-gene signature, that padlock probes were designed against the selected genes, and that the study used direct RNA in situ sequencing, so this is experimental targeted-panel use. [accepted_agree] |
| 10.64898/2026.02.03.703645 | 2026 | Single-cell Multiome Analysis of Chromatin State and Transcriptome in the Human Basal Ganglia | AGREE | Two initial subsets of genes were selected based on the snRNA-seq DEGs using NS Forest124 and snm3C-seq DMGs using Spapros separately | The packet explicitly states that Spapros was used to select an initial gene subset for the final 920-gene MERFISH panel. [accepted_agree] |
| 10.64898/2026.02.12.705594 | 2026 | A Multimodal Single-Cell Epigenomic and 3D Genome Atlas of the Human Basal Ganglia | AGREE | To design the gene panel, we used Spapros147 to prioritize genes differentially expressed across cell types | The packet explicitly states that Spapros was used to design the gene panel for this spatially validated study. [accepted_agree] |

#### PERSIST (5)

| DOI | Year | Title | Audit decision | Evidence (verbatim quote) | Audit reason / validation status |
| --- | --- | --- | --- | --- | --- |
| 10.1016/j.immuni.2026.03.003 | 2026 | Distinct tissue niches contribute to prostate tissue-resident memory CD8+ T cell differentiation and heterogeneity | AGREE | Next, supervised and unsupervised PERSIST96 was run using as mouse intestine scRNA-seq reference dataset and a mouse lung scRNA-seq dataset with the previously compiled set of genes as prior information, to yield a final list of 480 genes. | The paper explicitly states that supervised and unsupervised PERSIST was run with prior information to yield the final 480-gene list for the Xenium panel. [accepted_agree] |
| 10.1038/s41586-024-08466-x | 2025 | Tissue-resident memory CD8 T cell diversity is spatiotemporally imprinted | AGREE | Gene panel design made use of PERSIST10 | The evidence explicitly says gene panel design made use of PERSIST in this spatial study. [accepted_agree] |
| 10.1038/s41586-025-09921-z | 2026 | Neuro-epithelial circuits promote sensory convergence and intestinal immunity | AGREE | "... PERSIST ... was run ... to produce a final list of 480 genes ... Spatial transcriptomics was performed using the 10x Genomics Xenium platform ..." | The packet explicitly states PERSIST was run to produce the final 480-gene Xenium panel for the study, which is direct targeted-panel use. [accepted_agree] |
| 10.1084/jem.20240776 | 2025 | The XCL1-XCR1 axis supports intestinal tissue residency and antitumor immunity. | AGREE | Using this set as prior information to PERSIST ... unsupervised PERSIST filled in the remaining 79 genes ... supervised PERSIST ... adding an additional 70 genes ... capture 59 cell-type marker genes for 350 total targets | The packet explicitly states that PERSIST filled in and added genes for the final Xenium panels, which is direct targeted-panel use. [accepted_agree] |
| 10.1101/2024.03.20.585130 | 2024 | Functional Diversity of Memory CD8 T Cells is Spatiotemporally Imprinted | AGREE | "For the Xenium mouse 350 gene panel ... supervised PERSIST ... add[ed] an additional 70 genes ... [and] 59 cell type marker genes for 350 total targets." | The packet explicitly says supervised PERSIST was run to add genes for the Xenium mouse 350-gene panel, satisfying the targeted-panel and tool-driven selection requirements. [accepted_agree] |

#### geneBasis (4)

| DOI | Year | Title | Audit decision | Evidence (verbatim quote) | Audit reason / validation status |
| --- | --- | --- | --- | --- | --- |
| 10.1038/s41587-023-01747-2 | 2023 | Multimodal spatiotemporal phenotyping of human retinal organoid development | AGREE | The 100-gene panel ... was selected using geneBasisR48 | The packet explicitly states that the 100-gene panel was selected using geneBasisR. [accepted_agree] |
| 10.1101/2025.03.20.644368 | 2025 | Multi-omic human neural organoid cell atlas of the posterior brain | AGREE | To design the 500-gene panel ... an additional list of genes selected by geneBasis47 | The packet explicitly states that geneBasis-selected genes were added to the 500-gene |

| DOI | Year | Title | Audit decision | Evidence (verbatim quote) | Audit reason / validation status |
|---|---|---|---|---|---|
| | | | | | MERSCOPE panel. [accepted_agree] |
| 10.1101/2025.11.22.688128 | 2025 | A cross-species spatial transcriptomic atlas of the human and non-human primate basal ganglia | AGREE | These were used as a starter list for two gene selection tools, mFISHtools100 and geneBasis101. | The packet explicitly links geneBasis to the paper's gene-selection workflow for the spatial atlas panel. [accepted_agree] |
| 10.1101/2025.11.22.689869 | 2025 | Spatial patterning of transcriptional and regulatory programs in the primate subcortex | AGREE | A panel of 300 genes was constructed ... using ... GeneBasis134. | The packet explicitly states that the 300-gene panel was constructed using GeneBasis, satisfying targeted panel selection by the named tool. [accepted_agree] |

#### gpsFISH (2)

| DOI | Year | Title | Audit decision | Evidence (verbatim quote) | Audit reason / validation status |
|---|---|---|---|---|---|
| 10.1038/s41467-024-46437-y | 2024 | Mapping human tissues with highly multiplexed RNA in situ hybridization | AGREE | Human kidney gene selection was performed by gpsFISH82,83 | The evidence explicitly links gpsFISH to gene selection for the human kidney in situ experiment. [accepted_agree] |
| 10.1101/2025.08.07.667992 | 2025 | Single-cell multiomic human brain atlas reveals regulatory drivers of cortical regionality | AGREE | A 484-gene panel was selected with gpsFISH (26). | The packet explicitly says a 484-gene panel was selected with gpsFISH. [accepted_agree] |

#### COSG (1)

| DOI | Year | Title | Audit decision | Evidence (verbatim quote) | Audit reason / validation status |
|---|---|---|---|---|---|
| 10.1038/s41586-025-09996-8 | 2026 | Pyramidal neurons proportionately alter cortical interneuron subtypes | AGREE | "To design the MERFISH gene panel ... Top marker genes identified by COSG were combined ... to generate an initial gene panel ..." | The evidence explicitly links COSG marker genes to design of the MERFISH gene panel. [accepted_agree] |

#### SCMER (1)

| DOI | Year | Title | Audit decision | Evidence (verbatim quote) | Audit reason / validation status |
|---|---|---|---|---|---|
| 10.1038/s41467-023-40674-3 | 2023 | Spatial organization of the mouse retina at single cell resolution by MERFISH | AGREE | "We used SCMER ... to choose additional genes ... which brought up the total number in the second panel to 500 genes." | The evidence explicitly links SCMER to choosing additional genes for the paper's MERFISH panel, so this is direct experimental |

| DOI | Year | Title | Audit decision | Evidence (verbatim quote) | Audit reason / validation status |
|---|---|---|---|---|---|
| | | | | | panel-selection use. [accepted_agree] |

#### scGIST (1)

| DOI | Year | Title | Audit decision | Evidence (verbatim quote) | Audit reason / validation status |
|---|---|---|---|---|---|
| 10.1101/2025.08.19.671030 | 2025 | OpenFISH enables integrated high-resolution spatial transcriptomics and metabolomics on a single tissue section | AGREE | "We ran scGIST ... In total, we selected 109 genes ... for our final panel." | The evidence directly states scGIST was run and that its selected genes formed the final panel, meeting the targeted-panel use criterion. [accepted_agree] |

#### scGeneFit (1)

| DOI | Year | Title | Audit decision | Evidence (verbatim quote) | Audit reason / validation status |
|---|---|---|---|---|---|
| 10.64898/2025.12.19.695425 | 2025 | Exploiting pair correlation function to describe biological tissue structure | AGREE | The obtained clustering was then used as an input for the Python package scGeneFit ... and 40 markers. | The packet explicitly says scGeneFit was used to derive markers that informed the experimental antibody panel. [accepted_agree] |

#### scPNMF (1)

| DOI | Year | Title | Audit decision | Evidence (verbatim quote) | Audit reason / validation status |
|---|---|---|---|---|---|
| 10.1101/2024.10.08.617260 | 2024 | Spatial Single-Cell Mapping of Transcriptional Differences Across Genetic Backgrounds in Mouse Brains | AGREE | Encoding matrix was fit using a modified version of DPNMF/scPNMF to optimize the gene set. Resulting encoding matrix consisted of 6,654 genes and 35,919 probes. | The paper uses the DPNMF/scPNMF encoding matrix to optimize the gene set and generate the final probe/signature design for its own targeted spatial assay, so this is experimental tool use rather than background citation. [accepted_agree] |

### EXP-CITED (n = 52)

*Experimental spatial-transcriptomics paper that cited the tool without reporting that it used it for panel design.*

#### COSG (22)

| DOI | Year | Title | Audit decision | Evidence (verbatim quote) | Audit reason / validation status |
|---|---|---|---|---|---|
| 10.1002/ctm2.70425 | 2025 | Multi-omics profiling identifies <i>TNFRSF18</i> as a novel marker of exhausted $CD8^+$ T cells and reveals tumour-immune dynamics in colorectal cancer | AGREE | "TNFRSF18 was robustly recognized as a marker gene ... using both Seurat's FindMarkers function and COSG analysis." | The packet shows COSG was used to recognize a marker gene, but there is no nearby statement linking COSG to targeted spatial panel, probe, or target-gene selection. [accepted_agree] |
| 10.1007/s12672-024-01614-9 | 2024 | Computer-aided analysis reveals metallothionein-positive cancer-associated fibroblasts promote angiogenesis in gastric adenocarcinoma | AGREE | To characterize gene expression patterns, COSG algorithm was applied to revealed distinct gene signatures. | This experimental spatial paper uses COSG for gene-signature characterization only; there is no nearby link to targeted panel or probe selection. [accepted_agree] |
| 10.1016/j.brainres.2025.149688 | 2025 | Landscape of spatial disruption of homeostasis in the middle temporal gyrus of epileptic patients | DISAGREE | Abstract states the study investigates epilepsy using single-cell RNA sequencing and spatial transcriptomics; evidence quote does not mention COSG. | The packet clearly describes an experimental study using both scRNA-seq and spatial transcriptomics, but there is no close-context evidence linking COSG to targeted panel selection, so the conservative negative label is EXP-CITED. [accepted_disagree] |
| 10.1016/j.ccell.2025.03.017 | 2025 | Spatial determinants of antibody-drug conjugate SHR-A1811 efficacy in neoadjuvant treatment for HER2-positive breast cancer | AGREE | "Differential expression analyses between cell groups were carried out by COSG ..." | COSG is used for differential expression between cell groups, but there is no nearby statement linking it to targeted spatial panel or probe selection. [accepted_agree] |
| 10.1016/j.cell.2025.05.047 | 2025 | A Drosophila single-cell 3D spatiotemporal multi-omics atlas unveils panoramic key regulators of cell-type differentiation | AGREE | Top 200 marker genes generated by the COSG 99 were also examined for additional reference | The packet shows COSG used to generate marker genes for reference, with no nearby statement linking COSG to targeted spatial panel or probe selection. [accepted_agree] |
| 10.1016/j.celrep.2023.112643 | 2023 | Transcriptomic landscape of early hair follicle and epidermal development | AGREE | We also exploited an alternative way to identify cluster markers using the ... COSG pipeline70 | COSG is used to identify cluster markers, but there is no nearby link to targeted spatial panel or probe selection. [accepted_agree] |
| 10.1016/j.celrep.2023.113121 | 2023 | Distinct mechanisms for sebaceous gland self-renewal and regeneration provide durability in response to injury | AGREE | The COSG-identified top genes were used to establish the cell identity of each cluster | COSG-identified genes were used to assign cell identity, not explicitly to select a targeted spatial panel or probes. [accepted_agree] |

| DOI | Year | Title | Audit decision | Evidence (verbatim quote) | Audit reason / validation status |
|---|---|---|---|---|---|
| 10.1016/j.devcel.2025.03.009 | 2025 | Transcriptional conservation and evolutionary divergence of cell types across mammalian hypothalamus development | AGREE | "we used cosine similarity algorithm ... to designate cell subtype-specific genes" | The paper includes spatial transcriptomic data, but the evidence only links COSG to subtype-specific gene designation, not panel selection. [accepted_agree] |
| 10.1016/j.stem.2025.10.001 | 2025 | 3D-generation of high-purity midbrain dopaminergic progenitors and lineage-guided refinement of grafts supports Parkinson’s disease cell therapy | AGREE | "The top 100 differentially expressed genes ... were identified using the COSG ... package ..." | The paper has single-cell spatial transcriptomics, but the evidence only shows COSG was used to identify differentially expressed genes, not to select a targeted panel. [accepted_agree] |
| 10.1038/s12276-026-01636-z | 2026 | UBE2M as a bridge spanning neddylation and cell cycle regulation in colorectal adenocarcinoma | AGREE | we applied the COSG package (v0.9.0) to identify the top 500 marker genes of the target cell clusters | The packet only shows COSG used to identify marker genes for target clusters, without a nearby link to targeted spatial panel or probe selection. [accepted_agree] |
| 10.1038/s41421-024-00656-1 | 2024 | Single-cell multi-omics analysis of lineage development and spatial organization in the human fetal cerebellum | AGREE | COSG ... was used to calculate marker genes for each cluster. | COSG is used for cluster marker calculation only; there is no close-context link to targeted panel or probe selection. [accepted_agree] |
| 10.1038/s41467-026-68291-w | 2026 | Integrated multi-omic atlas reveals the hierarchy of spatiotemporal regulatory networks of mouse gastrulation | AGREE | the top 100 cell-type marker genes identified by the COSG (v 0.9.0) package63 were designated as zipcode genes | The packet only states that COSG marker genes were designated as zipcode genes; it does not explicitly tie COSG to a targeted spatial panel or probe-set selection event. [accepted_agree] |
| 10.1038/s41588-024-01662-5 | 2024 | Multi-omic profiling of clear cell renal cell carcinoma identifies metabolic reprogramming associated with disease progression | AGREE | Cell subpopulations were named based on top markers found by the COSG (v0.9.0) package. | COSG is used to determine top markers for naming subpopulations, but the packet does not link COSG to targeted spatial panel selection. [accepted_agree] |
| 10.1038/s41588-025-02139-9 | 2025 | Spatially resolved transcriptomic analysis of the adult human prostate | AGREE | Cell subpopulations were named based on the top markers determined by the COSG package54. | COSG is only used to determine top markers for naming cell subpopulations; there is no nearby link to targeted spatial panel selection. [accepted_agree] |
| 10.1038/s41593-025-02042-w | 2025 | FOXP genes regulate Purkinje cell diversity and cerebellar | AGREE | "Cell-type-specific genes were identified using the COSG | COSG is used to identify cell-type-specific genes, but the packet does |

| DOI | Year | Title | Audit decision | Evidence (verbatim quote) | Audit reason / validation status |
|---|---|---|---|---|---|
| | | morphogenesis | | package ..." | not link it to targeted spatial panel or probe selection. [accepted_agree] |
| 10.1093/neuonc/noaf179 | 2025 | A cycling, progenitor-like cell population at the base of atypical teratoid rhabdoid tumor subtype differentiation trajectories | AGREE | A connectivity map was generated by submitting the top 100 marker genes for IPC-like, NPC-like, OPC-like, and mesenchymal-like cell populations (COSGR46) to CLUE,47 | The packet only shows COSG used to obtain marker genes for a connectivity map; there is no nearby link to targeted spatial panel, probe, or target-gene selection. [accepted_agree] |
| 10.1101/2025.04.09.647847 | 2025 | <i>Tff2</i> marks gastric corpus progenitors that give rise to pyloric metaplasia/SPEM following injury | AGREE | "Marker genes were identified using the COSG package ..." | COSG is used to identify marker genes, but there is no nearby statement linking it to targeted spatial panel or probe selection. [accepted_agree] |
| 10.1101/2025.08.19.671030 | 2025 | OpenFISH enables integrated high-resolution spatial transcriptomics and metabolomics on a single tissue section | AGREE | we employed COSG53 to identify differentially expressed TE subfamilies ... The most highly and variably expressed TE loci ... were selected for probe design | The packet explicitly links COSG-derived TE subfamilies to subsequent locus selection for probe design in the paper's own spatial assay. [MANUAL DEMOTION 2026-05-20: panel-design audit attributes the final panel to scGIST; COSG was used for upstream TE/marker exploration only, not for panel selection. Demoted EXP-USED to EXP-CITED to keep the citation EXP-USED set aligned with panel-selection use.] [manual_demoted_post_audit] |
| 10.3389/fimmu.2025.1521038 | 2025 | Unraveling the spatial and signaling dynamics and splicing kinetics of immune infiltration in osteoarthritis synovium | AGREE | "We then applied COSG ... to determine marker genes for each cluster." | COSG is used to determine cluster marker genes, but the packet does not explicitly connect COSG to selection of a targeted spatial panel, probe set, or target list. [accepted_agree] |
| 10.64898/2026.02.26.705527 | 2026 | Protein-guided RNA barcoding links transcriptomes to synaptic architecture | AGREE | "Marker genes were identified using COSG ..." | The packet only says COSG was used to identify marker genes; there is no nearby link to design of a targeted spatial panel or probe set. [accepted_agree] |
| 10.7150/ijbs.114618 | 2025 | Single-cell and spatial transcriptomic analysis reveal cellular heterogeneity and cancer cell-intrinsic major | AGREE | "Cell type-specific gene signatures were defined on basis of the top 50 marker genes identified using ... COSG ..." | COSG is used to define cell type-specific signatures in a single-cell and spatial transcriptomic analysis, but not for targeted panel |

| DOI | Year | Title | Audit decision | Evidence (verbatim quote) | Audit reason / validation status |
|---|---|---|---|---|---|
| | | histocompatibility complex II expression in urothelial carcinoma | | | selection. [accepted_agree] |
| 10.7150/thno.99046 | 2024 | <i>DAB2</i><sup>+</sup> macrophages support <i>FAP</i><sup>+</sup> fibroblasts in shaping tumor barrier and inducing poor clinical outcomes in liver cancer | AGREE | "cell type-specific genes ... were initially screened using the R package COSG ... The top 30 ranked genes were then utilized for ssGSEA ..." | The study includes spatial transcriptome profiling, but COSG is used for cell type-specific genes and bulk-cohort scoring rather than targeted panel selection. [accepted_agree] |

#### NS-Forest v2 (7)

| DOI | Year | Title | Audit decision | Evidence (verbatim quote) | Audit reason / validation status |
|---|---|---|---|---|---|
| 10.1038/s41467-025-58874-4 | 2025 | Deconvolution of cell types and states in spatial multiomics utilizing TACIT | DISAGREE | Abstract describes spatial biology datasets and combined spatial transcriptomics; evidence only says "we develop TACIT ..." | The packet describes an experimental spatial biology study built around TACIT, but the provided evidence does not show explicit NS-Forest benchmarking or use, so the conservative negative label is EXP-CITED. [accepted_disagree] |
| 10.1101/2023.09.29.560114 | 2023 | Cell type-specific gene expression dynamics during human brain maturation | AGREE | The NS-Forest tool (v2.0) ... was used to identify combinations of marker genes uniquely defining each annotated cell type | NS-Forest is used to define marker-gene combinations for cell types, but the packet does not connect it to targeted spatial panel or probe design. [accepted_agree] |
| 10.1101/2024.01.31.576150 | 2024 | Charting the Spatial Transcriptome of the Human Cerebral Cortex at Single-Cell Resolution | AGREE | We used NS-Forest ... to identify the minimum set of marker genes whose combined expression identified the cells of one type with maximum accuracy | The packet shows NS-Forest used to identify marker genes, but there is no nearby statement linking those markers to final targeted panel/probe selection. [accepted_agree] |
| 10.1101/2025.05.23.655666 | 2025 | A Multimodal Spatial and Epigenomic Atlas of Human Adult Lung Topography | AGREE | To identify a minimal set of characterizing marker genes ... the validated NS-Forest method20 ... was used | The evidence only links NS-Forest to marker-gene identification from snRNA-seq, not to final MERFISH panel or probe selection. [accepted_agree] |
| 10.1126/sciadv.abn4965 | 2022 | A reference tissue atlas for the human kidney | AGREE | Evidence quote: NOT FOUND; abstract describes construction of a spatially specified human kidney atlas. | The packet clearly describes an experimental spatial atlas paper, and there is no evidence of targeted-panel use by NS-Forest, so the conservative negative label is EXP-CITED. [accepted_agree] |

| DOI | Year | Title | Audit decision | Evidence (verbatim quote) | Audit reason / validation status |
|---|---|---|---|---|---|
| 10.1186/s12864-024-10434-8 | 2024 | Spatial transcriptomic brain imaging reveals the effects of immunomodulation therapy on specific regional brain cells in a mouse dementia model | DISAGREE | "NSForest ... was applied and signature genes for the cell type were determined based on the cell type annotation information." | The paper is an experimental spatial transcriptomics study, but the evidence only says NS-Forest was used to determine cell-type signature genes, not to select a targeted spatial panel or probe set. [accepted_disagree] |
| 10.26508/lsa.202201701 | 2022 | Concordance of MERFISH spatial transcriptomics with bulk and single-cell RNA sequencing | AGREE | We performed MERFISH on the mouse liver and kidney and compared the resulting bulk and single-cell RNA statistics ... | This is a clear experimental spatial-transcriptomics paper; with the record assumed to be in the authoritative citing set and no close-context panel-selection use evidence, the conservative label is EXP-CITED. [accepted_agree] |

#### geneBasis (4)

| DOI | Year | Title | Audit decision | Evidence (verbatim quote) | Audit reason / validation status |
|---|---|---|---|---|---|
| 10.1038/s41467-024-46669-y | 2024 | Highly sensitive spatial transcriptomics using FISHnCHIPs of multiple co-expressed genes | AGREE | geneBasis: an iterative approach for unsupervised selection of targeted gene panels from scRNA-seq | The packet only contains a citation to geneBasis, not evidence that it was used for this paper's own targeted panel selection. [accepted_agree] |
| 10.1038/s41586-025-09875-2 | 2025 | Spatiotemporal cellular map of the developing human reproductive tract | DISAGREE | — | The packet provides no evidence text linking geneBasis to panel selection. The study is clearly experimental/spatial, so the conservative negative label is EXP-CITED rather than NEEDS_MANUAL. [accepted_disagree] |
| 10.1126/science.adt3439 | 2026 | Whole-embryo spatial transcriptomics at subcellular resolution from gastrulation to organogenesis | AGREE | Evidence quote: NOT FOUND; abstract describes whole-embryo spatial transcriptomics with weMERFISH. | This is a clear experimental spatial transcriptomics paper, and without a nearby tool-specific quote the conservative negative label is EXP-CITED. [accepted_agree] |
| 10.26508/lsa.202201701 | 2022 | Concordance of MERFISH spatial transcriptomics with bulk and single-cell RNA sequencing | AGREE | Missarova A ... (2021) GeneBasis: An iterative approach for unsupervised selection of targeted gene panels from ScRNA-seq. | The packet contains only a bibliographic citation to geneBasis, not evidence of direct use for this paper's panel selection. [accepted_agree] |

#### NS-Forest v4.0 (3)

| DOI | Year | Title | Audit decision | Evidence (verbatim quote) | Audit reason / validation status |
|---|---|---|---|---|---|
| 10.1101/2025.05.23.655666 | 2025 | A Multimodal Spatial and Epigenomic Atlas of Human Adult Lung Topography | AGREE | we applied NS-Forest ... to identify necessary and sufficient marker genes at each annotation level | The packet only reports marker-gene identification by NS-Forest, not explicit use for selecting the paper's final spatial panel or probes. [accepted_agree] |
| 10.1101/2025.11.22.689869 | 2025 | Spatial patterning of transcriptional and regulatory programs in the primate subcortex | AGREE | "Marker genes for every Cluster and Group were computed using NSForest ..." | The evidence shows NS-Forest was used to compute marker genes, but it does not explicitly tie the tool to targeted spatial panel, probe, or target-list design. [accepted_agree] |
| 10.64898/2025.12.22.695505 | 2025 | Diverse infections transcriptionally reprogram the intestinal epithelium and epithelial-immune cell interactions | AGREE | NOT FOUND | This is an experimental study using single-cell and spatial transcriptomics, but the packet provides no nearby evidence that NS-Forest drove panel or probe selection. [accepted_agree] |

#### PERSIST (3)

| DOI | Year | Title | Audit decision | Evidence (verbatim quote) | Audit reason / validation status |
|---|---|---|---|---|---|
| 10.1038/s41467-024-46669-y | 2024 | Highly sensitive spatial transcriptomics using FISHnCHIPs of multiple co-expressed genes | AGREE | Evidence quote: NOT FOUND; abstract describes FISHnCHIPs for spatially resolved analysis of intact tissue samples. | This is a clear experimental spatial transcriptomics paper, and without a nearby PERSIST-specific usage quote the conservative negative label is EXP-CITED. [accepted_agree] |
| 10.1038/s41477-025-02176-6 | 2026 | Imputation integrates single-cell and spatial gene expression data to resolve transcriptional networks in barley shoot meristem development | DISAGREE | We then used PERSIST to identify gene expression combinations suitable as barcodes for stages P1–P19 | PERSIST is used to identify gene-expression barcodes, but the packet does not explicitly connect that step to selecting the study's final targeted spatial panel or probe set. [accepted_disagree] |
| 10.1038/s41587-025-02811-9 | 2025 | Standardized metrics for assessment and reproducibility of imaging-based spatial transcriptomics datasets | AGREE | Evidence quote: NOT FOUND; abstract describes standardized metrics for imaging-based spatial transcriptomics datasets. | This is a clear experimental spatial transcriptomics assessment paper, and without a nearby PERSIST-specific usage quote the conservative negative label is |

| DOI | Year | Title | Audit decision | Evidence (verbatim quote) | Audit reason / validation status |
|---|---|---|---|---|---|
| | | | | | EXP-CITED. [accepted_agree] |

#### Spapros (3)

| DOI | Year | Title | Audit decision | Evidence (verbatim quote) | Audit reason / validation status |
|---|---|---|---|---|---|
| 10.1016/j.labinv.2025.104190 | 2025 | Rigor and Reproducibility of Spatial Transcriptomics Performed on Clinically Sourced Human Tissues | AGREE | In this study, we vet the technical reproducibility, data normalization methods, and assay sensitivity ... on one widely used spatial transcriptomics methodology. | This is an experimental spatial-transcriptomics reproducibility study; with authoritative citation status and no close-context Spapros-driven panel-selection evidence, the conservative label is EXP-CITED. [accepted_agree] |
| 10.1038/s41467-024-46669-y | 2024 | Highly sensitive spatial transcriptomics using FISHnCHIPs of multiple co-expressed genes | AGREE | We present FISHnCHIPs for highly sensitive in situ profiling of cell types and gene expression programs. | This is an experimental in situ spatial-transcriptomics paper; under the authoritative citing-set assumption and without local evidence that Spapros drove panel selection, the conservative label is EXP-CITED. [accepted_agree] |
| 10.1038/s41467-025-63798-0 | 2025 | Cassini: streamlined and scalable method for in situ profiling of RNA and protein | AGREE | NOT FOUND | This is an experimental in situ profiling methods paper, but the packet provides no nearby evidence that Spapros drove panel or probe selection. [accepted_agree] |

#### gpsFISH (3)

| DOI | Year | Title | Audit decision | Evidence (verbatim quote) | Audit reason / validation status |
|---|---|---|---|---|---|
| 10.1021/acsomega.5c05379 | 2025 | Enhanced RNA Preservation in Mouse Brain Tissue: A Strategy Combining Cardiac Perfusion with Hypersaline Immersion | AGREE | Evidence quote: NOT FOUND; abstract describes a preservation strategy for capillary-based spatial omics studies of mouse brain sections. | This is a clear experimental spatial transcriptomics paper, and without any nearby tool-use quote the conservative negative label is EXP-CITED. [accepted_agree] |
| 10.1101/2025.04.23.649965 | 2025 | From Transcripts to Cells: Dissecting Sensitivity, Signal Contamination, and Specificity in Xenium Spatial Transcriptomics | AGREE | NOT FOUND | This is an experimental Xenium performance/benchmark paper, but the packet provides no nearby evidence that gpsFISH drove panel selection or was directly |

| DOI | Year | Title | Audit decision | Evidence (verbatim quote) | Audit reason / validation status |
|---|---|---|---|---|---|
| | | | | | benchmarked. [accepted_agree] |
| 10.1186/s13059-025-03624-4 | 2025 | Comparison of spatial transcriptomics technologies using tumor cryosections | AGREE | NOT FOUND | This is an experimental spatial-transcriptomics technology-comparison paper, but the packet provides no nearby evidence that gpsFISH drove panel selection or was benchmarked specifically. [accepted_agree] |

#### SMaSH (2)

| DOI | Year | Title | Audit decision | Evidence (verbatim quote) | Audit reason / validation status |
|---|---|---|---|---|---|
| 10.1038/s41467-023-42658-9 | 2023 | Dbh+ catecholaminergic cardiomyocytes contribute to the structure and function of the cardiac conduction system in murine heart | DISAGREE | To identify specific and robust marker genes for each cell type, we used SMASH26, and in brief, kept the top 50 genes (Table S2) that contributed most to the classification of each cell type through identifying gene-wise Gini importance from the trained supervised ensemble machine learning classifier algorithm. | This Stereo-seq study is spatial, but Stereo-seq is whole-transcript capture rather than a targeted panel assay. The packet only shows SMaSH used to identify marker genes, not to choose a final targeted spatial panel, probe set, or target list. [accepted_disagree] |
| 10.1186/s12864-024-10434-8 | 2024 | Spatial transcriptomic brain imaging reveals the effects of immunomodulation therapy on specific regional brain cells in a mouse dementia model | AGREE | SMaSH: a scalable, general marker gene identification framework for single-cell RNA-sequencing | This is an experimental spatial-transcriptomics paper, but the packet only shows a citation to the SMaSH paper, not use for targeted panel selection. [accepted_agree] |

#### scGeneFit (2)

| DOI | Year | Title | Audit decision | Evidence (verbatim quote) | Audit reason / validation status |
|---|---|---|---|---|---|
| 10.1016/j.ccell.2025.04.003 | 2025 | Spatial and multiomics analysis of human and mouse lung adenocarcinoma precursors reveals TIM-3 as a putative target for precancer interception | DISAGREE | NOT FOUND | The packet is clearly an experimental spatial/multiomics paper, but no scGeneFit evidence was found. Under the addendum, the conservative negative label is EXP-CITED rather than NEEDS_MANUAL. [accepted_disagree] |
| 10.1038/s41467-025-58748-9 | 2025 | Hypoxia-induced Wnt5a-secreting fibroblasts promote colon cancer progression | AGREE | "scGeneFit ... was used to select marker genes that retain the classification of cell types ..." | The packet shows scGeneFit was used to select marker genes, but there is no nearby statement |

| DOI | Year | Title | Audit decision | Evidence (verbatim quote) | Audit reason / validation status |
|---|---|---|---|---|---|
| | | | | | linking it to targeted spatial panel or probe design. [accepted_agree] |

#### RankCorr (1)

| DOI | Year | Title | Audit decision | Evidence (verbatim quote) | Audit reason / validation status |
|---|---|---|---|---|---|
| 10.1186/s12864-024-10434-8 | 2024 | Spatial transcriptomic brain imaging reveals the effects of immunomodulation therapy on specific regional brain cells in a mouse dementia model | AGREE | we investigated the changes in spatial transcriptomic signatures and brain cell types using the 10x Genomics Visium platform | This is a clear experimental spatial-transcriptomics study using Visium; with authoritative citation status and no close-context tool-driven panel-selection statement, the conservative label is EXP-CITED. [accepted_agree] |

#### scGIST (1)

| DOI | Year | Title | Audit decision | Evidence (verbatim quote) | Audit reason / validation status |
|---|---|---|---|---|---|
| 10.1186/s13059-025-03624-4 | 2025 | Comparison of spatial transcriptomics technologies using tumor cryosections | AGREE | scGIST: gene panel design for spatial transcriptomics with prioritized gene sets | The packet only includes a bibliographic mention of scGIST, without a nearby link to panel use in the study. Since the paper is experimental/spatial, EXP-CITED is the conservative label. [accepted_agree] |

#### scMAGS (1)

| DOI | Year | Title | Audit decision | Evidence (verbatim quote) | Audit reason / validation status |
|---|---|---|---|---|---|
| 10.1186/s13059-024-03396-3 | 2024 | Characterization of regeneration initiating cells during Xenopus laevis tail regeneration | AGREE | "... discriminatory gene markers ... selected using scMAGS ..." | The paper used scMAGS-selected markers for downstream annotation enrichment, but the packet does not connect scMAGS to targeted spatial panel or probe selection. [accepted_agree] |

### COMP-BENCH (n = 98)

*Computational benchmarking or comparison paper that ran the tool as part of a benchmark.*

#### scGeneFit (23)

| DOI | Year | Title | Audit decision | Evidence (verbatim quote) | Audit reason / validation status |
|---|---|---|---|---|---|
| 10.1016/j.array.2025.100623 | 2025 | Two-stage copula-driven, structure-aware gene selection for single-cell analysis | AGREE | "... over other well-known existing methods ... and scGene-Fit [30]." | The packet explicitly states the method is compared with scGene-Fit and other methods. [accepted_agree] |
| 10.1016/j.cels.2026.101571 | 2026 | Identifying maximally informative signal-aware representations of single-cell data using the information bottleneck | AGREE | "We compared bioIB performance to scGeneFit ..." | The evidence explicitly compares bioIB against scGeneFit. [accepted_agree] |
| 10.1016/j.compbiomed.2023.106634 | 2023 | scMAGS: Marker gene selection from scRNA-seq data for spatial transcriptomics studies | AGREE | "the performance of scMAGS is compared to that of the existing methods scGeneFit ..." | The packet explicitly compares scMAGS against scGeneFit and other methods. [accepted_agree] |
| 10.1016/j.crmeth.2022.100332 | 2022 | Multiset multicover methods for discriminative marker selection | AGREE | "We tested G-PC and CEM-PC and compared them with eight prior methods: scGeneFit ..." | The packet explicitly lists scGeneFit among prior methods compared against the new method. [accepted_agree] |
| 10.1016/j.swevo.2025.102108 | 2025 | A multi-objective metaheuristic approach for generating cell type marker panels from single-cell transcriptomics | AGREE | "The performance of scMOCTYM is evaluated ... and compared against seven state-of-the-art methods." | The packet explicitly states the method was compared against several state-of-the-art methods, supporting COMP-BENCH. [accepted_agree] |
| 10.1038/s41467-023-37392-1 | 2023 | Predictive and robust gene selection for spatial transcriptomics | AGREE | "... we considered ... the scGeneFit method ..." | The evidence explicitly states PERSIST was compared with scGeneFit and other gene selection methods. [accepted_agree] |
| 10.1038/s41598-024-63492-z | 2024 | CellMarkerPipe: cell marker identification and evaluation pipeline in single cell transcriptomes | AGREE | the prioritized tools, i.e. Seurat, COSG, SC3, SCMarker, COMET, and scGeneFit, to be compared and benchmarked in this paper | The CellMarkerPipe paper explicitly benchmarks scGeneFit among prioritized tools. [accepted_agree] |
| 10.1093/bib/bbae626 | 2024 | CosGeneGate selects multi-functional and credible biomarkers for single-cell analysis | AGREE | "including ... scGeneFit ... and CosGeneGate." | The paper explicitly compares prediction accuracy using markers from scGeneFit and other models. [accepted_agree] |
| 10.1093/bioinformatics/btag051 | 2026 | Inference of marker genes of subtle cell state changes via iLR: iterative logistic regression | AGREE | "we compared iLR genes to genes identified using ... scGeneFit ..." | The packet explicitly compares iLR against scGeneFit on simulated scRNA-seq data. [accepted_agree] |

| DOI | Year | Title | Audit decision | Evidence (verbatim quote) | Audit reason / validation status |
|---|---|---|---|---|---|
| 10.1101/2021.04.08.438978 | 2021 | <tt>SMaSH:</tt> A scalable, general marker gene identification framework for single-cell RNA sequencing and Spatial Transcriptomics | AGREE | "we benchmarked it against ... scGeneFit ..." | The packet explicitly says SMaSH was benchmarked against scGeneFit. [accepted_agree] |
| 10.1101/2023.05.29.542760 | 2023 | Boosting Single-Cell RNA Sequencing Analysis with Simple Neural Attention | AGREE | We compared scANNA-selected genes against markers selected by state-of-the-art ML methods (scGeneFit ... ) | The scANNA paper explicitly compares against scGeneFit. [accepted_agree] |
| 10.1101/2023.06.19.545592 | 2024 | InGene: Finding influential genes from embeddings of nonlinear dimension reduction techniques | AGREE | InGene was compared with ... scGeneFit (16) | The paper explicitly compares InGene against scGeneFit and other methods. [accepted_agree] |
| 10.1101/gr.280637.125 | 2025 | Optimal marker genes for <i>c</i> -separated cell types with SepSolve | AGREE | "We compared the performance of SepSolve to ... scGeneFit ..." | The paper explicitly compares SepSolve with scGeneFit in a computational benchmarking setup. [accepted_agree] |
| 10.1109/mipro65660.2025.11132056 | 2025 | Accelerating Marker Gene Selection in Scrna-Seq | AGREE | We evaluated our implementation against scGeneFit on two datasets | The paper explicitly evaluates the proposed implementation against scGeneFit on two datasets. [accepted_agree] |
| 10.1186/s12859-020-03641-z | 202 | A rank-based marker selection method for high throughput scRNA-seq data | AGREE | "We ran scGeneFit ..." | The packet explicitly states scGeneFit was run as a comparison method in this computational benchmarking paper. [accepted_agree] |
| 10.1186/s12859-022-04860-2 | 2022 | SMaSH: a scalable, general marker gene identification framework for single-cell RNA-sequencing | AGREE | we benchmarked it against two recent standalone computational algorithms, scGeneFit [6] and RankCorr [7] | The SMaSH methods paper explicitly benchmarks against scGeneFit. [accepted_agree] |
| 10.1186/s13059-021-02548-z | 2021 | geneBasis: an iterative approach for unsupervised selection of targeted gene panels from scRNA-seq | AGREE | we applied geneBasis, as well as SCMER and scGeneFit | The geneBasis paper explicitly evaluates scGeneFit alongside other methods. [accepted_agree] |
| 10.1186/s13059-024-03174-1 | 2024 | Gene panel selection for targeted spatial transcriptomics | AGREE | We also compared gpsFISH with two previously published classification-based gene selection methods: RankCorr and scGenefit. | The gpsFISH paper explicitly compares against scGeneFit. [accepted_agree] |
| 10.1186/s13059-024-03185-y | 2024 | scGIST: gene panel design for spatial transcriptomics with prioritized gene sets | AGREE | We ran scGIST as well as scGeneFit, geneBasis, and SMaSH ... and calculated their accuracy | The scGIST paper explicitly evaluates scGeneFit as a baseline and reports accuracy across panel |

| DOI | Year | Title | Audit decision | Evidence (verbatim quote) | Audit reason / validation status |
|---|---|---|---|---|---|
| | | | | | sizes. [accepted_agree] |
| 10.1186/s13059-024-03410-8 | 2024 | scCTS: identifying the cell type-specific marker genes from population-level single-cell RNA-seq | AGREE | "We carefully analyze ... to compare scCTS with ... scGeneFit ..." | The packet explicitly says scCTS was compared with scGeneFit, which supports COMP-BENCH. [accepted_agree] |
| 10.1186/s44330-024-00015-2 | 2024 | Discovery of optimal cell type classification marker genes from single cell RNA sequencing data | AGREE | "To directly compare NS-Forest v4.0 ... RankCorr, scGeneFit, and MarkerMap were considered." | The paper directly compares NS-Forest with several marker-gene selection methods including scGeneFit, which fits COMP-BENCH. [accepted_agree] |
| 10.1371/journal.pcbi.1012643 | 2024 | Hierarchical marker genes selection in scRNA-seq analysis | AGREE | the hierarchical version of the scGeneFit performed even worse than the flat version | The computational paper explicitly compares hierarchical scGeneFit against the flat scGeneFit version. [accepted_agree] |
| 10.7554/elife.107531.1 | 2025 | CellCover Defines Marker Gene Panels Capturing Developmental Progression in Neocortical Neural Stem Cell Identity | AGREE | We compare CellCover with leading marker gene selection algorithms ... scGeneFit [15] | CellCover explicitly compares its performance with scGeneFit and other marker-gene algorithms. [accepted_agree] |

#### RankCorr (15)

| DOI | Year | Title | Audit decision | Evidence (verbatim quote) | Audit reason / validation status |
|---|---|---|---|---|---|
| 10.1016/j.crmeth.2022.100332 | 2022 | Multiset multicover methods for discriminative marker selection | AGREE | "We tested G-PC and CEM-PC and compared them with eight prior methods ... RankCorr ..." | The paper explicitly compares its methods with prior methods including RankCorr, fitting COMP-BENCH. [accepted_agree] |
| 10.1016/j.jbi.2023.104510 | 2023 | MAGNETO: Cell type marker panel generator from single-cell transcriptomic data | AGREE | including RANKCORR [23], SMaSH [27], Hypergate [22], sc2marker [17], and COMET [15] | MAGNETO is explicitly benchmarked against RankCorr and other state-of-the-art methods. [accepted_agree] |
| 10.1016/j.swevo.2025.102108 | 2025 | A multi-objective metaheuristic approach for generating cell type marker panels from single-cell transcriptomics | AGREE | The performance of scMOCTYM was evaluated by conducting comparisons with ... RANKCORR [17] | The computational marker-panel paper explicitly compares scMOCTYM against RankCorr. [accepted_agree] |

| DOI | Year | Title | Audit decision | Evidence (verbatim quote) | Audit reason / validation status |
|---|---|---|---|---|---|
| 10.1038/s41540-024-00339-3 | 2024 | MarkerMap: nonlinear marker selection for single-cell studies | AGREE | We benchmark MarkerMap against existing marker selection approaches and related methodologies from the wider explainable machine learning literature. | This single-cell marker-selection methods paper explicitly benchmarks MarkerMap against existing marker-selection approaches, so COMP-BENCH is appropriate for the named tool. [accepted_agree] |
| 10.1038/s43588-021-00070-7 | 2021 | Single-cell manifold-preserving feature selection for detecting rare cell populations | AGREE | The comparison passage explicitly includes "RankCorr" among the evaluated methods. | The packet explicitly lists RankCorr among comparison methods, supporting COMP-BENCH. [accepted_agree] |
| 10.1093/bib/bbaf478 | 2025 | Multi-view gene panel characterization for spatially resolved omics | AGREE | In addition, we compared panelScope-OA against several methods tested here. | This is a computational panel-design paper for spatially resolved omics that explicitly compares panelScope-OA against other methods, so COMP-BENCH is the correct conservative label. [accepted_agree] |
| 10.1101/2021.04.08.438978 | 2021 | <tt>SMaSH:</tt> A scalable, general marker gene identification framework for single-cell RNA sequencing and Spatial Transcriptomics | AGREE | we benchmarked it against two recent standalone computational algorithms, scGeneFit [8] and RankCorr [9] | The SMaSH paper explicitly benchmarks against RankCorr. [accepted_agree] |
| 10.1101/gr.275569.121 | 2021 | A machine learning method for the discovery of minimum marker gene combinations for cell type identification from single-cell RNA sequencing | AGREE | we compared it to two other marker gene selection tools ... RANKCORR | The paper explicitly compares NS-Forest v2.0 against RankCorr and other marker-gene selection tools. [accepted_agree] |
| 10.1186/s12859-022-04817-5 | 2022 | Detection of cell markers from single cell RNA-seq with sc2marker | AGREE | We evaluated the performances of competing methods ... RANKCORR [13] | The methods paper explicitly evaluates RankCorr as one of the competing methods. [accepted_agree] |
| 10.1186/s12859-022-04860-2 | 2022 | SMaSH: a scalable, general marker gene identification framework for single-cell RNA-sequencing | AGREE | we benchmarked it against two recent standalone computational algorithms, scGeneFit [6] and RankCorr [7] | The SMaSH methods paper explicitly benchmarks against RankCorr. [accepted_agree] |
| 10.1186/s13059-024-03174-1 | 2024 | Gene panel selection for targeted spatial transcriptomics | AGREE | We also compared gpsFISH with two previously published classification-based gene selection methods: RankCorr and scGenefit. | The gpsFISH methods paper explicitly compares its approach against RankCorr. [accepted_agree] |
| 10.1186/s13059-024-03183-0 | 2024 | A comparison of marker gene selection methods for single-cell RNA sequencing data | AGREE | "In this paper, we benchmark 59 computational methods for selecting marker genes in scRNA-seq data." | This is a computational comparison paper explicitly benchmarking marker-gene selection methods, which matches |

| DOI | Year | Title | Audit decision | Evidence (verbatim quote) | Audit reason / validation status |
|---|---|---|---|---|---|
| | | | | | COMP-BENCH for RankCorr. [accepted_agree] |
| 10.1186/s44330-024-00015-2 | 2024 | Discovery of optimal cell type classification marker genes from single cell RNA sequencing data | AGREE | four other methods: COMET [22], RankCorr [23], scGeneFit [24], and MarkerMap [25] were considered | The methods paper explicitly considers RankCorr as one of the comparator methods. [accepted_agree] |
| 10.21203/rs.3.rs-119885/v1 | 202 | Single-Cell Manifold Preserving Feature Selection (SCMER) | AGREE | "For comparison, we used ... RankCorr ..." | The evidence explicitly lists RankCorr among comparison methods, so COMP-BENCH is supported. [accepted_agree] |
| 10.7554/elife.107531.1 | 2025 | CellCover Defines Marker Gene Panels Capturing Developmental Progression in Neocortical Neural Stem Cell Identity | AGREE | We compare CellCover with leading marker gene selection algorithms ... RankCorr [21] ... to evaluate their effectiveness | This methods paper explicitly compares CellCover against RankCorr and other marker-gene selection algorithms. [accepted_agree] |

#### geneBasis (11)

| DOI | Year | Title | Audit decision | Evidence (verbatim quote) | Audit reason / validation status |
|---|---|---|---|---|---|
| 10.1038/s41467-023-37392-1 | 2023 | Predictive and robust gene selection for spatial transcriptomics | AGREE | we tested the recently proposed GeneBasis method41 | PERSIST is explicitly compared with GeneBasis in the benchmark experiments. [accepted_agree] |
| 10.1038/s41598-026-39168-1 | 2026 | CiCLoDS: Joint cell clustering and gene selection for single-cell spatial transcriptomics | AGREE | we benchmarked CiCLoDS against three competing methods: geneBasis6, scDCC9, and PCA | The CiCLoDS paper explicitly benchmarks against geneBasis. [accepted_agree] |
| 10.1093/bib/bbad389 | 2023 | FEED: a feature selection method based on gene expression decomposition for single cell clustering | AGREE | "We compared the performance of FEED with seven state-of-the-art gene selection methods ... including ... geneBasis ..." | The FEED paper explicitly compares performance against other gene-selection methods including geneBasis, which supports COMP-BENCH. [accepted_agree] |
| 10.1093/bib/bbaf478 | 2025 | Multi-view gene panel characterization for spatially resolved omics | AGREE | "Panel selection using geneBasis ... all parameters were set to default values ..." | This panel-comparison paper explicitly includes a geneBasis-derived panel in its evaluation framework. [accepted_agree] |
| 10.1093/bioinformatics/btad216 | 2023 | CellBRF: a feature selection method for single-cell clustering using cell balance and random forest | AGREE | We compare CellBRF with six state-of-the-art feature selection methods: ... geneBasisR | The CellBRF paper explicitly compares against geneBasisR. [accepted_agree] |

| DOI | Year | Title | Audit decision | Evidence (verbatim quote) | Audit reason / validation status |
|---|---|---|---|---|---|
| 10.1101/2025.07.14.664825 | 2025 | Identifying the Minimal Number of Protein Markers for Cell Type Annotation Using MiniMarS | AGREE | including ... geneBasisR (Missarova et al., 2021) | The MiniMarS paper explicitly compares marker-identification methods and names geneBasisR among them. [accepted_agree] |
| 10.1101/2025.10.02.680117 | 2025 | scGPD: single-cell informed gene panel design for targeted spatial transcriptomics | AGREE | we applied scGPD and three baseline gene selection methods: ... geneBasis [28] | The scGPD paper explicitly benchmarks against geneBasis as a baseline panel-selection method. [accepted_agree] |
| 10.1101/gr.280539.125 | 2025 | Label-free selection of marker genes in single-cell and spatial transcriptomics with geneCover | AGREE | We compared geneCover with five other methods: geneBasis, PERSIST, SCMER, and DUBStepR | geneCover explicitly benchmarks against geneBasis as one of the label-free comparison methods. [accepted_agree] |
| 10.1109/tcbbio.2025.3609721 | 2025 | Knowledge-Guided Gene Panel Selection for Label-Free Single-Cell RNA-Seq Data: A Reinforcement Learning Perspective | AGREE | "Our comparative analysis evaluated RIGPS against seven widely used baselines ... geneBasis [25] ..." | The packet explicitly describes comparative evaluation against multiple baselines including geneBasis, which supports COMP-BENCH. [accepted_agree] |
| 10.1186/s12859-025-06112-5 | 2025 | Redefining the high variable genes by optimized LOESS regression with positive ratio | AGREE | including ... genebasisR [11] | The GLP paper explicitly compares performance with geneBasisR. [accepted_agree] |
| 10.1186/s13059-024-03185-y | 2024 | scGIST: gene panel design for spatial transcriptomics with prioritized gene sets | AGREE | We ran scGIST as well as scGeneFit, geneBasis, and SMaSH ... and calculated their accuracy | The scGIST paper explicitly runs geneBasis as a baseline across datasets and panel sizes. [accepted_agree] |

#### PERSIST (8)

| DOI | Year | Title | Audit decision | Evidence (verbatim quote) | Audit reason / validation status |
|---|---|---|---|---|---|
| 10.1038/s41467-025-61476-9 | 2025 | Prioritizing perturbation-responsive gene patterns using interpretable deep learning | AGREE | we also evaluate PERSIST45, a recently proposed deep-learning-based method for gene selection | The paper explicitly benchmarks River against PERSIST in simulation comparisons. [accepted_agree] |
| 10.1038/s41540-024-00339-3 | 2024 | MarkerMap: nonlinear marker selection for single-cell studies | AGREE | We benchmark MarkerMap against existing marker selection approaches and related methodologies from the wider explainable machine learning literature. | This single-cell marker-selection methods paper explicitly benchmarks MarkerMap against existing marker-selection approaches, so COMP-BENCH is appropriate. [accepted_agree] |
| 10.1038/s41592-024-02496-z | 2024 | Probe set selection for targeted spatial transcriptomics | AGREE | We assessed Spapros performance against ten recently | This probe-set-selection methods paper explicitly benchmarks |

| DOI | Year | Title | Audit decision | Evidence (verbatim quote) | Audit reason / validation status |
| --- | --- | --- | --- | --- | --- |
| | | | | proposed gene selection methods ... in a large-scale benchmark using our evaluation pipeline. | Spapros against multiple gene-selection methods, so the named authoritative cited tool belongs in COMP-BENCH. [accepted_agree] |
| 10.1101/2025.04.03.646459 | 2025 | SpatialAgent: An autonomous AI agent for spatial biology | AGREE | For gene panel design, it was benchmarked against ... Persist (32) | The SpatialAgent paper explicitly benchmarks its gene-panel design against PERSIST. [accepted_agree] |
| 10.1101/2025.10.02.680117 | 2025 | scGPD: single-cell informed gene panel design for targeted spatial transcriptomics | AGREE | we applied scGPD and three baseline gene selection methods: PERSIST [26] | The scGPD paper explicitly benchmarks against PERSIST as a baseline gene-panel method. [accepted_agree] |
| 10.1101/2025.10.08.681071 | 2025 | Optimal Gene Panel Selection for Targeted Spatial Transcriptomics Experiments | AGREE | We benchmarked the performance of ReconST against seven existing methods: ... PERSIST [34] | ReconST is benchmarked against PERSIST and other existing methods for targeted spatial panel selection. [accepted_agree] |
| 10.1101/gr.280539.125 | 2025 | Label-free selection of marker genes in single-cell and spatial transcriptomics with geneCover | AGREE | We compared geneCover with five other methods: geneBasis, PERSIST, SCMER, and DUBStepR ... | The geneCover paper explicitly benchmarks against PERSIST. [accepted_agree] |
| 10.48550/arxiv.2512.17678 | 2025 | You Only Train Once: Differentiable Subset Selection for Omics Data | AGREE | "We benchmark our model against a suite of widely-used feature selection methods and the current state-of-the-art ..." | This is a computational subset-selection benchmarking paper, and the evidence explicitly states the model was benchmarked against widely used feature selection methods. [accepted_agree] |

#### SMaSH (8)

| DOI | Year | Title | Audit decision | Evidence (verbatim quote) | Audit reason / validation status |
| --- | --- | --- | --- | --- | --- |
| 10.1016/j.compbiomed.2023.106634 | 2023 | scMAGS: Marker gene selection from scRNA-seq data for spatial transcriptomics studies | AGREE | the performance of scMAGS is compared to that of the existing methods scGeneFit, SMaSH, and COSG | The scMAGS paper explicitly compares performance against SMaSH. [accepted_agree] |
| 10.1016/j.jbi.2023.104510 | 2023 | MAGNETO: Cell type marker panel generator from single-cell transcriptomic data | AGREE | "We compare the performance of our framework against ... SMaSH ..." | The packet explicitly says MAGNETO was compared against SMaSH and other approaches. [accepted_agree] |
| 10.1016/j.swevo.2025.102108 | 2025 | A multi-objective metaheuristic approach for generating cell type | AGREE | "The performance of scMOCTYM is evaluated ... and compared | The packet explicitly states the method was compared against seven state-of-the-art methods, |

| DOI | Year | Title | Audit decision | Evidence (verbatim quote) | Audit reason / validation status |
|---|---|---|---|---|---|
| | | marker panels from single-cell transcriptomics | | against seven state-of-the-art methods ..." | supporting COMP-BENCH for an authoritatively cited tool. [accepted_agree] |
| 10.1038/s41467-023-37392-1 | 2023 | Predictive and robust gene selection for spatial transcriptomics | AGREE | Our experiments compare PERSIST to several widely used and state-of-the-art gene selection methods ... SMaSH | PERSIST is explicitly benchmarked against SMaSH and other gene selection methods. [accepted_agree] |
| 10.1038/s41592-024-02496-z | 2024 | Probe set selection for targeted spatial transcriptomics | AGREE | We assessed Spapros performance against ten recently proposed gene selection methods ... in a large-scale benchmark using our evaluation pipeline. | This probe-set-selection methods paper explicitly benchmarks Spapros against multiple recently proposed gene-selection methods, including the authoritative cited tool set. [accepted_agree] |
| 10.1101/2023.05.29.542760 | 2023 | Boosting Single-Cell RNA Sequencing Analysis with Simple Neural Attention | AGREE | We compared scANNA-selected genes against markers selected by ... SMaSH | The scANNA paper explicitly compares its selected genes against SMaSH. [accepted_agree] |
| 10.1186/s13059-024-03183-0 | 2024 | A comparison of marker gene selection methods for single-cell RNA sequencing data | AGREE | "In this paper, we benchmark 59 computational methods for selecting marker genes in scRNA-seq data." | The paper is an explicit benchmarking study of marker-gene selection methods, so COMP-BENCH is appropriate for SMaSH. [accepted_agree] |
| 10.1186/s13059-024-03185-y | 2024 | scGIST: gene panel design for spatial transcriptomics with prioritized gene sets | AGREE | We ran scGIST as well as scGeneFit, geneBasis, and SMaSH ... and calculated their accuracy | The scGIST paper explicitly evaluates SMaSH as a baseline over multiple panel sizes. [accepted_agree] |

#### COSG (7)

| DOI | Year | Title | Audit decision | Evidence (verbatim quote) | Audit reason / validation status |
|---|---|---|---|---|---|
| 10.1016/j.compbiomed.2023.106634 | 2023 | scMAGS: Marker gene selection from scRNA-seq data for spatial transcriptomics studies | AGREE | "the performance of scMAGS is compared to that of ... COSG." | The paper explicitly compares scMAGS to COSG and other methods, which supports COMP-BENCH. [accepted_agree] |
| 10.1038/s41540-024-00339-3 | 2024 | MarkerMap: nonlinear marker selection for single-cell studies | AGREE | we quantified the quality of the learnt markers against those learnt through classical ... methods like COSG38 | MarkerMap explicitly quantifies marker quality against COSG and other classical methods. [accepted_agree] |
| 10.1038/s41598-024-63492-z | 2024 | CellMarkerPipe: cell marker identification and evaluation | AGREE | "... COSG ... to be compared and benchmarked in this paper." | The packet explicitly says COSG is among the prioritized tools compared and benchmarked in the |

| DOI | Year | Title | Audit decision | Evidence (verbatim quote) | Audit reason / validation status |
|---|---|---|---|---|---|
| | | pipeline in single cell transcriptomes | | | paper. [accepted_agree] |
| 10.1038/s42003-023-04820-3 | 2023 | MASI enables fast model-free standardization and integration of single-cell transcriptomics data | AGREE | we benchmarked 12 DE tests ... including ... COSG48 | The MASI paper explicitly benchmarks DE methods including COSG. [accepted_agree] |
| 10.1093/bib/bbae626 | 2024 | CosGeneGate selects multi-functional and credible biomarkers for single-cell analysis | AGREE | "including ... COSG ... and CosGeneGate." | The paper explicitly compares prediction accuracy using marker genes from COSG and other models. [accepted_agree] |
| 10.1093/nsr/nwaf030 | 2025 | DSCT: a novel deep-learning framework for rapid and accurate spatial transcriptomic cell typing | AGREE | ACOSG demonstrated optimal performance compared with the others | The DSCT paper explicitly compares performance across gene-selection algorithms and names COSG. [accepted_agree] |
| 10.1101/2025.10.08.681071 | 2025 | Optimal Gene Panel Selection for Targeted Spatial Transcriptomics Experiments | AGREE | We benchmarked the performance of ReconST against seven existing methods: ... COSG [36] | ReconST is explicitly benchmarked against COSG as one of the existing methods. [accepted_agree] |

#### SCMER (7)

| DOI | Year | Title | Audit decision | Evidence (verbatim quote) | Audit reason / validation status |
|---|---|---|---|---|---|
| 10.1038/s41467-023-44570-8 | 2024 | MarsGT: Multi-omics analysis for rare population inference using single-cell graph transformer | AGREE | we compared it with three clustering-like tools ... and SCMER1 | The MarsGT paper explicitly compares performance with SCMER. [accepted_agree] |
| 10.1038/s41467-024-46773-z | 2024 | DELVE: feature selection for preserving biological trajectories in single-cell data | AGREE | "We compared DELVE to eleven other feature selection approaches ..." | DELVE is presented as a feature-selection method and explicitly compared against other approaches, which supports COMP-BENCH for SCMER. [accepted_agree] |
| 10.1038/s41592-024-02496-z | 2024 | Probe set selection for targeted spatial transcriptomics | AGREE | general variation recovery (SelfE, SCMER, geneBasis, PCA, scPNMF, triku, unsupervised PERSIST (PERSISTus)) | The Spapros paper explicitly characterizes SCMER within its large-scale benchmark of gene selection methods. [accepted_agree] |

| DOI | Year | Title | Audit decision | Evidence (verbatim quote) | Audit reason / validation status |
|---|---|---|---|---|---|
| 10.1093/bib/bbad485 | 2023 | <i>methylClass</i>: an R package to construct DNA methylation-based classification models | AGREE | we used ... SCMER features | The methylation-classification paper explicitly evaluates SCMER-derived features against alternatives. [accepted_agree] |
| 10.1101/gr.280539.125 | 2025 | Label-free selection of marker genes in single-cell and spatial transcriptomics with geneCover | AGREE | We compared geneCover with five other methods: geneBasis, PERSIST, SCMER, and DUBStepR | The geneCover paper explicitly benchmarks against SCMER and other label-free methods. [accepted_agree] |
| 10.1186/s12864-025-11319-0 | 2025 | scMFG: a single-cell multi-omics integration method based on feature grouping | AGREE | compared our method with three clustering-like tools: GiniClust [45], RaceID [46], and SCMER [47] | The evaluation explicitly compares the proposed method against SCMER. [accepted_agree] |
| 10.1186/s13059-021-02548-z | 2021 | geneBasis: an iterative approach for unsupervised selection of targeted gene panels from scRNA-seq | AGREE | we applied geneBasis, as well as SCMER and scGeneFit (Methods), to four diverse systems | The paper explicitly evaluates geneBasis against SCMER and scGeneFit on several datasets. [accepted_agree] |

#### scPNMF (6)

| DOI | Year | Title | Audit decision | Evidence (verbatim quote) | Audit reason / validation status |
|---|---|---|---|---|---|
| 10.1038/s41592-024-02496-z | 2024 | Probe set selection for targeted spatial transcriptomics | AGREE | The benchmark passage explicitly lists "scPNMF" among assessed methods. | The packet explicitly places scPNMF among methods assessed in a large benchmark, supporting COMP-BENCH. [accepted_agree] |
| 10.1038/s41592-025-02624-3 | 2025 | Feature selection methods affect the performance of scRNA-seq data integration and querying | AGREE | "We evaluate the performance of variants of over 20 feature selection methods ..." | This is a computational benchmarking paper evaluating many feature selection methods, which supports COMP-BENCH for scPNMF. [accepted_agree] |
| 10.1038/s42003-025-07872-9 | 2025 | Infusing structural assumptions into dimensionality reduction for single-cell RNA sequencing data to identify small gene sets | AGREE | We also compared the BAE to ... scPNMF65, in terms of gene selection | The BAE paper explicitly compares its gene selection to scPNMF. [accepted_agree] |
| 10.1109/bibm62325.2024.10821945 | 2024 | Identification of cell-type-specific genes associated with autism by integrating gene expression and alternative polyadenylation profiles | AGREE | To further compare the performance of the RFECV set and the scPNMF set | The paper explicitly compares the RFECV feature set with the scPNMF feature set in a classification task. [accepted_agree] |
| 10.1186/s13059-024-03284-w | 2024 | scCDC: a computational method for gene-specific contamination | AGREE | the top 1000 variable genes identified by SeuratVST ... | The computational paper explicitly benchmarks feature-selection |

| DOI | Year | Title | Audit decision | Evidence (verbatim quote) | Audit reason / validation status |
|---|---|---|---|---|---|
| | | detection and correction in single-cell and single-nucleus RNA-seq data | | scPNMF ... were used for dimension reduction | pipelines including scPNMF. [accepted_agree] |
| 10.1186/s13059-026-03998-z | 2026 | Benchmarking LLM-based agents for single-cell omics analysis | AGREE | "Integrated 50 benchmarking tasks of single-cell omics analysis ... employing core analytical tools ..." | The packet describes a benchmarking paper covering many single-cell analysis tasks using core analytical tools, which supports a comparison/benchmark classification. [accepted_agree] |

#### NS-Forest v2 (5)

| DOI | Year | Title | Audit decision | Evidence (verbatim quote) | Audit reason / validation status |
|---|---|---|---|---|---|
| 10.1093/bib/bbae626 | 2024 | CosGeneGate selects multi-functional and credible biomarkers for single-cell analysis | AGREE | including genes from experts ... COSG [29], NSForest [27], scGeneFit [28] | CosGeneGate explicitly compares prediction accuracy using marker genes from NSForest and other models. [accepted_agree] |
| 10.1186/s13059-024-03410-8 | 2024 | scCTS: identifying the cell type-specific marker genes from population-level single-cell RNA-seq | AGREE | "We compare with ... NS-Forest ..." | The paper explicitly compares scCTS with NS-Forest and other methods, supporting COMP-BENCH. [accepted_agree] |
| 10.1186/s44330-024-00015-2 | 2024 | Discovery of optimal cell type classification marker genes from single cell RNA sequencing data | AGREE | Two of the methods were previously directly compared with NS-Forest v2.0 [13] | The methods paper directly compares NS-Forest v4.0 with earlier comparator methods and explicitly references prior direct comparison with NS-Forest v2.0. [accepted_agree] |
| 10.1371/journal.pone.0275070 | 2022 | Machine learning for cell type classification from single nucleus RNA sequencing data | AGREE | As a benchmark for comparison, we compared the F-beta classification scores of the top performing ML method, multinomial logistic regression, with those obtained using the minimum set of marker genes produced by NS-Forest [21] | This is a computational paper that explicitly benchmarks its top method against NS-Forest outputs. [accepted_agree] |
| 10.3389/fgene.2025.1583081 | 2025 | High-precision lung cancer subtype diagnosis on imbalanced exosomal data via Exo-LCClassifier | AGREE | the final Exo-LCClassifier model was compared with three methods (NS-Forest, SPLR, and ZIPLDA) | The final model is explicitly compared with NS-Forest as a benchmark method. [accepted_agree] |

#### Spapros (4)

| DOI | Year | Title | Audit decision | Evidence (verbatim quote) | Audit reason / validation status |
| --- | --- | --- | --- | --- | --- |
| 10.1093/bib/bbaf478 | 2025 | Multi-view gene panel characterization for spatially resolved omics | AGREE | "Panel was selected following Spapros's vignette (n = 200)." | This methods paper characterizes and compares multiple panel-construction approaches, and one panel was explicitly selected using Spapros. [accepted_agree] |
| 10.1093/bioinformatics/btaf194 | 2025 | Refinement strategies for Tangram for reliable single-cell to spatial mapping | AGREE | "We evaluated Tangram trained on ... (5) Spapros genes. We found that Spapros performed best ..." | The paper explicitly evaluates Tangram using several gene-set choices including Spapros genes, supporting comparison/benchmark treatment. [accepted_agree] |
| 10.1101/2025.04.03.646459 | 2025 | SpatialAgent: An autonomous AI agent for spatial biology | AGREE | To evaluate SpatialAgent's ability to autonomously design gene panels, we benchmarked its performance against computational baselines, human scientists, and a hybrid setting. | This is a computational spatial-biology methods paper with explicit benchmarking of gene-panel design performance against computational baselines; under the authoritative citing-set assumption, the conservative final label is COMP-BENCH. [accepted_agree] |
| 10.1101/2025.10.08.681071 | 2025 | Optimal Gene Panel Selection for Targeted Spatial Transcriptomics Experiments | AGREE | We benchmarked the performance of ReconST against seven existing methods: ... Spapros [15] | ReconST is explicitly benchmarked against Spapros as one of the existing targeted spatial panel-selection methods. [accepted_agree] |

#### gpsFISH (2)

| DOI | Year | Title | Audit decision | Evidence (verbatim quote) | Audit reason / validation status |
| --- | --- | --- | --- | --- | --- |
| 10.1093/bib/bbaf478 | 2025 | Multi-view gene panel characterization for spatially resolved omics | AGREE | "We conducted panel selection by following gpsFISH's vignette ..." | This methods paper characterizes and compares panels and explicitly says panel selection was conducted by following gpsFISH's vignette, supporting tool inclusion in a comparative framework. [accepted_agree] |
| 10.1109/tcbbio.2025.3609721 | 2025 | Knowledge-Guided Gene Panel Selection for Label-Free Single-Cell RNA-Seq Data: A Reinforcement Learning Perspective | AGREE | Our comparative analysis evaluated RIGPS against seven widely used baselines ... gpsFISH [48] | The comparative analysis explicitly includes gpsFISH as one of the evaluated baselines. [accepted_agree] |

#### scGIST (2)

| DOI | Year | Title | Audit decision | Evidence (verbatim quote) | Audit reason / validation status |
|---|---|---|---|---|---|
| 10.1101/2025.10.02.680117 | 2025 | scGPD: single-cell informed gene panel design for targeted spatial transcriptomics | AGREE | we applied scGPD and three baseline gene selection methods: PERSIST [26], scGIST [27], and geneBasis [28] | The scGPD paper explicitly applies scGIST as one of the baseline panel-selection methods. [accepted_agree] |
| 10.1109/tcbbio.2025.3609721 | 2025 | Knowledge-Guided Gene Panel Selection for Label-Free Single-Cell RNA-Seq Data: A Reinforcement Learning Perspective | AGREE | Our comparative analysis evaluated RIGPS against seven widely used baselines ... scGIST | The comparative analysis explicitly lists scGIST as one of the evaluated baselines. [accepted_agree] |

### COMP (n = 181)

*Computational methods or analysis paper that cited the tool without being an experimental deployment or benchmark.*

#### scGeneFit (30)

| DOI | Year | Title | Audit decision | Evidence (verbatim quote) | Audit reason / validation status |
|---|---|---|---|---|---|
| 10.1016/j.apsusc.2021.151061 | 2021 | Dissociation of air pollutants on the uniform surface of pentagonal BeP2 | AGREE | the investigation was carried out using density functional theory (DFT) for the dissociation of noxious gas molecules | This is a computational materials-science paper unrelated to single-cell or spatial transcriptomics; within this audit schema COMP is the conservative negative label for a computational paper with no relevant tool relation. [accepted_agree] |
| 10.1016/j.cels.2021.09.004 | 2021 | Cell-type modeling in spatial transcriptomics data elucidates spatially variable colocalization and communication between cell-types in mouse brain | AGREE | "We developed a neural network model ... STANN ..." | This is a computational spatial transcriptomics cell-type assignment paper, and the packet provides no explicit scGeneFit comparison or use evidence. [accepted_agree] |
| 10.1016/j.jbi.2023.104510 | 2023 | MAGNETO: Cell type marker panel generator from single-cell transcriptomic data | AGREE | "we propose a fully-automatic framework called MAGNETO ..." | This is a computational framework paper introducing MAGNETO, and the packet provides no explicit scGeneFit comparison or use evidence. [accepted_agree] |
| 10.1038/s41540-024-00339-3 | 2024 | MarkerMap: nonlinear marker selection for single-cell studies | DISAGREE | We benchmark MarkerMap's competitive performance against previously published approaches on real single cell gene expression data sets. | The packet says MarkerMap was benchmarked against published approaches, but the evidence does not explicitly mention scGeneFit. The conservative negative label is |

| DOI | Year | Title | Audit decision | Evidence (verbatim quote) | Audit reason / validation status |
|---|---|---|---|---|---|
| | | | | | COMP. [accepted_disagree] |
| 10.1038/s41592-024-02496-z | 2024 | Probe set selection for targeted spatial transcriptomics | DISAGREE | "We assessed Spapros performance against ten recently proposed gene selection methods ..." | The paper is a computational probe-selection methods paper, but the provided quote does not explicitly name scGeneFit among the benchmarked methods, so the conservative negative label is COMP. [accepted_disagree] |
| 10.1038/s41598-024-73356-1 | 2024 | scGAA: a general gated axial-attention model for accurate cell-type annotation of single-cell RNA-seq data | AGREE | we developed a new cell-type annotation model called scGAA | This is a computational scRNA-seq cell-type annotation paper; the packet does not explicitly benchmark or use scGeneFit. [accepted_agree] |
| 10.1038/s43588-024-00689-2 | 2024 | Biophysically interpretable inference of cell types from multimodal sequencing data | AGREE | we present meK-Means ... which integrates modalities and learns underlying, shared biophysical states | This is a computational multimodal inference paper, and the evidence does not explicitly benchmark or use scGeneFit. [accepted_agree] |
| 10.1093/bib/bbad424 | 2023 | Identifying phenotype-associated subpopulations through LP_SGL | AGREE | "We proposed LP_SGL ... to identify phenotype-associated subpopulations ..." | This is a computational subpopulation-identification paper introducing LP_SGL, and the packet provides no explicit scGeneFit comparison or use evidence. [accepted_agree] |
| 10.1093/bib/bbae018 | 2024 | scHybridBERT: integrating gene regulation and cell graph for spatiotemporal dynamics in single-cell clustering | AGREE | this study proposes the scHybridBERT architecture to conduct multi-view modeling of scRNA-seq data | This is a computational single-cell clustering paper, and the evidence does not explicitly benchmark or use scGeneFit. [accepted_agree] |
| 10.1093/bib/bbaf405 | 2025 | MuST: multiple-modality structure transformation for single-cell spatial transcriptomics | AGREE | "This paper introduces ... MuST, a novel methodology ..." | This is a computational spatial transcriptomics integration paper introducing MuST, and the packet provides no explicit scGeneFit comparison or use evidence. [accepted_agree] |

| DOI | Year | Title | Audit decision | Evidence (verbatim quote) | Audit reason / validation status |
|---|---|---|---|---|---|
| 10.1093/bib/bbaf478 | 2025 | Multi-view gene panel characterization for spatially resolved omics | DISAGREE | The evidence lists expert-curated, geneBasis, gpsFish, IO, Spapros, and o1 panels, but not scGeneFit. | The paper is computational and compares panels, but the provided evidence does not specifically support benchmarking of scGeneFit itself, so the conservative negative label is COMP. [accepted_disagree] |
| 10.1093/bib/bbag072 | 2026 | Imputing missing values in single-cell RNA-sequencing data: a statistical and machine learning-based approach | AGREE | we propose a novel method called single-cell dropout detection and imputation (scDDI) | This is a computational scRNA-seq imputation paper; the packet does not explicitly benchmark or use scGeneFit. [accepted_agree] |
| 10.1093/bioinformatics/btac263 | 2022 | Self-supervised learning of cell type specificity from immunohistochemical images | AGREE | We develop a method to predict cell type specificity of protein markers from unlabeled images | This is a computational method paper on predicting cell-type specificity from images; the packet does not explicitly benchmark or use scGeneFit. [accepted_agree] |
| 10.1101/2022.05.06.490859 | 2022 | Depth normalization for single-cell genomics count data | AGREE | Depth normalization for single-cell genomics count data | This is a computational normalization-methods paper for single-cell genomics; the packet does not explicitly benchmark or use scGeneFit. [accepted_agree] |
| 10.1101/2023.07.17.549344 | 2023 | Recurrent Composite Markers of Cell Types and States | AGREE | we developed Recurrent Composite Markers ... (RECOMBINE) | This is a computational marker-combination methods paper; the packet does not explicitly benchmark or use scGeneFit, so the conservative negative is COMP. [accepted_agree] |
| 10.1101/2024.04.24.590691 | 2024 | mastR: Marker Automated Screening Tool for multi-omics data | AGREE | We developed mastR, a novel computational tool for accurate marker identification from omics data. | This is a computational marker-identification methods paper; the packet does not explicitly benchmark or use scGeneFit, so the conservative negative is COMP. [accepted_agree] |
| 10.1101/2024.12.23.630040 | 2024 | CORTADO: Hill Climbing Optimization for Cell-Type Specific Marker Gene Discovery | AGREE | "we introduce CORTADO, a computational framework ..." | This is a computational marker-discovery paper introducing CORTADO, and the packet provides no explicit scGeneFit comparison or use evidence. [accepted_agree] |

| DOI | Year | Title | Audit decision | Evidence (verbatim quote) | Audit reason / validation status |
|---|---|---|---|---|---|
| 10.1101/2025.04.22.649977 | 2025 | spRefine Denoises and Imputes Spatial Transcriptomics with a Reference-Free Framework Powered by Genomic Language Model | AGREE | "we introduce spRefine, a deep learning framework ..." | This is a computational spatial transcriptomics denoising/imputation paper introducing spRefine, and the packet provides no explicit scGeneFit comparison or use evidence. [accepted_agree] |
| 10.1101/2025.06.27.661814 | 2025 | Fluctuation structure predicts genome-wide perturbation outcomes | AGREE | Here we present CIPHER ... a conceptual framework ... to predict transcriptome-wide perturbation outcomes | This is a computational perturbation-prediction paper; the packet does not explicitly benchmark or use scGeneFit, so the conservative negative is COMP. [accepted_agree] |
| 10.1101/gr.280539.125 | 2025 | Label-free selection of marker genes in single-cell and spatial transcriptomics with geneCover | AGREE | here, we introduce geneCover, a label-free combinatorial method that selects an optimal panel ... | This is a computational marker-panel selection paper for single-cell and spatial transcriptomics; the packet does not explicitly benchmark or use scGeneFit, so the conservative negative is COMP. [accepted_agree] |
| 10.1109/bibm62325.2024.10822193 | 2024 | MuST: Maximizing the Latent Capacity of Spatial Transcriptomics Data with Multi-modality Structure Transformation | AGREE | This paper introduces Multiple-modality Structure Transformation, named MuST | This is a computational spatial transcriptomics methods paper; the packet does not explicitly benchmark or use scGeneFit. [accepted_agree] |
| 10.1109/jbhi.2024.3370868 | 2024 | Learning Consistency and Specificity of Cells From Single-Cell Multi-Omic Data | AGREE | we propose a novel self-representation Learning-based Multi-omics data Integrative Clustering algorithm (sLMIC) | This is a computational single-cell multi-omics clustering paper; the packet provides no explicit scGeneFit benchmarking or targeted spatial panel-selection use. [accepted_agree] |
| 10.1111/biom.13926 | 2023 | Multiresolution Categorical Regression for Interpretable Cell-Type Annotation | AGREE | we propose a method for fitting the multinomial logistic regression model in high dimensions | This is a computational cell-type-annotation/statistical-methods paper; the packet does not explicitly benchmark or use scGeneFit. [accepted_agree] |
| 10.1145/3502871.3502881 | 2021 | Informative gene identification for Single-Cell RNA-Seq Data with Mutual Information based Firefly Algorithm | AGREE | In this paper, we perform supervised gene selection method based on firefly algorithm ... | This is a computational gene-selection paper for scRNA-seq; the packet does not explicitly benchmark or use scGeneFit, so the conservative negative is |

| DOI | Year | Title | Audit decision | Evidence (verbatim quote) | Audit reason / validation status |
|---|---|---|---|---|---|
| | | | | | COMP. [accepted_agree] |
| 10.1186/s12859-020-03625-z | 202 | A robust nonlinear low-dimensional manifold for single cell RNA-seq data | AGREE | We introduce the t-distributed Gaussian process latent variable model (tGPLVM) | This is a computational manifold-learning paper, and the evidence does not explicitly benchmark or use scGeneFit. [accepted_agree] |
| 10.1186/s13059-024-03183-0 | 2024 | A comparison of marker gene selection methods for single-cell RNA sequencing data | AGREE | In this paper, we benchmark 59 computational methods for selecting marker genes in scRNA-seq data. | This is a computational benchmark paper, but the provided evidence does not explicitly show scGeneFit among the benchmarked methods. The conservative label is COMP. [accepted_agree] |
| 10.12688/f1000research.122775.1 | 2022 | Spatial transcriptomics dimensionality reduction using wavelet bases | AGREE | we propose a gene expression dimensionality reduction algorithm that retains spatial structure. | This is a computational spatial-transcriptomics methods paper; the packet does not explicitly benchmark or use scGeneFit, so the conservative negative is COMP. [accepted_agree] |
| 10.1371/journal.pcbi.1009600 | 2022 | A copula based topology preserving graph convolution network for clustering of single-cell RNA-seq data | DISAGREE | Abstract discusses clustering of single-cell RNA-seq data; evidence only names the scGCConv package. | This is a computational clustering paper for single-cell RNA-seq rather than a spatial transcriptomics study, so COMP remains the conservative negative label. [accepted_disagree] |
| 10.48550/arxiv.2305.18962 | 2023 | Hyperbolic Diffusion Embedding and Distance for Hierarchical Representation Learning | AGREE | This paper presents a new method for hierarchical data embedding and distance. | This is a general hierarchical representation-learning paper, and the evidence does not explicitly benchmark or use scGeneFit. [accepted_agree] |
| 10.64898/2026.02.26.707870 | 2026 | PantheonOS: An Evolvable Multi-Agent Framework for Automatic Genomics Discovery | DISAGREE | combining e.g., HVG, DE, RF, SpaPROS or scGeneFit with fixed weighting schemes | PantheonOS mentions combining or selecting among methods including scGeneFit, but this is not an explicit benchmark or evaluation of scGeneFit itself. [accepted_disagree] |

#### COSG (29)

| DOI | Year | Title | Audit decision | Evidence (verbatim quote) | Audit reason / validation status |
|---|---|---|---|---|---|
| 10.1002/advs.202306329 | 2023 | Highly Accurate Estimation of Cell Type Abundance in Bulk Tissues | AGREE | a novel computational method named SCROAM is introduced | This is a computational deconvolution/cell-type abundance |

| DOI | Year | Title | Audit decision | Evidence (verbatim quote) | Audit reason / validation status |
|---|---|---|---|---|---|
| | | Based on Single-Cell Reference and Domain Adaptive Matching | | | estimation paper; the packet does not explicitly benchmark or use COSG. [accepted_agree] |
| 10.1002/advs.202505021 | 2025 | SCRIPT: Predicting Single-Cell Long-Range Cis-Regulation Based on Pretrained Graph Attention Networks | AGREE | SCRIPT ... is presented for inferring single-cell CRRs from transcriptomic and chromatin accessibility data. | This is a computational single-cell cis-regulation prediction paper; the packet does not explicitly benchmark or use COSG. [accepted_agree] |
| 10.1007/s12539-025-00762-y | 2025 | Clustering Single-Cell RNA-Seq Data with Low-Rank Matrix Factorization and Local Graph Regularization | AGREE | we introduce a novel clustering algorithm that integrates low-rank matrix decomposition with local graph regularization (LRMGC) | This is a computational single-cell clustering paper; the packet does not explicitly benchmark or use COSG, so the conservative negative is COMP. [accepted_agree] |
| 10.1016/j.cj.2026.02.021 | 2026 | PhytoCell: An ensemble learning framework for identifying cell states in plant scRNA-seq data | AGREE | PhytoCell ... combines feature selection engineering with machine learning to uncover cell markers | This is a computational plant scRNA-seq marker-identification and annotation paper; the packet does not explicitly benchmark or use COSG. [accepted_agree] |
| 10.1016/j.knosys.2025.114149 | 2025 | Dual-consistency graph-spectral embedding joint learning for multi-view single-cell clustering | AGREE | we propose a dual-consistency graph-spectral embedding joint learning multi-view clustering method named DcGSE | This is a computational single-cell clustering paper; the packet does not explicitly benchmark or use COSG, so the conservative negative is COMP. [accepted_agree] |
| 10.1038/s41398-025-03243-4 | 2025 | Systematic dissection of pleiotropic loci and critical regulons in excitatory neurons and microglia relevant to neuropsychiatric and ocular diseases | AGREE | we integrated bulk and single-cell multiomic profiles with GWAS summary statistics | This is a computational multi-omics/GWAS integration paper, and the evidence does not explicitly benchmark or use COSG. [accepted_agree] |
| 10.1038/s41467-026-70645-3 | 2026 | Ultra-precision deconvolution of spatial transcriptomics decodes immune heterogeneity and fate-defining programs in tissues | AGREE | We present UCASpatial, an ultra-precision spatial transcriptomics deconvolution algorithm | This is a computational spatial deconvolution paper, and the evidence does not explicitly benchmark or use COSG. [accepted_agree] |
| 10.1038/s41592-024-02496-z | 2024 | Probe set selection for targeted spatial transcriptomics | DISAGREE | "We assessed Spapros performance against ten recently proposed gene selection methods ..." | This is a computational probe-selection methods paper, but the provided evidence does not explicitly benchmark COSG, so the conservative negative label is COMP rather than COMP-BENCH. [accepted_disagree] |

| DOI | Year | Title | Audit decision | Evidence (verbatim quote) | Audit reason / validation status |
|---|---|---|---|---|---|
| 10.1038/s41598-024-78076-0 | 2024 | Embracing the informative missingness and silent gene in analyzing biologically diverse samples | AGREE | We developed the ABDS tool suite specifically for analyzing biologically diverse samples. | This is a computational bioinformatics tool-suite paper; the packet does not explicitly benchmark or use COSG. [accepted_agree] |
| 10.1038/s41598-025-27732-0 | 2025 | Assessment of chemical risk factor for breast cancer based on gene expression profiles | AGREE | the objective of our study was to characterize through a computational approach, a global gene expression profile | This is a computational toxicogenomics risk-assessment paper, and the evidence does not explicitly benchmark or use COSG. [accepted_agree] |
| 10.1038/s43587-025-01027-5 | 2025 | Integrating polygenic signals and single-cell multiomics identifies cell-type-specific regulomes critical for immune- and aging-related diseases | AGREE | Here, we present scMORE, a method that integrates single-cell transcriptomes and chromatin accessibility with GWAS summary statistics | This is a computational single-cell multi-omics integration paper; the packet does not explicitly benchmark or use COSG. [accepted_agree] |
| 10.1073/pnas.2517283123 | 2026 | Dissecting spatial patterning and signaling with directional diffusion in spatial multi-omics | AGREE | we present SpaDDM, a spatial multi-omics integration framework | This is a computational spatial multi-omics integration paper; the packet does not explicitly benchmark or use COSG. [accepted_agree] |
| 10.1093/bfgp/elad004 | 2023 | SCMcluster: a high-precision cell clustering algorithm integrating marker gene set with single-cell RNA sequencing data | AGREE | we propose a high-precision single-cell clustering algorithm-SCMcluster | This is a computational single-cell clustering methods paper; the packet does not explicitly benchmark or use COSG. [accepted_agree] |
| 10.1093/bib/bbac585 | 2023 | scGMAAE: Gaussian mixture adversarial autoencoders for diversification analysis of scRNA-seq data | AGREE | we design and develop a model named Gaussian mixture adversarial autoencoders (scGMAAE) | This is a computational scRNA-seq analysis method paper; the packet does not explicitly benchmark or use COSG. [accepted_agree] |
| 10.1093/bib/bbac625 | 2023 | scDCCA: deep contrastive clustering for single-cell RNA-seq data based on auto-encoder network | AGREE | Here, we propose a new deep contrastive clustering algorithm called scDCCA. | This is a computational single-cell clustering paper; the packet does not explicitly benchmark or use COSG, so the conservative negative is COMP. [accepted_agree] |
| 10.1093/bib/bbae422 | 2024 | ReCIDE: robust estimation of cell type proportions by integrating single-reference-based deconvolutions | AGREE | "In this study, we introduce ... ReCIDE ..." | This is a computational deconvolution framework paper introducing ReCIDE, and the packet provides no explicit COSG comparison or use evidence. [accepted_agree] |

| DOI | Year | Title | Audit decision | Evidence (verbatim quote) | Audit reason / validation status |
|---|---|---|---|---|---|
| 10.1093/bib/bbae572 | 2024 | Identifying cell types by lasso-constraint regularized Gaussian graphical model based on weighted distance penalty | AGREE | "In this paper, we propose a novel algorithm framework named WLGG." | This is a computational cell-type identification paper introducing WLGG, and the packet provides no explicit COSG comparison or use evidence. [accepted_agree] |
| 10.1093/bib/bbae703 | 2024 | Addressing scalability and managing sparsity and dropout events in single-cell representation identification with ZIGACL | AGREE | To tackle these challenges, the Zero-Inflated Graph Attention Collaborative Learning (ZIGACL) method has been developed. | This is a computational scRNA-seq representation-learning paper; the packet does not explicitly benchmark or use COSG. [accepted_agree] |
| 10.1093/bib/bbaf193 | 2025 | Multi-view clustering for single-cell RNA-seq data based on graph fusion | AGREE | "Here, we introduce scMCGF, a new multi-view clustering algorithm based on graph fusion." | This is a computational clustering paper introducing scMCGF, and the packet provides no explicit COSG comparison or use evidence. [accepted_agree] |
| 10.1093/bib/bbaf675 | 2025 | scHLens: a web server for hierarchically and interactively exploring single cell RNA-seq data | AGREE | "Here, we develop a hierarchical and interactive web server named scHLens." | This is a computational web-server paper for exploring scRNA-seq data, and the packet provides no explicit COSG comparison or use evidence. [accepted_agree] |
| 10.1093/bib/bbag094 | 2026 | scSCCNIA: similarity matrix based contrastive clustering with neighbor information aggregation for single-cell RNA sequencing data | AGREE | we propose a novel contrastive clustering framework called scSCCNIA | This is a computational scRNA-seq clustering methods paper; the packet does not explicitly benchmark or use COSG. [accepted_agree] |
| 10.1101/2023.07.05.547797 | 2023 | ABDS: tool suite for analyzing biologically diverse samples | AGREE | "We develop ABDS tool suite tailored specifically to analyzing biologically diverse samples." | This is a computational tool-suite paper, and the packet provides no explicit COSG comparison or use evidence. [accepted_agree] |
| 10.1101/2023.12.27.573479 | 2023 | ASCT: automatic single-cell toolbox in julia | AGREE | "ASCT ... is an automatic single-cell toolbox for analyzing single-cell RNA-Seq data." | This is a computational single-cell toolbox paper, and the packet does not provide explicit COSG comparison or use evidence. [accepted_agree] |
| 10.1101/2024.12.23.630040 | 2024 | CORTADO: Hill Climbing Optimization for Cell-Type Specific Marker Gene Discovery | DISAGREE | CORTADO was compared to four baseline methods across seven datasets using these metrics. | CORTADO is clearly a computational benchmarking paper, but the evidence does not explicitly identify COSG as one of the four baselines. The conservative negative label is |

| DOI | Year | Title | Audit decision | Evidence (verbatim quote) | Audit reason / validation status |
|---|---|---|---|---|---|
| | | | | | COMP. [accepted_disagree] |
| 10.1101/2025.03.05.641709 | 2025 | Leveraging cell-type specificity and similarity improves single-cell eQTL fine-mapping | AGREE | we propose a novel empirical Bayesian model named CASE ... to improve statistical power in eQTL fine-mapping | This is a computational single-cell eQTL fine-mapping paper; the packet does not explicitly benchmark or use COSG. [accepted_agree] |
| 10.1101/2025.11.18.689140 | 2025 | CBKMR: A Copula-based Bayesian Kernel Machine Regression Framework for Optimal Marker Detection in Omics Data | AGREE | We propose a copula-based Bayesian kernel machine regression (CBKMR) model ... | This is a computational omics marker-detection paper; the packet does not explicitly benchmark or use COSG, so the conservative negative is COMP. [accepted_agree] |
| 10.1101/2025.11.26.690796 | 2025 | LARIS enables accurate and efficient ligand and receptor interaction analysis in spatial transcriptomics | AGREE | we present LARIS ... an accurate and scalable method that identifies cell type-specific and spatially restricted ligand-receptor interactions | This is a computational spatial-transcriptomics ligand-receptor methods paper; the packet does not explicitly benchmark or use COSG, so the conservative negative is COMP. [accepted_agree] |
| 10.1186/s13059-024-03183-0 | 2024 | A comparison of marker gene selection methods for single-cell RNA sequencing data | DISAGREE | In this paper, we benchmark 59 computational methods for selecting marker genes in scRNA-seq data. | This is clearly a computational benchmark paper, but the provided evidence does not explicitly show COSG among the benchmarked methods. The conservative negative label is COMP. [accepted_disagree] |
| 10.1371/journal.pcbi.1014110 | 2026 | Clustering single-cell multi-omics data via weighted distance penalty and adaptive consistent graph regularization | AGREE | we introduce scWDAC ... a novel clustering method for single-cell multi-omics data. | This is a computational single-cell multi-omics clustering methods paper; the packet does not explicitly benchmark or use COSG. [accepted_agree] |

#### NS-Forest v2 (20)

| DOI | Year | Title | Audit decision | Evidence (verbatim quote) | Audit reason / validation status |
|---|---|---|---|---|---|
| 10.1038/s41467-023-37392-1 | 2023 | Predictive and robust gene selection for spatial transcriptomics | AGREE | We address this challenge by introducing ... PERSIST, to identify informative gene targets for spatial transcriptomics studies | This is a computational gene-selection paper for spatial transcriptomics, but the provided passage only introduces PERSIST and does not explicitly benchmark or use NS-Forest v2. [accepted_agree] |

| DOI | Year | Title | Audit decision | Evidence (verbatim quote) | Audit reason / validation status |
|---|---|---|---|---|---|
| 10.1038/s41597-022-01886-2 | 2023 | Brain Data Standards - A method for building data-driven cell-type ontologies | AGREE | a semi-automated pipeline to build a data-linked extension to the Cell Ontology | This is a data standard/ontology construction paper, and the evidence does not explicitly benchmark or use NS-Forest. [accepted_agree] |
| 10.1038/s41598-022-14192-z | 2022 | Cell type matching in single-cell RNA-sequencing data using FR-Match | AGREE | Here, we present significant enhancements to FR-Match (v2.0) | This is a computational cell-type matching paper centered on FR-Match, and the evidence does not explicitly benchmark or use NS-Forest. [accepted_agree] |
| 10.1038/s42003-024-06480-3 | 2024 | A point cloud segmentation framework for image-based spatial transcriptomics | DISAGREE | We simulated 34 cell-type marker genes selected with the NS-forest algorithm34. | This computational spatial-methods paper uses NS-Forest-selected genes only to construct a simulation setup; the packet does not link NS-Forest to the paper's own final targeted spatial panel or probe selection. [accepted_disagree] |
| 10.1093/bib/bbaa339 | 202 | FR-Match: robust matching of cell type clusters from single cell RNA sequencing data using the Friedman–Rafsky non-parametric test | AGREE | Here, we introduce a cluster-to-cluster cell type matching method —FR-Match ... | This is a computational single-cell cluster-matching paper; the packet does not explicitly benchmark or use NS-Forest v2, so the conservative negative is COMP. [accepted_agree] |
| 10.1093/bib/bbad389 | 2023 | FEED: a feature selection method based on gene expression decomposition for single cell clustering | AGREE | we present a Feature sElection method based on gene Expression Decomposition (FEED) ... | This is a computational feature-selection paper for single-cell clustering; the packet does not explicitly benchmark or use NS-Forest v2, so the conservative negative is COMP. [accepted_agree] |
| 10.1093/bib/bbaf478 | 2025 | Multi-view gene panel characterization for spatially resolved omics | DISAGREE | "We compared panelScope-OA against several methods tested here ..." plus the abstract describing panelScope/panelScope-OA. | The paper is a computational panel-characterization and optimization paper, but the provided evidence does not specifically support benchmarking of NS-Forest v2 itself, so the conservative negative label is COMP. [accepted_disagree] |
| 10.1093/nargab/lqad070 | 2023 | Single-cell reference mapping to construct and extend cell-type hierarchies | DISAGREE | "We do this using ... NS-Forest ... We ran NS-Forest ... on each dataset separately ..." | This packet supports direct use of NS-Forest within a computational method paper, but not explicit benchmarking or comparison of |

| DOI | Year | Title | Audit decision | Evidence (verbatim quote) | Audit reason / validation status |
|---|---|---|---|---|---|
| | | | | | NS-Forest itself. [accepted_disagree] |
| 10.1101/2024.04.08.588510 | 2024 | scRCA: a Siamese network-based pipeline for the annotation of cell types using imperfect single-cell RNA-seq reference data | AGREE | "Here, we built a Siamese network-based pipeline, termed scRCA ..." | This is a computational cell-type annotation paper introducing scRCA, and the packet provides no explicit NS-Forest comparison or use evidence. [accepted_agree] |
| 10.1101/2025.04.10.648034 | 2025 | Benchmarking single cell transcriptome matching methods for incremental growth of cell atlases | AGREE | This study benchmarks four computational tools for cell type annotation and matching – Azimuth, CellTypist, scArches, and FR-Match | This is a computational benchmarking paper, but the explicit benchmarked tools in the packet are Azimuth, CellTypist, scArches, and FR-Match, not NS-Forest. [accepted_agree] |
| 10.1101/2025.10.08.681071 | 2025 | Optimal Gene Panel Selection for Targeted Spatial Transcriptomics Experiments | AGREE | we introduce a novel method, named ReconST, to automatically design optimal gene panels for spatial transcriptomics profiling | This is a computational gene-panel-selection paper for targeted spatial transcriptomics; the packet does not explicitly benchmark or use NS-Forest v2, so the conservative negative is COMP. [accepted_agree] |
| 10.1101/2025.11.18.689140 | 2025 | CBKMR: A Copula-based Bayesian Kernel Machine Regression Framework for Optimal Marker Detection in Omics Data | AGREE | We propose a copula-based Bayesian kernel machine regression (CBKMR) model ... | This is a computational omics marker-detection paper; the packet does not explicitly benchmark or use NS-Forest v2, so the conservative negative is COMP. [accepted_agree] |
| 10.1109/tcbbio.2025.3609721 | 2025 | Knowledge-Guided Gene Panel Selection for Label-Free Single-Cell RNA-Seq Data: A Reinforcement Learning Perspective | DISAGREE | Our comparative analysis evaluated RIGPS against seven widely used baselines ... (7) gpsFISH [48] | The paper is computational and benchmark-oriented, but the provided evidence describes gpsFISH rather than NS-Forest. The conservative final label is COMP. [accepted_disagree] |
| 10.1109/tvcg.2025.3636102 | 2025 | CellScout: Visual Analytics for Mining Biomarkers in Cell State Discovery | AGREE | "we first designed a machine-learning algorithm ... to identify meaningful associations between cell populations and biomarkers." | This is a visual analytics / biomarker-mining methods paper, and the evidence does not explicitly show NS-Forest benchmarking or use. [accepted_agree] |
| 10.1186/s13059-021-02548-z | 2021 | geneBasis: an iterative approach for unsupervised selection of targeted gene panels from scRNA-seq | AGREE | We introduce an iterative approach, geneBasis, for selecting an optimal gene panel ... | This is a computational targeted-gene-panel selection paper; the packet does not explicitly benchmark or use NS-Forest v2, so the conservative negative is |

| DOI | Year | Title | Audit decision | Evidence (verbatim quote) | Audit reason / validation status |
|---|---|---|---|---|---|
| | | | | | COMP. [accepted_agree] |
| 10.1186/s13059-024-03174-1 | 2024 | Gene panel selection for targeted spatial transcriptomics | AGREE | Here we describe gpsFISH, a computational method performing gene selection through optimizing detection of known cell types. | This is a computational gene-panel-selection paper, but the provided passage only describes gpsFISH itself and does not explicitly benchmark or use NS-Forest v2. [accepted_agree] |
| 10.1186/s13059-024-03183-0 | 2024 | A comparison of marker gene selection methods for single-cell RNA sequencing data | DISAGREE | In this paper, we benchmark 59 computational methods for selecting marker genes in scRNA-seq data. | This is clearly a computational marker-selection comparison paper, but the evidence does not explicitly show NS-Forest among the benchmarked methods. The conservative negative label is COMP. [accepted_disagree] |
| 10.1371/journal.pone.0292961 | 2023 | On the use of QDE-SVM for gene feature selection and cell type classification from scRNA-seq data | AGREE | a quantum-inspired differential evolution ... was introduced to select a subset of genes ... | This is a computational gene-feature-selection paper for scRNA-seq; the packet does not explicitly benchmark or use NS-Forest v2, so the conservative negative is COMP. [accepted_agree] |
| 10.14569/ijacsa.2023.0140984 | 2023 | Analyzing RNA-Seq Gene Expression Data for Cancer Classification Through ML Approach | AGREE | the proposed ensemble classifier outperformed the existing machine-learning approaches | This is a machine-learning cancer classification paper, and the evidence does not explicitly benchmark or use NS-Forest. [accepted_agree] |
| 10.21203/rs.3.rs-4536158/v1 | 2024 | Spatial Deconvolution of Cell Types and Cell States at Scale Utilizing TACIT | DISAGREE | Spatial Deconvolution of Cell Types and Cell States at Scale Utilizing TACIT | TACIT is a computational method paper for spatial deconvolution/annotation. With no tool-specific evidence for NS-Forest, the conservative label is COMP, not EXP-CITED. [accepted_disagree] |

#### SCMER (17)

| DOI | Year | Title | Audit decision | Evidence (verbatim quote) | Audit reason / validation status |
|---|---|---|---|---|---|
| 10.1016/j.compbiomed.2025.109873 | 2025 | A novel coarsened graph learning method for scalable single-cell data analysis | AGREE | we propose a novel approach utilizing Feature-Aware Graph Coarsening via Hashing (FACH) | This is a computational graph-learning paper for single-cell data analysis, and the evidence does not explicitly benchmark or use |

| DOI | Year | Title | Audit decision | Evidence (verbatim quote) | Audit reason / validation status |
|---|---|---|---|---|---|
| | | | | | SCMER.<br>[accepted_agree] |
| 10.1038/s41467-025-67890-3 | 2025 | CellScope: high-performance cell atlas workflow with tree-structured representation | AGREE | "we present CellScope ... for constructing high-resolution cell atlases ..." | This is a computational cell-atlas workflow paper introducing CellScope, and the packet provides no explicit SCMER comparison or use evidence.<br>[accepted_agree] |
| 10.1038/s41467-026-68500-6 | 2026 | Robust characterization and interpretation of rare pathogenic cell populations from spatial omics using GARDEN | AGREE | "Here we show GARDEN, a computational framework ..." | This is a computational spatial-omics framework paper introducing GARDEN, and the packet provides no explicit SCMER comparison or use evidence.<br>[accepted_agree] |
| 10.1038/s41540-024-00339-3 | 2024 | MarkerMap: nonlinear marker selection for single-cell studies | AGREE | "Here we introduce MarkerMap ..." | This is a computational marker-selection paper introducing MarkerMap, and the packet provides no explicit SCMER comparison or use evidence.<br>[accepted_agree] |
| 10.1038/s41556-025-01755-1 | 2025 | CellNavi predicts genes directing cellular transitions by learning a gene graph-enhanced cell state manifold | AGREE | "we present CellNavi, a deep learning-based framework ..." | This is a computational framework paper introducing CellNavi, and the packet provides no explicit SCMER comparison or use evidence.<br>[accepted_agree] |
| 10.1038/s41598-026-39168-1 | 2026 | CiCLoDS: Joint cell clustering and gene selection for single-cell spatial transcriptomics | AGREE | We introduce CiCLoDS ... an unsupervised framework adapted for spatial transcriptomics to jointly optimize clustering and feature selection | This is a computational ST feature-selection/clustering paper, and the provided evidence does not explicitly benchmark or use SCMER.<br>[accepted_agree] |
| 10.1038/s42003-025-07872-9 | 2025 | Infusing structural assumptions into dimensionality reduction for single-cell RNA sequencing data to identify small gene sets | AGREE | "We propose the boosting autoencoder (BAE) approach ..." | This is a computational gene-selection/dimensionality-reduction paper introducing BAE, and the packet provides no explicit SCMER comparison or use evidence.<br>[accepted_agree] |
| 10.1093/bib/bbad389 | 2023 | FEED: a feature selection method based on gene expression decomposition for single cell clustering | DISAGREE | We compared the performance of FEED with seven state-of-the-art gene selection methods ... SCMarker and geneBasis | The paper is clearly computational, but the provided benchmark evidence names geneBasis rather than SCMER. The conservative final label is COMP, not COMP-BENCH.<br>[accepted_disagree] |

| DOI | Year | Title | Audit decision | Evidence (verbatim quote) | Audit reason / validation status |
|---|---|---|---|---|---|
| 10.1093/bib/bbaf478 | 2025 | Multi-view gene panel characterization for spatially resolved omics | DISAGREE | We curated eight gene panels ... including expert curation, random selection, computational methods, and the use of large language models. | The paper is clearly computational, but the provided evidence does not mention SCMER specifically. Conservatively, the final label should be COMP rather than COMP-BENCH. [accepted_disagree] |
| 10.1101/2023.07.17.549344 | 2023 | Recurrent Composite Markers of Cell Types and States | AGREE | we developed Recurrent Composite Markers for Biological Identities with Neighborhood Enrichment (RECOMBINE) | This is a computational marker-discovery paper, and the provided evidence does not explicitly benchmark or use SCMER. [accepted_agree] |
| 10.1101/2024.10.27.620174 | 2024 | Directing cellular transitions on gene graph-enhanced cell state manifold | AGREE | we present CellNavi, a deep learning-based framework designed to predict genes that drive cell state transitions | This is a computational framework for predicting genes that drive cell-state transitions, and the evidence does not explicitly benchmark or use SCMER. [accepted_agree] |
| 10.1101/2025.10.14.682419 | 2025 | scConcept: Contrastive pretraining for technology-agnostic single-cell representations beyond reconstruction | AGREE | We present scConcept ... a transformer-based contrastive learning framework | This is a computational single-cell foundation-model paper; the packet does not explicitly benchmark or use SCMER, so the conservative negative is COMP. [accepted_agree] |
| 10.1109/bibm58861.2023.10385799 | 2023 | Co-clustering of single-cell RNA-seq data based on weighted non-negative matrix tri-factorization combined with consensus clustering | AGREE | we propose scCO2, a method based on weighted non-negative matrix tri-factorization | This is a computational co-clustering paper, and the evidence does not explicitly benchmark or use SCMER. [accepted_agree] |
| 10.1109/tcbb.2022.3203185 | 2022 | A Clustering Method Unifying Cell-Type Recognition and Subtype Identification for Tumor Heterogeneity Analysis | AGREE | we proposed a cell similarity metric that unifies cell type recognition and subtype identification (UCRSI) | This is a computational single-cell clustering paper for tumor heterogeneity analysis; the packet does not explicitly benchmark or use SCMER, so the conservative negative is COMP. [accepted_agree] |
| 10.1186/s13059-024-03174-1 | 2024 | Gene panel selection for targeted spatial transcriptomics | AGREE | "Here we describe gpsFISH, a computational method ..." | This is a computational panel-selection paper about gpsFISH, and the packet provides no explicit SCMER comparison or use evidence. [accepted_agree] |
| 10.1186/s13059-025-03810-4 | 2025 | A novel gene expression stability metric to unveil homeostasis and regulation | AGREE | "Here, we introduce a new measure of gene expression | This is a computational methods paper introducing the gene homeostasis Z-index, and the |

| DOI | Year | Title | Audit decision | Evidence (verbatim quote) | Audit reason / validation status |
|---|---|---|---|---|---|
| | | | | stability, the gene homeostasis Z-index ..." | packet provides no explicit SCMER comparison or use evidence. [accepted_agree] |
| 10.64898/2026.02.25.707866 | 2026 | Information Geometry Reconciles Discrete and Continuous Variation in Single-Cell and Spatial Transcriptomic Analysis | AGREE | "we introduce GAIA ... an information-geometric framework ..." | This is a computational methods paper introducing GAIA, and the packet gives no explicit benchmarking or use evidence for SCMER, so the conservative negative label is COMP. [accepted_agree] |

#### RankCorr (16)

| DOI | Year | Title | Audit decision | Evidence (verbatim quote) | Audit reason / validation status |
|---|---|---|---|---|---|
| 10.1016/j.crmeth.2026.101329 | 2026 | Scalable nonparametric clustering with unified marker gene selection for single-cell RNA-seq data | AGREE | "we present NCLUSION ... to identify marker genes and cluster single-cell expression data simultaneously." | This is a computational clustering and marker-selection paper introducing NCLUSION, and the packet provides no explicit RankCorr comparison or use evidence. [accepted_agree] |
| 10.1038/s44320-025-00176-4 | 2025 | CANTAO: guiding clustering and annotation in single-cell RNA sequencing using average overlap | AGREE | "Here, we introduce CANTAO ..." | This is a computational clustering/annotation paper, and the packet provides no explicit RankCorr comparison or use evidence. [accepted_agree] |
| 10.1093/bioinformatics/btac683 | 2022 | AIscEA: unsupervised integration of single-cell gene expression and chromatin accessibility via their biological consistency | AGREE | We introduce AIscEA ... a computational method that integrates single-cell gene expression and chromatin accessibility | This is a computational integration paper, and the evidence does not explicitly benchmark or use RankCorr. [accepted_agree] |
| 10.1093/bioinformatics/btad216 | 2023 | CellBRF: a feature selection method for single-cell clustering using cell balance and random forest | AGREE | We develop CellBRF, a feature selection method that considers genes' relevance to cell types for single-cell clustering. | This is a computational feature-selection paper for single-cell clustering; the packet does not explicitly benchmark or use RankCorr, so the conservative negative is COMP. [accepted_agree] |
| 10.1093/bioinformatics/btag051 | 2026 | Inference of marker genes of subtle cell state changes via iLR: iterative logistic regression | AGREE | We present iterative logistic regression (iLR) for the identification of small sets of informative marker genes. | This is a computational method paper proposing iLR, and the evidence does not explicitly benchmark or use RankCorr. [accepted_agree] |

| DOI | Year | Title | Audit decision | Evidence (verbatim quote) | Audit reason / validation status |
| --- | --- | --- | --- | --- | --- |
| 10.1093/gigascience/giad113 | 2024 | Computational reproducibility of Jupyter notebooks from biomedical publications | AGREE | We address computational reproducibility at 2 levels | This is a computational reproducibility study, and the evidence does not explicitly benchmark or use RankCorr. [accepted_agree] |
| 10.1101/2023.07.17.549344 | 2023 | Recurrent Composite Markers of Cell Types and States | AGREE | we developed Recurrent Composite Markers ... (RECOMBINE) | This is a computational marker-combination methods paper; the packet does not explicitly benchmark or use RankCorr, so the conservative negative is COMP. [accepted_agree] |
| 10.1101/2024.03.23.586413 | 2024 | Algorithms for a Commons Cell Atlas | AGREE | we present a collection of data, algorithms, and tools to automate cataloging and analyzing cell types | This is a computational cell-atlas algorithms paper, and the evidence does not explicitly benchmark or use RankCorr. [accepted_agree] |
| 10.1101/2024.12.23.630040 | 2024 | CORTADO: Hill Climbing Optimization for Cell-Type Specific Marker Gene Discovery | DISAGREE | "CORTADO was compared to four baseline methods across seven datasets ..." | The paper is computational and compares CORTADO to baseline methods, but the provided evidence does not specifically identify RankCorr among them, so the conservative negative label is COMP. [accepted_disagree] |
| 10.1101/gr.280539.125 | 2025 | Label-free selection of marker genes in single-cell and spatial transcriptomics with geneCover | AGREE | we introduce geneCover, a label-free combinatorial method that selects an optimal panel | This is a computational methods paper introducing geneCover, and the provided evidence does not explicitly benchmark or use RankCorr. [accepted_agree] |
| 10.1101/gr.280637.125 | 2025 | Optimal marker genes for <i>c</i> -separated cell types with SepSolve | AGREE | we propose to find a small set of genes ... we formulate a linear program | This is a computational marker-gene selection paper; the packet does not explicitly benchmark or use RankCorr, so the conservative negative is COMP. [accepted_agree] |
| 10.1186/s13059-021-02548-z | 2021 | geneBasis: an iterative approach for unsupervised selection of targeted gene panels from scRNA-seq | AGREE | We introduce an iterative approach, geneBasis, for selecting an optimal gene panel | This is a computational methods paper on unsupervised targeted gene-panel selection; the packet does not explicitly benchmark or use RankCorr, so the conservative negative is COMP. [accepted_agree] |

| DOI | Year | Title | Audit decision | Evidence (verbatim quote) | Audit reason / validation status |
|---|---|---|---|---|---|
| 10.1186/s13059-023-02904-1 | 2023 | The shaky foundations of simulating single-cell RNA sequencing data | AGREE | "Here, we evaluated 16 scRNA-seq simulation methods ..." | This is a computational evaluation paper on scRNA-seq simulators, and the evidence does not explicitly mention RankCorr. [accepted_agree] |
| 10.1371/journal.pcbi.1012643 | 2024 | Hierarchical marker genes selection in scRNA-seq analysis | AGREE | Hierarchical Marker Genes Selection in scRNA-seq Analysis | This is a computational marker-selection methods paper; the packet does not show explicit benchmarking of RankCorr, so the conservative negative is COMP. [accepted_agree] |
| 10.3390/bioengineering12080829 | 2025 | DropDAE: Denosing Autoencoder with Contrastive Learning for Addressing Dropout Events in scRNA-seq Data | AGREE | we introduce DropDAE, a denoising autoencoder (DAE) model enhanced with contrastive learning ... | This is a computational scRNA-seq denoising/imputation paper; the packet does not explicitly benchmark or use RankCorr, so the conservative negative is COMP. [accepted_agree] |
| 10.3390/ijms25041998 | 2024 | uPAR (PLAUR) Marks Two Intra-Tumoral Subtypes of Glioblastoma: Insights from Single-Cell RNA Sequencing | AGREE | we identified PLAUR as a marker gene for two distinct subtypes in GBM | This is a computational marker-identification paper in glioblastoma, and the evidence does not explicitly benchmark or use RankCorr. [accepted_agree] |

#### PERSIST (14)

| DOI | Year | Title | Audit decision | Evidence (verbatim quote) | Audit reason / validation status |
|---|---|---|---|---|---|
| 10.1016/j.asoc.2025.114110 | 2025 | Metaheuristic-driven feature selection with dual feedback mechanisms for imbalanced cancer classification | AGREE | this study proposes DVFF ... a novel bacterial heuristic-based feature selection method | This is a computational feature-selection paper, and the packet does not provide explicit benchmarking or direct use evidence for PERSIST. [accepted_agree] |
| 10.1093/bib/bbaf478 | 2025 | Multi-view gene panel characterization for spatially resolved omics | AGREE | we developed a two-part framework: (i) panelScope ... and (ii) panelScope-OA | This is a computational panel-characterization/optimization paper, and the evidence does not explicitly benchmark or use PERSIST. [accepted_agree] |
| 10.1093/bioinformatics/btaf619 | 2025 | STAHD: a scalable and accurate method to detect spatial domains in high-resolution spatial transcriptomics data | AGREE | We present STAHD, a scalable and efficient framework for spatial domain detection in ST data. | This is a computational spatial-domain detection paper, and the evidence does not explicitly |

| DOI | Year | Title | Audit decision | Evidence (verbatim quote) | Audit reason / validation status |
| --- | --- | --- | --- | --- | --- |
| | | | | | benchmark or use PERSIST. [accepted_agree] |
| 10.1093/gigascience/giae089 | 2024 | stMMR: accurate and robust spatial domain identification from spatially resolved transcriptomics with multimodal feature representation | AGREE | "the authors developed a multi-modal deep learning method for identifying spatial domains from ST data ..." | This is a computational spatial-domain-identification paper, and the packet provides no explicit PERSIST comparison or use evidence. [accepted_agree] |
| 10.1093/nar/gkad157 | 2023 | Multi-task learning from multimodal single-cell omics with Matilda | AGREE | Here, we present Matilda, a multi-task learning method for integrative analysis of multimodal single-cell omics data. | This is a computational multimodal single-cell omics methods paper; the packet does not explicitly benchmark or use PERSIST, so the conservative negative is COMP. [accepted_agree] |
| 10.1101/2023.04.06.535943 | 2023 | CellCover Defines Marker Gene Panels Capturing Developmental Progression in Neocortical Neural Stem Cell Identity | AGREE | "we propose viewing marker gene panel selection as a variation of the minimal set-covering problem ..." | This is a computational marker-panel selection paper, and the evidence does not explicitly show PERSIST benchmarking or use. [accepted_agree] |
| 10.1101/2024.05.27.594872 | 2024 | Towards a universal spatial molecular atlas of the mouse brain | AGREE | here, we introduce FuseMap, a deep-learning-based framework for spatial transcriptomics ... | This is a computational spatial-transcriptomics integration and atlas-construction paper; the packet does not explicitly benchmark or use PERSIST, so the conservative negative is COMP. [accepted_agree] |
| 10.1101/2024.07.05.602236 | 2024 | SOFisher: Reinforcement Learning-Guided Experiment Designs for Spatial Omics | AGREE | we developed SOFisher ... to guide the selection of the next FOV position | This is a computational spatial-omics experiment-design paper about FOV selection; the packet does not explicitly benchmark or use PERSIST, so the conservative negative is COMP. [accepted_agree] |
| 10.1101/2024.08.04.606512 | 2024 | Identifying Differential Spatial Expression Patterns across Different Slices, Conditions and Developmental Stages with Interpretable Deep Learning | AGREE | we propose DSEP gene identification as a new task and develop River ... | This is a computational spatial-transcriptomics methods paper; the packet does not explicitly benchmark or use PERSIST, so the conservative negative is COMP. [accepted_agree] |

| DOI | Year | Title | Audit decision | Evidence (verbatim quote) | Audit reason / validation status |
|---|---|---|---|---|---|
| 10.1101/2025.10.14.682419 | 2025 | scConcept: Contrastive pretraining for technology-agnostic single-cell representations beyond reconstruction | AGREE | We present scConcept ... a transformer-based contrastive learning framework | This is a computational single-cell foundation-model paper; the packet does not explicitly benchmark or use PERSIST, so the conservative negative is COMP. [accepted_agree] |
| 10.1101/2025.11.18.689140 | 2025 | CBKMR: A Copula-based Bayesian Kernel Machine Regression Framework for Optimal Marker Detection in Omics Data | AGREE | We propose a copula-based Bayesian kernel machine regression (CBKMR) model ... | This is a computational omics marker-detection paper; the packet does not explicitly benchmark or use PERSIST, so the conservative negative is COMP. [accepted_agree] |
| 10.1186/s12864-024-11072-w | 2024 | Deciphering spatial domains from spatially resolved transcriptomics through spatially regularized deep graph networks | AGREE | "we introduce ... SR-DGN ... to learn spatially-consistent and informative spot representations." | This is a computational spatial-domain-identification paper, and the packet provides no explicit PERSIST comparison or use evidence. [accepted_agree] |
| 10.48550/arxiv.2301.00557 | 2023 | Learning to Maximize Mutual Information for Dynamic Feature Selection | AGREE | we develop a learning approach based on amortized optimization | This is a machine-learning feature-selection paper, and the evidence does not explicitly benchmark or use PERSIST. [accepted_agree] |
| 10.64898/2026.02.25.707866 | 2026 | Information Geometry Reconciles Discrete and Continuous Variation in Single-Cell and Spatial Transcriptomic Analysis | AGREE | we introduce GAIA ... an information-geometric framework for cell representation learning | This is a computational framework paper, and the evidence does not explicitly benchmark or use PERSIST. [accepted_agree] |

#### Spapros (11)

| DOI | Year | Title | Audit decision | Evidence (verbatim quote) | Audit reason / validation status |
|---|---|---|---|---|---|
| 10.1038/s41598-026-39168-1 | 2026 | CiCLoDS: Joint cell clustering and gene selection for single-cell spatial transcriptomics | AGREE | "We introduce CiCLoDS ... to jointly optimize clustering and feature selection ..." | This is a computational method paper introducing CiCLoDS; the evidence does not explicitly mention Spapros or benchmark it. [accepted_agree] |
| 10.1101/2023.07.05.547784 | 2023 | scARE: Attribution Regularization for Single Cell Representation Learning | AGREE | we propose scARE, a novel end-to-end generative deep learning model ... | This is a computational single-cell representation-learning paper; the packet does not explicitly benchmark or use Spapros, so the conservative negative is COMP. [accepted_agree] |

| DOI | Year | Title | Audit decision | Evidence (verbatim quote) | Audit reason / validation status |
|---|---|---|---|---|---|
| 10.1101/2024.07.04.601829 | 2024 | SPArrOW: a flexible, interactive and scalable pipeline for spatial transcriptomics analysis | AGREE | "here we present SPArrOW, a flexible, modular and scalable pipeline for processing spatial transcriptomics data" | This is a computational spatial transcriptomics processing pipeline paper, and the packet provides no explicit Spapros comparison or use evidence.<br>[accepted_agree] |
| 10.1101/2024.07.05.602236 | 2024 | SOFisher: Reinforcement Learning-Guided Experiment Designs for Spatial Omics | AGREE | we developed SOFisher, a reinforcement learning-based framework ... to guide the selection of the next FOV position | This is a computational reinforcement-learning experiment-design paper, and the evidence does not explicitly benchmark or use Spapros.<br>[accepted_agree] |
| 10.1101/2025.08.29.673037 | 2025 | ProbeST: a custom probe design pipeline for Spatial Transcriptomics | AGREE | we present ProbeST, an open-source computational pipeline for designing custom probe sets for any organism of interest | This is a computational probe-design pipeline for spatial transcriptomics; the packet does not explicitly benchmark or use Spapros, so the conservative negative is COMP.<br>[accepted_agree] |
| 10.1101/2025.10.14.682419 | 2025 | scConcept: Contrastive pretraining for technology-agnostic single-cell representations beyond reconstruction | AGREE | We present scConcept ... a transformer-based contrastive learning framework | This is a computational foundation-model paper, and the evidence does not explicitly benchmark or use Spapros.<br>[accepted_agree] |
| 10.1101/gr.280637.125 | 2025 | Optimal marker genes for <i>c</i> -separated cell types with SepSolve | AGREE | "we propose to find a small set of genes such that cell types are c-separated ..." | This is a computational marker-selection paper introducing SepSolve, and the packet provides no explicit Spapros comparison or use evidence.<br>[accepted_agree] |
| 10.1186/s13059-024-03303-w | 2024 | Gene count normalization in single-cell imaging-based spatially resolved transcriptomics | AGREE | "Here, we investigate the potential impact of different gene count normalization methods ..." | This is a computational analysis paper on normalization in imaging-based spatial transcriptomics, and the packet provides no explicit Spapros comparison or use evidence.<br>[accepted_agree] |
| 10.64898/2026.01.26.701684 | 2026 | ESFS: A Noise-Resilient Framework for Feature Selection and Marker Gene Discovery in Single-Cell Transcriptomics | AGREE | we introduce a modular software suite called Entropy Sorting Feature Selection (ESFS) | This is a computational feature-selection software paper, and the evidence does not explicitly benchmark or use Spapros.<br>[accepted_agree] |
| 10.64898/2026.02.26.707870 | 2026 | PantheonOS: An Evolvable Multi-Agent Framework for Automatic Genomics Discovery | DISAGREE | combining e.g., HVG, DE, RF, SpaPROS or scGeneFit with fixed weighting schemes | The PantheonOS passage cites SpaPROS as one of the competing/static strategies, but it |

| DOI | Year | Title | Audit decision | Evidence (verbatim quote) | Audit reason / validation status |
|---|---|---|---|---|---|
| | | | | | does not explicitly describe a benchmark or evaluation of Spapros itself. [accepted_disagree] |
| 10.7554/elife.107070.2 | 2026 | Evidence of off-target probe binding affecting 10x Genomics Xenium gene panels compromise accuracy of spatial transcriptomic profiling | AGREE | We developed a software tool, Off-target Probe Tracker (OPT) | This is a computational off-target probe analysis paper, and the evidence does not explicitly benchmark or use Spapros. [accepted_agree] |

#### SMaSH (10)

| DOI | Year | Title | Audit decision | Evidence (verbatim quote) | Audit reason / validation status |
|---|---|---|---|---|---|
| 10.1016/j.molp.2025.12.026 | 2025 | PlantscRNAdb 4.0: Improved marker identification and annotation under a cell-type uniformity for plants | AGREE | "we present PlantscRNAdb release 4.0 ... Using HCMarker, a novel computational tool developed in this study ..." | This is a computational database/resource paper introducing PlantscRNAdb 4.0 and HCMarker, and the evidence does not explicitly show SMaSH comparison or use. [accepted_agree] |
| 10.1038/s41540-024-00339-3 | 2024 | MarkerMap: nonlinear marker selection for single-cell studies | DISAGREE | "We benchmark MarkerMap against existing marker selection approaches ..." | This is a computational marker-selection paper, but the provided evidence only supports general benchmarking against existing approaches rather than a tool-specific benchmark of SMaSH, so the conservative negative label is COMP. [accepted_disagree] |
| 10.1093/bioinformatics/btag051 | 2026 | Inference of marker genes of subtle cell state changes via iLR: iterative logistic regression | AGREE | "We present iterative logistic regression (iLR) for the identification of small sets of informative marker genes." | This is a computational marker-gene paper introducing iLR, and the evidence does not explicitly show SMaSH comparison or use. [accepted_agree] |
| 10.1101/2024.04.24.590691 | 2024 | mastR: Marker Automated Screening Tool for multi-omics data | AGREE | "We developed mastR, a novel computational tool for accurate marker identification ..." | This is a computational marker-identification paper introducing mastR, and the evidence does not explicitly show SMaSH benchmarking or use. [accepted_agree] |

| DOI | Year | Title | Audit decision | Evidence (verbatim quote) | Audit reason / validation status |
|---|---|---|---|---|---|
| 10.1101/2025.11.18.689140 | 2025 | CBKMR: A Copula-based Bayesian Kernel Machine Regression Framework for Optimal Marker Detection in Omics Data | AGREE | We propose a copula-based Bayesian kernel machine regression (CBKMR) model | This is a computational marker-detection paper; the evidence does not explicitly benchmark or use SMaSH. [accepted_agree] |
| 10.1101/gr.280637.125 | 2025 | Optimal marker genes for <i>c</i> -separated cell types with SepSolve | AGREE | "we propose to find a small set of genes such that cell types are c-separated ..." | This is a computational marker-selection paper introducing SepSolve, and the packet provides no explicit SMaSH comparison or use evidence. [accepted_agree] |
| 10.1186/s13059-021-02548-z | 2021 | geneBasis: an iterative approach for unsupervised selection of targeted gene panels from scRNA-seq | AGREE | We introduce an iterative approach, geneBasis, for selecting an optimal gene panel | The packet is for the geneBasis methods paper and does not explicitly benchmark or use SMaSH in the provided evidence, so the conservative label is COMP. [accepted_agree] |
| 10.1371/journal.pcbi.1013354 | 2025 | Improving cell-type composition inference in spatial transcriptomics with SpaDAMA | AGREE | "we introduce a Domain-Adversarial Masked Autoencoder (SpaDAMA) method." | This is a computational spatial transcriptomics deconvolution paper, and the packet provides no explicit SMaSH comparison or use evidence. [accepted_agree] |
| 10.1371/journal.pone.0292961 | 2023 | On the use of QDE-SVM for gene feature selection and cell type classification from scRNA-seq data | AGREE | "QDE ... was introduced to select a subset of genes ..." | This is a computational scRNA-seq feature-selection paper, but the evidence only describes QDE-SVM and does not show explicit SMaSH use or benchmarking. [accepted_agree] |
| 10.3390/genes14030596 | 2023 | Cell Type Annotation Model Selection: General-Purpose vs. Pattern-Aware Feature Gene Selection in Single-Cell RNA-Seq Data | AGREE | "we compare and evaluate ... XGBoost and SVM, with information gain as a feature selection method." | This is a computational cell-type annotation / feature-selection paper, and the evidence does not explicitly mention SMaSH. [accepted_agree] |

#### scPNMF (9)

| DOI | Year | Title | Audit decision | Evidence (verbatim quote) | Audit reason / validation status |
|---|---|---|---|---|---|
| 10.1038/s41467-023-37392-1 | 2023 | Predictive and robust gene selection for spatial transcriptomics | AGREE | We address this challenge by introducing ... PERSIST, to identify informative gene targets for spatial transcriptomics studies | This is a computational gene-selection paper for spatial transcriptomics; the packet does not explicitly benchmark or use scPNMF, so the conservative |

| DOI | Year | Title | Audit decision | Evidence (verbatim quote) | Audit reason / validation status |
|---|---|---|---|---|---|
| | | | | | negative is COMP. [accepted_agree] |
| 10.1093/bib/bbad389 | 2023 | FEED: a feature selection method based on gene expression decomposition for single cell clustering | AGREE | "we present ... FEED ... which selects informative genes ..." | This is a computational feature-selection paper introducing FEED, and the packet provides no explicit scPNMF benchmarking or use evidence. [accepted_agree] |
| 10.1093/bioadv/vbag047 | 2026 | Sparse dimensionality reduction for analyzing single-cell-resolved interactions | AGREE | we present an end-to-end dimensionality reduction workflow, specifically tailored for single-cell cell–cell interaction data | This is a computational dimensionality-reduction paper for single-cell interaction data; the packet does not explicitly benchmark or use scPNMF, so the conservative negative is COMP. [accepted_agree] |
| 10.1101/2021.06.15.448478 | 2021 | Active feature selection discovers minimal gene sets for classifying cell types and disease states with single-cell mRNA-seq data | AGREE | we introduce an active learning method (ActiveSVM) that identifies minimal but highly-informative gene sets | This is a computational active feature-selection paper, and the evidence does not explicitly benchmark or use scPNMF. [accepted_agree] |
| 10.1101/2025.10.08.681071 | 2025 | Optimal Gene Panel Selection for Targeted Spatial Transcriptomics Experiments | AGREE | we introduce a novel method, named ReconST, to automatically design optimal gene panels | This is a computational gene-panel-selection paper for targeted spatial transcriptomics; the packet does not explicitly benchmark or use scPNMF, so the conservative negative is COMP. [accepted_agree] |
| 10.1109/cdc57313.2025.11312726 | 2022 | Orthogonal Nonnegative Matrix Factorization with Sparsity Constraints | DISAGREE | These include methods in ... [18] (PNMF). | The packet supports benchmarking against PNMF, but does not explicitly identify the named scPNMF tool. Conservatively, this should be COMP rather than COMP-BENCH. [accepted_disagree] |
| 10.1109/tcbb.2023.3298334 | 2023 | Multi-View Clustering With Graph Learning for scRNA-Seq Data | AGREE | In this study, a multi-view clustering with graph learning algorithm (MCGL) for scRNA-seq data is proposed ... | This is a computational single-cell clustering paper; the packet does not explicitly benchmark or use scPNMF, so the conservative negative is COMP. [accepted_agree] |

| DOI | Year | Title | Audit decision | Evidence (verbatim quote) | Audit reason / validation status |
|---|---|---|---|---|---|
| 10.1186/s13059-021-02548-z | 2021 | geneBasis: an iterative approach for unsupervised selection of targeted gene panels from scRNA-seq | AGREE | We introduce an iterative approach, geneBasis, for selecting an optimal gene panel | The packet is for the geneBasis methods paper and does not explicitly benchmark or use scPNMF in the provided evidence, so the conservative label is COMP. [accepted_agree] |
| 10.1615/jmachlearnmodelcomput.2023047230 | 2023 | EVALUATING THE COMPLEXITY OF GENE COEXPRESSION ESTIMATION FOR SINGLE-CELL DATA | AGREE | This study thoroughly tests the existing gene co-expression estimation methods on simulation datasets with known ground truth co-expression networks. | This is a computational single-cell benchmarking paper on gene coexpression estimation; the packet does not explicitly benchmark or use scPNMF, so the conservative negative is COMP. [accepted_agree] |

#### geneBasis (8)

| DOI | Year | Title | Audit decision | Evidence (verbatim quote) | Audit reason / validation status |
|---|---|---|---|---|---|
| 10.1038/s41592-024-02496-z | 2024 | Probe set selection for targeted spatial transcriptomics | DISAGREE | We assessed Spapros performance against ten recently proposed gene selection methods ... in a large-scale benchmark | The paper is clearly a computational benchmark study, but the provided evidence does not explicitly mention geneBasis. The conservative negative label is COMP. [accepted_disagree] |
| 10.1038/s41592-025-02795-z | 2025 | Spatial gene expression at single-cell resolution from histology using deep learning with GHIST | AGREE | "here, we introduce GHIST, a deep learning-based framework ..." | This is a computational framework paper introducing GHIST, and the packet provides no explicit geneBasis comparison or use evidence. [accepted_agree] |
| 10.1093/bioinformatics/btae518 | 2024 | scDiffusion: conditional generation of high-quality single-cell data using diffusion model | AGREE | "we developed scDiffusion, a generative model ... to generate high-quality scRNA-seq data ..." | This is a generative-model paper for scRNA-seq data, and the packet provides no explicit geneBasis comparison or use evidence. [accepted_agree] |
| 10.1101/2025.10.08.681071 | 2025 | Optimal Gene Panel Selection for Targeted Spatial Transcriptomics Experiments | AGREE | "we introduce a novel method, named ReconST, to automatically design optimal gene panels ..." | This is a computational panel-design paper introducing ReconST, and the packet provides no explicit geneBasis comparison or use evidence. [accepted_agree] |
| 10.1101/2025.10.14.682419 | 2025 | scConcept: Contrastive pretraining for technology-agnostic single-cell | AGREE | We present scConcept ... a transformer-based contrastive | This is a computational foundation-model paper, and the evidence |

| DOI | Year | Title | Audit decision | Evidence (verbatim quote) | Audit reason / validation status |
|---|---|---|---|---|---|
| | | representations beyond reconstruction | | learning framework | does not explicitly benchmark or use geneBasis. [accepted_agree] |
| 10.1109/tcbbio.2025.3624040 | 2025 | scMID: A Deep Multi-Omics Integration Framework for Comprehensive Single-Cell Data Analysis | AGREE | In this study, we propose an analysis method based on single-cell multi-omics data integration and dropout pattern (scMID). | This is a computational multi-omics integration paper; the packet does not explicitly benchmark or use geneBasis, so the conservative negative is COMP. [accepted_agree] |
| 10.1186/s13059-024-03174-1 | 2024 | Gene panel selection for targeted spatial transcriptomics | AGREE | "Here we describe gpsFISH, a computational method ..." | This is a computational gene-panel-selection paper about gpsFISH, and the packet provides no explicit geneBasis benchmarking or use evidence. [accepted_agree] |
| 10.64898/2026.02.18.706600 | 2026 | Comparative Biology at Single-Cell Resolution: Rigorous Matching of Atlases for Cross-Species Analysis | AGREE | "we introduce RIMA ... a method for quantitatively comparing transcriptomic atlases across species ..." | This is a computational cross-species atlas-matching paper, and the packet provides no explicit geneBasis comparison or use evidence. [accepted_agree] |

#### gpsFISH (4)

| DOI | Year | Title | Audit decision | Evidence (verbatim quote) | Audit reason / validation status |
|---|---|---|---|---|---|
| 10.1002/advs.202513210 | 2025 | CELLama: Foundation Model for Single Cell and Spatial Transcriptomics by Cell Embedding Leveraging Language Model Abilities | AGREE | "CELLama ... develops a framework ... enabling universal cell embedding." | This is a foundation-model paper for single-cell and spatial transcriptomics, and the packet provides no explicit gpsFISH comparison or use evidence. [accepted_agree] |
| 10.1038/s41592-025-02795-z | 2025 | Spatial gene expression at single-cell resolution from histology using deep learning with GHIST | AGREE | we introduce GHIST, a deep learning-based framework that predicts spatial gene expression at single-cell resolution | This is a computational spatial-gene-expression prediction paper, and the evidence does not explicitly benchmark or use gpsFISH. [accepted_agree] |
| 10.1101/2024.07.05.602236 | 2024 | SOFisher: Reinforcement Learning-Guided Experiment Designs for Spatial Omics | AGREE | "we developed SOFisher ... to guide the selection of the next FOV position ..." | The paper introduces SOFisher for field-of-view selection in spatial omics, not targeted gene-panel selection, and the packet does not show explicit gpsFISH benchmarking or use. [accepted_agree] |

| DOI | Year | Title | Audit decision | Evidence (verbatim quote) | Audit reason / validation status |
|---|---|---|---|---|---|
| 10.4208/csiam-ls.so-2025-0032 | 2026 | BRIDGE: An Interpretable Framework for Transcriptome-Scale Completion of Panel-Limited Spatial Transcriptomics | AGREE | We present BRIDGE, an interpretable framework that expands panel-limited spatial transcriptomics to transcriptome-level resolution ... | This is a computational spatial-transcriptomics imputation/completion paper; the packet does not explicitly benchmark or use gpsFISH, so the conservative negative is COMP. [accepted_agree] |

#### scMAGS (4)

| DOI | Year | Title | Audit decision | Evidence (verbatim quote) | Audit reason / validation status |
|---|---|---|---|---|---|
| 10.1093/bib/bbae626 | 2024 | CosGeneGate selects multi-functional and credible biomarkers for single-cell analysis | AGREE | "Here, we present a novel model, named CosGeneGate, to select marker genes ..." | This is a computational marker-selection paper introducing CosGeneGate, and the packet does not support a direct benchmark or use claim for scMAGS beyond authoritative citation inclusion. [accepted_agree] |
| 10.1101/2024.04.24.590691 | 2024 | mastR: Marker Automated Screening Tool for multi-omics data | AGREE | We developed mastR, a novel computational tool for accurate marker identification from omics data. | This is a computational marker-identification paper, and the packet does not support any direct benchmark or use claim for scMAGS. [accepted_agree] |
| 10.1101/2024.12.23.630040 | 2024 | CORTADO: Hill Climbing Optimization for Cell-Type Specific Marker Gene Discovery | AGREE | "we introduce CORTADO, a computational framework ..." | This is a computational marker-discovery paper introducing CORTADO, and the packet does not support benchmarking or direct use of scMAGS. [accepted_agree] |
| 10.3390/biom14040436 | 2024 | GTADC: A Graph-Based Method for Inferring Cell Spatial Distribution in Cancer Tissues | AGREE | "we propose a graph-based deep learning method, GTADC ... enhancing the accuracy of gene selection." | This is a computational method for inferring cell spatial distribution, and the packet does not support a benchmark or direct use claim for scMAGS. [accepted_agree] |

#### NS-Forest v4.0 (3)

| DOI | Year | Title | Audit decision | Evidence (verbatim quote) | Audit reason / validation status |
|---|---|---|---|---|---|
| 10.1093/bioadv/vbaf167 | 2024 | <i>ScaleSC</i> : a superfast and scalable single-cell RNA-seq data analysis pipeline powered by GPU | AGREE | "we developed ScaleSC, a GPU-accelerated solution for large-scale single-cell data processing." | This is a computational single-cell analysis pipeline paper introducing ScaleSC, and the packet gives no explicit NS-Forest v4.0 comparison or use evidence. [accepted_agree] |
| 10.1101/2025.08.14.670406 | 2025 | Constructing and Using Cell Type Populations of the Human Reference Atlas | AGREE | "HRApop ... quantified CTs ... using ... cell type annotation tools." | This is a computational/reference-atlas paper, and the packet provides no explicit NS-Forest v4.0 comparison or use evidence. [accepted_agree] |
| 10.1109/trustcom63139.2024.00294 | 2024 | scE(match): Privacy-Preserving Cluster Matching of Single-Cell Data | DISAGREE | The clients then perform feature selection either via dropout [22] (unsupervised) or NS-Forest [16], [49] (supervised), before communicating these selected features | The packet shows NS-Forest used as a supervised feature-selection component inside scE(match), not benchmarked as a baseline under evaluation. [accepted_disagree] |

#### geneCover (3)

| DOI | Year | Title | Audit decision | Evidence (verbatim quote) | Audit reason / validation status |
|---|---|---|---|---|---|
| 10.1093/bioinformatics/btag051 | 2026 | Inference of marker genes of subtle cell state changes via iLR: iterative logistic regression | AGREE | We present iterative logistic regression (iLR) for the identification of small sets of informative marker genes. | This is a computational marker-gene inference paper, and the packet does not support any direct benchmark or use claim for geneCover. [accepted_agree] |
| 10.1101/2025.11.24.690257 | 2025 | Mapping spatial gradients in spatial transcriptomics data with score matching | AGREE | We introduce SLOPER ... a generative model for learning spatial gradients (vector fields) from ST data. | This is a computational spatial-gradients paper, and the packet does not support any direct benchmark or use claim for geneCover. [accepted_agree] |
| 10.1101/gr.280803.125 | 2025 | Unified integration of spatial transcriptomics across platforms with LLOKI | AGREE | Here, we introduce LLOKI, a novel framework for integrating imaging-based ST data from diverse platforms without requiring shared gene panels. | This is a computational spatial transcriptomics integration paper, and the packet does not support any direct benchmark or use claim for geneCover. [accepted_agree] |

#### scGIST (3)

| DOI | Year | Title | Audit decision | Evidence (verbatim quote) | Audit reason / validation status |
|---|---|---|---|---|---|
| 10.1093/bib/bbaf478 | 2025 | Multi-view gene panel characterization for spatially resolved omics | AGREE | "we developed a two-part framework: (i) panelScope ... and (ii) panelScope-OA ..." | This is a computational panel-characterization paper introducing panelScope and panelScope-OA, and the provided evidence does not explicitly show scGIST use or benchmarking. [accepted_agree] |
| 10.1101/2024.07.05.602236 | 2024 | SOFisher: Reinforcement Learning-Guided Experiment Designs for Spatial Omics | AGREE | we developed SOFisher, a reinforcement learning-based framework ... to guide the selection of the next FOV position | This is a computational spatial-omics experiment-design paper about FOV selection; the packet does not show explicit benchmarking or use of scGIST, so the conservative negative is COMP. [accepted_agree] |
| 10.1101/2025.11.18.689140 | 2025 | CBKMR: A Copula-based Bayesian Kernel Machine Regression Framework for Optimal Marker Detection in Omics Data | AGREE | We propose a copula-based Bayesian kernel machine regression (CBKMR) model ... | This is a computational omics marker-detection paper; the packet does not explicitly benchmark or use scGIST, so the conservative negative is COMP. [accepted_agree] |

### REV (n = 36)

*Review, perspective, protocol overview, or other literature-synthesis article.*

#### NS-Forest v2 (6)

| DOI | Year | Title | Audit decision | Evidence (verbatim quote) | Audit reason / validation status |
|---|---|---|---|---|---|
| 10.1016/j.biosystems.2024.105142 | 2024 | Single-cell classification, analysis, and its application using deep learning techniques | AGREE | Review article | This is a review of single-cell classification and analysis using deep learning. [accepted_agree] |
| 10.1016/j.crbiot.2023.100127 | 2023 | CRISPR/Cas9 genetic screens in hepatocellular carcinoma gene discovery | AGREE | This review provides a summary of HCC carcinogenesis CRISPR/Cas9 screens | This is a review summarizing CRISPR/Cas9 screens in hepatocellular carcinoma. [accepted_agree] |
| 10.1101/gr.276129.121 | 2021 | New frontiers in single-cell genomics | AGREE | In this special issue of Genome Research, we have assembled a diverse collection of research and review articles ... | This is an editorial/introduction for a special issue on single-cell genomics. [accepted_agree] |

| DOI | Year | Title | Audit decision | Evidence (verbatim quote) | Audit reason / validation status |
|---|---|---|---|---|---|
| 10.1186/s41065-025-00528-y | 2025 | Global trends in machine learning applications for single-cell transcriptomics research | AGREE | This study aims to fill research gap through bibliometric and visual analysis | The paper is a bibliometric/trends analysis rather than a primary experimental or methods study. [accepted_agree] |
| 10.48550/arxiv.2301.08436 | 2023 | SpaceTx: A Roadmap for Benchmarking Spatial Transcriptomics Exploration of the Brain | AGREE | Panels were selected ... using a combination of four computational strategies ... NS-Forest37 ... | This is a roadmap/review paper, even though it summarizes prior NS-Forest-based panel selection work. [accepted_agree] |
| 10.48550/arxiv.2506.10037 | 2025 | The Cell Ontology in the age of single-cell omics | DISAGREE | Abstract says "we describe the wide variety of uses of CL ... detail ongoing work ... cover the challenges and future plans". | This is a descriptive overview/perspective on the Cell Ontology and its uses in single-cell omics rather than a primary methods or experimental paper, so REV is the better conservative label. [accepted_disagree] |

#### PERSIST (5)

| DOI | Year | Title | Audit decision | Evidence (verbatim quote) | Audit reason / validation status |
|---|---|---|---|---|---|
| 10.1002/aro2.70016 | 2025 | Technological Innovations and Applications of Spatial Transcriptomics in Livestock and Poultry Research | AGREE | This review offers a comprehensive examination of the evolution and limitations of spatial transcriptomics. | This is a review on spatial transcriptomics in livestock and poultry research. [accepted_agree] |
| 10.1002/eji.70119 | 2025 | Spatially Resolved Profiling of Compartmentalized Muscle and Brain Inflammation | AGREE | In this review, we highlight recent methodological and conceptual advances in spatially resolved profiling | This is a review on spatially resolved profiling of muscle and brain inflammation. [accepted_agree] |
| 10.1016/j.ajpath.2026.01.007 | 2026 | Redefining Pathobiology with Spatial Multi-Omics at the Intersection of Biology, Computation, and Histopathological Assessment | AGREE | The current review explores how spatial technologies ... are poised to refine diagnostic workflows | This is a review on spatial multi-omics and pathology. [accepted_agree] |
| 10.1016/j.biotechadv.2026.108807 | 2026 | Computational methods for spatial multi-omics integration | AGREE | "In this review, we systematically summarize existing spatial multi-omics integration methods ..." | The abstract explicitly states this is a review summarizing spatial multi-omics integration methods. [accepted_agree] |
| 10.48550/arxiv.2301.08436 | 2023 | SpaceTx: A Roadmap for Benchmarking Spatial Transcriptomics Exploration of the Brain | AGREE | we summarize the work of SpaceTx and highlight outstanding challenges | This is a roadmap/review-style paper about benchmarking spatial transcriptomics exploration of the brain. [accepted_agree] |

#### Spapros (5)

| DOI | Year | Title | Audit decision | Evidence (verbatim quote) | Audit reason / validation status |
|---|---|---|---|---|---|
| 10.1093/bib/bbaf723 | 2026 | Advances and challenges in single-cell RNA sequencing data analysis: a comprehensive review | AGREE | Advances and challenges in single-cell RNA sequencing data analysis: a comprehensive review | This is a comprehensive review of single-cell RNA-seq data analysis. [accepted_agree] |
| 10.1161/circresaha.125.325795 | 2026 | Advancing Cardiovascular Research With Single-Cell and Spatial Transcriptomics | AGREE | This review presents an overview of the recently developed applications of single-cell and spatial transcriptomics methods and their impact on cardiovascular research. | This is a review on single-cell and spatial transcriptomics in cardiovascular research. [accepted_agree] |
| 10.1177/01926233251391184 | 2025 | Neuro Omics: Translational Neuropathology Applications of Spatial Technologies From Nonclinical Models to Human Disease | AGREE | Meeting Report | This is a meeting report/review on spatial technologies in translational neuropathology. [accepted_agree] |
| 10.1186/s13073-024-01282-y | 2024 | Spatial multi-omics: novel tools to study the complexity of cardiovascular diseases | AGREE | This review focuses on the recent advancements in spatial multi-omics technologies | This is a review on spatial multi-omics in cardiovascular disease. [accepted_agree] |
| 10.48550/arxiv.2301.08436 | 2023 | SpaceTx: A Roadmap for Benchmarking Spatial Transcriptomics Exploration of the Brain | AGREE | "Here, we summarize the work of SpaceTx ... provide a roadmap for benchmarking ..." | The packet explicitly says this article summarizes SpaceTx and highlights challenges, which makes it a review/roadmap paper. [accepted_agree] |

#### gpsFISH (4)

| DOI | Year | Title | Audit decision | Evidence (verbatim quote) | Audit reason / validation status |
|---|---|---|---|---|---|
| 10.1016/j.ncrops.2024.100025 | 2024 | Applications for single-cell and spatial transcriptomics in plant research | AGREE | In this review, we offer a concise overview of the scRNA-seq and spatial transcriptomics experimental and computational procedures ... | This is a review on single-cell and spatial transcriptomics in plant research. [accepted_agree] |
| 10.1093/noajnl/vdaf193 | 2025 | Targeting the tumor microenvironment in pediatric gliomas: Advances and future directions in immunotherapy | AGREE | Evidence says "This review explores the distinctive features of the pediatric glioma TME ..." | This is a review article on tumor microenvironment targeting in pediatric gliomas and immunotherapy. [accepted_agree] |
| 10.1186/s13073-024-01282-y | 2024 | Spatial multi-omics: novel tools to study the complexity of cardiovascular diseases | AGREE | This review focuses on the recent advancements in spatial multi-omics technologies | This is a review on spatial multi-omics in cardiovascular disease. [accepted_agree] |
| 10.3390/ijms262411819 | 2025 | Why “Where” Matters as Much as “How Much”: Single-Cell and | AGREE | This review explains why the location of gene expression is as | This is a review on single-cell and spatial transcriptomics in plants. |

| DOI | Year | Title | Audit decision | Evidence (verbatim quote) | Audit reason / validation status |
|---|---|---|---|---|---|
| | | Spatial Transcriptomics in Plants | | important as its magnitude ... | [accepted_agree] |

#### scGeneFit (4)

| DOI | Year | Title | Audit decision | Evidence (verbatim quote) | Audit reason / validation status |
|---|---|---|---|---|---|
| 10.1038/s41587-019-0330-9 | 2019 | Exploring a world of a thousand dimensions | AGREE | news & views | This is a News & Views/editorial-style article, which belongs under REV. [accepted_agree] |
| 10.1371/journal.pcbi.1011288 | 2023 | The specious art of single-cell genomics | AGREE | Evidence quote: "PERSPECTIVE". | The packet explicitly marks this as a perspective article, which fits REV. [accepted_agree] |
| 10.3389/fanim.2025.1551244 | 2025 | Integrating omics approaches in livestock biotechnology: innovations in production and reproductive efficiency | AGREE | This review outlines prospect directions and potential novelties in livestock biotechnology | This is a review of omics integration in livestock biotechnology. [accepted_agree] |
| 10.3389/fsysb.2022.910243 | 2022 | Leveraging Systems Immunology to Optimize Diagnosis and Treatment of Inborn Errors of Immunity | AGREE | In this review, we will explore how systems immunology can contribute | This is a review on systems immunology for inborn errors of immunity. [accepted_agree] |

#### COSG (3)

| DOI | Year | Title | Audit decision | Evidence (verbatim quote) | Audit reason / validation status |
|---|---|---|---|---|---|
| 10.1016/j.jmb.2026.169684 | 2026 | Rising Star: Combining Bioinformatics and Experimental Biology to Decoding Non-coding RNAs and RNA Modifications | AGREE | Perspective Rising Star: Combining Bioinformatics and Experimental Biology to Decoding Non-coding RNAs and RNA Modifications | This is a perspective/profile article rather than a primary methods or experimental study. [accepted_agree] |
| 10.1093/bib/bbad416 | 2023 | An in-depth comparison of linear and non-linear joint embedding methods for bulk and single-cell multi-omics | AGREE | Review | The packet identifies this as a review/comparison overview rather than direct benchmarking or use of COSG. [accepted_agree] |
| 10.3390/ijms25031479 | 2024 | Application of Single-Cell Assay for Transposase-Accessible Chromatin with High Throughput Sequencing in Plant Science: Advances, Technical Challenges, and Prospects | AGREE | In this review, we provide an overview of recent progress in scATAC-seq and its application | This is a review article on plant scATAC-seq applications and prospects. [accepted_agree] |

#### NS-Forest v4.0 (2)

| DOI | Year | Title | Audit decision | Evidence (verbatim quote) | Audit reason / validation status |
|---|---|---|---|---|---|
| 10.1186/s41065-025-00528-y | 2025 | Global trends in machine learning applications for single-cell transcriptomics research | AGREE | This study aims to fill research gap through bibliometric and visual analysis | The paper is a bibliometric/trends analysis, so REV is appropriate for this tool-paper relation as well. [accepted_agree] |
| 10.48550/arxiv.2506.10037 | 2025 | The Cell Ontology in the age of single-cell omics | DISAGREE | The Cell Ontology in the age of single-cell omics | This arXiv paper is a perspective/review on the Cell Ontology and its uses in single-cell omics, not an OTHER-type protocol/resource report. [accepted_disagree] |

#### geneBasis (2)

| DOI | Year | Title | Audit decision | Evidence (verbatim quote) | Audit reason / validation status |
|---|---|---|---|---|---|
| 10.1016/j.pathol.2025.10.003 | 2025 | Spatial transcriptomics in bone research: navigating hype and hurdles | AGREE | Here, we review current ST platforms and outline their respective advantages and limitations ... | This is a review on spatial transcriptomics platforms and analysis in bone research. [accepted_agree] |
| 10.48550/arxiv.2301.08436 | 2023 | SpaceTx: A Roadmap for Benchmarking Spatial Transcriptomics Exploration of the Brain | AGREE | "Here, we summarize the work of SpaceTx ... provide a roadmap for benchmarking spatial transcriptomics methods ..." | The packet explicitly describes a roadmap/review article summarizing SpaceTx work and future benchmarking challenges. [accepted_agree] |

#### scMAGS (2)

| DOI | Year | Title | Audit decision | Evidence (verbatim quote) | Audit reason / validation status |
|---|---|---|---|---|---|
| 10.26599/bdma.2025.9020056 | 2025 | A Comprehensive Review of Cell-Type Deconvolution in Spatial Transcriptomic Data | AGREE | A Comprehensive Review of Cell-Type Deconvolution in Spatial Transcriptomic Data | The paper is explicitly a comprehensive review article. [accepted_agree] |
| 10.3389/fcell.2025.1712688 | 2025 | Decoding dental mesenchymal stem cells diversity: single-cell transcriptomics maps heterogeneity in molar development | AGREE | "TYPE Review" | The packet explicitly marks this paper as a review, so REV is appropriate. [accepted_agree] |

#### SCMER (1)

| DOI | Year | Title | Audit decision | Evidence (verbatim quote) | Audit reason / validation status |
|---|---|---|---|---|---|
| 10.1080/00268976.2023.2297818 | 2023 | The statistical physics of single cell genomics | AGREE | This paper delves into the remarkable similarities underlying the fundamental principles governing both systems | This is a review/conceptual article on the statistical physics of single-cell genomics. [accepted_agree] |

#### SMaSH (1)

| DOI | Year | Title | Audit decision | Evidence (verbatim quote) | Audit reason / validation status |
|---|---|---|---|---|---|
| 10.48550/arxiv.2312.06082 | 2023 | XAI meets Biology: A Comprehensive Review of Explainable AI in Bioinformatics Applications | AGREE | "This review provides a comprehensive analysis of various XAI techniques and their applications ..." | This is a review article on explainable AI in bioinformatics applications. [accepted_agree] |

#### scGIST (1)

| DOI | Year | Title | Audit decision | Evidence (verbatim quote) | Audit reason / validation status |
|---|---|---|---|---|---|
| 10.1016/j.ajpath.2026.01.007 | 2026 | Redefining Pathobiology with Spatial Multi-Omics at the Intersection of Biology, Computation, and Histopathological Assessment | AGREE | The current review explores how spatial technologies | This is a review article on spatial multi-omics and pathology. [accepted_agree] |

### NON-ST (n = 108)

*Paper outside spatial transcriptomics.*

#### COSG (66)

| DOI | Year | Title | Audit decision | Evidence (verbatim quote) | Audit reason / validation status |
|---|---|---|---|---|---|
| 10.1002/advs.202511856 | 2025 | Immunoregulatory Roles of Tumor-Originated Pericytes Identified by Single-Cell Analysis in Glioblastoma | DISAGREE | "... the COSG_markers function ... provided subgroup-specific genes." | The packet describes a single-cell glioblastoma study without spatial transcriptomics scope, and COSG is used only for subgroup-specific marker genes. [accepted_disagree] |
| 10.1002/pdi3.70027 | 2025 | Single-Cell Profiling of Pediatric High-Grade Gliomas Reveals OPC-Like Subpopulations Driving Tumorigenic Lineage Transitions | AGREE | Single-Cell Profiling of Pediatric High-Grade Gliomas Reveals OPC-Like Subpopulations ... | This is a non-spatial pediatric glioma single-cell profiling paper, not a spatial transcriptomics study. [accepted_agree] |

| DOI | Year | Title | Audit decision | Evidence (verbatim quote) | Audit reason / validation status |
|---|---|---|---|---|---|
| 10.1007/s00432-023-05403-9 | 2023 | Integration and deconvolution methodology deciphering prognosis-related signatures in lung adenocarcinoma | DISAGREE | Evidence describes a machine learning classifier using prognosis-related genes for LUAD risk stratification. | This is a lung adenocarcinoma prognosis/deconvolution study rather than a spatial transcriptomics or COSG-focused methods paper. [accepted_disagree] |
| 10.1007/s10815-024-03143-4 | 2024 | Rescue in vitro maturation using ovarian support cells of human oocytes from conventional stimulation cycles yields oocytes with improved nuclear maturation and transcriptomic resemblance to in vivo matured oocytes | AGREE | The OSC-IVM platform is a novel tool for rescue maturation of human oocytes ... | This is a non-spatial reproductive biology/oocyte maturation study, not a spatial transcriptomics study. [accepted_agree] |
| 10.1016/j.bbadis.2025.167918 | 2025 | Deciphering the regulatory networks of human male germline development from embryo to adulthood | AGREE | Deciphering the regulatory networks of human male germline development from embryo to adulthood | This is a germline development regulatory-network paper without a spatial transcriptomics study component. [accepted_agree] |
| 10.1016/j.bbrc.2025.151687 | 2025 | Nerve infiltration of bladder cancer predicts response to immunotherapy | AGREE | Evidence quote: NOT FOUND; abstract describes single-nucleus RNA-seq and denervation experiments. | The abstract describes public databases, mouse tumor models, and single-nucleus RNA-seq, but not a spatial transcriptomics study. [accepted_agree] |
| 10.1016/j.ccell.2025.01.001 | 2025 | Contrasting cytotoxic and regulatory T cell responses underlying distinct clinical outcomes to anti-PD-1 plus lenvatinib therapy in cancer | AGREE | Contrasting cytotoxic and regulatory T cell responses underlying distinct clinical outcomes to anti-PD-1 plus lenvatinib therapy in cancer | This is a cancer immunotherapy response paper, not a spatial transcriptomics study. [accepted_agree] |
| 10.1016/j.cell.2024.07.016 | 2024 | LAG-3 and PD-1 synergize on CD8+ T cells to drive T cell exhaustion and hinder autocrine IFN-γ-dependent anti-tumor immunity | AGREE | NOT FOUND | This is a non-spatial immunology/tumor-immunity paper; the packet does not indicate any spatial transcriptomics panel use of COSG. [accepted_agree] |
| 10.1016/j.celrep.2026.117076 | 2026 | Collagenesis orchestrates mechanoadaptive development and homeostasis of intervertebral discs | AGREE | Collagenesis orchestrates mechanoadaptive development and homeostasis of intervertebral discs | This is an intervertebral disc development/homeostasis paper, not a spatial transcriptomics study. [accepted_agree] |
| 10.1016/j.devcel.2023.11.007 | 2023 | Murine trophoblast organoids as a model for trophoblast development and CRISPR-Cas9 screening | AGREE | Murine trophoblast organoids as a model for trophoblast development and CRISPR-Cas9 screening | This is a non-spatial trophoblast organoid development and CRISPR-screening paper, not a spatial transcriptomics study. [accepted_agree] |
| 10.1016/j.forsciint.2025.112646 | 2025 | Single-nucleus RNA sequencing and machine learning identify | AGREE | Single-nucleus RNA sequencing and machine learning identify | This is a forensic biomarker-identification paper without a |

| DOI | Year | Title | Audit decision | Evidence (verbatim quote) | Audit reason / validation status |
|---|---|---|---|---|---|
| | | CACNA1A as a myocyte-specific biomarker for sudden unexplained death in schizophrenia | | CACNA1A as a myocyte-specific biomarker for sudden unexplained death in schizophrenia | spatial transcriptomics study component. [accepted_agree] |
| 10.1016/j.isci.2025.114328 | 2025 | United multi-omics and machine learning refine regulatory T cell-defined hepatocellular carcinoma subtypes | DISAGREE | This paper analyzes existing, publicly available data ... GEO ... HCCDB ... ArrayExpress ... and TCGA | This is a non-spatial hepatocellular carcinoma data-analysis study using existing datasets, not a computational methods paper about COSG. [accepted_disagree] |
| 10.1016/j.jare.2025.05.038 | 2025 | Dynamic heterogeneity towards drug resistance in AML cells is primarily driven by epigenomic mechanism unveiled by multi-omics analysis | AGREE | Dynamic heterogeneity towards drug resistance in AML cells is primarily driven by epigenomic mechanism ... | This is a non-spatial AML multi-omics drug-resistance paper, not a spatial transcriptomics study. [accepted_agree] |
| 10.1016/j.jgg.2024.11.005 | 2024 | Integrated analysis and systematic characterization of the regulatory network for human germline development | AGREE | Integrated analysis and systematic characterization of the regulatory network for human germline development | This is a human germline development regulatory-network analysis paper without a spatial transcriptomics study component. [accepted_agree] |
| 10.1016/j.jhazmat.2025.137375 | 2025 | Insight into perfluorooctanoic acid-induced impairment of mouse embryo implantation via single-cell RNA-seq | AGREE | Insight into perfluorooctanoic acid-induced impairment of mouse embryo implantation via single-cell RNA-seq | This is a non-spatial toxicology/reproduction scRNA-seq paper, not a spatial transcriptomics study. [accepted_agree] |
| 10.1016/j.lfs.2024.123236 | 2024 | Integrating multi-omics data reveals neuroblastoma subtypes in the tumor microenvironment | AGREE | Integrating multi-omics data reveals neuroblastoma subtypes in the tumor microenvironment | This is a neuroblastoma tumor microenvironment multi-omics paper without a spatial transcriptomics study component. [accepted_agree] |
| 10.1016/j.molp.2024.01.009 | 2024 | Callus proliferation-induced hypoxic microenvironment decreases shoot regeneration competence in Arabidopsis | AGREE | Callus proliferation-induced hypoxic microenvironment decreases shoot regeneration competence in Arabidopsis | This is a non-spatial plant regeneration biology paper, not a spatial transcriptomics study. [accepted_agree] |
| 10.1016/j.neuron.2025.05.002 | 2025 | An enhancer-AAV toolbox to target and manipulate distinct interneuron subtypes | AGREE | "we identified top cell-type-specific enhancers using ... COSG ... and selected the top enhancer candidates for in vivo testing." | This is a non-spatial experimental enhancer/AAV toolbox paper; COSG was used to identify enhancers for in vivo testing, not for targeted spatial panel selection. [accepted_agree] |
| 10.1016/j.scib.2023.09.021 | 2023 | Single-cell dynamics of liver development in postnatal pigs | AGREE | Single-cell dynamics of liver development in postnatal pigs | This is a postnatal pig liver single-cell atlas paper, not a spatial transcriptomics study. [accepted_agree] |

| DOI | Year | Title | Audit decision | Evidence (verbatim quote) | Audit reason / validation status |
|---|---|---|---|---|---|
| 10.1038/s41393-025-01077-x | 2025 | BDNF-GABA signaling in astrocytes: enhancing neural repair after SCI through MSC therapies | DISAGREE | STUDY DESIGN: An integrated bioinformatics data study. | This is an integrated bioinformatics disease study on spinal cord injury, not a computational methods paper and not a spatial transcriptomics study. [accepted_disagree] |
| 10.1038/s41393-025-01103-y | 2025 | Microglial landscape and signaling in spinal cord injury | DISAGREE | Study design: An integrated bioinformatics data study. | This is an integrated bioinformatics disease study on spinal cord injury, not a computational methods paper and not a spatial transcriptomics study. [accepted_disagree] |
| 10.1038/s41398-023-02665-2 | 2023 | Does decreased autophagy and dysregulation of LC3A in astrocytes play a role in major depressive disorder? | AGREE | Does decreased autophagy and dysregulation of LC3A in astrocytes play a role in major depressive disorder? | This is a non-spatial psychiatry/snRNA-seq paper, not a spatial transcriptomics study. [accepted_agree] |
| 10.1038/s41467-023-43462-1 | 2023 | Low-dose radiotherapy combined with dual PD-L1 and VEGFA blockade elicits antitumor response in hepatocellular carcinoma mediated by activated intratumoral CD8+ exhausted-like T cells | AGREE | Low-dose radiotherapy combined with dual PD-L1 and VEGFA blockade elicits antitumor response in hepatocellular carcinoma | This is an immuno-oncology hepatocellular carcinoma response paper without a spatial transcriptomics study component. [accepted_agree] |
| 10.1038/s41588-025-02475-w | 2026 | A microenvironment-determined risk continuum refines subtyping in meningioma and reveals determinants of machine learning-based tumor classification | DISAGREE | Abstract ... molecular meningioma classification ... immunohistochemistry ...; evidence quote: NOT FOUND | The abstract is a tumor-classification and immunohistochemistry study, not a spatial transcriptomics study, and the packet has no COSG passage. [accepted_disagree] |
| 10.1038/s41598-024-65090-5 | 2024 | Comprehensive analysis of endoplasmic reticulum stress related signature in head and neck squamous carcinoma | AGREE | Comprehensive analysis of endoplasmic reticulum stress related signature in head and neck squamous carcinoma | This is a head and neck squamous carcinoma signature-analysis paper without a spatial transcriptomics study component. [accepted_agree] |
| 10.1038/s43587-024-00631-1 | 2024 | Aging atlas reveals cell-type-specific effects of pro-longevity strategies | AGREE | Aging atlas reveals cell-type-specific effects of pro-longevity strategies | This is a non-spatial aging single-cell atlas paper, not a spatial transcriptomics study. [accepted_agree] |
| 10.1038/s44161-024-00464-6 | 2024 | Strategies for arterial graft optimization at the single-cell level | AGREE | Evidence quote: NOT FOUND; abstract describes single-cell-level analysis of arterial grafts. | This is a single-cell arterial-graft optimization study, not a spatial transcriptomics study. [accepted_agree] |
| 10.1080/07853890.2024.2443568 | 2024 | A diagnostic signatures for intervertebral disc degeneration using TNFAIP6 and COL6A2 | AGREE | Evidence quote: NOT FOUND; title and abstract describe single-cell | This is a diagnostic-signature study using single-cell and bulk RNA-seq, not a spatial transcriptomics |

| DOI | Year | Title | Audit decision | Evidence (verbatim quote) | Audit reason / validation status |
|---|---|---|---|---|---|
| | | based on single-cell RNA-seq and bulk RNA-seq analyses | | RNA-seq and bulk RNA-seq analyses. | study. [accepted_agree] |
| 10.1101/2023.03.27.534477 | | Improved rescue of immature oocytes obtained from conventional gonadotropin stimulation cycles via human induced pluripotent stem cell-derived ovarian support cell co-culture | AGREE | Improved rescue of immature oocytes obtained from conventional gonadotropin stimulation cycles | This is an ovarian support cell co-culture/oocyte rescue study rather than a spatial transcriptomics paper. [accepted_agree] |
| 10.1101/2023.11.09.566505 | 2023 | Targeting CD74-positive macrophages improves neoadjuvant therapy in cervical cancer as revealed by single-cell transcriptomics analysis | AGREE | Targeting CD74-positive macrophages improves neoadjuvant therapy in cervical cancer as revealed by single-cell transcriptomics analysis | This is a non-spatial cervical cancer single-cell analysis paper, not a spatial transcriptomics study. [accepted_agree] |
| 10.1101/2023.12.03.569806 | 2023 | Landscape and functional repertoires of long noncoding RNAs in the pan-cancer tumor microenvironment using single-nucleus total RNA sequencing | DISAGREE | Landscape and functional repertoires of long noncoding RNAs in the pan-cancer tumor microenvironment using single-nucleus total RNA sequencing | This pan-cancer single-nucleus RNA-seq analysis is not a spatial transcriptomics study, and the packet does not show any close-context COSG-linked targeted panel selection. [accepted_disagree] |
| 10.1101/2024.06.24.600550 | 2024 | Lipidic and senescent macrophages predict progression and response to combinatorial immunotherapy in triple-negative breast cancer | DISAGREE | we re-analyzed sc-RNA seq data ... and identified lipidic and senescent macrophages in TNBC patients | This is an experimental non-spatial triple-negative breast cancer single-cell study, not a computational methods paper and not a targeted spatial panel-selection study. [accepted_disagree] |
| 10.1101/2024.11.07.622485 | 2024 | <i>FOXP</i> Genes Regulate Purkinje Cell Diversity in Cerebellar Development and Evolution | DISAGREE | Cell type-specific genes were identified using the COSG package 56. | The packet only shows COSG used to identify cell-type-specific genes in a developmental single-cell study, with no spatial transcriptomics panel-selection link. [accepted_disagree] |
| 10.1101/2025.01.26.634963 | 2025 | Single-Cell Genomics Elucidates Molecular Variations and Regulatory Mechanisms in Circulating Immune Cells | AGREE | Single-Cell Genomics Elucidates Molecular Variations and Regulatory Mechanisms in Circulating Immune Cells | This is a circulating immune-cell multi-omics atlas, not a spatial transcriptomics study. [accepted_agree] |
| 10.1101/2025.05.09.652066 | 2025 | Characterization of hyperoxia-induced senescent lung macrophages in neonatal mice | DISAGREE | "Here, we reanalyzed scRNA-seq datasets ... of senescent lung cells ..." | This is a reanalysis of scRNA-seq datasets in neonatal lung macrophages, not a spatial transcriptomics study. [accepted_disagree] |

| DOI | Year | Title | Audit decision | Evidence (verbatim quote) | Audit reason / validation status |
|---|---|---|---|---|---|
| 10.1101/2025.07.09.663835 | 2025 | Granulin <sup>+</sup> macrophages promote lineage plasticity in prostate cancer through paracrine signaling loops | DISAGREE | (paraphrased) its signature genes were obtained using COSG[86] | The abstract describes a non-spatial-transcriptomics cancer study; COSG is only used to obtain signature genes, not for a targeted spatial panel. [accepted_disagree] |
| 10.1126/science.adt3130 | 2026 | Chinese Immune Multi-Omics Atlas | AGREE | Chinese Immune Multi-Omics Atlas | This is a circulating immune-cell multi-omics atlas, not a spatial transcriptomics study. [accepted_agree] |
| 10.1126/science.adu7265 | 2026 | Family single-cell atlases reveal pig pregnancy and fetal growth restriction critical cell types | AGREE | Evidence quote: NOT FOUND; abstract describes a 2.56 million-cell transcriptomic atlas. | The abstract clearly describes a large single-cell atlas of pig pregnancy and fetal growth restriction, not a spatial transcriptomics study. [accepted_agree] |
| 10.1136/jitc-2024-009024 | 2024 | Targeting tumor-associated macrophage-derived CD74 improves efficacy of neoadjuvant chemotherapy in combination with PD-1 blockade for cervical cancer | AGREE | Targeting tumor-associated macrophage-derived CD74 improves efficacy of neoadjuvant chemotherapy in combination with PD-1 blockade for cervical cancer | This is a cervical cancer therapy/immune microenvironment paper without a spatial transcriptomics study component. [accepted_agree] |
| 10.1136/jitc-2024-010871 | 2025 | Immunosuppressive microenvironment of liver restrains chemotherapeutic efficacy in triple-negative breast cancer | AGREE | Immunosuppressive microenvironment of liver restrains chemotherapeutic efficacy in triple-negative breast cancer | This is an immunotherapy/tumor microenvironment paper without a spatial transcriptomics study component. [accepted_agree] |
| 10.1186/s12864-024-11030-6 | 2024 | Cross-species single-cell analysis reveals divergence and conservation of peripheral blood mononuclear cells | AGREE | Cross-species single-cell analysis reveals divergence and conservation of peripheral blood mononuclear cells | This is a non-spatial cross-species PBMC single-cell analysis paper, not a spatial transcriptomics study. [accepted_agree] |
| 10.1186/s12883-026-04720-z | 2026 | Novel VSMC-associated biomarkers in intracranial aneurysm pathogenesis: a multi-omics and machine learning study | DISAGREE | publicly available gene expression datasets related to intracranial aneurysms were retrieved ... through comprehensive bioinformatics approaches | This is a non-spatial biomarker-discovery study using public expression datasets and machine learning, not a computational methods paper about COSG. [accepted_disagree] |
| 10.1186/s12967-023-03961-7 | 2023 | Abnormal signal pathways and tumor heterogeneity in osteosarcoma | DISAGREE | it should be noted that the study we examined is limited to bioinformatics analysis | This is a non-spatial osteosarcoma bioinformatics study, not a computational methods paper about COSG. [accepted_disagree] |
| 10.1186/s12967-023-04151-1 | 2023 | The potential crosstalk between tumor and plasma cells and its association with clinical outcome | DISAGREE | "COSG ... to obtain the top 100 marker genes ..." | The packet describes a tumor/plasma-cell crosstalk study without spatial transcriptomics scope, and COSG is only used to |

| DOI | Year | Title | Audit decision | Evidence (verbatim quote) | Audit reason / validation status |
|---|---|---|---|---|---|
| | | and immunotherapy response in bladder cancer | | | obtain marker genes. [accepted_disagree] |
| 10.1186/s12967-025-06150-w | 2025 | Single-cell profiling reveals a reduced epithelial defense system, decreased immune responses and the immune regulatory roles of different fibroblast subpopulations in chronic atrophic gastritis | AGREE | Evidence quote: NOT FOUND; abstract describes cellular changes in CAG lesions from single-cell profiling. | The abstract describes a single-cell profiling study in chronic atrophic gastritis without a spatial transcriptomics experiment. [accepted_agree] |
| 10.1186/s12967-025-06682-1 | 2025 | Pathway-based cancer transcriptome deciphers a high-resolution intrinsic heterogeneity within bladder cancer classification | AGREE | Pathway-based cancer transcriptome deciphers a high-resolution intrinsic heterogeneity within bladder cancer classification | This is a bladder cancer transcriptome classification paper, not a spatial transcriptomics study. [accepted_agree] |
| 10.1186/s13059-024-03340-5 | 2024 | Microsatellite instability at U2AF-binding polypyrimidic tract sites perturbs alternative splicing during colorectal cancer initiation | AGREE | Microsatellite instability at U2AF-binding polypyrimidic tract sites perturbs alternative splicing during colorectal cancer initiation | This is a colorectal cancer alternative-splicing study rather than a spatial transcriptomics study. [accepted_agree] |
| 10.1186/s13148-024-01800-0 | 2024 | Single-cell transcriptomics reveal the prognostic roles of epithelial and T cells and DNA methylation-based prognostic models in pancreatic cancer | AGREE | Single-cell transcriptomics reveal the prognostic roles of epithelial and T cells ... in pancreatic cancer | This is a non-spatial pancreatic cancer scRNA-seq/epigenetics prognostic-model paper, not a spatial transcriptomics study. [accepted_agree] |
| 10.1186/s40478-025-02038-4 | 2025 | Distinct characteristics of brain metastasis in lung adenocarcinoma: development of high-confidence cell lines | AGREE | Distinct characteristics of brain metastasis in lung adenocarcinoma: development of high-confidence cell lines | This is a brain-metastasis cell-line development paper without a spatial transcriptomics study component. [accepted_agree] |
| 10.21203/rs.3.rs-3960911/v1 | 2024 | Community effort to unravel the complex human neuronal landscape exemplified for the striatum | AGREE | Community effort to unravel the complex human neuronal landscape exemplified for the striatum | This is a neuronal landscape/community atlas paper without a spatial transcriptomics study component. [accepted_agree] |
| 10.21203/rs.3.rs-4241968/v1 | 2024 | Identification of Key Candidate Genes Related to Cartilage development during Murine embryonic limb development by single cell RNA-sequencing and Experimental Confirmation | AGREE | Identification of Key Candidate Genes Related to Cartilage development ... by single cell RNA-sequencing | This is a non-spatial murine embryonic limb/cartilage scRNA-seq study, not a spatial transcriptomics study. [accepted_agree] |
| 10.21203/rs.3.rs-4730227/v1 | 2024 | Single-cell profiling reveals phenotypic changes in chronic atrophic gastritis | AGREE | Evidence quote: NOT FOUND; abstract says "Single-cell profiling reveals ... chronic atrophic gastritis". | This is a single-cell profiling study of chronic atrophic gastritis, not a spatial transcriptomics study. [accepted_agree] |
| 10.21203/rs.3.rs-6389993/v1 | 2025 | Evaluating Deconvolution Methods Using Real Bulk RNA-expression | DISAGREE | "This study did not generate any new data. All scRNA-seq and bulk | This is a bulk RNA deconvolution benchmark using existing scRNA- |

| DOI | Year | Title | Audit decision | Evidence (verbatim quote) | Audit reason / validation status |
|---|---|---|---|---|---|
| | | Data for Robust Prognostic Insights in Pan-Cancer | | data used in this study are listed ..." | seq and bulk data, not a spatial transcriptomics study or a COSG-focused methods paper. [accepted_disagree] |
| 10.2147/itt.s451199 | 2024 | Characterizing CD8+ TEMRA Cells in CP/CPPS Patients: Insights from Targeted Single-Cell Transcriptomic and Functional Investigations | AGREE | Characterizing CD8+ TEMRA Cells in CP/CPPS Patients: Insights from Targeted Single-Cell Transcriptomic ... | This is a targeted single-cell immunology study, not a spatial transcriptomics study. [accepted_agree] |
| 10.31083/fbl48370 | 2026 | Neonatal Hyperoxia Induces Metabolic Reprogramming in Senescent Alveolar Macrophages, Leading to Persistent Lung Injury | AGREE | Neonatal Hyperoxia Induces Metabolic Reprogramming in Senescent Alveolar Macrophages, Leading to Persistent Lung Injury | This is a lung injury/macrophage biology paper without a spatial transcriptomics study component. [accepted_agree] |
| 10.3389/fcell.2025.1515681 | 2025 | Hypoxia signature derived from tumor-associated endothelial cells predict prognosis in gastric cancer | AGREE | Hypoxia signature derived from tumor-associated endothelial cells predict prognosis in gastric cancer | This is a non-spatial gastric cancer single-cell prognostic-signature paper, not a spatial transcriptomics study. [accepted_agree] |
| 10.3389/fimmu.2022.1050951 | 2022 | Complex interaction and heterogeneity among cancer stem cells in head and neck squamous cell carcinoma revealed by single-cell sequencing | DISAGREE | We depicted the landscape of HNSCC transcriptome profile by single-cell RNA-sequencing analysis of 20 HNSCC tissues | This is an experimental non-spatial single-cell cancer study, not a computational methods paper and not a targeted spatial panel-selection study. [accepted_disagree] |
| 10.3389/fimmu.2024.1310376 | 2024 | Revealing molecular and cellular heterogeneity in hypopharyngeal carcinogenesis through single-cell RNA and TCR/BCR sequencing | AGREE | Revealing molecular and cellular heterogeneity in hypopharyngeal carcinogenesis through single-cell RNA and TCR/BCR sequencing | This is a hypopharyngeal carcinogenesis single-cell/TCR-BCR sequencing paper without a spatial transcriptomics study component. [accepted_agree] |
| 10.3389/fimmu.2024.1363517 | 2024 | A division-of-labor mode contributes to the cardioprotective potential of mesenchymal stem/stromal cells in heart failure post myocardial infarction | AGREE | NOT FOUND | This is a non-spatial scRNA-seq/MSC cardioprotection paper, not a spatial transcriptomics study. [accepted_agree] |
| 10.3389/fonc.2025.1552841 | 2025 | Single-cell RNA profiling of colorectal granular-type laterally spreading tumor uncovers progression trajectory toward carcinoma and transcriptional signatures favoring lateral morphogenesis | AGREE | Single-cell RNA profiling of colorectal granular-type laterally spreading tumor uncovers progression trajectory toward carcinoma | This is a single-cell RNA profiling cancer paper without a spatial transcriptomics study component, so NON-ST is appropriate. [accepted_agree] |
| 10.3389/fphar.2025.1634985 | 2025 | Integrative analysis of lactylation related genes in prostate cancer: unveiling heterogeneity through | AGREE | Integrative analysis of lactylation related genes in prostate cancer ... | This is a non-spatial prostate cancer integrative single-cell/bulk analysis paper, not a spatial |

| DOI | Year | Title | Audit decision | Evidence (verbatim quote) | Audit reason / validation status |
|---|---|---|---|---|---|
| | | single-cell RNA-seq, bulk RNA-seq and machine learning | | | transcriptomics study. [accepted_agree] |
| 10.3390/ani14223263 | 2024 | Single-Nucleus RNA Sequencing Reveals the Transcriptome Profiling of Ovarian Cells in Adolescent Cyprinus carpio | AGREE | Single-Nucleus RNA Sequencing Reveals the Transcriptome Profiling of Ovarian Cells in Adolescent Cyprinus carpio | This is a single-nucleus ovary transcriptomics study with no spatial transcriptomics component, so NON-ST is appropriate. [accepted_agree] |
| 10.3390/biology15010095 | 2026 | Single-Cell Transcriptomic Profiling Reveals That Macrophage-Induced Angiogenesis Contributes to Immunotherapy Resistance in Hepatocellular Carcinoma | DISAGREE | We integrated single-cell RNA sequencing (scRNA-seq) data from 328 samples of 63 HCC patients | This is an experimental non-spatial single-cell hepatocellular carcinoma study, not a computational methods paper and not a targeted spatial panel-selection study. [accepted_disagree] |
| 10.48550/arxiv.2106.06468 | 2021 | Locally Sparse Neural Networks for Tabular Biomedical Data | DISAGREE | Locally Sparse Neural Networks for Tabular Biomedical Data | This is a general machine-learning paper for tabular biomedical data, not a spatial transcriptomics study or relevant single-cell methods paper. [accepted_disagree] |
| 10.64898/2026.02.12.703707 | 2026 | A stem cell aging clock links biological age deviation to clinical outcome in acute myeloid leukemia | DISAGREE | we developed a machine learning-based stem cell aging clock from 84 donors to predict chronological age from HSC/MPP transcriptomes | This is an experimental single-cell aging-clock study in hematopoietic stem cells, not a computational methods paper and not a spatial transcriptomics study. [accepted_disagree] |
| 10.7150/thno.112339 | 2025 | Pharmacological and genetic inhibition of ARG2 in CXCR2<sup>Hi</sup> myeloid-derived suppressor cells combats sepsis-induced lymphopenia | AGREE | Evidence quote: NOT FOUND; abstract describes key molecules within heterogeneous MDSCs in sepsis. | This is a sepsis/MDSC single-cell heterogeneity study, not a spatial transcriptomics study. [accepted_agree] |

#### NS-Forest v2 (12)

| DOI | Year | Title | Audit decision | Evidence (verbatim quote) | Audit reason / validation status |
|---|---|---|---|---|---|
| 10.1002/jlb.4cova0121-084rr | 2021 | Corticosteroid treatment in COVID-19 modulates host inflammatory responses and transcriptional signatures of immune dysregulation | AGREE | Evidence quote: NOT FOUND; abstract describes host inflammatory responses and transcriptional signatures in COVID-19. | This is a COVID-19 inflammatory-response transcriptomics study, not a spatial transcriptomics study. [accepted_agree] |
| 10.1016/j.jid.2023.06.211 | 2023 | Cross-Comparison of Inflammatory Skin Disease Transcriptomics Identifies PTEN as a Pathogenic | AGREE | the NS-Forest analysis identified minimum sets of marker genes ... validated in separate | This is a skin-disease transcriptomics/classifier paper without a spatial transcriptomics |

| DOI | Year | Title | Audit decision | Evidence (verbatim quote) | Audit reason / validation status |
|---|---|---|---|---|---|
| | | Disease Classifier in Cutaneous Lupus Erythematosus | | transcriptomics experiments and immunohistochemical experiments | study component. [accepted_agree] |
| 10.1016/j.xgen.2025.101007 | 2025 | Single-nucleus transcriptome atlas of orbitofrontal cortex in ALS with a deep learning-based decoding of alternative polyadenylation mechanisms | AGREE | Single-nucleus transcriptome atlas of orbitofrontal cortex in ALS | This is a single-nucleus transcriptome atlas paper, not a spatial transcriptomics study. [accepted_agree] |
| 10.1073/pnas.2304730120 | 2023 | split-intein Gal4 provides intersectional genetic labeling that is repressible by Gal80 | AGREE | "Using NS-Forest v2 to guide gene pair selection, we generated transgenic split-intein Gal4 lines ..." | This is a non-spatial experimental Drosophila labeling paper; NS-Forest guided gene-pair selection for transgenic lines, not spatial panel design. [accepted_agree] |
| 10.1093/intbio/zyae020 | 2024 | Machine learning ranking of plausible (un)explored synergistic gene combinations using sensitivity indices of time series measurements of Wnt signaling pathway | DISAGREE | Machine learning ranking of plausible (un)explored synergistic gene combinations using sensitivity indices of time series measurements of Wnt signaling pathway | This is a non-spatial signaling-pathway/gene-combination ranking paper, not a spatial transcriptomics study. [accepted_disagree] |
| 10.1101/2022.07.18.500480 | 2022 | Predicting the Reprogrammability of Human Cells Based on Transcriptome Data and SGD Classifier with Elastic-Net Regularization | DISAGREE | "constructed a ... classifier ... for predicting whether the cell lines are reprogrammable." | This is a transcriptome-based cell reprogramming prediction study, not a spatial or single-cell methods paper tied to NS-Forest. [accepted_disagree] |
| 10.1101/2025.08.02.668295 | 2025 | Human retinal organoid single-cell atlas allows to reconstruct retinal development at high resolution and identify nature restricted transcriptional states in vitro | DISAGREE | "We ... integrated existing ... single-cell RNA sequencing data to establish a human retinal organoid cell atlas ..." | This is a single-cell retinal organoid atlas paper, not a computational methods paper or spatial transcriptomics study. [accepted_disagree] |
| 10.3389/fimmu.2021.690470 | 2021 | Machine Learning-Based Single Cell and Integrative Analysis Reveals That Baseline mDC Predisposition Correlates With Hepatitis B Vaccine Antibody Response | AGREE | Marker genes were produced "for use in downstream quantitative PCR assays and semantic representations". | The packet clearly describes a non-spatial single-cell vaccine-response study; marker genes were for downstream assays, not spatial panel selection. [accepted_agree] |
| 10.3390/app12041850 | 2022 | Analyzing RNA-Seq Gene Expression Data Using Deep Learning Approaches for Cancer Classification | DISAGREE | Abstract and evidence describe RNA-Seq values converted to images and classified with deep learning across cancer types. | This is an RNA-seq deep learning cancer classification paper, not a single-cell or spatial transcriptomics paper. [accepted_disagree] |
| 10.3390/biology15070588 | 2026 | An AI-Enabled Single-Cell Transcriptomic Analysis Pipeline for Gene Signature Discovery in Natural Killer Cells Linked to | DISAGREE | Evidence and abstract describe an integrative single-cell analysis pipeline, not spatial transcriptomics. | This is an AI-enabled single-cell transcriptomic analysis pipeline for NK-cell signatures, not a spatial transcriptomics study. [accepted_disagree] |

| DOI | Year | Title | Audit decision | Evidence (verbatim quote) | Audit reason / validation status |
|---|---|---|---|---|---|
| | | Remission Outcomes in Chronic Myeloid Leukemia | | | |
| 10.64898/2026.03.26.714500 | 2026 | Single-cell lung eQTL dataset of Asian never-smokers highlights the roles of alveolar cells in lung cancer etiology | AGREE | Evidence quote: NOT FOUND; abstract describes single-cell lung eQTL analyses. | This is a single-cell eQTL study, not a spatial transcriptomics study. [accepted_agree] |
| 10.7717/peerj.16192 | 2023 | SaPt-CNN-LSTM-AR-EA: a hybrid ensemble learning framework for time series-based multivariate DNA sequence prediction | DISAGREE | SaPt-CNN-LSTM-AR-EA: a hybrid ensemble learning framework for time series-based multivariate DNA sequence prediction | This is a non-spatial DNA-sequence/time-series prediction methods paper, not a spatial transcriptomics or relevant computational citation context for NS-Forest v2. [accepted_disagree] |

#### scGeneFit (10)

| DOI | Year | Title | Audit decision | Evidence (verbatim quote) | Audit reason / validation status |
|---|---|---|---|---|---|
| 10.1016/j.acha.2023.03.001 | 2023 | Lie PCA: Density estimation for symmetric manifolds | DISAGREE | Lie PCA: Density estimation for symmetric manifolds | This is a general mathematics/machine-learning paper on symmetric manifolds, not a spatial transcriptomics study or relevant single-cell methods paper. [accepted_disagree] |
| 10.1016/j.cels.2025.101405 | 2025 | Identifying an optimal perturbation to induce a desired cell state by generative deep learning | AGREE | Identifying an optimal perturbation to induce a desired cell state by generative deep learning | This is a cell-state perturbation/generative deep learning paper, not a spatial transcriptomics study. [accepted_agree] |
| 10.1016/j.compbiomed.2021.105208 | 2022 | Ensemble feature selection for stable biomarker identification and cancer classification from microarray expression data | DISAGREE | Abstract describes stable biomarker identification and cancer classification from microarray expression data. | This is a microarray biomarker-identification and cancer-classification paper, not a single-cell or spatial transcriptomics methods paper. [accepted_disagree] |
| 10.1016/j.isci.2021.103292 | 2021 | How many markers are needed to robustly determine a cell's type? | DISAGREE | Abstract discusses marker replicability in BICCN datasets; evidence says only that the paper analyzes existing public data. | This paper analyzes marker replicability in existing single-cell atlas data and is not a spatial transcriptomics or computational benchmarking paper about scGeneFit. [accepted_disagree] |
| 10.1101/2024.10.15.618479 | 2024 | Interactive design and validation of antibody panels using single-cell RNA-seq atlases | DISAGREE | Abstract ... design of antibody panels ... We use Cytomarker to spatially profile human mammary | This paper is about antibody panel design and spatial profiling, not spatial transcriptomics; the packet |

| DOI | Year | Title | Audit decision | Evidence (verbatim quote) | Audit reason / validation status |
|---|---|---|---|---|---|
| | | | | tissue subpopulations ...; evidence quote: NOT FOUND | also lacks any explicit scGeneFit passage. [accepted_disagree] |
| 10.1109/tit.2019.2962681 | 2018 | SqueezeFit: Label-Aware Dimensionality Reduction by Semidefinite Programming | DISAGREE | SqueezeFit: Label-Aware Dimensionality Reduction by Semidefinite Programming | This is a general dimensionality-reduction/theory paper, not a spatial transcriptomics study or relevant single-cell methods paper. [accepted_disagree] |
| 10.1176/appi.ajp.20230602 | 2024 | Translational Insights From Cell Type Variation Across Amygdala Subnuclei in Rhesus Monkeys and Humans | DISAGREE | Together, these findings shed light on the composition of the amygdala and identify specific cell types ... | The packet does not indicate a spatial-transcriptomics study; this appears to be a cell-type variation/translational psychiatry paper rather than a spatial experiment, so the conservative negative label is NON-ST. [accepted_disagree] |
| 10.1186/s43046-025-00312-z | 2025 | Dual-modality treatment using gamma radiation and ZnO nanoparticles: effects on normal and malignant lung cells | AGREE | NOT FOUND | This is a non-spatial cancer radiotherapy/nanoparticle study, not a spatial transcriptomics study. [accepted_agree] |
| 10.3390/a16050254 | 2023 | Well-Separated Pair Decompositions for High-Dimensional Datasets | DISAGREE | Well-Separated Pair Decompositions for High-Dimensional Datasets | This is a general algorithms/mathematics paper on high-dimensional datasets, not a spatial transcriptomics study or a relevant single-cell methods paper. [accepted_disagree] |
| 10.48550/arxiv.2410.21107 | 2024 | Tree-Wasserstein Distance for High Dimensional Data with a Latent Feature Hierarchy | DISAGREE | Tree-Wasserstein Distance for High Dimensional Data with a Latent Feature Hierarchy | This is a general high-dimensional distance-learning paper, not a spatial transcriptomics study or relevant single-cell computational paper. [accepted_disagree] |

#### scPNMF (5)

| DOI | Year | Title | Audit decision | Evidence (verbatim quote) | Audit reason / validation status |
|---|---|---|---|---|---|
| 10.1080/01621459.2024.2342639 | 2024 | Dissecting Gene Expression Heterogeneity: Generalized Pearson Correlation Squares and the <i>K</i> -Lines Clustering Algorithm | DISAGREE | "we develop a K-lines clustering algorithm ..." | This is a statistics methodology paper on Pearson correlation squares and clustering, not a single-cell or spatial transcriptomics tool paper tied to scPNMF. [accepted_disagree] |

| DOI | Year | Title | Audit decision | Evidence (verbatim quote) | Audit reason / validation status |
|---|---|---|---|---|---|
| 10.1093/bioadv/vbag080 | 2026 | PeakPrime: a peak-guided primer design pipeline for target enrichment in 3′-end RNA-seq | DISAGREE | We present PeakPrime ... validate primers by qPCR and targeted 3′ RNA-seq ... While empirically tested in bulk RNA-seq | PeakPrime is presented and validated for targeted bulk 3′ RNA-seq, with only speculative extensions to spatial workflows, so this is not a spatial-transcriptomics study. [accepted_disagree] |
| 10.1172/jci153283 | 2021 | Genomic and transcriptomic profiling reveals distinct molecular subsets associated with outcomes in mantle cell lymphoma | AGREE | Genomic and transcriptomic profiling reveals distinct molecular subsets associated with outcomes in mantle cell lymphoma | This is a mantle cell lymphoma genomic/transcriptomic profiling study, not a spatial transcriptomics study. [accepted_agree] |
| 10.48550/arxiv.2104.10041 | 2021 | Particle swarm optimization in constrained maximum likelihood estimation a case study | DISAGREE | Abstract: "The aim of paper is to apply ... PSO to a constrained maximum likelihood estimation problem in pseudotime analysis ..." | This is a particle swarm optimization paper for pseudotime-related constrained likelihood estimation, not a single-cell marker-selection methods paper tied to scPNMF. [accepted_disagree] |
| 10.48550/arxiv.2210.02672 | 2023 | Orthogonal Non-negative Matrix Factorization: a Maximum-Entropy-Principle Approach | DISAGREE | Abstract describes ONMF methodology; evidence says methods compared include "PNMF". | The packet describes a generic orthogonal non-negative matrix factorization paper with no spatial transcriptomics study context, and the evidence refers to PNMF rather than scPNMF specifically. [accepted_disagree] |

#### NS-Forest v4.0 (3)

| DOI | Year | Title | Audit decision | Evidence (verbatim quote) | Audit reason / validation status |
|---|---|---|---|---|---|
| 10.1016/j.xgen.2025.101103 | 2025 | A high-resolution atlas of the brain predicts lineage and birth order underlying neuronal identity | AGREE | generate the first high-resolution, single-cell transcriptomic atlas of the adult Drosophila melanogaster central brain | This is a Drosophila central-brain single-cell atlas paper, not a spatial transcriptomics study. [accepted_agree] |
| 10.1038/s41588-025-02454-1 | 2026 | Single-cell atlas of the transcriptome and chromatin accessibility in the human retina | AGREE | NOT FOUND | This is a single-cell retina atlas paper, not a spatial transcriptomics study. [accepted_agree] |
| 10.1101/2025.04.10.648034 | 2025 | Benchmarking single cell transcriptome matching methods for incremental growth of cell atlases | DISAGREE | This study benchmarks four computational tools for cell type annotation and matching ... | This is a benchmark of single-cell transcriptome matching and atlas growth, not a spatial transcriptomics or targeted panel-selection study. [accepted_disagree] |

#### RankCorr (3)

| DOI | Year | Title | Audit decision | Evidence (verbatim quote) | Audit reason / validation status |
|---|---|---|---|---|---|
| 10.3389/fcell.2022.944325 | 2022 | Single-cell transcriptomics reveals male germ cells and Sertoli cells developmental patterns in dairy goats | AGREE | Evidence quote: NOT FOUND; abstract describes spermatogenesis and single-cell transcriptomics. | This is a single-cell developmental study in dairy goats and does not present a spatial transcriptomics study. [accepted_agree] |
| 10.3390/cancers15051557 | 2023 | Quantitative Evaluation of Stem-like Markers of Human Glioblastoma Using Single-Cell RNA Sequencing Datasets | DISAGREE | Using single-cell RNA sequencing datasets from 37 GBM patients, we obtained a large pool of 2173 GBM stem-like marker candidates. | This is a biological glioblastoma marker-evaluation study using scRNA-seq datasets, not a spatial transcriptomics study or a computational tool paper about RankCorr. [accepted_disagree] |
| 10.48550/arxiv.2503.00174 | 2025 | Optimal Transfer Learning for Missing Not-at-Random Matrix Completion | DISAGREE | Abstract discusses MNAR matrix completion and minimax estimation rather than transcriptomics tool use. | This is a matrix-completion / transfer-learning methods paper motivated by biology, not a single-cell or spatial transcriptomics paper. [accepted_disagree] |

#### geneBasis (3)

| DOI | Year | Title | Audit decision | Evidence (verbatim quote) | Audit reason / validation status |
|---|---|---|---|---|---|
| 10.1101/2021.10.20.465151 | 2021 | Immune, endothelial and neuronal network map in human lymph node and spleen | DISAGREE | Evidence quote: NOT FOUND; abstract describes CODEX imaging and 3D light sheet microscopy. | The abstract describes CODEX imaging and light-sheet microscopy rather than a spatial transcriptomics study, and the packet gives no geneBasis evidence. [accepted_disagree] |
| 10.1101/2024.10.15.618479 | 2024 | Interactive design and validation of antibody panels using single-cell RNA-seq atlases | DISAGREE | "Cytomarker has two options for initial panel creation ... or running geneBasis ..." | The paper is about antibody panel design and spatial profiling, not spatial transcriptomics; geneBasis is mentioned only as an option for initial antibody panel creation. [accepted_disagree] |
| 10.1186/s13058-023-01703-7 | 2023 | Single-cell RNA reveals a tumorigenic microenvironment in the interface zone of human breast tumors | AGREE | Evidence quote: NOT FOUND; abstract says "Single-cell RNA reveals a tumorigenic microenvironment ..." | This is a single-cell RNA study of the breast tumor interface zone, not a spatial transcriptomics study. [accepted_agree] |

#### SCMER (2)

| DOI | Year | Title | Audit decision | Evidence (verbatim quote) | Audit reason / validation status |
|---|---|---|---|---|---|
| 10.1016/j.neunet.2023.02.018 | 2023 | UDRN: Unified Dimensional Reduction Neural Network for feature selection and feature projection | DISAGREE | "This paper proposes a unified framework named Unified Dimensional Reduction Network (UDRN) ..." | This is a general dimensionality reduction / neural network methods paper rather than a single-cell or spatial transcriptomics tool paper tied to SCMER. [accepted_disagree] |
| 10.1093/narcan/zcac038 | 2022 | Single-cell CRISPR immune screens reveal immunological roles of tumor intrinsic factors | AGREE | Evidence quote: NOT FOUND; abstract describes single-cell CRISPR immune screens. | This is a single-cell CRISPR immune-screen study, not a spatial transcriptomics study. [accepted_agree] |

#### Spapros (2)

| DOI | Year | Title | Audit decision | Evidence (verbatim quote) | Audit reason / validation status |
|---|---|---|---|---|---|
| 10.1002/fft2.70022 | 2025 | Highly Targeted Metabolomics Coupled With Gene Expression Analysis by RT–qPCR Improves Beef Separation Based on Grass, Grain, or Grape Supplemented Diet | AGREE | Highly Targeted Metabolomics Coupled With Gene Expression Analysis by RT–qPCR Improves Beef Separation | This is a targeted metabolomics and RT-qPCR food-science paper, not a spatial transcriptomics study. [accepted_agree] |
| 10.1093/bioadv/vbag080 | 2026 | PeakPrime: a peak-guided primer design pipeline for target enrichment in 3′-end RNA-seq | DISAGREE | We present PeakPrime ... validate primers by qPCR and targeted 3′ RNA-seq ... While empirically tested in bulk RNA-seq | PeakPrime is a targeted bulk 3′ RNA-seq primer-design pipeline with only possible future spatial extensions, not a spatial-transcriptomics study. [accepted_disagree] |

#### SMaSH (1)

| DOI | Year | Title | Audit decision | Evidence (verbatim quote) | Audit reason / validation status |
|---|---|---|---|---|---|
| 10.1038/s41540-024-00441-6 | 2024 | Computational identification of surface markers for isolating distinct subpopulations from heterogeneous cancer cell populations | AGREE | Computational identification of surface markers for isolating distinct subpopulations from heterogeneous cancer cell populations | This is a non-spatial single-cell surface-marker identification paper for cancer subpopulations. [accepted_agree] |

#### gpsFISH (1)

| DOI | Year | Title | Audit decision | Evidence (verbatim quote) | Audit reason / validation status |
|---|---|---|---|---|---|
| 10.3390/ijms252312606 | 2024 | Combining Network Pharmacology and Transcriptomic Strategies to Explore the Pharmacological | AGREE | Combining Network Pharmacology and Transcriptomic Strategies to | This is a pharmacology/depression-mechanism paper, not a spatial |

| DOI | Year | Title | Audit decision | Evidence (verbatim quote) | Audit reason / validation status |
|---|---|---|---|---|---|
| | | Mechanism of Total Ginsenoside Ginseng Root and Its Impact on Antidepressant Effects | | Explore the Pharmacological Mechanism | transcriptomics study. [accepted_agree] |

### OTHER (n = 30)

*Residual category: corrections, non-research items, or otherwise ineligible records.*

#### NS-Forest v2 (6)

| DOI | Year | Title | Audit decision | Evidence (verbatim quote) | Audit reason / validation status |
|---|---|---|---|---|---|
| 10.1007/978-1-0716-4184-2_5 | 2024 | Homebuilt Imaging-Based Spatial Transcriptomics: Tertiary Lymphoid Structures as a Case Example | AGREE | Abstract describes a comprehensive DIY guide; evidence references a comparison figure within the guide. | This is a book chapter / protocol-style guide rather than a primary experimental or computational paper, so OTHER is appropriate. [accepted_agree] |
| 10.1038/s41592-024-02563-5 | 2025 | Human BioMolecular Atlas Program (HuBMAP): 3D Human Reference Atlas construction and usage | AGREE | Resource | This is a resource/infrastructure paper describing HuBMAP atlas construction and usage, which fits OTHER. [accepted_agree] |
| 10.1101/2024.07.03.601914 | 2024 | A DIY guide for image-based spatial transcriptomic: TLS as a case example | AGREE | A DIY guide for image-based spatial transcriptomic: TLS as a case example; Comparisons of three different marker gene selection methods ... | This is a DIY guide/protocol chapter rather than a primary experimental or benchmarking paper about NS-Forest v2. [accepted_agree] |
| 10.1371/journal.pbio.3002133 | 2023 | A guide to the BRAIN Initiative Cell Census Network data ecosystem | AGREE | Here, we provide a guide to the cellular and spatial approaches employed by the BICCN ... | This is a guide/resource article for the BICCN data ecosystem, which fits OTHER. [accepted_agree] |
| 10.48550/arxiv.2112.13289 | 2021 | Prevalence Threshold and bounds in the Accuracy of Binary Classification Systems | DISAGREE | Abstract defines accuracy of binary classification systems and discusses prevalence thresholds and accuracy metrics. | This is a general methodology/statistics manuscript on binary classification accuracy metrics, not a single-cell or spatial transcriptomics paper. [accepted_disagree] |
| 10.48550/arxiv.2204.09673 | 2022 | Guidelines for reporting cell types: the MIRACL standard | AGREE | Here we introduce the Minimal Information Reporting About a CelL (MIRACL) standard | This is a reporting standard/guideline paper, which fits OTHER rather than REV or COMP. [accepted_agree] |

#### COSG (5)

| DOI | Year | Title | Audit decision | Evidence (verbatim quote) | Audit reason / validation status |
|---|---|---|---|---|---|
| 10.1016/j.jmb.2026.169781 | 2026 | IMDD: A Database for Exploring Tissue-Specific Gene Expression Dynamics During Holometabolous Insects | AGREE | we have developed the Insect Metamorphic Development Database (IMDD), an interactive platform | This is a tissue-specific gene-expression database/resource paper, which fits OTHER. [accepted_agree] |
| 10.1016/j.molp.2025.12.026 | 2025 | PlantscRNAdb 4.0: Improved marker identification and annotation under a cell-type uniformity for plants | DISAGREE | we present PlantscRNAdb release 4.0, an updated comprehensive database ... | This PlantscRNAdb article is primarily a database/resource paper with associated tools, so it fits OTHER rather than COMP. [accepted_disagree] |
| 10.1038/s41597-026-06707-4 | 2026 | SexTumorDB: a comprehensive resource of sex-dependent tumor landscape at single-cell resolution | AGREE | we developed the SexTumorDB database, an integrated resource containing RNA sequencing profiles | This is a database/resource paper, which fits OTHER. [accepted_agree] |
| 10.1093/nar/gkac922 | 2022 | ImmCluster: an ensemble resource for immunology cell type clustering and annotations in normal and cancerous tissues | AGREE | we introduce the ImmCluster database | This is an immune-cell clustering and annotation database/resource paper, which fits OTHER. [accepted_agree] |
| 10.1093/nar/gkad820 | 2023 | SORC: an integrated spatial omics resource in cancer | AGREE | we have developed an integrated spatial omics resource in cancer (SORC) | This is a spatial-omics database/resource paper, which fits OTHER rather than a computational comparison or tool-use paper. [accepted_agree] |

#### SCMER (5)

| DOI | Year | Title | Audit decision | Evidence (verbatim quote) | Audit reason / validation status |
|---|---|---|---|---|---|
| 10.1007/978-1-0716-4184-2_5 | 2024 | Homebuilt Imaging-Based Spatial Transcriptomics: Tertiary Lymphoid Structures as a Case Example | AGREE | Here, we present a comprehensive do-it-yourself (DIY) guide to perform such experiments at reduced costs | This is a DIY/protocol chapter for imaging-based spatial transcriptomics rather than a primary benchmark or experimental paper about SCMER. [accepted_agree] |
| 10.1007/s00500-023-08435-y | 2023 | RETRACTED ARTICLE: Ensemble recurrent neural network with whale optimization algorithm-based DNA sequence classification for medical applications | AGREE | Evidence quote and title begin with "RETRACTED ARTICLE". | The packet is explicitly a retracted article, which is appropriately treated as OTHER. [accepted_agree] |
| 10.1101/2024.07.03.601914 | 2024 | A DIY guide for image-based spatial transcriptomic: TLS as a | AGREE | we present a comprehensive do-it-yourself guide to perform such | This is a DIY guide/protocol paper rather than a primary experimental |

| DOI | Year | Title | Audit decision | Evidence (verbatim quote) | Audit reason / validation status |
|---|---|---|---|---|---|
| | | case example | | experiments at reduced costs leveraging open-source approaches | or computational benchmark study about SCMER. [accepted_agree] |
| 10.1101/2024.11.30.626148 | 2024 | Unbiased learning of protein conformational representation via unsupervised random forest | DISAGREE | here, we present unsupervised random forest (URF) ... that identifies functionally critical features of biomolecules | This is a protein-biophysics representation-learning paper, which is out of scope for single-cell/spatial transcriptomics tool-paper screening. [accepted_disagree] |
| 10.18280/ria.360505 | 2022 | Diffusion Convolutional Recurrent Neural Network-Based Load Forecasting During COVID-19 Pandemic Situation | DISAGREE | we proffer Diffusion Convolutional Recurrent Neural Network (DCRNN) ... for load forecasting | This paper is an energy-load forecasting study during COVID-19 and is entirely out of scope for single-cell/spatial transcriptomics tool relations. [accepted_disagree] |

#### scGeneFit (4)

| DOI | Year | Title | Audit decision | Evidence (verbatim quote) | Audit reason / validation status |
|---|---|---|---|---|---|
| 10.1093/nar/gkad895 | 2023 | SCAN: Spatiotemporal Cloud Atlas for Neural cells | DISAGREE | "developed a comprehensive ... database (SCAN) ... providing a convenient and efficient data mining framework." | This is a database/resource paper for curated nervous system single-cell and spatial data, so OTHER is more precise than COMP. [accepted_disagree] |
| 10.1093/nar/gkaf1092 | 2025 | scBrainScope: cross-species multidimensional brain atlas | AGREE | we developed scBrainScope, a detailed and interactive transcriptomic atlas of the brain | This is a brain-atlas resource/database paper, which fits OTHER rather than a computational tool-use category. [accepted_agree] |
| 10.3150/21-bej1433 | 2022 | Estimation of Wasserstein distances in the Spiked Transport Model | DISAGREE | "We propose a new statistical model, the spiked transport model ..." | This is a statistics methodology paper on Wasserstein distance estimation, not a single-cell or spatial transcriptomics paper. [accepted_disagree] |
| 10.48550/arxiv.2005.03847 | 202 | Tree! I am no Tree! I am a Low Dimensional Hyperbolic Embedding | DISAGREE | we present a novel fast algorithm TREEREP ... given a hyperbolic metric ... | This is a general machine-learning/mathematical embedding paper, not a single-cell or spatial transcriptomics methods paper. [accepted_disagree] |

#### PERSIST (3)

| DOI | Year | Title | Audit decision | Evidence (verbatim quote) | Audit reason / validation status |
|---|---|---|---|---|---|
| 10.1007/978-1-0716-4184-2_5 | 2024 | Homebuilt Imaging-Based Spatial Transcriptomics: Tertiary Lymphoid Structures as a Case Example | AGREE | "Here, we present a comprehensive do-it-yourself (DIY) guide ..." | This is a DIY guide / protocol chapter for homebuilt imaging-based spatial transcriptomics, so OTHER is appropriate. [accepted_agree] |
| 10.1021/acschemneuro.4c00069 | 2024 | Effect of Different Staining Methods on Brain Cryosections | DISAGREE | In this study, four different staining methods were analyzed for their effect on RNA integrity in frozen sections of brain tissue. | This is a staining/sample-preparation methods paper for brain cryosections in a spatial-transcriptomics context, which fits OTHER rather than an experimental citation-screening label. [accepted_disagree] |
| 10.1101/2024.07.03.601914 | 2024 | A DIY guide for image-based spatial transcriptomic: TLS as a case example | AGREE | we present a comprehensive do-it-yourself guide to perform such experiments at reduced costs leveraging open-source approaches | This is a DIY guide/protocol paper rather than a primary experimental or computational benchmark study about PERSIST. [accepted_agree] |

#### SMaSH (2)

| DOI | Year | Title | Audit decision | Evidence (verbatim quote) | Audit reason / validation status |
|---|---|---|---|---|---|
| 10.1007/978-1-0716-4184-2_5 | 2024 | Homebuilt Imaging-Based Spatial Transcriptomics: Tertiary Lymphoid Structures as a Case Example | AGREE | Here, we present a comprehensive do-it-yourself (DIY) guide to perform such experiments at reduced costs | This is a DIY/protocol chapter for imaging-based spatial transcriptomics rather than a primary benchmark or experimental paper about SMaSH. [accepted_agree] |
| 10.1101/2024.07.03.601914 | 2024 | A DIY guide for image-based spatial transcriptomic: TLS as a case example | AGREE | we present a comprehensive do-it-yourself guide to perform such experiments at reduced costs leveraging open-source approaches | This is a DIY guide/protocol paper rather than a primary experimental or computational benchmark study about SMaSH. [accepted_agree] |

#### geneBasis (2)

| DOI | Year | Title | Audit decision | Evidence (verbatim quote) | Audit reason / validation status |
|---|---|---|---|---|---|
| 10.1007/978-1-0716-4184-2_5 | 2024 | Homebuilt Imaging-Based Spatial Transcriptomics: Tertiary Lymphoid Structures as a Case Example | AGREE | "Here, we present a comprehensive do-it-yourself (DIY) guide ..." | This is a DIY guide / protocol chapter for imaging-based spatial transcriptomics, so OTHER is appropriate. [accepted_agree] |

| DOI | Year | Title | Audit decision | Evidence (verbatim quote) | Audit reason / validation status |
|---|---|---|---|---|---|
| 10.1101/2024.07.03.601914 | 2024 | A DIY guide for image-based spatial transcriptomic: TLS as a case example | AGREE | we present a comprehensive do-it-yourself guide to perform such experiments at reduced costs leveraging open-source approaches | This is a DIY guide/protocol paper rather than a primary experimental or computational benchmark study about geneBasis. [accepted_agree] |

#### gpsFISH (2)

| DOI | Year | Title | Audit decision | Evidence (verbatim quote) | Audit reason / validation status |
|---|---|---|---|---|---|
| 10.1007/978-1-0716-4184-2_5 | 2024 | Homebuilt Imaging-Based Spatial Transcriptomics: Tertiary Lymphoid Structures as a Case Example | AGREE | Here, we present a comprehensive do-it-yourself (DIY) guide to perform such experiments | This is a methods/protocol-style DIY guide chapter rather than a primary experimental or computational benchmark paper. [accepted_agree] |
| 10.1101/2024.07.03.601914 | 2024 | A DIY guide for image-based spatial transcriptomic: TLS as a case example | AGREE | we present a comprehensive do-it-yourself guide to perform such experiments at reduced costs leveraging open-source approaches | This is a DIY guide/protocol paper rather than a primary experimental or computational benchmark study about gpsFISH. [accepted_agree] |

#### RankCorr (1)

| DOI | Year | Title | Audit decision | Evidence (verbatim quote) | Audit reason / validation status |
|---|---|---|---|---|---|
| 10.21203/rs.3.rs-914457/v1 | 2021 | Q-Mer Analysis: A Generalized Method for Analyzing RNA-Seq Data | DISAGREE | In this study, we developed a method to summarize RNA-Seq data without losing alignment information. | This is a general bulk RNA-seq methodology paper rather than a single-cell/spatial computational paper relevant to the tool-screening task, so OTHER is more appropriate than COMP. [accepted_disagree] |
