## supplementary_evidence_dossier_panel for "Lack of adoption of computational gene-panel-selection tools in targeted spatial transcriptomics"

### Panel-design classification evidence dossier

*Per-record audit dossier for the 1,646 panel-design audit rows in the current deduplicated corpus. For each record this document shows the final classification (TOOL / HYBRID-TOOL / VENDOR-MANUAL / VENDOR / MANUAL-PANEL / FILTERED), the GPT-5.4 audit decision (AGREE / DISAGREE / NEEDS_MANUAL), the verbatim PDF evidence quote used by the audit, the validation status, and a Fig. 2-scope flag indicating whether the record was excluded from the Fig. 2 analytic set by the temporary platform/scope exclusion list. Records that fall outside the current corpus after post-freeze manual_exclusions are not included (those are legacy audit rows from earlier corpus iterations, not used in any analysis).*

### Summary

**Audit rows in the live corpus: 1646 (of 1664 total in GPT_audit_FINAL.csv; 18 legacy rows outside the current corpus are excluded). Fig. 2 analytic set after temporary platform/scope exclusions: 1636 (10 records currently excluded).**

**Breakdown by classification (in current corpus):**

TOOL: 7 — Final panel was selected by one of the 18 canonical computational gene-panel-selection tools.

HYBRID-TOOL: 27 — Final panel combines a computational tool with manual, vendor, or other additions.

VENDOR-MANUAL: 50 — Vendor panel with manual additions, substitutions, or adaptations.

VENDOR: 99 — Fixed commercial or catalog panel.

MANUAL-PANEL: 304 — Panel curated from biological markers, differential expression, ranking, pathway lists, or literature — no canonical tool drove the final selection.

FILTERED: 1159 — No adjudicable targeted panel-design evidence in the available text (e.g. non-targeted spatial, computational paper without a new panel, supplement-only PDF).

### TOOL (n = 7)

*Final panel was selected by one of the 18 canonical computational gene-panel-selection tools.*

| DOI | Year | Title | Audit decision | Evidence (verbatim quote) | Validation status / scope flag |
|---|---|---|---|---|---|
| 10.1038/s41467-025-65704-0 | 2025 | Spatial single-cell atlas reveals regional variations in healthy and diseased human lung | AGREE | q1: panel_design=TOOL; tool_names=Spapros; uses_manual_additions=NO; uses_vendor_panel=NO. | accepted_agree \| Platform: HybISS;SCRINSHOT \| Tool(s): Spapros |
| 10.1038/s41588-024-01990-6 | 2024 | A temporal cortex cell atlas highlights gene expression dynamics during human brain maturation | AGREE | The packet says a subset of minimal markers was generated with NS-Forest v2 and used in the targeted MERFISH panel. | accepted_agree \| Platform: MERSCOPE \| Tool(s): NS-Forest v2 |
| 10.1101/2024.09.17.613303 | 2024 | Endothelial regulatory circuits control cranial neural crest migration and plasticity | AGREE | We used SPAPROS, a probe selection algorithm, on multiome-snRNA-seq, Smart-seq3, and organogenesis datasets. | accepted_agree \| Platform: MERSCOPE \| Tool(s): Spapros |

| DOI | Year | Title | Audit decision | Evidence (verbatim quote) | Validation status / scope flag |
|---|---|---|---|---|---|
| 10.1101/2024.10.08.617260 | 2024 | Spatial Single-Cell Mapping of Transcriptional Differences Across Genetic Backgrounds in Mouse Brains | AGREE | Encoding matrix was fit using a modified version of DPNMF/scPNMF to optimize the gene set. | accepted_agree \| Platform: MERFISH \| Tool(s): scPNMF |
| 10.1101/2025.10.29.684766 | 2025 | A dual genetic constraint underlies the conservation of early brains in vertebrates | AGREE | The packet says the brain bauplan signature was identified by running spapros on the integrated dataset. | accepted_agree \| Platform: in situ sequencing;Molecular Cartography \| Tool(s): Spapros |
| 10.3389/fncel.2023.1256619 | 2023 | Cochlear transcriptome analysis of an outbred mouse population (CFW). | AGREE | The packet states that NSForest marker genes were utilized as MERFISH probes and that NS-Forest was used to identify the minimum number of necessary and sufficient markers for each cluster. | accepted_agree \| Platform: MERSCOPE \| Tool(s): NS-Forest v2 |
| 10.64898/2026.02.12.705355 | 2026 | Multi-gene biomarkers reveal spatial organization and subpopulation-specific damage response in intrahepatic biliary epithelial cells | AGREE | q1: panel_design=TOOL; tool_names=NSForest; uses_manual_additions=NO; uses_vendor_panel=NO. | accepted_agree \| Platform: HCR-FISH \| Tool(s): NS-Forest v2 |

### HYBRID-TOOL (n = 27)

*Final panel combines a computational tool with manual, vendor, or other additions.*

| DOI | Year | Title | Audit decision | Evidence (verbatim quote) | Validation status / scope flag |
|---|---|---|---|---|---|
| 10.1016/j.cell.2024.02.030 | 2024 | Cellular architecture of evolving neuroinflammatory lesions and multiple sclerosis pathology | AGREE | The selection was made manually, based on the literature and single-cell datasets, in combination with different computational frameworks; for human samples, they employed the Xenium Human Brain Gene Expression panel with 266 genes. | accepted_agree \| Platform: Xenium \| Tool(s): NS-Forest v2 |
| 10.1016/j.immuni.2025.10.003 | 2025 | Single-cell spatial transcriptomic profiling defines a pathogenic inflammatory niche in chronic active multiple sclerosis lesions. | AGREE | Initially, the top 150 genes identified using Spapros were selected for identification of the major cell populations. Subsequently, an additional 250 genes ... were included based on previous literature. Furthermore, 100 additional genes were selected based on prior biological knowledge. | accepted_agree \| Platform: MERSCOPE \| Tool(s): Spapros |

| DOI | Year | Title | Audit decision | Evidence (verbatim quote) | Validation status / scope flag |
|---|---|---|---|---|---|
| 10.1016/j.immuni.2026.03.003 | 2026 | Distinct tissue niches contribute to prostate tissue-resident memory CD8+ T cell differentiation and heterogeneity | AGREE | q1: panel_design=HYBRID-TOOL; tool_names=PERSIST; uses_manual_additions=YES. | accepted_agree \| Platform: Xenium \| Tool(s): PERSIST |
| 10.1038/s41467-023-40674-3 | 2023 | Spatial organization of the mouse retina at single cell resolution by MERFISH. | AGREE | we used SCMER (single-cell manifold-preserving feature selection) tool to choose additional genes ... which brought up the total number in the second panel to 500 genes | accepted_agree \| Platform: MERFISH \| Tool(s): SCMER |
| 10.1038/s41467-024-46437-y | 2024 | Mapping human tissues with highly multiplexed RNA in situ hybridization | AGREE | Human kidney gene selection was performed by gpsFISH; a list of genes was selected by differential expression and curated marker genes were added manually. | accepted_agree \| Platform: Molecular Cartography \| Tool(s): gpsFISH |
| 10.1038/s41586-024-07171-z | 2024 | Spatially organized cellular communities form the developing human heart | AGREE | A list of 238 cell-subpopulation-specific genes was identified after applying NS-Forest2; 63 MERFISH target genes were manually selected from DGE and NS-Forest2 lists. | accepted_agree \| Platform: MERFISH \| Tool(s): NS-Forest v2 |
| 10.1038/s41586-024-08030-7 | 2024 | Temporally distinct 3D multi-omic dynamics in the developing human brain | AGREE | The packet says MERFISH gene selection used NSForest v2 and then added literature markers and genes with differential methylation. | accepted_agree \| Platform: MERFISH \| Tool(s): NS-Forest v2 |
| 10.1038/s41586-024-08466-x | 2025 | Tissue-resident memory CD8 T cell diversity is spatiotemporally imprinted | AGREE | citation prior lists PERSIST; q1: panel design made use of PERSIST plus manual additions and a vendor base set. | accepted_agree \| Platform: Xenium;MERSCOPE \| Tool(s): PERSIST |
| 10.1038/s41586-025-09921-z | 2026 | Neuro-epithelial circuits promote sensory convergence and intestinal immunity | AGREE | The packet states that PERSIST was run to produce the final list of 480 genes and that manually selected genes were added to the existing list. | accepted_agree \| Platform: Xenium \| Tool(s): PERSIST |
| 10.1038/s41586-025-09996-8 | 2026 | Pyramidal neurons proportionately alter cortical interneuron subtypes | AGREE | Top marker genes identified by COSG were combined with literature markers, and additional genes were added and tested on scRNA-seq. | accepted_agree \| Platform: MERFISH \| Tool(s): COSG |
| 10.1038/s41587-023-01747-2 | 2023 | Multimodal spatiotemporal phenotyping of human retinal organoid development | AGREE | The 100-gene panel was selected using geneBasisR, given the 4i gene panel and the top five marker genes of major cell types as the pre-selected genes. | accepted_agree \| Platform: Molecular Cartography \| Tool(s): geneBasis |

| DOI | Year | Title | Audit decision | Evidence (verbatim quote) | Validation status / scope flag |
|---|---|---|---|---|---|
| 10.1038/s42003-021-02517-z | 2021 | Comprehensive in situ mapping of human cortical transcriptomic cell types. | AGREE | The packet says the genes were manually and computationally curated and that NS-Forest was one of the main computational algorithms used. | accepted_agree \| Platform: in situ sequencing \| Tool(s): NS-Forest v2 |
| 10.1084/jem.20240776 | 2025 | The XCL1-XCR1 axis supports intestinal tissue residency and antitumor immunity. | AGREE | We created a base set of canonical immune marker genes, ligand-receptor pairs, spatially differentially expressed genes within mouse P14 T cells, and the 10X Genomics base human colon panel totaling 343 genes. Using this set as prior information to PERSIST, and a reference human immune cell scRNA-seq dataset, unsupervised PERSIST filled in the remaining 79 genes. | accepted_agree \| Platform: Xenium \| Tool(s): PERSIST |
| 10.1101/2023.06.29.547074 | 2023 | Single cell-resolution<i>in situ</i>sequencing elucidates spatial dynamics of multiple sclerosis lesion and disease evolution | AGREE | This selection was made manually in combination with different computational frameworks; Ref. 116 is NS-Forest. We employed the Xenium Human Brain Gene Expression panel with 266 genes. | accepted_agree \| Platform: Xenium \| Tool(s): NS-Forest v2 |
| 10.1101/2024.03.20.585130 | 2024 | Functional Diversity of Memory CD8 T Cells is Spatiotemporally Imprinted | AGREE | PERSIST was run to add genes, and the panel was built from a 10x Genomics base human colon panel plus immune markers, ligand-receptor pairs, and Xist. | accepted_agree \| Platform: Xenium;MERSCOPE \| Tool(s): PERSIST |
| 10.1101/2025.03.20.644368 | 2025 | Multi-omic human neural organoid cell atlas of the posterior brain | AGREE | The packet says the 500-gene MERSCOPE panel combined canonical markers with genes selected by geneBasis. | accepted_agree \| Platform: MERSCOPE \| Tool(s): geneBasis |
| 10.1101/2025.05.13.653495 | 2025 | A spatiotemporal cancer cell trajectory underlies glioblastoma heterogeneity | DISAGREE | The Xenium human brain panel was supplemented with additional probes, and the custom probes were largely selected using Spapros plus manual GB markers; Spapros is a canonical panel-selection tool. | corrected_high_risk_disagreement_canonical_tool \| Platform: Xenium \| Tool(s): Spapros |
| 10.1101/2025.06.13.659535 | 2025 | Non-disruptive 3D profiling of combinations of epigenetic marks in single cells | AGREE | gemma_q1_panel_design says NS-Forest was used to aid gene selection and the final set included literature-derived and methylation-derived additions. | accepted_agree \| Platform: Other / unspecified \| Tool(s): NS-Forest v2 |

| DOI | Year | Title | Audit decision | Evidence (verbatim quote) | Validation status / scope flag |
|---|---|---|---|---|---|
| 10.1101/2025.08.07.667992 | 2025 | Single-cell multiomic human brain atlas reveals regulatory drivers of cortical regionality | DISAGREE | A DART-FISH gene panel was selected with gpsFISH and then supplemented with curated marker genes; gpsFISH is a canonical panel-selection tool. | corrected_high_risk_disagreement_canonical_tool \| Platform: Other / unspecified \| Tool(s): gpsFISH |
| 10.1101/2025.08.19.671030 | 2025 | OpenFISH enables integrated high-resolution spatial transcriptomics and metabolomics on a single tissue section | AGREE | q1: panel_design=HYBRID-TOOL; tool_names=scGIST; uses_manual_additions=YES. | accepted_agree \| Platform: in situ sequencing \| Tool(s): scGIST |
| 10.1101/2025.08.29.673064 | 2025 | Spatial Analysis of T Cell Clonality in Autoimmune Kidney Disease Using TRV Probes | AGREE | Spapros was used to select 150 genes for the 10x Xenium panel; combined with 132 TRV variable genes designed with 10x Genomics and additional genes from a multi-tissue Xenium panel (Gemma4 packet quotes: "150 genes were selected using the Spapros algorithm ... with a single-cell kidney atlas and ANCA-GN T cell atlas." and "132 TRV variable genes with three probes each were designed in cooperation with 10x Genomics."). Revert 2026-05-21: previous evidence_used was cross-contaminated from 10.64898/2025.12.19.695425 and has been replaced; classification HYBRID-TOOL (tool: Spapros) is unchanged. | accepted_agree \| Platform: Xenium \| Tool(s): Spapros |
| 10.1101/2025.11.22.688128 | 2025 | A cross-species spatial transcriptomic atlas of the human and non-human primate basal ganglia | AGREE | The packet says basal ganglia marker genes were chosen by hand and used as a starter list for geneBasis. | accepted_agree \| Platform: Xenium;MERSCOPE \| Tool(s): geneBasis |
| 10.1101/2025.11.22.689869 | 2025 | Spatial patterning of transcriptional and regulatory programs in the primate subcortex | AGREE | Panel of 300 genes was constructed from computationally-derived and manually selected marker genes; panel search used mFISHtools and GeneBasis. | accepted_agree \| Platform: Xenium \| Tool(s): geneBasis |
| 10.64898/2025.12.19.695425 | 2025 | Exploiting pair correlation function to describe biological tissue structure | DISAGREE | The immune-oncology Xenium panel was extended with 100 additional genes, and scGeneFit was used before manual marker-list optimization; final classification restored to HYBRID-TOOL. | user_final_override_hybrid_tool \| Platform: Xenium \| Tool(s): scGeneFit |
| 10.64898/2026.01.10.694705 | 2026 | The Caudate Nucleus Exhibits Distinct Pathology and Cell Type- | AGREE | q1: panel_design=HYBRID-TOOL; tool_names=NSForest; | accepted_agree \| Platform: Xenium \| Tool(s): NS-Forest v2 |

| DOI | Year | Title | Audit decision | Evidence (verbatim quote) | Validation status / scope flag |
|---|---|---|---|---|---|
| | | Specific Responses Across Alzheimer’s Disease | | uses_manual_additions=YES. | |
| 10.64898/2026.02.03.703645 | 2026 | Single-cell Multiome Analysis of Chromatin State and Transcriptome in the Human Basal Ganglia | AGREE | Two initial subsets of genes were selected based on the snRNA-seq DEGs using NS Forest124 and snm3C-seq DMGs using Spapros separately | accepted_agree \| Platform: MERSCOPE \| Tool(s): NS-Forest v2;Spapros |
| 10.64898/2026.02.12.705594 | 2026 | A Multimodal Single-Cell Epigenomic and 3D Genome Atlas of the Human Basal Ganglia | AGREE | To design the gene panel, we used Spapros147 to prioritize genes differentially expressed across cell types | accepted_agree \| Platform: MERSCOPE \| Tool(s): Spapros |

### VENDOR-MANUAL (n = 50)

*Vendor panel with manual additions, substitutions, or adaptations.*

| DOI | Year | Title | Audit decision | Evidence (verbatim quote) | Validation status / scope flag |
|---|---|---|---|---|---|
| 10.1002/jmv.29009 | 2023 | Molecular and immune signatures, and pathological trajectories of fatal COVID-19 lungs defined by in situ spatial single-cell transcriptome analysis. | DISAGREE | paper_type=EXP, panel_scope=TARGETED, spatial=YES; chimeric padlock probes used 1 custom panel plus four predesigned panels (hImmune I1A, hImmune I3D, hLung L1C, hLung L2E, CARTANA AB). | validated_high_risk_disagreement \| Platform: CARTANA |
| 10.1016/j.ccell.2025.03.017 | 2025 | Spatial determinants of antibody-drug conjugate SHR-A1811 efficacy in neoadjuvant treatment for HER2-positive breast cancer | DISAGREE | q1: uses_vendor_panel=YES; uses_manual_additions=YES; panel_design=HYBRID-MANUAL. | validated_high_risk_disagreement \| Platform: Xenium |
| 10.1016/j.ccell.2025.10.004 | 2026 | Multimodal spatial-omics reveal co-evolution of alveolar progenitors and proinflammatory niches in progression of lung precursor lesions | DISAGREE | paper_type=EXP, panel_scope=TARGETED, spatial=YES; Xenium was used with a predesigned 5K Prime gene panel plus a custom-designed 100-gene add-on. | validated_high_risk_disagreement \| Platform: Xenium |
| 10.1016/j.celrep.2025.116114 | 2025 | A basophil-fibroblast pro-inflammatory axis fuels type 2 skin inflammation. | DISAGREE | The gene panel used for MERFISH was curated through the Vizgen Portal using mouse skin as a total FPKM reference. | validated_high_risk_disagreement \| Platform: MERSCOPE |
| 10.1016/j.jsbmb.2026.106993 | 2026 | Deletion of vitamin D receptor with calcium sensing receptor in keratinocytes promotes epidermal tumorigenesis by limiting dna | DISAGREE | As none of pre-designed panel accommodates our studies, we designed custom made probes (100 genes) using Xenium probe designer software including | validated_high_risk_disagreement \| Platform: Xenium |

| DOI | Year | Title | Audit decision | Evidence (verbatim quote) | Validation status / scope flag |
|---|---|---|---|---|---|
| | | repair and oxidative stress response genes. | | markers for epidermal and hair stem cells and their differentiation and proliferation markers. | |
| 10.1016/j.xgen.2025.101133 | 2026 | Cellular senescence in human liver under normal aging and cancer. | DISAGREE | q1 says the samples used the pre-set Xenium Human Multi-tissue and Cancer Panel plus 100 additional custom genes. | validated_high_risk_disagreement \| Platform: Xenium |
| 10.1016/j.xjidi.2025.100401 | 2025 | Immunoregulatory Microenvironment in the Elderly Skin. | DISAGREE | q1 says the probe hybridization mix used the 10x Genomics Xenium Human Skin Gene Expression Panel with an additional customized panel. | validated_high_risk_disagreement \| Platform: Xenium |
| 10.1038/s41380-025-03331-3 | 2026 | Single-cell resolution spatial transcriptomic signature of the retrosplenial cortex during memory consolidation. | DISAGREE | 297 DNA probes were pre-designed by 10x Genomics- 247 included from their mouse brain gene expression panel, and 50 custom probes. | validated_high_risk_disagreement \| Platform: Xenium |
| 10.1038/s41416-025-03088-0 | 2025 | Single-cell spatial analysis with Xenium reveals anti-tumour responses of CXCL13 + T and CXCL9+ cells after radiotherapy combined with anti-PD-L1 therapy. | DISAGREE | The Lung panel was kindly provided by Ayako S. The Immune panel was created using custom-ordered components. | validated_high_risk_disagreement \| Platform: Xenium |
| 10.1038/s41467-023-40156-6 | 2023 | Macrophage and neutrophil heterogeneity at single-cell spatial resolution in human inflammatory bowel disease. | DISAGREE | CosMx™ Human Universal Cell Characterization RNA Panel (1000-plex) + 20 custom genes | validated_high_risk_disagreement \| Platform: CosMx |
| 10.1038/s41467-024-45752-8 | 2024 | High resolution spatial profiling of kidney injury and repair using RNA hybridization-based in situ sequencing. | DISAGREE | We selected 50 custom kidney gene probes and added 150 pre-designed probes for immune and lung cell types. | validated_high_risk_disagreement \| Platform: CARTANA;HybISS |
| 10.1038/s41586-025-09602-x | 2025 | A parabrachial hub for need-state control of enduring pain | DISAGREE | q1: hybridization mix combined 1,000-plex mouse neuroscience RNA probe mix, custom probe mix, rRNA segmentation markers, RNA inhibitor, and Bruker buffer; uses_vendor_panel=YES. | validated_high_risk_disagreement \| Platform: CosMx |
| 10.1038/s41588-025-02080-x | 2025 | Spatial transcriptomics identifies molecular niche dysregulation associated with distal lung remodeling in pulmonary fibrosis | DISAGREE | 246 genes came from an early version of the Xenium human lung base panel and 97 genes came from a custom-designed panel curated from human lung single-cell data. | validated_high_risk_disagreement \| Platform: Xenium |
| 10.1038/s43587-024-00755-4 | 2025 | Clearance of p21 highly expressing senescent cells | DISAGREE | The Xenium Mouse Tissue Atlassing panel was supplemented | validated_high_risk_disagreement \| Platform: Xenium |

| DOI | Year | Title | Audit decision | Evidence (verbatim quote) | Validation status / scope flag |
|---|---|---|---|---|---|
| | | accelerates cutaneous wound healing. | | with a custom gene panel encompassing approximately 500 genes. | |
| 10.1038/s44321-025-00280-w | 2025 | Spatial transcriptomics elucidates localized immune responses in atherosclerotic coronary artery. | DISAGREE | The CosMx universal cell characterisation RNA panel contained a core of 950 probes and an additional set of 20 custom probes. | validated_high_risk_disagreement \| Platform: CosMx |
| 10.1073/pnas.2515006122 | 2025 | Spatial gene expression analysis reveals pathological niches in Japanese encephalitis virus neuroinvasion | DISAGREE | The analysis used a predesigned Mouse Brain Gene base panel (248 genes) for cell-type annotation, as well as 100 custom probes targeting three JEV genome regions, immune cell markers, and immune response genes. | validated_high_risk_disagreement \| Platform: Xenium |
| 10.1080/07366205.2025.2591484 | 2025 | Streamlining single-cell spatial transcriptomics for human kidney tissue. | DISAGREE | q1: 1000-plex CosMx TAP RNA panel with the addition of kidney-specific custom genes; q2: TARGETED spatial COMP paper. | validated_high_risk_disagreement \| Platform: CosMx |
| 10.1101/2022.10.06.510405 | 2022 | High resolution mapping of the breast cancer tumor microenvironment using integrated single cell, spatial and in situ analysis of FFPE tissue | DISAGREE | We used 280 genes from the Xenium Human Breast Panel with 33 add-on genes for a total of 313 genes ... selected and curated primarily based on single cell atlas data... | validated_high_risk_disagreement \| Platform: Xenium |
| 10.1101/2023.10.18.562892 | 2023 | <i>Ataxia-telangiectasia mutated</i>(<i>Atm</i>) disruption sensitizes spatially-directed H3.3K27M/TP53 diffuse midline gliomas to radiation therapy | DISAGREE | We supplemented 10xGenomics' standard mouse brain content with a custom panel containing padlock probes resulting in 298 brain- and DMG-specific mRNA transcript assays. | validated_high_risk_disagreement \| Platform: Xenium |
| 10.1101/2024.11.10.622609 | 2024 | Spatio-temporal dynamics of the fibrotic niche in cardiac repair | DISAGREE | The Xenium Mouse Tissue Atlassing Panel (379 genes) was applied and topped up with 96 custom genes based on DEGs of each cell (sub)type identified in our scRNA-seq data. | validated_high_risk_disagreement \| Platform: Xenium |
| 10.1101/2024.12.15.628587 | 2024 | Molecular and spatial analysis of ganglion cells on retinal flatmounts: diversity, topography, and perivascularity | DISAGREE | The custom 140-Gene MERSCOPE Gene Panel Mix (Vizgen, VZG179) was added to the tissue section. | validated_high_risk_disagreement \| Platform: MERSCOPE |
| 10.1101/2025.01.14.632658 | 2025 | Intraductal Papillary Mucinous Neoplasm Cellular Plasticity is | DISAGREE | q1: Human Universal Cell Characterization Panel (1000-plex, RNA) with additional custom | validated_high_risk_disagreement \| Platform: CosMx |

| DOI | Year | Title | Audit decision | Evidence (verbatim quote) | Validation status / scope flag |
|---|---|---|---|---|---|
| | | Linked with Repeat Element Dysregulation | | probes capturing retrotransposons including LINE1 ORF1/ORF2, HSATII, HERV-K, and HERV-H. | |
| 10.1101/2025.02.07.637114 | 2025 | Somatic mutations distinguish melanocyte subpopulations in human skin | DISAGREE | q1: targeted gene panel was based on the predesigned human skin panel (v1), and an additional 100 genes were incorporated to create a custom panel. | validated_high_risk_disagreement \| Platform: Xenium |
| 10.1101/2025.02.12.638004 | 2025 | Endurance exercise remodels pulmonary vein sleeve myocytes and promotes a proarrhythmic atrial substrate | DISAGREE | q1: supplemented the 10x Genomics standard mouse profiling panel with a 100-gene custom panel with padlock probes. | validated_high_risk_disagreement \| Platform: Xenium |
| 10.1101/2025.02.20.639291 | 2025 | Utilizing combined spatial transcriptomics to elucidate localized immune responses within human coronary arteries throughout the progression of atherosclerosis | DISAGREE | q1: CosMx Universal Cell Characterization RNA panel with a core of 950 probes and an additional set of 20 custom probes informed by the GeoMx DSP study. | validated_high_risk_disagreement \| Fig. 2: excluded (platform/scope) |
| 10.1101/2025.04.23.649965 | 2025 | From Transcripts to Cells: Dissecting Sensitivity, Signal Contamination, and Specificity in Xenium Spatial Transcriptomics | DISAGREE | Xenium profiling used the targeted Breast panel, Lung panel, custom immuno-oncology panel (Custom IO), and the 5K PRIME panel. | validated_high_risk_disagreement \| Platform: Xenium |
| 10.1101/2025.05.16.654515 | 2025 | Oncogenic drivers shape the tumor microenvironment in human gliomas | DISAGREE | q1: a custom gene panel of 100 genes was added to the pre-designed human brain gene panel. | validated_high_risk_disagreement \| Platform: Xenium |
| 10.1101/2025.09.02.673834 | 2025 | Streamlining Spatial Transcriptomics for Human Kidney Tissue | DISAGREE | q1: 1000-plex CosMx TAP panel with the addition of kidney-specific custom genes; q2: TARGETED spatial COMP paper. | validated_high_risk_disagreement \| Platform: CosMx |
| 10.1101/2025.10.01.679230 | 2025 | Latent plasticity of the human pancreas across development, health, and disease | DISAGREE | q1 vendor evidence: pre-designed Xenium Prime 5K Human Pan Tissue & Pathways Panel; q1 manual evidence: additional 100 custom probes selected from DE genes or differentiation-pattern genes absent from the core panel. | validated_high_risk_disagreement \| Platform: Xenium |
| 10.1101/2025.10.20.683239 | 2025 | Evaluating exon-skipping therapies targeting the central nervous system in Duchenne muscular dystrophy using high-resolution spatial transcriptomics | DISAGREE | q1 vendor evidence: pre-designed Xenium mouse brain panel with 247 genes; q1 manual evidence: custom add-on panel of 50 genes including Dmd isoforms and exon-junction probes. | validated_high_risk_disagreement \| Platform: Xenium |
| 10.1111/imm.13577 | 2023 | Transcriptomic profiling of cardiac tissues from SARS-CoV-2 patients | DISAGREE | The data included more than 1800 genes, with 22 add-in COVID-19- | validated_high_risk_disagreement \| Fig. 2: excluded (platform/scope) |

| DOI | Year | Title | Audit decision | Evidence (verbatim quote) | Validation status / scope flag |
|---|---|---|---|---|---|
| | | identifies DNA damage. | | related genes, 4 SARS-CoV-2 specific genes, 2 negative control genes, and 32 internal reference genes; the packet also cites the COVID-19 Immune Atlas panel with an additional SARS-CoV-2 probe panel. | |
| 10.1126/science.adt0559 | 2026 | IDH-mutant gliomas arise from glial progenitor cells harboring the initial driver mutation | DISAGREE | The assay used 266 probes from the predesigned Xenium Human Brain Panel along with 73 additional custom probes. | validated_high_risk_disagreement \| Platform: Xenium |
| 10.1158/2767-9764.crc-24-0498 | 2025 | A Melanoma Brain Metastasis CTC Signature and CTC:B-cell Clusters Associate with Secondary Liver Metastasis: A Melanoma Brain-Liver Metastasis Axis. | DISAGREE | the standard pre-designed human multi-tissue/cancer panel ... as well as a custom designed panel [30 customized genes] | validated_high_risk_disagreement \| Platform: Xenium |
| 10.1172/jci179395 | 2025 | Disruption of ataxia telangiectasia-mutated kinase enhances radiation therapy efficacy in spatially directed diffuse midline glioma models. | DISAGREE | We supplemented 10x Genomics' standard mouse brain content with a custom panel | validated_high_risk_disagreement \| Platform: Xenium |
| 10.1186/s12935-025-04161-7 | 2026 | Single-cell spatial analysis identifies ID1-high endothelial cells in tertiary lymphoid structures as predictors of durable response to immunotherapy in non-small cell lung cancer | DISAGREE | The slides were hybridized with a 430-gene probe set consisting of a 380-gene Human Immune-Oncology base panel plus 50 custom endothelial-focused genes. | validated_high_risk_disagreement \| Platform: Xenium |
| 10.1186/s13058-025-02183-7 | 2026 | Spatial and temporal dynamics of cancer-associated fibroblast niches in breast cancer | AGREE | The FFPE human infiltrating ductal carcinoma obtained from BioIVT with Custom Add-on Panel-Tissue Sample 1 by 10x Genomics was used for the analysis. | accepted_agree \| Platform: Xenium |
| 10.1186/s13059-025-03624-4 | 2025 | Comparison of spatial transcriptomics technologies using tumor cryosections. | DISAGREE | The human generic brain panel was hybridized with 70 add-on genes, and the tool citation only references scGIST in the surrounding discussion. | validated_high_risk_disagreement \| Platform: Xenium |
| 10.1186/s40364-024-00658-x | 2024 | Single cell spatial profiling of FFPE splenic tissue from a humanized mouse model of HIV infection. | DISAGREE | CosMx SMI was evaluated using a 1000-plex human RNA panel and then applied with 5 custom-designed HIV probes and one human-specific probe. | validated_high_risk_disagreement \| Platform: CosMx |
| 10.1186/s40478-025-02106-9 | 2025 | Genomic heterogeneity drives distinct infiltration patterns in glioblastoma. | DISAGREE | The panel consisted of the 1000-plex Universal Cell panel and a 30- | validated_high_risk_disagreement \| Platform: CosMx |

| DOI | Year | Title | Audit decision | Evidence (verbatim quote) | Validation status / scope flag |
|---|---|---|---|---|---|
| | | | | gene custom panel designed to characterize malignant states. | |
| 10.1523/eneuro.0046-25.2025 | 2025 | Single-Cell Approaches Define the Murine Leptomeninges: Cortical Brain Interface as a Distinct Cellular Neighborhood Composed of Neural and Non-neural Cell Types | DISAGREE | The probeset targeted 347 genes comprised of the 10X Genomics Adult Brain Panel plus a custom add-on probeset. | validated_high_risk_disagreement \| Platform: Xenium |
| 10.21203/rs.3.rs-9150312/v1 | 2026 | A Targeted Spatial Transcriptomics Framework for a High-Resolution Human Skin Cell Atlas | DISAGREE | "a 479-gene panel derived from the Xenium Human Skin Gene Expression Panel" plus evidence_manual: "curated with additional markers from published scRNA-seq datasets to optimise delineation of major skin cell populations." | validated_high_risk_disagreement \| Platform: Xenium |
| 10.2220/biomedres.46.129 | 2025 | Multihormone-producing capabilities of normal pancreatic islet cells-spatial transcriptomics analysis using Xenium. | DISAGREE | "Probe Hybridization Mix was prepared using a pre-designed panel ... and custom panel" and evidence_manual says the 100 genes were selected from differential expression results. | validated_high_risk_disagreement \| Platform: Xenium |
| 10.3389/fimmu.2025.1518771 | 2025 | Comprehensive sequencing of the lung neuroimmune landscape in response to asthmatic induction | DISAGREE | "the selected 493 genes included 300 genes provided by the Xenium Mouse Tissue Atlassing panel" and "92 custom genes selected from genes of interest". | validated_high_risk_disagreement \| Platform: Xenium |
| 10.3390/cancers16071444 | 2024 | scRNAseq and High-Throughput Spatial Analysis of Tumor and Normal Microenvironment in Solid Tumors Reveal a Possible Origin of Circulating Tumor Hybrid Cells | DISAGREE | The evidence says the datasets were generated using the pre-designed panel of 325 genes and the add-on panel of 100 additional genes. | validated_high_risk_disagreement \| Platform: Xenium |
| 10.3390/cells12091310 | 2023 | Structured Illumination Microscopy Improves Spot Detection Performance in Spatial Transcriptomics. | DISAGREE | Several CARTANA neuronal panels were used, and five PLPs were generated for Actb, Gapdh, Atp1a3, and Slc17a7. | validated_high_risk_disagreement \| Platform: CARTANA |
| 10.3390/cells15030253 | 2026 | Temporal and Spatial Gene Expression Dynamics in Neonatal HI Hippocampus with Focus on Arginase. | DISAGREE | The packet says sections were hybridized with the Spatial Genomics Mouse Brain Mapper Panel, supplemented with additional genes related to the ARG1 pathway. | validated_high_risk_disagreement \| Platform: seqFISH |
| 10.64898/2025.12.08.692193 | 2025 | Biomarker Quantification in Breast Cancer using Xenium In Situ | DISAGREE | The packet says a fully custom standalone Xenium panel of 280 genes was designed using the | validated_high_risk_disagreement \| Platform: Xenium |

| DOI | Year | Title | Audit decision | Evidence (verbatim quote) | Validation status / scope flag |
|---|---|---|---|---|---|
| | | | | cloud-based Xenium Panel Designer. | |
| 10.64898/2026.01.07.698201 | 2026 | Fishing with Two Lines: A Hybrid Approach to Spatial Transcriptomic Discovery | DISAGREE | The dual-slide setup combined a custom 480-gene V1 panel with the Xenium Prime 5K Human Pan Tissue and Pathways panel. | validated_high_risk_disagreement \| Platform: Xenium |
| 10.64898/2026.02.24.707661 | 2026 | STRATA: Spatial Regulon Field Theory Reveals Coupling Architecture of Human Skin and Its Homogenization in Melanoma | DISAGREE | The Xenium Human Skin Gene Expression dataset used 382 genes comprising 282 pre-designed genes plus 100 custom add-on panel genes, so this is vendor-manual rather than pure vendor. | corrected_high_risk_disagreement_vendor_manual \| Platform: Xenium |
| 10.64898/2026.03.06.710199 | 2026 | MERFISH 2.0, an ultra-sensitive single-cell spatial transcriptomics imaging chemistry across diverse tissue types | DISAGREE | Five 815-plex MERFISH 2.0 gene panels were custom-designed using Vizgen's gene panel design portal, so this is vendor-assisted/manual panel design rather than a stock vendor panel. | corrected_high_risk_disagreement_vendor_manual \| Platform: MERSCOPE |

### VENDOR (n = 99)

*Fixed commercial or catalog panel.*

| DOI | Year | Title | Audit decision | Evidence (verbatim quote) | Validation status / scope flag |
|---|---|---|---|---|---|
| 10.1002/advs.202503932 | 2025 | A Human Immuno-Lung Organoid Model to Study Macrophage-Mediated Lung Cell Senescence Upon SARS-CoV-2 Infection. | AGREE | paper_type=EXP, panel_scope=TARGETED, spatial=YES; CosMx spatial transcriptomics with a 1000-plex RNA panel and no manual additions. | accepted_agree \| Platform: CosMx |
| 10.1002/advs.202513242 | 2026 | Developing a Single-Cell Spatial Transcriptomics Workflow for In Vivo Evaluation of Implanted Biomaterials. | DISAGREE | Spatial transcriptomics experiments were carried out with Xenium analyzer with the mouse tissue at lassing panel. | validated_high_risk_disagreement \| Platform: Xenium |
| 10.1002/advs.202516965 | 2026 | Spatially Resolved Multiomics Reveals Metabolic Remodeling and Autophagy Activation in Adamantinomatous Craniopharyngiomas. | AGREE | paper_type=EXP, panel_scope=TARGETED, spatial=YES; CosMx SMI platform with a 6K mRNA panel. | accepted_agree \| Platform: CosMx |
| 10.1002/path.70035 | 2026 | Xenium-based spatial transcriptomic analyses uncover prognosis-associated | AGREE | paper_type=EXP, panel_scope=TARGETED, spatial=YES; profiling was | accepted_agree \| Platform: Xenium |

| DOI | Year | Title | Audit decision | Evidence (verbatim quote) | Validation status / scope flag |
|---|---|---|---|---|---|
| | | heterogeneity in the tumor microenvironment (TME) of angioimmunoblastic T-cell lymphoma (AITL). | | performed using the Xenium Prime 5K Human Pan Tissue & Pathways Panel. | |
| 10.1007/s00535-025-02320-1 | 2026 | TP53 mutation predicts resistance to immune checkpoint inhibitor-based therapy in intrahepatic cholangiocarcinoma. | AGREE | paper_type=EXP, panel_scope=TARGETED, spatial=YES; probe hybridization used the Xenium Human 5K Pan Tissue and Pathways Panel. | accepted_agree \| Platform: Xenium |
| 10.1007/s13402-025-01100-6 | 2025 | Integrating multi-modal transcriptomics identifies cellular subtypes with distinct roles in PDAC progression. | AGREE | paper_type=EXP, panel_scope=TARGETED, spatial=YES; one sample came from the 10x Genomics FFPE Human PDAC dataset using the Human Immuno-Oncology Profiling Panel. | accepted_agree \| Platform: Xenium |
| 10.1016/j.canlet.2026.218451 | 2026 | IGHG1<sup>+</sup> malignant epithelial Cell-myCAF crosstalk via MIF-CD74/APP-CD74 drives early brain metastasis in NSCLC: Delineated via primary tumor-brain metastasis single-cell and spatial transcriptomics. | AGREE | paper_type=EXP, panel_scope=TARGETED, spatial=YES; the CosMx Human 1K Universal Panel targeting 1000 genes was used. | accepted_agree \| Platform: CosMx |
| 10.1016/j.ccell.2025.04.003 | 2025 | Spatial and multiomics analysis of human and mouse lung adenocarcinoma precursors reveals TIM-3 as a putative target for precancer interception | AGREE | q1: panel_design=VENDOR; uses_vendor_panel=YES; uses_manual_additions=NO. | accepted_agree \| Platform: Xenium |
| 10.1016/j.celrep.2024.114216 | 2024 | Microglia-astrocyte crosstalk in the amyloid plaque niche of an Alzheimer's disease mouse model, as revealed by spatial transcriptomics. | AGREE | In situ hybridisation was performed with the RNA mouse neuroscience panel, including the 18s RNA probe. | accepted_agree \| Platform: CosMx |
| 10.1016/j.jcmgh.2025.101689 | 2026 | Acute Phase Response-driven Hepatic Niche Remodeling Promotes Fibrosis Resolution After Alcohol Cessation. | AGREE | Assay was performed in CosMx Spatial Molecular Imager using Mouse Universal Cell Characterization Panel, 1000-plex, RNA with Cell Segmentation Kit. | accepted_agree \| Platform: CosMx |
| 10.1016/j.jcmgh.2025.101717 | 2026 | Novel Acinar Metaplastic States Uncovered in Exocrine Pancreas Disease. | AGREE | Human pancreas was profiled using CosMx Spatial Molecular Imager with the Human 6k Discovery Panel. | accepted_agree \| Platform: CosMx |
| 10.1016/j.jhep.2026.03.022 | 2026 | Liver Sinusoidal Endothelial Cells and Laminin dictate | AGREE | Fixed samples were used for Molecular Cartography according to the manufacturer’s instructions; | accepted_agree \| Platform: Molecular Cartography |

| DOI | Year | Title | Audit decision | Evidence (verbatim quote) | Validation status / scope flag |
|---|---|---|---|---|---|
| | | cholangiocytes' fate in chronic liver disease. | | vendor panel created by Resolve for the Resolve Molecular Cartography method. | |
| 10.1016/j.kint.2025.08.042 | 2026 | Single-cell spatial transcriptomics reveal intraglomerular cell activation and ligand-receptor relationships in chronic, active antibody mediated rejection. | AGREE | CosMx Spatial Molecular Imaging with a 1000-plex probe panel was used. | accepted_agree \| Platform: CosMx |
| 10.1016/j.labinv.2025.104210 | 2025 | Clinicopathologic, Cellular, and Molecular Analyses of Pulmonary Neuroendocrine Carcinoma With High Expression of Hepatocyte Nuclear Factor 4 Alpha | AGREE | The gene list of the custom lung panel (design ID: 9GT3BT) was previously reported. | accepted_agree \| Platform: Xenium |
| 10.1016/j.labinv.2026.106079 | 2026 | High-Resolution Spatial Transcriptomic Characterization of Syncytial Variant of Nodular Sclerosis Classical Hodgkin Lymphoma. | AGREE | Spatial transcriptomic profiling was performed using the Xenium Prime 5K panel. | accepted_agree \| Platform: Xenium |
| 10.1016/j.modpat.2024.100681 | 2025 | Spatially Resolved Single-Cell Transcriptome Analysis of Mycosis Fungoides Reveals Distinct Biomarkers GNLY and FYB1 Compared With Psoriasis and Chronic Spongiotic Dermatitis. | AGREE | Single-cell transcriptomics ... were analyzed using CosMx spatial molecular imager utilizing surface markers, including CD3 and CD4. | accepted_agree \| Platform: CosMx |
| 10.1016/j.neuron.2025.07.025 | 2025 | Molecular and spatial analysis of ganglion cells on retinal flatmounts identifies perivascular neurons resilient to glaucoma. | AGREE | q1 says the 140-gene MERSCOPE Gene Panel Mix was added; q2 says the study targeted 140 genes in MERFISH on a targeted spatial paper. | accepted_agree \| Platform: MERSCOPE |
| 10.1016/j.saa.2026.127532 | 2026 | Accurate classification of ependymomas and medulloblastomas using Raman spectroscopy and pilot transcriptomic profiling. | AGREE | q1 and q2 both describe a pilot spatial-transcriptomics analysis using the Xenium In Situ v2 FFPE platform. | accepted_agree \| Platform: Xenium |
| 10.1016/j.tranon.2025.102573 | 2026 | Application of CTC-derived spheroid for drug screening toward personalized treatment in patients with breast cancer. | AGREE | q1 says FFPE samples were processed into Xenium in situ slides with the Xenium Human Breast Gene Expression Panel. | accepted_agree \| Platform: Xenium |
| 10.1016/j.xcrm.2026.102713 | 2026 | Single-cell spatial transcriptomics reveals tumor microenvironment heterogeneity in primary and lymph node-metastatic small cell lung cancer. | AGREE | q1 says the CosMx 1000-plex RNA panel was used strictly following the manufacturer protocol. | accepted_agree \| Platform: CosMx |

| DOI | Year | Title | Audit decision | Evidence (verbatim quote) | Validation status / scope flag |
|---|---|---|---|---|---|
| 10.1016/j.ydbio.2025.10.017 | 2026 | Poly(A) probe HCR RNA-FISH specifically marks pyriform nurse cells in the brown anole lizard ovary. | AGREE | q1 says the probes were designed and synthesized by Molecular Instruments targeting polyadenylated RNA. | accepted_agree \| Platform: HCR-FISH |
| 10.1017/s1551929521001346 | 2021 | Using MERSCOPE to Generate a Cell Atlas of the Mouse Brain that Includes Lowly Expressed Genes | AGREE | q1 says the MERFISH Mouse Brain Receptor Map was generated on Vizgen's MERSCOPE platform; q2 says it used a gene panel of 483 genes. | accepted_agree \| Platform: MERSCOPE |
| 10.1038/s41380-024-02651-0 | 2025 | Single-cell spatial transcriptomics reveals distinct patterns of dysregulation in non-neuronal and neuronal cells induced by the Trem2<sup>R47H</sup> Alzheimer's risk gene mutation. | AGREE | The gene panel was constructed based on the commercially available 300-gene set from Vizgen ... | accepted_agree \| Platform: MERSCOPE |
| 10.1038/s41467-024-47271-y | 2024 | Spatial transcriptomics reveals discrete tumour microenvironments and autocrine loops within ovarian cancer subclones. | AGREE | A 960 gene panel was applied to a serial section of the biopsy profiled by Visium from patient 5. | accepted_agree \| Platform: CosMx |
| 10.1038/s41467-025-65350-6 | 2025 | Single-nucleus and spatial transcriptomic profiling of human temporal cortex and white matter reveals key associations with AD pathology. | AGREE | The probes used belonged to a panel of 155 genes, designed for this study by CARTANA, based on their state-of-the-art guidelines. | accepted_agree \| Platform: CARTANA |
| 10.1038/s41586-025-09907-x | 2026 | Bidirectional CRISPR screens decode a GLIS3-dependent fibrotic cell circuit | AGREE | A customized 480-plex gene probe panel was designed and used for all human samples; uses_vendor_panel=YES; panel_design=VENDOR. | accepted_agree \| Platform: Xenium |
| 10.1038/s41586-026-10377-y | 2026 | A spatial atlas of the healthy human liver from live donors | AGREE | The evidence_vendor field says the predesigned pan cancer pathways 500-gene panel was used according to the MERSCOPE user guide. | accepted_agree \| Platform: MERSCOPE |
| 10.1038/s41598-023-42686-x | 2023 | Plasticity of cone photoreceptors in adult zebrafish revealed by thyroid hormone exposure. | AGREE | Transcript-selective probe sets were designed and generated by Molecular Instruments and can be ordered directly from their website. | accepted_agree \| Platform: HCR-FISH |
| 10.1038/s41598-025-26735-1 | 2025 | One section, two worlds: single-cell integration of MALDI-MSI and spatial transcriptomics on the same single tissue section. | DISAGREE | Probe hybridization, ligation, and amplification were carried out using the pre-designed Mouse Brain and Human Brain reagent in accordance with Protocol CG000749. | validated_high_risk_disagreement \| Platform: Xenium |

| DOI | Year | Title | Audit decision | Evidence (verbatim quote) | Validation status / scope flag |
|---|---|---|---|---|---|
| 10.1038/s42003-025-08518-6 | 2025 | Beyond the nuclear border: single-cell analysis of in situ sequenced human brain tissue using cellular features. | DISAGREE | We used the 10x Genomics Human Brain Panel (hBP), which includes 266 genes designed to identify the major cell types in the human CNS. | validated_high_risk_disagreement \| Platform: Xenium |
| 10.1038/s42003-026-09837-y | 2026 | Spatial profiling uncovers multicellular dynamics in early relapse of hepatitis B virus-associated follicular lymphoma. | AGREE | The study used the CosMx Spatial Molecular Imaging system with a 1,010-plex RNA panel. | accepted_agree \| Platform: CosMx |
| 10.1093/nargab/lqag007 | 2026 | SM3DD with segmented PCA: a comprehensive method for interpreting 3D spatial transcriptomics. | DISAGREE | q1: Adjacent serial tissue sections were profiled using the CosMx SMI 1000-plex assay; q2: TARGETED spatial COMP paper. | validated_high_risk_disagreement \| Platform: CosMx |
| 10.1093/neuonc/noaf146 | 2025 | Cranial radiotherapy profoundly affects glia without inducing widespread cellular senescence. | AGREE | 5 µm slices put on slide for spatial single cell transcriptomics using the CosMx Technology Access Program (Nanostring) according to manufacturer's recommendations with the Mouse RNA Neuroscience panel (1.0). | accepted_agree \| Platform: CosMx |
| 10.1093/neuonc/noag026 | 2026 | Spatial Transcriptomics Characterisation of Radionecrotic Changes in Glioblastoma Patients. | AGREE | We employed the off-the-shelf Xenium Human Brain Panel consisting of 266 genes. | accepted_agree \| Platform: Xenium |
| 10.1093/noajnl/vdad142 | 2023 | Spatial architecture of high-grade glioma reveals tumor heterogeneity within distinct domains. | NEEDS_MANUAL | CosMx uses '1000 RNA probes' and Xenium is analyzed as vendor platform data | pdf_validated_unclear_update \| Platform: CosMx |
| 10.1101/2021.10.11.463936 | 2021 | Highly resolved spatial transcriptomics for detection of rare events in cells | DISAGREE | Transcript-specific probes are designed using an algorithm developed specifically for MC by Resolve BioSciences that allows the design of probe sets of almost all genes of an organism. | validated_high_risk_disagreement \| Platform: Molecular Cartography |
| 10.1101/2023.01.04.519955 | 2023 | Spatial molecular imaging of the human type 2 diabetic islet | AGREE | an array of probes representing 990 human genes ... NanoString® ISH probes were prepared ... | accepted_agree \| Platform: CosMx |
| 10.1101/2023.06.29.546675 | 2023 | Applying high-resolution spatial transcriptomics to characterize the amyloid plaque cell niche in Alzheimer's Disease | AGREE | In situ hybridisation was performed with the RNA mouse neuroscience panel, including the 18s RNA probe. | accepted_agree \| Platform: CosMx |
| 10.1101/2023.12.13.571385 | 2023 | A Comparative Analysis of Imaging-Based Spatial Transcriptomics Platforms | AGREE | Two consecutive sections were processed using each platform's commercial iterations, utilizing pre- | accepted_agree \| Platform: Xenium;CosMx |

| DOI | Year | Title | Audit decision | Evidence (verbatim quote) | Validation status / scope flag |
|---|---|---|---|---|---|
| | | | | designed probe sets for general tissue profiling. | |
| 10.1101/2023.12.30.573739 | 2024 | SORBET: Automated cell-neighborhood analysis of spatial transcriptomics or proteomics for interpretable sample classification via GNN | DISAGREE | q1: sequenced using the CosMx Spatial Molecular Imager platform; 960 targeted transcripts; q2: TARGETED spatial COMP paper. | validated_high_risk_disagreement \| Platform: CosMx |
| 10.1101/2024.11.27.625536 | 2024 | Sub-cellular Imaging of the Entire Protein-Coding Human Transcriptome (18933-plex) on FFPE Tissue Using Spatial Molecular Imaging | AGREE | Bruker Spatial Biology has accomplished a commercial-grade panel (CosMx Spatial Molecular Imager Whole Transcriptome Panel). | accepted_agree \| Platform: CosMx |
| 10.1101/2025.02.07.25321742 | 2025 | Spatial biomarkers of response to neoadjuvant therapy in muscle-invasive bladder cancer: the DUTRENEO trial | AGREE | q1: Xenium Human Multi-tissue and Cancer Panel targeting 377 human markers; no manual additions or tool evidence. | accepted_agree \| Platform: Xenium |
| 10.1101/2025.02.11.637091 | 2025 | Optimisation of Xenium automated in situ sequencing for PAXgene-fixed tissue samples | AGREE | q1: assayed with the 379-plex 10x Genomics pre-designed Xenium Mouse Tissue Atlassing Panel (mMulti). | accepted_agree \| Platform: Xenium |
| 10.1101/2025.02.22.639181 | 2025 | A spatial transcriptomics atlas of live donors reveals unique zonation patterns in the healthy human liver | AGREE | q1: predesigned pan cancer pathways 500 gene panel used on the MERSCOPE platform. | accepted_agree \| Platform: MERSCOPE |
| 10.1101/2025.03.05.641608 | 2025 | Epithelial Plasticity and an Immune Suppressive Microenvironment Underpin Tumour Budding in Colorectal Cancer | AGREE | q1: CosMx SMI technology deploying the human 6K discovery RNA Panel (6,175 genes, including 20 negative control probes). | accepted_agree \| Platform: CosMx |
| 10.1101/2025.03.24.644582 | 2025 | Integrated single-cell and spatial analysis identifies context-dependent myeloid-T cell interactions in head and neck cancer immune checkpoint blockade response | AGREE | q1: Nanostring CosMx single-cell spatial omics with RNA gene panel including 435 ligands and receptors. | accepted_agree \| Platform: CosMx |
| 10.1101/2025.05.20.655231 | 2025 | Characterization of Bergmann Glia-like Progenitors in the Postnatal Mouse Cerebellum by <i>in vivo</i> Electroporation and Spatial Transcriptomics | AGREE | q1: Xenium Prime 5 K Mouse Pan Tissue and Pathways Panel, which targets 5001 genes, was used for the experiment. | accepted_agree \| Platform: Xenium |
| 10.1101/2025.05.22.655586 | 2025 | Spatial Biology and Organoid Technologies Reveal a Potential Therapy-Resistant Cancer Stem Cell Population in Pancreatic Ductal Adenocarcinoma | AGREE | q1: CosMx SMI instrument was used for slide scanning and detection of target molecules/genes in a 1000 plex panel. | accepted_agree \| Platform: CosMx |

| DOI | Year | Title | Audit decision | Evidence (verbatim quote) | Validation status / scope flag |
|---|---|---|---|---|---|
| 10.1101/2025.06.01.657310 | 2025 | Multimodal Validation of the Existence of Transitional Cerebellar Progenitors in the Human Fetal Cerebellum | AGREE | q1: Xenium Prime in situ analysis was performed using the Xenium Prime 5K Human Pan-Tissue and Pathways Assay Kit (10x Genomics, PN-1000671). | accepted_agree \| Platform: Xenium |
| 10.1101/2025.06.16.658716 | 2025 | The Spatial Atlas of Human Anatomy (SAHA): A Multimodal Subcellular-Resolution Reference Across Human Organs | AGREE | q1 vendor evidence: CosMx Human Universal Cell Characterization Panel, CosMx Human 6K Discovery Panel, Xenium Human Multi-Tissue and Cancer Panel, and Xenium Human Breast Panel; q1 tool evidence: NOT FOUND. | accepted_agree \| Platform: Xenium;CosMx |
| 10.1101/2025.06.23.661209 | 2025 | Comparative Analysis of Neonatal Hypomyelination Models Using Spatial Transcriptomics | NEEDS_MANUAL | 'CosMx, Novogene' and 'a predefined gene panel was analyzed' | pdf_validated_unclear_update \| Platform: CosMx |
| 10.1101/2025.07.07.663390 | 2025 | Decoding Lymphangioleiomyomatosis (LAM) Niche Environment via Integrative Analysis of Single Cell Multiomics and Spatial Transcriptomics | AGREE | q1 vendor evidence: the Xenium platform uses a panel of 377 human cancer-related genes pre-determined by 10x Genomics; no tool or manual-addition evidence found. | accepted_agree \| Platform: Xenium |
| 10.1101/2025.09.08.674825 | 2025 | A transcriptional map of human tonsil architecture: beyond the sum of (single cell) parts | AGREE | q1 vendor evidence: CosMx Human Universal Cell Characterization Panel (RNA, 1000 Plex) probes; no manual additions or tool evidence found. | accepted_agree \| Platform: CosMx |
| 10.1101/2025.10.31.685702 | 2025 | Spatial Transcriptomics Identify T Cell-Driven Mechanisms of Kidney Damage in Immune Checkpoint Inhibitor-Associated Acute Interstitial Nephritis | AGREE | q1 vendor evidence: samples were hybridized with the Xenium Prime 5K Human Pan Tissue & Pathways Panel; no manual additions or canonical tools found. | accepted_agree \| Platform: Xenium |
| 10.1101/2025.11.19.689127 | 2025 | Single-cell Spatial Transcriptional Profiling Uncovers Heterogeneous Cellular Responses to Pathogenic Tau in a Mouse Model of Neurodegeneration | AGREE | q1 vendor evidence: CosMx Mouse Neuroscience panel targeting 950 genes; no custom additions or canonical tools found. | accepted_agree \| Platform: CosMx |
| 10.1101/2025.11.24.690328 | 2025 | Spatial Transcriptomics Reveals <i>CXCL12</i> $^{+}$ Fibroblasts as Central Immune Organizers through CXCR4 Signaling in Abdominal Aortic Aneurysm | AGREE | Processed with the manufacturer protocol using the Xenium Human Gene Expression 5K panel containing probes for 5001 genes. | accepted_agree \| Platform: Xenium |

| DOI | Year | Title | Audit decision | Evidence (verbatim quote) | Validation status / scope flag |
|---|---|---|---|---|---|
| 10.1101/2025.11.28.691139 | 2025 | Tissue architecture and immune niches govern ctDNA release in colorectal cancer | AGREE | The study used the Xenium Human Colon Gene Expression Panel (352 genes). | accepted_agree \| Platform: Xenium |
| 10.1111/bpa.13322 | 2025 | Neuropathological and cerebrospinal fluid correlates of choroid plexus inflammation in progressive multiple sclerosis. | AGREE | The expression of 157 targeted inflammatory genes available for the Cartana system was assessed. | accepted_agree \| Platform: CARTANA |
| 10.1128/aem.00890-15 | 2015 | Use of Hybridization Chain Reaction-Fluorescent In Situ Hybridization To Track Gene Expression by Both Partners during Initiation of Symbiosis. | AGREE | The nucleic acid probes and hairpin sequences were obtained from Molecular Instruments, which also assisted with the probe design. | accepted_agree \| Platform: HCR-FISH |
| 10.1136/jitc-2025-012089 | 2025 | Spatial TCR clonality and clonal expansion in the in situ microenvironment of non-small cell lung cancer. | AGREE | padlock probe hybridization and rolling circle amplification (RCA) were performed according to the High Sensitivity Library Preparation Kit by Cartana | accepted_agree \| Platform: CARTANA |
| 10.1136/jitc-2025-012613 | 2025 | High Treg and PMN-MDSC densities are a hallmark of tertiary lymphoid structures in fatal cases of cervical cancer | DISAGREE | vendor = Xenium Human Immuno-Oncology panel | validated_high_risk_disagreement \| Platform: Xenium |
| 10.1136/jitc-2025-012991 | 2025 | Spatial transcriptomic profiling of metastatic renal cell carcinoma identifies chemokine-driven macrophage and CD8+ T-cell interactions predictive of immunotherapy response. | AGREE | Tissues were hybridized with 993-probe panels (CosMx Human Universal Cell Characterization Panel) | accepted_agree \| Platform: CosMx |
| 10.1158/1078-0432.ccr-23-0827 | 2023 | Trackable Intratumor Microdosing and Spatial Profiling Provide Early Insights into Activity of Investigational Agents in the Intact Tumor Microenvironment. | AGREE | using the NanoString Cancer Transcriptome Atlas (CTA) probe set | accepted_agree \| Platform: CosMx |
| 10.1158/1078-0432.ccr-25-2300 | 2026 | Integrated single-cell and spatial analysis reveals context-dependent myeloid-T cell interactions in response to immune checkpoint blockade in head and neck cancer. | AGREE | utilizing a 64-plex protein panel and a 1000-gene RNA panel | accepted_agree \| Platform: CosMx |
| 10.1158/2767-9764.crc-25-0314 | 2025 | In Situ Proinflammatory Effects of Dazostinag Alone or with Chemotherapy on the Tumor Microenvironment of Patients with Head and Neck Squamous Cell Carcinoma. | NEEDS_MANUAL | GeoMx DSP of '~1,800 target genes' and CosMx SMI are used | pdf_validated_unclear_update \| Fig. 2: excluded (platform/scope) |

| DOI | Year | Title | Audit decision | Evidence (verbatim quote) | Validation status / scope flag |
|---|---|---|---|---|---|
| 10.1165/rcmb.2025-0261oc | 2025 | Ectopic Expression of <i>MUC5B</i> in the Respiratory Bronchiole Initiates Endoplasmic Reticulum Stress in the IPF Lung. | AGREE | We included two slides using the 1000-plex human RNA panel and 24 custom genes | accepted_agree \| Platform: CosMx |
| 10.1186/s12915-024-01922-0 | 2024 | 3D exploration of gene expression in chicken embryos through combined RNA fluorescence in situ hybridization, immunofluorescence, and clearing. | AGREE | All probes were designed and purchased from Molecular Instruments; the probe sets consist of 19 to 20 pairs of oligos recognizing each target mRNA. | accepted_agree \| Platform: HCR-FISH |
| 10.1186/s12935-025-03635-y | 2025 | Characterizing tertiary lymphoid structures associated single-cell atlas in breast cancer patients. | AGREE | One dataset consists of single-cell resolution sequencing data from the Xenium platform, which in this dataset measured 313 genes. | accepted_agree \| Platform: Xenium |
| 10.1186/s13045-025-01773-4 | 2026 | A subcellular spatial atlas illuminates the microenvironmental remodeling of perineural invasion in distal cholangiocarcinoma | AGREE | Fourteen cores were subjected to spatial profiling of over 5,000 genes using the Xenium subcellular resolution spatial transcriptomics platform. | accepted_agree \| Platform: Xenium |
| 10.1186/s13073-024-01422-4 | 2024 | Spatial transcriptome profiling identifies DTX3L and BST2 as key biomarkers in esophageal squamous cell carcinoma tumorigenesis. | AGREE | The study used UV-photocleavable barcode-conjugated RNA probes sourced from the GeoMx Cancer Transcriptome Atlas and the CosMx Human 6K Discovery Panel. | accepted_agree \| Fig. 2: excluded (platform/scope) |
| 10.1200/jco.2024.42.4_suppl.468 | 2024 | Increased spatial coupling of integrin and collagen IV in the immunoresistant clear cell renal cell carcinoma tumor microenvironment. | AGREE | SMI was obtained using the human Immuno-oncology panel consisting of 978 RNA probes. | accepted_agree \| Platform: CosMx |
| 10.1242/dev.156869 | 2018 | Multidimensional quantitative analysis of mRNA expression within intact vertebrate embryos. | AGREE | For each target mRNA, a kit containing a DNA probe set and HCR amplifier was purchased from Molecular Instruments. | accepted_agree \| Platform: HCR-FISH |
| 10.1371/journal.pone.0281464 | 2023 | Induction of flight via midbrain projections to the cuneiform nucleus. | AGREE | CARTANA provided a list of genes and designed and provided the padlock probes. | accepted_agree \| Platform: CARTANA |
| 10.21203/rs.3.rs-4376986/v2 | 2024 | Spatial Single-Cell Transcriptomic Analysis in Breast Cancer Reveals Potential Biomarkers for PD1 Blockade Therapy | AGREE | "This was followed by overnight hybridization with Xenium DNA probes (Xenium Human Breast Gene Expression Probes 2000826, detailed in Supp. Table 3)" and uses_vendor_panel=YES. | accepted_agree \| Platform: Xenium |

| DOI | Year | Title | Audit decision | Evidence (verbatim quote) | Validation status / scope flag |
|---|---|---|---|---|---|
| 10.21203/rs.3.rs-4590465/v1 | 2024 | Single-Cell, Single-Nucleus and Xenium-Based Spatial Transcriptomics Analyses Reveal Inflammatory Activation and Altered Cell Interactions in the Hippocampus in Mice with Temporal Lobe Epilepsy. | AGREE | "Xenium-based (mouse brain panel, 247 genes) spatial transcriptomics analyses" and uses_vendor_panel=YES. | accepted_agree \| Platform: Xenium |
| 10.21203/rs.3.rs-5012685/v1 | 2024 | In situ spatial transcriptomic analysis of human skeletal muscle using the Xenium platform | AGREE | "probe hybridization was performed using the predesigned Human Multi-Tissue and Cancer Panel (10x Genomics, CA, USA; PN 1000626)" and uses_vendor_panel=YES. | accepted_agree \| Platform: Xenium |
| 10.21203/rs.3.rs-6772872/v1 | 2025 | Digital Pathology Predicts Immunotherapy Response and Maps Spatial Biomarkers in Gastric Cancer | AGREE | "processed according to the Xenium Prime In Situ Gene Expression... along with the Xenium Prime 5K Human Pan Tissue & Pathways Panel" and uses_vendor_panel=YES. | accepted_agree \| Platform: Xenium |
| 10.21203/rs.3.rs-7728048/v1 | 2025 | Deep Spatial Transcriptomic Profiling of Ovarian Clear Cell Carcinoma in the Real-World Setting | AGREE | "Overnight in situ probe hybridization with the probes from Xenium Prime 5K Human pan tissue & pathways panel... were performed according to the vendor's user guide." | accepted_agree \| Platform: Xenium |
| 10.21203/rs.3.rs-7971511/v1 | 2025 | Spatial transcriptomics links hepatocyte-macrophage interactions to viral signatures in seronegative hepatitis | AGREE | "processed using the Human Universal Cell Characterisation 1000-plex panel ... following the manufacturer's protocol" and uses_vendor_panel=YES. | accepted_agree \| Platform: CosMx |
| 10.3389/fimmu.2025.1728629 | 2025 | The local cellular response to Human Papillomavirus focuses on basal layer restoration as visualized in situ by specific cellular neighborhoods near infected cells | AGREE | "CosMx 1000-plex spatial transcriptomics" and uses_vendor_panel=YES. | accepted_agree \| Platform: CosMx |
| 10.3390/ijms27031297 | 2026 | IL-17 Cytokines Induce IκBζ in Dermal Fibroblasts to Promote Pro-Inflammatory Gene Expression in Psoriasis. | AGREE | Spatial transcriptomic profiling was performed using the CosMx Human 6000-plex RNA Panel following the manufacturer's protocol. | accepted_agree \| Platform: CosMx |
| 10.64898/2025.12.03.691993 | 2025 | geneSCOPE: gene Spatial Co-Occurrence of Pairwise Expression | DISAGREE | q1: analyzed an FFPE human lymph node section profiled with the 10x Genomics Xenium 5K | validated_high_risk_disagreement \| Platform: Xenium |

| DOI | Year | Title | Audit decision | Evidence (verbatim quote) | Validation status / scope flag |
|---|---|---|---|---|---|
| | | | | Pan-Tissue & Pathways gene panel (4644 genes in this case). | |
| 10.64898/2025.12.09.693138 | 2025 | A spatial atlas of colorectal cancer reveals the influence of stromal niches on tumour differentiation | AGREE | The packet says the CosMx Human Universal Cell Characterization Panel was utilised to profile 1000 genes in normal colon and tumour tissue specimens. | accepted_agree \| Platform: CosMx |
| 10.64898/2025.12.22.693132 | 2025 | Spatial transcriptomic profiling of decalcified murine musculoskeletal samples via Xenium Prime 5K | DISAGREE | Title names Xenium Prime 5K in a targeted spatial transcriptomic profiling paper; packet has no custom-addition evidence, so this is vendor panel/platform use. | validated_medium_risk_disagreement \| Platform: Xenium |
| 10.64898/2025.12.22.695505 | 2025 | Diverse infections transcriptionally reprogram the intestinal epithelium and epithelial-immune cell interactions | AGREE | gemma_q1_panel_design explicitly names the pre-designed Xenium Prime 5K Mouse Gene Expression Panel. | accepted_agree \| Platform: Xenium |
| 10.64898/2025.12.30.696765 | 2025 | Clinical feasibility of spatial transcriptomics using discarded tissue from diagnostic breast biopsies | AGREE | The packet says seqFISH was performed using Spatial Genomics with a 493-gene immune-oncology panel. | accepted_agree \| Platform: seqFISH |
| 10.64898/2026.01.23.701261 | 2026 | GxP Single-cell RNA-seq and Spatial Transcriptomics end-to-end pipeline for clinical research | DISAGREE | q1: Xenium Kidney (both non-diseased and cancer): Xenium Human Multi-Tissue and Cancer Panel; q2: targeted Xenium clinical pipeline context. | validated_high_risk_disagreement \| Platform: Xenium |
| 10.64898/2026.01.25.701559 | 2026 | Spatial pharmaco-multiomics reveals drug distribution, metabolic niches, and spatially constrained resistance in medulloblastoma | AGREE | The packet says tissue mRNA was targeted using the Xenium Human Brain Gene Expression Panel and hybridised overnight. | accepted_agree \| Platform: Xenium |
| 10.64898/2026.02.03.703526 | 2026 | GM-CSF regulates ILC states and myeloid cell signaling during ulceration in Crohn’s disease | NEEDS_MANUAL | Xenium In Situ is run under the 10x protocol and the text says 'Only genes included in the panel as cell type markers were used' | pdf_validated_unclear_update \| Platform: Xenium |
| 10.64898/2026.02.16.706245 | 2026 | Wayfarer: A multiscale framework for spatial analysis of tumor progression | DISAGREE | q1: Visium and Xenium (302 genes) data from adjacent sections; q2: targeted Xenium LUAD context. | validated_high_risk_disagreement \| Platform: Xenium |
| 10.64898/2026.02.18.700929 | 2026 | Combining xenium <i>in situ</i> spatial transcriptomics and imaging mass cytometry on a single tissue section | AGREE | Samples were analysed using the Xenium Human Multi-Tissue and Cancer Expression Panel (10X Genomics, PN-1000626) | accepted_agree \| Platform: Xenium |
| 10.64898/2026.03.03.709035 | 2026 | Convergent stromal and immune remodeling defines spatial tumor | AGREE | the samples were incubated ... with the Xenium Prime Human | accepted_agree \| Platform: Xenium |

| DOI | Year | Title | Audit decision | Evidence (verbatim quote) | Validation status / scope flag |
|---|---|---|---|---|---|
| | | dynamics in PARP inhibitor–resistant high-grade serous ovarian cancer | | Panel which targets 5000 human genes | |
| 10.64898/2026.03.17.712266 | 2026 | Cross-Modal Training Using Xenium Spatial Transcriptomics Enables DINO-DETR Based Detection of Vascular Niches in H&E Whole-Slide Images | DISAGREE | q1 says the study used the Xenium Human Immuno-Oncology Profiling Panel (380 genes) and followed the manufacturer's protocol. | validated_high_risk_disagreement \| Platform: Xenium |
| 10.64898/2026.03.17.712495 | 2026 | Spatio-temporal mapping of immune cell dynamics during human sequential lymph node metastasis | AGREE | Probes from the Xenium Prime 5K panel were hybridised overnight at 50°C. | accepted_agree \| Platform: Xenium |
| 10.64898/2026.03.20.713219 | 2026 | Hidden immune memory niches in inflammatory skin diseases | AGREE | All Xenium slides were processed using the Xenium Prime 5K Human Pan Tissue & Pathways Panel | accepted_agree \| Platform: Xenium |
| 10.64898/2026.03.23.713195 | 2026 | Spatial multi-omics of multiple myeloma uncovers niche-dependent pro-myeloma and immunosuppressive signaling in the bone marrow and extramedullary lesions | DISAGREE | Although the Xenium assay targets a curated panel of ~5,000 genes rather than the full transcriptome | validated_high_risk_disagreement \| Platform: Xenium |
| 10.70322/jrbtm.2025.10004 | 2025 | Enhanced Spatial Transcriptomics Analysis of Mouse Lung Tissues Reveals Cell-Specific Gene Expression Changes Associated with Pulmonary Hypertension. | AGREE | Xenium Mouse Tissue Atlassing Panel | accepted_agree \| Platform: Xenium |
| 10.7150/ijbs.114618 | 2025 | Single-cell and spatial transcriptomic analysis reveal cellular heterogeneity and cancer cell-intrinsic major histocompatibility complex II expression in urothelial carcinoma | AGREE | gemma_q1_panel_design cites Visium Spatial Gene Expression Reagent Kits and no manual or tool-based panel selection. | accepted_agree \| Fig. 2: excluded (platform/scope) |
| 10.7554/elife.107070 | 2025 | Evidence of off-target probe binding in the 10x Genomics Xenium v1 Human Breast Gene Expression Panel compromises accuracy of spatial transcriptomic profiling | DISAGREE | The study analyzes the 10x Genomics Xenium v1 Human Breast Gene Expression Panel and obtains probe FASTA from the publication. | validated_high_risk_disagreement \| Platform: Xenium |

### MANUAL-PANEL (n = 304)

*Panel curated from biological markers, differential expression, ranking, pathway lists, or literature — no canonical tool drove the final selection.*

| DOI | Year | Title | Audit decision | Evidence (verbatim quote) | Validation status / scope flag |
|---|---|---|---|---|---|
| 10.1002/advs.202415756 | 2025 | PLXDC1<sup>+</sup> Tumor-Associated Pancreatic Stellate Cells Promote Desmoplastic and Immunosuppressive Niche in Pancreatic Ductal Adenocarcinoma. | AGREE | paper_type=EXP, panel_scope=TARGETED, spatial=YES; selected EPCAM, KRT17, S100A2, PDPN, LRRC15, ACTA2, PLXDC1, CD68, SPP1, CD8A, TIGIT, and PDCD1 by expert choice. | accepted_agree \| Platform: in situ sequencing |
| 10.1002/advs.202519816 | 2026 | Spatial Transcriptomics of TMJ Reveals a Remodeling Fibroblast-Immune Microenvironment Driving Arthritis Pain. | AGREE | paper_type=EXP, panel_scope=TARGETED, spatial=YES; 96 target genes were selected based on cell-type or cell-status markers from prior scRNA-seq studies. | accepted_agree \| Platform: seqFISH |
| 10.1002/cne.25628 | 2024 | Cellular-resolution gene expression mapping reveals organization in the head ganglia of the gastropod, Berghia stephanieae. | AGREE | paper_type=EXP, panel_scope=TARGETED, spatial=YES; standard clustering and differential gene expression produced candidate genes explored with multiplexed in situ HCR. | accepted_agree \| Platform: HCR-FISH |
| 10.1002/cne.70016 | 2025 | Localization of Cholecystokinin/Sulfakinin Neuropeptides in Biomphalaria glabrata, an Intermediate Host for Schistosomiasis. | AGREE | paper_type=EXP, panel_scope=TARGETED, spatial=YES; hybridization used a probe set generated for the B. alexandrina CCK/SK transcript. | accepted_agree \| Platform: HCR-FISH |
| 10.1002/dvdy.573 | 2023 | Single cell sequencing of the mouse anterior palate reveals mesenchymal heterogeneity. | AGREE | paper_type=EXP, panel_scope=TARGETED, spatial=YES; one to two distinct markers per cluster were chosen to determine spatial localization. | accepted_agree \| Platform: HCR-FISH |
| 10.1002/jemt.24561 | 2024 | Coupling clearing and hybridization chain reaction approaches to investigate gene expression in organs inside intact insect heads. | AGREE | paper_type=EXP, panel_scope=TARGETED, spatial=YES; two genes were chosen by the authors and probe pairs were designed with HCR probe 3.0 Maker. | accepted_agree \| Platform: HCR-FISH |
| 10.1002/smtd.202501615 | 2026 | Efficient Spatial Gene Expression Profiling Using Split-Probe Ligation and Rolling Circle Amplification. | DISAGREE | Genes were manually curated from published literature and selected from scRNA-seq datasets, with DEGs from bulk RNA-seq added. | validated_high_risk_disagreement \| Platform: in situ sequencing |
| 10.1007/s00216-024-05327-4 | 2024 | Multiplexed in situ RNA imaging by combFISH. | DISAGREE | q1 says the authors selected 14 marker genes that were cell type-specific based on previous results. | validated_high_risk_disagreement \| Platform: in situ sequencing |

| DOI | Year | Title | Audit decision | Evidence (verbatim quote) | Validation status / scope flag |
|---|---|---|---|---|---|
| 10.1007/s00429-020-02194-4 | 2021 | Optimization and evaluation of fluorescence in situ hybridization chain reaction in cleared fresh-frozen brain tissues. | AGREE | paper_type=EXP, panel_scope=TARGETED, spatial=YES; probes were designed in the lab based on prior HCR guidance. | accepted_agree \| Platform: HCR-FISH |
| 10.1007/s11427-023-2420-7 | 2023 | Integrated neural tracing and in-situ barcoded sequencing reveals the logic of SCN efferent circuits in regulating circadian behaviors | AGREE | paper_type=EXP, panel_scope=TARGETED, spatial=YES; a panel of neuropeptide genes representing more than 5 percent of SCN neurons was detected alongside bcGFPs. | accepted_agree \| Platform: in situ sequencing |
| 10.1016/bs.mie.2016.03.020 | 2016 | RNA Imaging with Multiplexed Error-Robust Fluorescence In Situ Hybridization (MERFISH). | DISAGREE | Target regions were designed using transcript abundance, ENCODE RNA data, and Gencode annotations. | validated_high_risk_disagreement \| Platform: MERFISH |
| 10.1016/j.ccell.2025.02.009 | 2025 | Single-cell multi-stage spatial evolutional map of esophageal carcinogenesis | AGREE | paper_type=EXP, panel_scope=TARGETED, spatial=YES; gene panels of 300 genes were built for Xenium In Situ from pathway markers and cell-population markers. | accepted_agree \| Platform: Xenium;seqFISH |
| 10.1016/j.cell.2019.09.016 | 2019 | Optical Pooled Screens in Human Cells. | AGREE | We screened 3,063 sgRNAs targeting 963 genes using the LentiGuide-BC design and included all GO-annotated ubiquitin ligase and deubiquitinase enzymes plus 425 immune-related genes. | accepted_agree \| Platform: in situ sequencing |
| 10.1016/j.cell.2019.09.020 | 2019 | Multimodal Analysis of Cell Types in a Hypothalamic Node Controlling Social Behavior. | AGREE | We applied 58 probes comprising DEGs that marked the different VMHvl SMART-seq clusters. | accepted_agree \| Platform: seqFISH |
| 10.1016/j.cell.2019.09.023 | 2019 | High-Throughput Mapping of Long-Range Neuronal Projection Using In Situ Sequencing. | DISAGREE | BARseq was combined with FISH against Slc17a7 and Gad2 in the same cells. | validated_high_risk_disagreement \| Platform: in situ sequencing |
| 10.1016/j.cell.2019.11.025 | 2019 | A Spatiotemporal Organ-Wide Gene Expression and Cell Atlas of the Developing Human Heart. | AGREE | Half of the initial gene panel consisted of key spatial marker genes identified by ST, the other half marker genes for each scRNA-seq cluster, and additional cardiac-development genes were added. | accepted_agree \| Platform: in situ sequencing |

| DOI | Year | Title | Audit decision | Evidence (verbatim quote) | Validation status / scope flag |
|---|---|---|---|---|---|
| 10.1016/j.cell.2020.07.032 | 202 | Genome-Scale Imaging of the 3D Organization and Transcriptional Activity of Chromatin. | DISAGREE | Selected a 100-kb segment every ~3 Mb and adjusted spacing so each chromosome had at least 30 loci, yielding 1,041 target loci. | validated_high_risk_disagreement \| Platform: MERFISH |
| 10.1016/j.cell.2021.11.024 | 2021 | EASI-FISH for thick tissue defines lateral hypothalamus spatio-molecular organization. | AGREE | A combinatorial set of 24 marker-genes was selected for subsequent EASI-FISH, starting from DEGs, applying filtering criteria, manual inspection of Allen brain ISH data, and supplemented with markers from another dataset. | accepted_agree \| Platform: EASI-FISH |
| 10.1016/j.cell.2022.12.010 | 2023 | Molecular and spatial signatures of mouse brain aging at single-cell resolution | AGREE | We selected two sets of genes for MERFISH based on snRNA-seq results: cell-type markers differentially expressed between clusters and aging-related genes differentially expressed between ages, plus known marker genes curated from the literature. | accepted_agree \| Platform: MERFISH |
| 10.1016/j.cell.2024.03.013 | 2024 | Charting the cellular biogeography in colitis reveals fibroblast trajectories and coordinated spatial remodeling. | AGREE | To profile gene expression in the mouse colon, we selected a panel of 940 genes, including well-established cell-type markers and pathway genes largely compiled from KEGG. | accepted_agree \| Platform: MERFISH |
| 10.1016/j.cell.2024.10.023 | 2024 | Decoding transcriptional identity in developing human sensory neurons and organoid modeling | AGREE | The asmFISH-seq data on GW18 DRG contained 54 sensory neuron subtype marker genes identified in our snRNA-seq analysis. | accepted_agree \| Platform: seqFISH |
| 10.1016/j.cell.2025.05.047 | 2025 | A Drosophila single-cell 3D spatiotemporal multi-omics atlas unveils panoramic key regulators of cell-type differentiation | DISAGREE | q1: panel_design=MANUAL-PANEL; tool_names=NOT FOUND; uses_manual_additions=YES. | validated_high_risk_disagreement \| Platform: Other / unspecified |
| 10.1016/j.celrep.2025.115827 | 2025 | Brain-wide connectivity and novelty response of the dorsal endopiriform nucleus in mice | AGREE | A custom 500 gene panel was used, consisting of genes enriched in the CLA, EPd, PIR, and AI and cell-type markers used for clustering and differential expression analysis. | accepted_agree \| Platform: MERSCOPE |
| 10.1016/j.celrep.2025.116624 | 2025 | Spatial and single-cell transcriptomics reveal the reorganization of cerebellar microglia with aging. | AGREE | MERFISH imaging used a panel of 500 genes selected for relevance to aging, neuroinflammation, cellular stress, and cell-type specificity across brain regions. | accepted_agree \| Platform: MERFISH |

| DOI | Year | Title | Audit decision | Evidence (verbatim quote) | Validation status / scope flag |
|---|---|---|---|---|---|
| 10.1016/j.cels.2019.09.008 | 2019 | Single-Cell Analysis Reveals Regulatory Gene Expression Dynamics Leading to Lineage Commitment in Early T Cell Development. | AGREE | We used seqFISH to quantify transcripts of a curated panel of 65 regulatory and developmental state marker genes in pro-T cells. | accepted_agree \| Platform: seqFISH |
| 10.1016/j.chom.2026.01.018 | 2026 | An image-based transcriptomics atlas reveals the regional and microbiota-dependent molecular, cellular, and spatial structure of the murine gut. | AGREE | We created a MERFISH library targeting 1,922 genes, covering both cell-type markers as well as major receptor classes. | accepted_agree \| Platform: MERFISH |
| 10.1016/j.crmeth.2023.100476 | 2023 | Subcellular spatially resolved gene neighborhood networks in single cells. | DISAGREE | q1: "Twelve RNA markers were measured in HBM and HUC, and four additional RNA markers were measured in HCHs." | validated_high_risk_disagreement \| Platform: MERFISH;seqFISH |
| 10.1016/j.cub.2024.08.049 | 2024 | Three-dimensional molecular atlas highlights spatial and neurochemical complexity in the axial nerve cord of octopus arms. | AGREE | We selected specific markers to map and compare representatives from multiple neurochemical classes of neurons. | accepted_agree \| Platform: HCR-FISH |
| 10.1016/j.devcel.2022.04.016 | 2022 | Developmental landscape of human forebrain at a single-cell level identifies early waves of oligodendrogenesis. | AGREE | We targeted 50 genes that characterize the subsets of the populations identified by scRNA-seq and scATAC-seq. | accepted_agree \| Platform: in situ sequencing |
| 10.1016/j.devcel.2025.03.009 | 2025 | Transcriptional conservation and evolutionary divergence of cell types across mammalian hypothalamus development | DISAGREE | q1: panel_design=MANUAL-PANEL; tool_names=NOT FOUND; uses_manual_additions=YES. | validated_high_risk_disagreement \| Platform: HCR-FISH |
| 10.1016/j.diff.2024.100822 | 2024 | Redefining retinoic acid receptor expression in zebrafish embryos using Hybridization Chain Reaction. | AGREE | We generated HCR probes for and analyzed the expression of each rar and rxr gene transcript in embryos with either the pharyngeal endoderm or the CNCCs transgenically co-labeled. | accepted_agree \| Platform: HCR-FISH |
| 10.1016/j.ebiom.2019.09.009 | 2019 | Generation of in situ sequencing based OncoMaps to spatially resolve gene expression profiles of diagnostic and prognostic markers in breast cancer. | AGREE | In total 91 genes were selected for expression profiling in situ, including the OncotypeDX scoring, molecular subtyping, and genes involved in proliferation, DNA repair, EMT, invasiveness, stemness and angiogenesis. | accepted_agree \| Platform: in situ sequencing |
| 10.1016/j.isci.2018.12.024 | 2019 | Detection of Concordance between Transcriptional Levels of GPCRs and Receptor-Activity-Modifying Proteins. | AGREE | We used several criteria to select 14 representative GPCRs for examination in the MERFISH assay. | accepted_agree \| Platform: MERFISH |

| DOI | Year | Title | Audit decision | Evidence (verbatim quote) | Validation status / scope flag |
|---|---|---|---|---|---|
| 10.1016/j.jcmgh.2026.101730 | 2026 | Molecular and Chromatin Accessibility Programs Underlying Epithelial Injury and Impaired Regeneration in Neonatal Necrotizing Enterocolitis. | AGREE | A total of 495 probes were designed for capturing major signaling pathways or cell type genes. | accepted_agree \| Platform: MERSCOPE |
| 10.1016/j.jgg.2025.08.004 | 2025 | Spatiotemporal dynamics of neuron differentiation and migration in the developing human spinal cord. | AGREE | We focused on 129 TFs and genes, selected as top marker genes and crucial TFs of spinal cord subtypes, studied by relevant literature, from our scRNA-seq data. | accepted_agree \| Platform: in situ sequencing |
| 10.1016/j.jid.2025.04.041 | 2026 | High-Resolution Spatial Map of the Human Facial Sebaceous Gland Reveals Marker Genes and Decodes Sebocyte Differentiation. | AGREE | The gene panel for our MERFISH analyses included driver genes identified in the scRNA-seq sebocyte differentiation analysis as well as additional markers of interest. | accepted_agree \| Platform: MERFISH |
| 10.1016/j.jtos.2024.08.005 | 2024 | The effects of age and dysfunction on meibomian gland population dynamics. | AGREE | To validate the scRNA-seq findings spatially, we created a targeted gene panel based on canonical cell markers, lipid-associated genes, and differentially expressed genes in the single-cell data. | accepted_agree \| Platform: Molecular Cartography |
| 10.1016/j.lfs.2025.123776 | 2025 | Multi-omics analysis of outer membrane vesicles from P. goldsteinii in a psoriasis mouse model | AGREE | We conducted in situ sequencing on the frozen dorsal skin tissues from each group of mice, generating spatial expression maps for 16 genes. | accepted_agree \| Platform: in situ sequencing |
| 10.1016/j.neuron.2016.10.001 | 2016 | In Situ Transcription Profiling of Single Cells Reveals Spatial Organization of Cells in the Mouse Hippocampus. | AGREE | For the 125 gene set, 56 genes were selected because they showed spatially heterogeneous expression, another 44 were selected from a list of transcription factors, and 25 marker genes were selected from single-cell RNA-seq datasets. | accepted_agree \| Platform: seqFISH |
| 10.1016/j.omtm.2020.10.003 | 202 | <i>In Situ</i> Detection of Adeno-associated Viral Vector Genomes with SABER-FISH. | AGREE | q1 says SABER-FISH probes were designed in silico using ApE and lists custom oligonucleotide filtering criteria. | accepted_agree \| Platform: SABER-FISH |
| 10.1016/j.peptides.2024.171253 | 2024 | Expression patterns and behavioral effects of conopressin and APGWamide in the nudibranch Berghia stephanieae. | AGREE | q1 says the target genes were found by BLAST against a transcriptome and the probes were generated by Molecular | accepted_agree \| Platform: HCR-FISH |

| DOI | Year | Title | Audit decision | Evidence (verbatim quote) | Validation status / scope flag |
|---|---|---|---|---|---|
| | | | | Instruments or custom Python/IDT oligopools. | |
| 10.1016/j.xcrm.2024.101556 | 2024 | Transcriptomic and spatial dissection of human ex vivo right atrial tissue reveals proinflammatory microvascular changes in ischemic heart disease | AGREE | q1 says several putative marker genes were detected for identifying cell types in spatial transcriptomics on Resolve Molecular Cartography. | accepted_agree \| Platform: Molecular Cartography |
| 10.1016/j.xgen.2024.100629 | 2024 | Diversity of ribosomes at the level of rRNA variation associated with human health and disease. | AGREE | q1 says regions were selected so primers could be designed for the ES regions with abundant variants; q2 says the study designed a reverse-transcription primer for selected rRNA variant regions. | accepted_agree \| Platform: in situ sequencing |
| 10.1016/j.xgen.2025.101101 | 2026 | Transcript-guided targeted cell enrichment for scalable single-nucleus RNA sequencing. | DISAGREE | q1 says the EnrichSci workflow begins by analyzing existing snRNA-seq data to select a module of 5-10 genes specific to the target cell type. | validated_high_risk_disagreement \| Platform: HCR-FISH |
| 10.1016/j.xplc.2024.100984 | 2024 | Single-cell transcriptome atlases of soybean root and mature nodule reveal new regulatory programs that control the nodulation process. | AGREE | q1 says the authors identified highly expressed genes with Seurat V4 FindAllMarkers before probes were designed by Resolve Biosciences. | accepted_agree \| Platform: Molecular Cartography |
| 10.1016/j.ydbio.2024.05.012 | 2024 | Localization and origins of juvenile skeletogenic cells in the sea urchin Lytechinuspictus. | AGREE | q1 says six PMC-specific genes were selected for HCR-FISH and additional markers were chosen from single-cell data. | accepted_agree \| Platform: HCR-FISH |
| 10.1016/j.ydbio.2026.02.002 | 2026 | High-sensitivity in situ detection of Sox21 reveals transcript asymmetry from the zygote stage. | AGREE | q1 says Sox21 was tracked with HCR and Uhrf1/Rps6 were selected as internal controls. | accepted_agree \| Platform: HCR-FISH |
| 10.1016/j.ygcen.2026.114899 | 2026 | Multiplex in situ hybridization chain reaction reveals the spatial organization of six pituitary hormone mRNAs in the Japanese eel. | AGREE | q1 says DNA split probes were designed for six pituitary hormone mRNAs and q2 says 25-bp target regions were selected with GC filters. | accepted_agree \| Platform: HCR-FISH |
| 10.1016/j.yhbeh.2024.105578 | 2024 | Egg-laying hormone expression in identified neurons across developmental stages and reproductive states of the nudibranch Berghia stephanieae. | AGREE | q1 says a library of 30 HCR probe pairs was generated by Molecular Instruments from the complete mRNA sequence. | accepted_agree \| Platform: HCR-FISH |
| 10.1016/j.ymeth.2025.05.001 | 2025 | Optical pooled screening for the discovery of regulators of the alternative lengthening of telomeres pathway | DISAGREE | A set of sgRNAs targeted FANCM, DDX39A, and BLM, plus non-targeting and safe-targeting controls. | validated_high_risk_disagreement \| Platform: in situ sequencing |

| DOI | Year | Title | Audit decision | Evidence (verbatim quote) | Validation status / scope flag |
|---|---|---|---|---|---|
| 10.1021/acschembio.3c00431 | 2024 | Multiplex, Quantitative, High-Resolution Imaging of Protein:Protein Complexes via Hybridization Chain Reaction. | DISAGREE | Figure 4 demonstrates multiplex protein:protein imaging for three target complexes that localize to different compartments of A-431 adherent human cells. | validated_high_risk_disagreement \| Platform: HCR-FISH |
| 10.1021/cbmi.5c00220 | 2026 | Fast, Streamlined <i>In Situ</i> Genotyping of Bacterial Strains for Single-Cell and Subcellular Dynamics Profiling | DISAGREE | The 36 barcode sequences were selected from previously validated seqFISH21-compatible sequences. | validated_high_risk_disagreement \| Platform: seqFISH |
| 10.1038/nmeth.3895 | 2016 | Dense transcript profiling in single cells by image correlation decoding. | DISAGREE | We targeted transcripts for ten ribosomal proteins with corrFISH with two color binary codes. | validated_high_risk_disagreement \| Platform: seqFISH |
| 10.1038/s41396-020-00757-1 | 2021 | Experimentally-validated correlation analysis reveals new anaerobic methane oxidation partnerships with consortium-level heterogeneity in diazotrophy. | AGREE | A new FISH probe was designed in ARB ... designed to complement and target 16S rRNA sequences in a monophyletic 'Desulfococcus sp.' clade. | accepted_agree \| Platform: HCR-FISH |
| 10.1038/s41421-021-00266-1 | 2021 | Spatial transcriptome profiling by MERFISH reveals fetal liver hematopoietic stem cell niche architecture. | AGREE | we designed a MERFISH probe library that targets 45 cell-type marker genes selected from the scRNA-seq data and 95 niche-associated factors or signaling/marker genes from literature | accepted_agree \| Platform: MERFISH |
| 10.1038/s41422-023-00897-x | 2024 | Decoding the spatiotemporal regulation of transcription factors during human spinal cord development. | NEEDS_MANUAL | TF-seqFISH quantified 1085 TFs and 'developed a probe library consisting of 1085 mRNA-specific probes' | pdf_validated_unclear_update \| Platform: seqFISH |
| 10.1038/s41467-023-36137-4 | 2023 | Highly efficient and robust π-FISH rainbow for multiplexed in situ detection of diverse biomolecules. | DISAGREE | q1 states that 21 marker genes from scRNA-seq data were decoded by π-FISH rainbow. | validated_high_risk_disagreement \| Platform: HCR-FISH |
| 10.1038/s41467-023-37188-3 | 2023 | Spatial profiling of microbial communities by sequential FISH with error-robust encoding. | DISAGREE | Oligonucleotide probes were designed to target 16S or 23S rRNA using ARB, mathFISH, and DECIPHER. | validated_high_risk_disagreement \| Platform: seqFISH |
| 10.1038/s41467-023-39447-9 | 2023 | Spatial Transcriptomics-correlated Electron Microscopy maps transcriptional and ultrastructural responses to brain injury. | AGREE | Gene panel for this study consisted of 287 protein-coding genes and 98 blank probes ... known brain and immune cell type markers ... reactive markers from literature ... genes from cholesterol metabolic pathways. | accepted_agree \| Platform: MERFISH |
| 10.1038/s41467-023-43267-2 | 2023 | Fate specification is spatially intermingled across planarian stem | AGREE | Genes were selected based on known roles as FSTFs and | accepted_agree \| Platform: MERSCOPE |

| DOI | Year | Title | Audit decision | Evidence (verbatim quote) | Validation status / scope flag |
|---|---|---|---|---|---|
| | | cells. | | enrichment in differentiated tissues ... | |
| 10.1038/s41467-024-45907-7 | 2024 | A spatially-resolved transcriptional atlas of the murine dorsal pons at single-cell resolution. | AGREE | A MERSCOPE panel of 315 genes meeting at least one of the following criteria was assembled: highly variable genes obtained from DE analysis ... canonical glial, non-neuronal, and neuronal markers ... transcription factors, neuropeptides, and receptors | accepted_agree \| Platform: MERSCOPE |
| 10.1038/s41467-024-46669-y | 2024 | Highly sensitive spatial transcriptomics using FISHnCHIPs of multiple co-expressed genes | DISAGREE | q1: panel_design=MANUAL-PANEL; evidence_manual indicates multiple co-expressed genes for FISHnCHIPs. | validated_high_risk_disagreement \| Platform: Other / unspecified |
| 10.1038/s41467-024-51186-z | 2024 | Spatial transcriptomics defines injury specific microenvironments and cellular interactions in kidney regeneration and disease | AGREE | We selected 1300 target genes for seqFISH profiling based on key cell-type enriched markers from single-cell studies ... | accepted_agree \| Platform: seqFISH |
| 10.1038/s41467-024-55325-4 | 2025 | Spatial transcriptomics of healthy and fibrotic human liver at single-cell resolution. | AGREE | We designed a MERFISH panel targeting 317 genes ... supplemented with common markers of non-parenchymal liver cells | accepted_agree \| Platform: MERFISH |
| 10.1038/s41467-025-63429-8 | 2025 | Aging in mice alters regionally enriched striatal astrocytes | AGREE | We created a panel of 383 genes designed from our scRNAseq study ... known cell class markers, aging markers, and putative cell subtype markers | accepted_agree \| Platform: MERFISH |
| 10.1038/s41467-025-63467-2 | 2025 | Drought recovery in plants triggers a cell-state-specific immune activation. | AGREE | A custom-designed probe panel targeting ~1000 genes was used, including cell type cluster markers and RcS hub genes identified from snRNA-seq analyses | accepted_agree \| Platform: MERSCOPE |

| DOI | Year | Title | Audit decision | Evidence (verbatim quote) | Validation status / scope flag |
|---|---|---|---|---|---|
| 10.1038/s41467-025-63752-0 | 2025 | Molecular cartography of the human down syndrome and trisomic mouse brain. | AGREE | A meticulously chosen panel of 500 genes was used for MERFISH analysis. Among these, 132 genes served as canonical markers for identifying various cell types... The remaining genes in the panel encompassed biologically relevant signaling pathways (WNT, JAK/STAT, MAPK), ligand-receptor interactions, inflammatory markers, senescence-associated signatures, and extracellular matrix components related to DS. | manual_panel_source_correction (raw audit: VENDOR; Panel was manually curated for DS brain biology and cell-type identification; MERSCOPE is the imaging platform/kit rather than evidence of a fixed vendor gene panel.) \| Platform: MERSCOPE |
| 10.1038/s41467-025-63798-0 | 2025 | Cassini: streamlined and scalable method for in situ profiling of RNA and protein | DISAGREE | q1: panel_design=MANUAL-PANEL; evidence_manual=NOT FOUND; uses_manual_additions=YES. | validated_high_risk_disagreement \| Platform: Other / unspecified |
| 10.1038/s41467-025-64294-1 | 2025 | Tetrahedral DNA dendritic nanostructure-enhanced FISH for high-speed, sensitive spatial transcriptomics. | DISAGREE | q1 says the authors identified the top 10 marker genes for each cell type from previously reported single-cell RNA-seq datasets. | validated_high_risk_disagreement \| Platform: HCR-FISH |
| 10.1038/s41467-025-66048-5 | 2025 | A spatial atlas of the complement system uncovers unique expression patterns in postnatal brain development in mice. | AGREE | q1: 249-gene MERFISH probe library designed from canonical markers, region markers, neurotransmitter genes, neural-development/homeostasis genes, and complement genes; q1 tool/vendor evidence: not found. | accepted_agree \| Platform: MERFISH |
| 10.1038/s41467-025-66741-5 | 2025 | The ANTsX ecosystem for mapping the mouse brain. | DISAGREE | q2: "A set of 500 genes was selected to distinguish 5200 transcriptomic clusters." | validated_high_risk_disagreement \| Platform: MERFISH |
| 10.1038/s41467-026-68291-w | 2026 | Integrated multi-omic atlas reveals the hierarchy of spatiotemporal regulatory networks of mouse gastrulation | DISAGREE | q1: evidence_manual lists multiple RNAscope probes from Advanced Cell Diagnostics; panel_design=MANUAL-PANEL. | validated_high_risk_disagreement \| Platform: Other / unspecified |
| 10.1038/s41467-026-68995-z | 2026 | A spatiotemporal atlas of cerebrovascular development in zebrafish. | AGREE | q1: 36-gene panel for CapEC/AEC/VEC/AngEC states assembled from selected genes and prior marker references; q1 tool/vendor evidence: not found. | accepted_agree \| Platform: in situ sequencing |
| 10.1038/s41477-025-02176-6 | 2026 | Imputation integrates single-cell and spatial gene expression data to resolve transcriptional networks in barley shoot meristem development | AGREE | q1: panel_design=MANUAL-PANEL; tool_names=NOT FOUND. | accepted_agree \| Platform: Molecular Cartography |

| DOI | Year | Title | Audit decision | Evidence (verbatim quote) | Validation status / scope flag |
|---|---|---|---|---|---|
| 10.1038/s41559-023-02170-1 | 2023 | A lamprey neural cell type atlas illuminates the origins of the vertebrate brain. | AGREE | q1: 93 selected marker genes for ISS and four smRNA-FISH genes; no tool or vendor evidence. | accepted_agree \| Platform: in situ sequencing |
| 10.1038/s41559-024-02485-7 | 2024 | Sugar assimilation underlying dietary evolution of Neotropical bats. | AGREE | q1: probe sets target Slc5a1, Slc2a2, and Slc2a5 on bat intestinal tissue; no tool or vendor evidence. | accepted_agree \| Platform: HCR-FISH |
| 10.1038/s41586-019-1049-y | 2019 | Transcriptome-scale super-resolved imaging in tissues by RNA seqFISH. | DISAGREE | We randomly selected 10,000 genes while avoiding highly abundant housekeeping genes. | validated_high_risk_disagreement \| Platform: seqFISH |
| 10.1038/s41586-020-03126-2 | 2021 | Integrated spatial genomics reveals global architecture of single nuclei. | AGREE | q1: marker genes selected from previous single-cell imaging and RNA-seq studies in mESCs; q1 vendor/tool evidence: not found. | accepted_agree \| Platform: seqFISH |
| 10.1038/s41586-021-03705-x | 2021 | Spatially resolved cell atlas of the mouse primary motor cortex by MERFISH. | AGREE | q1: 258-gene MERFISH panel included canonical markers and genes selected by differential expression and mutual information from companion sc/snRNA-seq work. | accepted_agree \| Platform: MERFISH |
| 10.1038/s41586-022-04915-7 | 2021 | A transcriptomic axis predicts state modulation of cortical interneurons | DISAGREE | q1: 73-gene panel is a subset of a 99-gene prior panel picked using an unnamed algorithm on scRNA-seq data; q1 tool_names not found. | validated_high_risk_disagreement \| Platform: Other / unspecified |
| 10.1038/s41586-022-05425-2 | 2022 | Spatial genomics maps the structure, nature and evolution of cancer clones. | DISAGREE | q2: targeted ISS with a previously published 91-gene oncology panel and a novel 62-gene immune panel; q1 tool evidence refers only to BaSISS and layer-linking algorithms. | validated_high_risk_disagreement \| Platform: in situ sequencing |
| 10.1038/s41586-023-06569-5 | 2023 | Spatial atlas of the mouse central nervous system at molecular resolution. | AGREE | q1: 1,022 genes curated from reported marker genes and further supplemented with Allen Mouse Brain transcriptome database markers. | accepted_agree \| Platform: in situ sequencing;STARmap |
| 10.1038/s41586-023-06812-z | 2023 | A high-resolution transcriptomic and spatial atlas of cell types in the whole mouse brain. | DISAGREE | q1: select_N_markers from scrattch.bigcat chose the top 400 genes using a greedy algorithm on marker separability and significance. | validated_high_risk_disagreement \| Platform: MERSCOPE |
| 10.1038/s41586-024-07221-6 | 2025 | Whole-cortex in situ sequencing reveals input-dependent area identity. | AGREE | q1: final gene panel includes the 102-gene panel plus manually added Fezf2 and Hgf. | accepted_agree \| Platform: in situ sequencing |

| DOI | Year | Title | Audit decision | Evidence (verbatim quote) | Validation status / scope flag |
|---|---|---|---|---|---|
| 10.1038/s41586-024-08189-z | 2024 | A multi-omic atlas of human embryonic skeletal development. | AGREE | q1: probe panel design was based on fold-change thresholds in limb cell states; no tool or vendor evidence. | accepted_agree \| Platform: CARTANA |
| 10.1038/s41586-025-08838-x | 2025 | Spatial multi-omics reveals cell-type-specific nuclear compartments. | DISAGREE | Chromosomal DNA binding sequences were selected from mm10 newBalance DNA FISH probes at PaintSHOP resources. | validated_high_risk_disagreement \| Platform: seqFISH |
| 10.1038/s41586-025-09010-1 | 2025 | Spatial transcriptomics reveals human cortical layer and area specification. | AGREE | q1: initial 300-gene panel contained canonical fetal cortex markers, validated adult cortical-layer markers, SFARI genes, and manually curated top cluster markers. | accepted_agree \| Platform: MERSCOPE |
| 10.1038/s41586-025-09875-2 | 2025 | Spatiotemporal cellular map of the developing human reproductive tract | AGREE | q1: panel_design=MANUAL-PANEL; most unique marker genes were selected using tf-idf, geneBasis only evaluated completeness, and four genes were manually swapped. | accepted_agree \| Platform: CARTANA |
| 10.1038/s41587-021-01006-2 | 2022 | Integration of spatial and single-cell transcriptomic data elucidates mouse organogenesis. | AGREE | Genes were selected with scran findMarkers, then manually curated and manually added to the panel. | accepted_agree \| Platform: seqFISH |
| 10.1038/s41587-022-01455-3 | 2023 | Scalable in situ single-cell profiling by electrophoretic capture of mRNA using EEL FISH. | DISAGREE | q1: targeted 440 genes; optimized distribution of genes over available barcodes using prior knowledge from scRNA-seq; q2: TARGETED spatial COMP paper. | validated_high_risk_disagreement \| Platform: EEL FISH |
| 10.1038/s41587-022-01483-z | 2022 | High-plex imaging of RNA and proteins at subcellular resolution in fixed tissue by spatial molecular imaging. | DISAGREE | Genes for these cell states were primarily curated from the literature using hallmark genes for each set. The remaining genes were then chosen in a data-driven manner to maximize the contrast between expression profiles of different cell types. | validated_high_risk_disagreement \| Platform: CosMx |
| 10.1038/s41591-024-03073-9 | 2024 | Spatial multiomic landscape of the human placenta at molecular resolution. | AGREE | STARmap-ISS used 1,001 highly variable genes and STARmap-ISH used 48 manually curated landmark genes. | accepted_agree \| Platform: STARmap |

| DOI | Year | Title | Audit decision | Evidence (verbatim quote) | Validation status / scope flag |
|---|---|---|---|---|---|
| 10.1038/s41591-024-03215-z | 2024 | A multi-modal single-cell and spatial expression map of metastatic breast cancer biopsies across clinicopathological features. | AGREE | The packet says a dedicated panel of 297 genes was designed for MERFISH and ExSeq based on sc/snRNA-seq data and prior knowledge. | accepted_agree \| Platform: MERFISH;ExSeq |
| 10.1038/s41592-018-0095-y | 2018 | Transcript constellations in a tissue's universe | DISAGREE | The initial test targeted 160 cell-type- and neural-activity-regulated markers in adult mouse neocortex. | validated_high_risk_disagreement \| Platform: STARmap |
| 10.1038/s41592-018-0175-z | 2018 | Spatial organization of the somatosensory cortex revealed by osmFISH. | DISAGREE | The markers were chosen according to specificity and expression level to map all major cell types and most subtypes in 13 osmFISH cycles. | validated_high_risk_disagreement \| Platform: osmFISH |
| 10.1038/s41592-019-0404-0 | 2019 | SABER amplifies FISH: enhanced multiplexed imaging of RNA and DNA in cells and tissues. | DISAGREE | q1 says the study aimed to detect all seven major retinal cell classes using SABER probes against established markers; q2 says the paper is targeted spatial/in situ work. | validated_high_risk_disagreement \| Platform: SABER-FISH |
| 10.1038/s41592-019-0631-4 | 202 | Probabilistic cell typing enables fine mapping of closely related cell types in situ. | DISAGREE | q1 says the gene panel was selected by an algorithm that searches for a subset of genes and was manually curated before final selection. | validated_high_risk_disagreement \| Platform: in situ sequencing |
| 10.1038/s41592-020-0858-0 | 202 | Highly specific multiplexed RNA imaging in tissues with split-FISH. | DISAGREE | q1: randomly selected 317 genes as targets; q2: TARGETED spatial COMP paper. | validated_high_risk_disagreement \| Platform: SPLIT-FISH |
| 10.1038/s41592-020-0890-0 | 202 | 3D mapping and accelerated super-resolution imaging of the human genome using in situ sequencing. | DISAGREE | Barcodes were manually assigned to targets to maintain color-code diversity between neighboring targets. | validated_high_risk_disagreement \| Platform: in situ sequencing |
| 10.1038/s41592-023-01829-8 | 2023 | Spatiotemporally resolved transcriptomics reveals the subcellular RNA kinetic landscape. | DISAGREE | q1 says TEMPOmap mapped a curated list of 991 genes in HeLa cell cultures. | validated_high_risk_disagreement \| Platform: in situ sequencing |
| 10.1038/s41592-024-02184-y | 2024 | TISSUE: uncertainty-calibrated prediction of single-cell spatial transcriptomics improves downstream analyses. | DISAGREE | The packet says 'We curated a 140-gene panel for the MERFISH experiment.' | validated_high_risk_disagreement \| Platform: MERFISH |
| 10.1038/s41592-025-02867-0 | 2025 | Scalable spatial single-cell transcriptomics and translatomics in 3D thick tissue blocks. | DISAGREE | targeting a curated list of 1,017 genes (Supplementary Table 2). This gene list was compiled from reported cell-type marker genes in adult mouse central nervous system scRNA-seq datasets and | validated_high_risk_disagreement \| Platform: STARmap |

| DOI | Year | Title | Audit decision | Evidence (verbatim quote) | Validation status / scope flag |
|---|---|---|---|---|---|
| | | | | spatial transcriptomic mouse brain atlases | |
| 10.1038/s41593-020-0602-1 | 202 | Astrocyte layers in the mammalian cerebral cortex revealed by a single-cell in situ transcriptomic map | AGREE | The packet says the top 46 genes were selected from RNA-seq based on highest differential enrichment between cortical layers and/or astrocyte specificity. | accepted_agree \| Platform: Other / unspecified |
| 10.1038/s41593-021-00842-4 | 2021 | Integrating barcoded neuroanatomy with spatial transcriptional profiling enables identification of gene correlates of projections | DISAGREE | The authors extracted DE genes, filtered lowly expressed genes, ranked them by dataset support and fold change, and selected the top markers. | validated_high_risk_disagreement \| Platform: in situ sequencing |
| 10.1038/s41593-021-00938-x | 2021 | Decoding molecular and cellular heterogeneity of mouse nucleus accumbens | AGREE | The packet states that a panel of 253 genes was designed to transcriptionally profile distinct cell populations. | accepted_agree \| Platform: MERFISH |
| 10.1038/s41593-022-01251-x | 2023 | Integrative in situ mapping of single-cell transcriptional states and tissue histopathology in a mouse model of Alzheimer's disease. | DISAGREE | A curated list of 2,766 genes was extracted from previous bulk and single-cell RNA-seq studies and various AD-related databases. | validated_high_risk_disagreement \| Platform: STARmap |
| 10.1038/s41593-024-01764-7 | 2024 | A cross-disease resource of living human microglia identifies disease-enriched subsets and tool compounds recapitulating microglial states. | AGREE | The packet says the study designed a panel to detect cortical cell sub-types and included 39 markers for microglial subtypes. | accepted_agree \| Platform: MERSCOPE |
| 10.1038/s41593-025-02162-3 | 2026 | Spatial transcriptomics of the developing mouse brain immune landscape reveals effects of maternal immune activation and microbiome depletion | AGREE | The packet says the authors curated a panel of 500 genes for MERFISH. | accepted_agree \| Platform: MERSCOPE |
| 10.1038/s41598-018-22297-7 | 2018 | Multiplexed imaging of high-density libraries of RNAs with MERFISH and expansion microscopy. | DISAGREE | Among the 129 targeted RNAs, 106 were chosen in one abundance range and the remaining 23 spanned another range. | validated_high_risk_disagreement \| Platform: MERFISH |
| 10.1038/s41598-018-35001-6 | 2018 | Expression profiling and in situ screening of circular RNAs in human tissues. | DISAGREE | From the circRNAs identified in the frontal cortex, 129 high-level circRNAs and 127 linear targets were selected; CDR1as was added manually. | validated_high_risk_disagreement \| Platform: in situ sequencing |
| 10.1038/s41598-019-40026-6 | 2019 | A microfluidic platform towards automated multiplexed in situ sequencing. | DISAGREE | q1 says only one gene transcript, mouse beta-actin, was targeted and was chosen for stable | validated_high_risk_disagreement \| Platform: in situ sequencing |

| DOI | Year | Title | Audit decision | Evidence (verbatim quote) | Validation status / scope flag |
|---|---|---|---|---|---|
| | | | | expression and high transcript count. | |
| 10.1038/s41598-019-43943-8 | 2019 | Multiplexed detection of RNA using MERFISH and branched DNA amplification. | DISAGREE | Performed 5x5 amplification on a 130-RNA library previously measured using MERFISH. | validated_high_risk_disagreement \| Platform: MERFISH |
| 10.1038/s41598-022-11534-9 | 2022 | Direct RNA targeted in situ sequencing for transcriptomic profiling in tissue. | DISAGREE | To test the application and potential of the dRNA-HybISS, we targeted a panel of 50 genes curated based on previous publications to map cortical and hippocampal cell types. | validated_high_risk_disagreement \| Platform: in situ sequencing;CARTANA |
| 10.1038/s41598-023-36638-8 | 2023 | Reference-based cell type matching of in situ image-based spatial transcriptomics data on primary visual cortex of mouse brain | DISAGREE | q2: each experimental protocol assembled different probe panels with specific gene sets in their design (Supplementary Table S1). | validated_high_risk_disagreement \| Platform: MERFISH;ExSeq;BaristaSeq |
| 10.1038/s42003-025-08315-1 | 2025 | Spatial organization, chromatin accessibility and gene-regulatory programs defining mouse sensory neurons. | AGREE | The MERSCOPE panel included variable genes in different DRG neuron subtypes. | accepted_agree \| Platform: MERSCOPE |
| 10.1038/s44161-025-00717-y | 2025 | Spatial multiomics of acute myocardial infarction reveals immune cell infiltration through the endocardium. | AGREE | The packet says a 100-gene transcript panel was designed based on marker gene expression from scRNA-seq datasets and expert knowledge. | accepted_agree \| Platform: Molecular Cartography |
| 10.1053/j.gastro.2020.11.010 | 2021 | Single-Nucleus and In Situ RNA-Sequencing Reveal Cell Topographies in the Human Pancreas. | AGREE | We selected 98 marker genes (Supplementary Table 4) identified from sNuc-seq that distinguish pancreatic cell types, then applied ISS to tissue from 1 juvenile (1.5 years old) and 2 adult (30 and 53 years old) donors. | accepted_agree \| Platform: CARTANA |
| 10.1073/pnas.1612826113 | 2016 | High-throughput single-cell gene-expression profiling with multiplexed error-robust fluorescence in situ hybridization. | DISAGREE | Targeting regions were designed from transcriptome and abundance data using a custom probe-design pipeline with Tm, GC, and homology penalties. | validated_high_risk_disagreement \| Platform: MERFISH |
| 10.1073/pnas.1617699113 | 2016 | High-performance multiplexed fluorescence in situ hybridization in culture and tissue with matrix imprinting and clearing. | DISAGREE | The packet says the encoding probe set was designed as described previously. | validated_high_risk_disagreement \| Platform: MERFISH |
| 10.1073/pnas.1903808116 | 2019 | Imaging-based pooled CRISPR screening reveals regulators of lncRNA localization. | AGREE | (paraphrased) We selected 54 candidate genes involved in nuclear RNA regulation, including | accepted_agree \| Platform: MERFISH |

| DOI | Year | Title | Audit decision | Evidence (verbatim quote) | Validation status / scope flag |
|---|---|---|---|---|---|
| | | | | hnRNP family proteins, DExD/H box RNA helicases, and genes involved in RNA modification (Dataset S2). | |
| 10.1073/pnas.1912459116 | 2019 | Spatial transcriptome profiling by MERFISH reveals subcellular RNA compartmentalization and cell cycle-dependent gene expression. | AGREE | We then randomly selected 10,050 of these barcodes to uniquely encode 10,050 genes. | accepted_agree \| Platform: MERFISH |
| 10.1073/pnas.2309227120 | 2023 | Rapid and signal crowdedness-robust in situ sequencing through hybrid block coding. | DISAGREE | q1 says the genes included both marker and disease-related genes selected for cell classification and status characterization. | validated_high_risk_disagreement \| Platform: in situ sequencing |
| 10.1073/pnas.2313326120 | 2024 | Multiscale spatial mapping of cell populations across anatomical sites in healthy human skin and basal cell carcinoma. | AGREE | Based on published scRNAseq data (2, 17, 19, 20, 24–26) and our sequencing of facial skin, we defined a panel of 165 marker genes representing all skin cell populations (SI Appendix, Tables S2–S4). | accepted_agree \| Platform: in situ sequencing |
| 10.1073/pnas.2402514121 | 2024 | Multiplexed in situ hybridization reveals distinct lineage identities for major and minor vein initiation during maize leaf development. | AGREE | Twenty-seven genes (SI Appendix, Table S1) were selected as probes to hybridize to maize shoot apices comprising the shoot apical meristem and the first five leaf primordia. | accepted_agree \| Platform: Molecular Cartography |
| 10.1073/pnas.2421022122 | 2025 | Spatial profiling of the interplay between cell type- and vision-dependent transcriptomic programs in the visual cortex | AGREE | For these experiments, we selected a panel of 500 genes that included cortical area- and layer-specific markers, marker genes for all major cortical cell populations, and activity-regulated genes. | accepted_agree \| Platform: MERSCOPE |
| 10.1093/bib/bbaf277 | 2025 | Fusion of spatiotemporal and network models to prioritize multiscale effects in single-cell perturbations. | DISAGREE | q0: Perturb-STNet quantifies, ranks, and selects significant spatial and temporal differentially expressed regulators; q1: no canonical tool named. | validated_high_risk_disagreement \| Platform: MERFISH |
| 10.1093/bioinformatics/btad362 | 2022 | Optimization-based decoding of Imaging Spatial Transcriptomics data | DISAGREE | The sample was probed with a 20-gene library designed for microglia sub-classification and a 115-gene library designed for cell typing. | validated_high_risk_disagreement \| Platform: MERFISH |
| 10.1093/humrep/deag051 | 2026 | Spatial transcriptomics in the human adult ovary: insights into key signalling pathways during follicular atresia. | AGREE | We selected 80 genes to facilitate cell type identification in the ovary and to investigate the key signalling pathways WNT, TGFβ/BMP, NOTCH, and HH. | accepted_agree \| Platform: Molecular Cartography |

| DOI | Year | Title | Audit decision | Evidence (verbatim quote) | Validation status / scope flag |
|---|---|---|---|---|---|
| 10.1093/infdis/jiu455 | 2015 | Arginase I, polyamine, and prostaglandin E2 pathways suppress the inflammatory response and contribute to diffuse cutaneous leishmaniasis. | AGREE | The following host-specific cellular genes were quantified: ARGI, ODC, EP1, EP2, EP3, EP4, PLA2G4A, PTGS1, PTGS2, PTGES, SMS, and SRM. | accepted_agree \| Fig. 2: excluded (platform/scope) |
| 10.1093/nar/gkaa792 | 202 | Hybridization-based in situ sequencing (HybISS) for spatially resolved transcriptomics in human and mouse brain tissue. | DISAGREE | q1: gene panels were curated both manually and computationally; q2: TARGETED spatial COMP paper. | validated_high_risk_disagreement \| Platform: in situ sequencing;HybISS |
| 10.1093/nar/gkab120 | 2021 | Barcoded oligonucleotides ligated on RNA amplified for multiplexed and parallel in situ analyses. | DISAGREE | q1 states that differential gene expression analysis was used to select the 96 genes. | validated_high_risk_disagreement \| Platform: in situ sequencing |
| 10.1093/nar/gkaf461 | 2025 | Visualizing the transcription and replication of influenza A viral RNAs in cells by multiple direct RNA padlock probing and in situ sequencing (mudRapp-seq). | DISAGREE | The study visualizes 16 IAV mRNAs and vRNAs, and the PLPs were engineered with a two-nucleotide barcode. | validated_high_risk_disagreement \| Platform: in situ sequencing |
| 10.1093/plphys/kiab533 | 2022 | An in situ sequencing approach maps PLASTOCHRON1 at the boundary between indeterminate and determinate cells. | AGREE | To evaluate ISS and the simultaneous visualization of a panel of transcripts, we selected 90 genes (Supplemental Table S1) by cross-referencing in-house and published ISH patterns with tissue-specific expression levels obtained by laser microdissection of regions of the shoot apex. | accepted_agree \| Platform: CARTANA |
| 10.1097/hc9.0000000000000289 | 2023 | Defining spatiotemporal gene modules in liver regeneration using Analytical Dynamic Visual Spatial Omics Representation (ADViSOR) | DISAGREE | Genes relevant to liver function, proliferation, and inflammation were selected, and Molecular Cartography was used to capture probe images. | validated_high_risk_disagreement \| Platform: Molecular Cartography |
| 10.1098/rspb.2023.1686 | 2023 | Jaw size variation is associated with a novel craniofacial function for galanin receptor 2 in an adaptive radiation of pupfishes. | AGREE | We assayed galr2a expression in 2 dpf embryos and 8 dpf larvae from two independent lake populations for each of the three SSI species using fluorescent in-situ hybridization chain reaction (HCR) for galr2a and tropomyosin 3b (tpm3b). | accepted_agree \| Platform: HCR-FISH |
| 10.1099/jgv.0.002220 | 2026 | Remodelling of P-bodies and the cytoskeleton by <i>Orthohantavirus puumalaense</i> (Puumala virus). | AGREE | In order to undertake a comprehensive investigation of orthohantavirus infections by FISH, probe sets were designed to detect all three genomic RNA segments as well as the corresponding viral | accepted_agree \| Platform: seqFISH |

| DOI | Year | Title | Audit decision | Evidence (verbatim quote) | Validation status / scope flag |
|---|---|---|---|---|---|
| | | | | mRNAs. / Based on this rationale, we have designed a novel set of FISH probes... | |
| 10.1101/183020 | 2017 | High throughput expression profiling and in situ screening of circular RNAs in tissues | AGREE | From the circRNAs identified in the frontal cortex, we selected 129 circRNAs expressed at high levels as well as 127 linear RNA targets from the same genes for the design of the circRNA AmpliSeq panel. / The only exception ... CDR1as ... was added manually... | accepted_agree \| Platform: in situ sequencing |
| 10.1101/2020.02.03.931618 | 202 | Hybridization-based In Situ Sequencing (HybISS): spatial transcriptomic detection in human and mouse brain tissue | DISAGREE | q1: gene panels were curated both manually and computationally; q2: TARGETED spatial COMP paper. | validated_high_risk_disagreement \| Platform: in situ sequencing;HybISS |
| 10.1101/2020.02.07.938571 | 202 | SCRINSHOT, a spatial method for single-cell resolution mapping of cell states in tissue sections | DISAGREE | q1: used a panel of selective markers for club, goblet, basal, tuft and ionocytes; q2: TARGETED spatial COMP paper. | validated_high_risk_disagreement \| Platform: SCRINSHOT |
| 10.1101/2020.05.13.094268 | 202 | Expansion Sequencing: Spatially Precise<i>In Situ</i>Transcriptomics in Intact Biological Systems | DISAGREE | Initial marker genes were selected from literature and reference data; mfishtools is not one of the canonical panel-selection tools, so this is manual-panel rather than tool-based. | corrected_high_risk_disagreement_noncanonical_tool \| Platform: in situ sequencing;ExSeq |
| 10.1101/2020.06.04.105700 | 202 | Molecular, spatial and projection diversity of neurons in primary motor cortex revealed by in situ single-cell transcriptomics | AGREE | We selected a panel of 258 genes for MERFISH imaging ... by combining ... canonical marker genes ... differential gene expression analysis ... highest mutual information ... | accepted_agree \| Platform: MERFISH |
| 10.1101/2020.08.25.266460 | 202 | Integrating barcoded neuroanatomy with spatial transcriptional profiling reveals cadherin correlates of projections shared across the cortex | DISAGREE | The study extracted DE genes from multiple scRNA-seq datasets, ranked them, and selected the top markers for the panel. | validated_high_risk_disagreement \| Platform: in situ sequencing |
| 10.1101/2020.11.20.391896 | 202 | Highly multiplexed spatially resolved gene expression profiling of mouse organogenesis | AGREE | we used the scran function findMarkers ... Additionally, for each cell type, the panel of genes was manually curated ... Finally ... we manually added genes of interest... | accepted_agree \| Platform: seqFISH |
| 10.1101/2020.11.29.403055 | 202 | Global architecture of the nucleus in single cells by DNA seqFISH+ | AGREE | To select target regions ... candidate target regions ... masking coverage ... space out | accepted_agree \| Platform: seqFISH |

| DOI | Year | Title | Audit decision | Evidence (verbatim quote) | Validation status / scope flag |
|---|---|---|---|---|---|
| | | and multiplexed immunofluorescence | | approximately 2 Mb ... candidate genes related to mESCs pluripotency and differentiation were selected from previous studies... | |
| 10.1101/2020.12.02.408781 | 202 | Direct RNA targeted transcriptomic profiling in tissue using Hybridization-based RNA In Situ Sequencing (HybRISS) | DISAGREE | Genes were selected based on previous publications to delineate cell types in adult mouse brain sections. | validated_high_risk_disagreement \| Platform: in situ sequencing;HybISS |
| 10.1101/2020.12.07.414060 | 202 | Spatial organization of the transcriptome in individual neurons | AGREE | we selected 950 genes related to neuron projections, neurite development and synapses. We also included well-characterized cell type markers... | accepted_agree \| Platform: MERFISH |
| 10.1101/2021.02.24.432792 | 2021 | In situ single-cell activities of microbial populations revealed by spatial transcriptomics | AGREE | We generated a probe library targeting a set of 105 marker genes ... These included genes involved in biosynthetic capacity, anaerobic physiology, stress responses... | accepted_agree \| Platform: seqFISH |
| 10.1101/2021.06.04.447052 | 2021 | Spatial cell type mapping of the oligodendrocyte lineage in the mouse juvenile and adult CNS with in situ sequencing | AGREE | our targeted genes were selected from single-cell RNA sequencing data ... based on median expression values to target major markers... | accepted_agree \| Platform: in situ sequencing |
| 10.1101/2021.06.22.449468 | 2021 | Spatial transcriptomics using combinatorial fluorescence spectral and lifetime encoding, imaging and analysis | DISAGREE | q1: identified 10 genes from bulk sequencing data; genes chosen for housekeeping status or tumorigenesis; q2: TARGETED spatial COMP paper. | validated_high_risk_disagreement \| Platform: Other / unspecified |
| 10.1101/2021.07.17.452808 | 2021 | Decoding molecular and cellular heterogeneity of nucleus accumbens with high-throughput scRNA-seq and MERFISH | AGREE | we selected a panel of 253 gene targets ... including marker genes ... as well as genes with strong functional implications ... the remaining genes using ... mutual information (MI) analysis and ... differentially expressed (DE) gene analysis. | accepted_agree \| Platform: MERFISH |
| 10.1101/2021.08.11.455959 | 2021 | Multiplexed detection of SARS-CoV-2 genomic and subgenomic RNA using <i>in situ</i> hybridization | DISAGREE | We designed RNA FISH HCR probes using RajLab ProbeDesignHD software. / We then manually designed 52-mer oligos ... | validated_high_risk_disagreement \| Platform: HCR-FISH |
| 10.1101/2021.11.01.466826 | 2021 | Conservation and divergence in cortical cellular organization | AGREE | we use a combination of ... mutual information (MI) analysis and | accepted_agree \| Platform: MERFISH |

| DOI | Year | Title | Audit decision | Evidence (verbatim quote) | Validation status / scope flag |
|---|---|---|---|---|---|
| | | between human and mouse revealed by single-cell transcriptome imaging | | differentially expressed (DE) gene analysis ... resulting in ... marker gene candidates ... adding the manually and computationally selected genes provided by the Chan Zuckerberg Initiative SpaceTx Consortium... | |
| 10.1101/2021.11.03.467020 | 2021 | High-plex Multiomic Analysis in FFPE at Subcellular Level by Spatial Molecular Imaging | DISAGREE | Genes for these cell states were primarily curated from the literature. The remaining genes were then chosen in a data-driven manner to maximize the contrast between different cell type's expression profiles. | validated_high_risk_disagreement \| Platform: CosMx |
| 10.1101/2022.01.12.476082 | 2022 | Scalable <i>in situ</i> single-cell profiling by electrophoretic capture of mRNA | DISAGREE | q1: optimized distribution of genes over available barcodes using prior knowledge from single-cell RNA-seq; q2: TARGETED spatial COMP paper. | validated_high_risk_disagreement \| Platform: EEL FISH |
| 10.1101/2022.01.14.476072 | 2022 | Integrative <i>in situ</i> mapping of single-cell transcriptional states and tissue histopathology in an Alzheimer’s disease model | AGREE | A curated list of 2,766 genes were extracted from previous bulk and single-cell RNA-seq studies and diverse AD-related databases. | accepted_agree \| Platform: STARmap |
| 10.1101/2022.02.28.482278 | 2022 | Reconstructing the ancestral vertebrate brain using a lamprey neural cell type atlas | AGREE | we generated in situ sequencing (ISS) data for 93 selected marker genes ... and single-molecule RNA-FISH (smFISH) images for four genes... | accepted_agree \| Platform: CARTANA |
| 10.1101/2022.05.18.492475 | 2022 | Spatial Transcriptomics-correlated Electron Microscopy | AGREE | Genes included selection of known brain and immune cell type markers ... we included a panel of microglial, astrocytic and oligodendrocytic reactive markers from literature, and genes from cholesterol metabolic pathways. | accepted_agree \| Platform: MERFISH |
| 10.1101/2022.07.31.502046 | 2022 | Expansion Sequencing of RNA Barcoded Neurons in the Mammalian Brain: Progress and Implications for Molecularly Annotated Connectomics | DISAGREE | Tissues were then immunostained with antibodies targeted to the synaptic proteins Bassoon and Homer1, as well as to the virally-introduced GFP. | validated_high_risk_disagreement \| Platform: ExSeq |

| DOI | Year | Title | Audit decision | Evidence (verbatim quote) | Validation status / scope flag |
|---|---|---|---|---|---|
| 10.1101/2022.09.23.509133 | 2022 | Transcriptionally defined morphological subtypes of pancreatic ductal adenocarcinoma | AGREE | processed them for ISS with a curated list of 199 target transcripts ... Additional targets based on their reported or postulated role in PDAC development | accepted_agree \| Platform: CARTANA |
| 10.1101/2022.10.16.512401 | 2022 | Improved<i>in situ</i>sequencing for high-resolution targeted spatial transcriptomic analysis in tissue sections | DISAGREE | q1: genes selected manually based on existing literature; q2: TARGETED spatial COMP paper. | validated_high_risk_disagreement \| Platform: in situ sequencing |
| 10.1101/2023.03.06.531348 | 2023 | A molecularly defined and spatially resolved cell atlas of the whole mouse brain | AGREE | Our MERFISH gene panel was selected from marker genes differentially expressed between these subclasses and clusters ... we first included 123 subclass markers ... then added DE genes ... | accepted_agree \| Platform: MERFISH |
| 10.1101/2023.04.16.536509 | 2023 | Single-cell DNA Methylome and 3D Multi-omic Atlas of the Adult Mouse Brain | AGREE | selected genes based on cell type and spatial diversity ... mutual information ... pairwise differentially methylated gene analysis ... We selected 500 final genes as the gene panel based on rank in the RNA list... | accepted_agree \| Platform: MERFISH |
| 10.1101/2023.05.07.539762 | 2023 | High-resolution spatial multi-omics reveals cell-type specific nuclear compartments | DISAGREE | Chromosomal DNA binding sequences were selected from mm10 newBalance DNA FISH probes at PaintSHOP resources. | validated_high_risk_disagreement \| Platform: seqFISH |
| 10.1101/2023.08.08.551504 | 2023 | Multi-scale spatial mapping of cell populations across anatomical sites in healthy human skin and basal cell carcinoma | AGREE | Based on published scRNAseq data and our sequencing of facial skin, we defined a panel of 165 marker genes representing all skin cell populations. | accepted_agree \| Platform: in situ sequencing |
| 10.1101/2023.09.29.560114 | 2023 | Cell type-specific gene expression dynamics during human brain maturation | DISAGREE | Probes were ordered for RELN, FABP7, AQP4, RORB, CLSTN2, and TSHZ2. | validated_high_risk_disagreement \| Fig. 2: excluded (platform/scope) |
| 10.1101/2023.11.22.568325 | 2023 | Spatial and temporal profiling of the complement system uncovered novel functions of the alternative complement pathway in brain development | AGREE | We designed a 249-gene MERFISH probe library comprising canonical markers, cortical layer and region markers, neurotransmission genes, neurodevelopment/homeostasis genes, and complement genes. | accepted_agree \| Platform: MERFISH |

| DOI | Year | Title | Audit decision | Evidence (verbatim quote) | Validation status / scope flag |
|---|---|---|---|---|---|
| 10.1101/2024.01.27.575940 | 2024 | Re-activation of neurogenic niches in aging brain | AGREE | We implemented MERFISH with a gene list focused on determining the cell identities of neurons, glia, and cell intermediates appearing during canonical adult neurogenesis. | accepted_agree \| Platform: MERFISH |
| 10.1101/2024.02.05.578898 | 2024 | Multiplexed<i>in situ</i>hybridization reveals distinct lineage identities for major and minor vein initiation during maize leaf development | AGREE | We sought to generate a spatial expression atlas in maize shoot apices for genes in which an aberrant leaf venation phenotype has been described previously in either monocots or dicots. | accepted_agree \| Platform: Molecular Cartography |
| 10.1101/2024.03.29.587380 | 2024 | Control of behavioral uncertainty by divergent frontal circuits | AGREE | Probes for the DE genes of EGFP or tdTomato were selected according to the data of 10x single-cell sequencing. Probes for the layer specific genes were selected based on previous studies. | accepted_agree \| Platform: MERSCOPE |
| 10.1101/2024.04.14.589438 | 2024 | Three-Dimensional Molecular Atlas of Octopus Arm Neuroanatomy Highlights Spatial and Functional Complexity | AGREE | We selected these markers to identify representatives of key functional classes of neurons. | accepted_agree \| Platform: HCR-FISH |
| 10.1101/2024.05.17.594641 | 2024 | Deep-tissue spatial omics: imaging whole-embryo transcriptomics and subcellular structures at high spatial resolution | DISAGREE | the paper says it 'developed a probe library' and generated 254-gene / 120-gene libraries | pdf_validated_unclear_update \| Platform: HCR-FISH |
| 10.1101/2024.05.20.594955 | 2024 | Spatial omics of acute myocardial infarction reveals a novel mode of immune cell infiltration | AGREE | Based on marker gene expression from existing publicly available single-cell RNA-seq datasets and expert knowledge, we designed a 100 gene transcript panel specifically for this study. | accepted_agree \| Platform: Molecular Cartography |
| 10.1101/2024.06.05.597673 | 2024 | Spatial Single-cell Analysis Decodes Cortical Layer and Area Specification | AGREE | We curated a panel of three hundred genes for MERFISH analysis. | accepted_agree \| Platform: MERFISH |
| 10.1101/2024.07.10.602965 | 2024 | A multiomic atlas of human early skeletal development | AGREE | Probe panel design was based on fold-change thresholds in cell states of the limbs. | accepted_agree \| Platform: CARTANA |
| 10.1101/2024.07.16.603662 | 2024 | P-BODY AND CYTOSKELETON REMODELING BY ORTHOHANTAVIRUSES | AGREE | We systematically designed probes for all three genomic viral RNA segments as well as for the respective virus protein coding messenger RNA sequences. | accepted_agree \| Platform: seqFISH |

| DOI | Year | Title | Audit decision | Evidence (verbatim quote) | Validation status / scope flag |
|---|---|---|---|---|---|
| 10.1101/2024.10.15.618479 | 2024 | Interactive design and validation of antibody panels using single-cell RNA-seq atlases | DISAGREE | q1: 'We introduce Cytomarker, an interactive platform enabling human-in-the-loop design of antibody panels from single-cell transcriptomic data'; tool_names=Cytomarker. | validated_high_risk_disagreement \| Platform: Other / unspecified |
| 10.1101/2024.10.28.620681 | 2024 | Spatial Transcriptomics Reveals the Temporal Architecture of the Seminiferous Epithelial Cycle and Precise Sertoli-Germ Synchronization | AGREE | An initial panel of mouse transcription factors and signaling pathway genes (total 3073 genes) was selected from the Kegg Pathways Database. | accepted_agree \| Platform: seqFISH |
| 10.1101/2024.11.11.623014 | 2024 | Single-cell spatial transcriptomics of fixed, paraffin-embedded biopsies reveals colitis-associated cell networks | AGREE | Both panels were designed to detect colon-associated mucosal and submucosal cell subsets and differentially expressed genes identified by single-cell RNA sequencing of colon biopsies. | accepted_agree \| Platform: Xenium;MERSCOPE |
| 10.1101/2024.11.13.623440 | 2024 | An integrative layer-resolved atlas of the adult human meninges | DISAGREE | We created a three hundred gene panel using the single cell RNA sequencing data described previously. | validated_high_risk_disagreement \| Platform: MERSCOPE |
| 10.1101/2024.12.30.630829 | 2024 | Discovery of neuronal cell types by pairing whole cell reconstructions with RNA expression profiles | AGREE | q1: user-reviewed linked bioRxiv full text; relevant paper, manual selection; tool evidence not found. | accepted_agree \| Platform: EASI-FISH |
| 10.1101/2025.01.09.632272 | 2025 | Guidestar: a spike-in approach to improve RNA detection accuracy in imaging-based spatial transcriptomics | DISAGREE | q1: integrated library combining a 133-gene MERFISH panel with Guidestar probes; q2: targeted panel context. | validated_high_risk_disagreement \| Platform: MERFISH |
| 10.1101/2025.02.10.637514 | 2025 | SpaceBar enables clone tracing in spatial transcriptomic data | AGREE | q1: panel targeted both the barcode library and genes with interesting expression patterns in melanoma; melanoma portion curated from literature and RNA expression data. | accepted_agree \| Platform: seqFISH |
| 10.1101/2025.03.01.640978 | 2025 | Spatial and molecular insights into microglial roles in cerebellar aging | AGREE | q1: MERFISH imaging used a panel of 500 genes selected for relevance to aging, neuroinflammation, cellular stress, and cell-type specificity across brain regions. | accepted_agree \| Platform: MERFISH |

| DOI | Year | Title | Audit decision | Evidence (verbatim quote) | Validation status / scope flag |
|---|---|---|---|---|---|
| 10.1101/2025.03.27.641500 | 2025 | A factor-based analysis of individual human microglia uncovers regulators of an Alzheimer-related transcriptional signature | AGREE | q1: custom gene panel of 412 protein-coding genes with 164 added probes for regulators and targets; no vendor panel. | accepted_agree \| Platform: MERSCOPE |
| 10.1101/2025.03.27.645744 | 2025 | Uncovering the dark transcriptome in polarized neuronal compartments with mcDETECT | DISAGREE | q1: designed a 290-gene panel tailored for profiling distal RNA granules, comprising synaptic, dendritic, and axonal markers; q2: targeted spatial paper with panel-design evidence. | validated_high_risk_disagreement \| Platform: MERSCOPE |
| 10.1101/2025.04.17.647851 | 2025 | Molecular and Epigenetic Pathways Underlying Epithelial Damage and Repair in Necrotizing Enterocolitis via Multi-omics Approach | AGREE | q1: 500-gene panel targeting NEC pathways and cell-type genes; established marker genes and functional genes were selected. | accepted_agree \| Platform: MERSCOPE |
| 10.1101/2025.05.02.651832 | 2025 | Molecular and Spatial Organization of the Primary Olfactory System and its Responses to Social Odors | AGREE | q1: MERFISH probes were designed to preferentially target variable UTRs for the large odorant receptor repertoire. | accepted_agree \| Platform: MERFISH |
| 10.1101/2025.05.09.653118 | 2025 | In situ profiling of plasma cell clonality with image-based single-cell transcriptomics | NEEDS_MANUAL | "similarity-tree-based representation of relative homology to guide probe design"; "designed a 589-gene MERFISH library" | pdf_validated_unclear_update \| Platform: MERFISH |
| 10.1101/2025.05.23.655666 | 2025 | A Multimodal Spatial and Epigenomic Atlas of Human Adult Lung Topography | AGREE | q1: selection based on marker genes, ligands/receptors/TFs, ECM genes, and genes in distributed UMAP subdomains; no tool or vendor panel. | accepted_agree \| Platform: MERSCOPE |
| 10.1101/2025.06.11.659209 | 2025 | Logistic regression for estimating functional effects with spatial transcriptomics | DISAGREE | q2: "hybridizing with our custom 721-gene MERSCOPE gene panel" and q1: transcript counts for RORβ and five other layer-specific genes. | validated_high_risk_disagreement \| Platform: MERFISH |
| 10.1101/2025.07.01.662364 | 2025 | An Intrinsic-hoc Framework for Heterogeneous Cellular Senescence Elucidation Using Deep Graph Representation Learning and Experimental Validation | DISAGREE | The custom panel was curated based on human lung single-cell analysis data generated by the Lung Aging Lab. | validated_high_risk_disagreement \| Platform: Xenium |
| 10.1101/2025.07.16.665048 | 2025 | Barcoded Rabies In Situ Connectomics for high-throughput reconstruction of neural circuits | DISAGREE | q1 tool evidence: a modified greedy combinatorial search algorithm selected the top 80 genes; q1 manual evidence: | validated_high_risk_disagreement \| Platform: in situ sequencing |

| DOI | Year | Title | Audit decision | Evidence (verbatim quote) | Validation status / scope flag |
|---|---|---|---|---|---|
| | | | | sixteen additional genes were manually added; no canonical tool was named. | |
| 10.1101/2025.07.22.665939 | 2025 | Mapping mesenchymal diversity in the developing human intestine and organoids | AGREE | q1 manual evidence: custom fetal human small intestine Xenium probe panel designed based on conventional markers and previously acquired/published sequencing data; q1 tool/vendor evidence: NOT FOUND. | accepted_agree \| Platform: Xenium |
| 10.1101/2025.07.25.666577 | 2025 | Identification, Purification and Characterization of Mast Cells in Murine Liver Fibrosis: Novel Methods, Expression Signatures and Correlation with Disease Severity | AGREE | q1 manual evidence: complete list of target genes with Resolve Biosciences catalogue numbers is in Table 2; no tool evidence and no explicit vendor-panel selection is described. | accepted_agree \| Platform: Molecular Cartography |
| 10.1101/2025.08.09.25333297 | 2025 | Linking spatial omics to patient phenotypes at the population scale by BSNMani: Bayesian scalar-on-network regression with manifold learning | DISAGREE | q2: "A custom-designed panel of 140 genes was used to capture spatially resolved gene expression at single-cell resolution." | validated_high_risk_disagreement \| Platform: MERFISH |
| 10.1101/2025.08.11.669542 | 2025 | RESCUE: recovery of idiosyncratic expression patterns in spatial transcriptomics | DISAGREE | Using a custom MERFISH platform, the study generated spatial transcriptomics data targeting 130 genes relevant to aggression. | validated_high_risk_disagreement \| Platform: MERFISH |
| 10.1101/2025.08.14.670195 | 2025 | Macrophages drive a fibrogenic gene program of periductal fibroblasts in pediatric primary sclerosing cholangitis | AGREE | q1 manual evidence: a custom 400-gene liver immunology panel was used on the VizGen MERSCOPE platform; q1 tool evidence: NOT FOUND. | accepted_agree \| Platform: MERSCOPE |
| 10.1101/2025.08.19.670635 | 2025 | A Spatially Coordinated Keratinocyte-Fibroblast Circuit Recruits MMP9<sup>+</sup> Myeloid Cells to Drive IFN-I-Driven Inflammation in Photosensitive Autoimmunity | AGREE | q1 manual evidence: canonical markers, prior markers, cytokines/receptors, differentially expressed genes from prior studies, housekeeping genes, and in-house differential expression analyses; no canonical tool named. | accepted_agree \| Platform: seqFISH |

| DOI | Year | Title | Audit decision | Evidence (verbatim quote) | Validation status / scope flag |
|---|---|---|---|---|---|
| 10.1101/2025.09.13.676059 | 2025 | Aberrant host mRNA partitioning in Ebola virus condensates driven by RNA folding perpetuates species-dependent interferon response | DISAGREE | q1 evidence mentions 136 genes evaluated using the MERSCOPE Gene Panel Design Portal to check length and abundance constraints; q1 uses_vendor_panel=YES, but no explicit vendor panel source is named. | validated_high_risk_disagreement \| Platform: MERSCOPE |
| 10.1101/2025.09.22.676865 | 2025 | A single-cell spatial transcriptomic census of human skin anatomy | AGREE | q1 manual evidence: two 500-gene MERFISH panels composed of canonical markers and scRNA-seq-derived cell-type markers, plus ligand-receptor pairs; no tool evidence. | accepted_agree \| Platform: MERFISH |
| 10.1101/2025.09.25.678618 | 2025 | High resolution spatial profiling of the hematopoietic landscape of the murine lung | AGREE | q1 manual evidence: preliminary gene panel generated using cell-type specific markers, growth factors, cytokines, chemokines, and regulatory factors, then refined by abundance and specificity filters; no canonical tool named. | accepted_agree \| Platform: MERSCOPE |
| 10.1101/2025.10.09.681318 | 2025 | A Physically-Realistic Simulator and Cosine-Based Decoder for MERFISH Spatial Transcriptomics | DISAGREE | The final 140-gene MERFISH library was manually curated from vendor panel sources, immune gene lists, and RNA-seq upregulated genes. | validated_high_risk_disagreement \| Platform: MERFISH |
| 10.1101/2025.10.10.681751 | 2025 | GPU-accelerated, self-optimizing processing for 3D multiplexed iterative RNA-FISH experiments | NEEDS_MANUAL | "2380 MERFISH encoding probes were designed using PaintSHOP ... targeting 119 genes" | pdf_validated_unclear_update \| Platform: MERFISH |
| 10.1101/2025.10.16.682795 | 2025 | iss-nf: A Nextflow-based end-to-end <i>in situ</i> sequencing decoding workflow | DISAGREE | We designed a target gene panel (n = 242, Supplementary Table 1) with the company CARTANA, capturing relevant cell types, phenotypes and genes previously reported to be involved in disease onset, progression and therapy resistance. | manual_panel_source_correction (raw audit: VENDOR-MANUAL; Panel was manually selected with CARTANA support; company involvement is platform/design assistance rather than a fixed vendor panel or vendor panel plus additions.) \| Platform: CARTANA |
| 10.1101/2025.10.31.685953 | 2025 | A molecular map of the human spinal dorsal and ventral horn defines arrangement of neuronal types and glial sex differences | AGREE | q1 manual evidence: 480-custom gene Xenium spatial transcriptomics panel developed using the top expressing markers of each major cell type from snSeq; no tool or vendor panel evidence found. | accepted_agree \| Platform: Xenium |
| 10.1101/383943 | 2018 | Pooled optical screens in human cells | DISAGREE | The screen targeted 952 genes spanning NF-kB pathway | validated_high_risk_disagreement \| Platform: in situ sequencing |

| DOI | Year | Title | Audit decision | Evidence (verbatim quote) | Validation status / scope flag |
|---|---|---|---|---|---|
| | | | | components and GO-annotated ubiquitin ligase/deubiquitination genes. | |
| 10.1101/431957 | 2018 | A spatial atlas of inhibitory cell types in mouse hippocampus | DISAGREE | An algorithm searched for subset genes, then the final gene sets were chosen by manual curation of those lists. | validated_high_risk_disagreement \| Platform: in situ sequencing |
| 10.1101/432104 | 2018 | Single-cell in situ transcriptomic map of astrocyte cortical layer diversity | AGREE | We selected the top 46 genes from RNAseq that showed the highest differential enrichment between cortical layers and/or astrocyte specificity. | accepted_agree \| Platform: Other / unspecified |
| 10.1101/719930 | 2019 | Spatial and temporal transcriptomics reveal microglia-astroglia crosstalk in the amyloid-β plaque cell niche of Alzheimer's disease | AGREE | We used customized probes to map PIGs together with cell type markers such as Itgam, Slc1a3, Syp, and Plp1. | accepted_agree \| Platform: CARTANA |
| 10.1101/798512 | 2019 | Measuring chromosome conformation by fluorescence microscopy | DISAGREE | Fluorescently labeled 10 genomic loci over a 4 Mb stretch of chromosome 4 were chosen for the reconstruction experiment. | validated_high_risk_disagreement \| Platform: seqFISH |
| 10.1101/gr.261503.120 | 202 | Pooled protein tagging, cellular imaging, and in situ sequencing for monitoring drug action in real time. | AGREE | To design the intron-targeting sgRNA library, the authors started with 2889 genes from published metabolic enzyme lists, CRISPR library genes, and GO terms for histone modification and DNA methylation. | accepted_agree \| Platform: in situ sequencing |
| 10.1101/gr.278717.123 | 2024 | Single-cell spatial transcriptomics reveals a dystrophic trajectory following a developmental bifurcation of myoblast cell fates in facioscapulohumeral muscular dystrophy. | AGREE | The study performed MERFISH using a custom-designed 140-gene probe set including 24 previously identified DUX4 target genes. | accepted_agree \| Platform: MERFISH |
| 10.1101/gr.280637.125 | 2025 | Optimal marker genes for <i>c</i> -separated cell types with SepSolve | DISAGREE | SepSolve is not one of the canonical review tools; the title supports marker-gene/panel-selection relevance but not canonical TOOL classification. | validated_medium_risk_disagreement \| Fig. 2: excluded (platform/scope) |
| 10.1111/ede.12283 | 2019 | Novel approach to quantitative spatial gene expression uncovers genetic stochasticity in the developing Drosophila eye. | AGREE | Within the morphogenetic furrow, the study focuses on four genes: hairy, atonal, hedgehog, and Delta. | accepted_agree \| Platform: HCR-FISH |

| DOI | Year | Title | Audit decision | Evidence (verbatim quote) | Validation status / scope flag |
|---|---|---|---|---|---|
| 10.1111/jne.13429 | 2024 | Expression of novel androgen receptors in three GnRH neuron subtypes in the cichlid brain. | DISAGREE | We used a 50 gene custom panel specifically designed for A. burtoni. | validated_high_risk_disagreement \| Platform: Xenium |
| 10.1126/sciadv.abk0133 | 2022 | Computer-aided design of reversible hybridization chain reaction (CAD-HCR) enables multiplexed single-cell spatial proteomics imaging. | DISAGREE | q1 lists the proteins targeted in the assay, including Ezrin, beta-actin, alpha-tubulin, GM130, VDAC, lamin B1, KDM1/LSD1, SIRT1, and FBLL1. | validated_high_risk_disagreement \| Platform: HCR-FISH |
| 10.1126/sciadv.abn4965 | 2022 | A reference tissue atlas for the human kidney | AGREE | The CODEX data used 31 features: size plus 30 protein markers such as AQP1, CD34, THP, CK7, and CD104. | accepted_agree \| Platform: Other / unspecified |
| 10.1126/sciadv.ade9909 | 2023 | A single-cell resolution gene expression atlas of the larval zebrafish brain. | AGREE | We acquired FISH data for 290 genes that were previously identified as distinct, often cell type-specific markers. | accepted_agree \| Platform: HCR-FISH |
| 10.1126/sciadv.adr4026 | 2025 | Boosting multiplexing capabilities for error-robust spatial transcriptomic methods using a set exchange approach. | DISAGREE | q1: "we developed an R pipeline that evaluates alternative starting codebooks ... and iteratively exchanges sets between the codebook and the unused pool to improve the codebook size"; q2: targeted MERFISH panel scope. | validated_high_risk_disagreement \| Platform: MERFISH |
| 10.1126/science.aaa6090 | 2015 | RNA imaging. Spatially resolved, highly multiplexed RNA profiling in single cells. | DISAGREE | Central targeting sequences were designed from transcript abundance, ENCODE RNA data, and Gencode annotations. | validated_high_risk_disagreement \| Platform: MERFISH |
| 10.1126/science.aat5691 | 2018 | Three-dimensional intact-tissue sequencing of single-cell transcriptional states. | DISAGREE | We designed a panel of probes targeting 42 genes (tables S9 and S10) that mark key excitatory and inhibitory neuron types. | validated_high_risk_disagreement \| Platform: MERFISH;in situ sequencing;seqFISH;STARmap |
| 10.1126/science.aau5324 | 2018 | Molecular, spatial, and functional single-cell profiling of the hypothalamic preoptic region | AGREE | The gene panel consisted of 85 preselected genes plus 70 additional genes identified with scRNA-seq as neuronal cluster markers. | accepted_agree \| Platform: MERFISH |
| 10.1126/science.aax2656 | 2021 | Expansion sequencing: Spatially precise in situ transcriptomics in intact biological systems. | DISAGREE | We designed a panel of probes targeting 42 genes (tables S9 and S10) that mark key excitatory and inhibitory neuron types. | validated_high_risk_disagreement \| Platform: in situ sequencing;ExSeq |
| 10.1126/science.abd5059 | 202 | Cerebellar nuclei evolved by repeatedly duplicating a conserved cell-type set. | AGREE | Up to 20 marker genes were chosen to distinguish all cell types within each nucleus on coronal | accepted_agree \| Platform: STARmap |

| DOI | Year | Title | Audit decision | Evidence (verbatim quote) | Validation status / scope flag |
|---|---|---|---|---|---|
| | | | | sections spanning the anterior-posterior axis. | |
| 10.1126/science.abi4882 | 2021 | Spatial transcriptomics of planktonic and sessile bacterial populations at single-cell resolution. | AGREE | The study generated a probe library targeting 105 marker genes that capture core physiological aspects of the pathogen. | accepted_agree \| Platform: seqFISH |
| 10.1126/science.abm1741 | 2022 | Conservation and divergence of cortical cell organization in human and mouse revealed by MERFISH. | AGREE | The genes included 764 differentially expressed marker genes from single-nucleus data and additional expressed genes selected to increase gene coverage. | accepted_agree \| Platform: MERFISH |
| 10.1126/science.adf9941 | 2023 | Spatiotemporal molecular dynamics of the developing human thalamus | AGREE | The gene panel was designed using cluster markers from second-trimester scRNA-seq and snRNA-seq data for a 140-gene MERSCOPE panel. | accepted_agree \| Platform: MERSCOPE |
| 10.1126/science.adr0932 | 2025 | Highly multiplexed spatial transcriptomics in bacteria. | DISAGREE | We designed a MERFISH panel covering importers, HTCS genes, and other bacterial functions, targeting 159 operons in total. | validated_high_risk_disagreement \| Platform: MERFISH |
| 10.1126/science.adt3439 | 2026 | Whole-embryo spatial transcriptomics at subcellular resolution from gastrulation to organogenesis | AGREE | q1: panel_design=MANUAL-PANEL; tool_names=NOT FOUND. | accepted_agree \| Platform: Other / unspecified |
| 10.1126/science.adx3800 | 2025 | High-resolution spatial mapping of cell state and lineage dynamics in vivo with PEtracer. | AGREE | The 124-gene MERFISH panel was assembled from differentially expressed genes and manually selected canonical cell type markers. | accepted_agree \| Platform: MERFISH |
| 10.1128/jb.00685-19 | 202 | Single-Cell Analysis Reveals that the Enterococcal Sex Pheromone Response Results in Expression of Full-Length Conjugation Operon Transcripts in All Induced Cells. | AGREE | we designed fluorescence in situ hybridization chain reaction (HCR) probes for both early (QL, prgA, and prgB RNAs) and late QOp transcripts (prgJ, pcfC, and pcfG) | accepted_agree \| Platform: HCR-FISH |
| 10.1128/mbio.00796-16 | 2016 | Exposing the Three-Dimensional Biogeography and Metabolic States of Pathogens in Cystic Fibrosis Sputum via Hydrogel Embedding, Clearing, and rRNA Labeling. | AGREE | Aggregation measurements were performed from HCR-stained samples using probes listed in Table 1 | accepted_agree \| Platform: HCR-FISH |
| 10.1128/mbio.02622-19 | 2019 | Quantitative Visualization of Gene Expression in Mucoid and Nonmucoid Pseudomonas | AGREE | We designed and validated HCR v3.0 probe sets that could be used to (i) differentiate species by | accepted_agree \| Platform: HCR-FISH |

| DOI | Year | Title | Audit decision | Evidence (verbatim quote) | Validation status / scope flag |
|---|---|---|---|---|---|
| | | aeruginosa Aggregates Reveals Localized Peak Expression of Alginate in the Hypoxic Zone. | | targeting rRNAs and (ii) measure gene expression by targeting mRNAs. | |
| 10.1128/mbio.03751-21 | 2021 | Subcellular Detection of SARS-CoV-2 RNA in Human Tissue Reveals Distinct Localization in Alveolar Type 2 Pneumocytes and Alveolar Macrophages. | DISAGREE | We designed RNA FISH HCR probes using RajLab ProbeDesignHD software | validated_high_risk_disagreement \| Platform: HCR-FISH |
| 10.1128/msphere.00282-24 | 2024 | An approach to analyze spatiotemporal patterns of gene expression at single-cell resolution in <i>Candida albicans</i>-infected mouse tongues. | AGREE | As proof-of-concept, we first targeted the HWP1 transcript | accepted_agree \| Platform: HCR-FISH |
| 10.1128/msphere.00411-21 | 2021 | Imaging and Quantification of mRNA Molecules at Single-Cell Resolution in the Human Fungal Pathogen Candida albicans. | AGREE | We chose RTG3 and TYE7 which encode two transcription regulators | accepted_agree \| Platform: HCR-FISH |
| 10.1128/msphere.00657-25 | 2026 | Whole-tissue imaging reveals intrastrain diversity shapes the spatial organization of <i>Pseudomonas aeruginosa</i> in a murine infection model. | AGREE | We used HCR RNA-FISH probes targeting wbpL mRNA-A647 and 16S rRNA-A488 | accepted_agree \| Platform: HCR-FISH |
| 10.1128/spectrum.01115-24 | 2024 | Interplay between a polerovirus and a closterovirus decreases aphid transmission of the polerovirus. | DISAGREE | For the detection of BYV and BChV, we designed 25 (BYV) and 16 (BChV) primers with OligoMiner | validated_high_risk_disagreement \| Platform: SABER-FISH |
| 10.1136/gutjnl-2025-335793 | 2026 | Hybrid identity and distinct methylation profiles of incomplete intestinal metaplasia in the stomach | AGREE | we designed a custom panel comprising 293 genes enriched in epithelial cells | accepted_agree \| Platform: Xenium |
| 10.1158/1078-0432.ccr-23-3461 | 2024 | Spatial Transcriptome Mapping of the Desmoplastic Growth Pattern of Colorectal Liver Metastases by In Situ Sequencing Reveals a Biologically Relevant Zonation of the Desmoplastic Rim. | AGREE | Based on current literature, these genes were selected | accepted_agree \| Platform: in situ sequencing |
| 10.1161/atvbaha.125.324012 | 2026 | Single-Cell Multimodal Profiling Highlights Persistent Aortic Smooth Muscle Cell Changes in Diabetic Mice Despite Glycemic Control. | AGREE | We selected 479 genes based on DEG analysis from our scRNA-seq data and literature documenting cell type markers in the mouse aorta. | accepted_agree \| Platform: Xenium |
| 10.1176/appi.ajp.20230602 | 2024 | Translational Insights From Cell Type Variation Across Amygdala | DISAGREE | Probes for GAD1, NPFFR2, TSHZ1, GAD2, FOXP2, and SLC17A7 were hybridized in the | validated_high_risk_disagreement \| Platform: Other / unspecified |

| DOI | Year | Title | Audit decision | Evidence (verbatim quote) | Validation status / scope flag |
|---|---|---|---|---|---|
| | | Subnuclei in Rhesus Monkeys and Humans | | same rhesus individuals using RNAscope HiPlex v2. | |
| 10.1186/s12862-024-02237-y | 2024 | The adult shell matrix protein repertoire of the marine snail Crepidula is dominated by conserved genes that are also expressed in larvae. | DISAGREE | In situ expression patterns of 18 SMPs in 13 orthogroups were examined by HCR; SMPs were prioritized from prior differential expression in mantle, head, foot, and gill tissues. | validated_high_risk_disagreement \| Platform: HCR-FISH |
| 10.1186/s12915-019-0736-3 | 202 | Spatiotemporal mapping of RNA editing in the developing mouse brain using in situ sequencing reveals regional and cell-type-specific regulation. | AGREE | Padlock probes were designed to target 22 editing sites, and additional marker transcripts for interneurons, pyramidal neurons, oligodendrocytes, and astrocytes were included. | accepted_agree \| Platform: in situ sequencing |
| 10.1186/s12915-020-00874-5 | 202 | Automated identification of the mouse brain's spatial compartments from in situ sequencing data. | DISAGREE | The gene panels for the ISS experiments were selected based on single-cell RNA sequencing data from the hippocampus and using pci-Seq trimmed to two subsets of 82 and 97 genes. | validated_high_risk_disagreement \| Platform: in situ sequencing |
| 10.1186/s12915-022-01325-z | 2022 | Spatial and temporal heterogeneity in the lineage progression of fine oligodendrocyte subtypes. | AGREE | The 124 genes were selected from scRNA sequencing data based on median expression values, with additional differential expression analysis and some manually curated genes. | accepted_agree \| Platform: in situ sequencing |
| 10.1186/s12967-023-04384-0 | 2023 | Spatial tumour gene signature discriminates neoplastic from non-neoplastic compartments in colon cancer: unravelling predictive biomarkers for relapse. | AGREE | A panel of 176 genes was designed to examine pathways such as apoptosis, proliferation, angiogenesis, stemness, hypoxia, oxidative stress, invasion, and the microenvironment in relapsed versus non-relapsed patients. | accepted_agree \| Platform: in situ sequencing |
| 10.1186/s12974-025-03496-2 | 2025 | CXCR4 and CXCR6 dually limit T cell entry into the polyomavirus-infected brain | AGREE | The slides were loaded onto a MERFISH instrument to probe a set of 141 transcripts identifying resident/infiltrating cells and inflammation-associated mRNAs. | accepted_agree \| Platform: MERSCOPE |
| 10.1186/s13045-025-01716-z | 2025 | Deciphering cholangiocarcinoma heterogeneity and specific progenitor cell niche of extrahepatic cholangiocarcinoma at single-cell resolution. | AGREE | ISS was used to validate LY6D+ cancer cells and to inspect basal-like, stem cell, and proliferative marker genes in those cells. | accepted_agree \| Platform: in situ sequencing |
| 10.1186/s13059-024-03347-y | 2024 | StaVia: spatially and temporally aware cartography with higher- | DISAGREE | Panel chosen from scRNA-seq marker genes and known | validated_high_risk_disagreement \| Platform: MERFISH |

| DOI | Year | Title | Audit decision | Evidence (verbatim quote) | Validation status / scope flag |
|---|---|---|---|---|---|
| | | order random walks for cell atlases | | functional genes; 160 genes were used. | |
| 10.1186/s13059-025-03786-1 | 2025 | Systematic discovery of subcellular RNA patterns in the gut epithelium. | AGREE | We designed a MERFISH panel targeting 500 genes, including RNAs with unknown subcellular localization, APEX-seq targets, and tissue marker genes. | accepted_agree \| Platform: MERFISH |
| 10.1186/s13059-025-03920-z | 2025 | The differential subcellular localization of soybean transcripts, an additional regulatory mechanism of gene activity. | AGREE | The expression patterns of 23 marker genes were analyzed, and those genes had been previously identified from single-nucleus RNA-seq data. | accepted_agree \| Platform: Xenium;Molecular Cartography |
| 10.1186/s13227-024-00227-1 | 2024 | Hooked on zombie worms? Genetic blueprints of bristle formation in Osedax japonicus (Annelida). | AGREE | The comparative transcriptomic analysis focused on CS1, CS2, and NF70, and custom probe pairs were designed for each gene. | accepted_agree \| Platform: HCR-FISH |
| 10.1186/s13227-024-00231-5 | 2024 | Comparative Hox genes expression within the dimorphic annelid Streblospio benedicti reveals patterning variation during development. | AGREE | We identified 11 Hox genes using a homologous gene approach, and multiplex probes of those transcripts were then designed using HCR3.0 Probe Maker. | accepted_agree \| Platform: HCR-FISH |
| 10.1186/s13227-024-00235-1 | 2024 | Expression of distal limb patterning genes in Hypsibius exemplaris indicate regionalization and suggest distal identity of tardigrade legs. | AGREE | Seven distal limb patterning genes were identified from predicted protein sequences, and HCR probe sets were synthesized by Molecular Instruments based on those RNA sequences. | accepted_agree \| Platform: HCR-FISH |
| 10.1186/s13287-024-03854-6 | 2024 | Deciphering decidual deficiencies in recurrent spontaneous abortion and the therapeutic potential of mesenchymal stem cells at single-cell resolution. | AGREE | Padlock probes for the genes identified in this study were designed with PrimerQuest. | accepted_agree \| Platform: SCRINSHOT |
| 10.1186/s40478-025-02162-1 | 2025 | Spatial transcriptomics analysis uncovers ER stress in MANF-deficient Purkinje cells underlying alcohol-induced cerebellar neurodegeneration in mice. | AGREE | A customized 300-gene panel was designed using DEGs from Visium plus genes from prior publications, STRING, stress pathways, and cerebellum markers. | accepted_agree \| Platform: Xenium |
| 10.1242/bio.062391 | 2026 | Spatiotemporal analysis of genetic perturbations reveals a genetic cascade driving Tribolium gap gene initialization. | AGREE | The analysis focused on the core gap genes hb, Kr, mlpt, svb, and gt. | accepted_agree \| Platform: HCR-FISH |
| 10.1242/dev.140137 | 2016 | Mapping a multiplexed zoo of mRNA expression. | DISAGREE | When mapping the expression pattern for a new target mRNA, we balance brightness, robustness | validated_high_risk_disagreement \| Platform: HCR-FISH |

| DOI | Year | Title | Audit decision | Evidence (verbatim quote) | Validation status / scope flag |
|---|---|---|---|---|---|
| | | | | and cost considerations by using a probe set containing five DNA probes. | |
| 10.1242/dev.202307 | 2024 | HCR spectral imaging: 10-plex, quantitative, high-resolution RNA and protein imaging in highly autofluorescent samples. | DISAGREE | When assigning targets to Ch1-Ch10, higher-expression targets were assigned to channels that employ fluorophores with emission spectra strongly overlapping sample autofluorescence; other than these precautions, the targets were assigned randomly. | validated_high_risk_disagreement \| Platform: HCR-FISH |
| 10.1242/dev.202448 | 2024 | Open-source, high-throughput targeted in situ transcriptomics for developmental and tissue biology. | DISAGREE | q1: profiled a panel of 9 genes; genes label known cell types and include housekeeping mRNAs; q2: TARGETED spatial COMP paper. | validated_high_risk_disagreement \| Platform: in situ sequencing |
| 10.1242/dev.204814 | 2025 | Automated, high-throughput in situ hybridization of sea urchin (Lytechinus pictus) embryos. | DISAGREE | Using HT-HCR we probed three early developmental stages of sea urchin embryos, using a probe panel designed to target genes expressed at different developmental stages, in different spatial domains, and representing a wide range of functional categories. | validated_high_risk_disagreement \| Platform: HCR-FISH |
| 10.1242/dev.205254 | 2026 | Dynamic and non-uniform expression of key transcription factors provides insights into the emergence of neural crest cells at the neural plate border. | AGREE | HCR-FISH was used to probe expression of the neural crest markers snai2, sox8, foxd3, sox9, and sox10. | accepted_agree \| Platform: HCR-FISH |
| 10.1242/jcs.263407 | 2024 | ‘Iterative Bleaching Extends Multiplexity’ facilitates simultaneous identification of all major retinal cell types | AGREE | The panel of markers against proteins expressed in each cell type of the zebrafish retina was designed by the authors. | accepted_agree \| Platform: HCR-FISH |
| 10.1261/rna.079801.123 | 2024 | Computational analysis of super-resolved in situ sequencing data reveals genes modified by immune-tumor contact events. | AGREE | The interrogated genes included cell-type markers and genes known or suspected to be associated with cancer tissues. | accepted_agree \| Platform: ExSeq |
| 10.1262/jrd.2016-172 | 2017 | Visualization of the spatial arrangement of nuclear organization using three-dimensional fluorescence in situ hybridization in early mouse embryos: A new "EASI-FISH chamber glass" for mammalian embryos. | AGREE | As a 3D-FISH model, the study focused on the Zscan4 gene. | accepted_agree \| Platform: EASI-FISH |

| DOI | Year | Title | Audit decision | Evidence (verbatim quote) | Validation status / scope flag |
|---|---|---|---|---|---|
| 10.1371/journal.pbio.3000675 | 202 | SCRINSHOT enables spatial mapping of cell states in tissue sections with single-cell resolution. | DISAGREE | q1: used a panel of selective markers for club, goblet, basal, tuft, and ionocytes; q2: TARGETED spatial COMP paper. | validated_high_risk_disagreement \| Platform: SCRINSHOT |
| 10.1371/journal.pgen.1011782 | 2026 | The rewiring of a terminal selector regulatory cascade generates convergent neuronal laterality. | AGREE | The study systematically identified terminal identity genes by examining gcy genes and adopted a trio of gcy marker genes. | accepted_agree \| Platform: HCR-FISH |
| 10.1523/jneurosci.2049-22.2023 | 2023 | Effect of Selective Lesions of Nucleus Accumbens µ-Opioid Receptor-Expressing Cells on Heroin Self-Administration in Male and Female Rats: A Study with Novel <i>Oprm1-Cre</i> Knock-in Rats. | AGREE | The authors designed the Split-initiator DNA probes version 2.0 and had them synthesized by IDT. | accepted_agree \| Platform: HCR-FISH |
| 10.15252/msb.20199146 | 202 | Mammalian gene expression variability is explained by underlying cell state. | AGREE | The study measured 150 genes including Ca2+ signaling genes, cell-cycle genes, and genes correlating with cell state. | accepted_agree \| Platform: MERFISH |
| 10.18632/oncotarget.10245 | 2017 | A molecular pathology method for sequential fluorescence in situ hybridization for multi-gene analysis at the single-cell level | AGREE | Because not all FISH probes for disease genes are commercially available, the authors developed the probes used in the work themselves. | accepted_agree \| Platform: seqFISH |
| 10.21203/rs.3.rs-4468724/v1 | 2024 | A spatially resolved human glioblastoma atlas reveals distinct cellular and molecular patterns of anatomical niches | DISAGREE | "Recursive feature elimination (rfe) from the caret package in R was used for filtering functionally redundant genes." The packet also says this was a "custom panel of 348 genes" and tool_names=caret, which is not one of the 18 review tools. | validated_high_risk_disagreement \| Platform: Xenium |
| 10.21203/rs.3.rs-5374449/v1 | 2024 | Molecular Cartography of the Human and Mouse Down Syndrome Brain | AGREE | "A panel of 500 genes was curated to identify major brain cell types..." and evidence_manual is present; no tool or vendor panel is identified. | accepted_agree \| Platform: MERFISH |
| 10.21203/rs.3.rs-6289741/v1 | 2025 | Modular strategies for spatial mapping of diverse cell type data of the mouse brain | DISAGREE | q1: "A set of 500 genes were imaged that had been carefully chosen to distinguish the ~5200 clusters of our existing RNAseq taxonomy." | validated_high_risk_disagreement \| Platform: MERFISH |

| DOI | Year | Title | Audit decision | Evidence (verbatim quote) | Validation status / scope flag |
|---|---|---|---|---|---|
| 10.21203/rs.3.rs-6314842/v1 | 2025 | A Spatial Multi-Omic Framework Identifies Gliomas Permissive to TIL Expansion | AGREE | "a custom panel containing probes for 480 custom gene panel... is added to the tissue" and uses_manual_additions=YES. | accepted_agree \| Platform: Xenium |
| 10.21203/rs.3.rs-7895391/v1 | 2025 | A spatial and projection-based transcriptomic atlas of paraventricular hypothalamic cell types | AGREE | "503 genes specifically curated for the PVH region, comprised of top differentially expressed genes... canonical marker genes... and functionally relevant genes selected from the literature." | accepted_agree \| Platform: MERSCOPE |
| 10.26508/lsa.202201701 | 2023 | Concordance of MERFISH spatial transcriptomics with bulk and single-cell RNA sequencing. | AGREE | "The gene panel was designed by selecting the top 10 differentially expressed genes in each cell type according to Tabula Muris Senis cell atlas scRNA-seq data." | accepted_agree \| Platform: MERSCOPE |
| 10.3389/fimmu.2022.876321 | 2022 | Spatial Resolution of Mycobacterium tuberculosis Bacteria and Their Surrounding Immune Environments Based on Selected Key Transcripts in Mouse Lungs | AGREE | "a padlock probe panel against 33 immune transcripts of selected immune cell markers, activation molecules, and cytokines" | accepted_agree \| Platform: in situ sequencing |
| 10.3389/fimmu.2023.1332733 | 2023 | Immune mapping of human tuberculosis and sarcoidosis lung granulomas | AGREE | "Transcripts coding for chemokine receptors, cytokines, effector molecules, transcription factors, and surface molecules ... were targeted." | accepted_agree \| Platform: in situ sequencing |
| 10.3389/fnmol.2020.00075 | 202 | Modified <i>in situ</i> Hybridization Chain Reaction Using Short Hairpin DNAs. | DISAGREE | We designed split probes according to Choi et al. (2018). The sequence for probes were chosen in consideration of GC contents and homology without further optimization and evaluation. | validated_high_risk_disagreement \| Platform: HCR-FISH |
| 10.3389/fonc.2022.960734 | 2022 | Automated multimodal fluorescence microscopy for hyperplex spatial-proteomics: Coupling microfluidic-based immunofluorescence to high resolution, high sensitivity, three-dimensional analysis of histological slides | DISAGREE | The two employed probes were synthesized to detect Actin B1 and B-RAF mature RNA transcripts. | validated_high_risk_disagreement \| Platform: Other / unspecified |
| 10.3389/frdem.2025.1546433 | 2025 | Neurogenesis drives hippocampal formation-wide spatial transcription alterations in health and Alzheimer's disease | AGREE | The evidence says the tissue was analyzed using a panel of 158 genes, with the full list of genes described as manually selected probes from prior work. | accepted_agree \| Platform: CARTANA |

| DOI | Year | Title | Audit decision | Evidence (verbatim quote) | Validation status / scope flag |
|---|---|---|---|---|---|
| 10.3390/biomedicines10020484 | 2022 | In Situ Gene Expression in Native Cryofixed Bone Tissue. | AGREE | The packet says specific probes were designed and synthesized to target the mRNAs of beta-actin, and that primers were chosen using ApE software. | accepted_agree \| Platform: HCR-FISH |
| 10.3390/cells12040614 | 2023 | Retinoic Acid and POU Genes in Developing Amphioxus: A Focus on Neural Development. | AGREE | The packet says the expression patterns were assessed by chromogenic in situ hybridization and hybridization chain reaction, and that primers were designed for the genes. | accepted_agree \| Platform: HCR-FISH |
| 10.3390/cells13171435 | 2024 | Identification and Characterization of Multiple Paneth Cell Types in the Mouse Small Intestine | AGREE | The packet says the probe panel was designed from single-cell and bulk RNA-seq data and that probes were selected for all intestinal cell types, including differential markers. | accepted_agree \| Platform: Molecular Cartography |
| 10.3390/ijms22168539 | 2021 | Chromosomal Characterization of <i>Tripidium arundinaceum</i> Revealed by Oligo-FISH. | AGREE | The packet says the authors selected seven probes to identify the chromosomes, and that multiple oligo-FISH probes were used. | accepted_agree \| Platform: seqFISH |
| 10.3390/mps8020031 | 2025 | Mapping Gene Expression in Whole Larval Brains of <i>Bicyclus anynana</i> Butterflies. | DISAGREE | The packet says mapped the expression domains of ten genes and that probes against the 10 genes were designed using the referenced Excel sheets and downloaded CDS sequences. | validated_high_risk_disagreement \| Platform: HCR-FISH |
| 10.34133/bmef.0110 | 2025 | Graph-Based 3-Dimensional Spatial Gene Neighborhood Networks of Single Cells in Gels and Tissues. | DISAGREE | A library of 31 RNA targets was used in the 3D gel/tissue experiment. | validated_high_risk_disagreement \| Platform: MERFISH |
| 10.64898/2025.12.02.691876 | 2025 | Multiscale Spatial Transcriptomic Atlas of Human Basal Ganglia Cell-Type and Cellular Community Organization | AGREE | The packet says the panel comprised canonical cell type markers, housekeeping genes, and genes differentially expressed across disease states. | accepted_agree \| Platform: MERFISH |
| 10.64898/2025.12.06.692770 | 2025 | Brain-immune interactions generate pathogen-specific sickness states | AGREE | The packet says the MERFISH studies used a curated library of genes of interest based on published literature and that the library also included differentially expressed genes. | accepted_agree \| Platform: MERFISH |

| DOI | Year | Title | Audit decision | Evidence (verbatim quote) | Validation status / scope flag |
|---|---|---|---|---|---|
| 10.64898/2025.12.08.693033 | 2025 | Alveolar Type II Cell-derived MMP1 <sup>high</sup> basal cells promote destructive microcysts in idiopathic pulmonary fibrosis | AGREE | The packet says tissue sections were processed with Xenium and hybridized with a selected panel of RNA probes for the listed genes. | accepted_agree \| Platform: Xenium |
| 10.64898/2025.12.18.695268 | 2025 | A poly(A) isoform–aware single-cell and spatial atlas defines fibroblast niches in the human bladder | AGREE | The packet says a custom Xenium human lower urinary tract probe panel targeting 354 genes and 125 mRNA isoforms was designed. | accepted_agree \| Platform: Xenium |
| 10.64898/2025.12.23.696217 | 2025 | Spatially resolved translational dysregulation in <i>Grin2a</i> +/- mouse model of schizophrenia | AGREE | The packet says the 3,447-gene panel was compiled from canonical brain cell-type markers and genes differentially expressed in bulk RNA-seq, snRNA-seq, or proteomic data. | accepted_agree \| Platform: STARmap |
| 10.64898/2026.01.03.697464 | 2026 | Spatial dynamics of cellular and molecular plasticity in the maternal and postpartum mouse brain | AGREE | The packet says the authors used a custom 500-gene panel designed to profile brain cell types and pregnancy-related biological processes. | accepted_agree \| Platform: MERSCOPE |
| 10.64898/2026.02.08.704520 | 2026 | Mitochondrial Retrograde Signaling in <i>Arabidopsis thaliana</i> : heterogenous, spatial and polarised aspects | AGREE | First, we designed a 10x Xenium panel of 465 probes incorporating the top markers from each cluster | accepted_agree \| Platform: Xenium |
| 10.64898/2026.02.10.705126 | 2026 | X-CODE: a dual RNA barcoding system for multi-platform clonal tracking and spatial phenotyping | DISAGREE | q1: selected 27 VBSs for BU construction; 23 adapted from previously validated PLAYR insert probes and 4 developed following PLAYR criteria. | validated_high_risk_disagreement \| Platform: Xenium |
| 10.64898/2026.02.15.26346348 | 2026 | A network-based atlas of human skeletal muscle aging | AGREE | The gene list was sent to Vizgen Inc, and they selected 30-50 merfish probes per gene | accepted_agree \| Platform: Xenium;MERSCOPE |
| 10.64898/2026.02.20.707030 | 2026 | Single-phage profiling illuminates viral individuality during cell fate determination | AGREE | To capture the progression of phage development, we chose to target 18 genes | accepted_agree \| Platform: seqFISH |
| 10.64898/2026.03.05.709830 | 2026 | Cardiac-immune microniches programme macrophage states in the regenerating heart | DISAGREE | To enrich the panel for functional communication circuits, we adapted the NicheNet pipeline ... We assembled a 500-gene panel that combined five classes of genes | validated_high_risk_disagreement \| Platform: MERSCOPE |
| 10.7554/elife.101035 | 2025 | Molecular and spatial transcriptomic classification of midbrain dopamine neurons and | AGREE | We assembled a 500 gene panel which included 120 of the top | accepted_agree \| Platform: MERFISH |

| DOI | Year | Title | Audit decision | Evidence (verbatim quote) | Validation status / scope flag |
|---|---|---|---|---|---|
| | | their alterations in a LRRK2<sup>G2019S</sup> model of Parkinson's disease. | | DEGs identified within the snRNA-seq dataset | |
| 10.7554/elife.105388 | 2025 | Combined transcriptomic, connectivity, and activity profiling of the medial amygdala using highly amplified multiplexed in situ hybridization (hamFISH) | DISAGREE | The authors assembled a gene panel of cluster markers and previous-study genes, then finalized 31 genes for hamFISH. | validated_high_risk_disagreement \| Platform: Other / unspecified |
| 10.7554/elife.84262 | 2023 | Multimodal mapping of cell types and projections in the central nucleus of the amygdala. | AGREE | To identify marker genes for EASI-FISH, we first started with the differentially expressed genes | accepted_agree \| Platform: EASI-FISH |
| 10.7554/elife.87866 | 2024 | Rabies virus-based barcoded neuroanatomy resolved by single-cell RNA and in situ sequencing. | AGREE | we used padlock probes to target both the flanking regions of the barcodes and 104 marker genes | accepted_agree \| Platform: in situ sequencing |
| 10.7554/elife.90029 | 2024 | Three-dimensional single-cell transcriptome imaging of thick tissues. | DISAGREE | Two sets of genes were selected for MERFISH imaging: a previously selected 242-gene set and a previously selected 156-gene set. | validated_high_risk_disagreement \| Platform: MERFISH |

### FILTERED (n = 1159)

*No adjudicable targeted panel-design evidence in the available text (e.g. non-targeted spatial, computational paper without a new panel, supplement-only PDF).*

| DOI | Year | Title | Audit decision | Evidence (verbatim quote) | Validation status / scope flag |
|---|---|---|---|---|---|
| 10.1001/jama.2017.5640 | 2017 | An International Human Cell Atlas Consortium Takes Shape. | AGREE | gemma_final=FILTERED; doc_type=NON_ARTICLE; risk_flags=non_primary_document | accepted_agree |
| 10.1002/adma.201908410 | 202 | Combining Qdot Nanotechnology and DNA Nanotechnology for Sensitive Single-Cell Imaging. | AGREE | gemma_final=FILTERED; paper_type=EXP; spatial=NO; panel_scope=UNCLEAR; risk_flags=prefiltered_without_full_panel_qs | accepted_agree |
| 10.1002/advs.202303752 | 2024 | Characterization of the Nucleus Pulposus Progenitor Cells via Spatial Transcriptomics. | AGREE | gemma_final=FILTERED; paper_type=EXP; spatial=YES; panel_scope=NOT_TARGETED; notes=Pass-A abstract: not targeted | accepted_agree |

| DOI | Year | Title | Audit decision | Evidence (verbatim quote) | Validation status / scope flag |
|---|---|---|---|---|---|
| 10.1002/advs.202306329 | 2023 | Highly Accurate Estimation of Cell Type Abundance in Bulk Tissues Based on Single-Cell Reference and Domain Adaptive Matching | AGREE | Title only; paper_type COMP; spatial blank; panel_scope UNCLEAR; q1 panel-design fields are empty; gemma_final=FILTERED from abstract prefilter. | accepted_agree |
| 10.1002/advs.202505021 | 2025 | SCRIPT: Predicting Single-Cell Long-Range Cis-Regulation Based on Pretrained Graph Attention Networks | AGREE | paper.title='SCRIPT: Predicting Single-Cell Long-Range Cis-Regulation Based on Pretrained Graph Attention Networks'; citation_screening_prior.labels='COMP'; gemma_final.classification='FILTERED'; gemma_q1_panel_design blank; gemma_q2_paper_profile={'evidence_scope': '', 'evidence_spatial': '', 'evidence_type': '', 'panel_scope': '', 'paper_type': 'COMP', 'profile_confidence': '', 'spatial': ''} | accepted_agree |
| 10.1002/advs.202509090 | 2025 | SpaBatch: Deep Learning-Based Cross-Slice Integration and 3D Spatial Domain Identification in Spatial Transcriptomics. | AGREE | gemma_final=FILTERED; paper_type=COMP; spatial=YES; panel_scope=UNCLEAR; title=SpaBatch: Deep Learning-Based Cross-Slice Integration and 3D Spatial Domain Identification in Spatial Transcriptomics. | accepted_agree |
| 10.1002/advs.202511856 | 2025 | Immunoregulatory Roles of Tumor-Originated Pericytes Identified by Single-Cell Analysis in Glioblastoma | AGREE | paper.title='Immunoregulatory Roles of Tumor-Originated Pericytes Identified by Single-Cell Analysis in Glioblastoma'; citation_screening_prior.labels=''; gemma_final.classification='FILTERED'; gemma_q1_panel_design blank; gemma_q2_paper_profile={'evidence_scope': '', 'evidence_spatial': '', 'evidence_type': '', 'panel_scope': 'UNCLEAR', 'paper_type': 'EXP', 'profile_confidence': 'HIGH', 'spatial': 'NO'} | accepted_agree |
| 10.1002/advs.202513210 | 2025 | CELLama: Foundation Model for Single Cell and Spatial Transcriptomics by Cell Embedding Leveraging Language Model Abilities | AGREE | paper.title='CELLama: Foundation Model for Single Cell and Spatial Transcriptomics by Cell Embedding Leveraging Language Model Abilities'; citation_screening_prior.labels=''; | accepted_agree |

| DOI | Year | Title | Audit decision | Evidence (verbatim quote) | Validation status / scope flag |
|---|---|---|---|---|---|
| | | | | gemma_final.classification='FILTERED'; gemma_q1_panel_design blank; gemma_q2_paper_profile={'evidence_scope': '', 'evidence_spatial': '', 'evidence_type': '', 'panel_scope': 'UNCLEAR', 'paper_type': 'COMP', 'profile_confidence': 'HIGH', 'spatial': 'YES'} | |
| 10.1002/alz.076665 | 2023 | A complete pipeline for high-plex spatial proteomic profiling and analysis of neural cell phenotypes on the CosMx™ Spatial Molecular Imager and AtoMx™ Spatial Informatics Platform | AGREE | gemma_final=FILTERED; doc_type=NON_ARTICLE; risk_flags=non_primary_document | accepted_agree |
| 10.1002/alz.080729 | 2023 | Spatial Transcriptomics Analysis of Vulnerable Cell Populations in Middle Temporal Gyrus | AGREE | gemma_final=FILTERED; doc_type=NON_ARTICLE; risk_flags=non_primary_document | accepted_agree |
| 10.1002/alz.095162 | 2024 | Mapping the spatial immunometabolic responses of microglia in response to Alzheimer’s pathology | AGREE | gemma_final=FILTERED; doc_type=NON_ARTICLE; risk_flags=non_primary_document | accepted_agree |
| 10.1002/alz70855_107308 | 2025 | Multiomic Insights into Inflammation-Driven Microglial Dysregulation in AD | AGREE | gemma_final=FILTERED; doc_type=NON_ARTICLE; risk_flags=non_primary_document | accepted_agree |
| 10.1002/alz70859_099554 | 2025 | Drug Development. | AGREE | gemma_final=FILTERED; doc_type=NON_ARTICLE; risk_flags=non_primary_document | accepted_agree |
| 10.1002/aqc.3667 | 2021 | A 40-year chronology of the vulnerability of spinetail devil ray (<scp><i>Mobula mobular</i> </scp>) to eastern Pacific tuna fisheries and options for future conservation and management | AGREE | gemma_final=FILTERED; paper_type=EXP; spatial=NO; panel_scope=UNCLEAR; risk_flags=prefiltered_without_full_panel_qs | accepted_agree |
| 10.1002/aro2.70016 | 2025 | Technological Innovations and Applications of Spatial Transcriptomics in Livestock and Poultry Research | AGREE | paper.title='Technological Innovations and Applications of Spatial Transcriptomics in Livestock and Poultry Research'; citation_screening_prior.labels='REV'; gemma_final.classification='FILTERED'; gemma_q1_panel_design blank; gemma_q2_paper_profile={'evidence_scope': '', 'evidence_spatial': '', 'evidence_type': '', 'panel_scope': '', | accepted_agree |

| DOI | Year | Title | Audit decision | Evidence (verbatim quote) | Validation status / scope flag |
|---|---|---|---|---|---|
| | | | | 'paper_type': '', 'profile_confidence': '', 'spatial': ''} | |
| 10.1002/bies.202300130 | 2023 | Exploring the role of transcriptional and post-transcriptional processes in mRNA co-expression | AGREE | gemma_final=FILTERED; paper_type=EXP; spatial=NO; panel_scope=UNCLEAR; risk_flags=prefiltered_without_full_panel_qs | accepted_agree |
| 10.1002/ctd2.120 | 2022 | Single-cell nucleic acid sequencing reinforces to reveal signal-dependent pathways contributing to cellular heterogeneity | AGREE | gemma_final=FILTERED; paper_type=OTHER; spatial=UNKNOWN; panel_scope=UNCLEAR; risk_flags=prefiltered_without_full_panel_qs | accepted_agree |
| 10.1002/ctd2.70077 | 2025 | Significance of stereologically spatiotemporal cells in molecular medicine | AGREE | gemma_final=FILTERED; paper_type=OTHER; spatial=YES; panel_scope=NOT_TARGETED; risk_flags=prefiltered_without_full_panel_qs | accepted_agree |
| 10.1002/ctm2.1127 | 2022 | Re-exploring tissue regeneration by novel spatial transcriptomics technologies | AGREE | gemma_final=FILTERED; paper_type=OTHER; spatial=YES; panel_scope=UNCLEAR; risk_flags=prefiltered_without_full_panel_qs | accepted_agree |
| 10.1002/ctm2.1264 | 2023 | Spatial transcriptomics herald a new era of transcriptome research | AGREE | gemma_final=FILTERED; paper_type=OTHER; spatial=YES; panel_scope=UNCLEAR; risk_flags=prefiltered_without_full_panel_qs | accepted_agree |
| 10.1002/ctm2.70119 | 2024 | Statistical and computational methods for enabling the clinical and translational application of spatial transcriptomics | AGREE | gemma_final=FILTERED; paper_type=OTHER; spatial=YES; panel_scope=UNCLEAR; risk_flags=prefiltered_without_full_panel_qs | accepted_agree |
| 10.1002/ctm2.70425 | 2025 | Multi-omics profiling identifies <i>TNFRSF18</i> as a novel marker of exhausted CD8$^{+}$ T cells and reveals tumour-immune dynamics in colorectal cancer | AGREE | paper.title='Multi-omics profiling identifies <i>TNFRSF18</i> as a novel marker of exhausted CD8$^{+}$ T cells and reveals tumour-immune dynamics in colorectal cancer'; citation_screening_prior.labels=''; gemma_final.classification='FILTERED'; gemma_q1_panel_design blank; gemma_q2_paper_profile={'evidence_scope': '', 'evidence_spatial': '', 'evidence_type': '', 'panel_scope': 'UNCLEAR', 'paper_type': 'EXP', | accepted_agree |

| DOI | Year | Title | Audit decision | Evidence (verbatim quote) | Validation status / scope flag |
|---|---|---|---|---|---|
| | | | | 'profile_confidence': 'HIGH', 'spatial': 'NO'} | |
| 10.1002/cyto.a.23955 | 202 | PySpacell: A Python Package for Spatial Analysis of Cell Images. | AGREE | gemma_final=FILTERED; paper_type=COMP; spatial=YES; panel_scope=UNCLEAR; title=PySpacell: A Python Package for Spatial Analysis of Cell Images. | accepted_agree |
| 10.1002/cyto.a.23986 | 202 | Special Issue on Machine Learning for Single Cell Data | AGREE | gemma_final=FILTERED; doc_type=EDITORIAL; risk_flags=prefiltered_without_full_panel_qs | accepted_agree |
| 10.1002/cyto.a.24884 | 2024 | Points2Regions: Fast, interactive clustering of imaging-based spatial transcriptomics data | AGREE | gemma_final=FILTERED; paper_type=COMP; spatial=YES; panel_scope=UNCLEAR; title=Points2Regions: Fast, interactive clustering of imaging-based spatial transcriptomics data | accepted_agree |
| 10.1002/cyto.a.24939 | 2025 | Nuclear-Free Zoning | AGREE | gemma_final=FILTERED; paper_type=OTHER; spatial=YES; panel_scope=UNCLEAR; risk_flags=prefiltered_without_full_panel_qs | accepted_agree |
| 10.1002/eji.70119 | 2025 | Spatially Resolved Profiling of Compartmentalized Muscle and Brain Inflammation | AGREE | paper.title='Spatially Resolved Profiling of Compartmentalized Muscle and Brain Inflammation'; citation_screening_prior.labels='REV'; gemma_final.classification='FILTERED'; gemma_q1_panel_design blank; gemma_q2_paper_profile={'evidence_scope': '', 'evidence_spatial': '', 'evidence_type': '', 'panel_scope': '', 'paper_type': '', 'profile_confidence': '', 'spatial': ''} | accepted_agree |
| 10.1002/epi.70079 | 2026 | Spatial transcriptomics in epilepsy research: Early successes, opportunities, and challenges | AGREE | paper_type=OTHER; spatial=YES; panel_scope=UNCLEAR; gemma_final_notes=Pass-A abstract: OTHER → FILTERED | accepted_agree |
| 10.1002/fft2.70022 | 2025 | Highly Targeted Metabolomics Coupled With Gene Expression Analysis by RT–qPCR Improves Beef Separation Based on Grass, Grain, or Grape Supplemented Diet | AGREE | paper.title='Highly Targeted Metabolomics Coupled With Gene Expression Analysis by RT–qPCR Improves Beef Separation Based on Grass, Grain, or Grape Supplemented Diet'; citation_screening_prior.labels='N | accepted_agree |

| DOI | Year | Title | Audit decision | Evidence (verbatim quote) | Validation status / scope flag |
|---|---|---|---|---|---|
| | | | | ON-ST'; gemma_final.classification='FILTERED'; gemma_q1_panel_design blank; gemma_q2_paper_profile={'evidence_scope': '', 'evidence_spatial': '', 'evidence_type': '', 'panel_scope': '', 'paper_type': '', 'profile_confidence': '', 'spatial': 'NO'} | |
| 10.1002/hon.70093_19 | 2025 | 19 \| MULTI-OMIC PROFILING OF DARK ZONE LYMPHOMAS UNCOVERS DYSREGULATED GENE REGULATORY NETWORKS REFLECTING DZ-LZ B-CELL TRANSITION DYNAMICS | AGREE | doc_type=NON_ARTICLE; gemma_final_notes=Q0: NON_ARTICLE → FILTERED | accepted_agree |
| 10.1002/hon.70094_172 | 2025 | 172 \| AGGRESSIVE B-CELL LYMPHOMAS RETAIN ATR-DEPENDENT DETERMINANTS OF T-CELL EXCLUSION FROM THE GERMINAL CENTER DARK ZONE | AGREE | doc_type=NON_ARTICLE; gemma_final_notes=Q0: NON_ARTICLE → FILTERED | accepted_agree |
| 10.1002/hon.70096_477 | 2025 | 477 \| LONGITUDINAL MULTI-OMICS INVESTIGATION IN IBRUTINIB-BORTEZOMIB-R-CHOP-TREATED DLBCL PATIENTS | AGREE | doc_type=NON_ARTICLE; gemma_final_notes=Q0: NON_ARTICLE → FILTERED | accepted_agree |
| 10.1002/imt2.70054 | 2025 | Gut microbiota and tuberculosis | AGREE | paper.title='Gut microbiota and tuberculosis'; citation_screening_prior.labels='REV'; gemma_final.classification='FILTERED'; gemma_q1_panel_design blank; gemma_q2_paper_profile={'evidence_scope': '', 'evidence_spatial': '', 'evidence_type': '', 'panel_scope': '', 'paper_type': '', 'profile_confidence': '', 'spatial': ''} | accepted_agree |
| 10.1002/jlb.4cova0121-084rr | 2021 | Corticosteroid treatment in COVID-19 modulates host inflammatory responses and transcriptional signatures of immune dysregulation | AGREE | Title: 'Corticosteroid treatment in COVID-19 modulates host inflammatory responses and transcriptional signatures of immune dysregulation'; packet fields show non-spatial prior and Gemma's FILTERED prefilter. | accepted_agree |

| DOI | Year | Title | Audit decision | Evidence (verbatim quote) | Validation status / scope flag |
|---|---|---|---|---|---|
| 10.1002/jmri.29610 | 2025 | Quantitative MRI for Monitoring Metabolic Dysfunction-Associated Steatotic Liver Disease: A Test-Retest Repeatability Study. | AGREE | paper_type=EXP; spatial=NO; panel_scope=UNCLEAR; gemma_final_notes=Pass-A abstract: non-spatial → FILTERED | accepted_agree |
| 10.1002/mog2.88 | 2024 | Deciphering transcriptional bursts and enhancer dynamics: Advancing cancer therapeutics through single-cell global run-on sequencing | AGREE | paper_type=COMP; spatial=NO; panel_scope=UNCLEAR; gemma_final_notes=Pass-A abstract: COMP → FILTERED | accepted_agree |
| 10.1002/pdi3.70027 | 2025 | Single-Cell Profiling of Pediatric High-Grade Gliomas Reveals OPC-Like Subpopulations Driving Tumorigenic Lineage Transitions | AGREE | paper.title='Single-Cell Profiling of Pediatric High-Grade Gliomas Reveals OPC-Like Subpopulations Driving Tumorigenic Lineage Transitions'; citation_screening_prior.labels='NON-ST'; gemma_final.classification='FILTERED'; gemma_q1_panel_design blank; gemma_q2_paper_profile={'evidence_scope': '', 'evidence_spatial': '', 'evidence_type': '', 'panel_scope': '', 'paper_type': '', 'profile_confidence': '', 'spatial': 'NO'} | accepted_agree |
| 10.1002/qub2.70011 | 2025 | Spatial transcriptomics iterative hierarchical clustering (stIHC): A novel method for identifying spatial gene co-expression modules. | AGREE | paper_type=COMP; spatial=YES; panel_scope=UNCLEAR; gemma_final_notes=Pass-A abstract: COMP → FILTERED | accepted_agree |
| 10.1002/smll.202512096 | 2026 | High Sensitivity Cardiac Troponin I Detection via MP-Locked Aptamer and Multimeric DNAzyme-Coupled Hyperbranched Hybridization Chain Reaction. | AGREE | paper_type=EXP; spatial=NO; panel_scope=UNCLEAR; gemma_final_notes=Pass-A abstract: non-spatial → FILTERED | accepted_agree |
| 10.1002/smtd.202400505 | 2024 | Orthogonal DNA Self-Assembly-Based Expansion Microscopy Platform for Amplified, Multiplexed Biomarker Imaging. | AGREE | paper_type=COMP; spatial=NO; panel_scope=UNCLEAR; gemma_final_notes=Pass-A abstract: COMP → FILTERED | accepted_agree |
| 10.1002/smtd.202400975 | 2025 | UCS: A Unified Approach to Cell Segmentation for Subcellular Spatial Transcriptomics. | AGREE | paper_type=COMP; spatial=YES; panel_scope=UNCLEAR; gemma_final_notes=Pass-A abstract: COMP → FILTERED | accepted_agree |
| 10.1002/smtd.202570019 | 2025 | Integrating Metabolic RNA Labeling-Based Time-Resolved Single-Cell RNA Sequencing with | AGREE | paper_type=EXP; spatial=YES; panel_scope=NOT_TARGETED; gemma_final_notes=Pass-A | accepted_agree |

| DOI | Year | Title | Audit decision | Evidence (verbatim quote) | Validation status / scope flag |
|---|---|---|---|---|---|
| | | Spatial Transcriptomics for Spatiotemporal Transcriptomic Analysis (Small Methods 3/2025) | | abstract: not targeted → FILTERED | |
| 10.1002/wrna.1842 | 2024 | Spatial analysis toolkits for <scp>RNA</scp> in situ sequencing | AGREE | paper_type=OTHER; spatial=YES; panel_scope=TARGETED; gemma_final_notes=Pass-A abstract: OTHER → FILTERED | accepted_agree |
| 10.1007/978-1-0716-4184-2_5 | 2024 | Homebuilt Imaging-Based Spatial Transcriptomics: Tertiary Lymphoid Structures as a Case Example | AGREE | Title is 'Homebuilt Imaging-Based Spatial Transcriptomics: Tertiary Lymphoid Structures as a Case Example'; gemma notes say OTHER -> FILTERED and q1 panel-design fields are empty. | accepted_agree |
| 10.1007/978-1-0716-4276-4_5 | 2025 | Spatial-Omics Methods and Applications | AGREE | Citation prior labels the item REV; title is 'Spatial-Omics Methods and Applications'; q1 panel-design fields are empty. | accepted_agree |
| 10.1007/978-3-031-71452-8_10 | 2024 | Performance Metrics of Binary Classifiers | NEEDS_MANUAL | spatial=NO; packet does not support an in-scope spatial panel-design record. Original audit evidence: q0: main article on performance metrics of binary classifiers; q1/q2: no panel-design evidence. | validated_high_risk_manual_filtered |
| 10.1007/978-3-031-81728-1_38 | 2025 | Single-Cell Genomics | AGREE | gemma_q0_document_type=REVIEW; doc_evidence says the chapter is an overview of single-cell technologies and methods. | accepted_agree |
| 10.1007/978-3-031-90252-9_38 | 2025 | GeneCover: A Combinatorial Approach for Label-Free Marker Gene Selection | AGREE | Title: 'GeneCover: A Combinatorial Approach for Label-Free Marker Gene Selection'; packet fields show Q0: NON_ARTICLE and prior tools listing GeneCover-related tool names, with Gemma's FILTERED prefilter. | accepted_agree |
| 10.1007/978-3-031-90252-9_42 | 2025 | Unified Integration of Spatial Transcriptomics Across Platforms | AGREE | Title: 'Unified Integration of Spatial Transcriptomics Across Platforms'; packet fields show Q0: NON_ARTICLE and Gemma's FILTERED prefilter. | accepted_agree |
| 10.1007/978-3-031-90252-9_53 | 2025 | Optimal Marker Genes for c-Separated Cell Types | AGREE | Title: 'Optimal Marker Genes for c-Separated Cell Types'; packet fields show Q0: NON_ARTICLE and Gemma's FILTERED prefilter. | accepted_agree |

| DOI | Year | Title | Audit decision | Evidence (verbatim quote) | Validation status / scope flag |
|---|---|---|---|---|---|
| 10.1007/978-3-032-10024-5_11 | 2026 | Next-Generation Sequencing Methods Applied to Organoids in Cancer Research | AGREE | gemma_q0_document_type=REVIEW; doc_evidence says the chapter reviews NGS methods for organoid cancer studies. | accepted_agree |
| 10.1007/978-3-662-65902-1_4 | 2022 | Approaches to Marker Gene Identification from Single-Cell RNA-Sequencing Data | AGREE | gemma_q0_document_type=REVIEW; doc_evidence says the chapter reviews marker selection methods. | accepted_agree |
| 10.1007/s00262-025-03952-1 | 2025 | Distinct maturity and spatial distribution of tertiary lymphoid structures in head and neck squamous cell carcinoma: implications for tumor immunity and clinical outcomes | AGREE | paper_type=EXP; spatial=YES; panel_scope=NOT_TARGETED; gemma_final_notes=Pass-A abstract: not targeted → FILTERED | accepted_agree |
| 10.1007/s00292-026-01551-0 | 2026 | Half a century of gene expression analysis: The path to modern spatial transcriptomics in autopsy tissue | AGREE | Title is 'Half a century of gene expression analysis: The path to modern spatial transcriptomics in autopsy tissue'; gemma notes say OTHER -> FILTERED and q1 panel-design fields are empty. | accepted_agree |
| 10.1007/s00299-025-03641-4 | 2025 | Introducing chromosome 1Ns from Leymus mollis improves grain protein content of recipient wheat. | AGREE | paper_type=EXP; spatial=NO; panel_scope=UNCLEAR; gemma_final_notes=Pass-A abstract: non-spatial → FILTERED | accepted_agree |
| 10.1007/s00425-016-2517-3 | 2016 | Molecular cytogenetic characterization of a new wheat-rye 1BL•1RS translocation line expressing superior stripe rust resistance and enhanced grain yield. | AGREE | paper_type=EXP; spatial=NO; panel_scope=UNCLEAR; gemma_final_notes=Pass-A abstract: non-spatial → FILTERED | accepted_agree |
| 10.1007/s00432-023-05403-9 | 2023 | Integration and deconvolution methodology deciphering prognosis-related signatures in lung adenocarcinoma | AGREE | Title only; paper_type COMP; spatial blank; panel_scope UNCLEAR; q1 panel-design fields are empty; gemma_final=FILTERED from abstract prefilter. | accepted_agree |
| 10.1007/s00500-023-08435-y | 2023 | RETRACTED ARTICLE: Ensemble recurrent neural network with whale optimization algorithm-based DNA sequence classification for medical applications | AGREE | Title only; paper_type COMP; spatial blank; panel_scope UNCLEAR; q1 panel-design fields are empty; gemma_final=FILTERED from abstract prefilter. | accepted_agree |
| 10.1007/s00702-020-02286-9 | 2021 | Promise and challenges of dystonia brain banking: establishing a human tissue | AGREE | paper_type=OTHER; spatial=NO; panel_scope=UNCLEAR; | accepted_agree |

| DOI | Year | Title | Audit decision | Evidence (verbatim quote) | Validation status / scope flag |
|---|---|---|---|---|---|
| | | repository for studies of X-Linked Dystonia-Parkinsonism. | | gemma_final_notes=Pass-A abstract: OTHER → FILTERED | |
| 10.1007/s10142-024-01415-x | 2024 | A review on advancements in feature selection and feature extraction for high-dimensional NGS data analysis | AGREE | title: A review on advancements in feature selection and feature extraction for high-dimensional NGS data analysis; q0 doc_type=REVIEW; q1 panel-design fields are empty; citation prior labels=REV and Gemma_final=FILTERED. | accepted_agree |
| 10.1007/s10565-025-10037-3 | 2025 | Activation of CaMKII+ neurons in the paramedian raphe nucleus promotes general anesthesia in male mice | AGREE | paper_type=EXP; spatial=NO; panel_scope=UNCLEAR; gemma_final_notes=Pass-A abstract: non-spatial → FILTERED | accepted_agree |
| 10.1007/s10571-024-01484-3 | 2024 | A Bibliometric Analysis of the Spatial Transcriptomics Literature from 2006 to 2023 | AGREE | citation_screening_prior.labels=REV; citation_screening_prior.tools=PERSIST; gemma_final_notes=citation-screening prior pre-filter: labels=['REV']; exp_used_tools=['(none)']; all_tools=['PERSIST'] | accepted_agree |
| 10.1007/s10722-025-02391-2 | 2025 | Analysis of the molecular karyotype of Kengyilia grandiglumis on the Qinghai-Tibet Plateau via sequential FISH and GISH | AGREE | paper_type=EXP; spatial=NO; panel_scope=UNCLEAR; gemma_final_notes=Pass-A abstract: non-spatial → FILTERED | accepted_agree |
| 10.1007/s10815-024-03143-4 | 2024 | Rescue in vitro maturation using ovarian support cells of human oocytes from conventional stimulation cycles yields oocytes with improved nuclear maturation and transcriptomic resemblance to in vivo matured oocytes | AGREE | paper.title='Rescue in vitro maturation using ovarian support cells of human oocytes from conventional stimulation cycles yields oocytes with improved nuclear maturation and transcriptomic resemblance to in vivo matured oocytes'; gemma_q2_paper_profile.spatial=NO; gemma_final=FILTERED. | accepted_agree |
| 10.1007/s11032-024-01493-6 | 2024 | Introgression of tetraploid <i>Thinopyrum elongatum</i> 6EL segments enhances the stripe rust resistance of adult wheat plants. | AGREE | paper_type=EXP; spatial=NO; panel_scope=UNCLEAR; gemma_final_notes=Pass-A abstract: non-spatial → FILTERED | accepted_agree |

| DOI | Year | Title | Audit decision | Evidence (verbatim quote) | Validation status / scope flag |
|---|---|---|---|---|---|
| 10.1007/s11427-023-2561-0 | 2024 | Single-cell omics: experimental workflow, data analyses and applications | AGREE | citation_screening_prior: labels=REV; gemma_final source=citation_screening_prefilter -> FILTERED; q1 panel-design fields are empty. | accepted_agree |
| 10.1007/s12032-025-03042-3 | 2025 | Next-generation oncology: integrative therapeutic frontiers at the crossroads of precision genomics, immuno-engineering, and tumor microenvironment modulation | NEEDS_MANUAL | Final manual decision: FILTERED because no local PDF was available and the packet did not provide usable final gene-panel source evidence. Original evidence: paper_type=OTHER; spatial=YES; panel_scope=TARGETED; q1 panel-design fields are empty; gemma_final=FILTERED from abstract prefilter. | manual_final_filter_after_pdf_review |
| 10.1007/s12035-020-01910-9 | 202 | Sub-region-Specific Optic Nerve Head Glial Activation in Glaucoma | AGREE | paper_type=EXP; spatial=NO; panel_scope=UNCLEAR; q1 panel-design fields are empty; title is not a spatial panel-design study. | accepted_agree |
| 10.1007/s12288-020-01287-8 | 202 | A Novel PDGFRβ Rearrangement in Systemic Mastocytosis with an Associated Haematologic Neoplasm | AGREE | paper_type/doc_type evidence: EDITORIAL ((paraphrased) CORRESPONDENCE<br>Dear Editor,); gemma_final notes: Q0: EDITORIAL → FILTERED | accepted_agree |
| 10.1007/s12539-025-00762-y | 2025 | Clustering Single-Cell RNA-Seq Data with Low-Rank Matrix Factorization and Local Graph Regularization | AGREE | paper.title='Clustering Single-Cell RNA-Seq Data with Low-Rank Matrix Factorization and Local Graph Regularization'; citation_screening_prior.labels='COMP'; gemma_final.classification='FILTERED'; gemma_q1_panel_design blank; gemma_q2_paper_profile={'evidence_scope': '', 'evidence_spatial': '', 'evidence_type': '', 'panel_scope': '', 'paper_type': 'COMP', 'profile_confidence': '', 'spatial': ''} | accepted_agree |
| 10.1007/s12672-024-01614-9 | 2024 | Computer-aided analysis reveals metallothionein-positive cancer-associated fibroblasts promote angiogenesis in gastric adenocarcinoma | AGREE | gemma_q1_panel_design = NOT FOUND and gemma_q2_panel_scope = NOT_TARGETED. | accepted_agree |
| 10.1007/s40472-024-00450-8 | 2024 | Single Cell Sequencing and Spatial Transcriptomics in Kidney | NEEDS_MANUAL | PDF: kidney-transplant review; discusses Visium and other whole- | pdf_validated_unclear_update |

| DOI | Year | Title | Audit decision | Evidence (verbatim quote) | Validation status / scope flag |
|---|---|---|---|---|---|
| | | Transplantation | | transcriptome methods. | |
| 10.1007/s40620-025-02233-5 | 2025 | Identification of conserved gene expression changes across common glomerular diseases by spatial transcriptomics. | AGREE | paper_type=EXP; spatial=YES; panel_scope=NOT_TARGETED; q1 panel-design fields are empty; gemma_final=FILTERED. | accepted_agree |
| 10.1016/b978-0-12-814919-5.00012-9 | 2019 | Single-Cell Transcriptomics: Technology and Applications | AGREE | paper_type/doc_type evidence: REVIEW (CHAPTER 12 Single-Cell Transcriptomics: Technology and Applications); gemma_final notes: Q0: REVIEW → FILTERED | accepted_agree |
| 10.1016/bs.acr.2024.06.006 | 2024 | Current computational methods for spatial transcriptomics in cancer biology | AGREE | paper.title='Current computational methods for spatial transcriptomics in cancer biology'; gemma_q0_document_type=REVIEW; gemma_final=FILTERED. | accepted_agree |
| 10.1016/bs.mcb.2025.10.012 | 2026 | Utilisation of spatial methodologies for the investigation of the tumour microenvironment | AGREE | gemma_final: Pass-A abstract prefilter -> FILTERED; q1 panel-design fields are empty; q2 paper_type=OTHER, spatial=YES, panel_scope=TARGETED. | accepted_agree |
| 10.1016/j.acha.2023.03.001 | 2023 | Lie PCA: Density estimation for symmetric manifolds | AGREE | title='Lie PCA: Density estimation for symmetric manifolds'; citation_screening_prior.labels='COMP'; gemma_final.source='citation_screening_prefilter' | accepted_agree |
| 10.1016/j.ajpath.2026.01.007 | 2026 | Redefining Pathobiology with Spatial Multi-Omics at the Intersection of Biology, Computation, and Histopathological Assessment | AGREE | title: Redefining Pathobiology with Spatial Multi-Omics at the Intersection of Biology, Computation, and Histopathological Assessment; q0 doc_type=REVIEW; q1 panel-design fields are empty; citation prior labels=REV and Gemma_final=FILTERED. | accepted_agree |
| 10.1016/j.ajt.2025.07.057 | 2025 | Urinary Single-Cell RNA Sequencing from a Pig Kidney to Living Human Xenotransplantation Reveals Dynamic Cellular Changes | AGREE | paper_type and q0 indicate NON_ARTICLE; q1 panel-design fields are empty; gemma_final=FILTERED. | accepted_agree |
| 10.1016/j.annonc.2023.09.1306 | 2023 | 2278P Endothelial cell heterogeneity defined by single-cell spatial transcriptomic analysis of breast cancers | AGREE | q0=NON_ARTICLE; q1 panel-design fields are empty; gemma_final=FILTERED. | accepted_agree |

| DOI | Year | Title | Audit decision | Evidence (verbatim quote) | Validation status / scope flag |
|---|---|---|---|---|---|
| 10.1016/j.apsusc.2021.151061 | 2021 | Dissociation of air pollutants on the uniform surface of pentagonal BeP2 | AGREE | Title: 'Dissociation of air pollutants on the uniform surface of pentagonal BeP2'; packet fields show paper_type=COMP and Gemma's FILTERED prefilter. | accepted_agree |
| 10.1016/j.ard.2025.05.024 | 2025 | OP0001 COMPARATIVE ANALYSIS OF SYNOVIAL TISSUE TOPOGRAPHY BY SINGLE CELL TRANSCRIPTOMICS AND SPATIAL TRANSCRIPTOMICS REVEALS DISCRETE AND OVERLAPPING DISEASE PATHWAYS DRIVING CHECKPOINT INHIBITOR-INDUCED AND SPONTANEOUS INFLAMMATORY ARTHRITIS | AGREE | q0=NON_ARTICLE; q1 panel-design fields are empty; gemma_final=FILTERED. | accepted_agree |
| 10.1016/j.ard.2025.05.032 | 2025 | OP0009 SPATIAL TRANSCRIPTOMIC-BASED PHENOTYPING OF THE FIBROBLAST NICHES IN SYSTEMIC SCLEROSIS-ASSOCIATED PRIMARY HEART INVOLVEMENT | AGREE | q0=NON_ARTICLE; q1 panel-design fields are empty; gemma_final=FILTERED. | accepted_agree |
| 10.1016/j.ard.2025.05.638 | 2025 | POS0251 PROFIBROTIC MONOCYTE-DERIVED ALVEOLAR MACROPHAGES ARE ASSOCIATED WITH DISEASE SEVERITY IN PATIENTS WITH SYSTEMIC SCLEROSIS-ASSOCIATED INTERSTITIAL LUNG DISEASE | AGREE | q0=NON_ARTICLE; q1 panel-design fields are empty; gemma_final=FILTERED. | accepted_agree |
| 10.1016/j.array.2025.100623 | 2025 | Two-stage copula-driven, structure-aware gene selection for single-cell analysis | AGREE | paper.title='Two-stage copula-driven, structure-aware gene selection for single-cell analysis'; citation_screening_prior.labels='COMP-BENCH'; gemma_final.classification='FILTERED'; gemma_q1_panel_design blank; gemma_q2_paper_profile={'evidence_scope': '', 'evidence_spatial': '', 'evidence_type': '', 'panel_scope': '', 'paper_type': 'COMP', 'profile_confidence': '', 'spatial': ''} | accepted_agree |
| 10.1016/j.asoc.2025.114110 | 2025 | Metaheuristic-driven feature selection with dual feedback | AGREE | paper.title='Metaheuristic-driven feature selection with dual | accepted_agree |

| DOI | Year | Title | Audit decision | Evidence (verbatim quote) | Validation status / scope flag |
|---|---|---|---|---|---|
| | | mechanisms for imbalanced cancer classification | | feedback mechanisms for imbalanced cancer classification'; citation_screening_prior.labels='COMP'; gemma_final.classification='FILTERED'; gemma_q1_panel_design blank; gemma_q2_paper_profile={'evidence_scope': '', 'evidence_spatial': '', 'evidence_type': '', 'panel_scope': '', 'paper_type': 'COMP', 'profile_confidence': '', 'spatial': ''} | |
| 10.1016/j.atherosclerosis.2022.06.322 | 2022 | Single-cell and spatially resolved transcriptome analysis reveals cellular heterogeneities and novel regulators of atherosclerotic plaque destabilization | AGREE | q0=NON_ARTICLE (POSTER VIEWING SESSION); q1 panel-design fields are empty; gemma_final=FILTERED. | accepted_agree |
| 10.1016/j.atherosclerosis.2023.06.725 | 2023 | Disease-associated cell states in atherosclerosis defined by spatial transcriptomics | AGREE | q0=NON_ARTICLE; q1 panel-design fields are empty; gemma_final=FILTERED. | accepted_agree |
| 10.1016/j.bbadis.2023.166985 | 2023 | The in situ transcriptomic landscape of breast tumour-associated and normal adjacent endothelial cells | AGREE | paper_type=EXP, panel_scope=NOT_TARGETED, spatial=YES; the WTA panel consisted of 18,676 total targets. | accepted_agree \| Fig. 2: excluded (platform/scope) |
| 10.1016/j.bbadis.2025.167918 | 2025 | Deciphering the regulatory networks of human male germline development from embryo to adulthood | AGREE | paper.title='Deciphering the regulatory networks of human male germline development from embryo to adulthood'; citation_screening_prior.labels='NON-ST'; gemma_final.classification='FILTERED'; gemma_q1_panel_design blank; gemma_q2_paper_profile={'evidence_scope': '', 'evidence_spatial': '', 'evidence_type': '', 'panel_scope': '', 'paper_type': '', 'profile_confidence': '', 'spatial': 'NO'} | accepted_agree |
| 10.1016/j.bbrc.2025.151687 | 2025 | Nerve infiltration of bladder cancer predicts response to immunotherapy | AGREE | citation_screening_prior.labels=NON-ST; gemma_final.classification=FILTERED; gemma_q1_panel_design blank. | accepted_agree |
| 10.1016/j.biosystems.2024.105142 | 2024 | Single-cell classification, analysis, and its application using deep | AGREE | paper.title='Single-cell classification, analysis, and its | accepted_agree |

| DOI | Year | Title | Audit decision | Evidence (verbatim quote) | Validation status / scope flag |
|---|---|---|---|---|---|
| | | learning techniques | | application using deep learning techniques'; gemma_q0_document_type=REVIEW; gemma_final=FILTERED. | |
| 10.1016/j.biotechadv.2026.108807 | 2026 | Computational methods for spatial multi-omics integration | AGREE | citation_screening_prior: labels=REV; gemma_final source=citation_screening_prefilter -> FILTERED; q1 panel-design fields are empty. | accepted_agree |
| 10.1016/j.bpj.2018.11.105 | 2019 | In Situ Imaging of Transcriptome and Genome in Single Cells | AGREE | q0=NON_ARTICLE; q1 panel-design fields are empty; gemma_final=FILTERED. | accepted_agree |
| 10.1016/j.bpj.2018.11.2796 | 2019 | A Cartographic View of Membrane Targeting and Association of the C2 Domain from Protein Kinase C | AGREE | q0=NON_ARTICLE; q1 panel-design fields are empty; gemma_final=FILTERED. | accepted_agree |
| 10.1016/j.bpj.2019.11.2578 | 202 | Large-Scale Species-Specific Microbial Identification by Fluorescence In Situ Hybridization | AGREE | q0=NON_ARTICLE; q1 panel-design fields are empty; gemma_final=FILTERED. | accepted_agree |
| 10.1016/j.bpj.2021.11.1246 | 2022 | Molecular cartography: charting the sea of molecular organization in live synapses with nanoscale precision | AGREE | doc_type=NON_ARTICLE; no panel-design evidence is present. | accepted_agree |
| 10.1016/j.bpj.2023.11.3284 | 2024 | Single-virus transcriptomics of bacteriophage lambda | AGREE | q0=NON_ARTICLE; q1 panel-design fields are empty; gemma_final=FILTERED. | accepted_agree |
| 10.1016/j.bpj.2023.11.3292 | 2024 | Single-molecule spatial transcriptome profiling of the mammalian heart | AGREE | q0=NON_ARTICLE; q1 panel-design fields are empty; gemma_final=FILTERED. | accepted_agree |
| 10.1016/j.brainres.2025.149688 | 2025 | Landscape of spatial disruption of homeostasis in the middle temporal gyrus of epileptic patients | AGREE | citation_screening_prior.labels=COMP; gemma_q2_paper_profile.spatial=NO; gemma_final.classification=FILTERED. | accepted_agree |
| 10.1016/j.cancergen.2019.04.058 | 2019 | 52. A rare (7;12) translocation resulting in a rearrangement of the IKZF1 locus with concurrent deletion of CDKN2A, CDKN2B and PAX5 loci: an unannotated genetic abnormality in pediatric B-lymphoblastic leukemia/lymphoma | AGREE | q0=NON_ARTICLE; q1 panel-design fields are empty; gemma_final=FILTERED. | accepted_agree |
| 10.1016/j.ccell.2023.01.010 | 2023 | Spatial profiling technologies illuminate the tumor microenvironment | AGREE | title='Spatial profiling technologies illuminate the tumor microenvironment'; | accepted_agree |

| DOI | Year | Title | Audit decision | Evidence (verbatim quote) | Validation status / scope flag |
|---|---|---|---|---|---|
| | | | | citation_screening_prior.labels='REV'; gemma_final.source='citation_screening_prefilter' | |
| 10.1016/j.ccell.2025.01.001 | 2025 | Contrasting cytotoxic and regulatory T cell responses underlying distinct clinical outcomes to anti-PD-1 plus lenvatinib therapy in cancer | AGREE | citation_screening_prior.labels=NON-ST; gemma_final.classification=FILTERED; gemma_q1_panel_design blank. | accepted_agree |
| 10.1016/j.ceb.2024.102358 | 2024 | Molecular cartography within axons | AGREE | paper_type=OTHER; spatial=NO; panel_scope=UNCLEAR; q1 panel-design fields are empty; gemma_final=FILTERED. | accepted_agree |
| 10.1016/j.cell.2017.03.008 | 2017 | Tridimensional Visualization and Analysis of Early Human Development. | AGREE | paper_type=EXP; spatial=NO; panel_scope=UNCLEAR; q1 panel-design fields are empty; gemma_final=FILTERED. | accepted_agree |
| 10.1016/j.cell.2017.06.021 | 2017 | One (Cell) in a Million | AGREE | paper_type/doc_type evidence: EDITORIAL (Matthew Pavlovich Editor, Trends in Biotechnology); gemma_final notes: Q0: EDITORIAL → FILTERED | accepted_agree |
| 10.1016/j.cell.2020.06.038 | 202 | Spatial Transcriptomics and In Situ Sequencing to Study Alzheimer's Disease. | AGREE | paper_type=EXP; spatial=YES; panel_scope=NOT_TARGETED; q1 panel-design fields are empty; gemma_final=FILTERED. | accepted_agree |
| 10.1016/j.cell.2022.09.035 | 2022 | Spatially resolved epigenomic profiling of single cells in complex tissues | NEEDS_MANUAL | PDF: spatially resolved epigenomic profiling with selected loci, not a gene-panel design study. | pdf_validated_unclear_update |
| 10.1016/j.cell.2022.10.014 | 2022 | Omics goes spatial epigenomics. | AGREE | Title: Omics goes spatial epigenomics.; paper_type=OTHER; panel_scope=TARGETED; spatial=YES; gemma_final notes say Pass-A abstract: OTHER -> FILTERED. | accepted_agree |
| 10.1016/j.cell.2022.10.017 | 2022 | The phenotypic landscape of essential human genes | AGREE | Title: The phenotypic landscape of essential human genes; paper_type=EXP; spatial=NO; panel_scope=UNCLEAR; gemma_final notes say non-spatial -> FILTERED. | accepted_agree |

| DOI | Year | Title | Audit decision | Evidence (verbatim quote) | Validation status / scope flag |
|---|---|---|---|---|---|
| 10.1016/j.cell.2023.11.006 | 2023 | Molecular cartography uncovers evolutionary and microenvironmental dynamics in sporadic colorectal tumors. | AGREE | paper_type=EXP, spatial=YES, scope=NOT_TARGETED; no panel-design evidence found. | accepted_agree |
| 10.1016/j.cell.2024.07.016 | 2024 | LAG-3 and PD-1 synergize on CD8+ T cells to drive T cell exhaustion and hinder autocrine IFN-γ-dependent anti-tumor immunity | AGREE | paper.title='LAG-3 and PD-1 synergize on CD8+ T cells to drive T cell exhaustion and hinder autocrine IFN-γ-dependent anti-tumor immunity'; gemma_q2_paper_profile.spatial=NO; gemma_final=FILTERED. | accepted_agree |
| 10.1016/j.cell.2025.05.022 | 2025 | Perturb-Multimodal: A platform for pooled genetic screens with imaging and sequencing in intact mammalian tissue | AGREE | Title: Perturb-Multimodal: A platform for pooled genetic screens with imaging and sequencing in intact mammalian tissue; paper_type=COMP; spatial=YES; panel_scope=UNCLEAR; gemma_final notes say COMP -> FILTERED. | accepted_agree |
| 10.1016/j.cell.2025.05.027 | 2025 | STAMP: Single-cell transcriptomics analysis and multimodal profiling through imaging | AGREE | Title: STAMP: Single-cell transcriptomics analysis and multimodal profiling through imaging; paper_type=COMP; spatial=YES; panel_scope=UNCLEAR; gemma_final notes say COMP -> FILTERED. | accepted_agree |
| 10.1016/j.cell.2025.08.023 | 2025 | Microglia-astrocyte crosstalk regulates synapse remodeling via Wnt signaling | AGREE | Title: Microglia-astrocyte crosstalk regulates synapse remodeling via Wnt signaling; paper_type=EXP; spatial=NO; panel_scope=UNCLEAR; gemma_final notes say non-spatial -> FILTERED. | accepted_agree |
| 10.1016/j.cell.2025.08.029 | 2025 | Creeping fat-derived mechanosensitive fibroblasts drive intestinal fibrosis in Crohn's disease strictures | AGREE | Title: Creeping fat-derived mechanosensitive fibroblasts drive intestinal fibrosis in Crohn's disease strictures; paper_type=EXP; spatial=NO; panel_scope=UNCLEAR; gemma_final notes say non-spatial -> FILTERED. | accepted_agree |
| 10.1016/j.celrep.2021.110251 | 2022 | Single-cell analysis of early chick hypothalamic development reveals that hypothalamic cells are | AGREE | Title: Single-cell analysis of early chick hypothalamic development reveals that hypothalamic cells are | accepted_agree |

| DOI | Year | Title | Audit decision | Evidence (verbatim quote) | Validation status / scope flag |
|---|---|---|---|---|---|
| | | induced from prethalamic-like progenitors. | | induced from prethalamic-like progenitors.; paper_type=EXP; spatial=NO; panel_scope=UNCLEAR; gemma_final notes say non-spatial -> FILTERED. | |
| 10.1016/j.celrep.2023.112643 | 2023 | Transcriptomic landscape of early hair follicle and epidermal development | AGREE | title='Transcriptomic landscape of early hair follicle and epidermal development'; citation_screening_prior.labels=''; gemma_final.source='abstract_prefilter' | accepted_agree |
| 10.1016/j.celrep.2023.113121 | 2023 | Distinct mechanisms for sebaceous gland self-renewal and regeneration provide durability in response to injury | AGREE | title='Distinct mechanisms for sebaceous gland self-renewal and regeneration provide durability in response to injury'; citation_screening_prior.labels=''; gemma_final.source='abstract_prefilter' | accepted_agree |
| 10.1016/j.celrep.2025.116125 | 2025 | The neurexin gene family regulates olfactory glomerular formation | AGREE | Title: The neurexin gene family regulates olfactory glomerular formation; paper_type=EXP; spatial=NO; panel_scope=UNCLEAR; gemma_final notes say non-spatial -> FILTERED. | accepted_agree |
| 10.1016/j.celrep.2025.116717 | 2026 | Transcriptomic and spatial organization of mouse spinal cord projection neurons | AGREE | Title: Transcriptomic and spatial organization of mouse spinal cord projection neurons; paper_type=EXP; spatial=NO; panel_scope=UNCLEAR; gemma_final notes say non-spatial -> FILTERED. | accepted_agree |
| 10.1016/j.celrep.2026.117076 | 2026 | Collagenesis orchestrates mechanoadaptive development and homeostasis of intervertebral discs | AGREE | citation_screening_prior: non-spatial prior; gemma_final source=citation_screening_prefilter -> FILTERED; q1 panel-design fields are empty; q2 spatial=NO. | accepted_agree |
| 10.1016/j.cels.2018.03.010 | 2018 | Network Visualization and Analysis of Spatially Aware Gene Expression Data with InsituNet. | AGREE | Title: Network Visualization and Analysis of Spatially Aware Gene Expression Data with InsituNet.; paper_type=COMP; spatial=YES; panel_scope=UNCLEAR; gemma_final notes say COMP -> FILTERED. | accepted_agree |

| DOI | Year | Title | Audit decision | Evidence (verbatim quote) | Validation status / scope flag |
|---|---|---|---|---|---|
| 10.1016/j.cels.2021.09.004 | 2022 | Cell-type modeling in spatial transcriptomics data elucidates spatially variable colocalization and communication between cell-types in mouse brain | AGREE | Title: Cell-type modeling in spatial transcriptomics data elucidates spatially variable colocalization and communication between cell-types in mouse brain; citation_screening_prior says computational-only prior with scGeneFit; paper_type=COMP; gemma_final notes cite the computational-only prefilter. | accepted_agree |
| 10.1016/j.cels.2024.04.006 | 2024 | Accurate single-molecule spot detection for image-based spatial transcriptomics with weakly supervised deep learning. | NEEDS_MANUAL | Final manual decision: FILTERED because the paper is a downstream analysis/modeling/QC method or dataset-use paper, with no final panel-design source. Original evidence: No panel-design evidence is present in the packet. | manual_final_filter_after_pdf_review |
| 10.1016/j.cels.2024.07.001 | 2024 | PanIN and CAF transitions in pancreatic carcinogenesis revealed with spatial data integration | AGREE | Title: PanIN and CAF transitions in pancreatic carcinogenesis revealed with spatial data integration; paper_type=EXP; spatial=YES; panel_scope=NOT_TARGETED; gemma_final notes say not targeted -> FILTERED. | accepted_agree |
| 10.1016/j.cels.2025.101195 | 2025 | Spatial transcriptomics-aided localization for single-cell transcriptomics with STALocator. | AGREE | Title: Spatial transcriptomics-aided localization for single-cell transcriptomics with STALocator.; paper_type=COMP; spatial=YES; panel_scope=UNCLEAR; gemma_final notes say COMP -> FILTERED. | accepted_agree |
| 10.1016/j.cels.2025.101243 | 2025 | SpaGRN: Investigating spatially informed regulatory paths for spatially resolved transcriptomics data. | AGREE | Title: SpaGRN: Investigating spatially informed regulatory paths for spatially resolved transcriptomics data.; paper_type=COMP; spatial=YES; panel_scope=UNCLEAR; gemma_final notes say COMP -> FILTERED. | accepted_agree |
| 10.1016/j.cels.2025.101405 | 2025 | Identifying an optimal perturbation to induce a desired cell state by generative deep learning | AGREE | citation_screening_prior.labels=COMP; gemma_q2_paper_profile.spatial=NO; gemma_final.classification=FILTERED. | accepted_agree |

| DOI | Year | Title | Audit decision | Evidence (verbatim quote) | Validation status / scope flag |
|---|---|---|---|---|---|
| 10.1016/j.cels.2026.101571 | 2026 | Identifying maximally informative signal-aware representations of single-cell data using the information bottleneck | AGREE | citation_screening_prior: computational-only prior; gemma_final source=citation_screening_prefilter -> FILTERED; q1 panel-design fields are empty; q2 paper_type=COMP. | accepted_agree |
| 10.1016/j.chempr.2018.05.014 | 2018 | DNA-Coded Fluorescence for Color Painting RNAs | AGREE | Title: DNA-Coded Fluorescence for Color Painting RNAs; paper_type=COMP; spatial=YES; panel_scope=UNCLEAR; gemma_final notes say COMP -> FILTERED. | accepted_agree |
| 10.1016/j.chom.2026.01.004 | 2026 | Temporal transcriptional rhythms govern coral-symbiont function and microbiome dynamics | AGREE | paper_type=EXP, spatial=NO, scope=UNCLEAR; no panel-design evidence found. | accepted_agree |
| 10.1016/j.cj.2026.02.021 | 2026 | PhytoCell: An ensemble learning framework for identifying cell states in plant scRNA-seq data | AGREE | citation_screening_prior: computational-only prior; gemma_final source=citation_screening_prefilter -> FILTERED; q1 panel-design fields are empty; q2 paper_type=COMP. | accepted_agree |
| 10.1016/j.compbiomed.2021.105208 | 2022 | Ensemble feature selection for stable biomarker identification and cancer classification from microarray expression data | AGREE | Title only; paper_type COMP; spatial blank; panel_scope blank; q1 panel-design fields are empty; gemma_final=FILTERED from citation-screening prefilter. | accepted_agree |
| 10.1016/j.compbiomed.2023.106634 | 2023 | scMAGS: Marker gene selection from scRNA-seq data for spatial transcriptomics studies | AGREE | title: scMAGS: Marker gene selection from scRNA-seq data for spatial transcriptomics studies; q2 paper_type=COMP; q1 panel-design fields are empty; citation prior labels=COMP\|COMP-BENCH and Gemma_final=FILTERED. | accepted_agree |
| 10.1016/j.compbiomed.2025.109873 | 2025 | A novel coarsened graph learning method for scalable single-cell data analysis | AGREE | citation_screening_prior.labels=NON-ST; gemma_final.classification=FILTERED; gemma_q1_panel_design blank. | accepted_agree |
| 10.1016/j.conb.2021.09.011 | 2021 | Transcriptomic evidence for dense peptidergic networks within forebrains of four widely divergent tetrapods | AGREE | Title: 'Transcriptomic evidence for dense peptidergic networks within forebrains of four widely divergent tetrapods'; packet fields show | accepted_agree |

| DOI | Year | Title | Audit decision | Evidence (verbatim quote) | Validation status / scope flag |
| --- | --- | --- | --- | --- | --- |
| | | | | doc_type=REVIEW and Gemma's FILTERED prefilter. | |
| 10.1016/j.crbiot.2023.100127 | 2023 | CRISPR/Cas9 genetic screens in hepatocellular carcinoma gene discovery | AGREE | title='CRISPR/Cas9 genetic screens in hepatocellular carcinoma gene discovery'; citation_screening_prior.labels='REV'; gemma_final.source='citation_screening_prefilter' | accepted_agree |
| 10.1016/j.crmeth.2021.100061 | 2021 | Systematic detection of m6A-modified transcripts at single-molecule and single-cell resolution | AGREE | Title: Systematic detection of m6A-modified transcripts at single-molecule and single-cell resolution; paper_type=EXP; spatial=NO; panel_scope=UNCLEAR; gemma_final notes say non-spatial -> FILTERED. | accepted_agree |
| 10.1016/j.crmeth.2022.100201 | 2022 | Robust and sensitive <i>in situ</i> RNA detection using Yn-situ. | AGREE | Title: Robust and sensitive in situ RNA detection using Yn-situ.; paper_type=COMP; spatial=YES; panel_scope=UNCLEAR; gemma_final notes say COMP -> FILTERED. | accepted_agree |
| 10.1016/j.crmeth.2022.100332 | 2022 | Multiset multicover methods for discriminative marker selection | AGREE | citation_screening_prior: computational-only prior; gemma_final source=citation_screening_prefilter -> FILTERED; q1 panel-design fields are empty; q2 paper_type=COMP. | accepted_agree |
| 10.1016/j.crmeth.2023.100507 | 2023 | An information theoretic approach to detecting spatially varying genes. | NEEDS_MANUAL | PDF: Maxspin is a tool for detecting spatially varying genes; benchmarks Visium/CosMx/MERFISH. | pdf_validated_unclear_update |
| 10.1016/j.crmeth.2023.100573 | 2023 | Three-dimensional molecular cartography of human cerebral organoids revealed by double-barcoded spatial transcriptomics. | AGREE | paper_type=COMP, spatial=YES, scope=NOT_TARGETED; no panel-design evidence found. | accepted_agree |
| 10.1016/j.crmeth.2024.100886 | 2024 | WEST is an ensemble method for spatial transcriptomics analysis | AGREE | Title: WEST is an ensemble method for spatial transcriptomics analysis; paper_type=COMP; spatial=YES; panel_scope=UNCLEAR; gemma_final notes say COMP -> FILTERED. | accepted_agree |

| DOI | Year | Title | Audit decision | Evidence (verbatim quote) | Validation status / scope flag |
|---|---|---|---|---|---|
| 10.1016/j.crmeth.2026.101306 | 2026 | An integrative spatial multi-omic workflow for unified analysis of tumor tissue | AGREE | Title: An integrative spatial multi-omic workflow for unified analysis of tumor tissue; paper_type=COMP; spatial=YES; panel_scope=UNCLEAR; gemma_final notes say COMP -> FILTERED. | accepted_agree |
| 10.1016/j.crmeth.2026.101329 | 2026 | Scalable nonparametric clustering with unified marker gene selection for single-cell RNA-seq data | AGREE | citation_screening_prior labels=COMP and all_tools=RankCorr; gemma_final source=citation_screening_prefilter -> FILTERED; q1 panel-design fields are empty. | accepted_agree |
| 10.1016/j.cropro.2024.106633 | 2024 | Efficacy, persistence and residue levels of fungicides for Botrytis control in wild blueberry | AGREE | abstract_prefilter: non-spatial | accepted_agree |
| 10.1016/j.csbj.2022.11.051 | 2022 | Temporal progress of gene expression analysis with RNA-Seq data: A review on the relationship between computational methods | AGREE | title='Temporal progress of gene expression analysis with RNA-Seq data: A review on the relationship between computational methods'; citation_screening_prior.labels='REV'; gemma_final.source='citation_screening_prefilter' | accepted_agree |
| 10.1016/j.cub.2014.12.059 | 2015 | Oncogenic properties of apoptotic tumor cells in aggressive B cell lymphoma. | AGREE | paper_type=EXP, spatial=YES, scope=NOT_TARGETED; no panel-design evidence found. | accepted_agree |
| 10.1016/j.devcel.2023.11.007 | 2023 | Murine trophoblast organoids as a model for trophoblast development and CRISPR-Cas9 screening | AGREE | title='Murine trophoblast organoids as a model for trophoblast development and CRISPR-Cas9 screening'; citation_screening_prior.labels='NON-ST'; gemma_final.source='citation_screening_prefilter' | accepted_agree |
| 10.1016/j.ecoleng.2025.107526 | 2025 | The effectiveness of a fish-friendly pumping station for critically endangered European eel; An assessment using live eels, fish-mounted sensors and passive sensors | AGREE | abstract_prefilter: non-spatial | accepted_agree |
| 10.1016/j.ejca.2025.115811 | 2025 | Transcriptional memory states of malignant cutaneous T-cell lymphoma cells inform targeted therapy | AGREE | Q0 NON_ARTICLE: Abstracts, EORTC-CLTG 2025 | accepted_agree |

| DOI | Year | Title | Audit decision | Evidence (verbatim quote) | Validation status / scope flag |
|---|---|---|---|---|---|
| 10.1016/j.ejca.2025.115843 | 2025 | Spatial transcriptomic characterization of non-lesional, non-tumor lesional and tumor skin in mycosis fungoides using CosMx spatial molecular imaging | NEEDS_MANUAL | doc_type=NON_ARTICLE; non-article packet with no panel-source evidence. Original audit evidence: Q0 NON_ARTICLE abstract; title mentions CosMx spatial molecular imaging; q1 panel fields are empty | validated_medium_risk_manual_filtered |
| 10.1016/j.ejso.2025.110918 | 2025 | Spatial Transcriptomics Analysis of Histopathological Growth Patterns in Colorectal Liver Metastases Using In Situ Sequencing | NEEDS_MANUAL | doc_type=NON_ARTICLE; non-article packet with no panel-source evidence. Original audit evidence: Q0 NON_ARTICLE abstract; title mentions in situ sequencing; q1 panel fields are empty | validated_medium_risk_manual_filtered |
| 10.1016/j.ekir.2021.01.037 | 2021 | Pharmacokinetics of Vancomycin in Pediatric Patients Receiving Intermittent Hemodialysis or Hemodiafiltration | AGREE | abstract_prefilter: non-spatial | accepted_agree |
| 10.1016/j.engappai.2025.113687 | 2026 | CrossToGene: A bidirectional cross-modality interaction framework for predicting spatial gene expression from histopathological images | AGREE | abstract_prefilter: COMP; title is a prediction framework, not panel selection | accepted_agree |
| 10.1016/j.envint.2021.106893 | 2022 | Dose-response metabolomics and pathway sensitivity to map molecular cartography of bisphenol A exposure. | AGREE | abstract_prefilter: non-spatial | accepted_agree |
| 10.1016/j.esmoop.2025.105715 | 2025 | 137P Multidimensional characterization of tumor microenvironment heterogeneity by HER2 status in gastric cancer | AGREE | Q0 NON_ARTICLE: abstracts | accepted_agree |
| 10.1016/j.esmoop.2025.105716 | 2025 | 138P Prognostic analysis of KRAS mutations in metastasic colorectal cancer: A real-world cohort study | AGREE | Q0 NON_ARTICLE: abstracts | accepted_agree |
| 10.1016/j.euroneuro.2024.08.339 | 2024 | T29. SPATIAL-MOLECULAR SEX DIFFERENCES IN THE HUMAN VENTROMEDIAL AND ARCUATE HYPOTHALAMUS | NEEDS_MANUAL | doc_type=NON_ARTICLE; non-article packet with no panel-source evidence. Original audit evidence: Q0 NON_ARTICLE abstract; title indicates spatial-molecular study; q1 panel fields are empty | validated_medium_risk_manual_filtered |
| 10.1016/j.euroneuro.2025.08.508 | 2025 | TRANSCRIPTOMIC AND CHROMATIN DYNAMICS OF THE HUMAN PTSD BRAIN AT SINGLE CELL RESOLUTION | AGREE | Q0 NON_ARTICLE: Abstracts | accepted_agree |
| 10.1016/j.fertnstert.2023.08.744 | 2023 | USING IN SITU GENE EXPRESSION TO CREATE A | NEEDS_MANUAL | doc_type=NON_ARTICLE; non-article packet with no panel-source | validated_medium_risk_manual_filtered |

| DOI | Year | Title | Audit decision | Evidence (verbatim quote) | Validation status / scope flag |
|---|---|---|---|---|---|
| | | THREE-DIMENSIONAL MAP OF HUMAN BLASTOCYSTS | | evidence. Original audit evidence: Q0 NON_ARTICLE oral abstract; title mentions in situ gene expression; q1 panel fields are empty | |
| 10.1016/j.forsciint.2025.112646 | 2025 | Single-nucleus RNA sequencing and machine learning identify CACNA1A as a myocyte-specific biomarker for sudden unexplained death in schizophrenia | AGREE | citation_screening_prior.labels=NON-ST; gemma_final.classification=FILTERED; gemma_q1_panel_design blank. | accepted_agree |
| 10.1016/j.gendis.2024.101205 | 2024 | In situ transcriptomic analysis of spermatocytes in non-obstructive azoospermia reveals senescence-like states in arrested spermatocytes | NEEDS_MANUAL | spatial=NO; packet does not support an in-scope spatial panel-design record. Original audit evidence: q1 panel_design: NOT FOUND; q2 panel_scope: UNCLEAR; q2 spatial: NO. | validated_high_risk_manual_filtered |
| 10.1016/j.heliyon.2023.e15306 | 2023 | TissUUmaps 3: Improvements in interactive visualization, exploration, and quality assessment of large-scale spatial omics data. | AGREE | abstract_prefilter: COMP; title is TissUUmaps 3 visualization tool | accepted_agree |
| 10.1016/j.ibneur.2023.08.2132 | 2023 | SPATIALLY RESOLVED SINGLE-CELL GENOMICS & MOLECULAR AND CELLULAR ARCHITECTURE OF THE BRAIN | AGREE | Title: 'SPATIALLY RESOLVED SINGLE-CELL GENOMICS & MOLECULAR AND CELLULAR ARCHITECTURE OF THE BRAIN'; packet fields show Q0: NON_ARTICLE and Gemma's FILTERED prefilter. | accepted_agree |
| 10.1016/j.identj.2025.105294 | 2025 | P21/P16 Senescence Heterogeneity In Diabetic Bone | AGREE | Q0 NON_ARTICLE: Aim or purpose text is unrelated to panel design | accepted_agree |
| 10.1016/j.identj.2025.106075 | 2025 | Construction And Mechanism Exploration Of Tumor Budding Organoids (TBO) | AGREE | Q0 NON_ARTICLE: Aim or purpose text is unrelated to panel design | accepted_agree |
| 10.1016/j.identj.2025.106129 | 2025 | Lycium Barbarum Glycopetide Remodeling Periodontal Immunopathology By Macrophage Metabolic Reprogramming | AGREE | abstract_prefilter: non-spatial | accepted_agree |
| 10.1016/j.ijrobp.2025.06.3281 | 2025 | Integrating Histopathology and Spatial Transcriptomics for Tumor Microenvironment Analysis and Personalized Radiotherapy | NEEDS_MANUAL | doc_type=NON_ARTICLE; non-article packet with no panel-source evidence. Original audit evidence: Q0 NON_ARTICLE poster Q&A abstract; q1 panel fields are empty | validated_medium_risk_manual_filtered |

| DOI | Year | Title | Audit decision | Evidence (verbatim quote) | Validation status / scope flag |
|---|---|---|---|---|---|
| 10.1016/j.imbio.2025.153071 | 2025 | Spatial transcriptomics maps complement-driven cell-to-cell interactions during acute kidney injury | NEEDS_MANUAL | panel_scope=NOT_TARGETED with no panel-source evidence; untargeted/no-panel records are out of scope. Original audit evidence: abstract_prefilter: not targeted; title is spatial transcriptomics study; q1 panel fields are empty | validated_medium_risk_manual_filtered |
| 10.1016/j.iotech.2025.101343 | 2025 | 346P Deciphering immune resistance mechanisms in NSCLC through patient-specific immunotherapy profiling | AGREE | Q0 NON_ARTICLE: abstracts | accepted_agree |
| 10.1016/j.irbm.2018.04.004 | 2018 | Improvements of Liver MR Imaging Clinical Protocols to Simultaneously Quantify Steatosis and Iron Overload | AGREE | gemma_final=FILTERED; paper_type=EXP; spatial=NO; panel_scope=UNCLEAR; title=Improvements of Liver MR Imaging Clinical Protocols to Simultaneously Quantify Steatosis and Iron Overload | accepted_agree |
| 10.1016/j.isci.2021.103292 | 2021 | How many markers are needed to robustly determine a cell's type? | AGREE | Title: 'How many markers are needed to robustly determine a cell's type?'; packet fields show paper_type=COMP, spatial blank/NO, and Gemma's FILTERED prefilter. | accepted_agree |
| 10.1016/j.isci.2025.114328 | 2025 | United multi-omics and machine learning refine regulatory T cell-defined hepatocellular carcinoma subtypes | AGREE | citation_screening_prior.labels=NON-ST; gemma_final.classification=FILTERED; gemma_q1_panel_design blank. | accepted_agree |
| 10.1016/j.jaci.2022.03.034 | 2022 | Distinct disease-specific Tfh cell populations in 2 different fibrotic diseases: IgG<sub>4</sub>-related disease and Kimura disease. | AGREE | The Visium Spatial Gene Expression Slide & Reagent Kit was used to generate sequencing libraries; q2 panel_scope is NOT_TARGETED. | accepted_agree |
| 10.1016/j.jare.2025.05.038 | 2025 | Dynamic heterogeneity towards drug resistance in AML cells is primarily driven by epigenomic mechanism unveiled by multi-omics analysis | AGREE | citation_screening_prior.labels=NON-ST; gemma_final.classification=FILTERED; gemma_q1_panel_design blank. | accepted_agree |
| 10.1016/j.jbc.2023.104751 | 2023 | A DNAzyme-enhanced nonlinear hybridization chain reaction for sensitive detection of microRNA. | AGREE | gemma_final=FILTERED; paper_type=EXP; spatial=NO; panel_scope=UNCLEAR; title=A DNAzyme-enhanced nonlinear | accepted_agree |

| DOI | Year | Title | Audit decision | Evidence (verbatim quote) | Validation status / scope flag |
|---|---|---|---|---|---|
| | | | | hybridization chain reaction for sensitive detection of microRNA. | |
| 10.1016/j.jbi.2023.104510 | 2023 | MAGNETO: Cell type marker panel generator from single-cell transcriptomic data | AGREE | title: MAGNETO: Cell type marker panel generator from single-cell transcriptomic data; q2 paper_type=COMP; q1 panel-design fields are empty; citation prior labels=COMP\|COMP-BENCH and Gemma_final=FILTERED. | accepted_agree |
| 10.1016/j.jcmgh.2025.101590 | 2025 | Single Cell Profiling in the Sox10<sup>Dom</sup> Hirschsprung Mouse Implicates Hox Genes in Enteric Neuron Trajectory Allocation. | AGREE | gemma_final=FILTERED; paper_type=EXP; spatial=NO; panel_scope=UNCLEAR; title=Single Cell Profiling in the Sox10 Dom Hirschsprung Mouse Implicates Hox Genes in Enteric Neuron Trajectory Allocation. | accepted_agree |
| 10.1016/j.jgg.2016.09.006 | 2016 | A functional centromere lacking CentO sequences in a newly formed ring chromosome in rice. | AGREE | gemma_final=FILTERED; paper_type=EXP; spatial=NO; panel_scope=UNCLEAR; title=A functional centromere lacking CentO sequences in a newly formed ring chromosome in rice. | accepted_agree |
| 10.1016/j.jgg.2023.04.001 | 2023 | VT3D: a visualization toolbox for 3D transcriptomic data. | AGREE | gemma_final=FILTERED; paper_type=COMP; spatial=YES; panel_scope=UNCLEAR; title=VT3D: a visualization toolbox for 3D transcriptomic data. | accepted_agree |
| 10.1016/j.jgg.2024.11.005 | 2024 | Integrated analysis and systematic characterization of the regulatory network for human germline development | AGREE | paper.title='Integrated analysis and systematic characterization of the regulatory network for human germline development'; gemma_q2_paper_profile.spatial=NO; gemma_final=FILTERED. | accepted_agree |
| 10.1016/j.jgg.2024.11.009 | 2025 | KanCell: dissecting cellular heterogeneity in biological tissues through integrated single-cell and spatial transcriptomics. | AGREE | gemma_final=FILTERED; paper_type=COMP; spatial=YES; panel_scope=UNCLEAR; title=KanCell: dissecting cellular heterogeneity in biological tissues through integrated single-cell and spatial transcriptomics. | accepted_agree |

| DOI | Year | Title | Audit decision | Evidence (verbatim quote) | Validation status / scope flag |
|---|---|---|---|---|---|
| 10.1016/j.jhazmat.2025.137375 | 2025 | Insight into perfluorooctanoic acid-induced impairment of mouse embryo implantation via single-cell RNA-seq | AGREE | citation_screening_prior.labels=COMP; gemma_q2_paper_profile.spatial=NO; gemma_final.classification=FILTERED. | accepted_agree |
| 10.1016/j.jid.2023.06.211 | 2023 | Cross-Comparison of Inflammatory Skin Disease Transcriptomics Identifies PTEN as a Pathogenic Disease Classifier in Cutaneous Lupus Erythematosus | AGREE | title='Cross-Comparison of Inflammatory Skin Disease Transcriptomics Identifies PTEN as a Pathogenic Disease Classifier in Cutaneous Lupus Erythematosus'; citation_screening_prior.labels='NON-ST'; gemma_final.source='citation_screening_prefilter' | accepted_agree |
| 10.1016/j.jid.2024.07.006 | 2024 | How Can Spatial Transcriptomic Profiling Advance Our Understanding of Skin Diseases? | AGREE | paper.title='How Can Spatial Transcriptomic Profiling Advance Our Understanding of Skin Diseases?'; gemma_q0_document_type=REVIEW; gemma_final=FILTERED. | accepted_agree |
| 10.1016/j.jid.2025.06.1542 | 2025 | LB1244 Spatial transcriptomics analysis of Merkel cell carcinoma reveals novel immune cell subsets and hypoxia as correlates of immunotherapy response | AGREE | gemma_final=FILTERED; doc_type=; spatial=; panel_scope=; title=LB1244 Spatial transcriptomics analysis of Merkel cell carcinoma reveals novel immune cell subsets and hypoxia as correlates of immunotherapy response | accepted_agree |
| 10.1016/j.jmb.2026.169684 | 2026 | Rising Star: Combining Bioinformatics and Experimental Biology to Decoding Non-coding RNAs and RNA Modifications | AGREE | citation_screening_prior labels=REV and all_tools=COSG; gemma_final source=citation_screening_prefilter -> FILTERED; q1 panel-design fields are empty. | accepted_agree |
| 10.1016/j.jmb.2026.169781 | 2026 | IMDD: A Database for Exploring Tissue-Specific Gene Expression Dynamics During Holometabolous Insects | AGREE | gemma_final source=abstract_prefilter -> FILTERED; q2 paper_type=OTHER, spatial=NO, panel_scope=UNCLEAR; q1 panel-design fields are empty. | accepted_agree |
| 10.1016/j.jneumeth.2025.110548 | 2025 | Immuno-laser capture microdissection of perfusion-fixed mouse brain tissue coupled to RNA-seq. | AGREE | gemma_final=FILTERED; paper_type=OTHER; spatial=YES; panel_scope=NOT_TARGETED; title=Immuno-laser capture microdissection of perfusion-fixed | accepted_agree |

| DOI | Year | Title | Audit decision | Evidence (verbatim quote) | Validation status / scope flag |
|---|---|---|---|---|---|
| | | | | mouse brain tissue coupled to RNA-seq. | |
| 10.1016/j.jobcr.2025.10.015 | 2025 | Artificial Intelligence–driven spatial transcriptomics in OSCC: Mapping the tumor microenvironment and personalizing therapy | AGREE | gemma_final=FILTERED; paper_type=OTHER; spatial=YES; panel_scope=UNCLEAR; title=Artificial Intelligence-driven spatial transcriptomics in OSCC: Mapping the tumor microenvironment and personalizing therapy | accepted_agree |
| 10.1016/j.jpain.2024.01.050 | 2024 | Cell-Type Map of Pain and Non-Pain Somatosensory States in the Spinal Cord | AGREE | gemma_final=FILTERED; doc_type=NON_ARTICLE; title=Cell-Type Map of Pain and Non-Pain Somatosensory States in the Spinal Cord | accepted_agree |
| 10.1016/j.jtho.2016.11.1778 | 2017 | P3.02b-110 ROS1 Translocation as a Bystander Mutation in T790M EGFR Mutated NSCLC | AGREE | gemma_final=FILTERED; doc_type=NON_ARTICLE; title=P3.02b-110 ROS1 Translocation as a Bystander Mutation in T790M EGFR Mutated NSCLC | accepted_agree |
| 10.1016/j.jtho.2017.09.1549 | 2017 | P3.02-019 Clinical Validation of a Real Time PCR Assay for the Detection of ROS1 Fusion in Chinese Non-Small Cell Lung Cancer | AGREE | gemma_final=FILTERED; doc_type=NON_ARTICLE; title=P3.02-019 Clinical Validation of a Real Time PCR Assay for the Detection of ROS1 Fusion in Chinese Non-Small Cell Lung Cancer | accepted_agree |
| 10.1016/j.jtho.2019.08.569 | 2019 | MA09.06 Adaptive Mechanisms of Resistance to Targeted Therapy in EGFR Mutant Brain Metastasis | AGREE | gemma_final=FILTERED; doc_type=NON_ARTICLE; title=MA09.06 Adaptive Mechanisms of Resistance to Targeted Therapy in EGFR Mutant Brain Metastasis | accepted_agree |
| 10.1016/j.kint.2025.10.005 | 2026 | REMODELing mechanistic trials for kidney disease: a multimodal, tissue-centered approach to understand the renal mechanism of action of semaglutide | AGREE | gemma_final=FILTERED; paper_type=OTHER; spatial=NO; panel_scope=UNCLEAR; title=REMODELing mechanistic trials for kidney disease: a multimodal, tissue-centered approach to understand the renal mechanism of action of semaglutide | accepted_agree |

| DOI | Year | Title | Audit decision | Evidence (verbatim quote) | Validation status / scope flag |
|---|---|---|---|---|---|
| 10.1016/j.knosys.2025.114149 | 2025 | Dual-consistency graph-spectral embedding joint learning for multi-view single-cell clustering | AGREE | gemma_final.source=abstract_prefilter; gemma_final.notes=non-spatial; gemma_q2_paper_profile.panel_scope=UNCLEAR. | accepted_agree |
| 10.1016/j.labinv.2025.104190 | 2025 | Rigor and Reproducibility of Spatial Transcriptomics Performed on Clinically Sourced Human Tissues | AGREE | gemma_q2_paper_profile.panel_scope=NOT_TARGETED; gemma_q2_paper_profile.paper_type=EXP; gemma_final.classification=FILTERED. | accepted_agree |
| 10.1016/j.labinv.2025.105793 | 2026 | 1495 KDM6A deficiency in urine urothelial cells can be identified by immunocytochemistry and is associated with increased bladder cancer risk | AGREE | gemma_final=FILTERED; paper_type=EXP; spatial=UNKNOWN; panel_scope=UNCLEAR; title=KDM6A deficiency in urine urothelial cells can be identified by immunocytochemistry and is associated with increased bladder cancer risk | accepted_agree |
| 10.1016/j.lfs.2024.123236 | 2024 | Integrating multi-omics data reveals neuroblastoma subtypes in the tumor microenvironment | AGREE | paper.title='Integrating multi-omics data reveals neuroblastoma subtypes in the tumor microenvironment'; gemma_q2_paper_profile.spatial=NO; gemma_final=FILTERED. | accepted_agree |
| 10.1016/j.lungcan.2015.12.011 | 2016 | Sorafenib treatment for patients with RET fusion-positive non-small cell lung cancer. | AGREE | gemma_final=FILTERED; paper_type=EXP; spatial=NO; panel_scope=UNCLEAR; title=Sorafenib treatment for patients with RET fusion-positive non-small cell lung cancer. | accepted_agree |
| 10.1016/j.matbio.2020.04.004 | 202 | Spatial-omics: Novel approaches to probe cell heterogeneity and extracellular matrix biology | AGREE | gemma_final=FILTERED; paper_type=OTHER; spatial=YES; panel_scope=UNCLEAR; title=Spatial-omics: Novel approaches to probe cell heterogeneity and extracellular matrix biology | accepted_agree |
| 10.1016/j.mcpro.2025.101320 | 2025 | 2.5 Nanoscale molecular cartography of the cellular membranes | AGREE | gemma_final=FILTERED; doc_type=NON_ARTICLE; title=2.5 Nanoscale molecular cartography of the cellular membranes | accepted_agree |

| DOI | Year | Title | Audit decision | Evidence (verbatim quote) | Validation status / scope flag |
|---|---|---|---|---|---|
| 10.1016/j.mcpro.2026.101536 | 2026 | mosna reveals different types of cellular interactions predictive of response to immunotherapies and survival in cancer. | AGREE | gemma_final=FILTERED; paper_type=COMP; spatial=YES; panel_scope=UNCLEAR; title=mosna reveals different types of cellular interactions predictive of response to immunotherapies and survival in cancer. | accepted_agree |
| 10.1016/j.modpat.2025.100866 | 2025 | Histology-Based Virtual RNA Inference Identifies Pathways Associated With Metastasis Risk in Colorectal Cancer. | AGREE | gemma_final=FILTERED; paper_type=COMP; spatial=YES; panel_scope=NOT_TARGETED; title=Histology-Based Virtual RNA Inference Identifies Pathways Associated With Metastasis Risk in Colorectal Cancer. | accepted_agree |
| 10.1016/j.molp.2024.01.009 | 2024 | Callus proliferation-induced hypoxic microenvironment decreases shoot regeneration competence in Arabidopsis | AGREE | paper.title='Callus proliferation-induced hypoxic microenvironment decreases shoot regeneration competence in Arabidopsis'; gemma_q2_paper_profile.spatial=NO; gemma_final=FILTERED. | accepted_agree |
| 10.1016/j.molp.2025.12.026 | 2025 | PlantscRNAdb 4.0: Improved marker identification and annotation under a cell-type uniformity for plants | AGREE | title: PlantscRNAdb 4.0: Improved marker identification and annotation under a cell-type uniformity for plants; q0 doc_type is blank, q2 paper_type=OTHER, q2 panel_scope=UNCLEAR, q2 spatial=NO, and q1 panel-design fields are empty. | accepted_agree |
| 10.1016/j.ncrops.2024.100025 | 2024 | Applications for single-cell and spatial transcriptomics in plant research | AGREE | paper.title='Applications for single-cell and spatial transcriptomics in plant research'; gemma_q0_document_type=REVIEW; gemma_q2_paper_profile.spatial=YES; gemma_final=FILTERED. | accepted_agree |
| 10.1016/j.neunet.2023.02.018 | 2023 | UDRN: Unified Dimensional Reduction Neural Network for feature selection and feature projection | AGREE | title='UDRN: Unified Dimensional Reduction Neural Network for feature selection and feature projection'; citation_screening_prior.labels='COMP'; gemma_final.source='citation_screening_prefilter' | accepted_agree |
| 10.1016/j.neuron.2017.04.023 | 2017 | Integrating Results across Methodologies Is Essential for | AGREE | Title, paper_type=OTHER, spatial=YES, | accepted_agree |

| DOI | Year | Title | Audit decision | Evidence (verbatim quote) | Validation status / scope flag |
|---|---|---|---|---|---|
| | | Producing Robust Neuronal Taxonomies. | | panel_scope=TARGETED, gemma_final=FILTERED. | |
| 10.1016/j.neuron.2017.05.008 | 2017 | seqFISH Accurately Detects Transcripts in Single Cells and Reveals Robust Spatial Organization in the Hippocampus. | NEEDS_MANUAL | PDF: Matters Arising response reuses the 125/249-gene seqFISH data; no new panel selection. | pdf_validated_unclear_update |
| 10.1016/j.neuron.2023.12.014 | 2024 | Mapping of multiple neurotransmitter receptor subtypes and distinct protein complexes to the connectome. | AGREE | Title, paper_type=EXP, spatial=NO, panel_scope=UNCLEAR, gemma_final=FILTERED. | accepted_agree |
| 10.1016/j.neuron.2024.06.004 | 2024 | Cellular and circuit architecture of the lateral septum for reward processing | AGREE | Title, paper_type=EXP, spatial=NO, panel_scope=UNCLEAR, gemma_final=FILTERED. | accepted_agree |
| 10.1016/j.neuron.2025.05.002 | 2025 | An enhancer-AAV toolbox to target and manipulate distinct interneuron subtypes | AGREE | citation_screening_prior.labels=NON-ST; gemma_final.classification=FILTERED; gemma_q1_panel_design blank. | accepted_agree |
| 10.1016/j.nmd.2024.07.510 | 2024 | 306P Evaluating genetic therapies targeting the central nervous system in Duchenne muscular dystrophy using high-resolution spatial transcriptomics | AGREE | Risk flag non_primary_document and gemma_final=FILTERED. | accepted_agree |
| 10.1016/j.ohx.2022.e00343 | 2022 | An open source 16-channel fluidics system for automating sequential fluorescent <i>in situ</i> hybridization (FISH)-based imaging. | AGREE | Title, paper_type=COMP, spatial=YES, panel_scope=UNCLEAR, gemma_final=FILTERED. | accepted_agree |
| 10.1016/j.oooo.2023.12.645 | 2024 | SPATIAL LOCALIZATION OF MACROPHAGES AND GALR2/GALANIN EXPRESSION IN ORAL SQUAMOUS CELL CARCINOMA (OSCC) | AGREE | Risk flag non_primary_document and gemma_final=FILTERED. | accepted_agree |
| 10.1016/j.pathol.2025.10.003 | 2025 | Spatial transcriptomics in bone research: navigating hype and hurdles | AGREE | gemma_q0_document_type.doc_type=REVIEW; citation_screening_prior.labels=NON-ST; gemma_final.classification=FILTERED. | accepted_agree |
| 10.1016/j.patter.2024.101022 | 2024 | CellContrast: Reconstructing spatial relationships in single-cell RNA sequencing data via deep contrastive learning. | AGREE | Title, paper_type=COMP, spatial=YES, panel_scope=UNCLEAR, gemma_final=FILTERED. | accepted_agree |

| DOI | Year | Title | Audit decision | Evidence (verbatim quote) | Validation status / scope flag |
|---|---|---|---|---|---|
| 10.1016/j.placenta.2025.07.037 | 2025 | Single-cell spatial mapping of trophoblast subpopulations in the human placenta using HybISS | AGREE | Risk flag non_primary_document and gemma_final=FILTERED. | accepted_agree |
| 10.1016/j.pld.2017.05.003 | 2017 | High molecular karyotype variation revealed in indigenous <i>Elymus nutans</i> in the Qinghai Plateau. | AGREE | Title, paper_type=EXP, spatial=NO, panel_scope=UNCLEAR, gemma_final=FILTERED. | accepted_agree |
| 10.1016/j.rbmo.2015.04.003 | 2015 | Altered segregation pattern and numerical chromosome abnormalities interrelate in spermatozoa from Robertsonian translocation carriers | AGREE | Title, paper_type=EXP, spatial=NO, panel_scope=UNCLEAR, gemma_final=FILTERED. | accepted_agree |
| 10.1016/j.rmr.2024.01.068 | 2024 | Identification of abnormal airway niches in the fibrotic lung using spatial transcriptomics | AGREE | Risk flag non_primary_document and gemma_final=FILTERED. | accepted_agree |
| 10.1016/j.scib.2023.09.021 | 2023 | Single-cell dynamics of liver development in postnatal pigs | AGREE | title='Single-cell dynamics of liver development in postnatal pigs'; citation_screening_prior.labels='NON-ST'; gemma_final.source='citation_screening_prefilter' | accepted_agree |
| 10.1016/j.scienta.2017.12.064 | 2018 | Nucleolar activity and physical location of ribosomal DNA loci in Vitis vinifera L. by silver staining and sequential FISH | AGREE | q2 says spatial is NO and panel scope is UNCLEAR; q1 only describes a 45S rDNA probe for chromosome mapping. | accepted_agree |
| 10.1016/j.scienta.2020.109633 | 202 | Development of a set of chromosome-specific oligonucleotide markers and karyotype analysis in the Japanese morning glory Ipomoea nil | AGREE | Title, paper_type=EXP, spatial=NO, panel_scope=UNCLEAR, gemma_final=FILTERED. | accepted_agree |
| 10.1016/j.semcancer.2026.01.002 | 2026 | Integrative spatial omics and artificial intelligence: transforming cancer research with omics data and AI | AGREE | Title, paper_type=OTHER, spatial=YES, panel_scope=UNCLEAR, gemma_final=FILTERED. | accepted_agree |
| 10.1016/j.stem.2025.10.001 | 2025 | 3D-generation of high-purity midbrain dopaminergic progenitors and lineage-guided refinement of grafts supports Parkinson’s disease cell therapy | NEEDS_MANUAL | panel_scope=NOT_TARGETED with no target-gene panel source evidence; untargeted/no-panel records are out of scope. Original audit evidence: q1: panel_design=UNCLEAR; q2: panel_scope=NOT_TARGETED; evidence_tool/evidence_manual/evidence_vendor are NOT FOUND. | validated_high_risk_manual_filtered |

| DOI | Year | Title | Audit decision | Evidence (verbatim quote) | Validation status / scope flag |
|---|---|---|---|---|---|
| 10.1016/j.swevo.2025.102108 | 2025 | A multi-objective metaheuristic approach for generating cell type marker panels from single-cell transcriptomics | NEEDS_MANUAL | spatial=NO; packet does not support an in-scope spatial panel-design record. Original audit evidence: We propose a multi-objective metaheuristic approach called scMOCTYM for generating cell type marker panels from single-cell transcriptomic data | validated_high_risk_manual_filtered |
| 10.1016/j.ultrasmedbio.2019.07.444 | 2019 | New guidelines on gastrointestinal ultrasound (GIUS) | AGREE | Title, paper_type=OTHER, spatial=NO, panel_scope=UNCLEAR, gemma_final=FILTERED. | accepted_agree |
| 10.1016/j.urolonc.2024.01.048 | 2024 | EPITHELIAL TO MESENCHYMAL TRANSITION (EMT) AND PD-1/PD-L1 SIGNALING PROVIDE RECIPROCAL FEEDBACK IN RENAL CELL CARCINOMA PROGRESSION WITH IMPLICATIONS FOR IMMUNOTHERAPY RESPONSE | AGREE | Risk flag non_primary_document and gemma_final=FILTERED. | accepted_agree |
| 10.1016/j.urolonc.2024.01.049 | 2024 | PROFILING THE TUMOR IMMUNE MICROENVIRONMENT OF PAPILLARY RENAL CELL CARCINOMA | AGREE | Risk flag non_primary_document and gemma_final=FILTERED. | accepted_agree |
| 10.1016/j.xgen.2023.100384 | 2023 | A unified pipeline for FISH spatial transcriptomics. | NEEDS_MANUAL | Pipeline for MERFISH/seqFISH/targeted ISS; text says "novel seqFISH decoding algorithm" and QC metrics, not panel design. | pdf_validated_unclear_update |
| 10.1016/j.xgen.2025.101005 | 2025 | Galaxy single-cell & spatial omics community update: Navigating new frontiers in 2025 | AGREE | title: Galaxy single-cell & spatial omics community update: Navigating new frontiers in 2025; q0 doc_type=REVIEW; q1 panel-design fields are empty; citation prior labels=REV and Gemma_final=FILTERED. | accepted_agree |
| 10.1016/j.xgen.2025.101007 | 2025 | Single-nucleus transcriptome atlas of orbitofrontal cortex in ALS with a deep learning-based decoding of alternative polyadenylation mechanisms | AGREE | citation_screening_prior.labels=NON-ST; gemma_final.classification=FILTERED; gemma_q1_panel_design blank. | accepted_agree |
| 10.1016/j.xgen.2025.101103 | 2025 | A high-resolution atlas of the brain predicts lineage and birth order underlying neuronal identity | AGREE | gemma_final.source=abstract_prefilter; gemma_final.notes=non-spatial; | accepted_agree |

| DOI | Year | Title | Audit decision | Evidence (verbatim quote) | Validation status / scope flag |
|---|---|---|---|---|---|
| | | | | gemma_q2_paper_profile.panel_scope=UNCLEAR. | |
| 10.1016/j.xgen.2025.101105 | 2026 | Robust integration and annotation of single-cell and spatial omics data using interpretable gene programs. | AGREE | Title, paper_type=COMP, spatial=YES, panel_scope=UNCLEAR, gemma_final=FILTERED. | accepted_agree |
| 10.1016/j.xhgg.2025.100447 | 2025 | Integrating spatial transcriptomics and snRNA-seq data enhances differential gene expression analysis results of AD-related phenotypes | AGREE | Title, paper_type=COMP, spatial=YES, panel_scope=UNCLEAR, gemma_final=FILTERED. | accepted_agree |
| 10.1017/s1551929521000894 | 2021 | MERFISH Enables Mapping of Cellular Diversity with Spatial Context at Subcellular Resolution | NEEDS_MANUAL | Final manual decision: FILTERED because no local PDF was available and the packet did not provide usable final gene-panel source evidence. Original evidence: q1 has no panel-design evidence; q2 only says the paper reports a MERFISH measurement of 417 genes across a human colon cancer sample. | manual_final_filter_after_pdf_review |
| 10.1017/s2633903x2300017x | 2023 | Performant web-based interactive visualization tool for spatially-resolved transcriptomics experiments. | AGREE | Title, paper_type=COMP, spatial=YES, panel_scope=UNCLEAR, gemma_final=FILTERED. | accepted_agree |
| 10.1021/acs.analchem.8b02572 | 2018 | Magnetism-Resolved Separation and Fluorescence Quantification for Near-Simultaneous Detection of Multiple Pathogens | AGREE | Title: Magnetism-Resolved Separation and Fluorescence Quantification for Near-Simultaneous Detection of Multiple Pathogens; gemma_final notes: Pass-A abstract: non-spatial -> FILTERED. | accepted_agree |
| 10.1021/acs.bioconjchem.1c00375 | 2022 | A Photoacoustic Contrast Agent for miR-21 via NIR Fluorescent Hybridization Chain Reaction. | AGREE | Title: A Photoacoustic Contrast Agent for miR-21 via NIR Fluorescent Hybridization Chain Reaction.; gemma_final notes: Pass-A abstract: non-spatial -> FILTERED. | accepted_agree |
| 10.1021/acs.energyfuels.0c02744 | 202 | Molecular Cartography of A1 and A2 Asphaltene Subfractions from Classical Molecular Dynamics Simulations | AGREE | Title: Molecular Cartography of A1 and A2 Asphaltene Subfractions from Classical Molecular Dynamics Simulations; gemma_final notes: Pass-A abstract: COMP -> FILTERED. | accepted_agree |

| DOI | Year | Title | Audit decision | Evidence (verbatim quote) | Validation status / scope flag |
|---|---|---|---|---|---|
| 10.1021/acs.jcim.1c01269 | 2022 | SURFMAP: A Software for Mapping in Two Dimensions Protein Surface Features. | AGREE | Title: SURFMAP: A Software for Mapping in Two Dimensions Protein Surface Features.; gemma_final notes: Pass-A abstract: COMP -> FILTERED. | accepted_agree |
| 10.1021/acs.jnatprod.5c01111 | 2026 | Paraisariamides: Cycloheptapeptide Toxins from Entomopathogenic Fungi (<i>Paraisaria</i> spp.) That Inhibit Total Protein Synthesis. | AGREE | q1 and q2 contain no spatial/panel-design evidence; the paper is about toxin isolation and structure elucidation. | accepted_agree |
| 10.1021/acschemneuro.4c00069 | 2024 | Effect of Different Staining Methods on Brain Cryosections | AGREE | The packet says the library construction protocol consisted of microregion RNA extraction and Smart-seq2 library construction, with panel_design = NOT FOUND. | accepted_agree |
| 10.1021/acsmedchemlett.5c00735 | 2025 | Precision Therapeutics Across Cancer, Immunology, and Pain | AGREE | Title: Precision Therapeutics Across Cancer, Immunology, and Pain; gemma_final notes: Pass-A abstract: OTHER -> FILTERED. | accepted_agree |
| 10.1021/acsnano.3c10216 | 2024 | Enhancing Specific Fluorescence In Situ Hybridization with Quantum Dots for Single-Molecule RNA Imaging in Formalin-Fixed Paraffin-Embedded Tumor Tissues. | NEEDS_MANUAL | Final manual decision: FILTERED because no local PDF was available and the packet did not provide usable final gene-panel source evidence. Original evidence: Initially, we used PaintSHOP software to obtain specific target sites for mRNA; we selected 20 consecutive specific sites for each gene and QD readout probe for binding with the split probes. | manual_final_filter_after_pdf_review |
| 10.1021/acsnano.4c18470 | 2025 | Advancements in Nanomaterials and Molecular Probes for Spatial Omics | AGREE | gemma_q0_document_type.doc_type=REVIEW; citation_screening_prior.labels=NON-ST; gemma_final.classification=FILTERED. | accepted_agree |
| 10.1021/acsomega.5c05379 | 2025 | Enhanced RNA Preservation in Mouse Brain Tissue: A Strategy Combining Cardiac Perfusion with Hypersaline Immersion | AGREE | gemma_final.source=abstract_prefilter; gemma_q2_paper_profile.paper_type=OTHER; gemma_q2_paper_profile.spatial=YES. | accepted_agree |
| 10.1021/jacs.9b11748 | 202 | <i>In Situ</i> Genetically Cascaded Amplification for | AGREE | Title: In Situ Genetically Cascaded Amplification for Imaging RNA | accepted_agree |

| DOI | Year | Title | Audit decision | Evidence (verbatim quote) | Validation status / scope flag |
|---|---|---|---|---|---|
| | | Imaging RNA Subcellular Locations. | | Subcellular Locations.; gemma_final notes: Pass-A abstract: COMP -> FILTERED. | |
| 10.1038/d41586-019-02477-9 | 2019 | Starfish enterprise: finding RNA patterns in single cells | AGREE | Title: Starfish enterprise: finding RNA patterns in single cells; gemma_final notes: Q0: NON_ARTICLE -> FILTERED. | accepted_agree |
| 10.1038/ncomms8294 | 2015 | Proximity-dependent initiation of hybridization chain reaction. | AGREE | Title: Proximity-dependent initiation of hybridization chain reaction.; gemma_final notes: Pass-A abstract: COMP -> FILTERED. | accepted_agree |
| 10.1038/nmeth.3899 | 2016 | Nanoscale imaging of RNA with expansion microscopy. | AGREE | Title: Nanoscale imaging of RNA with expansion microscopy.; gemma_final notes: Pass-A abstract: COMP -> FILTERED. | accepted_agree |
| 10.1038/nmeth.4086 | 2016 | Neuroscience: Hi-res spatial gene expression in the brain | AGREE | Title: Neuroscience: Hi-res spatial gene expression in the brain; gemma_final notes: Q0: NON_ARTICLE -> FILTERED. | accepted_agree |
| 10.1038/nmeth.4087 | 2016 | Stem cells: Man-made mouse eggs | AGREE | Title: Stem cells: Man-made mouse eggs; gemma_final notes: Q0: NON_ARTICLE -> FILTERED. | accepted_agree |
| 10.1038/nmeth.4088 | 2016 | Structural biology: Laser ablation for growing protein crystals | AGREE | Title: Structural biology: Laser ablation for growing protein crystals; gemma_final notes: Q0: NON_ARTICLE -> FILTERED. | accepted_agree |
| 10.1038/nmeth.4089 | 2016 | Molecular biology: Annotating RNAs by the company they keep | AGREE | Title: Molecular biology: Annotating RNAs by the company they keep; gemma_final notes: Q0: NON_ARTICLE -> FILTERED. | accepted_agree |
| 10.1038/nmeth.4108 | 2017 | Rapidly evolving homing CRISPR barcodes. | AGREE | Title: Rapidly evolving homing CRISPR barcodes.; gemma_final notes: Pass-A abstract: COMP -> FILTERED. | accepted_agree |
| 10.1038/nrg.2017.107 | 2018 | Gene expression: SPOTting single RNA molecules transcriptome-wide | AGREE | Title: Gene expression: SPOTting single RNA molecules transcriptome-wide; gemma_final notes: Q0: NON_ARTICLE -> FILTERED. | accepted_agree |
| 10.1038/nrg.2017.99 | 2017 | Functional genomics: Shining a light on genetic screen strategies | AGREE | paper_type/doc_type evidence: EDITORIAL (RESEARCH HIGHLIGHTS); gemma_final notes: Q0: EDITORIAL → FILTERED | accepted_agree |

| DOI | Year | Title | Audit decision | Evidence (verbatim quote) | Validation status / scope flag |
|---|---|---|---|---|---|
| 10.1038/s12276-026-01636-z | 2026 | UBE2M as a bridge spanning neddylation and cell cycle regulation in colorectal adenocarcinoma | AGREE | gemma_final notes say Pass-A abstract: non-spatial -> FILTERED; q2 paper_type=EXP, spatial=NO, panel_scope=UNCLEAR; q1 panel-design fields are empty. | accepted_agree |
| 10.1038/s41377-024-01734-5 | 2025 | Optical sorting: past, present and future | AGREE | gemma_final.source=abstract_prefilter; gemma_final.notes=non-spatial; gemma_q1_panel_design blank. | accepted_agree |
| 10.1038/s41379-018-0008-8 | 2018 | MYB and MYBL1 in adenoid cystic carcinoma: diversity in the mode of genomic rearrangement and transcripts. | AGREE | Title: MYB and MYBL1 in adenoid cystic carcinoma: diversity in the mode of genomic rearrangement and transcripts.; gemma_final notes: Pass-A abstract: non-spatial -> FILTERED. | accepted_agree |
| 10.1038/s41380-025-03014-z | 2025 | A single-cell genomic atlas for the effects of chronic ethanol exposure in the mouse dorsal striatum | AGREE | Title: A single-cell genomic atlas for the effects of chronic ethanol exposure in the mouse dorsal striatum; gemma_final notes: Pass-A abstract: non-spatial -> FILTERED. | accepted_agree |
| 10.1038/s41392-025-02409-2 | 2025 | Tumor microenvironment delineates differential responders to trastuzumab emtansine in HER2-positive metastatic breast cancer patients previously treated with pyrotinib: an exploratory biomarker analysis of a phase II study (NJMU-BC02) | AGREE | citation_screening_prior.labels=NON-ST; gemma_final.classification=FILTERED; gemma_q1_panel_design blank. | accepted_agree |
| 10.1038/s41393-025-01077-x | 2025 | BDNF-GABA signaling in astrocytes: enhancing neural repair after SCI through MSC therapies | AGREE | citation_screening_prior.labels=COMP; gemma_q2_paper_profile.spatial=NO; gemma_final.classification=FILTERED. | accepted_agree |
| 10.1038/s41393-025-01103-y | 2025 | Microglial landscape and signaling in spinal cord injury | AGREE | paper.title='Microglial landscape and signaling in spinal cord injury'; citation_screening_prior.labels='COMP'; gemma_final.classification='FILTERED'; gemma_q1_panel_design blank; gemma_q2_paper_profile.paper_type='COMP'; gemma_q2_paper_profile.spatial=''. | accepted_agree |

| DOI | Year | Title | Audit decision | Evidence (verbatim quote) | Validation status / scope flag |
|---|---|---|---|---|---|
| 10.1038/s41398-023-02665-2 | 2023 | Does decreased autophagy and dysregulation of LC3A in astrocytes play a role in major depressive disorder? | AGREE | title='Does decreased autophagy and dysregulation of LC3A in astrocytes play a role in major depressive disorder?'; citation_screening_prior.labels='NON-ST'; gemma_final.source='citation_screening_prefilter' | accepted_agree |
| 10.1038/s41398-025-03243-4 | 2025 | Systematic dissection of pleiotropic loci and critical regulons in excitatory neurons and microglia relevant to neuropsychiatric and ocular diseases | AGREE | paper.title='Systematic dissection of pleiotropic loci and critical regulons in excitatory neurons and microglia relevant to neuropsychiatric and ocular diseases'; citation_screening_prior.labels='COMP'; gemma_final.classification='FILTERED'; gemma_q1_panel_design blank; gemma_q2_paper_profile.paper_type='COMP'; gemma_q2_paper_profile.spatial=''. | accepted_agree |
| 10.1038/s41413-025-00429-w | 2025 | Advances in spatial transcriptomics and its application in the musculoskeletal system | AGREE | paper.title='Advances in spatial transcriptomics and its application in the musculoskeletal system'; citation_screening_prior.labels='REV'; gemma_q0_document_type.doc_type='REVIEW'; gemma_final.classification='FILTERED'; gemma_q1_panel_design blank. | accepted_agree |
| 10.1038/s41417-023-00651-w | 2023 | A novel subclonal rearrangement of the STRN3::PDGFRB gene in de novo acute myeloid leukemia with NPM1 mutation and its leukemogenic effects | AGREE | Title: A novel subclonal rearrangement of the STRN3::PDGFRB gene in de novo acute myeloid leukemia with NPM1 mutation and its leukemogenic effects; gemma_final notes: Pass-A abstract: non-spatial -> FILTERED. | accepted_agree |
| 10.1038/s41421-024-00656-1 | 2024 | Single-cell multi-omics analysis of lineage development and spatial organization in the human fetal cerebellum | AGREE | Q2: not a targeted panel-design paper -> FILTERED | accepted_agree |

| DOI | Year | Title | Audit decision | Evidence (verbatim quote) | Validation status / scope flag |
|---|---|---|---|---|---|
| 10.1038/s41422-020-0353-2 | 202 | Reconstruction of cell spatial organization from single-cell RNA sequencing data based on ligand-receptor mediated self-assembly. | AGREE | Title: Reconstruction of cell spatial organization from single-cell RNA sequencing data based on ligand-receptor mediated self-assembly.; gemma_final notes: Pass-A abstract: COMP -> FILTERED. | accepted_agree |
| 10.1038/s41422-023-00901-4 | 2024 | It's me, hi, I solved the problem, it's TF-seqFISH | AGREE | paper_type/doc_type evidence: EDITORIAL (RESEARCH HIGHLIGHT); gemma_final notes: Q0: EDITORIAL → FILTERED | accepted_agree |
| 10.1038/s41431-020-00740-6 | 202 | Abstracts from the 53rd European Society of Human Genetics (ESHG) Conference: Oral Presentations | AGREE | Title: Abstracts from the 53rd European Society of Human Genetics (ESHG) Conference: Oral Presentations; gemma_final notes: Pass-A abstract: OTHER -> FILTERED. | accepted_agree |
| 10.1038/s41467-018-05944-5 | 2018 | High-resolution in situ transcriptomics of Pseudomonas aeruginosa unveils genotype independent patho-phenotypes in cystic fibrosis lungs. | NEEDS_MANUAL | spatial=NO; packet does not support an in-scope spatial panel-design record. Original audit evidence: No panel-design evidence was provided in the packet. | validated_high_risk_manual_filtered |
| 10.1038/s41467-021-21453-4 | 2021 | Optimal marker gene selection for cell type discrimination in single cell analyses. | AGREE | paper_type=COMP; spatial=UNCLEAR; gemma_final=FILTERED (Pass-A abstract: COMP -> FILTERED); q1 panel-design fields are blank. | accepted_agree |
| 10.1038/s41467-021-22645-8 | 2021 | Spatiotemporal patterning of photoresponsive DNA-based hydrogels to tune local cell responses. | AGREE | paper_type=EXP; spatial=NO; gemma_final=FILTERED (Pass-A abstract: non-spatial -> FILTERED); q1 panel-design fields are blank. | accepted_agree |
| 10.1038/s41467-021-23807-4 | 2021 | Cell segmentation-free inference of cell types from in situ transcriptomics data. | NEEDS_MANUAL | SSAM analyzes osmFISH, MERFISH, and smFISH images; text is about segmentation-free inference, not panel design. | pdf_validated_unclear_update |
| 10.1038/s41467-021-24405-0 | 2021 | Author Correction: Cell segmentation-free inference of cell types from in situ transcriptomics data | AGREE | paper_type/doc_type evidence: CORRECTION (Author Correction: Cell segmentation-free inference of cell types from in situ transcriptomics data); gemma_final notes: Q0: CORRECTION → FILTERED | accepted_agree |
| 10.1038/s41467-021-26044-x | 2021 | ClusterMap for multi-scale clustering analysis of spatial gene | AGREE | paper_type=COMP; spatial=YES; gemma_final=FILTERED (Pass-A | accepted_agree |

| DOI | Year | Title | Audit decision | Evidence (verbatim quote) | Validation status / scope flag |
|---|---|---|---|---|---|
| | | expression | | abstract: COMP -> FILTERED); q1 panel-design fields are blank. | |
| 10.1038/s41467-022-32111-8 | 2022 | Knowledge-graph-based cell-cell communication inference for spatially resolved transcriptomic data with SpaTalk. | AGREE | paper_type=COMP; spatial=YES; gemma_final=FILTERED (Pass-A abstract: COMP -> FILTERED); q1 panel-design fields are blank. | accepted_agree |
| 10.1038/s41467-023-37392-1 | 2023 | Predictive and robust gene selection for spatial transcriptomics | DISAGREE | This is the computational methods paper introducing PERSIST itself. Although it designs gene targets for spatial transcriptomics inside the method study, that is not a paper-level experimental adoption event and should remain filtered from the panel-design adoption set. | validated_high_risk_disagreement |
| 10.1038/s41467-023-38121-4 | 2023 | Reconstruction of the cell pseudo-space from single-cell RNA sequencing data with scSpace. | AGREE | paper_type=COMP; spatial=YES; gemma_final=FILTERED (Pass-A abstract: COMP -> FILTERED); q1 panel-design fields are blank. | accepted_agree |
| 10.1038/s41467-023-39895-3 | 2023 | Leveraging spatial transcriptomics data to recover cell locations in single-cell RNA-seq with CeLEry. | AGREE | paper_type=COMP; spatial=YES; gemma_final=FILTERED (Pass-A abstract: COMP -> FILTERED); q1 panel-design fields are blank. | accepted_agree |
| 10.1038/s41467-023-41825-2 | 2023 | Author Correction: Spatial organization of the mouse retina at single cell resolution by MERFISH | AGREE | paper_type/doc_type evidence: CORRECTION (Author Correction: Spatial organization of the mouse retina at single cell resolution by MERFISH); gemma_final notes: Q0: CORRECTION → FILTERED | accepted_agree |
| 10.1038/s41467-023-42658-9 | 2023 | Dbh+ catecholaminergic cardiomyocytes contribute to the structure and function of the cardiac conduction system in murine heart | DISAGREE | Stereo-seq is a whole-transcript spatial method and the paper profile marks panel_scope as NOT_TARGETED, so SMaSH marker identification here is not a targeted panel-design event. | accepted_disagree |

| DOI | Year | Title | Audit decision | Evidence (verbatim quote) | Validation status / scope flag |
|---|---|---|---|---|---|
| 10.1038/s41467-023-43462-1 | 2023 | Low-dose radiotherapy combined with dual PD-L1 and VEGFA blockade elicits antitumor response in hepatocellular carcinoma mediated by activated intratumoral CD8+ exhausted-like T cells | AGREE | title='Low-dose radiotherapy combined with dual PD-L1 and VEGFA blockade elicits antitumor response in hepatocellular carcinoma mediated by activated intratumoral CD8+ exhausted-like T cells'; citation_screening_prior.labels='NON-ST'; gemma_final.source='citation_screening_prefilter' | accepted_agree |
| 10.1038/s41467-023-44570-8 | 2024 | MarsGT: Multi-omics analysis for rare population inference using single-cell graph transformer | AGREE | paper.title='MarsGT: Multi-omics analysis for rare population inference using single-cell graph transformer'; gemma_final=FILTERED. | accepted_agree |
| 10.1038/s41467-024-46773-z | 2024 | DELVE: feature selection for preserving biological trajectories in single-cell data | AGREE | paper.title='DELVE: feature selection for preserving biological trajectories in single-cell data'; gemma_final=FILTERED. | accepted_agree |
| 10.1038/s41467-024-47883-4 | 2024 | Cross-modality mapping using image varifolds to align tissue-scale atlases to molecular-scale measures with application to 2D brain sections. | NEEDS_MANUAL | Cross-modality atlas mapping to MERFISH/BARseq; no evidence of gene-panel selection or curation. | pdf_validated_unclear_update |
| 10.1038/s41467-024-48185-5 | 2024 | A non-methanogenic archaeon within the order Methanocellales | AGREE | paper_type=EXP; spatial=NO; gemma_final=FILTERED (Pass-A abstract: non-spatial -> FILTERED); q1 panel-design fields are blank. | accepted_agree |
| 10.1038/s41467-024-48981-z | 2024 | Sopa: a technology-invariant pipeline for analyses of image-based spatial omics. | AGREE | paper_type=COMP; spatial=YES; gemma_final=FILTERED (Pass-A abstract: COMP -> FILTERED); q1 panel-design fields are blank. | accepted_agree |
| 10.1038/s41467-024-49457-w | 2024 | Intracellular spatial transcriptomic analysis toolkit (InSTAnT). | AGREE | paper_type=COMP; spatial=YES; gemma_final=FILTERED (Pass-A abstract: COMP -> FILTERED); q1 panel-design fields are blank. | accepted_agree |
| 10.1038/s41467-024-50837-5 | 2024 | Tissue characterization at an enhanced resolution across spatial omics platforms with deep generative model. | AGREE | paper_type=COMP; spatial=YES; gemma_final=FILTERED (Pass-A abstract: COMP -> FILTERED); q1 panel-design fields are blank. | accepted_agree |

| DOI | Year | Title | Audit decision | Evidence (verbatim quote) | Validation status / scope flag |
|---|---|---|---|---|---|
| 10.1038/s41467-025-58089-7 | 2025 | SIMVI disentangles intrinsic and spatial-induced cellular states in spatial omics data. | AGREE | paper_type=COMP; spatial=YES; gemma_final=FILTERED (Pass-A abstract: COMP -> FILTERED); q1 panel-design fields are blank. | accepted_agree |
| 10.1038/s41467-025-58511-0 | 2025 | Spatially resolved atlas of breast cancer uncovers intercellular machinery of venular niche governing lymphocyte extravasation. | AGREE | paper_type=EXP; spatial=YES; panel_scope=NOT_TARGETED; gemma_final=FILTERED (Pass-A abstract: not targeted -> FILTERED). | accepted_agree |
| 10.1038/s41467-025-58748-9 | 2025 | Hypoxia-induced Wnt5a-secreting fibroblasts promote colon cancer progression | AGREE | paper.title='Hypoxia-induced Wnt5a-secreting fibroblasts promote colon cancer progression'; gemma_final.classification='FILTERED'; gemma_q2_paper_profile.paper_type='EXP'; gemma_q2_paper_profile.spatial='NO'; gemma_q2_paper_profile.panel_scope='UNCLEAR'; gemma_q1_panel_design blank. | accepted_agree |
| 10.1038/s41467-025-58874-4 | 2025 | Deconvolution of cell types and states in spatial multiomics utilizing TACIT | AGREE | paper.title='Deconvolution of cell types and states in spatial multiomics utilizing TACIT'; citation_screening_prior.labels='COMP'; citation_screening_prior.tools='NS-Forest v2'; gemma_final.classification='FILTERED'; gemma_q1_panel_design blank; gemma_q2_paper_profile.paper_type='COMP'. | accepted_agree |
| 10.1038/s41467-025-60482-1 | 2025 | Divergent combinations of enhancers encode spatial gene expression. | AGREE | paper_type=COMP; spatial=YES; gemma_final=FILTERED (Pass-A abstract: COMP -> FILTERED); q1 panel-design fields are blank. | accepted_agree |
| 10.1038/s41467-025-61476-9 | 2025 | Prioritizing perturbation-responsive gene patterns using interpretable deep learning | AGREE | paper.title='Prioritizing perturbation-responsive gene patterns using interpretable deep learning'; citation_screening_prior.labels='COMP-BENCH'; citation_screening_prior.tools='PERSIST'; gemma_final.classification='FILTE | accepted_agree |

| DOI | Year | Title | Audit decision | Evidence (verbatim quote) | Validation status / scope flag |
|---|---|---|---|---|---|
| | | | | RED'; gemma_q1_panel_design blank. | |
| 10.1038/s41467-025-63050-9 | 2025 | FUSION: a web-based application for in-depth exploration of multi-omics data with brightfield histology. | AGREE | paper_type=COMP; spatial=YES; gemma_final=FILTERED (Pass-A abstract: COMP -> FILTERED); q1 panel-design fields are blank. | accepted_agree |
| 10.1038/s41467-025-64259-4 | 2025 | Data-driven fine-grained region discovery in the mouse brain with transformers. | AGREE | paper_type=COMP; spatial=YES; gemma_final=FILTERED (Pass-A abstract: COMP -> FILTERED); q1 panel-design fields are blank. | accepted_agree |
| 10.1038/s41467-025-66584-0 | 2025 | The molecular cartography of malignant and benign sebaceous tumours. | AGREE | paper_type=EXP; spatial=NO; gemma_final=FILTERED (Pass-A abstract: non-spatial -> FILTERED); q1 panel-design fields are blank. | accepted_agree |
| 10.1038/s41467-025-67890-3 | 2025 | CellScope: high-performance cell atlas workflow with tree-structured representation | AGREE | paper.title='CellScope: high-performance cell atlas workflow with tree-structured representation'; citation_screening_prior.labels='COMP'; citation_screening_prior.tools='SCMER'; gemma_final.classification='FILTERED'; gemma_q1_panel_design blank; gemma_q2_paper_profile.paper_type='COMP'. | accepted_agree |
| 10.1038/s41467-026-68500-6 | 2026 | Robust characterization and interpretation of rare pathogenic cell populations from spatial omics using GARDEN | AGREE | citation_screening_prior labels=COMP and all_tools=SCMER; gemma_final source=citation_screening_prefilter -> FILTERED; q1 panel-design fields are empty. | accepted_agree |
| 10.1038/s41467-026-70090-2 | 2026 | Reconstructing single-cell resolution from spatial transcriptomics with CellRefiner. | AGREE | paper_type=COMP; spatial=YES; gemma_final=FILTERED (Pass-A abstract: COMP -> FILTERED); q1 panel-design fields are blank. | accepted_agree |
| 10.1038/s41467-026-70528-7 | 2026 | FineST: contrastive learning integrates histology and spatial transcriptomics for nuclei-resolved ligand-receptor analysis. | AGREE | paper_type=COMP; spatial=YES; gemma_final=FILTERED (Pass-A abstract: COMP -> FILTERED); q1 panel-design fields are blank. | accepted_agree |
| 10.1038/s41467-026-70645-3 | 2026 | Ultra-precision deconvolution of spatial transcriptomics decodes | AGREE | citation_screening_prior labels=COMP and all_tools=COSG; gemma_final | accepted_agree |

| DOI | Year | Title | Audit decision | Evidence (verbatim quote) | Validation status / scope flag |
|---|---|---|---|---|---|
| | | immune heterogeneity and fate-defining programs in tissues | | source=citation_screening_prefilter -> FILTERED; q1 panel-design fields are empty. | |
| 10.1038/s41522-023-00423-2 | 2023 | Quantitative whole-tissue 3D imaging reveals bacteria in close association with mouse jejunum mucosa. | DISAGREE | q1: universal degenerate HCR v3.0 probe set plus two new probes for E. coli and Bacteroides/Parabacteroides; q2: targeted bacterial 16S rRNA visualization, not gene-panel selection. | validated_high_risk_disagreement |
| 10.1038/s41540-024-00339-3 | 2024 | MarkerMap: nonlinear marker selection for single-cell studies | AGREE | Title is 'MarkerMap: nonlinear marker selection for single-cell studies'; citation prior marks it computational-only and q1 panel-design fields are empty. | accepted_agree |
| 10.1038/s41540-024-00441-6 | 2024 | Computational identification of surface markers for isolating distinct subpopulations from heterogeneous cancer cell populations | AGREE | paper.title='Computational identification of surface markers for isolating distinct subpopulations from heterogeneous cancer cell populations'; gemma_q2_paper_profile.spatial=NO; gemma_final=FILTERED. | accepted_agree |
| 10.1038/s41551-025-01512-7 | 2025 | Spatial profiling of gene editing by in situ sequencing in mice and macaques. | DISAGREE | q1: ABEmax/ABE8e constructs introducing synonymous mutations in Actb, Gapdh, and Tubb genes; q2: targeted ISS used to map editing events. | validated_high_risk_disagreement |
| 10.1038/s41556-025-01755-1 | 2025 | CellNavi predicts genes directing cellular transitions by learning a gene graph-enhanced cell state manifold | AGREE | paper.title='CellNavi predicts genes directing cellular transitions by learning a gene graph-enhanced cell state manifold'; citation_screening_prior.labels='COMP'; citation_screening_prior.tools='SCMER'; gemma_final.classification='FILTERED'; gemma_q1_panel_design blank. | accepted_agree |
| 10.1038/s41556-025-01781-z | 2025 | CoCo-ST detects global and local biological structures in spatial transcriptomics datasets. | AGREE | paper_type=COMP; spatial=YES; gemma_final=FILTERED (Pass-A abstract: COMP -> FILTERED); q1 panel-design fields are blank. | accepted_agree |
| 10.1038/s41568-025-00886-9 | 2025 | A guide to transcriptomic deconvolution in cancer | AGREE | title: A guide to transcriptomic deconvolution in cancer; q0 doc_type=REVIEW; q1 panel-design fields are empty; citation | accepted_agree |

| DOI | Year | Title | Audit decision | Evidence (verbatim quote) | Validation status / scope flag |
|---|---|---|---|---|---|
| | | | | prior labels=REV and Gemma_final=FILTERED. | |
| 10.1038/s41575-023-00784-1 | 2023 | A Roadmap for the Human Gut Cell Atlas | AGREE | title='A Roadmap for the Human Gut Cell Atlas'; citation_screening_prior.labels='REV'; gemma_final.source='citation_screening_prefilter' | accepted_agree |
| 10.1038/s41576-021-00444-7 | 2022 | Temporal modelling using single-cell transcriptomics | AGREE | title='Temporal modelling using single-cell transcriptomics'; citation_screening_prior.labels='REV'; gemma_final.source='citation_screening_prefilter' | accepted_agree |
| 10.1038/s41576-022-00552-y | 2022 | Spatial epigenomics in single cells | AGREE | Q0 doc_type=NON_ARTICLE; gemma_final=FILTERED (Q0: NON_ARTICLE -> FILTERED); risk_flags includes non_primary_document. | accepted_agree |
| 10.1038/s41576-023-00586-w | 2023 | Best practices for single-cell analysis across modalities | AGREE | title='Best practices for single-cell analysis across modalities'; citation_screening_prior.labels='REV'; gemma_final.source='citation_screening_prefilter' | accepted_agree |
| 10.1038/s41576-025-00868-5 | 2025 | Spatial multi-omics of nuclear architecture with two-layer seqFISH. | AGREE | paper_type/doc_type evidence: EDITORIAL (Research highlights); gemma_final notes: Q0: EDITORIAL → FILTERED | accepted_agree |
| 10.1038/s41580-024-00768-2 | 2024 | Profiling cell identity and tissue architecture with single-cell and spatial transcriptomics | AGREE | title: Profiling cell identity and tissue architecture with single-cell and spatial transcriptomics; q0 doc_type=REVIEW; q1 panel-design fields are empty; citation prior labels=REV and Gemma_final=FILTERED. | accepted_agree |
| 10.1038/s41586-021-03634-9 | 2021 | Exploring tissue architecture using spatial transcriptomics | AGREE | Title: 'Exploring tissue architecture using spatial transcriptomics'; packet fields show doc_type=REVIEW and Gemma's FILTERED prefilter. | accepted_agree |
| 10.1038/s41586-021-03969-3 | 2021 | Isoform cell-type specificity in the mouse primary motor cortex. | DISAGREE | q1 panel_design: UNCLEAR / not found; q2 panel_scope=NOT_TARGETED | validated_high_risk_disagreement |

| DOI | Year | Title | Audit decision | Evidence (verbatim quote) | Validation status / scope flag |
|---|---|---|---|---|---|
| | | | | with Visium/Spatial Gene Expression on spinal cord. | |
| 10.1038/s41586-022-05385-7 | 2022 | The neurons that restore walking after paralysis. | AGREE | q2 panel_scope=NOT_TARGETED and q1 has no recoverable panel-design source beyond cross-referencing validated probes. | accepted_agree |
| 10.1038/s41586-024-07336-w | 2024 | Paternal microbiome perturbations impact offspring fitness | AGREE | gemma_final=FILTERED; paper_type=EXP; spatial=NO; panel_scope=UNCLEAR; title=Paternal microbiome perturbations impact offspring fitness; notes=Pass-A abstract: non-spatial -> FILTERED. | accepted_agree |
| 10.1038/s41586-024-08350-8 | 2025 | Brain-wide cell-type-specific transcriptomic signatures of healthy ageing in mice | AGREE | gemma_final=FILTERED; paper_type=EXP; spatial=NO; panel_scope=UNCLEAR; title=Brain-wide cell-type-specific transcriptomic signatures of healthy ageing in mice; notes=Pass-A abstract: non-spatial -> FILTERED. | accepted_agree |
| 10.1038/s41586-025-09420-1 | 2025 | Amygdala–liver signalling orchestrates glycaemic responses to stress | AGREE | gemma_final=FILTERED; paper_type=EXP; spatial=NO; panel_scope=UNCLEAR; title=Amygdala-liver signalling orchestrates glycaemic responses to stress; notes=Pass-A abstract: non-spatial -> FILTERED. | accepted_agree |
| 10.1038/s41586-025-09644-1 | 2025 | Continuous cell-type diversification in mouse visual cortex development | AGREE | gemma_final=FILTERED; paper_type=EXP; spatial=NO; panel_scope=UNCLEAR; title=Continuous cell-type diversification in mouse visual cortex development; notes=Pass-A abstract: non-spatial -> FILTERED. | accepted_agree |
| 10.1038/s41586-025-09916-w | 2026 | MAPK-driven epithelial cell plasticity drives colorectal cancer therapeutic resistance | AGREE | gemma_final=FILTERED; paper_type=EXP; spatial=NO; panel_scope=UNCLEAR; title=MAPK-driven epithelial cell plasticity drives colorectal cancer therapeutic resistance; notes=Pass-A abstract: non-spatial -> FILTERED. | accepted_agree |
| 10.1038/s41587-019-0207-y | 2019 | Immuno-SABER enables highly multiplexed and amplified protein | AGREE | gemma_final=FILTERED; paper_type=COMP; spatial=YES; | accepted_agree |

| DOI | Year | Title | Audit decision | Evidence (verbatim quote) | Validation status / scope flag |
|---|---|---|---|---|---|
| | | imaging in tissues. | | panel_scope=UNCLEAR; title=Immuno-SABER enables highly multiplexed and amplified protein imaging in tissues.; notes=Pass-A abstract: COMP -> FILTERED. | |
| 10.1038/s41587-019-0330-9 | 2019 | Exploring a world of a thousand dimensions | AGREE | Title: 'Exploring a world of a thousand dimensions'; packet fields show labels=REV and Gemma's FILTERED prefilter. | accepted_agree |
| 10.1038/s41587-021-01044-w | 2022 | Cell segmentation in imaging-based spatial transcriptomics. | AGREE | gemma_final=FILTERED; paper_type=COMP; spatial=YES; panel_scope=UNCLEAR; title=Cell segmentation in imaging-based spatial transcriptomics.; notes=Pass-A abstract: COMP -> FILTERED. | accepted_agree |
| 10.1038/s41587-022-01251-z | 2022 | Integrative spatial analysis of cell morphologies and transcriptional states with MUSE. | AGREE | gemma_final=FILTERED; paper_type=COMP; spatial=YES; panel_scope=UNCLEAR; title=Integrative spatial analysis of cell morphologies and transcriptional states with MUSE.; notes=Pass-A abstract: COMP -> FILTERED. | accepted_agree |
| 10.1038/s41587-023-01681-3 | 2023 | Spatial tropism profiling of AAV vectors by ultrasensitive sequential FISH in tissue | AGREE | gemma_final=FILTERED; doc_type=NON_ARTICLE; doc_evidence=Research briefing; risk_flags=non_primary_document. | accepted_agree |
| 10.1038/s41587-024-02391-0 | 2025 | Multiplexed, image-based pooled screens in primary cells and tissues with PerturbView. | AGREE | gemma_final=FILTERED; paper_type=COMP; spatial=YES; panel_scope=UNCLEAR; title=Multiplexed, image-based pooled screens in primary cells and tissues with PerturbView.; notes=Pass-A abstract: COMP -> FILTERED. | accepted_agree |
| 10.1038/s41587-024-02516-5 | 2025 | NIS-Seq enables cell-type-agnostic optical perturbation screening. | AGREE | gemma_final=FILTERED; paper_type=EXP; spatial=NO; panel_scope=UNCLEAR; title=NIS-Seq enables cell-type-agnostic optical perturbation screening.; notes=Pass-A abstract: non-spatial -> FILTERED. | accepted_agree |
| 10.1038/s41587-025-02811-9 | 2025 | Standardized metrics for assessment and reproducibility of | AGREE | gemma_final=FILTERED; paper_type=OTHER; spatial=YES; | accepted_agree |

| DOI | Year | Title | Audit decision | Evidence (verbatim quote) | Validation status / scope flag |
|---|---|---|---|---|---|
| | | imaging-based spatial transcriptomics datasets. | | panel_scope=TARGETED; title=Standardized metrics for assessment and reproducibility of imaging-based spatial transcriptomics datasets.; notes=Pass-A abstract: OTHER -> FILTERED. | |
| 10.1038/s41588-022-01256-z | 2023 | SPICEMIX enables integrative single-cell spatial modeling of cell identity. | AGREE | gemma_final=FILTERED; paper_type=COMP; spatial=YES; panel_scope=UNCLEAR; title=SPICEMIX enables integrative single-cell spatial modeling of cell identity.; notes=Pass-A abstract: COMP -> FILTERED. | accepted_agree |
| 10.1038/s41588-024-01662-5 | 2024 | Multi-omic profiling of clear cell renal cell carcinoma identifies metabolic reprogramming associated with disease progression | AGREE | gemma_q1_panel_design = NOT FOUND; gemma_q2_panel_scope = NOT_TARGETED. | accepted_agree |
| 10.1038/s41588-024-01802-x | 2024 | Single-cell multi-omic and spatial profiling of human kidneys implicates the fibrotic microenvironment in kidney disease progression. | AGREE | gemma_final=FILTERED; paper_type=EXP; spatial=YES; panel_scope=NOT_TARGETED; title=Single-cell multi-omic and spatial profiling of human kidneys implicates the fibrotic microenvironment in kidney disease progression.; notes=Pass-A abstract: not targeted -> FILTERED. | accepted_agree |
| 10.1038/s41588-025-02139-9 | 2025 | Spatially resolved transcriptomic analysis of the adult human prostate | AGREE | gemma_q1_panel_design = NOT FOUND; gemma_q2_panel_scope = NOT_TARGETED; gemma_final = FILTERED. | accepted_agree |
| 10.1038/s41588-025-02454-1 | 2026 | Single-cell atlas of the transcriptome and chromatin accessibility in the human retina | AGREE | gemma_final notes say Pass-A abstract: non-spatial -> FILTERED; q2 paper_type=EXP, spatial=NO, panel_scope=UNCLEAR; q1 panel-design fields are empty. | accepted_agree |
| 10.1038/s41588-025-02475-w | 2026 | A microenvironment-determined risk continuum refines subtyping in meningioma and reveals determinants of machine learning-based tumor classification | AGREE | gemma_final notes say Pass-A abstract: non-spatial -> FILTERED; q2 paper_type=EXP, spatial=NO, panel_scope=UNCLEAR; q1 panel-design fields are empty. | accepted_agree |
| 10.1038/s41588-026-02555-5 | 2026 | A refined blueprint for human skin | AGREE | paper_type/doc_type evidence: EDITORIAL (News & views); | accepted_agree |

| DOI | Year | Title | Audit decision | Evidence (verbatim quote) | Validation status / scope flag |
|---|---|---|---|---|---|
| | | | | gemma_final notes: Q0: EDITORIAL → FILTERED | |
| 10.1038/s41592-020-01019-w | 202 | Author Correction: Highly specific multiplexed RNA imaging in tissues with split-FISH | AGREE | paper_type/doc_type evidence: CORRECTION (Author Correction: Highly specific multiplexed RNA imaging in tissues with split-FISH); gemma_final notes: Q0: CORRECTION → FILTERED | accepted_agree |
| 10.1038/s41592-021-01264-7 | 2021 | Deep learning and alignment of spatially resolved single-cell transcriptomes with Tangram. | AGREE | gemma_final=FILTERED; paper_type=COMP; spatial=YES; panel_scope=UNCLEAR; title=Deep learning and alignment of spatially resolved single-cell transcriptomes with Tangram.; notes=Pass-A abstract: COMP -> FILTERED. | accepted_agree |
| 10.1038/s41592-022-01575-3 | 2022 | Cell type-specific inference of differential expression in spatial transcriptomics. | AGREE | gemma_final=FILTERED; paper_type=COMP; spatial=YES; panel_scope=UNCLEAR; title=Cell type-specific inference of differential expression in spatial transcriptomics.; notes=Pass-A abstract: COMP -> FILTERED. | accepted_agree |
| 10.1038/s41592-022-01601-4 | 2022 | ISSAAC-seq enables sensitive and flexible multimodal profiling of chromatin accessibility and gene expression in single cells. | AGREE | gemma_final=FILTERED; paper_type=COMP; spatial=YES; panel_scope=UNCLEAR; title=ISSAAC-seq enables sensitive and flexible multimodal profiling of chromatin accessibility and gene expression in single cells.; notes=Pass-A abstract: COMP -> FILTERED. | accepted_agree |
| 10.1038/s41592-022-01604-1 | 2022 | Light-Seq: light-directed in situ barcoding of biomolecules in fixed cells and tissues for spatially indexed sequencing. | AGREE | gemma_final=FILTERED; paper_type=COMP; spatial=YES; panel_scope=NOT_TARGETED; title=Light-Seq: light-directed in situ barcoding of biomolecules in fixed cells and tissues for spatially indexed sequencing.; notes=Pass-A abstract: COMP -> FILTERED. | accepted_agree |
| 10.1038/s41592-022-01718-6 | 2022 | Spatially resolved epigenomics | AGREE | gemma_final=FILTERED; doc_type=NON_ARTICLE; doc_evidence=Research highlights; risk_flags=non_primary_document. | accepted_agree |

| DOI | Year | Title | Audit decision | Evidence (verbatim quote) | Validation status / scope flag |
|---|---|---|---|---|---|
| 10.1038/s41592-023-01922-y | 2023 | Assembly-free phylogenetic trees | AGREE | gemma_final=FILTERED; doc_type=NON_ARTICLE; doc_evidence=Research highlights; risk_flags=non_primary_document. | accepted_agree |
| 10.1038/s41592-023-01976-y | 2023 | DNA-barcoded signal amplification for imaging mass cytometry enables sensitive and highly multiplexed tissue imaging. | AGREE | paper title; gemma_final notes/source; empty q1 panel-design fields; q2 profile says EXP/non-spatial and no panel evidence. | accepted_agree |
| 10.1038/s41592-023-01977-x | 2023 | Multielement Z-tag imaging by X-ray fluorescence microscopy for next-generation multiplex imaging. | AGREE | paper title; gemma_final notes/source; empty q1 panel-design fields; q2 profile says COMP/non-spatial and no panel evidence. | accepted_agree |
| 10.1038/s41592-024-02212-x | 2025 | SpatialData: an open and universal data framework for spatial omics. | AGREE | paper title; gemma_final notes/source; empty q1 panel-design fields; q2 profile says COMP/spatial and no panel evidence. | accepted_agree |
| 10.1038/s41592-024-02284-9 | 2024 | Accurate and efficient integrative reference-informed spatial domain detection for spatial transcriptomics. | AGREE | paper title; gemma_final notes/source; empty q1 panel-design fields; q2 profile says COMP/spatial and no panel evidence. | accepted_agree |
| 10.1038/s41592-024-02294-7 | 2024 | Deconwolf enables high-performance deconvolution of widefield fluorescence microscopy images. | AGREE | paper title; gemma_final notes/source; empty q1 panel-design fields; q2 profile says COMP/spatial and no panel evidence. | accepted_agree |
| 10.1038/s41592-024-02408-1 | 2024 | Mapping cellular interactions from spatially resolved transcriptomics data. | AGREE | paper title; gemma_final notes/source; empty q1 panel-design fields; q2 profile says COMP/spatial and no panel evidence. | accepted_agree |
| 10.1038/s41592-024-02496-z | 2024 | Probe set selection for targeted spatial transcriptomics. | DISAGREE | This is the computational methods paper introducing Spapros itself. Although it includes a demonstration SCRINSHOT experiment with a Spapros-generated panel, that is not a paper-level experimental adoption event and should remain filtered from the panel-design adoption set. | validated_high_risk_disagreement |

| DOI | Year | Title | Audit decision | Evidence (verbatim quote) | Validation status / scope flag |
|---|---|---|---|---|---|
| 10.1038/s41592-024-02512-2 | 2025 | Efficient and highly amplified imaging of nucleic acid targets in cellular and histopathological samples with pSABER. | AGREE | paper title; gemma_final notes/source; empty q1 panel-design fields; q2 profile says COMP/spatial and no panel evidence. | accepted_agree |
| 10.1038/s41592-024-02532-y | 2024 | Considerations for building and using integrated single-cell atlases | AGREE | paper_type/doc_type evidence: REVIEW; citation-screening prior labels=['REV']; gemma_final notes: citation-screening prior pre-filter | accepted_agree |
| 10.1038/s41592-024-02563-5 | 2025 | Human BioMolecular Atlas Program (HuBMAP): 3D Human Reference Atlas construction and usage | AGREE | paper.title='Human BioMolecular Atlas Program (HuBMAP): 3D Human Reference Atlas construction and usage'; gemma_final.classification='FILTERED'; gemma_q2_paper_profile.paper_type='OTHER'; gemma_q2_paper_profile.spatial='YES'; gemma_q2_paper_profile.panel_scope='UNCLEAR'; gemma_q1_panel_design blank. | accepted_agree |
| 10.1038/s41592-025-02617-2 | 2025 | Optimizing Xenium In Situ data utility by quality assessment and best-practice analysis workflows. | AGREE | paper title; gemma_final notes/source; empty q1 panel-design fields; q2 profile says OTHER/targeted and no panel evidence. | accepted_agree |
| 10.1038/s41592-025-02624-3 | 2025 | Feature selection methods affect the performance of scRNA-seq data integration and querying | AGREE | paper.title='Feature selection methods affect the performance of scRNA-seq data integration and querying'; gemma_final.classification='FILTERED'; gemma_q2_paper_profile.paper_type='COMP'; gemma_q2_paper_profile.spatial='NO'; gemma_q1_panel_design blank. | accepted_agree |
| 10.1038/s41592-025-02697-0 | 2025 | Cell simulation as cell segmentation | AGREE | paper title; gemma_final notes/source; empty q1 panel-design fields; q2 profile says COMP/spatial and no panel evidence. | accepted_agree |
| 10.1038/s41592-025-02726-y | 2025 | Characterizing protein sequence determinants of nuclear | AGREE | paper title; gemma_final notes/source; empty q1 panel-design fields; q2 profile says | accepted_agree |

| DOI | Year | Title | Audit decision | Evidence (verbatim quote) | Validation status / scope flag |
|---|---|---|---|---|---|
| | | condensates by high-throughput pooled imaging with CondenSeq. | | COMP/spatial and no panel evidence. | |
| 10.1038/s41592-025-02795-z | 2025 | Spatial gene expression at single-cell resolution from histology using deep learning with GHIST | AGREE | title: Spatial gene expression at single-cell resolution from histology using deep learning with GHIST; q2 paper_type=COMP, panel_scope=UNCLEAR, spatial=YES; q1 panel-design fields are empty. | accepted_agree |
| 10.1038/s41592-025-02814-z | 2025 | Nicheformer: a foundation model for single-cell and spatial omics. | AGREE | paper title; gemma_final notes/source; empty q1 panel-design fields; q2 profile says COMP/spatial and no panel evidence. | accepted_agree |
| 10.1038/s41592-025-02815-y | 2025 | From slices to deep dishes: spatial transcriptomics and translatomics of thick tissue blocks | AGREE | paper_type/doc_type evidence: EDITORIAL (News & views); gemma_final notes: Q0: EDITORIAL → FILTERED | accepted_agree |
| 10.1038/s41593-023-01254-2 | 2023 | Precise mapping of molecular and cellular brain changes in Alzheimer’s disease | AGREE | gemma_final notes/source; paper_profile is empty because the packet is non-article; no panel-design evidence is present. | accepted_agree |
| 10.1038/s41593-024-01806-0 | 2024 | Opportunities and challenges of single-cell and spatially resolved genomics methods for neuroscience discovery | AGREE | citation_screening_prior: labels=REV; gemma_final source=citation_screening_prefilter -> FILTERED; q1 panel-design fields are empty. | accepted_agree |
| 10.1038/s41593-025-02042-w | 2025 | FOXP genes regulate Purkinje cell diversity and cerebellar morphogenesis | AGREE | paper.title='FOXP genes regulate Purkinje cell diversity and cerebellar morphogenesis'; gemma_final.classification='FILTERED'; gemma_q2_paper_profile.paper_type='EXP'; gemma_q2_paper_profile.spatial='NO'; gemma_q1_panel_design blank. | accepted_agree |
| 10.1038/s41596-025-01248-3 | 2025 | Spatially resolved in situ profiling of mRNA life cycle at transcriptome scale in intact cells and tissues using STARmap PLUS, RIBOmap and TEMPOmap | AGREE | paper.title='Spatially resolved in situ profiling of mRNA life cycle at transcriptome scale in intact cells and tissues using STARmap PLUS, RIBOmap and TEMPOmap'; citation_screening_prior.labels='REV'; citation_screening_prior.tools='gps | accepted_agree |

| DOI | Year | Title | Audit decision | Evidence (verbatim quote) | Validation status / scope flag |
|---|---|---|---|---|---|
| | | | | FISH'; gemma_q0_document_type.doc_type='REVIEW'; gemma_final.classification='FILTERED'. | |
| 10.1038/s41597-022-01886-2 | 2023 | Brain Data Standards - A method for building data-driven cell-type ontologies | AGREE | title='Brain Data Standards - A method for building data-driven cell-type ontologies'; citation_screening_prior.labels='COMP'; gemma_final.source='citation_screening_prefilter' | accepted_agree |
| 10.1038/s41597-026-06707-4 | 2026 | SexTumorDB: a comprehensive resource of sex-dependent tumor landscape at single-cell resolution | AGREE | gemma_final notes say Pass-A abstract: non-spatial -> FILTERED; q2 paper_type=OTHER, spatial=NO, panel_scope=UNCLEAR; q1 panel-design fields are empty. | accepted_agree |
| 10.1038/s41597-026-06937-6 | 2026 | STHELAR, a multi-tissue dataset linking spatial transcriptomics and histology for cell type annotation. | AGREE | paper title; gemma_final notes/source; empty q1 panel-design fields; q2 profile says OTHER/targeted and no panel evidence. | accepted_agree |
| 10.1038/s41598-018-21541-4 | 2018 | Creating a 3D microbial and chemical snapshot of a human habitat. | NEEDS_MANUAL | spatial=NO; packet does not support an in-scope spatial panel-design record. Original audit evidence: gemma_q1_panel_design is UNCLEAR with evidence_classification=NOT FOUND and the paper profile does not describe a targeted gene panel. | validated_high_risk_manual_filtered |
| 10.1038/s41598-019-41951-2 | 2019 | Modeling Spatial Correlation of Transcripts with Application to Developing Pancreas. | AGREE | paper title; gemma_final notes/source; empty q1 panel-design fields; q2 profile says COMP/spatial and no panel evidence. | accepted_agree |
| 10.1038/s41598-020-75708-z | 202 | Spatial transcriptomics inferred from pathology whole-slide images links tumor heterogeneity to survival in breast and lung cancer | AGREE | paper title; gemma_final notes/source; empty q1 panel-design fields; q2 profile says COMP/spatial and no panel evidence. | accepted_agree |
| 10.1038/s41598-020-76439-x | 202 | ProbeDealer is a convenient tool for designing probes for highly | NEEDS_MANUAL | Final manual decision: FILTERED because the available evidence concerns probe/barcode/oligo | manual_final_filter_after_pdf_review |

| DOI | Year | Title | Audit decision | Evidence (verbatim quote) | Validation status / scope flag |
|---|---|---|---|---|---|
| | | multiplexed fluorescence in situ hybridization. | | design or single-gene detection, not final gene-panel selection. Original evidence: q1: ProbeDealer designs probes for multiplexed FISH; q2: default secondary probe sequences for MERFISH and seqFISH. | |
| 10.1038/s41598-022-14192-z | 2022 | Cell type matching in single-cell RNA-sequencing data using FR-Match | AGREE | title='Cell type matching in single-cell RNA-sequencing data using FR-Match'; citation_screening_prior.labels='COMP'; gemma_final.source='citation_screening_prefilter' | accepted_agree |
| 10.1038/s41598-022-19984-x | 2022 | Nondestructive protein sampling with electroporation facilitates profiling of spatial differential protein expression in breast tumors in vivo. | NEEDS_MANUAL | panel_scope=NOT_TARGETED with no target-gene panel source evidence; untargeted/no-panel records are out of scope. Original audit evidence: gemma_q1_panel_design is UNCLEAR and evidence_classification is NOT FOUND; the paper profile is not a targeted panel-design study. | validated_high_risk_manual_filtered |
| 10.1038/s41598-024-53557-4 | 2024 | Author Correction: Direct RNA targeted in situ sequencing for transcriptomic profiling in tissue | AGREE | paper_type/doc_type evidence: CORRECTION (Author Correction: Direct RNA targeted in situ sequencing for transcriptomic profiling in tissue); gemma_final notes: Q0: CORRECTION → FILTERED | accepted_agree |
| 10.1038/s41598-024-63492-z | 2024 | CellMarkerPipe: cell marker identification and evaluation pipeline in single cell transcriptomes | AGREE | citation_screening_prior: computational-only prior; gemma_final source=citation_screening_prefilter -> FILTERED; q1 panel-design fields are empty; q2 paper_type=COMP. | accepted_agree |
| 10.1038/s41598-024-65090-5 | 2024 | Comprehensive analysis of endoplasmic reticulum stress related signature in head and neck squamous carcinoma | AGREE | citation-screening prior labels=['NON-ST']; gemma_final notes: non-spatial prior pre-filter | accepted_agree |
| 10.1038/s41598-024-73356-1 | 2024 | scGAA: a general gated axial-attention model for accurate cell-type annotation of single-cell RNA-seq data | AGREE | paper_type/profile evidence: COMP; citation-screening prior labels=['COMP']; gemma_final | accepted_agree |

| DOI | Year | Title | Audit decision | Evidence (verbatim quote) | Validation status / scope flag |
|---|---|---|---|---|---|
| | | | | notes: computational-only prior pre-filter | |
| 10.1038/s41598-024-78076-0 | 2024 | Embracing the informative missingness and silent gene in analyzing biologically diverse samples | AGREE | paper_type/profile evidence: COMP; citation-screening prior labels=['COMP']; gemma_final notes: computational-only prior pre-filter | accepted_agree |
| 10.1038/s41598-025-14807-1 | 2025 | Single cell spatial transcriptomics links Wnt signaling disruption to extracellular matrix development in a cleft palate model. | AGREE | paper title; gemma_final notes/source; paper_profile marks NOT_TARGETED; q1 panel-design fields are empty. | accepted_agree |
| 10.1038/s41598-025-27732-0 | 2025 | Assessment of chemical risk factor for breast cancer based on gene expression profiles | AGREE | paper.title='Assessment of chemical risk factor for breast cancer based on gene expression profiles'; citation_screening_prior.labels='COMP'; citation_screening_prior.tools='COSG'; gemma_final.classification='FILTERED'; gemma_q1_panel_design blank; gemma_q2_paper_profile.paper_type='COMP'. | accepted_agree |
| 10.1038/s41598-026-39168-1 | 2026 | CiCLoDS: Joint cell clustering and gene selection for single-cell spatial transcriptomics | AGREE | paper title; citation-screening prior says computational-only; q1 panel-design fields are empty; no canonical tool evidence in packet. | accepted_agree |
| 10.1038/s42003-023-04820-3 | 2023 | MASI enables fast model-free standardization and integration of single-cell transcriptomics data | AGREE | title='MASI enables fast model-free standardization and integration of single-cell transcriptomics data'; citation_screening_prior.labels='COMP-BENCH'; gemma_final.source='citation_screening_prefilter' | accepted_agree |
| 10.1038/s42003-024-06349-5 | 2024 | SpatialPrompt: spatially aware scalable and accurate tool for spot deconvolution and domain identification in spatial transcriptomics. | AGREE | paper title; gemma_final notes/source; empty q1 panel-design fields; q2 profile says COMP/spatial and no panel evidence. | accepted_agree |
| 10.1038/s42003-024-06480-3 | 2024 | A point cloud segmentation framework for image-based spatial transcriptomics | DISAGREE | This computational segmentation paper uses NS-Forest-selected genes only to construct a simulation setup; it does not report the paper's own targeted panel-design event, so it should be | accepted_disagree |

| DOI | Year | Title | Audit decision | Evidence (verbatim quote) | Validation status / scope flag |
|---|---|---|---|---|---|
| | | | | filtered from the panel-design adoption set. | |
| 10.1038/s42003-025-07872-9 | 2025 | Infusing structural assumptions into dimensionality reduction for single-cell RNA sequencing data to identify small gene sets | AGREE | title: Infusing structural assumptions into dimensionality reduction for single-cell RNA sequencing data to identify small gene sets; q2 paper_type=COMP; q1 panel-design fields are empty; citation prior labels=COMP\|COMP-BENCH and Gemma_final=FILTERED. | accepted_agree |
| 10.1038/s42003-025-08485-y | 2025 | Cluster-independent multiscale marker identification in single-cell RNA-seq data using localized marker detector (LMD) | AGREE | paper.title='Cluster-independent multiscale marker identification in single-cell RNA-seq data using localized marker detector (LMD)'; citation_screening_prior.labels='COMP-BENCH'; citation_screening_prior.tools='COSG'; gemma_final.classification='FILTERED'; gemma_q1_panel_design blank. | accepted_agree |
| 10.1038/s43587-024-00631-1 | 2024 | Aging atlas reveals cell-type-specific effects of pro-longevity strategies | AGREE | citation-screening prior labels= ['NON-ST']; gemma_final notes: non-spatial prior pre-filter | accepted_agree |
| 10.1038/s43587-025-01027-5 | 2025 | Integrating polygenic signals and single-cell multiomics identifies cell-type-specific regulomes critical for immune- and aging-related diseases | AGREE | paper.title='Integrating polygenic signals and single-cell multiomics identifies cell-type-specific regulomes critical for immune- and aging-related diseases'; citation_screening_prior.labels='COMP'; citation_screening_prior.tools='COSG'; gemma_final.classification='FILTERED'; gemma_q1_panel_design blank. | accepted_agree |
| 10.1038/s43588-021-00070-7 | 2021 | Single-cell manifold-preserving feature selection for detecting rare cell populations | AGREE | Title: 'Single-cell manifold-preserving feature selection for detecting rare cell populations'; packet fields show COMP prior, no spatial evidence, and Gemma's FILTERED prefilter. | accepted_agree |
| 10.1038/s43588-024-00689-2 | 2024 | Biophysically interpretable inference of cell types from multimodal sequencing data | AGREE | paper_type/profile evidence: COMP; citation-screening prior labels=['COMP']; gemma_final | accepted_agree |

| DOI | Year | Title | Audit decision | Evidence (verbatim quote) | Validation status / scope flag |
|---|---|---|---|---|---|
| | | | | notes: computational-only prior pre-filter | |
| 10.1038/s44161-024-00464-6 | 2024 | Strategies for arterial graft optimization at the single-cell level | AGREE | citation-screening prior labels=['NON-ST']; gemma_final notes: non-spatial prior pre-filter | accepted_agree |
| 10.1038/s44320-023-00006-5 | 2023 | Identifying Spatial Co-occurrence in Healthy and InflAmed tissues (ISCHIA) | AGREE | paper title; gemma_final notes/source; empty q1 panel-design fields; q2 profile says COMP/spatial and no panel evidence. | accepted_agree |
| 10.1038/s44320-025-00176-4 | 2025 | CANTAO: guiding clustering and annotation in single-cell RNA sequencing using average overlap | AGREE | paper.title='CANTAO: guiding clustering and annotation in single-cell RNA sequencing using average overlap'; citation_screening_prior.labels='COMP'; citation_screening_prior.tools='RankCorr'; gemma_final.classification='FILTERED'; gemma_q1_panel_design blank. | accepted_agree |
| 10.1039/d5sc09638c | 2026 | cShot: spatial dynamic imaging of cell cycle-dependent miRNA heterogeneity using dynamic DNA patterns. | AGREE | paper title; gemma_final notes/source; empty q1 panel-design fields; q2 profile says COMP/spatial and no panel evidence. | accepted_agree |
| 10.1042/bsr20171266 | 2018 | Increased sympathovagal imbalance evaluated by heart rate variability is associated with decreased T2* MRI and left ventricular function in transfusion-dependent thalassemia patients | AGREE | Title; gemma_final=FILTERED; gemma_q2_paper_profile spatial=NO, panel_scope=UNCLEAR; risk_flags=prefiltered_without_full_panel_qs. | accepted_agree |
| 10.1042/bsr20221680 | 2022 | Spatial RNA sequencing methods show high resolution of single cell in cancer metastasis and the formation of tumor microenvironment | AGREE | title='Spatial RNA sequencing methods show high resolution of single cell in cancer metastasis and the formation of tumor microenvironment'; citation_screening_prior.labels='REV'; gemma_final.source='citation_screening_prefilter' | accepted_agree |
| 10.1053/j.gastro.2024.07.039 | 2024 | A MERFISHing Expedition for Colitis | AGREE | gemma_q0_document_type doc_type=NON_ARTICLE, doc_confidence=HIGH, doc_evidence=NOT FOUND; | accepted_agree |

| DOI | Year | Title | Audit decision | Evidence (verbatim quote) | Validation status / scope flag |
|---|---|---|---|---|---|
| | | | | gemma_final=FILTERED; risk_flags=non_primary_document. | |
| 10.1053/j.gastro.2025.09.043 | 2026 | Single-Cell Analysis of Chemotherapy-induced Remodeling Reveals CD276-driven Basal-like Chemoresistance in Pancreatic Cancer. | AGREE | Title; gemma_q2_paper_profile spatial=YES, panel_scope=NOT_TARGETED, paper_type=EXP; gemma_final=FILTERED; risk_flags=prefiltered_without_full_panel_qs. | accepted_agree |
| 10.1073/pnas.1424409112 | 2015 | Molecular cartography of the human skin surface in 3D. | AGREE | q2 spatial: NO; q1 panel_design: NOT FOUND. | accepted_agree |
| 10.1073/pnas.2004997117 | 202 | Profile of Xiaowei Zhuang, winner of the 2020 Vilcek Prize in Biomedical Science | AGREE | Title; gemma_q2_paper_profile spatial=YES, panel_scope=TARGETED, paper_type=OTHER; gemma_final=FILTERED; risk_flags=prefiltered_without_full_panel_qs. | accepted_agree |
| 10.1073/pnas.2304730120 | 2023 | split-intein Gal4 provides intersectional genetic labeling that is repressible by Gal80 | AGREE | title='split-intein Gal4 provides intersectional genetic labeling that is repressible by Gal80'; citation_screening_prior.labels='NON-ST'; gemma_final.source='citation_screening_prefilter' | accepted_agree |
| 10.1073/pnas.2503952122 | 2025 | A unified framework for identification of cell-type-specific spatially variable genes in spatial transcriptomic studies. | AGREE | Title; gemma_q2_paper_profile spatial=YES, panel_scope=UNCLEAR, paper_type=COMP; gemma_final=FILTERED; risk_flags=prefiltered_without_full_panel_qs. | accepted_agree |
| 10.1073/pnas.2511483123 | 2026 | Lipid nanoparticle GM-CSF replacement for autoimmune pulmonary alveolar proteinosis | AGREE | Title; gemma_final=FILTERED; gemma_q2_paper_profile spatial=NO, panel_scope=UNCLEAR, paper_type=EXP; risk_flags=prefiltered_without_full_panel_qs. | accepted_agree |
| 10.1073/pnas.2517283123 | 2026 | Dissecting spatial patterning and signaling with directional diffusion in spatial multi-omics | AGREE | citation_screening_prior labels=COMP and all_tools=COSG; gemma_final source=citation_screening_prefilter -> FILTERED; q1 panel-design fields are empty. | accepted_agree |

| DOI | Year | Title | Audit decision | Evidence (verbatim quote) | Validation status / scope flag |
|---|---|---|---|---|---|
| 10.1080/00268976.2023.2297818 | 2023 | The statistical physics of single cell genomics | AGREE | title='The statistical physics of single cell genomics'; citation_screening_prior.labels='REV'; gemma_final.source='citation_screening_prefilter' | accepted_agree |
| 10.1080/01621459.2024.2342639 | 2024 | Dissecting Gene Expression Heterogeneity: Generalized Pearson Correlation Squares and the <i>K</i> -Lines Clustering Algorithm | AGREE | paper_type/profile evidence: COMP; citation-screening prior labels=['COMP']; gemma_final notes: computational-only prior pre-filter | accepted_agree |
| 10.1080/07853890.2024.2443568 | 2024 | A diagnostic signatures for intervertebral disc degeneration using TNFAIP6 and COL6A2 based on single-cell RNA-seq and bulk RNA-seq analyses | AGREE | citation-screening prior labels= ['NON-ST']; gemma_final notes: non-spatial prior pre-filter | accepted_agree |
| 10.1080/19336934.2024.2409968 | 2024 | Optimization of hybridization chain reaction for imaging single RNA molecules in <i>Drosophila</i> larvae. | AGREE | Title; gemma_q2_paper_profile spatial=YES, panel_scope=UNCLEAR, paper_type=OTHER; gemma_final=FILTERED; risk_flags=prefiltered_without_full_panel_qs. | accepted_agree |
| 10.1080/19336934.2024.2428499 | 2025 | A fast <i>in situ</i> hybridization chain reaction method in <i>Drosophila</i> embryos and ovaries. | AGREE | Title; gemma_q2_paper_profile spatial=YES, panel_scope=UNCLEAR, paper_type=OTHER; gemma_final=FILTERED; risk_flags=prefiltered_without_full_panel_qs. | accepted_agree |
| 10.1080/21541264.2023.2199669 | 2023 | Understanding spatiotemporal coupling of gene expression using single molecule RNA imaging technologies | AGREE | Title; gemma_q2_paper_profile spatial=YES, panel_scope=UNCLEAR, paper_type=OTHER; gemma_final=FILTERED; risk_flags=prefiltered_without_full_panel_qs. | accepted_agree |
| 10.1089/cmb.2024.0532 | 2024 | SpatialCTD: A Large-Scale Tumor Microenvironment Spatial Transcriptomic Dataset to Evaluate Cell Type Deconvolution for Immuno-Oncology. | AGREE | Title; gemma_q2_paper_profile spatial=YES, panel_scope=UNCLEAR, paper_type=COMP; gemma_final=FILTERED; risk_flags=prefiltered_without_full_panel_qs. | accepted_agree |

| DOI | Year | Title | Audit decision | Evidence (verbatim quote) | Validation status / scope flag |
|---|---|---|---|---|---|
| 10.1089/genedge.6.01.157 | 2024 | Vizgen, Ultivue Merge, Further Consolidating Spatial Biology | AGREE | gemma_q0_document_type doc_type=NON_ARTICLE, doc_evidence=FORM 10-Q, doc_confidence=HIGH; gemma_final=FILTERED; risk_flags=non_primary_document. | accepted_agree |
| 10.1093/abt/tbaf017 | 2025 | Reprogramming the tumor microenvironment: synergistic mechanisms of antibody–drug conjugates and immune checkpoint inhibitors | AGREE | paper.title='Reprogramming the tumor microenvironment: synergistic mechanisms of antibody-drug conjugates and immune checkpoint inhibitors'; citation_screening_prior.labels='REV'; citation_screening_prior.tools='PERSIST'; gemma_q0_document_type.doc_type='REVIEW'; gemma_final.classification='FILTERED'. | accepted_agree |
| 10.1093/ajcp/aqy102.314 | 2018 | Use of Mate-Pair Sequencing (MPseq) to Elucidate a Complex BCR-ABL1 Rearrangement Observed in a Newly Diagnosed Case of Chronic Myeloid Leukemia | AGREE | Title; gemma_final=FILTERED; gemma_q2_paper_profile spatial=NO, panel_scope=UNCLEAR, paper_type=EXP; risk_flags=prefiltered_without_full_panel_qs. | accepted_agree |
| 10.1093/bfgp/elad004 | 2023 | SCMcluster: a high-precision cell clustering algorithm integrating marker gene set with single-cell RNA sequencing data | AGREE | title='SCMcluster: a high-precision cell clustering algorithm integrating marker gene set with single-cell RNA sequencing data'; citation_screening_prior.labels='COMP'; gemma_final.source='citation_screening_prefilter' | accepted_agree |
| 10.1093/bfgp/elaf018 | 2025 | VARGG: a deep learning framework advancing precise spatial domain identification and cellular heterogeneity analysis in spatial transcriptomics | AGREE | Title; gemma_q2_paper_profile spatial=YES, panel_scope=UNCLEAR, paper_type=COMP; gemma_final=FILTERED; risk_flags=prefiltered_without_full_panel_qs. | accepted_agree |
| 10.1093/bib/bbaa339 | 202 | FR-Match: robust matching of cell type clusters from single cell RNA sequencing data using the Friedman–Rafsky non-parametric test | AGREE | Title: 'FR-Match: robust matching of cell type clusters from single cell RNA sequencing data using the Friedman–Rafsky non-parametric test'; packet fields show | accepted_agree |

| DOI | Year | Title | Audit decision | Evidence (verbatim quote) | Validation status / scope flag |
|---|---|---|---|---|---|
| | | | | paper_type=COMP and Gemma's FILTERED prefilter. | |
| 10.1093/bib/bbac191 | 2022 | Artificial intelligence and machine learning approaches using gene expression and variant data for personalized medicine | AGREE | title='Artificial intelligence and machine learning approaches using gene expression and variant data for personalized medicine'; citation_screening_prior.labels='REV'; gemma_final.source='citation_screening_prefilter' | accepted_agree |
| 10.1093/bib/bbac563 | 2023 | spaCI: deciphering spatial cellular communications through adaptive graph model. | AGREE | The colon dataset (Colon cancer 2) contains a MERFISH measurement of a 500 gene panel. | accepted_agree |
| 10.1093/bib/bbac585 | 2023 | scGMAAE: Gaussian mixture adversarial autoencoders for diversification analysis of scRNA-seq data | AGREE | title='scGMAAE: Gaussian mixture adversarial autoencoders for diversification analysis of scRNA-seq data'; citation_screening_prior.labels='COMP'; gemma_final.source='citation_screening_prefilter' | accepted_agree |
| 10.1093/bib/bbac625 | 2023 | scDCCA: deep contrastive clustering for single-cell RNA-seq data based on auto-encoder network | AGREE | title='scDCCA: deep contrastive clustering for single-cell RNA-seq data based on auto-encoder network'; citation_screening_prior.labels='COMP'; gemma_final.source='citation_screening_prefilter' | accepted_agree |
| 10.1093/bib/bbad338 | 2023 | SpaRx: elucidate single-cell spatial heterogeneity of drug responses for personalized treatment. | AGREE | the 853 and 500 genes are selected based on the RNA panels used in the NanoString CosMx and the Vizgen MERSCOPE data, respectively. | accepted_agree |
| 10.1093/bib/bbad389 | 2023 | FEED: a feature selection method based on gene expression decomposition for single cell clustering | AGREE | Title is 'FEED: a feature selection method based on gene expression decomposition for single cell clustering'; citation prior marks it computational-only and q1 panel-design fields are empty. | accepted_agree |
| 10.1093/bib/bbad416 | 2023 | An in-depth comparison of linear and non-linear joint embedding methods for bulk and single-cell multi-omics | AGREE | title='An in-depth comparison of linear and non-linear joint embedding methods for bulk and single-cell multi-omics'; citation_screening_prior.labels='REV'; | accepted_agree |

| DOI | Year | Title | Audit decision | Evidence (verbatim quote) | Validation status / scope flag |
|---|---|---|---|---|---|
| | | | | gemma_final.source='citation_screening_prefilter' | |
| 10.1093/bib/bbad424 | 2023 | Identifying phenotype-associated subpopulations through LP_SGL | AGREE | title='Identifying phenotype-associated subpopulations through LP_SGL'; citation_screening_prior.labels='COMP'; gemma_final.source='citation_screening_prefilter' | accepted_agree |
| 10.1093/bib/bbad485 | 2023 | <i>methylClass</i>: an R package to construct DNA methylation-based classification models | AGREE | title='<i>methylClass</i>: an R package to construct DNA methylation-based classification models'; citation_screening_prior.labels='COMP-BENCH'; gemma_final.source='citation_screening_prefilter' | accepted_agree |
| 10.1093/bib/bbad525 | 2024 | scCorrector: a robust method for integrating multi-study single-cell data. | NEEDS_MANUAL | Integrates scRNA-seq with MERFISH and says it can "extend the gene coverage" of MERFISH; no panel-design source is given. | pdf_validated_unclear_update |
| 10.1093/bib/bbae018 | 2024 | scHybridBERT: integrating gene regulation and cell graph for spatiotemporal dynamics in single-cell clustering | AGREE | paper_type/profile evidence: COMP; citation-screening prior labels=['COMP']; gemma_final notes: computational-only prior pre-filter | accepted_agree |
| 10.1093/bib/bbae422 | 2024 | ReCIDE: robust estimation of cell type proportions by integrating single-reference-based deconvolutions | AGREE | paper_type/profile evidence: COMP; citation-screening prior labels=['COMP']; gemma_final notes: computational-only prior pre-filter | accepted_agree |
| 10.1093/bib/bbae572 | 2024 | Identifying cell types by lasso-constraint regularized Gaussian graphical model based on weighted distance penalty | AGREE | paper_type/profile evidence: COMP; citation-screening prior labels=['COMP']; gemma_final notes: computational-only prior pre-filter | accepted_agree |
| 10.1093/bib/bbae626 | 2024 | CosGeneGate selects multi-functional and credible biomarkers for single-cell analysis | AGREE | Title is 'CosGeneGate selects multi-functional and credible biomarkers for single-cell analysis'; citation prior marks it computational-only and q1 panel-design fields are empty. | accepted_agree |
| 10.1093/bib/bbae703 | 2024 | Addressing scalability and managing sparsity and dropout | AGREE | paper_type/profile evidence: COMP; citation-screening prior labels=['COMP']; gemma_final | accepted_agree |

| DOI | Year | Title | Audit decision | Evidence (verbatim quote) | Validation status / scope flag |
|---|---|---|---|---|---|
| | | events in single-cell representation identification with ZIGACL | | notes: computational-only prior pre-filter | |
| 10.1093/bib/bbaf020 | 2024 | Graph contrastive learning of subcellular-resolution spatial transcriptomics improves cell type annotation and reveals critical molecular pathways. | NEEDS_MANUAL | Benchmarks MERFISH, CosMx SMI, and Xenium datasets; focuses on cell-type annotation, not choosing a gene panel. | pdf_validated_unclear_update |
| 10.1093/bib/bbaf085 | 2025 | Learning directed acyclic graphs for ligands and receptors based on spatially resolved transcriptomic data of ovarian cancer | AGREE | Title; gemma_q2_paper_profile spatial=YES, panel_scope=UNCLEAR, paper_type=COMP; gemma_final=FILTERED; risk_flags=prefiltered_without_full_panel_qs. | accepted_agree |
| 10.1093/bib/bbaf193 | 2025 | Multi-view clustering for single-cell RNA-seq data based on graph fusion | AGREE | paper.title='Multi-view clustering for single-cell RNA-seq data based on graph fusion'; citation_screening_prior.labels='COMP'; citation_screening_prior.tools='COSG'; gemma_final.classification='FILTERED'; gemma_q1_panel_design blank. | accepted_agree |
| 10.1093/bib/bbaf306 | 2025 | Artifacts in spatial transcriptomics data: their detection, importance, prevalence, and prevention. | AGREE | Title; gemma_q2_paper_profile spatial=YES, panel_scope=UNCLEAR, paper_type=COMP; gemma_final=FILTERED; risk_flags=prefiltered_without_full_panel_qs. | accepted_agree |
| 10.1093/bib/bbaf405 | 2025 | MuST: multiple-modality structure transformation for single-cell spatial transcriptomics | AGREE | paper.title='MuST: multiple-modality structure transformation for single-cell spatial transcriptomics'; citation_screening_prior.labels='COMP'; citation_screening_prior.tools='scGeneFit'; gemma_final.classification='FILTERED'; gemma_q1_panel_design blank; gemma_q2_paper_profile.paper_type='COMP'. | accepted_agree |
| 10.1093/bib/bbaf478 | 2025 | Multi-view gene panel characterization for spatially | DISAGREE | This is a computational panel-characterization and benchmarking | validated_high_risk_disagreement |

| DOI | Year | Title | Audit decision | Evidence (verbatim quote) | Validation status / scope flag |
|---|---|---|---|---|---|
| | | resolved omics | | paper. Although panelScope-OA generates optimized panels inside the method study, this is not an experimental paper-level adoption event and should be filtered from the adoption set. | |
| 10.1093/bib/bbaf528 | 2025 | Denoising spatially resolved transcriptomics with consistency of heterogeneous spatial coordinates, transcription, and morphology. | AGREE | Title; gemma_q2_paper_profile spatial=YES, panel_scope=UNCLEAR, paper_type=COMP; gemma_final=FILTERED; risk_flags=prefiltered_without_full_panel_qs. | accepted_agree |
| 10.1093/bib/bbaf631.003 | 2025 | CoSMIM: a covariance-based multi-scale framework for integrated spatial microenvironment analysis | AGREE | Title; gemma_q2_paper_profile spatial=YES, panel_scope=UNCLEAR, paper_type=COMP; gemma_final=FILTERED; risk_flags=prefiltered_without_full_panel_qs. | accepted_agree |
| 10.1093/bib/bbaf631.005 | 2025 | Image-free estimation of cell locations and types in spatial transcriptomics via quadtree partitioning and probabilistic clustering | AGREE | Title: 'Image-free estimation of cell locations and types in spatial transcriptomics via quadtree partitioning and probabilistic clustering'; packet fields show Q0: NON_ARTICLE and Gemma's FILTERED prefilter. | accepted_agree |
| 10.1093/bib/bbaf675 | 2025 | scHLens: a web server for hierarchically and interactively exploring single cell RNA-seq data | AGREE | paper.title='scHLens: a web server for hierarchically and interactively exploring single cell RNA-seq data'; citation_screening_prior.labels='COMP'; citation_screening_prior.tools='COSG'; gemma_final.classification='FILTERED'; gemma_q1_panel_design blank. | accepted_agree |
| 10.1093/bib/bbaf723 | 2026 | Advances and challenges in single-cell RNA sequencing data analysis: a comprehensive review | AGREE | citation_screening_prior labels=REV and all_tools=Spapros; gemma_final source=citation_screening_prefilter -> FILTERED; q1 panel-design fields are empty. | accepted_agree |
| 10.1093/bib/bbag072 | 2026 | Imputing missing values in single-cell RNA-sequencing data: a | AGREE | citation_screening_prior labels=COMP and | accepted_agree |

| DOI | Year | Title | Audit decision | Evidence (verbatim quote) | Validation status / scope flag |
|---|---|---|---|---|---|
| | | statistical and machine learning-based approach | | all_tools=scGeneFit; gemma_final source=citation_screening_prefilter -> FILTERED; q1 panel-design fields are empty. | |
| 10.1093/bib/bbag094 | 2026 | scSCCNIA: similarity matrix based contrastive clustering with neighbor information aggregation for single-cell RNA sequencing data | AGREE | citation_screening_prior labels=COMP and all_tools=COSG; gemma_final source=citation_screening_prefilter -> FILTERED; q1 panel-design fields are empty. | accepted_agree |
| 10.1093/bioadv/vbac048 | 2022 | Sparcle: assigning transcripts to cells in multiplexed images. | NEEDS_MANUAL | Sparcle assigns transcripts to cells in MERFISH/smFISH/STARmap datasets; the text emphasizes segmentation/assignment, not panel design. | pdf_validated_unclear_update |
| 10.1093/bioadv/vbae064 | 2024 | Uncover spatially informed variations for single-cell spatial transcriptomics with STew. | AGREE | Title; gemma_q2_paper_profile spatial=YES, panel_scope=UNCLEAR, paper_type=COMP; gemma_final=FILTERED; risk_flags=prefiltered_without_full_panel_qs. | accepted_agree |
| 10.1093/bioadv/vbae081 | 2024 | CellsFromSpace: a fast, accurate, and reference-free tool to deconvolve and annotate spatially distributed omics data. | AGREE | Title; gemma_q2_paper_profile spatial=YES, panel_scope=UNCLEAR, paper_type=COMP; gemma_final=FILTERED; risk_flags=prefiltered_without_full_panel_qs. | accepted_agree |
| 10.1093/bioadv/vbaf167 | 2024 | <i>ScaleSC</i> : a superfast and scalable single-cell RNA-seq data analysis pipeline powered by GPU | AGREE | paper_type/profile evidence: COMP; citation-screening prior labels=['COMP']; gemma_final notes: computational-only prior pre-filter | accepted_agree |
| 10.1093/bioadv/vbag047 | 2026 | Sparse dimensionality reduction for analyzing single-cell-resolved interactions | AGREE | citation_screening_prior labels=COMP and all_tools=scPNMF; gemma_final source=citation_screening_prefilter -> FILTERED; q1 panel-design fields are empty. | accepted_agree |
| 10.1093/bioadv/vbag080 | 2026 | PeakPrime: a peak-guided primer design pipeline for target enrichment in 3′-end RNA-seq | AGREE | title: PeakPrime: a peak-guided primer design pipeline for target enrichment in 3'-end RNA-seq; q2 paper_type=COMP; q1 panel-design fields are empty; citation | accepted_agree |

| DOI | Year | Title | Audit decision | Evidence (verbatim quote) | Validation status / scope flag |
|---|---|---|---|---|---|
| | | | | prior labels=COMP and Gemma_final=FILTERED. | |
| 10.1093/bioinformatics/btac263 | 2022 | Self-supervised learning of cell type specificity from immunohistochemical images | AGREE | title='Self-supervised learning of cell type specificity from immunohistochemical images'; citation_screening_prior.labels='COMP'; gemma_final.source='citation_screening_prefilter' | accepted_agree |
| 10.1093/bioinformatics/btac299 | 2022 | SpatialExperiment: infrastructure for spatially-resolved transcriptomics data in R using Bioconductor. | AGREE | Title; gemma_q2_paper_profile spatial=YES, panel_scope=UNCLEAR, paper_type=COMP; gemma_final=FILTERED; risk_flags=prefiltered_without_full_panel_qs. | accepted_agree |
| 10.1093/bioinformatics/btac605 | 2022 | SD2: spatially resolved transcriptomics deconvolution through integration of dropout and spatial information. | AGREE | Title: 'SD2: spatially resolved transcriptomics deconvolution through integration of dropout and spatial information.' Packet metadata: paper_type=COMP, spatial=YES, panel_scope=UNCLEAR, gemma_q1_panel_design empty. | accepted_agree |
| 10.1093/bioinformatics/btac683 | 2022 | AIscEA: unsupervised integration of single-cell gene expression and chromatin accessibility via their biological consistency | AGREE | title='AIscEA: unsupervised integration of single-cell gene expression and chromatin accessibility via their biological consistency'; citation_screening_prior.labels='COMP'; gemma_final.source='citation_screening_prefilter' | accepted_agree |
| 10.1093/bioinformatics/btad216 | 2023 | CellBRF: a feature selection method for single-cell clustering using cell balance and random forest | AGREE | citation_screening_prior: computational-only prior; gemma_final source=citation_screening_prefilter -> FILTERED; q1 panel-design fields are empty; q2 paper_type=COMP. | accepted_agree |
| 10.1093/bioinformatics/btad269 | 2023 | CLARIFY: cell-cell interaction and gene regulatory network refinement from spatially resolved transcriptomics. | NEEDS_MANUAL | CLARIFY infers GRNs/CCIs from seqFISH and MERFISH datasets; it is downstream analysis, not panel selection. | pdf_validated_unclear_update |
| 10.1093/bioinformatics/btad550 | 2023 | MoleculeExperiment enables consistent infrastructure for | AGREE | Title: 'MoleculeExperiment enables consistent infrastructure for | accepted_agree |

| DOI | Year | Title | Audit decision | Evidence (verbatim quote) | Validation status / scope flag |
|---|---|---|---|---|---|
| | | molecule-resolved spatial omics data in bioconductor. | | molecule-resolved spatial omics data in Bioconductor.' Packet metadata: paper_type=COMP, spatial=YES, panel_scope=UNCLEAR, gemma_q1_panel_design empty. | |
| 10.1093/bioinformatics/btae518 | 2024 | scDiffusion: conditional generation of high-quality single-cell data using diffusion model | AGREE | paper_type/profile evidence: COMP; citation-screening prior labels=['COMP']; gemma_final notes: computational-only prior pre-filter | accepted_agree |
| 10.1093/bioinformatics/btaf142 | 2025 | HISSTA: a human in situ single-cell transcriptome atlas. | NEEDS_MANUAL | PDF: HISSTA is a database/atlas; marker genes are for annotation, not panel design. | pdf_validated_unclear_update |
| 10.1093/bioinformatics/btaf194 | 2025 | Refinement strategies for Tangram for reliable single-cell to spatial mapping | AGREE | title: Refinement strategies for Tangram for reliable single-cell to spatial mapping; citation prior labels=COMP-BENCH; q1 panel-design fields are empty; gemma_final=FILTERED. | accepted_agree |
| 10.1093/bioinformatics/btaf246 | 2025 | TissueViewer: a web-based multiplexed image viewer. | AGREE | Title: 'TissueViewer: a web-based multiplexed image viewer.' Packet metadata: paper_type=COMP, spatial=YES, panel_scope=UNCLEAR, gemma_q1_panel_design empty. | accepted_agree |
| 10.1093/bioinformatics/btaf451 | 2025 | Mitigation of multi-scale biases in cell-type deconvolution for spatially resolved transcriptomics using HarmoDecon. | AGREE | Title: 'Mitigation of multi-scale biases in cell-type deconvolution for spatially resolved transcriptomics using HarmoDecon.' Packet metadata: paper_type=COMP, spatial=YES, panel_scope=UNCLEAR, gemma_q1_panel_design empty. | accepted_agree |
| 10.1093/bioinformatics/btaf619 | 2025 | STAHD: a scalable and accurate method to detect spatial domains in high-resolution spatial transcriptomics data | AGREE | title: STAHD: a scalable and accurate method to detect spatial domains in high-resolution spatial transcriptomics data; citation prior labels=COMP; q1 panel-design fields are empty; gemma_final=FILTERED. | accepted_agree |
| 10.1093/bioinformatics/btag051 | 2026 | Inference of marker genes of subtle cell state changes via iLR: iterative logistic regression | NEEDS_MANUAL | spatial=NO; packet does not support an in-scope spatial panel-design record. Original audit evidence: Inference of marker | validated_high_risk_manual_filtered |

| DOI | Year | Title | Audit decision | Evidence (verbatim quote) | Validation status / scope flag |
|---|---|---|---|---|---|
| | | | | genes of subtle cell state changes via iLR: iterative logistic regression | |
| 10.1093/bioinformatics/bty718 | 2019 | A Bayesian model for single cell transcript expression analysis on MERFISH data. | NEEDS_MANUAL | doc_type=SUPPLEMENT; packet does not provide usable main-article panel-design evidence. Original audit evidence: q0: SUPPLEMENT; q1/q2 fields are empty. | validated_high_risk_manual_filtered |
| 10.1093/biomtc/ujae066 | 2024 | An interpretable Bayesian clustering approach with feature selection for analyzing spatially resolved transcriptomics data | AGREE | Title: 'An interpretable Bayesian clustering approach with feature selection for analyzing spatially resolved transcriptomics data.' Packet metadata: paper_type=COMP, spatial=YES, panel_scope=UNCLEAR, gemma_q1_panel_design empty. | accepted_agree |
| 10.1093/brain/awae030 | 2024 | Characterization of cortico-meningeal translocator protein expression in multiple sclerosis. | NEEDS_MANUAL | Final manual decision: FILTERED because no local PDF was available and the packet did not provide usable final gene-panel source evidence. Original evidence: q1 panel_design: NOT FOUND; q2 spatial: YES. | manual_final_filter_after_pdf_review |
| 10.1093/cei/uxae077 | 2024 | Implementing distinct spatial proteogenomic technologies: opportunities, challenges, and key considerations | AGREE | paper_type/doc_type evidence: REVIEW; citation-screening prior labels=['REV']; gemma_final notes: citation-screening prior pre-filter | accepted_agree |
| 10.1093/ckj/sfw140 | 2017 | Achieving high convection volumes in postdilution online hemodiafiltration: A prospective multicenter study | AGREE | Title: 'Achieving high convection volumes in postdilution online hemodiafiltration: A prospective multicenter study.' Packet metadata: paper_type=EXP, spatial=NO, panel_scope=UNCLEAR, gemma_q1_panel_design empty. | accepted_agree |
| 10.1093/ecco-jcc/jjad212.0024 | 2024 | OP24 SPP1 in colitis-associated colon cancer | AGREE | Title: 'OP24 SPP1 in colitis-associated colon cancer.' Packet metadata: Q0=NON_ARTICLE, gemma_final=FILTERED. | accepted_agree |

| DOI | Year | Title | Audit decision | Evidence (verbatim quote) | Validation status / scope flag |
|---|---|---|---|---|---|
| 10.1093/ecco-jcc/jjad212.0127 | 2024 | DOP87 Characterization of pathologic fibroblast and macrophage cell populations in colonic Crohn’s Disease with spatial transcriptomics | AGREE | Title: 'DOP87 Characterization of pathologic fibroblast and macrophage cell populations in colonic Crohn's Disease with spatial transcriptomics.' Packet metadata: Q0=NON_ARTICLE, gemma_final=FILTERED. | accepted_agree |
| 10.1093/ecco-jcc/jjad212.0227 | 2024 | P097 Spatially resolved insights into fistulating Crohn's disease pathogenesis: Unveiling molecular heterogeneity | AGREE | Title: 'P097 Spatially resolved insights into fistulating Crohn's disease pathogenesis: Unveiling molecular heterogeneity.' Packet metadata: Q0=NON_ARTICLE, gemma_final=FILTERED. | accepted_agree |
| 10.1093/ecco-jcc/jjaf231.1584 | 2026 | Obesity-Driven Fibroblast Remodelling in Intestinal Fibrosis: A Multi-Omic Approach | AGREE | Title: 'Obesity-Driven Fibroblast Remodelling in Intestinal Fibrosis: A Multi-Omic Approach.' Packet metadata: Q0=NON_ARTICLE, gemma_final=FILTERED. | accepted_agree |
| 10.1093/ecco-jcc/jjaf231.236 | 2026 | P0055 Novel KRT17+ epithelium-neutrophil interactions emerge in inflamed IBD mucosa | AGREE | Title: 'P0055 Novel KRT17+ epithelium-neutrophil interactions emerge in inflamed IBD mucosa.' Packet metadata: Q0=NON_ARTICLE, gemma_final=FILTERED. | accepted_agree |
| 10.1093/geroni/igad104.0731 | 2023 | SPATIAL PROFILING CELLULAR SENESCENCE IN THE HUMAN BRAIN ACROSS EIGHT DECADES OF LIFE | AGREE | Title: 'SPATIAL PROFILING CELLULAR SENESCENCE IN THE HUMAN BRAIN ACROSS EIGHT DECADES OF LIFE.' Packet metadata: Q0=NON_ARTICLE, gemma_final=FILTERED. | accepted_agree |
| 10.1093/geroni/igad104.1555 | 2023 | SINGLE-CELL AND SPATIAL OMICS FOR MAPPING CELLULAR SENESCENCE IN HEALTH, AGING AND DISEASE | NEEDS_MANUAL | PDF: review/perspective on single-cell and spatial omics for cellular senescence. | pdf_validated_unclear_update |
| 10.1093/geroni/igae098.1144 | 2024 | HIGH RESOLUTION SPATIAL PROTEOGENOMICS TO ASSESS TARGET ENGAGEMENT AND COMPARE SENOLYTIC EFFICACY IN THE BRAIN OF TAU TRANSGENIC MICE | AGREE | Title: 'HIGH RESOLUTION SPATIAL PROTEOGENOMICS TO ASSESS TARGET ENGAGEMENT AND COMPARE SENOLYTIC EFFICACY IN THE BRAIN OF TAU TRANSGENIC MICE.' Packet metadata: Q0=NON_ARTICLE, gemma_final=FILTERED. | accepted_agree |
| 10.1093/gigascience/giad113 | 2024 | Computational reproducibility of Jupyter notebooks from biomedical | AGREE | paper_type/profile evidence: COMP; citation-screening prior | accepted_agree |

| DOI | Year | Title | Audit decision | Evidence (verbatim quote) | Validation status / scope flag |
|---|---|---|---|---|---|
| | | publications | | labels=['COMP']; gemma_final notes: computational-only prior pre-filter | |
| 10.1093/gigascience/giae089 | 2024 | stMMR: accurate and robust spatial domain identification from spatially resolved transcriptomics with multimodal feature representation | AGREE | paper_type/profile evidence: COMP; citation-screening prior labels=['COMP']; gemma_final notes: computational-only prior pre-filter | accepted_agree |
| 10.1093/gigascience/giae103 | 2025 | Unveiling patterns in spatial transcriptomics data: a novel approach utilizing graph attention autoencoder and multiscale deep subspace clustering network. | AGREE | Title: 'Unveiling patterns in spatial transcriptomics data: a novel approach utilizing graph attention autoencoder and multiscale deep subspace clustering network.' Packet metadata: paper_type=COMP, spatial=YES, panel_scope=NOT_TARGETED, gemma_q1_panel_design empty. | accepted_agree |
| 10.1093/gigascience/giaf002 | 2025 | Dimensionality reduction for visualizing spatially resolved profiling data using SpaSNE. | AGREE | Title: 'Dimensionality reduction for visualizing spatially resolved profiling data using SpaSNE.' Packet metadata: paper_type=COMP, spatial=YES, panel_scope=UNCLEAR, gemma_q1_panel_design empty. | accepted_agree |
| 10.1093/gigascience/giaf090 | 2025 | SPEX: A modular end-to-end platform for high-plex tissue spatial omics analysis | AGREE | Title: 'SPEX: A modular end-to-end platform for high-plex tissue spatial omics analysis.' Packet metadata: paper_type=COMP, spatial=YES, panel_scope=UNCLEAR, gemma_q1_panel_design empty. | accepted_agree |
| 10.1093/gpbjnl/qzag006 | 2026 | FAST: Scalable Factor Analysis for Spatial Dimension Reduction of Multi-section Spatial Transcriptomics. | AGREE | Title: 'FAST: Scalable Factor Analysis for Spatial Dimension Reduction of Multi-section Spatial Transcriptomics.' Packet metadata: paper_type=COMP, spatial=YES, panel_scope=UNCLEAR, gemma_q1_panel_design empty. | accepted_agree |
| 10.1093/humrep/deaf097.300 | 2025 | O-300 Characterization of the transcriptional changes of human endometrium across the window of implantation during natural cycles reveals consistent dynamics | AGREE | Title: 'O-300 Characterization of the transcriptional changes of human endometrium across the window of implantation during natural cycles reveals consistent dynamics.' Packet metadata: Q0=NON_ARTICLE, gemma_final=FILTERED. | accepted_agree |

| DOI | Year | Title | Audit decision | Evidence (verbatim quote) | Validation status / scope flag |
|---|---|---|---|---|---|
| 10.1093/humrep/deaf097.304 | 2025 | O-304 The effects of placental invasion on the maternal cells in the decidua of early pregnancy | AGREE | gemma_q0_document_type.doc_type=NON_ARTICLE; doc_evidence=41st Annual Meeting of the European Society of Human Reproduction and Embryology | accepted_agree |
| 10.1093/ibd/izae020.151 | 2024 | VEDOLIZUMAB RESPONSE AND NON-RESPONSE IN ULCERATIVE COLITIS: INSIGHTS FROM SPATIAL TRANSCRIPTOMICS | AGREE | gemma_q0_document_type.doc_type=NON_ARTICLE; doc_evidence=Abstract citation ID: izae020.151 | accepted_agree |
| 10.1093/ibd/izae282.139 | 2025 | SPATIAL TRANSCRIPTOMICS REVEAL ACTIVATED T CELLS NEIGHBORING EPITHELIAL CELLS IN IMMUNE CHECKPOINT INHIBITOR ASSOCIATED COLITIS | AGREE | gemma_q0_document_type.doc_type=NON_ARTICLE; doc_evidence=(paraphrased) Crohn's & Colitis Congress; Abstract citation ID: izae282.139 | accepted_agree |
| 10.1093/ibd/izag006.095 | 2026 | DISTINCTIONS IN CD4+ MACROPHAGE BEHAVIOR SIGNIFY CROHN'S DISEASE WITH PERIANAL COMPLICATIONS | AGREE | gemma_q0_document_type.doc_type=NON_ARTICLE; doc_evidence=Abstract citation ID: izag006.093 | accepted_agree |
| 10.1093/ibd/izag006.112 | 2026 | TIME-RESOLVED CELL-CELL COMMUNICATION INFERENCE DURING ULCERATIVE COLITIS PROGRESSION | AGREE | Title: 'TIME-RESOLVED CELL-CELL COMMUNICATION INFERENCE DURING ULCERATIVE COLITIS PROGRESSION'; packet fields show Q0: NON_ARTICLE and Gemma's FILTERED prefilter. | accepted_agree |
| 10.1093/ibd/izag006.129 | 2026 | IDENTIFYING DISTINCT INFLAMMATORY FIBROBLAST STATES BY SPATIAL TRANSCRIPTOMICS TO PREDICT COLECTOMY IN PEDIATRIC ACUTE SEVERE COLITIS | AGREE | gemma_q0_document_type.doc_type=NON_ARTICLE; doc_evidence=Abstract citation ID: izag006.129 | accepted_agree |
| 10.1093/icb/icad007 | 2023 | A Fast And Versatile Method for Simultaneous HCR, Immunohistochemistry And Edu Labeling (SHInE). | AGREE | gemma_final.classification=FILTERED; gemma_q2_paper_profile.paper_type=OTHER; spatial=YES; panel_scope=UNCLEAR; doc_evidence=Pass-A abstract: OTHER -> FILTERED | accepted_agree |
| 10.1093/intbio/zyae020 | 2024 | Machine learning ranking of plausible (un)explored synergistic | AGREE | paper_type/profile evidence: COMP; citation-screening prior | accepted_agree |

| DOI | Year | Title | Audit decision | Evidence (verbatim quote) | Validation status / scope flag |
|---|---|---|---|---|---|
| | | gene combinations using sensitivity indices of time series measurements of Wnt signaling pathway | | labels=['COMP']; gemma_final notes: computational-only prior pre-filter | |
| 10.1093/jimmun/vkaf283.1103 | 2025 | A lymph node slice culture model to spatially profile early vaccine responses in human tissue 3287 | AGREE | gemma_q0_document_type.doc_type=NON_ARTICLE; doc_evidence=Abstract Description: | accepted_agree |
| 10.1093/jimmun/vkaf283.1463 | 2025 | Intra-tumoral bacterial signals are associated with distinct immune niches in metastatic brain tumors 3692 | AGREE | gemma_q0_document_type.doc_type=NON_ARTICLE; doc_evidence=Abstracts | accepted_agree |
| 10.1093/jimmun/vkaf283.1992 | 2025 | Multi-omic spatial profiling of nasal polyps identifies multiple immune-epithelial cell-ligand receptor interactions 4292 | AGREE | gemma_q0_document_type.doc_type=NON_ARTICLE; doc_evidence=Abstract citation ID: vkaf283.1992 | accepted_agree |
| 10.1093/jimmun/vkaf283.2557 | 2025 | A Deep Dive into the Functional Heterogeneity of Interstitial Macrophages 9108 | AGREE | gemma_q0_document_type.doc_type=NON_ARTICLE; doc_evidence=Abstract citation ID: vkaf283.2557 | accepted_agree |
| 10.1093/jimmun/vkaf283.2571 | 2025 | Identification and functional role of medulla germinal centers during SARS-CoV-2 infection and reinfection in a macaque model using Xenium in situ spatial transcriptomics 9127 | AGREE | gemma_q0_document_type.doc_type=NON_ARTICLE; doc_evidence=Abstract citation ID: vkaf283.2571 | accepted_agree |
| 10.1093/jimmun/vkaf283.2679 | 2025 | Tertiary lymphoid follicles in the human nasal mucosa are inductive sites for humoral immunity 9253 | AGREE | gemma_q0_document_type.doc_type=NON_ARTICLE; doc_evidence=Abstract citation ID: vkaf283.2679 | accepted_agree |
| 10.1093/jimmun/vkaf283.448 | 2025 | Spatial biology discovery approaches to dissect in situ modification of site-specific tissue HIV reservoirs 2542 | AGREE | gemma_q0_document_type.doc_type=NON_ARTICLE; doc_evidence=Abstract citation ID: vkaf283.449 | accepted_agree |
| 10.1093/jimmun/vkaf283.872 | 2025 | Defining molecular determinants of T cell spatial localization and clonal response throughout PDAC malignant progression 3030 | AGREE | gemma_q0_document_type.doc_type=NON_ARTICLE; doc_evidence=Abstract citation ID: vkaf283.872 | accepted_agree |
| 10.1093/nar/gkab630 | 2021 | Correction to 'Hybridization-based <i>in situ</i> sequencing (HybISS) for spatially resolved transcriptomics in human and mouse brain tissue' | AGREE | gemma_q0_document_type.doc_type=CORRECTION | accepted_agree |

| DOI | Year | Title | Audit decision | Evidence (verbatim quote) | Validation status / scope flag |
|---|---|---|---|---|---|
| 10.1093/nar/gkac922 | 2022 | ImmCluster: an ensemble resource for immunology cell type clustering and annotations in normal and cancerous tissues | AGREE | title='ImmCluster: an ensemble resource for immunology cell type clustering and annotations in normal and cancerous tissues'; citation_screening_prior.labels=''; gemma_final.source='abstract_prefilter' | accepted_agree |
| 10.1093/nar/gkad1023 | 2023 | Probabilistic cell/domain-type assignment of spatial transcriptomics data with SpatialAnno. | AGREE | gemma_final.classification=FILTERED; gemma_q2_paper_profile.paper_type=COMP; panel_scope=UNCLEAR; spatial=YES; profile_confidence=HIGH | accepted_agree |
| 10.1093/nar/gkad157 | 2023 | Multi-task learning from multimodal single-cell omics with Matilda | AGREE | title='Multi-task learning from multimodal single-cell omics with Matilda'; citation_screening_prior.labels='COMP'; gemma_final.source='citation_screening_prefilter' | accepted_agree |
| 10.1093/nar/gkad169 | 2023 | The spatial landscape of gene expression isoforms in tissue sections. | AGREE | gemma_q2_paper_profile.paper_type=COMP; panel_scope=NOT_TARGETED; spatial=YES; profile_confidence=HIGH | accepted_agree |
| 10.1093/nar/gkad337 | 2023 | vissE.cloud: a webserver to visualise higher order molecular phenotypes from enrichment analysis. | NEEDS_MANUAL | PDF: vissE.Cloud is a webserver for gene-set enrichment analysis and visualization. | pdf_validated_unclear_update |
| 10.1093/nar/gkad820 | 2023 | SORC: an integrated spatial omics resource in cancer | AGREE | title='SORC: an integrated spatial omics resource in cancer'; citation_screening_prior.labels=''; gemma_final.source='abstract_prefilter' | accepted_agree |
| 10.1093/nar/gkad895 | 2023 | SCAN: Spatiotemporal Cloud Atlas for Neural cells | AGREE | title='SCAN: Spatiotemporal Cloud Atlas for Neural cells'; citation_screening_prior.labels=''; gemma_final.source='abstract_prefilter' | accepted_agree |
| 10.1093/nar/gkae583 | 2024 | Imaging and quantification of human and viral circular RNAs. | NEEDS_MANUAL | PDF: ampFISH/smFISH probe pairs were designed for circRNAs; no gene-panel selection. | pdf_validated_unclear_update |

| DOI | Year | Title | Audit decision | Evidence (verbatim quote) | Validation status / scope flag |
|---|---|---|---|---|---|
| 10.1093/nar/gkaf1092 | 2025 | scBrainScope: cross-species multidimensional brain atlas | AGREE | title: scBrainScope: cross-species multidimensional brain atlas; q2 spatial=YES, panel_scope=UNCLEAR, paper_type=COMP; q1 panel-design fields are empty; gemma_final=FILTERED. | accepted_agree |
| 10.1093/nar/gkaf1456 | 2026 | SCALE: unsupervised multiscale domain identification in spatial omics data. | NEEDS_MANUAL | Final manual decision: FILTERED because the paper is a downstream analysis/modeling/QC method or dataset-use paper, with no final panel-design source. Original evidence: q1: panel_design=UNCLEAR; q2: targeted MERFISH/Xenium context only. | manual_final_filter_after_pdf_review |
| 10.1093/nar/gkaf1477 | 2026 | Microenvironment-aware spatial modeling for accurate inference of cell identity. | NEEDS_MANUAL | PDF: computational model for cell-identity inference from spatial patterns; no panel design. | pdf_validated_unclear_update |
| 10.1093/nar/gkx1206 | 2018 | Efficient in situ barcode sequencing using padlock probe-based BaristaSeq. | NEEDS_MANUAL | PDF: a response about seqFISH barcode data; no new panel selection in this paper. | pdf_validated_unclear_update |
| 10.1093/narcan/zcac038 | 2022 | Single-cell CRISPR immune screens reveal immunological roles of tumor intrinsic factors | AGREE | title='Single-cell CRISPR immune screens reveal immunological roles of tumor intrinsic factors'; citation_screening_prior.labels='NON-ST'; gemma_final.source='citation_screening_prefilter' | accepted_agree |
| 10.1093/nargab/lqad070 | 2023 | Single-cell reference mapping to construct and extend cell-type hierarchies | AGREE | title='Single-cell reference mapping to construct and extend cell-type hierarchies'; citation_screening_prior.labels='COMP-BENCH'; gemma_final.source='citation_screening_prefilter' | accepted_agree |
| 10.1093/nargab/lqae100 | 2024 | SIRV: spatial inference of RNA velocity at the single-cell resolution. | AGREE | Pass-A abstract: COMP → FILTERED | accepted_agree |
| 10.1093/nargab/lqaf114 | 2024 | PRISM: a Python package for interactive and integrated analysis of multiplexed tissue microarrays | NEEDS_MANUAL | PDF: PRISM is a Python package for TMA masking, segmentation, clustering, and spatial analysis. | pdf_validated_unclear_update |
| 10.1093/nargab/lqag039 | 2025 | SpNeigh: spatial neighborhood and differential expression analysis | AGREE | Pass-A abstract: COMP → FILTERED | accepted_agree |

| DOI | Year | Title | Audit decision | Evidence (verbatim quote) | Validation status / scope flag |
|---|---|---|---|---|---|
| | | for high-resolution spatial transcriptomics | | | |
| 10.1093/ndt/gfad063b_6014 | 2023 | #6014 SINGLE-CELL SPATIAL TRANSCRIPTOMICS OF THE KIDNEY IN HEALTH AND DISEASE DEFINES INJURY-SPECIFIC DOMAINS AND IDENTIFIES NOVEL CELL-CELL INTERACTIONS | AGREE | Q0: NON_ARTICLE → FILTERED | accepted_agree |
| 10.1093/ndt/gfaf116.0968 | 2025 | #271 Tubular epithelial cells are the primary source of Flt3L in acute kidney injury, independent of immune cells | AGREE | Q0: NON_ARTICLE → FILTERED | accepted_agree |
| 10.1093/neuonc/noac174.100 | 2022 | P02.07.B Patient-derived glioblastoma organoids: Elucidating the mechanisms of glioblastoma therapeutic resistance in the context of tumor microenvironment | AGREE | Q0: NON_ARTICLE → FILTERED | accepted_agree |
| 10.1093/neuonc/noac209.1113 | 2022 | TMIC-70. HARMONIZED SINGLE-CELL LANDSCAPE, INTERCELLULAR CROSSTALK AND TUMOR ARCHITECTURE OF GLIOBLASTOMA | AGREE | Q0: NON_ARTICLE → FILTERED | accepted_agree |
| 10.1093/neuonc/noac209.456 | 2022 | EPCO-21. THE SPATIAL ORGANIZATION OF H3-K27M MUTANT DIFFUSE MIDLINE GLIOMA | AGREE | Q0: NON_ARTICLE → FILTERED | accepted_agree |
| 10.1093/neuonc/noad179.1122 | 2023 | TMIC-56. DECIPHERING THE INTRICATE GLIOBLASTOMA TUMOR MICROENVIRONMENT AND EGFRVIII TRANSCRIPT DISTRIBUTION THROUGH SINGLE-CELL SPATIAL PROFILING | AGREE | Q0: NON_ARTICLE → FILTERED | accepted_agree |
| 10.1093/neuonc/noad179.1123 | 2023 | TMIC-57. CHARTING THE SPATIAL ARCHITECTURE OF GLIOBLASTOMA USING GBMAP: INSIGHTS FROM IN SITU SEQUENCING | AGREE | Q0: NON_ARTICLE → FILTERED | accepted_agree |
| 10.1093/neuonc/noad179.1148 | 2023 | TMIC-83. THE MOLECULAR AND SPATIAL ARCHITECTURE OF AN IMMUNE-RICH GLIOMA SUBTYPE | AGREE | Q0: NON_ARTICLE → FILTERED | accepted_agree |

| DOI | Year | Title | Audit decision | Evidence (verbatim quote) | Validation status / scope flag |
|---|---|---|---|---|---|
| 10.1093/neuonc/noae144.091 | 2024 | P01.03.A DECIPHERING THE TUMOUR ECOSYSTEM IN GLIOMAS USING SPATIAL TRANSCRIPTOMICS | AGREE | Q0: NON_ARTICLE → FILTERED | accepted_agree |
| 10.1093/neuonc/noae165.0043 | 2024 | EPCO-44. SPATIAL TRANSCRIPTOMICS ENABLES STATISTICAL MODELING OF SEX DIFFERENCE IN GENE CO-EXPRESSION NETWORK AND IMMUNE LANDSCAPE OF GLIOBLASTOMA | AGREE | Q0: NON_ARTICLE → FILTERED | accepted_agree |
| 10.1093/neuonc/noae165.0054 | 2024 | EPCO-55. SPATIAL HETEROGENEITY IN THE TUMOR CORE AND INFILTRATING MARGIN OF<i>IDH</i>WT GLIOBLASTOMA | AGREE | Pass-A abstract: not targeted → FILTERED | accepted_agree |
| 10.1093/neuonc/noae165.0155 | 2024 | BIOS-07. GRADUAL METABOLIC ALTERATIONS RESTRUCTURE THE SPATIAL ARCHITECTURE AND IMMUNITY IN GLIOBLASTOMA | AGREE | Q0: NON_ARTICLE → FILTERED | accepted_agree |
| 10.1093/neuonc/noae165.1003 | 2024 | EXTH-73. TARGETING THE CELL ADHESION CLUSTER OF DIFFERENTIATION 44 TRANSMEMBRANE GLYCOPROTEIN SENSITIZES CRANIOPHARYNGIOMAS TO IMMUNOTHERAPY | AGREE | Q0: NON_ARTICLE → FILTERED | accepted_agree |
| 10.1093/neuonc/noaf085 | 2025 | SOX2 commands LIM homeobox transcription factors in choroid plexus development and tumorigenesis | NEEDS_MANUAL | Final manual decision: FILTERED because manual review found no usable evidence for final gene-panel source selection. Original evidence: q1 panel_design: NOT FOUND; q2 spatial: YES. | manual_final_filter_after_pdf_review |
| 10.1093/neuonc/noaf179 | 2025 | A cycling, progenitor-like cell population at the base of atypical teratoid rhabdoid tumor subtype differentiation trajectories | AGREE | title: A cycling, progenitor-like cell population at the base of atypical teratoid rhabdoid tumor subtype differentiation trajectories; q2 paper_type=EXP, spatial=NO, panel_scope=UNCLEAR; q1 panel-design fields are empty. | accepted_agree |
| 10.1093/neuonc/noaf193.025 | 2025 | KS03.4.A SPATIAL MULTI-OMIC PROFILING REVEALS DETERMINANTS OF TIL | AGREE | Q0: NON_ARTICLE → FILTERED | accepted_agree |

| DOI | Year | Title | Audit decision | Evidence (verbatim quote) | Validation status / scope flag |
|---|---|---|---|---|---|
| | | EXPANDABILITY IN HIGH-GRADE GLIOMAS | | | |
| 10.1093/neuonc/noaf193.027 | 2025 | KS03.6.A THE RECURRENT TISSUE STATES ACROSS ADULT GLIOMAS AS REVEALED BY SPATIAL TRANSCRIPTOMICS | AGREE | Q0: NON_ARTICLE → FILTERED | accepted_agree |
| 10.1093/neuonc/noaf193.052 | 2025 | OS05.4.A THE GLIOBLASTOMA HYPOXIC NICHE REWIRES LIPID-LADEN MACROPHAGE PHENOTYPES IN AN ONTOGENY DEPENDENT MANNER | AGREE | Pass-A abstract: not targeted → FILTERED | accepted_agree |
| 10.1093/neuonc/noaf193.135 | 2025 | P02.08.A A SPATIALLY RESOLVED HUMAN GLIOBLASTOMA ATLAS REVEALS DISTINCT CELLULAR AND MOLECULAR PATTERNS OF ANATOMICAL NICHES | AGREE | Q0: NON_ARTICLE → FILTERED | accepted_agree |
| 10.1093/neuonc/noaf193.142 | 2025 | P02.15.B GLIOBLASTOMA IN THE CONNECTOME: NEURAL PHENOTYPES PREDICT TUMOR INTEGRATION AND CONNECTIVITY | AGREE | Q0: NON_ARTICLE → FILTERED | accepted_agree |
| 10.1093/neuonc/noaf193.191 | 2025 | P04.05.B DEEP LEARNING ANALYSIS OF SPATIALLY RESOLVED SINGLE-CELL TRANSCRIPTOMICS UNCOVERS CELLULAR REORGANIZATION IN GLIOBLASTOMA AT RECURRENCE | AGREE | q0 says NON_ARTICLE; q1 panel-design fields are empty; q2 profile is empty. | accepted_agree |
| 10.1093/neuonc/noaf193.202 | 2025 | P04.16.A DECIPHERING NEURAL-IMMUNE RECIPROCAL INTERACTIONS IN GLIOBLASTOMA USING 3D SINGEL-CELL SPATIAL TRANSCRIPTOMICS | AGREE | q0 says NON_ARTICLE; q1 panel-design fields are empty; q2 profile is empty. | accepted_agree |
| 10.1093/neuonc/noaf201.0016 | 2025 | EPCO-16. CHROMATIN REMODELING WITH COMBINED FACT AND BET INHIBITION DISRUPTS ONCOGENIC TRANSCRIPTION IN DIFFUSE MIDLINE GLIOMA | AGREE | q0 says NON_ARTICLE; q1 panel-design fields are empty; q2 profile is empty. | accepted_agree |
| 10.1093/neuonc/noaf201.0024 | 2025 | EPCO-24. DISSECTING THE ECOSYSTEM OF PEDIATRIC | AGREE | q0 says NON_ARTICLE; q1 panel-design fields are empty; q2 profile is empty. | accepted_agree |

| DOI | Year | Title | Audit decision | Evidence (verbatim quote) | Validation status / scope flag |
|---|---|---|---|---|---|
| | | H3K27 ALTERED DIFFUSE MIDLINE GLIOMA | | | |
| 10.1093/neuonc/noaf201.0053 | 2025 | EPCO-54. DEFINING THE MULTI-SCALE ORGANIZATION AND REGULATION OF EGFR EXTRACHROMOSOMAL DNA IN GLIOBLASTOMA WITH MULTI-MODAL MERFISH | AGREE | q0 says NON_ARTICLE; q1 panel-design fields are empty; q2 profile is empty. | accepted_agree |
| 10.1093/neuonc/noaf201.0113 | 2025 | BIOM-25. GRAMMA-GLUTAMYL-HYDROLASE (GGH) AS A TARGETABLE DRIVER IN GLIOBLASTOMA: A COMPREHENSIVE OMICS-DRIVEN BIOINFORMATICS-TO-BENCH PIPELINE INTEGRATING CRISPR, ANIMAL MODELS, AND ADVANCED SPATIAL/MULTIPLEX/3D MOLECULAR IMAGING | AGREE | q0 says NON_ARTICLE; q1 panel-design fields are empty; q2 profile is empty. | accepted_agree |
| 10.1093/neuonc/noaf201.0160 | 2025 | BIOM-72. INTEGRATIVE SINGLE-NUCLEI SPATIAL TRANSCRIPTOMIC PROFILING OF PRIMARY AND RECURRENT HUMAN IDH-WILDTYPE GLIOBLASTOMA REVEALS CELL-SPECIFIC EGFRVIII EXPRESSION AND ANXA1 SIGNALING AXIS | AGREE | q0 says NON_ARTICLE; q1 panel-design fields are empty; q2 profile is empty. | accepted_agree |
| 10.1093/neuonc/noaf201.0225 | 2025 | CNSC-17. RAPID RAMAN DISCRIMINATION OF GLIOBLASTOMA EDEMA, INFILTRATION, AND ENHANCEMENT VIA SINGLE-CELL AND SPATIAL OMICS | AGREE | q0 says NON_ARTICLE; q1 panel-design fields are empty; q2 profile is empty. | accepted_agree |
| 10.1093/neuonc/noaf201.0288 | 2025 | CNSC-81. H3G34-MUTANT DIFFUSE HEMISPHERIC GLIOMAS CONTAIN CELLULAR NICHES OF GABAERGIC INTERNEURON-LIKE CELLS WITH DUAL FREQUENCY AUTONOMOUS RHYTHMIC CALCIUM ACTIVITY | AGREE | q0 says NON_ARTICLE; q1 panel-design fields are empty; q2 profile is empty. | accepted_agree |
| 10.1093/neuonc/noaf201.0326 | 2025 | STEM-18. GENETIC LINEAGE TRACING OF MEDULLOBLASTOMA UNDER | AGREE | q0 says NON_ARTICLE; q1 panel-design fields are empty; q2 profile is empty. | accepted_agree |

| DOI | Year | Title | Audit decision | Evidence (verbatim quote) | Validation status / scope flag |
|---|---|---|---|---|---|
| | | THERAPY WITH SPATIAL CONTEXT | | | |
| 10.1093/neuonc/noaf201.0865 | 2025 | IMMU-67. On-Treatment Spatiotemporal Profiling of Glioblastoma During Oncolytic ImmunoTherapy | AGREE | q0 says NON_ARTICLE; q1 panel-design fields are empty; q2 profile is empty. | accepted_agree |
| 10.1093/neuonc/noaf201.1003 | 2025 | PATH-51. Identification of molecular markers predicting malignant transformation in low grade gliomas | AGREE | q0 says NON_ARTICLE; q1 panel-design fields are empty; q2 profile is empty. | accepted_agree |
| 10.1093/neuonc/noaf201.1815 | 2025 | TMIC-60. Spatial profiling of IDH-wildtype glioblastoma using CosMx Spatial Molecular Imaging technology to establish a large-scale FFPE tissue cohort | AGREE | q0 says NON_ARTICLE; q1 panel-design fields are empty; q2 profile is empty. | accepted_agree |
| 10.1093/neuonc/noaf201.1841 | 2025 | TMIC-86. Meningioma brain-invasion is linked to inflammatory induced tumor-neuronal connectivity | AGREE | q0 says NON_ARTICLE; q1 panel-design fields are empty; q2 profile is empty. | accepted_agree |
| 10.1093/neuonc/noaf201.1846 | 2025 | TMIC-91. Deciphering spatio-temporal trajectories of tissue adaptation after surgical resection of glioblastoma | AGREE | q0 says NON_ARTICLE; q1 panel-design fields are empty; q2 profile is empty. | accepted_agree |
| 10.1093/noajnl/vdaf193 | 2025 | Targeting the tumor microenvironment in pediatric gliomas: Advances and future directions in immunotherapy | AGREE | doc_type=REVIEW; title: Targeting the tumor microenvironment in pediatric gliomas: Advances and future directions in immunotherapy; q1 panel-design fields are empty. | accepted_agree |
| 10.1093/noajnl/vdaf236.007 | 2025 | 10189-CB-6 Rapid Raman Discrimination of Glioblastoma Edema, Infiltration, and Enhancement via Single-Cell and Spatial Omics | AGREE | q0 says NON_ARTICLE; q1 panel-design fields are empty; q2 profile is empty. | accepted_agree |
| 10.1093/nsr/nwaf030 | 2025 | DSCT: a novel deep-learning framework for rapid and accurate spatial transcriptomic cell typing | AGREE | title: DSCT: a novel deep-learning framework for rapid and accurate spatial transcriptomic cell typing; citation prior labels=COMP-BENCH; q1 panel-design fields are empty; gemma_final=FILTERED. | accepted_agree |
| 10.1093/oncolo/oyaf276.053 | 2025 | 52Spatial transcriptomic mapping reveals immune-inflamed niches in sarcomatoid chromophobe renal cell carcinoma | AGREE | q0 says NON_ARTICLE; q1 panel-design fields are empty; q2 profile is empty. | accepted_agree |

| DOI | Year | Title | Audit decision | Evidence (verbatim quote) | Validation status / scope flag |
|---|---|---|---|---|---|
| 10.1096/fasebj.2019.33.1_supplement.221.2 | 2019 | Dynamics and Spatial Genomics of the Nascent Transcriptome by Intron seqFISH | NEEDS_MANUAL | Final manual decision: FILTERED because no local PDF was available and the packet did not provide usable final gene-panel source evidence. Original evidence: To multiplex TAS detection for 10,421 genes, we use sequential rounds of hybridization to generate a unique temporal barcode sequence on each gene, which we then decode by aligning images from multiple round of barcoding hybridization. | manual_final_filter_after_pdf_review |
| 10.1097/01.ju.0001009560.23593.56.02 | 2024 | PD16-02 INCREASED SPATIAL COUPLING OF INTEGRIN AND COLLAGEN IV IN THE IMMUNORESISTANT CLEAR CELL RENAL CELL CARCINOMA TUMOR MICROENVIRONMENT | AGREE | q0 says NON_ARTICLE; q1 panel-design fields are empty; q2 profile is empty. | accepted_agree |
| 10.1097/01.ju.0001009560.23593.56.04 | 2024 | PD16-04 SPATIAL ANALYSIS OF PRIMARY AND METASTATIC LESIONS IN SARCOMATOID RENAL CELL CARCINOMA REVEALS INSIGHT TO TUMOR BIOLOGY AND IMMUNE MICROENVIRONMENT | AGREE | q0 says NON_ARTICLE; q1 panel-design fields are empty; q2 profile is empty. | accepted_agree |
| 10.1097/01.ju.0001009560.23593.56.05 | 2024 | PD16-05 EPITHELIAL TO MESENCHYMAL TRANSITION (EMT) AND PD-1/PD-L1 SIGNALING PROVIDE RECIPROCAL FEEDBACK IN RENAL CELL CARCINOMA PROGRESSION WITH IMPLICATIONS FOR IMMUNOTHERAPY RESPONSE | AGREE | q0 says NON_ARTICLE; q1 panel-design fields are empty; q2 profile is empty. | accepted_agree |
| 10.1097/01.tp.0001066488.16290.8e | 2024 | 442.1: Spatial transcriptomics of kidney biopsies following adoptive regulatory T cell transfer reveals a marginal zone B cell signature and novel intra-graft immune interactions. | AGREE | title: 442.1: Spatial transcriptomics of kidney biopsies following adoptive regulatory T cell transfer reveals a marginal zone B cell signature and novel intra-graft immune interactions.; final notes: Q0: NON_ARTICLE → FILTERED; packet flags: non_primary_document | accepted_agree |
| 10.1097/01.tp.0001066624.94591.08 | 2024 | 447.7: Multi-omic profiling in a human decedent receiving a pig | AGREE | title: 447.7: Multi-omic profiling in a human decedent receiving a pig | accepted_agree |

| DOI | Year | Title | Audit decision | Evidence (verbatim quote) | Validation status / scope flag |
| --- | --- | --- | --- | --- | --- |
| | | kidney xenograft reveals concerted immune responses preceding a biopsy confirmed rejection event. | | kidney xenograft reveals concerted immune responses preceding a biopsy confirmed rejection event.; final notes: Q0: NON_ARTICLE → FILTERED; packet flags: non_primary_document | |
| 10.1097/cd9.0000000000000027 | 2021 | Single-Cell Sequencing of the Cardiovascular System: Challenges in Translation | AGREE | title: Single-Cell Sequencing of the Cardiovascular System: Challenges in Translation; final notes: Pass-A abstract: OTHER → FILTERED; packet flags: prefiltered_without_full_panel_qs | accepted_agree |
| 10.1097/hc9.0000000000000813 | 2025 | Single-cell atlas of human pediatric liver reveals age-related hepatic gene signatures | AGREE | title: Single-cell atlas of human pediatric liver reveals age-related hepatic gene signatures; final notes: Pass-A abstract: non-spatial → FILTERED; packet flags: prefiltered_without_full_panel_qs | accepted_agree |
| 10.1097/hep.0000000000001360 | 2026 | AI-based phenotyping of hepatic fiber morphology to inform molecular alterations in metabolic dysfunction-associated steatotic liver disease. | AGREE | paper_type/doc_type evidence: EDITORIAL (JOINT GI SOCIETY STATEMENT); gemma_final notes: Q0: EDITORIAL → FILTERED | accepted_agree |
| 10.1097/hs9.0000000000000193 | 2019 | Efficacy of Tyrosine Kinase Inhibitor Therapy in a Chemotherapy-refractory B-cell Precursor Acute Lymphoblastic Leukemia With <i>ZC3HAV1-ABL2</i> Fusion | AGREE | title: Efficacy of Tyrosine Kinase Inhibitor Therapy in a Chemotherapy-refractory B-cell Precursor Acute Lymphoblastic Leukemia With <i>ZC3HAV1-ABL2</i> Fusion; final notes: Pass-A abstract: non-spatial → FILTERED; packet flags: prefiltered_without_full_panel_qs | accepted_agree |
| 10.1097/ju.0000000000003272.12 | 2023 | PD17-12 SPATIAL MOLECULAR IMAGING IN SARCOMATOID RENAL CELL CARCINOMA REVEALS CROSSTALK BETWEEN IMMUNE CELLS AND TUMOR CELLS | AGREE | title: PD17-12 SPATIAL MOLECULAR IMAGING IN SARCOMATOID RENAL CELL CARCINOMA REVEALS CROSSTALK BETWEEN IMMUNE CELLS AND TUMOR CELLS; final notes: Q0: NON_ARTICLE → FILTERED; packet flags: non_primary_document | accepted_agree |
| 10.1097/mat.0000000000000255 | 2015 | Intensive hemodialysis preserved cardiac injury | AGREE | title: Intensive hemodialysis preserved cardiac injury; final notes: Pass-A abstract: non-spatial → FILTERED; packet flags: prefiltered_without_full_panel_qs | accepted_agree |

| DOI | Year | Title | Audit decision | Evidence (verbatim quote) | Validation status / scope flag |
|---|---|---|---|---|---|
| 10.1097/pas.0000000000001408 | 202 | Clinicopathologic and Molecular Analysis of the TFEB Fusion Variant Reveals New Members of TFEB Translocation Renal Cell Carcinomas (RCCs): Expanding the Genomic Spectrum. | AGREE | title: Clinicopathologic and Molecular Analysis of the TFEB Fusion Variant Reveals New Members of TFEB Translocation Renal Cell Carcinomas (RCCs): Expanding the Genomic Spectrum.; final notes: Pass-A abstract: non-spatial → FILTERED; packet flags: prefiltered_without_full_panel_qs | accepted_agree |
| 10.1097/tp.0000000000005592 | 2025 | Blind Spots in HLA Class II Detection Generate Challenges for Antibody and RNA-based Assays | AGREE | Q0 document_type: REVIEW; no panel-design evidence is present. | accepted_agree |
| 10.1101/143321 | 2017 | SpatialDE - Identification of spatially variable genes | AGREE | title: SpatialDE - Identification of spatially variable genes; final notes: Pass-A abstract: COMP → FILTERED; packet flags: prefiltered_without_full_panel_qs | accepted_agree |
| 10.1101/180323 | 2017 | Efficient<i>in situ</i>barcode sequencing using padlock probe-based BaristaSeq | NEEDS_MANUAL | PDF: BaristaSeq is a padlock-probe barcode sequencing method; no panel selection source. | pdf_validated_unclear_update |
| 10.1101/2020.01.14.906313 | 202 | A Bayesian nonparametric semi-supervised model for integration of multiple single-cell experiments | AGREE | Data from seqFISH+ consists of cell specific counts of 10,000 genes and cell centroid coordinates for each cell. | accepted_agree |
| 10.1101/2020.02.12.945345 | 202 | Spage2vec: Unsupervised detection of spatial gene expression constellations | AGREE | title: Spage2vec: Unsupervised detection of spatial gene expression constellations; final notes: Pass-A abstract: COMP → FILTERED; packet flags: prefiltered_without_full_panel_qs | accepted_agree |
| 10.1101/2020.02.13.948521 | 202 | Reconstruction of cell spatial organization based on ligand-receptor mediated self-assembly | AGREE | title: Reconstruction of cell spatial organization based on ligand-receptor mediated self-assembly; final notes: Pass-A abstract: COMP → FILTERED; packet flags: prefiltered_without_full_panel_qs | accepted_agree |
| 10.1101/2020.07.14.201384 | 202 | Validation and tuning of<i>in situ</i>transcriptomics image processing workflows with crowdsourced annotations | AGREE | title: Validation and tuning of<i>in situ</i>transcriptomics image processing workflows with crowdsourced annotations; final notes: Pass-A abstract: COMP → FILTERED; packet flags: prefiltered_without_full_panel_qs | accepted_agree |

| DOI | Year | Title | Audit decision | Evidence (verbatim quote) | Validation status / scope flag |
|---|---|---|---|---|---|
| 10.1101/2020.08.29.272831 | 202 | Deep learning and alignment of spatially-resolved whole transcriptomes of single cells in the mouse brain with Tangram | AGREE | title: Deep learning and alignment of spatially-resolved whole transcriptomes of single cells in the mouse brain with Tangram; final notes: Pass-A abstract: COMP → FILTERED; packet flags: prefiltered_without_full_panel_qs | accepted_agree |
| 10.1101/2020.09.09.290064 | 202 | Integrative Analysis of Spatial and Single-Cell Transcriptomics Reveals Principles of Tissue Organization and Intercellular Communication in Mouse Olfactory Bulb | AGREE | We apply STANN to assign cell-types in a recent ST dataset (SeqFISH+) of mouse olfactory bulb (MOB). | accepted_agree |
| 10.1101/2020.10.05.326777 | 202 | Bayesian segmentation of spatially resolved transcriptomics data | AGREE | title: Bayesian segmentation of spatially resolved transcriptomics data; final notes: Pass-A abstract: COMP → FILTERED; packet flags: prefiltered_without_full_panel_qs | accepted_agree |
| 10.1101/2020.11.17.387795 | 202 | scDesign2: a transparent simulator that generates high-fidelity single-cell gene expression count data with gene correlations captured | NEEDS_MANUAL | scDesign2 generates synthetic MERFISH/pciSeq data; it is a simulator, not a panel-selection method. | pdf_validated_unclear_update |
| 10.1101/2020.11.29.383067 | 202 | SPICEMIX: Integrative single-cell spatial modeling of cell identity | AGREE | title: SPICEMIX: Integrative single-cell spatial modeling of cell identity; final notes: Pass-A abstract: COMP → FILTERED; packet flags: prefiltered_without_full_panel_qs | accepted_agree |
| 10.1101/2021.03.01.433040 | 2021 | ISTDECO: In Situ Transcriptomics Decoding by Deconvolution | AGREE | title: ISTDECO: In Situ Transcriptomics Decoding by Deconvolution; final notes: Pass-A abstract: COMP → FILTERED; packet flags: prefiltered_without_full_panel_qs | accepted_agree |
| 10.1101/2021.04.08.438978 | 2021 | <tt>SMaSH:</tt> A scalable, general marker gene identification framework for single-cell RNA sequencing and Spatial Transcriptomics | AGREE | citation_screening_prior: computational-only prior; gemma_final source=citation_screening_prefilter -> FILTERED; q1 panel-design fields are empty; q2 paper_type=COMP. | accepted_agree |
| 10.1101/2021.06.15.448478 | 2021 | Active feature selection discovers minimal gene sets for classifying cell types and disease states with single-cell mRNA-seq data | AGREE | Title: 'Active feature selection discovers minimal gene sets for classifying cell types and disease states with single-cell mRNA-seq | accepted_agree |

| DOI | Year | Title | Audit decision | Evidence (verbatim quote) | Validation status / scope flag |
|---|---|---|---|---|---|
| | | | | data'; packet fields show paper_type=COMP and Gemma's FILTERED prefilter. | |
| 10.1101/2021.07.11.451750 | 2021 | Learning cell communication from spatial graphs of cells | NEEDS_MANUAL | Cell-communication inference on MERFISH brain data; no evidence of choosing or modifying the gene panel. | pdf_validated_unclear_update |
| 10.1101/2021.10.12.464086 | 2021 | PoSTcode: Probabilistic image-based spatial transcriptomics decoder | AGREE | title: PoSTcode: Probabilistic image-based spatial transcriptomics decoder; final notes: Pass-A abstract: COMP → FILTERED; packet flags: prefiltered_without_full_panel_qs | accepted_agree |
| 10.1101/2021.10.20.465151 | 2021 | Immune, endothelial and neuronal network map in human lymph node and spleen | AGREE | Title: 'Immune, endothelial and neuronal network map in human lymph node and spleen'; packet fields show paper_type=EXP, spatial=NO, and Gemma's FILTERED prefilter. | accepted_agree |
| 10.1101/2022.03.14.484288 | 2022 | Dual spatially resolved transcriptomics for SARS-CoV-2 host-pathogen colocalization studies in humans | AGREE | title: Dual spatially resolved transcriptomics for SARS-CoV-2 host-pathogen colocalization studies in humans; final notes: Pass-A abstract: not targeted → FILTERED; packet flags: prefiltered_without_full_panel_qs | accepted_agree |
| 10.1101/2022.03.28.486139 | 2022 | Reference-based cell type matching of spatial transcriptomics data | NEEDS_MANUAL | Compares cell-type matching across MERFISH/smFISH/BaristaSeq/ExSeq; mentions protocols have different probe panels, but does not select one. | pdf_validated_unclear_update |
| 10.1101/2022.04.14.488419 | 2022 | Supervised spatial inference of dissociated single-cell data with SageNet | AGREE | title: Supervised spatial inference of dissociated single-cell data with SageNet; final notes: Pass-A abstract: COMP → FILTERED; packet flags: prefiltered_without_full_panel_qs | accepted_agree |
| 10.1101/2022.05.06.490859 | 2022 | Depth normalization for single-cell genomics count data | AGREE | title='Depth normalization for single-cell genomics count data'; citation_screening_prior.labels='COMP'; gemma_final.source='citation_screening_prefilter' | accepted_agree |

| DOI | Year | Title | Audit decision | Evidence (verbatim quote) | Validation status / scope flag |
|---|---|---|---|---|---|
| 10.1101/2022.06.27.497696 | 2022 | Modeling and inference of spatial intercellular communications and multilayer signaling regulations using stMLnet | AGREE | title: Modeling and inference of spatial intercellular communications and multilayer signaling regulations using stMLnet; final notes: Pass-A abstract: COMP → FILTERED; packet flags: prefiltered_without_full_panel_qs | accepted_agree |
| 10.1101/2022.07.18.500480 | 2022 | Predicting the Reprogrammability of Human Cells Based on Transcriptome Data and SGD Classifier with Elastic-Net Regularization | AGREE | title='Predicting the Reprogrammability of Human Cells Based on Transcriptome Data and SGD Classifier with Elastic-Net Regularization'; citation_screening_prior.labels='COMP'; gemma_final.source='citation_screening_prefilter' | accepted_agree |
| 10.1101/2022.08.18.504464 | 2022 | SiGra: Single-cell spatial elucidation through image-augmented graph transformer | AGREE | Title: 'SiGra: Single-cell spatial elucidation through image-augmented graph transformer'; gemma_final=FILTERED; gemma_q1 empty; gemma_q2 says spatial=YES, paper_type=COMP. | accepted_agree |
| 10.1101/2022.10.21.513201 | 2022 | MorphNet Predicts Cell Morphology from Single-Cell Gene Expression | NEEDS_MANUAL | MorphNet is trained on MERFISH gene expression to generate morphology; that is downstream modeling, not panel design. | pdf_validated_unclear_update |
| 10.1101/2022.10.24.513598 | 2022 | Spatially resolved human kidney multi-omics single cell atlas highlights the key role of the fibrotic microenvironment in kidney disease progression | AGREE | Title: 'Spatially resolved human kidney multi-omics single cell atlas...'; gemma_final=FILTERED; gemma_q1 empty; gemma_q2 says spatial=YES, paper_type=EXP, panel_scope=NOT_TARGETED/UNCLEAR. | accepted_agree |
| 10.1101/2022.10.26.513573 | 2022 | The BRAIN Initiative Cell Census Network Data Ecosystem: A User’s Guide | AGREE | title='The BRAIN Initiative Cell Census Network Data Ecosystem: A User’s Guide'; citation_screening_prior.labels=''; gemma_final.source='abstract_prefilter' | accepted_agree |
| 10.1101/2022.11.04.515095 | 2022 | Integrated transcriptome and lineage analyses reveal novel catecholaminergic cardiomyocytes | DISAGREE | Stereo-seq is a whole-transcript spatial method and the paper profile marks panel_scope as NOT_TARGETED, so SMaSH | accepted_disagree |

| DOI | Year | Title | Audit decision | Evidence (verbatim quote) | Validation status / scope flag |
|---|---|---|---|---|---|
| | | contributing to the cardiac conduction system in murine heart | | marker identification here is not a targeted panel-design event. | |
| 10.1101/2023.01.12.523851 | 2023 | Integration of spatial and single-cell data across modalities with weak linkage | AGREE | Title: 'Integration of spatial and single-cell data across modalities with weak linkage'; gemma_final=FILTERED; gemma_q1 empty; gemma_q2 says spatial=YES, paper_type=COMP. | accepted_agree |
| 10.1101/2023.01.18.524520 | 2023 | A 3-dimensional molecular cartography of human cerebral organoids revealed by double-barcoded spatial transcriptomics | AGREE | panel_scope: NOT_TARGETED; evidence_scope: interrogate gene expression in an unbiased manner; evidence_classification: NOT FOUND | accepted_agree |
| 10.1101/2023.01.30.526264 | 2023 | Programmable peroxidase-assisted signal amplification enables flexible detection of nucleic acid targets in cellular and histopathological specimens | AGREE | Title: 'Programmable peroxidase-assisted signal amplification...'; gemma_final=FILTERED; gemma_q1 empty; gemma_q2 says spatial=YES, paper_type=COMP. | accepted_agree |
| 10.1101/2023.02.18.528138 | 2023 | Conformational analysis of chromosome structures reveals vital role of chromosome morphology in gene function | NEEDS_MANUAL | Chromosome-structure analysis validated with DNA-MERFISH data; no gene-panel selection evidence. | pdf_validated_unclear_update |
| 10.1101/2023.03.13.532376 | 2023 | InClust+: the multimodal version of inClust for multimodal data integration, imputation, and cross modal generation | NEEDS_MANUAL | Final manual decision: FILTERED because the paper is a downstream analysis/modeling/QC method or dataset-use paper, with no final panel-design source. Original evidence: q1: panel_design=UNCLEAR; q2: MERFISH with 153 common genes, but no selection source. | manual_final_filter_after_pdf_review |
| 10.1101/2023.03.16.532947 | 2023 | <i>mosna</i> reveals different types of cellular interactions predictive of response to immunotherapies and survival in cancer | AGREE | Title: 'mosna reveals different types of cellular interactions...'; gemma_final=FILTERED; gemma_q1 empty; gemma_q2 says spatial=YES, paper_type=COMP. | accepted_agree |

| DOI | Year | Title | Audit decision | Evidence (verbatim quote) | Validation status / scope flag |
|---|---|---|---|---|---|
| 10.1101/2023.03.27.534291 | 2023 | i-stLearn: An interactive platform for spatial transcriptomics analysis | AGREE | Title: 'i-stLearn: An interactive platform for spatial transcriptomics analysis'; gemma_final=FILTERED; gemma_q1 empty; gemma_q2 says spatial=YES, paper_type=COMP. | accepted_agree |
| 10.1101/2023.03.27.534477 | — | Improved rescue of immature oocytes obtained from conventional gonadotropin stimulation cycles via human induced pluripotent stem cell-derived ovarian support cell co-culture | AGREE | Title: 'Improved rescue of immature oocytes obtained from conventional gonadotropin stimulation cycles via human induced pluripotent stem cell-derived ovarian support cell co-culture'; packet fields show labels=NON-ST and Gemma's FILTERED prefilter. | accepted_agree |
| 10.1101/2023.03.28.534622 | 2023 | A Universal Method for Crossing Molecular and Atlas Modalities using Simplex-Based Image Varifolds and Quadratic Programming | NEEDS_MANUAL | Atlas-to-MERFISH mapping paper; text discusses "MERFISH spatial transcriptomics" and image registration, not panel construction. | pdf_validated_unclear_update |
| 10.1101/2023.05.14.540710 | 2023 | Robust and interpretable prediction of gene markers and cell types from spatial transcriptomics data | AGREE | Title: 'Robust and interpretable prediction of gene markers and cell types from spatial transcriptomics data'; gemma_final=FILTERED; gemma_q1 empty; gemma_q2 says spatial=YES, paper_type=COMP. | accepted_agree |
| 10.1101/2023.05.29.542760 | 2023 | Boosting Single-Cell RNA Sequencing Analysis with Simple Neural Attention | AGREE | citation_screening_prior: computational-only prior; gemma_final source=citation_screening_prefilter -> FILTERED; q1 panel-design fields are empty; q2 paper_type=COMP. | accepted_agree |
| 10.1101/2023.06.19.545592 | 2024 | InGene: Finding influential genes from embeddings of nonlinear dimension reduction techniques | AGREE | paper_type/profile evidence: COMP; citation-screening prior labels=['COMP-BENCH']; gemma_final notes: computational-only prior pre-filter | accepted_agree |
| 10.1101/2023.06.26.546618 | 2023 | Dendrimeric DNA Coordinate Barcoding Design for Spatial RNA Sequencing | AGREE | Title: 'Dendrimeric DNA Coordinate Barcoding Design for Spatial RNA Sequencing'; gemma_final=FILTERED; gemma_q1 empty; gemma_q2 | accepted_agree |

| DOI | Year | Title | Audit decision | Evidence (verbatim quote) | Validation status / scope flag |
|---|---|---|---|---|---|
| | | | | says spatial=YES, paper_type=COMP. | |
| 10.1101/2023.07.05.547784 | 2023 | scARE: Attribution Regularization for Single Cell Representation Learning | AGREE | title='scARE: Attribution Regularization for Single Cell Representation Learning'; citation_screening_prior.labels='COMP'; gemma_final.source='citation_screening_prefilter' | accepted_agree |
| 10.1101/2023.07.05.547797 | 2023 | ABDS: tool suite for analyzing biologically diverse samples | AGREE | title='ABDS: tool suite for analyzing biologically diverse samples'; citation_screening_prior.labels='COMP'; gemma_final.source='citation_screening_prefilter' | accepted_agree |
| 10.1101/2023.07.11.548486 | 2023 | FAST: a fast and scalable factor analysis for spatially aware dimension reduction of multi-section spatial transcriptomics data | AGREE | Title: 'FAST: a fast and scalable factor analysis for spatially aware dimension reduction...'; gemma_final=FILTERED; gemma_q1 empty; gemma_q2 says spatial=YES, paper_type=COMP. | accepted_agree |
| 10.1101/2023.07.13.548611 | 2023 | Pooled tagging and hydrophobic targeting of endogenous proteins for unbiased mapping of unfolded protein responses | AGREE | Title: 'Pooled tagging and hydrophobic targeting of endogenous proteins...'; gemma_final=FILTERED; gemma_q1 empty; gemma_q2 says spatial=YES, paper_type=COMP. | accepted_agree |
| 10.1101/2023.07.17.549344 | 2023 | Recurrent Composite Markers of Cell Types and States | AGREE | title: Recurrent Composite Markers of Cell Types and States; q2 paper_type=COMP; q1 panel-design fields are empty; citation prior labels=COMP and Gemma_final=FILTERED. | accepted_agree |
| 10.1101/2023.08.28.554970 | 2023 | SIMVI reveals intrinsic and spatial-induced states in spatial omics data | AGREE | Title: 'SIMVI reveals intrinsic and spatial-induced states in spatial omics data'; gemma_final=FILTERED; gemma_q1 empty; gemma_q2 says spatial=YES, paper_type=COMP. | accepted_agree |
| 10.1101/2023.10.09.23296701 | 2023 | Feasibility of Inferring Spatial Transcriptomics from Single-Cell Histological Patterns for Studying | AGREE | Title: 'Feasibility of Inferring Spatial Transcriptomics from Single-Cell Histological Patterns...'; gemma_final=FILTERED; | accepted_agree |

| DOI | Year | Title | Audit decision | Evidence (verbatim quote) | Validation status / scope flag |
|---|---|---|---|---|---|
| | | Colon Cancer Tumor Heterogeneity | | gemma_q1 empty; gemma_q2 says spatial=YES, paper_type=COMP, panel_scope=NOT_TARGETED. | |
| 10.1101/2023.11.04.565623 | 2023 | Exploring building blocks of cell organization by estimating network motifs using graph isomorphism network | AGREE | In a case study using STARmap Plus technologies, TrimNN identified several biologically meaningful large-size network motifs. | accepted_agree |
| 10.1101/2023.11.08.566343 | 2023 | MEM-GAN: A Pseudo Membrane Generator for Single-cell Imaging in Fluorescent Microscopy | NEEDS_MANUAL | Mem-GAN is evaluated on CosMx and adapted to MERFISH/FISHseq for membrane generation and segmentation. | pdf_validated_unclear_update |
| 10.1101/2023.11.09.566505 | 2023 | Targeting CD74-positive macrophages improves neoadjuvant therapy in cervical cancer as revealed by single-cell transcriptomics analysis | AGREE | title='Targeting CD74-positive macrophages improves neoadjuvant therapy in cervical cancer as revealed by single-cell transcriptomics analysis'; citation_screening_prior.labels='NON-ST'; gemma_final.source='citation_screening_prefilter' | accepted_agree |
| 10.1101/2023.11.10.566642 | 2023 | CGCom: a framework for inferring Cell-cell Communication based on Graph Neural Network | AGREE | The authors' seqFISH dataset provided spatially resolved expressions of 387 target genes. | accepted_agree |
| 10.1101/2023.12.03.569806 | 2023 | Landscape and functional repertoires of long noncoding RNAs in the pan-cancer tumor microenvironment using single-nucleus total RNA sequencing | AGREE | title='Landscape and functional repertoires of long noncoding RNAs in the pan-cancer tumor microenvironment using single-nucleus total RNA sequencing'; citation_screening_prior.labels='COMP'; gemma_final.source='citation_screening_prefilter' | accepted_agree |
| 10.1101/2023.12.26.573143 | 2023 | Highly multiplexed, image-based pooled screens in primary cells and tissues with PerturbView | AGREE | Title: 'Highly multiplexed, image-based pooled screens in primary cells and tissues with PerturbView'; gemma_final=FILTERED; gemma_q1 empty; gemma_q2 says spatial=YES, paper_type=COMP. | accepted_agree |
| 10.1101/2023.12.27.573479 | 2023 | ASCT: automatic single-cell toolbox in julia | AGREE | title='ASCT: automatic single-cell toolbox in julia'; citation_screening_prior.labels='COMP'; | accepted_agree |

| DOI | Year | Title | Audit decision | Evidence (verbatim quote) | Validation status / scope flag |
|---|---|---|---|---|---|
| | | | | gemma_final.source='citation_screening_prefilter' | |
| 10.1101/2024.01.31.576150 | 2024 | Charting the Spatial Transcriptome of the Human Cerebral Cortex at Single-Cell Resolution | AGREE | paper title (`Charting the Spatial Transcriptome of the Human Cerebral Cortex at Single-Cell Resolution`); `paper_type=EXP`; `citation_screening_prior.summary`; empty `gemma_q1_panel_design`; `gemma_final.classification=FILTERED`. | accepted_agree |
| 10.1101/2024.03.23.586413 | 2024 | Algorithms for a Commons Cell Atlas | AGREE | paper title (`Algorithms for a Commons Cell Atlas`); `paper_type=COMP`; `citation_screening_prior.summary`; empty `gemma_q1_panel_design`; `gemma_final.classification=FILTERED`. | accepted_agree |
| 10.1101/2024.04.08.588510 | 2024 | scRCA: a Siamese network-based pipeline for the annotation of cell types using imperfect single-cell RNA-seq reference data | AGREE | paper title (`scRCA: a Siamese network-based pipeline for the annotation of cell types using imperfect single-cell RNA-seq reference data`); `paper_type=COMP`; `citation_screening_prior.summary`; empty `gemma_q1_panel_design`; `gemma_final.classification=FILTERED`. | accepted_agree |
| 10.1101/2024.04.08.588641 | 2024 | Empirical Bayes spline model learns multi-way genomic interactions from single cell 3D genome data | AGREE | We introduce MINTsC to learn Multi-way INTeractions from single cell Hi-C. | accepted_agree |
| 10.1101/2024.04.24.590691 | 2024 | mastR: Marker Automated Screening Tool for multi-omics data | AGREE | title: mastR: Marker Automated Screening Tool for multi-omics data; q2 paper_type=COMP; q1 panel-design fields are empty; citation prior labels=COMP and Gemma_final=FILTERED. | accepted_agree |
| 10.1101/2024.05.27.594872 | 2024 | Towards a universal spatial molecular atlas of the mouse brain | AGREE | paper title (`Towards a universal spatial molecular atlas of the mouse brain`); `paper_type=COMP`; `citation_screening_prior.summary`; empty | accepted_agree |

| DOI | Year | Title | Audit decision | Evidence (verbatim quote) | Validation status / scope flag |
|---|---|---|---|---|---|
| | | | | `gemma_q1_panel_design`; `gemma_final.classification=FILTERED`. | |
| 10.1101/2024.05.28.596283 | 2024 | Beyond variability: a novel gene expression stability metric to unveil homeostasis and regulation | AGREE | paper title (`Beyond variability: a novel gene expression stability metric to unveil homeostasis and regulation`); `paper_type=COMP`; `citation_screening_prior.summary`; empty `gemma_q1_panel_design`; `gemma_final.classification=FILTERED`. | accepted_agree |
| 10.1101/2024.05.31.596908 | 2024 | SpaNorm: spatially-aware normalisation for spatial transcriptomics data | AGREE | Title: 'SpaNorm: spatially-aware normalisation for spatial transcriptomics data'; gemma_final=FILTERED; gemma_q1 empty; gemma_q2 says spatial=YES, paper_type=COMP. | accepted_agree |
| 10.1101/2024.06.12.598686 | 2024 | Inferring single-cell spatial gene expression with tissue morphology via explainable deep learning | AGREE | The 379 genes measured by the Xenium panel were sequenced using the FFPE protocol. | accepted_agree |
| 10.1101/2024.06.24.600550 | 2024 | Lipidic and senescent macrophages predict progression and response to combinatorial immunotherapy in triple-negative breast cancer | AGREE | paper title (`Lipidic and senescent macrophages predict progression and response to combinatorial immunotherapy in triple-negative breast cancer`); `paper_type=COMP`; `citation_screening_prior.summary`; empty `gemma_q1_panel_design`; `gemma_final.classification=FILTERED`. | accepted_agree |
| 10.1101/2024.06.26.600708 | 2024 | MuCST: restoring and integrating heterogeneous morphology images and spatial transcriptomics data with contrastive learning | AGREE | Title: 'MuCST: restoring and integrating heterogeneous morphology images and spatial transcriptomics data with contrastive learning'; gemma_final=FILTERED; gemma_q1 empty; gemma_q2 says spatial=YES, paper_type=COMP. | accepted_agree |
| 10.1101/2024.07.03.601914 | 2024 | A DIY guide for image-based spatial transcriptomic: TLS as a case example | AGREE | Title is 'A DIY guide for image-based spatial transcriptomic: TLS as a case example'; gemma notes | accepted_agree |

| DOI | Year | Title | Audit decision | Evidence (verbatim quote) | Validation status / scope flag |
|---|---|---|---|---|---|
| | | | | say OTHER -> FILTERED and q1 panel-design fields are empty. | |
| 10.1101/2024.07.04.601829 | 2024 | SPArrOW: a flexible, interactive and scalable pipeline for spatial transcriptomics analysis | AGREE | paper title (`SPArrOW: a flexible, interactive and scalable pipeline for spatial transcriptomics analysis`); `paper_type=COMP`; `citation_screening_prior.summary` ; empty `gemma_q1_panel_design`; `gemma_final.classification=FILTERED`. | accepted_agree |
| 10.1101/2024.07.05.602236 | 2024 | SOFisher: Reinforcement Learning-Guided Experiment Designs for Spatial Omics | AGREE | Title is 'SOFisher: Reinforcement Learning-Guided Experiment Designs for Spatial Omics'; citation prior marks it computational-only and q1 panel-design fields are empty. | accepted_agree |
| 10.1101/2024.07.09.602652 | 2024 | Visual analysis of spatial transcriptomics data with RedeViz | AGREE | Title: 'Visual analysis of spatial transcriptomics data with RedeViz'; gemma_final=FILTERED; gemma_q1 empty; gemma_q2 says spatial=YES, paper_type=COMP. | accepted_agree |
| 10.1101/2024.07.23.604776 | 2024 | Rapid and memory-efficient analysis and quality control of large spatial transcriptomics datasets | AGREE | Title: 'Rapid and memory-efficient analysis and quality control of large spatial transcriptomics datasets'; gemma_final=FILTERED; gemma_q1 empty; gemma_q2 says spatial=YES, paper_type=COMP. | accepted_agree |
| 10.1101/2024.08.04.606512 | 2024 | Identifying Differential Spatial Expression Patterns across Different Slices, Conditions and Developmental Stages with Interpretable Deep Learning | AGREE | paper title (`Identifying Differential Spatial Expression Patterns across Different Slices, Conditions and Developmental Stages with Interpretable Deep Learning`); `paper_type=COMP`; `citation_screening_prior.summary` ; empty `gemma_q1_panel_design`; `gemma_final.classification=FILTERED`. | accepted_agree |
| 10.1101/2024.10.19.619181 | 2024 | SPATIAL TRANSCRIPTOMIC PROFILING OF ADVANCED OVARIAN CLEAR CELL CARCINOMA REVEALS INTRA- | AGREE | Title: 'SPATIAL TRANSCRIPTOMIC PROFILING OF ADVANCED OVARIAN CLEAR CELL CARCINOMA...'; | accepted_agree |

| DOI | Year | Title | Audit decision | Evidence (verbatim quote) | Validation status / scope flag |
|---|---|---|---|---|---|
| | | TUMOR HETEROGENEITY IN EPITHELIAL-MESENCHYMAL GRADIENT | | gemma_final=FILTERED; gemma_q1 empty; gemma_q2 says spatial=YES, paper_type=COMP, panel_scope=UNCLEAR. | |
| 10.1101/2024.10.27.620174 | 2024 | Directing cellular transitions on gene graph-enhanced cell state manifold | AGREE | paper title (`Directing cellular transitions on gene graph-enhanced cell state manifold`); `paper_type=COMP`; `citation_screening_prior.summary`; empty `gemma_q1_panel_design`; `gemma_final.classification=FILTERED`. | accepted_agree |
| 10.1101/2024.11.07.622485 | 2024 | <i>FOXP</i> Genes Regulate Purkinje Cell Diversity in Cerebellar Development and Evolution | AGREE | paper title (`<i>FOXP</i> Genes Regulate Purkinje Cell Diversity in Cerebellar Development and Evolution`); `paper_type=EXP`; `citation_screening_prior.summary`; empty `gemma_q1_panel_design`; `gemma_final.classification=FILTERED`. | accepted_agree |
| 10.1101/2024.11.21.624653 | 2024 | Identifying spatially variable genes by projecting to morphologically relevant curves | AGREE | Title: 'Identifying spatially variable genes by projecting to morphologically relevant curves'; gemma_final=FILTERED; gemma_q1 empty; gemma_q2 says spatial=YES, paper_type=COMP. | accepted_agree |
| 10.1101/2024.11.25.625243 | 2024 | NoButter: An R package for reducing transcript dispersion in CosMx Spatial Molecular Imaging Data | NEEDS_MANUAL | PDF: "R package for reducing transcript dispersion in CosMx SMI data". | pdf_validated_unclear_update |
| 10.1101/2024.11.30.626148 | 2024 | Unbiased learning of protein conformational representation via unsupervised random forest | AGREE | paper title (`Unbiased learning of protein conformational representation via unsupervised random forest`); `paper_type=COMP`; `citation_screening_prior.summary`; empty `gemma_q1_panel_design`; `gemma_final.classification=FILTERED`. | accepted_agree |
| 10.1101/2024.12.23.630040 | 2024 | CORTADO: Hill Climbing Optimization for Cell-Type Specific | AGREE | Title is 'CORTADO: Hill Climbing Optimization for Cell-Type Specific | accepted_agree |

| DOI | Year | Title | Audit decision | Evidence (verbatim quote) | Validation status / scope flag |
|---|---|---|---|---|---|
| | | Marker Gene Discovery | | Marker Gene Discovery'; citation prior marks it computational-only and q1 panel-design fields are empty. | |
| 10.1101/2025.01.07.631783 | 2025 | BellaVista: Open-Source Visualization for Imaging-Based Spatial Transcriptomics | AGREE | Title: 'BellaVista: Open-Source Visualization for Imaging-Based Spatial Transcriptomics'; gemma_final=FILTERED; gemma_q1 empty; gemma_q2 says spatial=YES, paper_type=COMP. | accepted_agree |
| 10.1101/2025.01.18.633701 | 2025 | SpatialFormer: Universal Spatial Representation Learning from Subcellular Molecular to Multicellular Landscapes | AGREE | The gene panels of 10X Genomics were chosen to accurately type cells, and identify select immune, proliferation, and tumor markers. | accepted_agree |
| 10.1101/2025.01.26.634963 | 2025 | Single-Cell Genomics Elucidates Molecular Variations and Regulatory Mechanisms in Circulating Immune Cells | AGREE | title: Single-Cell Genomics Elucidates Molecular Variations and Regulatory Mechanisms in Circulating Immune Cells; citation prior labels=NON-ST; spatial=NO; q1 panel-design fields are empty. | accepted_agree |
| 10.1101/2025.01.30.635803 | 2025 | Identification of High-Risk Cells in Single-Cell Spatially Resolved Transcriptomics Data Using DEGAS Spatial Smoothing | AGREE | DEGAS was applied to our newly generated Type II Diabetes Xenium dataset. | accepted_agree |
| 10.1101/2025.02.02.636138 | 2025 | scBSP: A fast and accurate tool for identifying spatially variable features from high-resolution spatial omics data | AGREE | paper title; gemma_final notes/source; q2 profile says COMP/spatial with no panel-selection evidence; empty q1 panel-design fields. | accepted_agree |
| 10.1101/2025.02.05.636735 | 2025 | Φ-Space ST: a platform-agnostic method to identify cell states in spatial transcriptomics studies | AGREE | paper title; gemma_final notes/source; q2 profile says COMP/spatial with no panel-selection evidence; empty q1 panel-design fields. | accepted_agree |
| 10.1101/2025.02.12.637849 | 2025 | Optimal marker genes for <i>c</i>-separated cell types | AGREE | paper title; citation-screening prior marks computational-only; gemma_final notes/source; q2 panel fields are absent/empty. | accepted_agree |
| 10.1101/2025.02.13.638045 | 2025 | Tissue Reassembly with Generative AI | AGREE | paper title; gemma_final notes/source; q2 profile says COMP/spatial with no panel-selection evidence; empty q1 panel-design fields. | accepted_agree |

| DOI | Year | Title | Audit decision | Evidence (verbatim quote) | Validation status / scope flag |
|---|---|---|---|---|---|
| 10.1101/2025.02.16.638209 | 2025 | Extending scope and power of circular RNA research with circtools 2.0 | AGREE | The module identifies 50-bp long target sequences and positions two adjacent 20-bp padlock probes for each target sequence. | accepted_agree |
| 10.1101/2025.02.17.638195 | 2025 | Joint imputation and deconvolution of gene expression across spatial transcriptomics platforms | AGREE | paper title; gemma_final notes/source; q2 profile says COMP/spatial with no panel-selection evidence; empty q1 panel-design fields. | accepted_agree |
| 10.1101/2025.02.24.640007 | 2025 | Geospatially informed representation of spatial genomics data with SpatialFeatureExperiment | AGREE | paper title; gemma_final notes/source; q2 profile says COMP/spatial with no panel-selection evidence; empty q1 panel-design fields. | accepted_agree |
| 10.1101/2025.03.05.641709 | 2025 | Leveraging cell-type specificity and similarity improves single-cell eQTL fine-mapping | AGREE | title: Leveraging cell-type specificity and similarity improves single-cell eQTL fine-mapping; citation prior labels=COMP; q1 panel-design fields are empty; gemma_final=FILTERED. | accepted_agree |
| 10.1101/2025.03.06.641951 | 2025 | Sequencing-free whole genome spatial transcriptomics at molecular resolution in intact tissue | AGREE | paper title; gemma_final notes/source; q2 profile says COMP/spatial and panel_scope=NOT_TARGETED; empty q1 panel-design fields. | accepted_agree |
| 10.1101/2025.03.09.642280 | 2025 | Decipher cell communication with attention: CLARA | NEEDS_MANUAL | CLARA uses a mouse embryo SeqFISH dataset and a human cartilage dataset to infer communication; no panel design is described. | pdf_validated_unclear_update |
| 10.1101/2025.03.14.643160 | 2025 | Segger: Fast and accurate cell segmentation of imaging-based spatial transcriptomics data | AGREE | We applied segger and leading alternative cell-segmentation methods to a widely used 10x Genomics Xenium dataset derived from FFPE human breast cancer tissue (313 gene panel). | accepted_agree |
| 10.1101/2025.03.25.645278 | 2025 | <i>In Situ</i> Graphene-Seq: Spatial Transcriptomics and Chronic Electrophysiological Characterization of Tissue Microenvironments | DISAGREE | Final manual decision: FILTERED because the paper is a downstream analysis/modeling/QC method or dataset-use paper, with no final panel-design source. Original evidence: Q1 panel_design is UNCLEAR and no tool or vendor source is found; the | manual_final_filter_after_pdf_review |

| DOI | Year | Title | Audit decision | Evidence (verbatim quote) | Validation status / scope flag |
|---|---|---|---|---|---|
| | | | | packet only identifies targeted in situ graphene-seq. | |
| 10.1101/2025.04.03.646459 | 2025 | SpatialAgent: An autonomous AI agent for spatial biology | AGREE | title: SpatialAgent: An autonomous AI agent for spatial biology; q2 paper_type=COMP; q1 panel-design fields are empty; citation prior labels=COMP-BENCH and Gemma_final=FILTERED. | accepted_agree |
| 10.1101/2025.04.09.647847 | 2025 | <i>Tff2</i> marks gastric corpus progenitors that give rise to pyloric metaplasia/SPEM following injury | DISAGREE | 10x Genomics Visium Spatial Gene Expression with a human whole transcriptome probe set is untargeted, so it is out of scope for panel-design classification. | corrected_high_risk_disagreement_untargeted_method |
| 10.1101/2025.04.10.648034 | 2025 | Benchmarking single cell transcriptome matching methods for incremental growth of cell atlases | AGREE | title: Benchmarking single cell transcriptome matching methods for incremental growth of cell atlases; q2 paper_type=COMP, panel_scope=UNCLEAR, spatial=NO; q1 panel-design fields are empty. | accepted_agree |
| 10.1101/2025.04.22.649977 | 2025 | spRefine Denoises and Imputes Spatial Transcriptomics with a Reference-Free Framework Powered by Genomic Language Model | AGREE | title: spRefine Denoises and Imputes Spatial Transcriptomics with a Reference-Free Framework Powered by Genomic Language Model; citation prior labels=COMP; q1 panel-design fields are empty; gemma_final=FILTERED. | accepted_agree |
| 10.1101/2025.05.08.652741 | 2025 | Popari: Modeling multisample variation in spatial transcriptomics | AGREE | paper title; gemma_final notes/source; q2 profile says COMP/spatial with no panel-selection evidence; empty q1 panel-design fields. | accepted_agree |
| 10.1101/2025.05.09.652066 | 2025 | Characterization of hyperoxia-induced senescent lung macrophages in neonatal mice | AGREE | title: Characterization of hyperoxia-induced senescent lung macrophages in neonatal mice; citation prior labels=COMP; q2 paper_type=COMP; q1 panel-design fields are empty. | accepted_agree |
| 10.1101/2025.05.16.654329 | 2025 | High-Resolution Spatial Profiling of Microglia Reveals Proximity Associated Immunometabolic Reprogramming in Alzheimer’s Disease | AGREE | gemma_q0_document_type doc_type=NON_ARTICLE; gemma_final notes/source; no q1 panel-design evidence. | accepted_agree |
| 10.1101/2025.05.21.653987 | 2025 | SCALE: Unsupervised Multi-Scale Domain Identification in Spatial | NEEDS_MANUAL | Final manual decision: FILTERED because the paper is a | manual_final_filter_after_pdf_review |

| DOI | Year | Title | Audit decision | Evidence (verbatim quote) | Validation status / scope flag |
|---|---|---|---|---|---|
| | | Omics Data | | downstream analysis/modeling/QC method or dataset-use paper, with no final panel-design source. Original evidence: q1: panel_design=UNCLEAR; q2: targeted Xenium/MERFISH context only. | |
| 10.1101/2025.05.28.656712 | 2025 | Quantitative molecular cartography of emergency myelopoiesis reveals conserved modules of hematopoietic activation | AGREE | paper title; gemma_final notes/source; q2 profile says EXP/non-spatial with no panel evidence; empty q1 panel-design fields. | accepted_agree |
| 10.1101/2025.05.30.656369 | 2025 | HCR-Proxy resolves site-specific proximal RNA proteomes at subcompartmental nanoscale resolution | AGREE | paper title; gemma_final notes/source; q2 profile says COMP/spatial with no panel-selection evidence; empty q1 panel-design fields. | accepted_agree |
| 10.1101/2025.05.30.656960 | 2025 | EnrichMap: Spatially informed enrichment analysis for functional interpretation of spatial transcriptomics | AGREE | paper title; gemma_final notes/source; q2 profile says COMP/spatial with no panel-selection evidence; empty q1 panel-design fields. | accepted_agree |
| 10.1101/2025.06.23.660674 | 2025 | Tracing colorectal malignancy transformation from cell to tissue scale | DISAGREE | CosMx Human WTx panel targets 19,867 genes and is described as whole-transcriptome spatial molecular imaging, so it is untargeted and out of scope for panel-design classification. | corrected_high_risk_disagreement_untargeted_method |
| 10.1101/2025.06.27.661814 | 2025 | Fluctuation structure predicts genome-wide perturbation outcomes | AGREE | title: Fluctuation structure predicts genome-wide perturbation outcomes; citation prior labels=COMP; q1 panel-design fields are empty; gemma_final=FILTERED. | accepted_agree |
| 10.1101/2025.07.03.662904 | 2025 | SpatialMap: A Scalable Deep Learning Method for Cell Typing in Subcellular Spatial Transcriptomics | NEEDS_MANUAL | Final manual decision: FILTERED because the paper is a downstream analysis/modeling/QC method or dataset-use paper, with no final panel-design source. Original evidence: q1: panel_design=UNCLEAR; q2: targeted CosMx/MERFISH/Xenium context; no panel-selection evidence. | manual_final_filter_after_pdf_review |
| 10.1101/2025.07.03.663047 | 2025 | Validation of Morphology-Guided Computational Enhancement for | AGREE | paper title; gemma_final notes/source; q2 profile says | accepted_agree |

| DOI | Year | Title | Audit decision | Evidence (verbatim quote) | Validation status / scope flag |
|---|---|---|---|---|---|
| | | Single-Cell Resolution Spatial Transcriptomics | | COMP/spatial but panel_scope=NOT_TARGETED; empty q1 panel-design fields. | |
| 10.1101/2025.07.09.663835 | 2025 | Granulin <sup>+</sup> macrophages promote lineage plasticity in prostate cancer through paracrine signaling loops | AGREE | title: Granulin <sup>+</sup> macrophages promote lineage plasticity in prostate cancer through paracrine signaling loops; q2 paper_type=EXP, spatial=NO, panel_scope=UNCLEAR; q1 panel-design fields are empty. | accepted_agree |
| 10.1101/2025.07.14.664825 | 2025 | Identifying the Minimal Number of Protein Markers for Cell Type Annotation Using MiniMarS | AGREE | title: Identifying the Minimal Number of Protein Markers for Cell Type Annotation Using MiniMarS; citation prior labels=COMP-BENCH; q1 panel-design fields are empty; gemma_final=FILTERED. | accepted_agree |
| 10.1101/2025.07.24.665835 | 2025 | GraphTME: a framework for predicting response to immune checkpoint inhibitors by interpreting cell-cell interactions in the tumour microenvironment using spatial transcriptomics of tumour tissue | NEEDS_MANUAL | Final manual decision: FILTERED because the paper is a downstream analysis/modeling/QC method or dataset-use paper, with no final panel-design source. Original evidence: q1: panel_design=UNCLEAR; q2: MERFISH dataset context only. | manual_final_filter_after_pdf_review |
| 10.1101/2025.08.02.668295 | 2025 | Human retinal organoid single-cell atlas allows to reconstruct retinal development at high resolution and identify nature restricted transcriptional states in vitro | AGREE | title: Human retinal organoid single-cell atlas allows to reconstruct retinal development at high resolution and identify nature restricted transcriptional states in vitro; citation prior labels=COMP; q1 panel-design fields are empty; gemma_final=FILTERED. | accepted_agree |
| 10.1101/2025.08.08.669400 | 2025 | A scalable approach for tissue microarray profiling using spatial single cell transcriptomics | AGREE | paper title; gemma_final notes/source; q2 profile says OTHER/spatial with panel_scope=TARGETED, but q1 panel-design fields are empty and no panel evidence is provided. | accepted_agree |
| 10.1101/2025.08.14.670318 | 2025 | GRIDGENE: Guided Region Identification based on Density of GENEs – a transcript density-based approach to characterize tissues by spatial transcriptomics | AGREE | CosMx data includes measurements for 999 genes from a standard panel. In contrast, Xenium Issue cores are circular, 1500 um in diameter, and are profiled for 480 genes using a custom panel. | accepted_agree |

| DOI | Year | Title | Audit decision | Evidence (verbatim quote) | Validation status / scope flag |
|---|---|---|---|---|---|
| 10.1101/2025.08.14.670406 | 2025 | Constructing and Using Cell Type Populations of the Human Reference Atlas | AGREE | title: Constructing and Using Cell Type Populations of the Human Reference Atlas; q2 paper_type=OTHER, spatial=NO, panel_scope=UNCLEAR; q1 panel-design fields are empty. | accepted_agree |
| 10.1101/2025.08.22.671846 | 2025 | Energy-Regularized Graph Learning for Multiscale Spatial Representation | AGREE | paper title; gemma_final notes/source; q2 profile says COMP/spatial with no panel-selection evidence; empty q1 panel-design fields. | accepted_agree |
| 10.1101/2025.08.29.673037 | 2025 | ProbeST: a custom probe design pipeline for Spatial Transcriptomics | AGREE | title: ProbeST: a custom probe design pipeline for Spatial Transcriptomics; citation prior labels=COMP; q1 panel-design fields are empty; gemma_final=FILTERED. | accepted_agree |
| 10.1101/2025.09.04.674228 | 2025 | Impact of Data Quality on Deep Learning Prediction of Spatial Transcriptomics from Histology Images | AGREE | paper title; gemma_final notes/source; q2 profile says COMP/spatial with no panel-selection evidence; empty q1 panel-design fields. | accepted_agree |
| 10.1101/2025.09.06.674629 | 2025 | S3R: Modeling spatially varying associations with Spatially Smooth Sparse Regression | AGREE | paper title; gemma_final notes/source; q2 profile says COMP/spatial with no panel-selection evidence; empty q1 panel-design fields. | accepted_agree |
| 10.1101/2025.09.08.674969 | 2025 | Stem-like Prostate Remodeling in Obesity Mediates Resistance to 5α-Reductase Inhibition Therapy in BPH | NEEDS_MANUAL | Final manual decision: FILTERED because manual review found no usable evidence for final gene-panel source selection. Original evidence: q1 tool/vendor/manual evidence: NOT FOUND; q2 only states that Xenium spatial profiling was performed, without naming how the panel was selected. | manual_final_filter_after_pdf_review |
| 10.1101/2025.09.24.678295 | 2025 | Super-resolved spatial transcriptomics reveals early hippocampal RNA localization changes in a mouse model of Alzheimer's disease | NEEDS_MANUAL | Final manual decision: FILTERED because the evidence is spatial profiling context or untargeted spatial transcriptomics without an in-scope panel-design source. Original evidence: q1 tool/vendor/manual evidence: NOT FOUND; q2 only says the study used a targeted panel of 101 genes with no panel-source detail. | manual_final_filter_after_pdf_review |

| DOI | Year | Title | Audit decision | Evidence (verbatim quote) | Validation status / scope flag |
|---|---|---|---|---|---|
| 10.1101/2025.09.24.678397 | 2025 | Mapping disease critical spatially variable gene programs by integrating spatial transcriptomics with human genetics | AGREE | paper title; gemma_final notes/source; q2 profile says COMP/spatial with no panel-selection evidence; empty q1 panel-design fields. | accepted_agree |
| 10.1101/2025.09.30.679413 | 2025 | Exploring the correspondence between gene expression and thalamic nuclei using the THALMANAC resource | AGREE | paper title; gemma_final notes/source; q2 profile says OTHER/spatial with panel_scope=TARGETED, but no panel-selection evidence is provided in q1. | accepted_agree |
| 10.1101/2025.10.01.679918 | 2025 | SpacGPA: annotating spatial transcriptomes through <i>de novo</i> interpretable gene programs | AGREE | paper title; gemma_final notes/source; q2 profile says COMP/spatial with no panel-selection evidence; empty q1 panel-design fields. | accepted_agree |
| 10.1101/2025.10.02.680117 | 2025 | scGPD: single-cell informed gene panel design for targeted spatial transcriptomics | DISAGREE | This is the computational methods paper introducing scGPD. Even though it designs panels inside the method study, that is not a paper-level experimental adoption event and should remain filtered from the panel-design adoption set. | corrected_medium_risk_disagreement_canonical_tool |
| 10.1101/2025.10.08.681071 | 2025 | Optimal Gene Panel Selection for Targeted Spatial Transcriptomics Experiments | DISAGREE | This is a computational methods and benchmarking paper introducing ReconST. Even though it designs panels inside the method study, that is not a paper-level experimental adoption event and should remain filtered from the panel-design adoption set. | corrected_medium_risk_disagreement_canonical_tool |
| 10.1101/2025.10.14.682419 | 2025 | scConcept: Contrastive pretraining for technology-agnostic single-cell representations beyond reconstruction | AGREE | Title is 'scConcept: Contrastive pretraining for technology-agnostic single-cell representations beyond reconstruction'; citation prior marks it computational-only and q1 panel-design fields are empty. | accepted_agree |
| 10.1101/2025.10.17.683188 | 2025 | Spatially varying cell-specific gene regulation network inference | AGREE | paper.title='Spatially varying cell-specific gene regulation network inference'; gemma_final.notes=Pass-A abstract: COMP → FILTERED | accepted_agree |
| 10.1101/2025.10.31.685785 | 2025 | STARCall integrates image stitching, alignment, and read | AGREE | paper.title='STARCall integrates image stitching, alignment, and read calling to enable scalable | accepted_agree |

| DOI | Year | Title | Audit decision | Evidence (verbatim quote) | Validation status / scope flag |
|---|---|---|---|---|---|
| | | calling to enable scalable analysis of in situ sequencing data | | analysis of in situ sequencing data'; gemma_final.notes=Pass-A abstract: COMP → FILTERED | |
| 10.1101/2025.11.08.687410 | 2025 | Can Lightweight LLM Agents Improve Spatial Transcriptomics Annotation? | NEEDS_MANUAL | Final manual decision: FILTERED because the paper is a downstream analysis/modeling/QC method or dataset-use paper, with no final panel-design source. Original evidence: No panel-design evidence is present in the packet. | manual_final_filter_after_pdf_review |
| 10.1101/2025.11.18.689140 | 2025 | CBKMR: A Copula-based Bayesian Kernel Machine Regression Framework for Optimal Marker Detection in Omics Data | AGREE | Title is 'CBKMR: A Copula-based Bayesian Kernel Machine Regression Framework for Optimal Marker Detection in Omics Data'; citation prior marks it computational-only and q1 panel-design fields are empty. | accepted_agree |
| 10.1101/2025.11.20.689462 | 2025 | USHER: Guiding Foundation Model Representations through Distribution Shifts | AGREE | Typical Xenium gene panels are small (300-500 genes). | accepted_agree |
| 10.1101/2025.11.24.690257 | 2025 | Mapping spatial gradients in spatial transcriptomics data with score matching | AGREE | title: Mapping spatial gradients in spatial transcriptomics data with score matching; q2 spatial=YES, panel_scope=UNCLEAR, paper_type=COMP; q1 panel-design fields are empty. | accepted_agree |
| 10.1101/2025.11.24.690325 | 2025 | TissueNarrator: Generative Modeling of Spatial Transcriptomics with Large Language Models | NEEDS_MANUAL | Final manual decision: FILTERED because the paper is a downstream analysis/modeling/QC method or dataset-use paper, with no final panel-design source. Original evidence: No panel-design evidence is present in the packet. | manual_final_filter_after_pdf_review |
| 10.1101/2025.11.26.690796 | 2025 | LARIS enables accurate and efficient ligand and receptor interaction analysis in spatial transcriptomics | AGREE | title: LARIS enables accurate and efficient ligand and receptor interaction analysis in spatial transcriptomics; citation prior labels=COMP; q1 panel-design fields are empty; gemma_final=FILTERED. | accepted_agree |
| 10.1101/324855 | 2018 | starmap: Immersive visualisation of single cell data using smartphone-enabled virtual reality | AGREE | paper.title='starmap: Immersive visualisation of single cell data using smartphone-enabled virtual reality'; gemma_final.notes=Pass-A abstract: COMP → FILTERED | accepted_agree |

| DOI | Year | Title | Audit decision | Evidence (verbatim quote) | Validation status / scope flag |
|---|---|---|---|---|---|
| 10.1101/401810 | 2018 | SABER enables highly multiplexed and amplified detection of DNA and RNA in cells and tissues | DISAGREE | Final manual decision: FILTERED because the available evidence concerns probe/barcode/oligo design or single-gene detection, not final gene-panel selection. Original evidence: q1 only states that OligoMiner was used to design orthogonal oligopaint probe sequences; q1 does not describe the target-gene selection rationale. | manual_final_filter_after_pdf_review |
| 10.1101/507566 | 2018 | Highly multiplexed<i>in situ</i>protein imaging with signal amplification by Immuno-SABER | AGREE | paper.title='Highly multiplexed<i>in situ</i>protein imaging with signal amplification by Immuno-SABER'; gemma_final.notes=Pass-A abstract: COMP → FILTERED | accepted_agree |
| 10.1101/701680 | 2019 | Giotto, a pipeline for integrative analysis and visualization of single-cell spatial transcriptomic data | AGREE | paper.title='Giotto, a pipeline for integrative analysis and visualization of single-cell spatial transcriptomic data'; gemma_final.notes=Pass-A abstract: COMP → FILTERED | accepted_agree |
| 10.1101/722819 | 2019 | In Situ Transcriptome Accessibility Sequencing (INSTA-seq) | AGREE | paper.title='In Situ Transcriptome Accessibility Sequencing (INSTA-seq)'; gemma_final.notes=Pass-A abstract: COMP → FILTERED | accepted_agree |
| 10.1101/gr.271288.120 | 2021 | Characterizing spatial gene expression heterogeneity in spatially resolved single-cell transcriptomic data with nonuniform cellular densities. | AGREE | paper.title='Characterizing spatial gene expression heterogeneity in spatially resolved single-cell transcriptomic data with nonuniform cellular densities.'; gemma_final.notes=Pass-A abstract: COMP → FILTERED | accepted_agree |
| 10.1101/gr.275569.121 | 2021 | A machine learning method for the discovery of minimum marker gene combinations for cell type identification from single-cell RNA sequencing | AGREE | Title: 'A machine learning method for the discovery of minimum marker gene combinations for cell type identification from single-cell RNA sequencing'; packet fields show paper_type=COMP and Gemma's FILTERED prefilter. | accepted_agree |
| 10.1101/gr.276129.121 | 2021 | New frontiers in single-cell genomics | AGREE | Title: 'New frontiers in single-cell genomics'; packet fields show doc_type=REVIEW and Gemma's FILTERED prefilter. | accepted_agree |
| 10.1101/gr.279857.124 | 2025 | Dissecting multilayer cell-cell communications with signaling | AGREE | paper.title='Dissecting multilayer cell-cell communications with | accepted_agree |

| DOI | Year | Title | Audit decision | Evidence (verbatim quote) | Validation status / scope flag |
|---|---|---|---|---|---|
| | | feedback loops from spatial transcriptomics data. | | signaling feedback loops from spatial transcriptomics data.'; gemma_final.notes=Pass-A abstract: COMP → FILTERED | |
| 10.1101/gr.280539.125 | 2025 | Label-free selection of marker genes in single-cell and spatial transcriptomics with geneCover | AGREE | Title is 'Label-free selection of marker genes in single-cell and spatial transcriptomics with geneCover'; gemma notes say COMP -> FILTERED and q1 panel-design fields are empty. | accepted_agree |
| 10.1101/gr.280803.125 | 2025 | Unified integration of spatial transcriptomics across platforms with LLOKI | AGREE | title: Unified integration of spatial transcriptomics across platforms with LLOKI; citation prior labels=COMP; q1 panel-design fields are empty; gemma_final=FILTERED. | accepted_agree |
| 10.1101/gr.281664.125 | 2025 | RECOMB 2025 Special Issue | AGREE | doc_type=EDITORIAL; title: RECOMB 2025 Special Issue; q1 panel-design fields are empty; gemma_final=FILTERED. | accepted_agree |
| 10.1109/bibm58861.2023.10385799 | 2023 | Co-clustering of single-cell RNA-seq data based on weighted non-negative matrix tri-factorization combined with consensus clustering | AGREE | title='Co-clustering of single-cell RNA-seq data based on weighted non-negative matrix tri-factorization combined with consensus clustering'; citation_screening_prior.labels='COMP'; gemma_final.source='citation_screening_prefilter' | accepted_agree |
| 10.1109/bibm62325.2024.10821945 | 2024 | Identification of cell-type-specific genes associated with autism by integrating gene expression and alternative polyadenylation profiles | AGREE | paper title (`Identification of cell-type-specific genes associated with autism by integrating gene expression and alternative polyadenylation profiles`); `paper_type=EXP`; `citation_screening_prior.summary`; empty `gemma_q1_panel_design`; `gemma_final.classification=FILTERED`. | accepted_agree |
| 10.1109/bibm62325.2024.10822193 | 2024 | MuST: Maximizing the Latent Capacity of Spatial Transcriptomics Data with Multi-modality Structure Transformation | AGREE | paper title (`MuST: Maximizing the Latent Capacity of Spatial Transcriptomics Data with Multi-modality Structure Transformation`); `paper_type=COMP`; | accepted_agree |

| DOI | Year | Title | Audit decision | Evidence (verbatim quote) | Validation status / scope flag |
|---|---|---|---|---|---|
| | | | | `citation_screening_prior.summary`; empty `gemma_q1_panel_design`; `gemma_final.classification=FILTERED`. | |
| 10.1109/bibm66473.2025.11356348 | 2025 | Subcellular Resolution of Whole Spatial Transcriptome Framework via Transferring Distribution with SRGene | AGREE | paper.title='Subcellular Resolution of Whole Spatial Transcriptome Framework via Transferring Distribution with SRGene'; gemma_final.notes=Pass-A abstract: COMP → FILTERED | accepted_agree |
| 10.1109/cdc57313.2025.11312726 | 2022 | Orthogonal Nonnegative Matrix Factorization with Sparsity Constraints | AGREE | title='Orthogonal Nonnegative Matrix Factorization with Sparsity Constraints'; citation_screening_prior.labels="; gemma_final.source='abstract_prefilter' | accepted_agree |
| 10.1109/icccr49711.2021.9349286 | 2021 | Rapid and Accurate Regional Star-Map Simulated Method | AGREE | paper.title='Rapid and Accurate Regional Star-Map Simulated Method'; gemma_final.notes=Pass-A abstract: COMP → FILTERED | accepted_agree |
| 10.1109/iceeict53079.2022.9768589 | 2022 | Automated region split algorithm with pre-trained CNN model for tuna classification | AGREE | paper.title='Automated region split algorithm with pre-trained CNN model for tuna classification'; gemma_final.notes=Pass-A abstract: COMP → FILTERED | accepted_agree |
| 10.1109/icpr.2018.8546069 | 2018 | Sequential Fish Catch Forecasting Using Bayesian State Space Models | AGREE | paper.title='Sequential Fish Catch Forecasting Using Bayesian State Space Models'; gemma_final.notes=Pass-A abstract: COMP → FILTERED | accepted_agree |
| 10.1109/jbhi.2024.3370868 | 2024 | Learning Consistency and Specificity of Cells From Single-Cell Multi-Omic Data | AGREE | paper title (`Learning Consistency and Specificity of Cells From Single-Cell Multi-Omic Data`); `paper_type=COMP`; `citation_screening_prior.summary`; empty `gemma_q1_panel_design`; `gemma_final.classification=FILTERED`. | accepted_agree |
| 10.1109/mipro65660.2025.11132056 | 2025 | Accelerating Marker Gene Selection in Scrna-Seq | AGREE | title: Accelerating Marker Gene Selection in Scrna-Seq; citation prior labels=COMP-BENCH; q1 panel-design fields are empty; q2 | accepted_agree |

| DOI | Year | Title | Audit decision | Evidence (verbatim quote) | Validation status / scope flag |
|---|---|---|---|---|---|
| | | | | paper_type=COMP; gemma_final=FILTERED. | |
| 10.1109/oceanschennai45887.2022.9775327 | 2022 | Sequential Fish Catch Counter Using Vision-based Fish Detection and Tracking | AGREE | paper.title='Sequential Fish Catch Counter Using Vision-based Fish Detection and Tracking'; gemma_final.notes=Pass-A abstract: COMP → FILTERED | accepted_agree |
| 10.1109/ojcoms.2025.3593088 | 2025 | STARMAP: Spaceborne Target Acquisition Radar With Meta-RL Assisted Placement | AGREE | paper.title='STARMAP: Spaceborne Target Acquisition Radar With Meta-RL Assisted Placement'; gemma_final.notes=Pass-A abstract: COMP → FILTERED | accepted_agree |
| 10.1109/tcbb.2022.3203185 | 2022 | A Clustering Method Unifying Cell-Type Recognition and Subtype Identification for Tumor Heterogeneity Analysis | AGREE | title='A Clustering Method Unifying Cell-Type Recognition and Subtype Identification for Tumor Heterogeneity Analysis'; citation_screening_prior.labels='COMP'; gemma_final.source='citation_screening_prefilter' | accepted_agree |
| 10.1109/tcbb.2023.3298334 | 2023 | Multi-View Clustering With Graph Learning for scRNA-Seq Data | AGREE | title='Multi-View Clustering With Graph Learning for scRNA-Seq Data'; citation_screening_prior.labels='COMP'; gemma_final.source='citation_screening_prefilter' | accepted_agree |
| 10.1109/tcbbio.2025.3609721 | 2025 | Knowledge-Guided Gene Panel Selection for Label-Free Single-Cell RNA-Seq Data: A Reinforcement Learning Perspective | AGREE | Title is 'Knowledge-Guided Gene Panel Selection for Label-Free Single-Cell RNA-Seq Data: A Reinforcement Learning Perspective'; citation prior marks it computational-only and q1 panel-design fields are empty. | accepted_agree |
| 10.1109/tcbbio.2025.3624040 | 2025 | scMID: A Deep Multi-Omics Integration Framework for Comprehensive Single-Cell Data Analysis | AGREE | title: scMID: A Deep Multi-Omics Integration Framework for Comprehensive Single-Cell Data Analysis; citation prior labels=COMP; q1 panel-design fields are empty; q2 paper_type=COMP; gemma_final=FILTERED. | accepted_agree |
| 10.1109/tcbbio.2026.3676606 | 2026 | DyGFormer:Transformer With Resistance-Distance Bias for | AGREE | paper.title='DyGFormer:Transformer With Resistance-Distance Bias for Semi-Supervised Cell-Type | accepted_agree |

| DOI | Year | Title | Audit decision | Evidence (verbatim quote) | Validation status / scope flag |
|---|---|---|---|---|---|
| | | Semi-Supervised Cell-Type Identification. | | Identification.'; gemma_final.notes=Pass-A abstract: COMP → FILTERED | |
| 10.1109/tit.2019.2962681 | 2018 | SqueezeFit: Label-Aware Dimensionality Reduction by Semidefinite Programming | AGREE | Title: 'SqueezeFit: Label-Aware Dimensionality Reduction by Semidefinite Programming'; packet fields show paper_type=COMP and Gemma's FILTERED prefilter. | accepted_agree |
| 10.1109/trustcom63139.2024.00294 | 2024 | scE(match): Privacy-Preserving Cluster Matching of Single-Cell Data | AGREE | paper title (`scE(match): Privacy-Preserving Cluster Matching of Single-Cell Data`); `paper_type=COMP`; `citation_screening_prior.summary`; empty `gemma_q1_panel_design`; `gemma_final.classification=FILTERED`. | accepted_agree |
| 10.1109/tvcg.2025.3636102 | 2025 | CellScout: Visual Analytics for Mining Biomarkers in Cell State Discovery | AGREE | title: CellScout: Visual Analytics for Mining Biomarkers in Cell State Discovery; citation prior labels=COMP; q1 panel-design fields are empty; q2 paper_type=COMP; gemma_final=FILTERED. | accepted_agree |
| 10.1111/1462-2920.14109 | 2018 | Metabolic potential and in situ transcriptomic profiles of previously uncharacterized key microbial groups involved in coupled carbon, nitrogen and sulfur cycling in anoxic marine zones | AGREE | paper.title='Metabolic potential and in situ transcriptomic profiles of previously uncharacterized key microbial groups involved in coupled carbon, nitrogen and sulfur cycling in anoxic marine zones'; gemma_final.notes=Pass-A abstract: non-spatial → FILTERED | accepted_agree |
| 10.1111/aor.12785 | 2017 | Adsorption as a Contributor for Inflammatory Mediators Removal by Different Hemofiltration Membranes: A Pilot Study | AGREE | paper.title='Adsorption as a Contributor for Inflammatory Mediators Removal by Different Hemofiltration Membranes: A Pilot Study'; gemma_final.notes=Pass-A abstract: non-spatial → FILTERED | accepted_agree |
| 10.1111/biom.13926 | 2023 | Multiresolution Categorical Regression for Interpretable Cell-Type Annotation | AGREE | title='Multiresolution Categorical Regression for Interpretable Cell-Type Annotation'; citation_screening_prior.labels='COMP'; gemma_final.source='citation_screening_prefilter' | accepted_agree |

| DOI | Year | Title | Audit decision | Evidence (verbatim quote) | Validation status / scope flag |
|---|---|---|---|---|---|
| 10.1111/febs.15572 | 202 | Spage2vec: Unsupervised representation of localized spatial gene expression signatures | AGREE | title: Spage2vec: Unsupervised representation of localized spatial gene expression signatures; final notes: Pass-A abstract: COMP -> FILTERED; q2 profile: spatial=YES, scope=UNCLEAR, panel fields empty. | accepted_agree |
| 10.1111/gtc.70098 | 2026 | Developing a Robust Multiround HCR-FISH Method Modified for Caenorhabditis elegans. | AGREE | title: Developing a Robust Multiround HCR-FISH Method Modified for Caenorhabditis elegans.; final notes: Pass-A abstract: OTHER -> FILTERED; q2 profile: spatial=YES, scope=TARGETED, panel fields empty. | accepted_agree |
| 10.1111/imcb.12256 | 2019 | A method for transcriptome-wide gene expression quantification in intact tissues | AGREE | title: A method for transcriptome-wide gene expression quantification in intact tissues; final notes: Pass-A abstract: OTHER -> FILTERED; q2 profile: spatial=YES, scope=TARGETED, panel fields empty. | accepted_agree |
| 10.1111/jnc.70315 | 2025 | Considerations for Spatial Omics, Metabolite Analyses, and Tissue-Harvesting Artifacts | AGREE | title: Considerations for Spatial Omics, Metabolite Analyses, and Tissue-Harvesting Artifacts; final notes: Pass-A abstract: OTHER -> FILTERED; q2 profile: spatial=YES, scope=UNCLEAR, panel fields empty. | accepted_agree |
| 10.1111/ped.15859 | 2025 | Monitoring measurable residual disease in NUP98::NSD1-positive acute myeloid leukemia. | AGREE | title: Monitoring measurable residual disease in NUP98::NSD1-positive acute myeloid leukemia.; final notes: Pass-A abstract: non-spatial -> FILTERED; q2 profile: spatial=NO. | accepted_agree |
| 10.1111/tpj.14276 | 2019 | In situ transcriptomic and metabolomic study of the loss of photosynthesis in the leaves of mixotrophic plants exploiting fungi. | NEEDS_MANUAL | spatial=NO; packet does not support an in-scope spatial panel-design record. Original audit evidence: q1 panel_design: NOT FOUND; q2 spatial: NOT FOUND. | validated_high_risk_manual_filtered |
| 10.1111/tpj.70049 | 2025 | ExPOSE: a comprehensive toolkit to perform expansion microscopy in plant protoplast systems. | AGREE | title: ExPOSE: a comprehensive toolkit to perform expansion microscopy in plant protoplast systems.; final notes: Pass-A abstract: OTHER -> FILTERED; q2 profile: spatial=YES, | accepted_agree |

| DOI | Year | Title | Audit decision | Evidence (verbatim quote) | Validation status / scope flag |
|---|---|---|---|---|---|
| | | | | scope=UNCLEAR, panel fields empty. | |
| 10.1126/science.371.6528.477-l | 2021 | Identifying transcript location in cells | AGREE | title: Identifying transcript location in cells; final notes: Pass-A abstract: COMP -> FILTERED; q2 profile: spatial=YES, scope=NOT_TARGETED, panel fields empty. | accepted_agree |
| 10.1126/science.aay3446 | 202 | In situ genome sequencing resolves DNA sequence and structure in intact biological samples | AGREE | title: In situ genome sequencing resolves DNA sequence and structure in intact biological samples; final notes: Pass-A abstract: COMP -> FILTERED; q2 profile: spatial=YES, scope=UNCLEAR, panel fields empty. | accepted_agree |
| 10.1126/science.abq4964 | 2023 | The dawn of spatial omics | AGREE | title='The dawn of spatial omics'; citation_screening_prior.labels='REV'; gemma_final.source='citation_screening_prefilter' | accepted_agree |
| 10.1126/science.adq2084 | 2025 | Deep-tissue transcriptomics and subcellular imaging at high spatial resolution. | AGREE | title: Deep-tissue transcriptomics and subcellular imaging at high spatial resolution.; final notes: Pass-A abstract: COMP -> FILTERED; q2 profile: spatial=YES, scope=UNCLEAR, panel fields empty. | accepted_agree |
| 10.1126/science.adt3130 | 2026 | Chinese Immune Multi-Omics Atlas | AGREE | citation_screening_prior labels=NON-ST and all_tools=COSG; gemma_final source=citation_screening_prefilter -> FILTERED; q1 panel-design fields are empty. | accepted_agree |
| 10.1126/science.adu7265 | 2026 | Family single-cell atlases reveal pig pregnancy and fetal growth restriction critical cell types | AGREE | citation_screening_prior labels=NON-ST and all_tools=COSG; gemma_final source=citation_screening_prefilter -> FILTERED; q1 panel-design fields are empty. | accepted_agree |
| 10.1126/scitranslmed.adh1315 | 2025 | Dysfunctional CD11c−CD21− extrafollicular memory B cells are enriched in the periphery and tumors of patients with cancer | AGREE | title: Dysfunctional CD11c−CD21− extrafollicular memory B cells are enriched in the periphery and tumors of patients with cancer; final notes: Pass-A abstract: non- | accepted_agree |

| DOI | Year | Title | Audit decision | Evidence (verbatim quote) | Validation status / scope flag |
|---|---|---|---|---|---|
| | | | | spatial -> FILTERED; q2 profile: spatial=NO. | |
| 10.1128/aem.01807-20 | 202 | Active Soil Nitrifying Communities Revealed by <i>In Situ</i> Transcriptomics and Microcosm-Based Stable-Isotope Probing. | AGREE | title: Active Soil Nitrifying Communities Revealed by In Situ Transcriptomics and Microcosm-Based Stable-Isotope Probing.; final notes: Pass-A abstract: non-spatial -> FILTERED; q2 profile: spatial=NO. | accepted_agree |
| 10.1128/mbio.00004-24 | 2024 | Deep-sea in situ and laboratory multi-omics provide insights into the sulfur assimilation of a deep-sea Chloroflexotabacterium | AGREE | title: Deep-sea in situ and laboratory multi-omics provide insights into the sulfur assimilation of a deep-sea Chloroflexotabacterium; final notes: Pass-A abstract: non-spatial -> FILTERED; q2 profile: spatial=NO. | accepted_agree |
| 10.1128/msystems.00601-21 | 2021 | Chemical gradients of plant substrates in an atta texana fungus garden | AGREE | title: Chemical gradients of plant substrates in an atta texana fungus garden; final notes: Pass-A abstract: non-spatial -> FILTERED; q2 profile: spatial=NO. | accepted_agree |
| 10.1136/annrheumdis-2023-eular.728 | 2023 | AB0167 SINGLE-CELL SPATIAL TRANSCRIPTOME ANALYSIS IDENTIFIES DISEASE-SPECIFIC FIBROBLASTS IN RHEUMATOID VASCULITIS | AGREE | title: AB0167 SINGLE-CELL SPATIAL TRANSCRIPTOME ANALYSIS IDENTIFIES DISEASE-SPECIFIC FIBROBLASTS IN RHEUMATOID VASCULITIS; final notes: Q0: NON_ARTICLE -> FILTERED; q1 panel fields empty. | accepted_agree |
| 10.1136/annrheumdis-2023-eular.767 | 2023 | POS0288 DEGRANULATING PR3+ MYELOID CELLS CHARACTERIZE PROLIFERATIVE LUPUS NEPHRITIS | AGREE | title: POS0288 DEGRANULATING PR3+ MYELOID CELLS CHARACTERIZE PROLIFERATIVE LUPUS NEPHRITIS; final notes: Pass-A abstract: non-spatial -> FILTERED; q2 profile: spatial=NO. | accepted_agree |
| 10.1136/jitc-2021-sitc2021.901 | 2021 | 901 Bacillus Calmette-Guerin can subvert patients antitumor immune response by downregulating HLA-I expression on cancer cells | AGREE | title: 901 Bacillus Calmette-Guerin can subvert patients antitumor immune response by downregulating HLA-I expression on cancer cells; final notes: Q0: NON_ARTICLE -> FILTERED; q1 panel fields empty. | accepted_agree |

| DOI | Year | Title | Audit decision | Evidence (verbatim quote) | Validation status / scope flag |
|---|---|---|---|---|---|
| 10.1136/jitc-2021-sitc2021.920 | 2021 | 920 A single-cell spatially resolved MERFISH map of the colorectal tumor immune microenvironment | AGREE | title: 920 A single-cell spatially resolved MERFISH map of the colorectal tumor immune microenvironment; final notes: Q0: NON_ARTICLE -> FILTERED; q1 panel fields empty. | accepted_agree |
| 10.1136/jitc-2024-009024 | 2024 | Targeting tumor-associated macrophage-derived CD74 improves efficacy of neoadjuvant chemotherapy in combination with PD-1 blockade for cervical cancer | AGREE | paper title (`Targeting tumor-associated macrophage-derived CD74 improves efficacy of neoadjuvant chemotherapy in combination with PD-1 blockade for cervical cancer`); `paper_type=unspecified`; `citation_screening_prior.summary`; empty `gemma_q1_panel_design`; `gemma_final.classification=FILTERED`. | accepted_agree |
| 10.1136/jitc-2024-010871 | 2025 | Immunosuppressive microenvironment of liver restrains chemotherapeutic efficacy in triple-negative breast cancer | AGREE | title: Immunosuppressive microenvironment of liver restrains chemotherapeutic efficacy in triple-negative breast cancer; citation prior labels=NON-ST; q2 spatial=NO; q1 panel-design fields are empty. | accepted_agree |
| 10.1136/jitc-2025-sitc2025.1346 | 2025 | 1346 Molecular and immunologic correlates of response in a phase II study of neoadjuvant lenvatinib plus pembrolizumab in Merkel cell carcinoma | AGREE | title: 1346 Molecular and immunologic correlates of response in a phase II study of neoadjuvant lenvatinib plus pembrolizumab in Merkel cell carcinoma; final notes: Q0: NON_ARTICLE -> FILTERED; q1 panel fields empty. | accepted_agree |
| 10.1136/lupus-2021-lupus21century.4 | 2021 | 104 Distinct spatial profile of inflammatory gene expression in the brain of a mouse model of neuropsychiatric lupus | AGREE | title: 104 Distinct spatial profile of inflammatory gene expression in the brain of a mouse model of neuropsychiatric lupus; final notes: Q0: NON_ARTICLE -> FILTERED; q1 panel fields empty. | accepted_agree |
| 10.1136/lupus-2022-lupus21century.13 | 2022 | 401 Type I interferon in NPSLE | AGREE | title: 401 Type I interferon in NPSLE; final notes: Q0: NON_ARTICLE -> FILTERED; q1 panel fields empty. | accepted_agree |
| 10.1136/lupus-2023-kcr.201 | 2023 | LP-097 Degranulating PR3+ myeloid cells characterize proliferative lupus nephritis | AGREE | Packet fields show either a prefilter/non-primary classification | accepted_agree |

| DOI | Year | Title | Audit decision | Evidence (verbatim quote) | Validation status / scope flag |
|---|---|---|---|---|---|
| | | | | or empty panel-design evidence fields. | |
| 10.1136/lupus-2023-kcr.83 | 2023 | LSO-101 A spatially-resolved single cell resolution atlas of pediatric lupus nephritisuncovers complex interactions between stromal and infiltrating immune cells | AGREE | Packet fields show either a prefilter/non-primary classification or empty panel-design evidence fields. | accepted_agree |
| 10.1136/lupus-2023-lupus21century.76 | 2024 | 1011 Degranulating PR3+ myeloid cells characterize proliferative lupus nephritis | AGREE | Packet fields show either a prefilter/non-primary classification or empty panel-design evidence fields. | accepted_agree |
| 10.1136/thorax-2025-btsabstracts.112 | 2025 | S106 Spatial transcriptomic profiling of airway inflammation and structural changes in asthma | AGREE | Packet fields show either a prefilter/non-primary classification or empty panel-design evidence fields. | accepted_agree |
| 10.1136/thorax-2025-btsabstracts.419 | 2025 | P238 Spatial transcriptomic profiling at single-cell resolution reveals lung responses to sterile injury | AGREE | Packet fields show either a prefilter/non-primary classification or empty panel-design evidence fields. | accepted_agree |
| 10.1139/gen-2017-0046 | 2017 | Characterization of genome in tetraploid StY species of Elymus (Triticeae: Poaceae) using sequential FISH and GISH. | AGREE | Packet fields show either a prefilter/non-primary classification or empty panel-design evidence fields. | accepted_agree |
| 10.1139/gen-2023-0116 | 2024 | Development and application of whole-chromosome painting of chromosomes 7A and 8A of <i>Arachis duranensis</i> based on chromosome-specific single-copy oligonucleotides. | AGREE | Packet fields show either a prefilter/non-primary classification or empty panel-design evidence fields. | accepted_agree |
| 10.1145/3502871.3502881 | 2021 | Informative gene identification for Single-Cell RNA-Seq Data with Mutual Information based Firefly Algorithm | AGREE | gemma_final: Pass-A abstract prefilter -> FILTERED; q1 panel-design fields are empty; q2 paper_type=COMP, spatial=NO, panel_scope=UNCLEAR. | accepted_agree |
| 10.1145/3638569.3638571 | 2023 | A Multiclass Method for Selecting Differentially-Expressed and Cell-Type-Discriminative Genes from scRNA-Seq Data | AGREE | title='A Multiclass Method for Selecting Differentially-Expressed and Cell-Type-Discriminative Genes from scRNA-Seq Data'; citation_screening_prior.labels=''; gemma_final.source='abstract_prefilter' | accepted_agree |
| 10.1145/3698587.3701381 | 2024 | Graph learning of disentangled representation for accurately | AGREE | Packet fields show either a prefilter/non-primary classification | accepted_agree |

| DOI | Year | Title | Audit decision | Evidence (verbatim quote) | Validation status / scope flag |
|---|---|---|---|---|---|
| | | aligning multiple spatial slices | | or empty panel-design evidence fields. | |
| 10.1145/3728424.3760768 | 2025 | PRESTIGE-ST: Patch Resolution and Encoder STrategies for Inference of Gene Expression from Spatial Transcriptomics | AGREE | Title: 'PRESTIGE-ST: Patch Resolution and Encoder STrategies for Inference of Gene Expression from Spatial Transcriptomics'; packet fields show Q0: NON_ARTICLE and Gemma's FILTERED prefilter. | accepted_agree |
| 10.1145/3774976.3774990 | 2025 | Multimodal Fusion of Gene Expression and Alternative Polyadenylation Profiles for Spatial Clustering | AGREE | title: Multimodal Fusion of Gene Expression and Alternative Polyadenylation Profiles for Spatial Clustering; q2 spatial=YES; q2 panel_scope=UNCLEAR; q2 paper_type=COMP; q1 panel-design fields are empty. | accepted_agree |
| 10.1146/annurev-pathmechdis-111523-023417 | 2024 | Challenges and Opportunities in the Clinical Translation of High-Resolution Spatial Transcriptomics | AGREE | Gemma notes say title pre-filter: review/perspective/commentary; q1 panel-design fields are empty. | accepted_agree |
| 10.1158/0008-5472.c.7700640 | 2025 | Data from spatialGE Is a User-Friendly Web Application That Facilitates Spatial Transcriptomics Data Analysis | AGREE | Packet fields show either a prefilter/non-primary classification or empty panel-design evidence fields. | accepted_agree |
| 10.1158/0008-5472.can-24-2346 | 2025 | spatialGE Is a User-Friendly Web Application That Facilitates Spatial Transcriptomics Data Analysis. | AGREE | Packet fields show either a prefilter/non-primary classification or empty panel-design evidence fields. | accepted_agree |
| 10.1158/1078-0432.22444764 | 2023 | Supplementary Figure 2 from A Prospective PCR-Based Screening for the <i>EML4-ALK</i> Oncogene in Non–Small Cell Lung Cancer | AGREE | Packet fields show either a prefilter/non-primary classification or empty panel-design evidence fields. | accepted_agree |
| 10.1158/1078-0432.30028528 | 2025 | Supplementary Table S2 from SARC028 Samples Reveal an Interplay between TGF-β, IFN Signaling, and Low HLA Class I Expression as Contributors to Ewing Sarcoma Checkpoint Blockade Resistance | AGREE | Packet fields show either a prefilter/non-primary classification or empty panel-design evidence fields. | accepted_agree |
| 10.1158/2159-8290.25956905.v1 | 2024 | Supplementary_Table_S4 from Glioblastoma-Infiltrating <i>CD8</i><sup>+</sup> T Cells Are Predominantly a Clonally Expanded | AGREE | Packet fields show either a prefilter/non-primary classification or empty panel-design evidence fields. | accepted_agree |

| DOI | Year | Title | Audit decision | Evidence (verbatim quote) | Validation status / scope flag |
|---|---|---|---|---|---|
| | | <i>GZMK</i><sup>+ </sup> Effector Population | | | |
| 10.1158/2159-8290.30050709 | 2025 | Supplemental Table S9 from Cytotoxic NK Cells Impede Response to Checkpoint Immunotherapy in Melanoma with an Immune-Excluded Phenotype | AGREE | Packet fields show either a prefilter/non-primary classification or empty panel-design evidence fields. | accepted_agree |
| 10.1159/000445859 | 2016 | Heterogeneity of a Constitutional Complex Chromosomal Rearrangement in 2q | AGREE | Packet fields show either a prefilter/non-primary classification or empty panel-design evidence fields. | accepted_agree |
| 10.1159/000508803 | 2021 | Impact of Medium Cut-Off Dialyzers on Patient-Reported Outcomes: COREXH Registry | AGREE | Packet fields show either a prefilter/non-primary classification or empty panel-design evidence fields. | accepted_agree |
| 10.1161/circresaha.125.325795 | 2026 | Advancing Cardiovascular Research With Single-Cell and Spatial Transcriptomics | AGREE | Packet fields show either a prefilter/non-primary classification or empty panel-design evidence fields. | accepted_agree |
| 10.11646/zootaxa.4081.1.1 | 2016 | Revision of the palaearctic species of Eupelmus (Eupelmus) Dalman (Hymenoptera: Chalcidoidea: Eupelmidae) | AGREE | Packet fields show either a prefilter/non-primary classification or empty panel-design evidence fields. | accepted_agree |
| 10.1172/jci153283 | 2021 | Genomic and transcriptomic profiling reveals distinct molecular subsets associated with outcomes in mantle cell lymphoma | AGREE | Title: 'Genomic and transcriptomic profiling reveals distinct molecular subsets associated with outcomes in mantle cell lymphoma'; packet fields show spatial=NO and Gemma's FILTERED prefilter. | accepted_agree |
| 10.1177/01926233251391184 | 2025 | Neuro Omics: Translational Neuropathology Applications of Spatial Technologies From Nonclinical Models to Human Disease | AGREE | title: Neuro Omics: Translational Neuropathology Applications of Spatial Technologies From Nonclinical Models to Human Disease; citation prior labels=REV; q1 panel-design fields are empty; gemma_final=FILTERED. | accepted_agree |
| 10.1177/2472630319868188 | 2019 | Life Sciences Discovery and Technology Highlights | AGREE | Packet fields show either a prefilter/non-primary classification or empty panel-design evidence fields. | accepted_agree |
| 10.1182/blood-2018-99-118412 | 2018 | HDP101, a Novel B-Cell Maturation Antigen (BCMA)-Targeted Antibody Conjugated to α-Amanitin, Is Active Against | AGREE | Packet fields show either a prefilter/non-primary classification or empty panel-design evidence fields. | accepted_agree |

| DOI | Year | Title | Audit decision | Evidence (verbatim quote) | Validation status / scope flag |
|---|---|---|---|---|---|
| | | Myeloma with Preferential Efficacy Against Pre-Clinical Models of Deletion 17p | | | |
| 10.1182/blood-2018-99-118700 | 2018 | Molecular Resistance Mechanisms in Multiple Myeloma | AGREE | Abstract prefilter marked non-spatial. | accepted_agree |
| 10.1182/blood-2022-168191 | 2022 | Spatial-Temporal Multiomic Analysis of Tumor-Immune Interactions in Patients with AML Receiving Pembrolizumab and Decitabine | AGREE | invites applicants for full-time clinical positions in the Leukemia and Benign Hematology services. | accepted_agree |
| 10.1182/blood-2023-185522 | 2023 | Genome-Scale High-Resolution Spatial Mapping of the Pro-Tumorigenic Cellular Niche in Classic Hodgkin Lymphoma | AGREE | Abstract prefilter marked not targeted. | accepted_agree |
| 10.1182/blood-2023-190096 | 2023 | Characterization of Niche Cells and Signals Regulating Stepwise Embryonic Hematopoiesis Using Slide-Seq and Merfish Spatial Transcriptomics | AGREE | The 65th ASH Annual Meeting Abstracts<br>POSTER ABSTRACTS | accepted_agree |
| 10.1182/blood-2024-201171 | 2024 | Spatially Resolved Single-Cell Transcriptomics of T Cells in Follicular Lymphoma Reveals Distinct Immune Ecosystems | AGREE | The 66th ASH Annual Meeting Abstracts | accepted_agree |
| 10.1182/blood-2024-201862 | 2024 | Social Stress Impacts Multicellular Molecular Mechanisms in the Brain Affecting Nociception and Neuroimmune Activity in Sickle Cell Disease | AGREE | The 66th ASH Annual Meeting Abstracts<br>ORAL ABSTRACTS | accepted_agree |
| 10.1182/blood-2024-203052 | 2024 | Tracking Bone Marrow Regeneration after CAR-T Cell Therapy in Myeloma at Spatial Single-Cell Resolution Reveals Determinants of Prolonged Cytopenia | AGREE | POSTER ABSTRACTS | accepted_agree |
| 10.1182/blood-2024-203714 | 2024 | Single-Cell Spatial Transcriptomics Identifies Lymphoid Aggregates with Location-Specific Alterations of Adaptive Immune Cells in Human MDS Bone Marrow | AGREE | The 66th ASH Annual Meeting Abstracts | accepted_agree |
| 10.1182/blood-2024-204183 | 2024 | ICAM4 Mediates Human Erythroid Centered Island Formation | AGREE | The 66th ASH Annual Meeting Abstracts<br>POSTER ABSTRACTS | accepted_agree |
| 10.1182/blood-2024-205264 | 2024 | Development of a Spatial Multiomics Platform to Integrate | AGREE | The 66th ASH Annual Meeting Abstracts | accepted_agree |

| DOI | Year | Title | Audit decision | Evidence (verbatim quote) | Validation status / scope flag |
|---|---|---|---|---|---|
| | | Genomic, Transcriptomic and Proteomic Features for Translational Research in Multiple Myeloma | | | |
| 10.1182/blood-2024-205266 | 2024 | Mapping the Human Bone Marrow in Myeloproliferative Neoplasia Using Spatial Transcriptomics | AGREE | The 66th ASH Annual Meeting Abstracts<br>ORAL ABSTRACTS | accepted_agree |
| 10.1182/blood-2024-208339 | 2024 | Spatial Transcriptomics Characterize Dynamic and Polarized Niche for Embryonic Hematopoiesis at Single-Cell Level | AGREE | The 66th ASH Annual Meeting Abstracts<br>ORAL ABSTRACTS | accepted_agree |
| 10.1182/blood-2024-208450 | 2024 | Characterization of Multicellular Niches Supporting Hematopoietic Stem Cells within Distinct Zones | AGREE | Abstract prefilter marked not targeted. | accepted_agree |
| 10.1182/blood-2024-208838 | 2024 | Integrated Omics to Dissect the Landscape in Chemosensitive and Refractory Extranodal Natural-Killer T-Cell Lymphoma (ENKTCL): A Global Study of the PETAL (PEripheral T-Cell Lymphoma) Consortium | AGREE | The 66th ASH Annual Meeting Abstracts | accepted_agree |
| 10.1182/blood-2024-208880 | 2024 | Biological and Prognostic Subgroups of Plasmablastic Lymphoma Defined By EBV Status and <i>MYC</i> Rearrangement-an L.L.M.P.P. Study | AGREE | The 66th ASH Annual Meeting Abstracts | accepted_agree |
| 10.1182/blood-2024-211058 | 2024 | Bone Marrow Microenvironment Dictates the Progression of Clonal Hematopoiesis through Stromal-Macrophage Cell-Cell Interactions | AGREE | The 66th ASH Annual Meeting Abstracts<br>ORAL ABSTRACTS | accepted_agree |
| 10.1182/blood-2024-211070 | 2024 | Myofibroblast to T-Zone Reticular Cell Ratio Identifies Unicentric Castleman Disease Lymph Nodes | AGREE | The 66th ASH Annual Meeting Abstracts<br>POSTER ABSTRACTS | accepted_agree |
| 10.1182/blood-2024-211509 | 2024 | Case Report: Diagnosis and Management of Myeloid/Lymphoid Neoplasms with <i> <i>ETV6::FLT3</i> </i> Rearrangement | AGREE | Abstract prefilter marked non-spatial. | accepted_agree |
| 10.1182/blood-2024-211658 | 2024 | Unveiling the Tumor Microenvironment in Extranodal NK/T-Cell Lymphoma (ENKTCL) Patients with in Situ Spatial Transcriptomics | AGREE | POSTER ABSTRACTS | accepted_agree |

| DOI | Year | Title | Audit decision | Evidence (verbatim quote) | Validation status / scope flag |
|---|---|---|---|---|---|
| 10.1182/blood-2024-211863 | 2024 | <i>Spatial Characterization of Molecular Pathways Suggests the Dysregulation of LGR5 Expressing Fibroblasts in Patients with sclerodermatous Chronic Graft-Vs-Host Disease</i> | AGREE | POSTER ABSTRACTS | accepted_agree |
| 10.1182/blood-2025-145 | 2025 | Primary central nervous system lymphoma reveals a pro-tumoral brain-specific myeloid landscape. | AGREE | gemma_q0_document_type doc_type=NON_ARTICLE; gemma_final notes/source; empty q1 panel-design fields. | accepted_agree |
| 10.1182/blood-2025-146 | 2025 | Spatial multi-omics reveal distinct immunosuppressive lipid-laden macrophages in primary CNS lymphoma compared to systemic DLBCL | AGREE | gemma_q0_document_type doc_type=NON_ARTICLE; gemma_final notes/source; empty q1 panel-design fields. | accepted_agree |
| 10.1182/blood-2025-2311 | 2025 | Multiplex detection of gastrointestinal microbiota using short probes in graft vs. host disease | AGREE | gemma_q0_document_type doc_type=NON_ARTICLE; title; gemma_final notes/source; empty q1 panel-design fields. | accepted_agree |
| 10.1182/blood-2025-2572 | 2025 | Spatial single-cell multiomics of the bone marrow microenvironment in NPM1-mutant Acute Myeloid Leukemia | AGREE | gemma_q0_document_type doc_type=NON_ARTICLE; gemma_final notes/source; empty q1 panel-design fields. | accepted_agree |
| 10.1182/blood-2025-3151 | 2025 | Spatial transcriptomics identified dynamic and spatial-resolved niche cell and signal architecture for hematopoietic stem cell specification | AGREE | gemma_q0_document_type doc_type=NON_ARTICLE; gemma_final notes/source; empty q1 panel-design fields. | accepted_agree |
| 10.1182/blood-2025-350 | 2025 | Humanized PDX models reveal Lymphoma–Host interactions and enhanced T-cell therapies via epigenetic modulation | AGREE | gemma_q0_document_type doc_type=NON_ARTICLE; gemma_final notes/source; empty q1 panel-design fields. | accepted_agree |
| 10.1182/blood-2025-3527 | 2025 | Single-cell profiling of natural killer T-cell lymphoma reveals stratified immune features and therapeutic strategies | AGREE | gemma_q0_document_type doc_type=NON_ARTICLE; gemma_final notes/source; empty q1 panel-design fields. | accepted_agree |
| 10.1182/blood-2025-3542 | 2025 | Genomic and immune determinants of response to CD3 x CD20 bispecific antibodies in aggressive B-cell lymphoma | AGREE | gemma_q0_document_type doc_type=NON_ARTICLE; gemma_final notes/source; empty q1 panel-design fields. | accepted_agree |
| 10.1182/blood-2025-3552 | 2025 | Multimodal single-cell and spatial analyses reveal a distinct immune ecosystem in pediatric classic Hodgkin lymphoma | AGREE | gemma_q0_document_type doc_type=NON_ARTICLE; gemma_final notes/source; empty q1 panel-design fields. | accepted_agree |

| DOI | Year | Title | Audit decision | Evidence (verbatim quote) | Validation status / scope flag |
|---|---|---|---|---|---|
| 10.1182/blood-2025-3558 | 2025 | Ruxolitinib increases immunogenicity and primes the tumor microenvironment for immune checkpoint blockade in 9p24.1 amplified lymphomas | AGREE | gemma_q0_document_type doc_type=NON_ARTICLE; gemma_final notes/source; empty q1 panel-design fields. | accepted_agree |
| 10.1182/blood-2025-3761 | 2025 | Spatial transcriptomic dissection of the myeloproliferative neoplasm niche uncovers mutant clone interactions and microenvironmental rewiring | AGREE | gemma_q0_document_type doc_type=NON_ARTICLE; gemma_final notes/source; empty q1 panel-design fields. | accepted_agree |
| 10.1182/blood-2025-3949 | 2025 | Spatially resolved transcriptomics reveals immunosuppressive niches and clonal diversity in extramedullary disease in multiple myeloma refractory to immunotherapy | AGREE | gemma_q0_document_type doc_type=NON_ARTICLE; gemma_final notes/source; empty q1 panel-design fields. | accepted_agree |
| 10.1182/blood-2025-462 | 2025 | Distinct blood endothelial cells shape spatial niches in angioimmunoblastic T-cell lymphoma | AGREE | gemma_q0_document_type doc_type=NON_ARTICLE; gemma_final notes/source; empty q1 panel-design fields. | accepted_agree |
| 10.1182/blood-2025-4955 | 2025 | Cellular senescence associated with aging and chemotherapy exposure primes bone marrow for immune dysfunction and myeloma persistence | AGREE | gemma_q0_document_type doc_type=NON_ARTICLE; gemma_final notes/source; empty q1 panel-design fields. | accepted_agree |
| 10.1182/blood-2025-5730 | 2025 | Evolution of t(11;14) biology in multiple myeloma and it's role on outcomes to venetoclax treatment | AGREE | paper title; gemma_final notes/source; q2 paper_type=EXP with panel_scope=UNCLEAR; empty q1 panel-design fields. | accepted_agree |
| 10.1182/blood-2025-5804 | 2025 | Spatial biomarkers of CAR-T treatment response in relapsed refractory multiple myeloma | AGREE | gemma_q0_document_type doc_type=NON_ARTICLE; gemma_final notes/source; empty q1 panel-design fields. | accepted_agree |
| 10.1182/blood-2025-5864 | 2025 | Spatial communication modules identify prognostic cell-cell signaling hubs in chronic graft-versus-host disease | AGREE | gemma_q0_document_type doc_type=NON_ARTICLE; gemma_final notes/source; empty q1 panel-design fields. | accepted_agree |
| 10.1182/blood-2025-588 | 2025 | CSF-1R+ macrophages orchestrate human skin chronic graft-versus-host disease | AGREE | gemma_q0_document_type doc_type=NON_ARTICLE; gemma_final notes/source; empty q1 panel-design fields. | accepted_agree |
| 10.1182/blood-2025-860 | 2025 | Multi-omics single-cell spatial profiling of paediatric and adult acute myeloid leukaemia links | AGREE | gemma_q0_document_type doc_type=NON_ARTICLE; | accepted_agree |

| DOI | Year | Title | Audit decision | Evidence (verbatim quote) | Validation status / scope flag |
|---|---|---|---|---|---|
| | | spatial organization, molecular genetics and microenvironmental interactions | | gemma_final notes/source; empty q1 panel-design fields. | |
| 10.1182/blood.v130.suppl_1.4335.4335 | 2017 | Overexpression of EZH2 in Multiple Myeloma Is Associated with Poor Prognosis Regardless of Treatment with Novel Agents or High-Dose Chemotherapy | AGREE | paper title; gemma_final notes/source; q2 paper_type=EXP with panel_scope=UNCLEAR; empty q1 panel-design fields. | accepted_agree |
| 10.1183/13993003.congress-2019.pa1333 | 2019 | STARMAP: an observational study to assess disease-relevant outcomes using home-monitoring devices in patients with idiopathic pulmonary fibrosis (IPF) | AGREE | paper.title='STARMAP: an observational study to assess disease-relevant outcomes using home-monitoring devices in patients with idiopathic pulmonary fibrosis (IPF)'; gemma_final='Pass-A abstract: non-spatial -> FILTERED'; gemma_q2_spatial=NO | accepted_agree |
| 10.1183/23120541.00561-2022 | 2023 | ERS International Congress 2022: highlights from the Basic and Translational Science Assembly | AGREE | title='ERS International Congress 2022: highlights from the Basic and Translational Science Assembly'; citation_screening_prior.labels='REV'; gemma_final.source='citation_screening_prefilter' | accepted_agree |
| 10.1186/s12859-020-03625-z | 202 | A robust nonlinear low-dimensional manifold for single cell RNA-seq data | AGREE | Title: 'A robust nonlinear low-dimensional manifold for single cell RNA-seq data'; packet fields show paper_type=COMP and Gemma's FILTERED prefilter. | accepted_agree |
| 10.1186/s12859-020-03641-z | 202 | A rank-based marker selection method for high throughput scRNA-seq data | AGREE | Title: 'A rank-based marker selection method for high throughput scRNA-seq data'; packet fields show paper_type=COMP-BENCH and Gemma's FILTERED prefilter. | accepted_agree |
| 10.1186/s12859-021-04302-5 | 2021 | Matisse: a MATLAB-based analysis toolbox for in situ sequencing expression maps. | AGREE | Here we present MATLAB-based Analysis toolbox for in situ sequencing (ISS) expression maps (Matisse). | accepted_agree |
| 10.1186/s12859-022-04817-5 | 2022 | Detection of cell markers from single cell RNA-seq with sc2marker | AGREE | title='Detection of cell markers from single cell RNA-seq with sc2marker'; citation_screening_prior.labels='COMP-BENCH'; | accepted_agree |

| DOI | Year | Title | Audit decision | Evidence (verbatim quote) | Validation status / scope flag |
|---|---|---|---|---|---|
| | | | | gemma_final.source='citation_screening_prefilter' | |
| 10.1186/s12859-022-04860-2 | 2022 | SMaSH: a scalable, general marker gene identification framework for single-cell RNA-sequencing | AGREE | citation_screening_prior: computational-only prior; gemma_final source=citation_screening_prefilter -> FILTERED; q1 panel-design fields are empty; q2 paper_type=COMP. | accepted_agree |
| 10.1186/s12859-024-05656-2 | 2024 | InClust+: the deep generative framework with mask modules for multimodal data integration, imputation, and cross-modal generation. | NEEDS_MANUAL | Final manual decision: FILTERED because the paper is a downstream analysis/modeling/QC method or dataset-use paper, with no final panel-design source. Original evidence: q1: panel_design=UNCLEAR; q2: MERFISH/scRNA integration context only. | manual_final_filter_after_pdf_review |
| 10.1186/s12859-025-06099-z | 2025 | TransST: transfer learning embedded spatial factor modeling of spatial transcriptomics data | AGREE | paper.title='TransST: transfer learning embedded spatial factor modeling of spatial transcriptomics data'; gemma_final='Pass-A abstract: COMP -> FILTERED'; gemma_q2_paper_profile=spatial YES, scope UNCLEAR | accepted_agree |
| 10.1186/s12859-025-06112-5 | 2025 | Redefining the high variable genes by optimized LOESS regression with positive ratio | AGREE | title: Redefining the high variable genes by optimized LOESS regression with positive ratio; citation prior labels=COMP-BENCH; q2 paper_type=COMP; q1 panel-design fields are empty; gemma_final=FILTERED. | accepted_agree |
| 10.1186/s12859-025-06243-9 | 2025 | Winnow-KAN: single-cell RNA-seq location recovery with small-gene-set spatial transcriptomics | AGREE | paper.title='Winnow-KAN: single-cell RNA-seq location recovery with small-gene-set spatial transcriptomics'; gemma_final='Pass-A abstract: COMP -> FILTERED'; gemma_q2_paper_profile=spatial YES, scope UNCLEAR | accepted_agree |
| 10.1186/s12864-016-3231-z | 2016 | Fluorescence chromosome banding and FISH mapping in perennial ryegrass, Lolium perenne L. | AGREE | paper.title='Fluorescence chromosome banding and FISH mapping in perennial ryegrass, Lolium perenne L.'; gemma_final='Pass-A abstract: | accepted_agree |

| DOI | Year | Title | Audit decision | Evidence (verbatim quote) | Validation status / scope flag |
|---|---|---|---|---|---|
| | | | | non-spatial -> FILTERED'; gemma_q2_spatial=NO | |
| 10.1186/s12864-024-10434-8 | 2024 | Spatial transcriptomic brain imaging reveals the effects of immunomodulation therapy on specific regional brain cells in a mouse dementia model | AGREE | Q2: not a targeted panel-design paper -> FILTERED | accepted_agree |
| 10.1186/s12864-024-10469-x | 2024 | HyperGCN: an effective deep representation learning framework for the integrative analysis of spatial transcriptomics data. | AGREE | paper.title='HyperGCN: an effective deep representation learning framework for the integrative analysis of spatial transcriptomics data.'; gemma_final='Pass-A abstract: COMP -> FILTERED'; gemma_q2_paper_profile=spatial YES, scope UNCLEAR | accepted_agree |
| 10.1186/s12864-024-11030-6 | 2024 | Cross-species single-cell analysis reveals divergence and conservation of peripheral blood mononuclear cells | AGREE | paper title (`Cross-species single-cell analysis reveals divergence and conservation of peripheral blood mononuclear cells`); `paper_type=unspecified`; `citation_screening_prior.summary`; empty `gemma_q1_panel_design`; `gemma_final.classification=FILTERED`. | accepted_agree |
| 10.1186/s12864-024-11072-w | 2024 | Deciphering spatial domains from spatially resolved transcriptomics through spatially regularized deep graph networks | AGREE | paper title (`Deciphering spatial domains from spatially resolved transcriptomics through spatially regularized deep graph networks`); `paper_type=COMP`; `citation_screening_prior.summary`; empty `gemma_q1_panel_design`; `gemma_final.classification=FILTERED`. | accepted_agree |
| 10.1186/s12864-025-11319-0 | 2025 | scMFG: a single-cell multi-omics integration method based on feature grouping | AGREE | title: scMFG: a single-cell multi-omics integration method based on feature grouping; citation prior labels=COMP-BENCH; q2 paper_type=COMP; q1 panel-design fields are empty; gemma_final=FILTERED. | accepted_agree |
| 10.1186/s12864-025-11818-0 | 2025 | NuclampFISH enables cell sorting based on nuclear RNA expression for chromatin analysis. | DISAGREE | Final manual decision: FILTERED because the available evidence concerns probe/barcode/oligo | manual_final_filter_after_pdf_review |

| DOI | Year | Title | Audit decision | Evidence (verbatim quote) | Validation status / scope flag |
|---|---|---|---|---|---|
| | | | | design or single-gene detection, not final gene-panel selection. Original evidence: q1 says EEF2 mRNA was labeled with smFISH, clampFISH, and HCR; no broader panel-selection rationale is provided. | |
| 10.1186/s12870-022-03496-x | 2022 | Molecular cytogenetics and development of St-chromosome-specific molecular markers of novel stripe rust resistant wheat-Thinopyrum intermedium and wheat-Thinopyrum ponticum substitution lines. | AGREE | paper.title='Molecular cytogenetics and development of St-chromosome-specific molecular markers of novel stripe rust resistant wheat-Thinopyrum intermedium and wheat-Thinopyrum ponticum substitution lines.'; gemma_final='Pass-A abstract: non-spatial -> FILTERED'; gemma_q2_spatial=NO | accepted_agree |
| 10.1186/s12883-026-04720-z | 2026 | Novel VSMC-associated biomarkers in intracranial aneurysm pathogenesis: a multi-omics and machine learning study | AGREE | citation_screening_prior labels=COMP and all_tools=COSG; gemma_final source=citation_screening_prefilter -> FILTERED; q1 panel-design fields are empty. | accepted_agree |
| 10.1186/s12900-016-0055-7 | 2016 | Structuprint: a scalable and extensible tool for two-dimensional representation of protein surfaces. | AGREE | paper.title='Structuprint: a scalable and extensible tool for two-dimensional representation of protein surfaces.'; gemma_final='Pass-A abstract: COMP -> FILTERED'; gemma_q2_paper_profile=spatial NO, scope UNCLEAR | accepted_agree |
| 10.1186/s12918-015-0177-8 | 2015 | Systems biology surveillance decrypts pathological transcriptome remodeling. | AGREE | paper.title='Systems biology surveillance decrypts pathological transcriptome remodeling.'; gemma_final='Pass-A abstract: non-spatial -> FILTERED'; gemma_q2_spatial=NO | accepted_agree |
| 10.1186/s12967-023-03961-7 | 2023 | Abnormal signal pathways and tumor heterogeneity in osteosarcoma | AGREE | title='Abnormal signal pathways and tumor heterogeneity in osteosarcoma'; citation_screening_prior.labels='COMP'; gemma_final.source='citation_screening_prefilter' | accepted_agree |

| DOI | Year | Title | Audit decision | Evidence (verbatim quote) | Validation status / scope flag |
|---|---|---|---|---|---|
| 10.1186/s12967-023-04151-1 | 2023 | The potential crosstalk between tumor and plasma cells and its association with clinical outcome and immunotherapy response in bladder cancer | NEEDS_MANUAL | panel_scope=NOT_TARGETED with no target-gene panel source evidence; untargeted/no-panel records are out of scope. Original audit evidence: No panel-design evidence is present in the packet. | validated_high_risk_manual_filtered |
| 10.1186/s12967-025-06150-w | 2025 | Single-cell profiling reveals a reduced epithelial defense system, decreased immune responses and the immune regulatory roles of different fibroblast subpopulations in chronic atrophic gastritis | AGREE | title: Single-cell profiling reveals a reduced epithelial defense system, decreased immune responses and the immune regulatory roles of different fibroblast subpopulations in chronic atrophic gastritis; citation prior labels=NON-ST; q2 spatial=NO; q1 panel-design fields are empty. | accepted_agree |
| 10.1186/s12967-025-06682-1 | 2025 | Pathway-based cancer transcriptome deciphers a high-resolution intrinsic heterogeneity within bladder cancer classification | AGREE | title: Pathway-based cancer transcriptome deciphers a high-resolution intrinsic heterogeneity within bladder cancer classification; citation prior labels=NON-ST; q2 spatial=NO; q1 panel-design fields are empty. | accepted_agree |
| 10.1186/s13058-023-01703-7 | 2023 | Single-cell RNA reveals a tumorigenic microenvironment in the interface zone of human breast tumors | AGREE | title='Single-cell RNA reveals a tumorigenic microenvironment in the interface zone of human breast tumors'; citation_screening_prior.labels='NON-ST'; gemma_final.source='citation_screening_prefilter' | accepted_agree |
| 10.1186/s13059-021-02544-3 | 2021 | Feature selection revisited in the single-cell era | AGREE | Citation prior labels the item REV; title is 'Feature selection revisited in the single-cell era'; q1 panel-design fields are empty. | accepted_agree |
| 10.1186/s13059-021-02548-z | 2021 | geneBasis: an iterative approach for unsupervised selection of targeted gene panels from scRNA-seq | AGREE | Title is 'geneBasis: an iterative approach for unsupervised selection of targeted gene panels from scRNA-seq'; citation prior marks it computational-only and q1 panel-design fields are empty. | accepted_agree |
| 10.1186/s13059-023-02904-1 | 2023 | The shaky foundations of simulating single-cell RNA sequencing data | AGREE | Title only; paper_type COMP; spatial blank; panel_scope blank; q1 panel-design fields are empty; gemma_final=FILTERED from citation-screening prefilter. | accepted_agree |

| DOI | Year | Title | Audit decision | Evidence (verbatim quote) | Validation status / scope flag |
|---|---|---|---|---|---|
| 10.1186/s13059-023-03080-y | 2023 | Dual spatially resolved transcriptomics for human host-pathogen colocalization studies in FFPE tissue sections. | AGREE | paper.title='Dual spatially resolved transcriptomics for human host-pathogen colocalization studies in FFPE tissue sections.'; gemma_final='Pass-A abstract: COMP -> FILTERED'; gemma_q2_paper_profile=spatial YES, scope NOT_TARGETED | accepted_agree |
| 10.1186/s13059-023-03123-4 | 2023 | Challenges and opportunities to computationally deconvolve heterogeneous tissue with varying cell sizes using single-cell RNA-sequencing datasets | AGREE | Title only; gemma_final notes title pre-filter: review/perspective/commentary; q1 panel-design fields are empty. | accepted_agree |
| 10.1186/s13059-024-03174-1 | 2024 | Gene panel selection for targeted spatial transcriptomics. | AGREE | paper.title='Gene panel selection for targeted spatial transcriptomics.'; citation_screening_prior.summary=labels=['COMP', 'COMP-BENCH']; exp_used_tools=['(none)']; all_tools=['NS-Forest v2', 'RankCorr', 'SCMER', 'geneBasis', 'scGeneFit']; gemma_final='Pass-A abstract: computational-only prior -> FILTERED' | accepted_agree |
| 10.1186/s13059-024-03183-0 | 2024 | A comparison of marker gene selection methods for single-cell RNA sequencing data | AGREE | Title is 'A comparison of marker gene selection methods for single-cell RNA sequencing data'; citation prior marks it computational-only and q1 panel-design fields are empty. | accepted_agree |
| 10.1186/s13059-024-03185-y | 2024 | scGIST: gene panel design for spatial transcriptomics with prioritized gene sets | AGREE | title: scGIST: gene panel design for spatial transcriptomics with prioritized gene sets; q2 paper_type=COMP; q1 panel-design fields are empty; citation prior labels=COMP\|COMP-BENCH and Gemma_final=FILTERED. | accepted_agree |
| 10.1186/s13059-024-03217-7 | 2024 | Bento: a toolkit for subcellular analysis of spatial transcriptomics data. | AGREE | paper.title='Bento: a toolkit for subcellular analysis of spatial transcriptomics data.'; gemma_final='Pass-A abstract: COMP -> FILTERED'; gemma_q2_paper_profile=spatial YES, scope UNCLEAR | accepted_agree |
| 10.1186/s13059-024-03227-5 | 2024 | Merit of integrating in situ transcriptomics and anatomical | NEEDS_MANUAL | 'Mathematical algorithm-based reconstruction using scRNA-seq | pdf_validated_unclear_update |

| DOI | Year | Title | Audit decision | Evidence (verbatim quote) | Validation status / scope flag |
|---|---|---|---|---|---|
| | | information for cell annotation and lineage construction in single-cell analyses of Populus | | data' and lcmRNA-seq/in situ transcriptomes, not a targeted panel study | |
| 10.1186/s13059-024-03228-4 | 2024 | Response to: “Merit of integrating in situ transcriptomics and anatomical information for cell annotation and lineage construction in single-cell analyses of Populus” | AGREE | paper_type/doc_type evidence: EDITORIAL (CORRESPONDENCE Liang et al. Genome Biology (2024) 25:86); gemma_final notes: Q0: EDITORIAL → FILTERED | accepted_agree |
| 10.1186/s13059-024-03284-w | 2024 | scCDC: a computational method for gene-specific contamination detection and correction in single-cell and single-nucleus RNA-seq data | AGREE | paper title (`scCDC: a computational method for gene-specific contamination detection and correction in single-cell and single-nucleus RNA-seq data`); `paper_type=COMP`; `citation_screening_prior.summary`; empty `gemma_q1_panel_design`; `gemma_final.classification=FILTERED`. | accepted_agree |
| 10.1186/s13059-024-03303-w | 2024 | Gene count normalization in single-cell imaging-based spatially resolved transcriptomics | AGREE | paper title (`Gene count normalization in single-cell imaging-based spatially resolved transcriptomics`); `paper_type=COMP`; `citation_screening_prior.summary`; empty `gemma_q1_panel_design`; `gemma_final.classification=FILTERED`. | accepted_agree |
| 10.1186/s13059-024-03340-5 | 2024 | Microsatellite instability at U2AF-binding polypyrimidic tract sites perturbs alternative splicing during colorectal cancer initiation | AGREE | Title only; paper_type blank; spatial=NO; panel_scope blank; q1 panel-design fields are empty; gemma_final=FILTERED from citation-screening prefilter. | accepted_agree |
| 10.1186/s13059-024-03396-3 | 2024 | Characterization of regeneration initiating cells during Xenopus laevis tail regeneration | NEEDS_MANUAL | panel_scope=NOT_TARGETED with no target-gene panel source evidence; untargeted/no-panel records are out of scope. Original audit evidence: No panel-design evidence is present in the packet. | validated_high_risk_manual_filtered |
| 10.1186/s13059-024-03410-8 | 2024 | scCTS: identifying the cell type-specific marker genes from population-level single-cell RNA-seq | AGREE | citation_screening_prior: computational-only prior; gemma_final source=citation_screening_prefilter -> FILTERED; q1 panel-design | accepted_agree |

| DOI | Year | Title | Audit decision | Evidence (verbatim quote) | Validation status / scope flag |
|---|---|---|---|---|---|
| | | | | fields are empty; q2 paper_type=COMP. | |
| 10.1186/s13059-025-03747-8 | 2025 | Cross-expression meta-analysis of mouse brain slices reveals coordinated gene expression across spatially adjacent cells | AGREE | paper.title='Cross-expression meta-analysis of mouse brain slices reveals coordinated gene expression across spatially adjacent cells'; gemma_final='Pass-A abstract: COMP -> FILTERED'; gemma_q2_paper_profile=spatial YES, scope UNCLEAR | accepted_agree |
| 10.1186/s13059-025-03810-4 | 2025 | A novel gene expression stability metric to unveil homeostasis and regulation | AGREE | title: A novel gene expression stability metric to unveil homeostasis and regulation; citation prior labels=COMP; q2 paper_type=COMP; q1 panel-design fields are empty; gemma_final=FILTERED. | accepted_agree |
| 10.1186/s13059-025-03846-6 | 2026 | Neighborhood nonnegative matrix factorization identifies patterns and spatially-variable genes in large-scale spatial transcriptomics data | NEEDS_MANUAL | Final manual decision: FILTERED because the paper is a downstream analysis/modeling/QC method or dataset-use paper, with no final panel-design source. Original evidence: q1: panel_design=UNCLEAR; q2: panel_scope=UNCLEAR. | manual_final_filter_after_pdf_review |
| 10.1186/s13059-025-03908-9 | 2026 | RNA2seg: a generalist model for cell segmentation in image-based spatial transcriptomics. | AGREE | paper.title='RNA2seg: a generalist model for cell segmentation in image-based spatial transcriptomics.'; gemma_final='Pass-A abstract: COMP -> FILTERED'; gemma_q2_paper_profile=spatial YES, scope UNCLEAR | accepted_agree |
| 10.1186/s13059-026-03998-z | 2026 | Benchmarking LLM-based agents for single-cell omics analysis | AGREE | gemma_final notes say Pass-A abstract: non-spatial -> FILTERED; q2 paper_type=COMP, spatial=NO, panel_scope=UNCLEAR; q1 panel-design fields are empty. | accepted_agree |
| 10.1186/s13073-024-01282-y | 2024 | Spatial multi-omics: novel tools to study the complexity of cardiovascular diseases | AGREE | citation_screening_prior: labels=REV; gemma_final source=citation_screening_prefilter -> FILTERED; q1 panel-design fields are empty. | accepted_agree |

| DOI | Year | Title | Audit decision | Evidence (verbatim quote) | Validation status / scope flag |
|---|---|---|---|---|---|
| 10.1186/s13148-024-01800-0 | 2024 | Single-cell transcriptomics reveal the prognostic roles of epithelial and T cells and DNA methylation-based prognostic models in pancreatic cancer | AGREE | Title only; paper_type blank; spatial=NO; panel_scope blank; q1 panel-design fields are empty; gemma_final=FILTERED from citation-screening prefilter. | accepted_agree |
| 10.1186/s13148-025-01930-z | 2025 | Multiple highly methylated CpG sites as potential epigenetic markers for the diagnosis of prostate cancer. | AGREE | paper.title='Multiple highly methylated CpG sites as potential epigenetic markers for the diagnosis of prostate cancer.'; gemma_final='Pass-A abstract: not targeted -> FILTERED'; gemma_q2_paper_profile=spatial YES, scope NOT_TARGETED | accepted_agree |
| 10.1186/s40168-019-0724-4 | 2019 | Reproducible changes in the gut microbiome suggest a shift in microbial and host metabolism during spaceflight | AGREE | paper.title='Reproducible changes in the gut microbiome suggest a shift in microbial and host metabolism during spaceflight'; gemma_final='Pass-A abstract: non-spatial -> FILTERED'; gemma_q2_spatial=NO | accepted_agree |
| 10.1186/s40478-025-02038-4 | 2025 | Distinct characteristics of brain metastasis in lung adenocarcinoma: development of high-confidence cell lines | AGREE | title: Distinct characteristics of brain metastasis in lung adenocarcinoma: development of high-confidence cell lines; citation prior labels=NON-ST; q2 spatial=NO; q1 panel-design fields are empty. | accepted_agree |
| 10.1186/s40793-022-00400-w | 2022 | In-situ sequencing reveals the effect of storage on lacustrine sediment microbiome demographics and functionality. | AGREE | paper.title='In-situ sequencing reveals the effect of storage on lacustrine sediment microbiome demographics and functionality.'; gemma_final='Pass-A abstract: non-spatial -> FILTERED'; gemma_q2_spatial=NO | accepted_agree |
| 10.1186/s41065-025-00528-y | 2025 | Global trends in machine learning applications for single-cell transcriptomics research | AGREE | title: Global trends in machine learning applications for single-cell transcriptomics research; q0 doc_type=REVIEW; q1 panel-design fields are empty; citation prior labels=REV and Gemma_final=FILTERED. | accepted_agree |
| 10.1186/s43046-025-00312-z | 2025 | Dual-modality treatment using gamma radiation and ZnO nanoparticles: effects on normal and malignant lung cells | AGREE | title: Dual-modality treatment using gamma radiation and ZnO nanoparticles: effects on normal and malignant lung cells; citation prior labels=NON-ST; q2 | accepted_agree |

| DOI | Year | Title | Audit decision | Evidence (verbatim quote) | Validation status / scope flag |
|---|---|---|---|---|---|
| | | | | spatial=NO; q1 panel-design fields are empty. | |
| 10.1186/s44330-024-00015-2 | 2024 | Discovery of optimal cell type classification marker genes from single cell RNA sequencing data | AGREE | title: Discovery of optimal cell type classification marker genes from single cell RNA sequencing data; q2 paper_type=COMP; q1 panel-design fields are empty; citation prior labels=COMP-BENCH and Gemma_final=FILTERED. | accepted_agree |
| 10.1186/s44330-026-00055-w | 2026 | Next generation statistical framework for next generation spatial transcriptomics data | AGREE | paper.title='Next generation statistical framework for next generation spatial transcriptomics data'; gemma_final='Pass-A abstract: COMP -> FILTERED'; gemma_q2_paper_profile=spatial YES, scope UNCLEAR | accepted_agree |
| 10.1200/jco.2024.42.16_suppl.e23546 | 2024 | Effect of molecular cartography on recurrence-specific drivers through analysis of matched primary and recurrent rhabdomyosarcoma tumors. | AGREE | paper.title='Effect of molecular cartography on recurrence-specific drivers through analysis of matched primary and recurrent rhabdomyosarcoma tumors.'; gemma_final='Pass-A abstract: non-spatial -> FILTERED'; gemma_q2_spatial=NO | accepted_agree |
| 10.1210/jendso/bvad131 | 2023 | In Situ Spatial Reconstruction of Distinct Normal and Pathological Cell Populations Within the Human Adrenal Gland. | AGREE | The packet's paper profile is NOT_TARGETED and contains no panel-design evidence. | accepted_agree |
| 10.1210/jendso/bvae163.2226 | 2024 | 8773 CD44 Variant Isoforms Expressed Within PAX7/SOX2-Positive Pituitary Neuroendocrine Tumors Attribute to Therapy Resistance and Recurrence of Cushing’s Disease | AGREE | paper.title='8773 CD44 Variant Isoforms Expressed Within PAX7/SOX2-Positive Pituitary Neuroendocrine Tumors Attribute to Therapy Resistance and Recurrence of Cushing's Disease'; gemma_final='Q0: NON_ARTICLE -> FILTERED' | accepted_agree |
| 10.1210/jendso/bvae163.2394 | 2024 | SAT-075 Revealing the Heterogeneity and Behavior of Pituitary Neuroendocrine Tumors in Cushing’s Disease Patients using Spatial Multiomics | AGREE | paper.title='SAT-075 Revealing the Heterogeneity and Behavior of Pituitary Neuroendocrine Tumors in Cushing's Disease Patients using Spatial Multiomics'; gemma_final='Q0: NON_ARTICLE -> FILTERED' | accepted_agree |
| 10.1210/jendso/bvaf149.1842 | 2025 | MON-264 Molecular Mechanisms of Progesterone Receptor in Inguinal Hernia Formation via | AGREE | title: MON-264 Molecular Mechanisms of Progesterone Receptor in Inguinal Hernia | accepted_agree |

| DOI | Year | Title | Audit decision | Evidence (verbatim quote) | Validation status / scope flag |
|---|---|---|---|---|---|
| | | Skeletal Muscle Fibrosis and Atrophy | | Formation via Skeletal Muscle Fibrosis and Atrophy; q0 doc_type: NON_ARTICLE; final notes: Q0: NON_ARTICLE → FILTERED | |
| 10.1210/jendso/bvaf149.1847 | 2025 | MON-271 Vitamin D Receptor with Calcium Sensing Receptor Control Epidermal Tumorigenesis Through Oxidative Stress And DNA Repair Genes With Tumor Suppressor P53 Signaling | AGREE | title: MON-271 Vitamin D Receptor with Calcium Sensing Receptor Control Epidermal Tumorigenesis Through Oxidative Stress And DNA Repair Genes With Tumor Suppressor P53 Signaling; q0 doc_type: NON_ARTICLE; final notes: Q0: NON_ARTICLE → FILTERED | accepted_agree |
| 10.1210/jendso/bvaf149.482 | 2025 | MON-737 Use of scRNA seq. and Xenium In Situ to determine the Cellular and Molecular Impact of Abaloparatide on the Distinct sites of Fracture healing | AGREE | title: MON-737 Use of scRNA seq. and Xenium In Situ to determine the Cellular and Molecular Impact of Abaloparatide on the Distinct sites of Fracture healing; q0 doc_type: NON_ARTICLE; final notes: Q0: NON_ARTICLE → FILTERED | accepted_agree |
| 10.1227/neu.0000000000003360_494 | 2025 | 494 Multi-Omics Integration and Mapping of Invasion Markers Reveal Tumor-Specific Druggable Targets in Primary Human Glioblastoma | AGREE | title: 494 Multi-Omics Integration and Mapping of Invasion Markers Reveal Tumor-Specific Druggable Targets in Primary Human Glioblastoma; q1 panel fields empty; q2 profile empty; final notes: Pass-A abstract: not targeted → FILTERED | accepted_agree |
| 10.1242/bio.058990 | 2022 | Modulation of retinoid-X-receptors differentially regulates expression of apolipoprotein genes apoc1 and apoeb by zebrafish microglia. | AGREE | title: Modulation of retinoid-X-receptors differentially regulates expression of apolipoprotein genes apoc1 and apoeb by zebrafish microglia.; q1 panel fields empty; q2 profile empty; final notes: Pass-A abstract: non-spatial → FILTERED | accepted_agree |
| 10.1242/bio.059147 | 2022 | Tissue-specific transcriptome profiles identify functional differences key to understanding whole plant response to life in variable salinity | AGREE | title: Tissue-specific transcriptome profiles identify functional differences key to understanding whole plant response to life in variable salinity; q1 panel fields empty; q2 profile empty; final notes: Pass-A abstract: non-spatial → FILTERED | accepted_agree |

| DOI | Year | Title | Audit decision | Evidence (verbatim quote) | Validation status / scope flag |
|---|---|---|---|---|---|
| 10.1242/dev.138560 | 2016 | Single-molecule RNA detection at depth by hybridization chain reaction and tissue hydrogel embedding and clearing. | AGREE | title: Single-molecule RNA detection at depth by hybridization chain reaction and tissue hydrogel embedding and clearing.; q1 panel fields empty; q2 profile empty; final notes: Pass-A abstract: COMP → FILTERED | accepted_agree |
| 10.1242/dev.165753 | 2018 | Third-generation <i>in situ</i> hybridization chain reaction: multiplexed, quantitative, sensitive, versatile, robust. | AGREE | title: Third-generation <i>in situ</i> hybridization chain reaction: multiplexed, quantitative, sensitive, versatile, robust.; q1 panel fields empty; q2 profile empty; final notes: Pass-A abstract: COMP → FILTERED | accepted_agree |
| 10.1242/dev.199847 | 2021 | Hybridization chain reaction enables a unified approach to multiplexed, quantitative, high-resolution immunohistochemistry and in situ hybridization. | AGREE | title: Hybridization chain reaction enables a unified approach to multiplexed, quantitative, high-resolution immunohistochemistry and in situ hybridization.; q1 panel fields empty; q2 profile empty; final notes: Pass-A abstract: COMP → FILTERED | accepted_agree |
| 10.1242/dev.202429 | 2024 | BirthSeq, a new method to isolate and analyze dated cells in different vertebrates | AGREE | title: BirthSeq, a new method to isolate and analyze dated cells in different vertebrates; q1 panel fields empty; q2 profile empty; final notes: Pass-A abstract: COMP → FILTERED | accepted_agree |
| 10.1242/dev.204775 | 2025 | One probe fits all: a highly customizable modular RNA in situ hybridization platform expanding the application of SABER DNA probes. | AGREE | title: One probe fits all: a highly customizable modular RNA in situ hybridization platform expanding the application of SABER DNA probes.; q1 panel fields empty; q2 profile empty; final notes: Pass-A abstract: COMP → FILTERED | accepted_agree |
| 10.1242/dev.204822 | 2025 | In preprints: unfolding the spatial genomics frontier of mouse gastrulation | AGREE | title: In preprints: unfolding the spatial genomics frontier of mouse gastrulation; q1 panel fields empty; q2 profile empty; final notes: Pass-A abstract: OTHER → FILTERED | accepted_agree |
| 10.1261/rna.079239.122 | 2022 | High-throughput imaging of mRNA at the single-cell level in human primary immune cells. | AGREE | title: High-throughput imaging of mRNA at the single-cell level in human primary immune cells.; q1 panel fields empty; q2 profile empty; final notes: Pass-A abstract: COMP → FILTERED | accepted_agree |

| DOI | Year | Title | Audit decision | Evidence (verbatim quote) | Validation status / scope flag |
|---|---|---|---|---|---|
| 10.12688/f1000research.122775.1 | 2022 | Spatial transcriptomics dimensionality reduction using wavelet bases | AGREE | title='Spatial transcriptomics dimensionality reduction using wavelet bases'; citation_screening_prior.labels='COMP'; gemma_final.source='citation_screening_prefilter' | accepted_agree |
| 10.13523/j.cb.2304056 | 2023 | Analysis of Patent Trends for the Spatial Omics Technology | AGREE | title: Analysis of Patent Trends for the Spatial Omics Technology; q1 panel fields empty; q2 profile empty; final notes: Pass-A abstract: OTHER → FILTERED | accepted_agree |
| 10.13523/j.cb.2312103 | 2024 | Recent Advances in Spatiotemporal Omics Technology | AGREE | title: Recent Advances in Spatiotemporal Omics Technology; q1 panel fields empty; q2 profile empty; final notes: Pass-A abstract: OTHER → FILTERED | accepted_agree |
| 10.1360/ssv-2019-0224 | 202 | The “toolbox” for chromatin conformation capture | AGREE | title: The “toolbox” for chromatin conformation capture; q1 panel fields empty; q2 profile empty; final notes: Pass-A abstract: OTHER → FILTERED | accepted_agree |
| 10.1360/ssv-2025-0140 | 2025 | Current progress of imaging-based 3D spatial transcriptomics | AGREE | title: Current progress of imaging-based 3D spatial transcriptomics; q1 panel fields empty; q2 profile empty; final notes: Pass-A abstract: OTHER → FILTERED | accepted_agree |
| 10.1360/tb-2025-0014 | 2025 | Advances in rolling circle amplification-based in situ sequencing for spatial transcriptomics technologies | AGREE | title: Advances in rolling circle amplification-based in situ sequencing for spatial transcriptomics technologies; q1 panel fields empty; q2 profile empty; final notes: Pass-A abstract: OTHER → FILTERED | accepted_agree |
| 10.1371/journal.pbio.3002133 | 2023 | A guide to the BRAIN Initiative Cell Census Network data ecosystem | AGREE | Title only; paper_type OTHER; spatial=YES; panel_scope=UNCLEAR; q1 panel-design fields are empty; gemma_final=FILTERED from abstract prefilter. | accepted_agree |
| 10.1371/journal.pbio.3003477 | 2025 | Cell2Spatial is a computational framework that maps single cells to spatial transcriptomic spots to reconstruct tissue architecture. | AGREE | title: Cell2Spatial is a computational framework that maps single cells to spatial transcriptomic spots to reconstruct | accepted_agree |

| DOI | Year | Title | Audit decision | Evidence (verbatim quote) | Validation status / scope flag |
|---|---|---|---|---|---|
| | | | | tissue architecture.; q1 panel fields empty; q2 profile empty; final notes: Pass-A abstract: COMP → FILTERED | |
| 10.1371/journal.pcbi.1009274 | 2021 | Validation and tuning of in situ transcriptomics image processing workflows with crowdsourced annotations. | AGREE | title: Validation and tuning of in situ transcriptomics image processing workflows with crowdsourced annotations.; q1 panel fields empty; q2 profile empty; final notes: Pass-A abstract: COMP → FILTERED | accepted_agree |
| 10.1371/journal.pcbi.1009600 | 2022 | A copula based topology preserving graph convolution network for clustering of single-cell RNA-seq data | AGREE | title='A copula based topology preserving graph convolution network for clustering of single-cell RNA-seq data'; citation_screening_prior.labels='COMP'; gemma_final.source='citation_screening_prefilter' | accepted_agree |
| 10.1371/journal.pcbi.1010366 | 2022 | De novo spatiotemporal modelling of cell-type signatures in the developmental human heart using graph convolutional neural networks. | AGREE | q2 describes de novo spatiotemporal analysis of existing ISS data using a graph convolutional model. | accepted_agree |
| 10.1371/journal.pcbi.1011288 | 2023 | The specious art of single-cell genomics | AGREE | Title only; gemma_final notes title/labels pre-filter as review; q1 panel-design fields are empty. | accepted_agree |
| 10.1371/journal.pcbi.1011324 | 2023 | Curated single cell multimodal landmark datasets for R/Bioconductor. | AGREE | title='Curated single cell multimodal landmark datasets for R/Bioconductor.'; gemma_final=FILTERED from abstract_prefilter; q0 doc_type=blank; q1 panel fields empty; q2 paper_type=OTHER, spatial=YES, panel_scope=TARGETED | accepted_agree |
| 10.1371/journal.pcbi.1012643 | 2024 | Hierarchical marker genes selection in scRNA-seq analysis | AGREE | citation_screening_prior: computational-only prior; gemma_final source=citation_screening_prefilter -> FILTERED; q1 panel-design fields are empty; q2 paper_type=COMP. | accepted_agree |

| DOI | Year | Title | Audit decision | Evidence (verbatim quote) | Validation status / scope flag |
|---|---|---|---|---|---|
| 10.1371/journal.pcbi.1013212.s001 | 2025 | Downstream analysis on STARmap data of the mouse placenta. | NEEDS_MANUAL | doc_type=SUPPLEMENT; packet does not provide usable main-article panel-design evidence. Original audit evidence: q0: SUPPLEMENT; q1/q2 fields are empty. | validated_high_risk_manual_filtered |
| 10.1371/journal.pcbi.1013212.s002 | 2025 | Downstream analysis on MERFISH data of mouse hypothalamic preoptic region. | AGREE | The packet is titled downstream analysis on MERFISH data and contains no panel-design evidence. | accepted_agree |
| 10.1371/journal.pcbi.1013212.s003 | 2025 | Downstream analysis on seqFISH+ data of mouse secondary somatosensory cortex. | AGREE | The packet title states downstream analysis on seqFISH+ data and gives no panel-design evidence. | accepted_agree |
| 10.1371/journal.pcbi.1013354 | 2025 | Improving cell-type composition inference in spatial transcriptomics with SpaDAMA | AGREE | title: Improving cell-type composition inference in spatial transcriptomics with SpaDAMA; citation prior labels=COMP; q2 paper_type=COMP; q1 panel-design fields are empty; gemma_final=FILTERED. | accepted_agree |
| 10.1371/journal.pcbi.1013759 | 2025 | Randomized Spatial PCA (RASP): A computationally efficient method for dimensionality reduction of high-resolution spatial transcriptomics data. | AGREE | title='Randomized Spatial PCA (RASP): A computationally efficient method for dimensionality reduction of high-resolution spatial transcriptomics data.'; gemma_final=FILTERED from abstract_prefilter; q0 doc_type=blank; q1 panel fields empty; q2 paper_type=COMP, spatial=YES, panel_scope=UNCLEAR | accepted_agree |
| 10.1371/journal.pcbi.1013759.s001 | 2025 | Includes supplemental methods and results. | AGREE | title='Includes supplemental methods and results.'; gemma_final=FILTERED from abstract_prefilter; q0 doc_type=blank; q1 panel fields empty; q2 paper_type=COMP, spatial=YES, panel_scope=UNCLEAR | accepted_agree |
| 10.1371/journal.pcbi.1013810 | 2025 | MAGNET: Multi-view graph autoencoder with cell-gene attention for cell interaction network reconstruction from spatial transcriptomics. | NEEDS_MANUAL | Final manual decision: FILTERED because the paper is a downstream analysis/modeling/QC method or dataset-use paper, with no final panel-design source. Original evidence: q1: | manual_final_filter_after_pdf_review |

| DOI | Year | Title | Audit decision | Evidence (verbatim quote) | Validation status / scope flag |
|---|---|---|---|---|---|
| | | | | panel_design=UNCLEAR; q2: benchmark datasets only. | |
| 10.1371/journal.pcbi.1014110 | 2026 | Clustering single-cell multi-omics data via weighted distance penalty and adaptive consistent graph regularization | AGREE | citation_screening_prior labels=COMP and all_tools=COSG; gemma_final source=citation_screening_prefilter -> FILTERED; q1 panel-design fields are empty. | accepted_agree |
| 10.1371/journal.pntd.0013120 | 2025 | Probing orthobunyavirus reassortment using Bunyamwera and Batai viruses as models. | AGREE | title='Probing orthobunyavirus reassortment using Bunyamwera and Batai viruses as models.'; gemma_final=FILTERED from abstract_prefilter; q0 doc_type=blank; q1 panel fields empty; q2 paper_type=EXP, spatial=NO, panel_scope=UNCLEAR | accepted_agree |
| 10.1371/journal.pone.0140227 | 2015 | Development and Molecular Cytogenetic Identification of a Novel Wheat-Leymus mollis Lm#7Ns (7D) Disomic Substitution Line with Stripe Rust Resistance. | AGREE | title='Development and Molecular Cytogenetic Identification of a Novel Wheat-Leymus mollis Lm#7Ns (7D) Disomic Substitution Line with Stripe Rust Resistance.'; gemma_final=FILTERED from abstract_prefilter; q0 doc_type=blank; q1 panel fields empty; q2 paper_type=EXP, spatial=NO, panel_scope=UNCLEAR | accepted_agree |
| 10.1371/journal.pone.0275070 | 2022 | Machine learning for cell type classification from single nucleus RNA sequencing data | AGREE | title='Machine learning for cell type classification from single nucleus RNA sequencing data'; citation_screening_prior.labels='COMP-BENCH'; gemma_final.source='citation_screening_prefilter' | accepted_agree |
| 10.1371/journal.pone.0292961 | 2023 | On the use of QDE-SVM for gene feature selection and cell type classification from scRNA-seq data | AGREE | citation_screening_prior: computational-only prior; gemma_final source=citation_screening_prefilter -> FILTERED; q1 panel-design fields are empty; q2 paper_type=COMP. | accepted_agree |
| 10.14288/1.0451898 | 2026 | Topic-guided cell segmentation for image-based spatial transcriptomics | AGREE | Title is 'Topic-guided cell segmentation for image-based spatial transcriptomics'; q2 marks | accepted_agree |

| DOI | Year | Title | Audit decision | Evidence (verbatim quote) | Validation status / scope flag |
|---|---|---|---|---|---|
| | | | | paper_type=COMP and q1 panel-design fields are empty. | |
| 10.14293/apmc13-2025-0012 | 2025 | Discovering cell type and tissue architecture with imaging-based spatial transcriptomics | AGREE | title='Discovering cell type and tissue architecture with imaging-based spatial transcriptomics'; gemma_final=FILTERED from abstract_prefilter; q0 doc_type=blank; q1 panel fields empty; q2 paper_type=COMP, spatial=YES, panel_scope=UNCLEAR | accepted_agree |
| 10.14293/apmc13-2025-0332 | 2025 | Spatially Guided Single-Cell Analysis Integrating Spatial Transcriptomics and Spatial Cell Sorting for In-Depth Profiling | AGREE | title='Spatially Guided Single-Cell Analysis Integrating Spatial Transcriptomics and Spatial Cell Sorting for In-Depth Profiling'; gemma_final=FILTERED from gemma4; q0 doc_type=NON_ARTICLE; q1 panel fields empty; q2 paper_type=blank, spatial=blank, panel_scope=blank | accepted_agree |
| 10.14569/ijacsa.2023.0140984 | 2023 | Analyzing RNA-Seq Gene Expression Data for Cancer Classification Through ML Approach | AGREE | Title only; paper_type COMP; spatial blank; panel_scope blank; q1 panel-design fields are empty; gemma_final=FILTERED from citation-screening prefilter. | accepted_agree |
| 10.1515/pcr-2015-0014 | 2015 | From cartography of the Universe to molecular cartography – the use of map projections | AGREE | title='From cartography of the Universe to molecular cartography – the use of map projections'; gemma_final=FILTERED from abstract_prefilter; q0 doc_type=blank; q1 panel fields empty; q2 paper_type=OTHER, spatial=NO, panel_scope=UNCLEAR | accepted_agree |
| 10.15347/wjs/2021.004 | 2021 | A broad introduction to RNA-Seq | AGREE | title='A broad introduction to RNA-Seq'; gemma_final=FILTERED from abstract_prefilter; q0 doc_type=blank; q1 panel fields empty; q2 paper_type=OTHER, spatial=NO, panel_scope=UNCLEAR | accepted_agree |
| 10.15761/nnr.1000116 | 2019 | Cryotherapy in Tension Headache: analysis of the frequency of symptoms | AGREE | title='Cryotherapy in Tension Headache: analysis of the frequency of symptoms'; gemma_final=FILTERED from | accepted_agree |

| DOI | Year | Title | Audit decision | Evidence (verbatim quote) | Validation status / scope flag |
|---|---|---|---|---|---|
| | | | | abstract_prefilter; q0 doc_type=blank; q1 panel fields empty; q2 paper_type=EXP, spatial=NO, panel_scope=UNCLEAR | |
| 10.15826/izv2.2022.24.2.026 | 2022 | Xenia by A. Dragomoshchenko and Ja. Iwaszkiewicz: The Typological Similarity of Genre Innovation | AGREE | title='Xenia by A. Dragomoshchenko and Ja. Iwaszkiewicz: The Typological Similarity of Genre Innovation'; gemma_final=FILTERED from abstract_prefilter; q0 doc_type=blank; q1 panel fields empty; q2 paper_type=OTHER, spatial=NO, panel_scope=UNCLEAR | accepted_agree |
| 10.1590/s1678-86212018000300264 | 2018 | Editorial: Edição especial ENCAC 2017 | AGREE | title='Editorial: Edição especial ENCAC 2017'; gemma_final=FILTERED from title_prefilter; q0 doc_type=EDITORIAL; q1 panel fields empty; q2 paper_type=blank, spatial=blank, panel_scope=blank | accepted_agree |
| 10.1615/jmachlearnmodelcomput.2023047230 | 2023 | EVALUATING THE COMPLEXITY OF GENE COEXPRESSION ESTIMATION FOR SINGLE-CELL DATA | AGREE | Title only; paper_type COMP; spatial=NO; panel_scope=UNCLEAR; q1 panel-design fields are empty; gemma_final=FILTERED from abstract prefilter. | accepted_agree |
| 10.1681/asn.20223311s114b | 2022 | Genome-Wide CRISPR Screen Reveals That Elamipretide (SS-31) Mediated PLSCR3 Activation Mitigates Mitochondrial Dysfunction During AKI | AGREE | title='Genome-Wide CRISPR Screen Reveals That Elamipretide (SS-31) Mediated PLSCR3 Activation Mitigates Mitochondrial Dysfunction During AKI'; gemma_final=FILTERED from gemma4; q0 doc_type=NON_ARTICLE; q1 panel fields empty; q2 paper_type=blank, spatial=blank, panel_scope=blank | accepted_agree |
| 10.1681/asn.20223311s114c | 2022 | Downregulation of Syndecan-1 and Alternative Complement in Renal Proximal Tubular Epithelial Cells by Crotamine/Sirna Complexes | AGREE | title='Downregulation of Syndecan-1 and Alternative Complement in Renal Proximal Tubular Epithelial Cells by Crotamine/Sirna Complexes'; gemma_final=FILTERED from gemma4; q0 doc_type=NON_ARTICLE; q1 | accepted_agree |

| DOI | Year | Title | Audit decision | Evidence (verbatim quote) | Validation status / scope flag |
|---|---|---|---|---|---|
| | | | | panel fields empty; q2 paper_type=blank, spatial=blank, panel_scope=blank | |
| 10.1681/asn.20240hhbthkj | 2024 | Multimodal Spatial Transcriptomic Characterization of Mouse Kidney Injury and Repair | AGREE | title='Multimodal Spatial Transcriptomic Characterization of Mouse Kidney Injury and Repair'; gemma_final=FILTERED from gemma4; q0 doc_type=NON_ARTICLE; q1 panel fields empty; q2 paper_type=blank, spatial=blank, panel_scope=blank | accepted_agree |
| 10.1681/asn.2024drdrywbz | 2024 | Mapping the Microenvironment: A Single-Cell Spatial Atlas of Diabetic Nephropathy | AGREE | title='Mapping the Microenvironment: A Single-Cell Spatial Atlas of Diabetic Nephropathy'; gemma_final=FILTERED from gemma4; q0 doc_type=NON_ARTICLE; q1 panel fields empty; q2 paper_type=blank, spatial=blank, panel_scope=blank | accepted_agree |
| 10.1681/asn.2024eb9gnfwk | 2024 | Single-Cell and Spatial Transcriptomics Identify Immediate Early Genes as Central Transcriptomic Factors during Renal Endothelial Regeneration In Vivo | AGREE | title='Single-Cell and Spatial Transcriptomics Identify Immediate Early Genes as Central Transcriptomic Factors during Renal Endothelial Regeneration In Vivo'; gemma_final=FILTERED from gemma4; q0 doc_type=NON_ARTICLE; q1 panel fields empty; q2 paper_type=blank, spatial=blank, panel_scope=blank | accepted_agree |
| 10.1681/asn.2024fnrwxc0m | 2024 | Single-Cell Spatial Atlas of Immune and Kidney Cell States in Human Rapidly Progressive Glomerulonephritis | AGREE | title='Single-Cell Spatial Atlas of Immune and Kidney Cell States in Human Rapidly Progressive Glomerulonephritis'; gemma_final=FILTERED from gemma4; q0 doc_type=NON_ARTICLE; q1 panel fields empty; q2 paper_type=blank, spatial=blank, panel_scope=blank | accepted_agree |

| DOI | Year | Title | Audit decision | Evidence (verbatim quote) | Validation status / scope flag |
|---|---|---|---|---|---|
| 10.1681/asn.2024tqndbj8a | 2024 | Cell States and Niches in Diabetic Kidneys Identified by Spatial Mapping of Targeted Gene Panel at Single-Cell Resolution | AGREE | title='Cell States and Niches in Diabetic Kidneys Identified by Spatial Mapping of Targeted Gene Panel at Single-Cell Resolution'; gemma_final=FILTERED from gemma4; q0 doc_type=NON_ARTICLE; q1 panel fields empty; q2 paper_type=blank, spatial=blank, panel_scope=blank | accepted_agree |
| 10.17160/josha.12.3.1044 | 2025 | Frontiers of Scientific Discovery: Protein Evolution, Spatial Transcriptomics, and Antibody Research | AGREE | title='Frontiers of Scientific Discovery: Protein Evolution, Spatial Transcriptomics, and Antibody Research'; gemma_final=FILTERED from abstract_prefilter; q0 doc_type=blank; q1 panel fields empty; q2 paper_type=OTHER, spatial=YES, panel_scope=TARGETED | accepted_agree |
| 10.17912/micropub.biology.001274 | 2024 | Visualization of gene expression in <i>Pristionchus pacificus</i> with smFISH and in situ HCR. | NEEDS_MANUAL | Final manual decision: FILTERED because the available evidence concerns probe/barcode/oligo design or single-gene detection, not final gene-panel selection. Original evidence: HCR probe maker, a probe design software, was downloaded; no gene-selection source is stated. | manual_final_filter_after_pdf_review |
| 10.18280/ria.360505 | 2022 | Diffusion Convolutional Recurrent Neural Network-Based Load Forecasting During COVID-19 Pandemic Situation | AGREE | title='Diffusion Convolutional Recurrent Neural Network-Based Load Forecasting During COVID-19 Pandemic Situation'; citation_screening_prior.labels='COMP'; gemma_final.source='citation_screening_prefilter' | accepted_agree |
| 10.18609/ioi.2021.036 | 2021 | Cancer in 4D: the importance of spatial and temporal heterogeneity through breast cancer molecular cartography | AGREE | title: Cancer in 4D: the importance of spatial and temporal heterogeneity through breast cancer molecular cartography; final notes: Pass-A abstract: OTHER -> FILTERED; q2 profile: spatial=YES, scope=TARGETED, panel fields empty. | accepted_agree |

| DOI | Year | Title | Audit decision | Evidence (verbatim quote) | Validation status / scope flag |
|---|---|---|---|---|---|
| 10.20517/2394-4722.2023.131 | 2024 | Prognostic value of clonal evolution identified by sequential FISH in untreated chronic lymphocytic leukaemia | AGREE | title: Prognostic value of clonal evolution identified by sequential FISH in untreated chronic lymphocytic leukaemia; final notes: Pass-A abstract: non-spatial -> FILTERED; q2 profile: spatial=NO, scope=UNCLEAR, panel fields empty. | accepted_agree |
| 10.20538/1682-0363-2025-4-111-119 | 2026 | Modern methods of DNA probe synthesis for fluorescence in situ hybridization (FISH): technologies and applications | AGREE | title: Modern methods of DNA probe synthesis for fluorescence in situ hybridization (FISH): technologies and applications; final notes: Pass-A abstract: OTHER -> FILTERED; q2 profile: spatial=YES, scope=UNCLEAR, panel fields empty. | accepted_agree |
| 10.20868/upm.thesis.43380 | 2016 | Genomics of Specificity in the Symbiotic Interaction between Rhizobium leguminosarum and Legumes | AGREE | title: Genomics of Specificity in the Symbiotic Interaction between Rhizobium leguminosarum and Legumes; final notes: Pass-A abstract: non-spatial -> FILTERED; q2 profile: spatial=NO, scope=UNCLEAR, panel fields empty. | accepted_agree |
| 10.20944/preprints202410.2601.v1 | 2024 | Single Cell Sequencing: Genomic and Transcriptomics Approaches in Cancer Cell Biology | AGREE | title: Single Cell Sequencing: Genomic and Transcriptomics Approaches in Cancer Cell Biology; final notes: Pass-A abstract: OTHER -> FILTERED; q2 profile: spatial=YES, scope=UNCLEAR, panel fields empty. | accepted_agree |
| 10.20944/preprints202504.1450.v1 | 2025 | A Practical Guide to Spatial Transcriptomics: Lessons from over 1000 Samples | AGREE | title: A Practical Guide to Spatial Transcriptomics: Lessons from over 1000 Samples; final notes: Pass-A abstract: OTHER -> FILTERED; q2 profile: spatial=YES, scope=UNCLEAR, panel fields empty. | accepted_agree |
| 10.21203/rs.3.rs-119885/v1 | 202 | Single-Cell Manifold Preserving Feature Selection (SCMER) | AGREE | Title: 'Single-Cell Manifold Preserving Feature Selection (SCMER)'; packet fields show paper_type=COMP-BENCH and Gemma's FILTERED prefilter. | accepted_agree |
| 10.21203/rs.3.rs-1242718/v1 | 2022 | Molecular Harvesting of Proteins with Electroporation In Vivo | AGREE | The paper profile is NOT_TARGETED and no panel- | accepted_agree |

| DOI | Year | Title | Audit decision | Evidence (verbatim quote) | Validation status / scope flag |
|---|---|---|---|---|---|
| | | Facilitates the Profiling of Spatial Differential Protein Expression in Tumors | | design evidence is provided. | |
| 10.21203/rs.3.rs-2716740/v1 | 2023 | A novel subclonal rearrangement of the STRN3::PDGFRB genes in de novo acute myeloid leukemia with NPM1 mutation and its leukemogenic effects | AGREE | title: A novel subclonal rearrangement of the STRN3::PDGFRB genes in de novo acute myeloid leukemia with NPM1 mutation and its leukemogenic effects; final notes: Pass-A abstract: non-spatial -> FILTERED; q2 profile: spatial=NO, scope=UNCLEAR, panel fields empty. | accepted_agree |
| 10.21203/rs.3.rs-2826640/v1 | 2023 | SpatialCTD: a large-scale TME spatial transcriptomic dataset to evaluate cell type deconvolution for immuno-oncology | AGREE | title: SpatialCTD: a large-scale TME spatial transcriptomic dataset to evaluate cell type deconvolution for immuno-oncology; final notes: Pass-A abstract: COMP -> FILTERED; q2 profile: spatial=YES, scope=UNCLEAR, panel fields empty. | accepted_agree |
| 10.21203/rs.3.rs-3583635/v1 | 2024 | Generative Self-Supervised Graphs Enhance Integration, Imputation and Domains Identification of Spatial Transcriptomics | AGREE | title: Generative Self-Supervised Graphs Enhance Integration, Imputation and Domains Identification of Spatial Transcriptomics; final notes: Pass-A abstract: COMP -> FILTERED; q2 profile: spatial=YES, scope=UNCLEAR, panel fields empty. | accepted_agree |
| 10.21203/rs.3.rs-3657512/v1 | 2023 | Introgression of tetraploid Thinopyrum elongatum 6EL segments enhances the stripe rust resistance of adult wheat plants | AGREE | title: Introgression of tetraploid Thinopyrum elongatum 6EL segments enhances the stripe rust resistance of adult wheat plants; final notes: Pass-A abstract: non-spatial -> FILTERED; q2 profile: spatial=NO, scope=UNCLEAR, panel fields empty. | accepted_agree |

| DOI | Year | Title | Audit decision | Evidence (verbatim quote) | Validation status / scope flag |
|---|---|---|---|---|---|
| 10.21203/rs.3.rs-38569/v3 | 2021 | Physical mapping of repetitive oligonucleotides facilitates the establishment of a genome map-based karyotype to identify chromosomal variations in peanut | AGREE | title: Physical mapping of repetitive oligonucleotides facilitates the establishment of a genome map-based karyotype to identify chromosomal variations in peanut; final notes: Pass-A abstract: non-spatial -> FILTERED; q2 profile: spatial=NO, scope=UNCLEAR, panel fields empty. | accepted_agree |
| 10.21203/rs.3.rs-3960911/v1 | 2024 | Community effort to unravel the complex human neuronal landscape exemplified for the striatum | AGREE | Title only; paper_type blank; spatial=NO; panel_scope blank; q1 panel-design fields are empty; gemma_final=FILTERED from citation-screening prefilter. | accepted_agree |
| 10.21203/rs.3.rs-4241968/v1 | 2024 | Identification of Key Candidate Genes Related to Cartilage development during Murine embryonic limb development by single cell RNA-sequencing and Experimental Confirmation | AGREE | Title only; paper_type COMP; spatial=NO; panel_scope blank; q1 panel-design fields are empty; gemma_final=FILTERED from abstract prefilter. | accepted_agree |
| 10.21203/rs.3.rs-4419408/v1 | 2024 | ABDS: a bioinformatics tool suite for analyzing biologically diverse samples | AGREE | Title only; paper_type COMP; spatial blank; panel_scope blank; q1 panel-design fields are empty; gemma_final=FILTERED from abstract prefilter. | accepted_agree |
| 10.21203/rs.3.rs-4428586/v1 | 2024 | Accurate, Scalable and Cross-platform Cell Identification for High-resolution Spatial Transcriptomics | AGREE | title: Accurate, Scalable and Cross-platform Cell Identification for High-resolution Spatial Transcriptomics; final notes: Pass-A abstract: COMP -> FILTERED; q2 profile: spatial=YES, scope=UNCLEAR, panel fields empty. | accepted_agree |
| 10.21203/rs.3.rs-4536158/v1 | 2024 | Spatial Deconvolution of Cell Types and Cell States at Scale Utilizing TACIT | AGREE | Title only; paper_type COMP; spatial=YES; panel_scope=UNCLEAR; q1 panel-design fields are empty; gemma_final=FILTERED from abstract prefilter. | accepted_agree |
| 10.21203/rs.3.rs-4730227/v1 | 2024 | Single-cell profiling reveals phenotypic changes in chronic atrophic gastritis | AGREE | Title only; paper_type blank; spatial=NO; panel_scope blank; q1 panel-design fields are empty; gemma_final=FILTERED from citation-screening prefilter. | accepted_agree |

| DOI | Year | Title | Audit decision | Evidence (verbatim quote) | Validation status / scope flag |
|---|---|---|---|---|---|
| 10.21203/rs.3.rs-4909620/v1 | 2025 | Thor: a platform for cell-level investigation of spatial transcriptomics and histology | AGREE | title: Thor: a platform for cell-level investigation of spatial transcriptomics and histology; final notes: Pass-A abstract: COMP -> FILTERED; q2 profile: spatial=YES, scope=UNCLEAR, panel fields empty. | accepted_agree |
| 10.21203/rs.3.rs-5969552/v1 | 2025 | Single Cell Spatial Transcriptomics of the Murine Embryonic Palate Links Pax9 to Patterning and Organization of Extracellular Matrix Components | AGREE | title: Single Cell Spatial Transcriptomics of the Murine Embryonic Palate Links Pax9 to Patterning and Organization of Extracellular Matrix Components; final notes: Pass-A abstract: not targeted -> FILTERED; q2 profile: spatial=YES, scope=NOT_TARGETED, panel fields empty. | accepted_agree |
| 10.21203/rs.3.rs-6080581/v1 | 2025 | Decoding cellular transcriptional regulatory networks governing wheat inflorescence development | NEEDS_MANUAL | Final manual decision: FILTERED because manual review found no usable evidence for final gene-panel source selection. Original evidence: evidence_tool=NOT FOUND, evidence_manual=NOT FOUND, evidence_vendor=NOT FOUND; panel_scope=TARGETED but no panel-design rationale is provided. | manual_final_filter_after_pdf_review |
| 10.21203/rs.3.rs-6389993/v1 | 2025 | Evaluating Deconvolution Methods Using Real Bulk RNA-expression Data for Robust Prognostic Insights in Pan-Cancer | AGREE | title: Evaluating Deconvolution Methods Using Real Bulk RNA-expression Data for Robust Prognostic Insights in Pan-Cancer; citation prior labels=COMP; q2 paper_type=COMP; q1 panel-design fields are empty. | accepted_agree |
| 10.21203/rs.3.rs-6930366/v1 | 2025 | CellLoop: Identifying single-cell 3D genome chromatin loops | NEEDS_MANUAL | Final manual decision: FILTERED because the paper is a downstream analysis/modeling/QC method or dataset-use paper, with no final panel-design source. Original evidence: q1: panel_design=UNCLEAR; q2: MERFISH mentioned, but no target-gene selection evidence. | manual_final_filter_after_pdf_review |
| 10.21203/rs.3.rs-7243521/v1 | 2025 | Deciphering Microenvironmental Heterogeneity by Scalable Niche Guided Module Discovery | AGREE | title: Deciphering Microenvironmental Heterogeneity by Scalable Niche Guided Module | accepted_agree |

| DOI | Year | Title | Audit decision | Evidence (verbatim quote) | Validation status / scope flag |
|---|---|---|---|---|---|
| | | | | Discovery; final notes: Pass-A abstract: COMP -> FILTERED; q2 profile: spatial=YES, scope=UNCLEAR, panel fields empty. | |
| 10.21203/rs.3.rs-8566824/v1 | 2026 | Microenvironment-aware transcriptome reconstruction in spatial transcriptomics | NEEDS_MANUAL | Final manual decision: FILTERED because no local PDF was available and the packet did not provide usable final gene-panel source evidence. Original evidence: No panel-design evidence is present in the packet. | manual_final_filter_after_pdf_review |
| 10.21203/rs.3.rs-914457/v1 | 2021 | Q-Mer Analysis: A Generalized Method for Analyzing RNA-Seq Data | AGREE | Title: 'Q-Mer Analysis: A Generalized Method for Analyzing RNA-Seq Data'; packet fields show paper_type=COMP and Gemma's FILTERED prefilter. | accepted_agree |
| 10.21203/rs.3.rs-990495/v1 | 2021 | CCST: Cell clustering for spatial transcriptomics data with graph neural network | NEEDS_MANUAL | CCST clusters MERFISH, seqFISH+, and 10x Visium datasets; no panel-design or panel-selection step is described. | pdf_validated_unclear_update |
| 10.2139/ssrn.3766496 | 2021 | Characterizing Tissue Composition Through Combined Analysis of Morphologies and Transcriptional States | AGREE | title: Characterizing Tissue Composition Through Combined Analysis of Morphologies and Transcriptional States; final notes: Pass-A abstract: COMP -> FILTERED; q2 profile: spatial=YES, scope=UNCLEAR, panel fields empty. | accepted_agree |
| 10.2139/ssrn.6556561 | 2026 | Unveiling Finer-Grained Cellular Structures and Rare Pathological Niches through Scalable Spatial Omics Multi-Integration | AGREE | title: Unveiling Finer-Grained Cellular Structures and Rare Pathological Niches through Scalable Spatial Omics Multi-Integration; final notes: Pass-A abstract: COMP -> FILTERED; q2 profile: spatial=YES, scope=UNCLEAR, panel fields empty. | accepted_agree |
| 10.2147/itt.s451199 | 2024 | Characterizing CD8+ TEMRA Cells in CP/CPPS Patients: Insights from Targeted Single-Cell Transcriptomic and Functional Investigations | AGREE | Title only; paper_type blank; spatial=NO; panel_scope blank; q1 panel-design fields are empty; gemma_final=FILTERED from citation-screening prefilter. | accepted_agree |
| 10.21608/ejh.2019.16977.1168 | 2019 | Histology Revolution: from inefficient, two-dimensional, and low-resolution techniques to high- | AGREE | title: Histology Revolution: from inefficient, two-dimensional, and low-resolution techniques to high- | accepted_agree |

| DOI | Year | Title | Audit decision | Evidence (verbatim quote) | Validation status / scope flag |
|---|---|---|---|---|---|
| | | throughput, three-dimensional and high-resolution techniques | | throughput, three-dimensional and high-resolution techniques; final notes: Pass-A abstract: OTHER -> FILTERED; q2 profile: spatial=YES, scope=UNCLEAR, panel fields empty. | |
| 10.2215/cjn.0000000000000146 | 2023 | Computational Pathology Fusing Spatial Technologies | AGREE | title: Computational Pathology Fusing Spatial Technologies; final notes: Pass-A abstract: OTHER -> FILTERED; q2 profile: spatial=YES, scope=UNCLEAR, panel fields empty. | accepted_agree |
| 10.24963/ijcai.2023/531 | 2023 | Pseudo-Labeling Enhanced by Privileged Information and Its Application to In Situ Sequencing Images | AGREE | Title: 'Pseudo-Labeling Enhanced by Privileged Information and Its Application to In Situ Sequencing Images'; packet fields show Q0: NON_ARTICLE and Gemma's FILTERED prefilter. | accepted_agree |
| 10.26434/chemrxiv-2023-n20f0 | 2023 | Spatial Transcriptomics: Emerging Technologies in Tissue Gene Expression Profiling | AGREE | title: Spatial Transcriptomics: Emerging Technologies in Tissue Gene Expression Profiling; final notes: Pass-A abstract: OTHER -> FILTERED; q2 profile: spatial=YES, scope=UNCLEAR, panel fields empty. | accepted_agree |
| 10.26434/chemrxiv-2025-m950n | 2025 | Multiplexed RNA imaging and in situ profiling in living cells | AGREE | paper.title='Multiplexed RNA imaging and in situ profiling in living cells'; gemma_final='Pass-A abstract: OTHER → FILTERED'; gemma_q2_spatial=YES; gemma_q2_paper_type=OTHER; empty q1 panel-design fields. | accepted_agree |
| 10.26599/bdma.2025.9020056 | 2025 | A Comprehensive Review of Cell-Type Deconvolution in Spatial Transcriptomic Data | AGREE | title: A Comprehensive Review of Cell-Type Deconvolution in Spatial Transcriptomic Data; citation prior labels=REV; q1 panel-design fields are empty; gemma_final=FILTERED. | accepted_agree |
| 10.29296/24999490-2024-02-02 | 2024 | SEQUENCING OF SINGLE CELLS: APPLICATION AND PERSPECTIVES | AGREE | paper.title='SEQUENCING OF SINGLE CELLS: APPLICATION AND PERSPECTIVES'; gemma_final='Pass-A abstract: OTHER → FILTERED'; gemma_q2_spatial=YES; gemma_q2_paper_type=OTHER; empty q1 panel-design fields. | accepted_agree |

| DOI | Year | Title | Audit decision | Evidence (verbatim quote) | Validation status / scope flag |
|---|---|---|---|---|---|
| 10.3103/s0096392519020056 | 2019 | Some Peculiarities in Application of Denaturating and Non-Denaturating In Situ Hybridization on Chromosomes of Cereals | AGREE | paper.title='Some Peculiarities in Application of Denaturating and Non-Denaturating In Situ Hybridization on Chromosomes of Cereals'; gemma_final='Pass-A abstract: OTHER → FILTERED'; gemma_q2_spatial=YES; gemma_q2_paper_type=OTHER; empty q1 panel-design fields. | accepted_agree |
| 10.31083/fbl48370 | 2026 | Neonatal Hyperoxia Induces Metabolic Reprogramming in Senescent Alveolar Macrophages, Leading to Persistent Lung Injury | AGREE | paper.title='Neonatal Hyperoxia Induces Metabolic Reprogramming in Senescent Alveolar Macrophages, Leading to Persistent Lung Injury'; citation_screening_prior.labels='NON-ST'; gemma_final.classification='FILTERED'; gemma_q1_panel_design blank; gemma_q2_paper_profile={'evidence_scope': '', 'evidence_spatial': '', 'evidence_type': '', 'panel_scope': '', 'paper_type': '', 'profile_confidence': '', 'spatial': 'NO'} | accepted_agree |
| 10.3150/21-bej1433 | 2022 | Estimation of Wasserstein distances in the Spiked Transport Model | AGREE | title='Estimation of Wasserstein distances in the Spiked Transport Model'; citation_screening_prior.labels='COMP'; gemma_final.source='citation_screening_prefilter' | accepted_agree |
| 10.31635/ccschem.022.202202171 | 2022 | Construction of a Stimuli-Responsive DNAzyme-Braked DNA Nanomachine for the Amplified Imaging of miRNAs in Living Cells and Mice | AGREE | paper.title='Construction of a Stimuli-Responsive DNAzyme-Braked DNA Nanomachine for the Amplified Imaging of miRNAs in Living Cells and Mice'; gemma_final='Pass-A abstract: non-spatial → FILTERED'; gemma_q2_spatial=NO; gemma_q2_paper_type=EXP; empty q1 panel-design fields. | accepted_agree |
| 10.33140/jcei.08.01.04 | 2023 | FISH or not to FISH - Therapeutic Implications of sequential FISH studies in diagnosing B-cell “Double-Hit” and “Triple Hit” lymphomas | AGREE | paper.title='FISH or not to FISH - Therapeutic Implications of sequential FISH studies in diagnosing B-cell “Double-Hit” and “Triple Hit” lymphomas'; | accepted_agree |

| DOI | Year | Title | Audit decision | Evidence (verbatim quote) | Validation status / scope flag |
|---|---|---|---|---|---|
| | | | | gemma_final='Pass-A abstract: non-spatial → FILTERED'; gemma_q2_spatial=NO; gemma_q2_paper_type=EXP; empty q1 panel-design fields. | |
| 10.3354/esr01305 | 2024 | Vulnerability of the Critically Endangered leatherback turtle to fisheries bycatch in the eastern Pacific Ocean. II. Assessment of mitigation measures | AGREE | paper.title='Vulnerability of the Critically Endangered leatherback turtle to fisheries bycatch in the eastern Pacific Ocean. II. Assessment of mitigation measures'; gemma_final='Pass-A abstract: non-spatial → FILTERED'; gemma_q2_spatial=NO; gemma_q2_paper_type=EXP; empty q1 panel-design fields. | accepted_agree |
| 10.3354/meps13032 | 2019 | Ecological Assessment of the Sustainable Impacts of Fisheries (EASI-Fish): a flexible vulnerability assessment approach to quantify the cumulative impacts of fishing in data-limited settings | AGREE | paper.title='Ecological Assessment of the Sustainable Impacts of Fisheries (EASI-Fish): a flexible vulnerability assessment approach to quantify the cumulative impacts of fishing in data-limited settings'; gemma_final='Pass-A abstract: COMP → FILTERED'; gemma_q2_spatial=NO; gemma_q2_paper_type=COMP; empty q1 panel-design fields. | accepted_agree |
| 10.3389/fams.2024.1403901 | 2024 | Analysis of community connectivity in spatial transcriptomics data. | AGREE | paper.title='Analysis of community connectivity in spatial transcriptomics data.'; gemma_final='Pass-A abstract: COMP → FILTERED'; gemma_q2_spatial=YES; gemma_q2_paper_type=COMP; empty q1 panel-design fields. | accepted_agree |
| 10.3389/fanim.2025.1551244 | 2025 | Integrating omics approaches in livestock biotechnology: innovations in production and reproductive efficiency | AGREE | title: Integrating omics approaches in livestock biotechnology: innovations in production and reproductive efficiency; citation prior labels=REV; q1 panel-design fields are empty; gemma_final=FILTERED. | accepted_agree |
| 10.3389/fbinf.2025.1713975 | 2025 | SpaLLM: a general framework for spatial domain identification with large language models. | AGREE | paper.title='SpaLLM: a general framework for spatial domain identification with large language models.'; gemma_final='Pass-A abstract: COMP → FILTERED'; | accepted_agree |

| DOI | Year | Title | Audit decision | Evidence (verbatim quote) | Validation status / scope flag |
|---|---|---|---|---|---|
| | | | | gemma_q2_spatial=YES; gemma_q2_paper_type=COMP; empty q1 panel-design fields. | |
| 10.3389/fcell.2022.944325 | 2022 | Single-cell transcriptomics reveals male germ cells and Sertoli cells developmental patterns in dairy goats | AGREE | Title: 'Single-cell transcriptomics reveals male germ cells and Sertoli cells developmental patterns in dairy goats'; packet fields show paper_type=COMP, spatial=NO, and Gemma's FILTERED prefilter. | accepted_agree |
| 10.3389/fcell.2025.1515681 | 2025 | Hypoxia signature derived from tumor-associated endothelial cells predict prognosis in gastric cancer | AGREE | title: Hypoxia signature derived from tumor-associated endothelial cells predict prognosis in gastric cancer; citation prior labels=NON-ST; q2 spatial=NO; q1 panel-design fields are empty. | accepted_agree |
| 10.3389/fcell.2025.1697930 | 2025 | Thyroid hormone receptor beta-2 (TRβ2) overexpression modulates photoreceptor phenotype diversity in a ligand-dependent manner. | AGREE | paper.title='Thyroid hormone receptor beta-2 (TRβ2) overexpression modulates photoreceptor phenotype diversity in a ligand-dependent manner.'; gemma_final='Pass-A abstract: non-spatial → FILTERED'; gemma_q2_spatial=NO; gemma_q2_paper_type=EXP; empty q1 panel-design fields. | accepted_agree |
| 10.3389/fcell.2025.1712688 | 2025 | Decoding dental mesenchymal stem cells diversity: single-cell transcriptomics maps heterogeneity in molar development | AGREE | title: Decoding dental mesenchymal stem cells diversity: single-cell transcriptomics maps heterogeneity in molar development; citation prior labels=REV; q1 panel-design fields are empty; gemma_final=FILTERED. | accepted_agree |
| 10.3389/fcimb.2021.694470 | 2021 | Tissue/Biofluid Specific Molecular Cartography of <i>Leishmania donovani</i> Infected BALB/c Mice: Deciphering Systemic Reprogramming. | AGREE | paper.title='Tissue/Biofluid Specific Molecular Cartography of <i>Leishmania donovani</i> Infected BALB/c Mice: Deciphering Systemic Reprogramming.'; gemma_final='Pass-A abstract: non-spatial → FILTERED'; gemma_q2_spatial=NO; gemma_q2_paper_type=EXP; empty q1 panel-design fields. | accepted_agree |
| 10.3389/fdgth.2025.1535018 | 2025 | The data scientist as a mainstay of the tumor board: global | AGREE | paper.title='The data scientist as a mainstay of the tumor board: global implications and | accepted_agree |

| DOI | Year | Title | Audit decision | Evidence (verbatim quote) | Validation status / scope flag |
|---|---|---|---|---|---|
| | | implications and opportunities for the global south | | opportunities for the global south'; gemma_final='Pass-A abstract: OTHER → FILTERED'; gemma_q2_spatial=NO; gemma_q2_paper_type=OTHER; empty q1 panel-design fields. | |
| 10.3389/fgene.2021.785290 | 2022 | Analysis and Visualization of Spatial Transcriptomic Data | AGREE | Title: 'Analysis and Visualization of Spatial Transcriptomic Data'; packet field doc_type=REVIEW and Gemma's FILTERED prefilter. | accepted_agree |
| 10.3389/fgene.2022.785877 | 2022 | SSAM-lite: A Light-Weight Web App for Rapid Analysis of Spatially Resolved Transcriptomics Data | AGREE | paper.title='SSAM-lite: A Light-Weight Web App for Rapid Analysis of Spatially Resolved Transcriptomics Data'; gemma_final='Pass-A abstract: COMP → FILTERED'; gemma_q2_spatial=YES; gemma_q2_paper_type=COMP; empty q1 panel-design fields. | accepted_agree |
| 10.3389/fgene.2022.893522 | 2022 | SpatialMap: Spatial Mapping of Unmeasured Gene Expression Profiles in Spatial Transcriptomic Data Using Generalized Linear Spatial Models. | AGREE | paper.title='SpatialMap: Spatial Mapping of Unmeasured Gene Expression Profiles in Spatial Transcriptomic Data Using Generalized Linear Spatial Models.'; gemma_final='Pass-A abstract: COMP → FILTERED'; gemma_q2_spatial=YES; gemma_q2_paper_type=COMP; empty q1 panel-design fields. | accepted_agree |
| 10.3389/fgene.2024.1356709 | 2024 | Bayesian hidden mark interaction model for detecting spatially variable genes in imaging-based spatially resolved transcriptomics data. | AGREE | we developed a generalized energy-based framework to model gene expression measured from imaging-based SRT platforms | accepted_agree |
| 10.3389/fgene.2025.1583081 | 2025 | High-precision lung cancer subtype diagnosis on imbalanced exosomal data via Exo-LCClassifier | AGREE | title: High-precision lung cancer subtype diagnosis on imbalanced exosomal data via Exo-LCClassifier; citation prior labels=COMP-BENCH; q2 paper_type=COMP; q1 panel-design fields are empty; gemma_final=FILTERED. | accepted_agree |
| 10.3389/fgene.2025.1695803 | 2025 | KSRV: a Kernel PCA-Based framework for inferring spatial RNA velocity at single-cell resolution. | AGREE | paper.title='KSRV: a Kernel PCA-Based framework for inferring spatial RNA velocity at single-cell resolution.'; gemma_final='Pass-A | accepted_agree |

| DOI | Year | Title | Audit decision | Evidence (verbatim quote) | Validation status / scope flag |
|---|---|---|---|---|---|
| | | | | abstract: COMP → FILTERED'; gemma_q2_spatial=YES; gemma_q2_paper_type=COMP; empty q1 panel-design fields. | |
| 10.3389/fimmu.2021.690470 | 2021 | Machine Learning-Based Single Cell and Integrative Analysis Reveals That Baseline mDC Predisposition Correlates With Hepatitis B Vaccine Antibody Response | AGREE | Title: 'Machine Learning-Based Single Cell and Integrative Analysis Reveals That Baseline mDC Predisposition Correlates With Hepatitis B Vaccine Antibody Response'; packet fields show spatial=NO and Gemma's FILTERED prefilter. | accepted_agree |
| 10.3389/fimmu.2022.1050951 | 2022 | Complex interaction and heterogeneity among cancer stem cells in head and neck squamous cell carcinoma revealed by single-cell sequencing | AGREE | Title: 'Complex interaction and heterogeneity among cancer stem cells in head and neck squamous cell carcinoma revealed by single-cell sequencing'; packet fields show paper_type=EXP, spatial blank/NO, and Gemma's FILTERED prefilter. | accepted_agree |
| 10.3389/fimmu.2024.1310376 | 2024 | Revealing molecular and cellular heterogeneity in hypopharyngeal carcinogenesis through single-cell RNA and TCR/BCR sequencing | AGREE | Title only; paper_type blank; spatial=NO; panel_scope blank; q1 panel-design fields are empty; gemma_final=FILTERED from citation-screening prefilter. | accepted_agree |
| 10.3389/fimmu.2024.1363517 | 2024 | A division-of-labor mode contributes to the cardioprotective potential of mesenchymal stem/stromal cells in heart failure post myocardial infarction | AGREE | Title only; paper_type blank; spatial=NO; panel_scope blank; q1 panel-design fields are empty; gemma_final=FILTERED from citation-screening prefilter. | accepted_agree |
| 10.3389/fimmu.2025.1521038 | 2025 | Unraveling the spatial and signaling dynamics and splicing kinetics of immune infiltration in osteoarthritis synovium | AGREE | gemma_q1_panel_design = NOT FOUND and gemma_q2_panel_scope = NOT_TARGETED. | accepted_agree |
| 10.3389/fimmu.2025.1530327 | 2025 | Cellular and molecular determinants mediating the dysregulated germinal center immune dynamics in systemic lupus erythematosus | NEEDS_MANUAL | panel_scope=NOT_TARGETED with no target-gene panel source evidence; untargeted/no-panel records are out of scope. Original audit evidence: gemma_q1_panel_design contains no evidence_tool, evidence_manual, or evidence_vendor; panel_scope is NOT_TARGETED. | validated_high_risk_manual_filtered \| Fig. 2: excluded (platform/scope) |
| 10.3389/fimmu.2025.1699915 | 2025 | Mapping TAM–tumor crosstalk in glioma via ligand–receptor multi- | AGREE | paper.title='Mapping TAM–tumor crosstalk in glioma via ligand– | accepted_agree |

| DOI | Year | Title | Audit decision | Evidence (verbatim quote) | Validation status / scope flag |
|---|---|---|---|---|---|
| | | omics: mechanisms of immune evasion | | receptor multi-omics: mechanisms of immune evasion'; gemma_q0_document_type=REVIEW; gemma_final=FILTERED. | |
| 10.3389/fmars.2021.627724 | 2021 | Three-Dimensional Molecular Cartography of the Caribbean Reef-Building Coral Orbicella faveolata | AGREE | paper.title='Three-Dimensional Molecular Cartography of the Caribbean Reef-Building Coral Orbicella faveolata'; gemma_final='Pass-A abstract: non-spatial → FILTERED'; gemma_q2_spatial=NO; gemma_q2_paper_type=EXP; empty q1 panel-design fields. | accepted_agree |
| 10.3389/fmicb.2018.03141 | 2018 | Genomic and in-situ Transcriptomic Characterization of the Candidate Phylum NPL-UPL2 From Highly Alkaline Highly Reducing Serpentinized Groundwater | NEEDS_MANUAL | spatial=NO; packet does not support an in-scope spatial panel-design record. Original audit evidence: evidence_tool=NOT FOUND, evidence_manual=NOT FOUND, evidence_vendor=NOT FOUND, spatial=NO, panel_scope=UNCLEAR. | validated_high_risk_manual_filtered |
| 10.3389/fnmol.2025.1716825 | 2025 | One RNA-binding protein, many decisions: integrating the transcript life cycle into neuronal regulation | AGREE | paper.title='One RNA-binding protein, many decisions: integrating the transcript life cycle into neuronal regulation'; gemma_final='Pass-A abstract: OTHER → FILTERED'; gemma_q2_spatial=YES; gemma_q2_paper_type=OTHER; empty q1 panel-design fields. | accepted_agree |
| 10.3389/fonc.2025.1552841 | 2025 | Single-cell RNA profiling of colorectal granular-type laterally spreading tumor uncovers progression trajectory toward carcinoma and transcriptional signatures favoring lateral morphogenesis | AGREE | paper.title='Single-cell RNA profiling of colorectal granular-type laterally spreading tumor uncovers progression trajectory toward carcinoma and transcriptional signatures favoring lateral morphogenesis'; gemma_q2_paper_profile.spatial=NO; gemma_final=FILTERED. | accepted_agree |

| DOI | Year | Title | Audit decision | Evidence (verbatim quote) | Validation status / scope flag |
|---|---|---|---|---|---|
| 10.3389/fphar.2025.1634985 | 2025 | Integrative analysis of lactylation related genes in prostate cancer: unveiling heterogeneity through single-cell RNA-seq, bulk RNA-seq and machine learning | AGREE | paper.title='Integrative analysis of lactylation related genes in prostate cancer: unveiling heterogeneity through single-cell RNA-seq, bulk RNA-seq and machine learning'; gemma_q2_paper_profile.spatial=NO; gemma_final=FILTERED. | accepted_agree |
| 10.3389/fphys.2018.01491 | 2018 | Analysis of in situ Transcriptomes Reveals Divergent Adaptive Response to Hyper- and Hypo-Salinity in the Hong Kong Oyster, Crassostrea hongkongensis | AGREE | paper.title='Analysis of in situ Transcriptomes Reveals Divergent Adaptive Response to Hyper- and Hypo-Salinity in the Hong Kong Oyster, Crassostrea hongkongensis'; gemma_final='Pass-A abstract: non-spatial → FILTERED'; gemma_q2_spatial=NO; gemma_q2_paper_type=EXP; empty q1 panel-design fields. | accepted_agree |
| 10.3389/fphys.2021.624037 | 2021 | Transcriptomic Identification of Draxin-Responsive Targets During Cranial Neural Crest EMT. | AGREE | paper.title='Transcriptomic Identification of Draxin-Responsive Targets During Cranial Neural Crest EMT.'; gemma_final='Pass-A abstract: non-spatial → FILTERED'; gemma_q2_spatial=NO; gemma_q2_paper_type=EXP; empty q1 panel-design fields. | accepted_agree |
| 10.3389/fphys.2022.882413 | 2022 | Optimization of Whole Mount RNA Multiplexed <i>in situ</i> Hybridization Chain Reaction With Immunohistochemistry, Clearing and Imaging to Visualize Octopus Embryonic Neurogenesis. | AGREE | paper.title='Optimization of Whole Mount RNA Multiplexed <i>in situ</i> Hybridization Chain Reaction With Immunohistochemistry, Clearing and Imaging to Visualize Octopus Embryonic Neurogenesis.'; gemma_final='Pass-A abstract: OTHER → FILTERED'; gemma_q2_spatial=YES; gemma_q2_paper_type=OTHER; empty q1 panel-design fields. | accepted_agree |
| 10.3389/fsysb.2022.910243 | 2022 | Leveraging Systems Immunology to Optimize Diagnosis and Treatment of Inborn Errors of Immunity | AGREE | Title: 'Leveraging Systems Immunology to Optimize Diagnosis and Treatment of Inborn Errors of Immunity'; packet fields show doc_type=REVIEW and Gemma's FILTERED prefilter. | accepted_agree |

| DOI | Year | Title | Audit decision | Evidence (verbatim quote) | Validation status / scope flag |
|---|---|---|---|---|---|
| 10.3390/a16050254 | 2023 | Well-Separated Pair Decompositions for High-Dimensional Datasets | AGREE | Title only; paper_type COMP; spatial blank; panel_scope blank; q1 panel-design fields are empty; gemma_final=FILTERED from citation-screening prefilter. | accepted_agree |
| 10.3390/agriculture15171887 | 2025 | Identification and Comparative Analysis of Genetic Effects of 2Ns Chromosome Introgression from Psathyrostachys huashanica and Leymus mollis into Common Wheat | AGREE | gemma_final: Pass-A abstract prefilter -> FILTERED; q1 panel-design fields are empty; q2 spatial=NO. | accepted_agree |
| 10.3390/ani14223263 | 2024 | Single-Nucleus RNA Sequencing Reveals the Transcriptome Profiling of Ovarian Cells in Adolescent Cyprinus carpio | AGREE | Title only; paper_type COMP; spatial=NO; panel_scope blank; q1 panel-design fields are empty; gemma_final=FILTERED from abstract prefilter. | accepted_agree |
| 10.3390/app12041850 | 2022 | Analyzing RNA-Seq Gene Expression Data Using Deep Learning Approaches for Cancer Classification | AGREE | Title: 'Analyzing RNA-Seq Gene Expression Data Using Deep Learning Approaches for Cancer Classification'; packet fields show paper_type=COMP and Gemma's FILTERED prefilter. | accepted_agree |
| 10.3390/bioengineering12080829 | 2025 | DropDAE: Denosing Autoencoder with Contrastive Learning for Addressing Dropout Events in scRNA-seq Data | AGREE | paper.title='DropDAE: Denosing Autoencoder with Contrastive Learning for Addressing Dropout Events in scRNA-seq Data'; gemma_final=FILTERED. | accepted_agree |
| 10.3390/biology15010095 | 2026 | Single-Cell Transcriptomic Profiling Reveals That Macrophage-Induced Angiogenesis Contributes to Immunotherapy Resistance in Hepatocellular Carcinoma | AGREE | paper.title='Single-Cell Transcriptomic Profiling Reveals That Macrophage-Induced Angiogenesis Contributes to Immunotherapy Resistance in Hepatocellular Carcinoma'; citation_screening_prior.labels='COMP'; gemma_final.classification='FILTERED'; gemma_q1_panel_design blank; gemma_q2_paper_profile= {'evidence_scope': '', 'evidence_spatial': '', 'evidence_type': '', 'panel_scope': '', 'paper_type': 'COMP', 'profile_confidence': '', 'spatial': ''} | accepted_agree |
| 10.3390/biology15070588 | 2026 | An AI-Enabled Single-Cell Transcriptomic Analysis Pipeline for Gene Signature Discovery in | AGREE | paper.title='An AI-Enabled Single-Cell Transcriptomic Analysis Pipeline for Gene Signature | accepted_agree |

| DOI | Year | Title | Audit decision | Evidence (verbatim quote) | Validation status / scope flag |
|---|---|---|---|---|---|
| | | Natural Killer Cells Linked to Remission Outcomes in Chronic Myeloid Leukemia | | Discovery in Natural Killer Cells Linked to Remission Outcomes in Chronic Myeloid Leukemia'; citation_screening_prior.labels='COMP'; gemma_final.classification='FILTERED'; gemma_q1_panel_design blank; gemma_q2_paper_profile={'evidence_scope': '', 'evidence_spatial': '', 'evidence_type': '', 'panel_scope': '', 'paper_type': 'COMP', 'profile_confidence': '', 'spatial': ''} | |
| 10.3390/biom14040436 | 2024 | GTADC: A Graph-Based Method for Inferring Cell Spatial Distribution in Cancer Tissues | AGREE | Title only; paper_type COMP; spatial blank; panel_scope blank; q1 panel-design fields are empty; gemma_final=FILTERED from abstract prefilter. | accepted_agree |
| 10.3390/biomedicines13030755 | 2025 | Ethanol Induces Craniofacial Defects in Bmp Mutants Independent of <i>nkx2.3</i> by Elevating Cranial Neural Crest Cell Apoptosis. | AGREE | gemma_final: Pass-A abstract prefilter -> FILTERED; q1 panel-design fields are empty; q2 spatial=NO. | accepted_agree |
| 10.3390/cancers13133198 | 2021 | High-Resolution Cartography of the Transcriptome and Methylome Landscapes of Diffuse Gliomas. | AGREE | gemma_final: Pass-A abstract prefilter -> FILTERED; q1 panel-design fields are empty; q2 spatial=NO. | accepted_agree |
| 10.3390/cancers14153628 | 2022 | Techniques for Profiling the Cellular Immune Response and Their Implications for Interventional Oncology | AGREE | Title: 'Techniques for Profiling the Cellular Immune Response and Their Implications for Interventional Oncology'; packet fields show doc_type=REVIEW and Gemma's FILTERED prefilter. | accepted_agree |
| 10.3390/cancers15051557 | 2023 | Quantitative Evaluation of Stem-like Markers of Human Glioblastoma Using Single-Cell RNA Sequencing Datasets | AGREE | Title only; paper_type COMP; spatial blank; panel_scope blank; q1 panel-design fields are empty; gemma_final=FILTERED from citation-screening prefilter. | accepted_agree |
| 10.3390/cancers15143615 | 2023 | Single-Cell Transcriptomics for Unlocking Personalized Cancer Immunotherapy: Toward Targeting the Origin of Tumor Development Immunogenicity | AGREE | Title only; gemma_final notes labels=['REV']; exp_used_tools=['(none)']; all_tools=['scGeneFit']; q1 panel-design fields are empty. | accepted_agree |
| 10.3390/cells12101442 | 2023 | Transcriptome-Powered Pluripotent Stem Cell | AGREE | Title only; gemma_final notes labels=['REV']; exp_used_tools= | accepted_agree |

| DOI | Year | Title | Audit decision | Evidence (verbatim quote) | Validation status / scope flag |
|---|---|---|---|---|---|
| | | Differentiation for Regenerative Medicine | | ['(none)']; all_tools=['NS-Forest v2']; q1 panel-design fields are empty. | |
| 10.3390/cells12162042 | 2023 | Spatial Transcriptomic Technologies | AGREE | Title only; gemma_final notes labels=['REV']; exp_used_tools=['(none)']; all_tools=['SMaSH']; q1 panel-design fields are empty. | accepted_agree |
| 10.3390/cells14131000 | 2025 | Defining Keypoints to Align H&E Images and Xenium DAPI-Stained Images Automatically. | AGREE | We present Xenium-Align, a keypoint identification method that automatically generates keypoint files for image registration in Xenium Explorer. | accepted_agree |
| 10.3390/genes14030596 | 2023 | Cell Type Annotation Model Selection: General-Purpose vs. Pattern-Aware Feature Gene Selection in Single-Cell RNA-Seq Data | AGREE | Title only; paper_type COMP; spatial blank; panel_scope blank; q1 panel-design fields are empty; gemma_final=FILTERED from citation-screening prefilter. | accepted_agree |
| 10.3390/genes17030310 | 2026 | Leveraging Spot-Gene Heterogeneous Graphs for Unified Spatially Resolved Transcriptomics Domain Detection on Single-Slice and Multi-Slice Data. | AGREE | gemma_final: Pass-A abstract prefilter -> FILTERED; q1 panel-design fields are empty; q2 paper_type=COMP. | accepted_agree |
| 10.3390/ijms21051861 | 202 | Characterization and Evaluation of Resistance to Powdery Mildew of Wheat-<i>Aegilops geniculata</i> Roth 7M<sup>g</sup> (7A) Alien Disomic Substitution Line W16998. | AGREE | gemma_final: Pass-A abstract prefilter -> FILTERED; q1 panel-design fields are empty; q2 spatial=NO. | accepted_agree |
| 10.3390/ijms23137056 | 2022 | Development and Molecular Cytogenetic Identification of Two Wheat-<i>Aegilops geniculata</i> Roth 7M<sup>g</sup> Chromosome Substitution Lines with Resistance to Fusarium Head Blight, Powdery Mildew and Stripe Rust. | AGREE | gemma_final: Pass-A abstract prefilter -> FILTERED; q1 panel-design fields are empty; q2 spatial=NO. | accepted_agree |
| 10.3390/ijms24032458 | 2023 | Big Data in Gastroenterology Research | AGREE | Title only; gemma_final notes labels=['REV']; exp_used_tools=['(none)']; all_tools=['NS-Forest v2']; q1 panel-design fields are empty. | accepted_agree |
| 10.3390/ijms24076726 | 2023 | Molecular Cytological Analysis and Specific Marker Development in Wheat-<i>Psathyrostachys huashanica</i> Keng 3Ns | AGREE | gemma_final: Pass-A abstract prefilter -> FILTERED; q1 panel-design fields are empty; q2 spatial=NO. | accepted_agree |

| DOI | Year | Title | Audit decision | Evidence (verbatim quote) | Validation status / scope flag |
|---|---|---|---|---|---|
| | | Additional Line with Elongated Glume. | | | |
| 10.3390/ijms25031479 | 2024 | Application of Single-Cell Assay for Transposase-Accessible Chromatin with High Throughput Sequencing in Plant Science: Advances, Technical Challenges, and Prospects | AGREE | Title only; doc_type=REVIEW; spatial blank; panel_scope blank; q1 panel-design fields are empty; gemma_final=FILTERED from citation-screening prefilter. | accepted_agree |
| 10.3390/ijms25041998 | 2024 | uPAR (PLAUR) Marks Two Intra-Tumoral Subtypes of Glioblastoma: Insights from Single-Cell RNA Sequencing | AGREE | Title only; paper_type blank; spatial=NO; panel_scope blank; q1 panel-design fields are empty; gemma_final=FILTERED from citation-screening prefilter. | accepted_agree |
| 10.3390/ijms252312606 | 2024 | Combining Network Pharmacology and Transcriptomic Strategies to Explore the Pharmacological Mechanism of Total Ginsenoside Ginseng Root and Its Impact on Antidepressant Effects | AGREE | Title only; paper_type blank; spatial=NO; panel_scope blank; q1 panel-design fields are empty; gemma_final=FILTERED from citation-screening prefilter. | accepted_agree |
| 10.3390/ijms252413706 | 2024 | Advancements in the Application of scRNA-Seq in Breast Research: A Review | AGREE | Title only; doc_type=REVIEW; spatial blank; panel_scope blank; q1 panel-design fields are empty; gemma_final=FILTERED from citation-screening prefilter. | accepted_agree |
| 10.3390/ijms262411819 | 2025 | Why “Where” Matters as Much as “How Much”: Single-Cell and Spatial Transcriptomics in Plants | AGREE | citation_screening_prior: labels=REV; tools=gpsFISH only in screening prior; q1 panel-design fields are empty. | accepted_agree |
| 10.3390/metabo10100406 | 202 | Metabolites from microbes isolated from the skin of the panamanian rocket frog colostethus panamansis (Anura: Dendrobatidae) | AGREE | gemma_final: Pass-A abstract prefilter -> FILTERED; q1 panel-design fields are empty; q2 spatial=NO. | accepted_agree |
| 10.3390/s23135990 | 2023 | Recent Approaches to Design and Analysis of Electrical Impedance Systems for Single Cells Using Machine Learning | AGREE | Title only; gemma_final notes labels=['REV']; exp_used_tools=['(none)']; all_tools=['SCMER']; q1 panel-design fields are empty. | accepted_agree |
| 10.3390/vaccines14020158 | 2026 | Advances in Spatial Transcriptomics for Infectious Disease Research: Insight for Vaccine Development | AGREE | gemma_final: Pass-A abstract prefilter -> FILTERED; q1 panel-design fields are empty; q2 paper_type=OTHER. | accepted_agree |
| 10.3732/apps.1700053 | 2017 | An efficient pipeline to generate data for studies in plastid | AGREE | gemma_final: Pass-A abstract prefilter -> FILTERED; q1 panel- | accepted_agree |

| DOI | Year | Title | Audit decision | Evidence (verbatim quote) | Validation status / scope flag |
|---|---|---|---|---|---|
| | | population genomics and phylogeography. | | design fields are empty; q2 paper_type=COMP. | |
| 10.3760/cma.j.cn112151-20210609-00426 | 2022 | [The interpretation trap of DDIT3 split FISH in liposarcoma]. | AGREE | gemma_final: Pass-A abstract prefilter -> FILTERED; q1 panel-design fields are empty; q2 spatial=NO. | accepted_agree |
| 10.38212/2224-6614.3385 | 2021 | Molecular cartography of residue pesticides on grape surface in 3D by ambient ionization tandem mass spectrometry. | AGREE | gemma_final: Pass-A abstract prefilter -> FILTERED; q1 panel-design fields are empty; q2 spatial=NO. | accepted_agree |
| 10.3929/ethz-b-000591173 | 2022 | Sphere-sequencing unveils local tissue microenvironments at single cell resolution | AGREE | gemma_final: Pass-A abstract prefilter -> FILTERED; q1 panel-design fields are empty; q2 paper_type=OTHER. | accepted_agree |
| 10.4049/jimmunol.208.supp.172.18 | 2022 | Cell type-specific differential expression in Alzheimer's disease and cancer using spatial transcriptomics | AGREE | gemma_final: Pass-A abstract prefilter -> FILTERED; q1 panel-design fields are empty; q2 paper_type=COMP. | accepted_agree |
| 10.4049/jimmunol.210.supp.219.08 | 2023 | Harnessing spatial transcriptomics to study fate decisions and lineage specification in T cell development | AGREE | gemma_final: Pass-A abstract prefilter -> FILTERED; q1 panel-design fields are empty; q2 panel_scope=NOT_TARGETED. | accepted_agree |
| 10.4103/1673-5374.327335 | 2021 | Guardians of the eye: new tales about retinal microglia and other ocular macrophages | AGREE | gemma_final: Pass-A abstract prefilter -> FILTERED; q1 panel-design fields are empty; q2 paper_type=OTHER. | accepted_agree |
| 10.4208/csiam-ls.so-2025-0032 | 2026 | BRIDGE: An Interpretable Framework for Transcriptome-Scale Completion of Panel-Limited Spatial Transcriptomics | AGREE | citation_screening_prior: labels=COMP; exp_used_tools=(none); all_tools=gpsFISH; q1 panel-design fields are empty. | accepted_agree |
| 10.4239/wjd.v16.i9.107663 | 2025 | Spatial transcriptomics meets diabetic kidney disease: Illuminating the path to precision medicine | AGREE | paper.title='Spatial transcriptomics meets diabetic kidney disease: Illuminating the path to precision medicine'; gemma_q0_document_type=REVIEW; gemma_final=FILTERED. | accepted_agree |
| 10.46421/entac.v18i.749 | 202 | AVALIAÇÃO DE ELEMENTOS DE CONTROLE SOLAR MÓVEIS PROJETADOS COM MODELAGEM PARAMÉTRICA | AGREE | gemma_final: Pass-A abstract prefilter -> FILTERED; q1 panel-design fields are empty; q2 paper_type=OTHER. | accepted_agree |
| 10.47184/tp.2024.01.07 | 2024 | Integration of spatially resolved transcriptomics into pathological research: Opportunities and challenges | AGREE | gemma_final: Pass-A abstract prefilter -> FILTERED; q1 panel-design fields are empty; q2 | accepted_agree |

| DOI | Year | Title | Audit decision | Evidence (verbatim quote) | Validation status / scope flag |
|---|---|---|---|---|---|
| | | | | paper_type=OTHER, spatial=YES, panel_scope=UNCLEAR. | |
| 10.47570/joci.2024.002 | 2024 | ExSeq: Expansion Sequencing for Single-Cell Spatial Transcriptomics | AGREE | gemma_final: Pass-A abstract prefilter -> FILTERED; q1 panel-design fields are empty; q2 paper_type=COMP. | accepted_agree |
| 10.48550/arxiv.2005.03847 | 202 | Tree! I am no Tree! I am a Low Dimensional Hyperbolic Embedding | AGREE | Title: 'Tree! I am no Tree! I am a Low Dimensional Hyperbolic Embedding'; packet fields show paper_type=COMP and Gemma's FILTERED prefilter. | accepted_agree |
| 10.48550/arxiv.2104.10041 | 2021 | Particle swarm optimization in constrained maximum likelihood estimation a case study | AGREE | Title: 'Particle swarm optimization in constrained maximum likelihood estimation a case study'; packet fields show paper_type=COMP and Gemma's FILTERED prefilter. | accepted_agree |
| 10.48550/arxiv.2106.06468 | 2021 | Locally Sparse Neural Networks for Tabular Biomedical Data | AGREE | Title: 'Locally Sparse Neural Networks for Tabular Biomedical Data'; packet fields show paper_type=COMP and Gemma's FILTERED prefilter. | accepted_agree |
| 10.48550/arxiv.2112.13289 | 2021 | Prevalence Threshold and bounds in the Accuracy of Binary Classification Systems | AGREE | Title: 'Prevalence Threshold and bounds in the Accuracy of Binary Classification Systems'; packet fields show paper_type=COMP and Gemma's FILTERED prefilter. | accepted_agree |
| 10.48550/arxiv.2204.09673 | 2022 | Guidelines for reporting cell types: the MIRACL standard | AGREE | Title: 'Guidelines for reporting cell types: the MIRACL standard'; packet fields show paper_type=OTHER, panel_scope=UNCLEAR, spatial=NO, and Gemma's FILTERED prefilter. | accepted_agree |
| 10.48550/arxiv.2210.02672 | 2023 | Orthogonal Non-negative Matrix Factorization: a Maximum-Entropy-Principle Approach | AGREE | Title only; paper_type COMP; spatial=NO; panel_scope=UNCLEAR; q1 panel-design fields are empty; gemma_final=FILTERED from abstract prefilter. | accepted_agree |
| 10.48550/arxiv.2301.00557 | 2023 | Learning to Maximize Mutual Information for Dynamic Feature Selection | AGREE | Title only; paper_type COMP; spatial blank; panel_scope blank; q1 panel-design fields are empty; gemma_final=FILTERED from citation-screening prefilter. | accepted_agree |

| DOI | Year | Title | Audit decision | Evidence (verbatim quote) | Validation status / scope flag |
|---|---|---|---|---|---|
| 10.48550/arxiv.2301.08436 | 2023 | SpaceTx: A Roadmap for Benchmarking Spatial Transcriptomics Exploration of the Brain | AGREE | Citation prior labels the item REV; title is 'SpaceTx: A Roadmap for Benchmarking Spatial Transcriptomics Exploration of the Brain'; q1 panel-design fields are empty. | accepted_agree |
| 10.48550/arxiv.2305.18962 | 2023 | Hyperbolic Diffusion Embedding and Distance for Hierarchical Representation Learning | AGREE | Title only; paper_type COMP; spatial blank; panel_scope blank; q1 panel-design fields are empty; gemma_final=FILTERED from citation-screening prefilter. | accepted_agree |
| 10.48550/arxiv.2312.06082 | 2023 | XAI meets Biology: A Comprehensive Review of Explainable AI in Bioinformatics Applications | AGREE | Title only; gemma_final notes labels=['REV']; exp_used_tools=['(none)']; all_tools=['SMaSH']; q1 panel-design fields are empty. | accepted_agree |
| 10.48550/arxiv.2410.21107 | 2024 | Tree-Wasserstein Distance for High Dimensional Data with a Latent Feature Hierarchy | AGREE | Title only; paper_type COMP; spatial blank; panel_scope blank; q1 panel-design fields are empty; gemma_final=FILTERED from abstract prefilter. | accepted_agree |
| 10.48550/arxiv.2503.00174 | 2025 | Optimal Transfer Learning for Missing Not-at-Random Matrix Completion | AGREE | paper.title='Optimal Transfer Learning for Missing Not-at-Random Matrix Completion'; gemma_final=FILTERED. | accepted_agree |
| 10.48550/arxiv.2506.10037 | 2025 | The Cell Ontology in the age of single-cell omics | AGREE | title: The Cell Ontology in the age of single-cell omics; q2 paper_type=OTHER, panel_scope=UNCLEAR, spatial=NO; q1 panel-design fields are empty. | accepted_agree |
| 10.48550/arxiv.2512.17678 | 2025 | You Only Train Once: Differentiable Subset Selection for Omics Data | AGREE | paper.title='You Only Train Once: Differentiable Subset Selection for Omics Data'; gemma_final=FILTERED. | accepted_agree |
| 10.53771/ijlsra.2025.9.1.0045 | 2025 | Unraveling tumor microenvironments with spatial genomics: Implications for precision breast cancer therapy | AGREE | Title only; paper_type=OTHER; spatial=YES; panel_scope=UNCLEAR; q1 panel-design fields are empty; gemma_final=FILTERED from abstract prefilter. | accepted_agree |
| 10.5525/gla.thesis.85717 | 2026 | At the frontier of immunology: exploring cellular crosstalk across time and space | AGREE | Title only; paper_type=COMP; spatial=YES; panel_scope=UNCLEAR; q1 panel-design fields are empty; | accepted_agree |

| DOI | Year | Title | Audit decision | Evidence (verbatim quote) | Validation status / scope flag |
|---|---|---|---|---|---|
| | | | | gemma_final=FILTERED from abstract prefilter. | |
| 10.55627/mmc.002.002.0160 | 2022 | The Neurons That Restore Walking After Paralysis | AGREE | paper_type/doc_type evidence: EDITORIAL (Editor’s Choice Editor's Selection of the Important Research Investigations in the Field of Molecular Medicine Communications from Around the World); gemma_final notes: Q0: EDITORIAL → FILTERED | accepted_agree |
| 10.56093/sr.v53i2.1 | 2025 | Application of Next-Generation Sequencing (NGS) Technologies for Efficient Breeding, Quality Assurance and Testing of Seeds | AGREE | Title only; paper_type=OTHER; spatial=UNCLEAR; panel_scope=UNCLEAR; q1 panel-design fields are empty; gemma_final=FILTERED from abstract prefilter. | accepted_agree |
| 10.59717/j.xinn-life.2023.100006 | 2023 | Spatial transcriptomics in cancer research: Opportunities and challenges | AGREE | Title only; paper_type=OTHER; spatial=YES; panel_scope=UNCLEAR; q1 panel-design fields are empty; gemma_final=FILTERED from abstract prefilter. | accepted_agree |
| 10.6019/tol.spatial-transcriptomics-neuro-w.2026.00001.1 | 2026 | Integrating single-cell and spatial transcriptomics to map the rules of neurological conditions | AGREE | Title only; paper_type=OTHER; spatial=YES; panel_scope=UNCLEAR; q1 panel-design fields are empty; gemma_final=FILTERED from abstract prefilter. | accepted_agree |
| 10.6084/m9.figshare.31311538.v1 | 2026 | Additional file 1 of Neighborhood nonnegative matrix factorization identifies patterns and spatially-variable genes in large-scale spatial transcriptomics data | AGREE | Title only; paper_type=OTHER; spatial=YES; panel_scope=TARGETED; q1 panel-design fields are empty; gemma_final=FILTERED from abstract prefilter. | accepted_agree |
| 10.61173/tsjz3r50 | 2025 | Analysis of Research Methods for Heterogeneous Cellular Senescence Biomarkers | AGREE | Title only; paper_type=OTHER; spatial=YES; panel_scope=TARGETED; q1 panel-design fields are empty; gemma_final=FILTERED from abstract prefilter. | accepted_agree |

| DOI | Year | Title | Audit decision | Evidence (verbatim quote) | Validation status / scope flag |
|---|---|---|---|---|---|
| 10.61373/gp025k.0062 | 2025 | Xuyu Qian: Understanding human brain development and diseases using human-based approaches | AGREE | Title only; paper_type=OTHER; spatial=YES; panel_scope=TARGETED; q1 panel-design fields are empty; gemma_final=FILTERED from abstract prefilter. | accepted_agree |
| 10.61841/nn-pbs-12-1-1 | 2026 | SPACESEQUEST: A UNIFIED PIPELINE FOR SPATIAL TRANSCRIPTOMICS DATA ANALYSIS | AGREE | Title only; paper_type=COMP; spatial=YES; panel_scope=UNCLEAR; q1 panel-design fields are empty; gemma_final=FILTERED from abstract prefilter. | accepted_agree |
| 10.64898/2025.12.01.691603 | 2025 | BioLACE: unifying spatial geometry and marker priors for cohesive cell-type clustering in spatial transcriptomics | AGREE | Title only; paper_type=COMP; spatial=YES; panel_scope=UNCLEAR; q1 panel-design fields are empty; gemma_final=FILTERED from abstract prefilter. | accepted_agree |
| 10.64898/2025.12.09.693237 | 2025 | Riemannian Metric Learning for Alignment of Spatial Multiomics | AGREE | Title only; paper_type=COMP; spatial=YES; panel_scope=UNCLEAR; q1 panel-design fields are empty; gemma_final=FILTERED from abstract prefilter. | accepted_agree |
| 10.64898/2025.12.09.693287 | 2025 | NicheAgent: LLM-Guided Zero-Shot Niche Identification for Spatial Transcriptomics | AGREE | Title only; paper_type=COMP; spatial=YES; panel_scope=UNCLEAR; q1 panel-design fields are empty; gemma_final=FILTERED from abstract prefilter. | accepted_agree |
| 10.64898/2025.12.11.693759 | 2025 | MisTIC: Missegmented Transcript Inference Correction for Improved Spatial Transcriptomics Analysis | AGREE | gemma_final notes say Pass-A abstract: COMP -> FILTERED; q2 paper_type=COMP, spatial=YES, panel_scope=UNCLEAR; q1 panel-design fields are empty. | accepted_agree |
| 10.64898/2025.12.15.694341 | 2025 | STUltra: scalable and accurate integration for subcellular-level spatial omics data | AGREE | Title only; paper_type=COMP; spatial=YES; panel_scope=UNCLEAR; q1 panel-design fields are empty; gemma_final=FILTERED from abstract prefilter. | accepted_agree |
| 10.64898/2025.12.22.696108 | 2025 | FlashDeconv enables atlas-scale, multi-resolution spatial deconvolution via structure-preserving sketching | AGREE | Title only; paper_type=COMP; spatial=YES; panel_scope=NOT_TARGETED; q1 panel-design fields are empty; | accepted_agree |

| DOI | Year | Title | Audit decision | Evidence (verbatim quote) | Validation status / scope flag |
|---|---|---|---|---|---|
| | | | | gemma_final=FILTERED from abstract prefilter. | |
| 10.64898/2025.12.28.696778 | 2025 | LLM-scCurator: Data-centric feature distillation for zero-shot cell-type annotation | AGREE | Title only; paper_type=COMP; spatial=YES; panel_scope=UNCLEAR; q1 panel-design fields are empty; gemma_final=FILTERED from abstract prefilter. | accepted_agree |
| 10.64898/2026.01.07.697982 | 2026 | CellWHISPER disentangles direct cell–cell communication from structural proximity | AGREE | Title only; paper_type=COMP; spatial=YES; panel_scope=UNCLEAR; q1 panel-design fields are empty; gemma_final=FILTERED from abstract prefilter. | accepted_agree |
| 10.64898/2026.01.15.699623 | 2026 | DeepSpaceDB 2.0: an interactive spatial transcriptomics database for large-scale Xenium data exploration | AGREE | Title only; paper_type=OTHER; spatial=YES; panel_scope=TARGETED; q1 panel-design fields are empty; gemma_final=FILTERED from abstract prefilter. | accepted_agree |
| 10.64898/2026.01.26.701684 | 2026 | ESFS: A Noise-Resilient Framework for Feature Selection and Marker Gene Discovery in Single-Cell Transcriptomics | AGREE | paper.title='ESFS: A Noise-Resilient Framework for Feature Selection and Marker Gene Discovery in Single-Cell Transcriptomics'; citation_screening_prior.labels='COMP'; gemma_final.classification='FILTERED'; gemma_q1_panel_design blank; gemma_q2_paper_profile={'evidence_scope': '', 'evidence_spatial': '', 'evidence_type': '', 'panel_scope': '', 'paper_type': 'COMP', 'profile_confidence': '', 'spatial': ''} | accepted_agree |
| 10.64898/2026.02.09.704806 | 2026 | Parsimonious cell co-localization scoring for spatial transcriptomics | AGREE | q2 describes a neighborhood-product co-localization score on a pruned spatial neighbor graph. | accepted_agree |
| 10.64898/2026.02.12.703707 | 2026 | A stem cell aging clock links biological age deviation to clinical outcome in acute myeloid leukemia | AGREE | paper.title='A stem cell aging clock links biological age deviation to clinical outcome in acute myeloid leukemia'; citation_screening_prior.labels='COMP'; gemma_final.classification='FILTERED'; gemma_q1_panel_design | accepted_agree |

| DOI | Year | Title | Audit decision | Evidence (verbatim quote) | Validation status / scope flag |
|---|---|---|---|---|---|
| | | | | blank; gemma_q2_paper_profile= {'evidence_scope': '', 'evidence_spatial': '', 'evidence_type': '', 'panel_scope': '', 'paper_type': 'COMP', 'profile_confidence': '', 'spatial': ''} | |
| 10.64898/2026.02.13.705867 | 2026 | jazzPanda: A hybrid approach to find spatial marker genes in imaging-based spatial transcriptomics data | NEEDS_MANUAL | PDF: jazzPanda finds spatial marker genes via a hybrid analytical approach. | pdf_validated_unclear_update |
| 10.64898/2026.02.18.706570 | 2026 | Spartan: Spatial Activation Aware Transcriptomic Analysis Network | AGREE | Title only; paper_type=COMP; spatial=YES; panel_scope=UNCLEAR; q1 panel-design fields are empty; gemma_final=FILTERED from abstract prefilter. | accepted_agree |
| 10.64898/2026.02.18.706600 | 2026 | Comparative Biology at Single-Cell Resolution: Rigorous Matching of Atlases for Cross-Species Analysis | AGREE | paper.title='Comparative Biology at Single-Cell Resolution: Rigorous Matching of Atlases for Cross-Species Analysis'; citation_screening_prior.labels='COMP'; gemma_final.classification='FILTERED'; gemma_q1_panel_design blank; gemma_q2_paper_profile= {'evidence_scope': '', 'evidence_spatial': '', 'evidence_type': '', 'panel_scope': '', 'paper_type': 'COMP', 'profile_confidence': '', 'spatial': ''} | accepted_agree |
| 10.64898/2026.02.19.706743 | 2026 | OT-knn: a neighborhood-aware optimal transport framework for aligning spatial transcriptomics data | AGREE | title: OT-knn: a neighborhood-aware optimal transport framework for aligning spatial transcriptomics data; q0 notes: Pass-A abstract: COMP → FILTERED; q1 panel-design fields are empty; q2 panel_scope=UNCLEAR; q2 spatial=YES; q2 paper_type=COMP. | accepted_agree |
| 10.64898/2026.02.21.707168 | 2026 | Pixel2Gene enables histology-guided reconstruction and prediction of spatial gene expression | AGREE | title: Pixel2Gene enables histology-guided reconstruction and prediction of spatial gene expression; q0 notes: Pass-A abstract: COMP → FILTERED; q1 panel-design fields are empty; q2 panel_scope=UNCLEAR; q2 | accepted_agree |

| DOI | Year | Title | Audit decision | Evidence (verbatim quote) | Validation status / scope flag |
|---|---|---|---|---|---|
| | | | | spatial=YES; q2 paper_type=COMP. | |
| 10.64898/2026.02.23.705895 | 2026 | RSTG: Robust Generation of High Quality Spatial Transcriptomics Data using Beta Divergence Based AutoEncoder | AGREE | title: RSTG: Robust Generation of High Quality Spatial Transcriptomics Data using Beta Divergence Based AutoEncoder; q0 notes: Pass-A abstract: COMP → FILTERED; q1 panel-design fields are empty; q2 panel_scope=UNCLEAR; q2 spatial=YES; q2 paper_type=COMP. | accepted_agree |
| 10.64898/2026.02.25.707866 | 2026 | Information Geometry Reconciles Discrete and Continuous Variation in Single-Cell and Spatial Transcriptomic Analysis | AGREE | title: Information Geometry Reconciles Discrete and Continuous Variation in Single-Cell and Spatial Transcriptomic Analysis; q2 paper_type=COMP; q1 panel-design fields are empty; citation prior labels=COMP and Gemma_final=FILTERED. | accepted_agree |
| 10.64898/2026.02.26.705527 | 2026 | Protein-guided RNA barcoding links transcriptomes to synaptic architecture | NEEDS_MANUAL | panel_scope=NOT_TARGETED with no target-gene panel source evidence; untargeted/no-panel records are out of scope. Original audit evidence: q1: panel_design=UNCLEAR; evidence_tool/evidence_manual/evidence_vendor are NOT FOUND; q2: panel_scope=NOT_TARGETED. | validated_high_risk_manual_filtered |
| 10.64898/2026.02.26.707418 | 2026 | SpatialCompassV (SCOMV): <i>De novo</i> cell and gene spatial pattern classification and spatially differential gene identification | NEEDS_MANUAL | PDF: SCOMV clusters genes/cells by spatial pattern and finds spatially differential genes. | pdf_validated_unclear_update |
| 10.64898/2026.02.26.707870 | 2026 | PantheonOS: An Evolvable Multi-Agent Framework for Automatic Genomics Discovery | AGREE | title: PantheonOS: An Evolvable Multi-Agent Framework for Automatic Genomics Discovery; q0 notes: citation-screening prior pre-filter: computational-only prior; q1 panel-design fields are empty; q2 panel_scope is not populated; title and prior do not show panel selection. | accepted_agree |
| 10.64898/2026.02.26.708370 | 2026 | STCS: A Platform-Agnostic Framework for Cell-Level | AGREE | title: STCS: A Platform-Agnostic Framework for Cell-Level | accepted_agree |

| DOI | Year | Title | Audit decision | Evidence (verbatim quote) | Validation status / scope flag |
|---|---|---|---|---|---|
| | | Reconstruction in Sequencing-Based Spatial Transcriptomics | | Reconstruction in Sequencing-Based Spatial Transcriptomics; q0 notes: Pass-A abstract: COMP → FILTERED; q1 panel-design fields are empty; q2 panel_scope=NOT_TARGETED; q2 spatial=YES; q2 paper_type=COMP. | |
| 10.64898/2026.03.01.708931 | 2026 | RankMap: Rank-based reference mapping for fast and robust cell type annotation in spatial and single-cell transcriptomics | AGREE | title: RankMap: Rank-based reference mapping for fast and robust cell type annotation in spatial and single-cell transcriptomics; q0 notes: Pass-A abstract: COMP → FILTERED; q1 panel-design fields are empty; q2 panel_scope=UNCLEAR; q2 spatial=YES; q2 paper_type=COMP. | accepted_agree |
| 10.64898/2026.03.12.711372 | 2026 | Multi-scale spatial testing recovers gene programs missed by existing detection methods | AGREE | title: Multi-scale spatial testing recovers gene programs missed by existing detection methods; q0 notes: Pass-A abstract: COMP → FILTERED; q1 panel-design fields are empty; q2 panel_scope=UNCLEAR; q2 spatial=YES; q2 paper_type=COMP. | accepted_agree |
| 10.64898/2026.03.17.712359 | 2026 | STiLE: Automated Tissue Microarray Dearraying for Spatial Transcriptomics | AGREE | title: STiLE: Automated Tissue Microarray Dearraying for Spatial Transcriptomics; q0 notes: Pass-A abstract: COMP → FILTERED; q1 panel-design fields are empty; q2 panel_scope=UNCLEAR; q2 spatial=YES; q2 paper_type=COMP. | accepted_agree |
| 10.64898/2026.03.24.713944 | 2026 | Allos: an integrated Python toolkit for isoform-level single-cell and spatial in-situ transcriptomics | AGREE | title: Allos: an integrated Python toolkit for isoform-level single-cell and spatial in-situ transcriptomics; q0 notes: Pass-A abstract: COMP → FILTERED; q1 panel-design fields are empty; q2 panel_scope=UNCLEAR; q2 spatial=YES; q2 paper_type=COMP. | accepted_agree |

| DOI | Year | Title | Audit decision | Evidence (verbatim quote) | Validation status / scope flag |
|---|---|---|---|---|---|
| 10.64898/2026.03.25.713520 | 2026 | CosMxScope: Scalable Reconstruction and Digital Pathology Integration of Imaging-Based Spatial Transcriptomics Data | NEEDS_MANUAL | PDF: CosMxScope reconstructs whole-slide images and visualizes transcript coordinates. | pdf_validated_unclear_update |
| 10.64898/2026.03.26.714500 | 2026 | Single-cell lung eQTL dataset of Asian never-smokers highlights the roles of alveolar cells in lung cancer etiology | AGREE | paper.title='Single-cell lung eQTL dataset of Asian never-smokers highlights the roles of alveolar cells in lung cancer etiology'; citation_screening_prior.labels='NON-ST'; gemma_final.classification='FILTERED'; gemma_q1_panel_design blank; gemma_q2_paper_profile={'evidence_scope': '', 'evidence_spatial': '', 'evidence_type': '', 'panel_scope': '', 'paper_type': '', 'profile_confidence': '', 'spatial': 'NO'} | accepted_agree |
| 10.64898/2026.03.28.715026 | 2026 | GraphBG: Fast Bayesian Domain Detection via Spectral Graph Convolutions for Multi-slice and Multi-modal Spatial Transcriptomics | AGREE | title: GraphBG: Fast Bayesian Domain Detection via Spectral Graph Convolutions for Multi-slice and Multi-modal Spatial Transcriptomics; q0 notes: Pass-A abstract: COMP → FILTERED; q1 panel-design fields are empty; q2 panel_scope=UNCLEAR; q2 spatial=YES; q2 paper_type=COMP. | accepted_agree |
| 10.7150/ijbs.117249 | 2025 | Spatial Cell Atlas of Lateral Septum Reveals Changes Underlying Anxiety and Fear Learning Deficits in Mice with Abnormal Immunity. | NEEDS_MANUAL | Final manual decision: FILTERED because manual review found no usable evidence for final gene-panel source selection. Original evidence: Integrating multiplexed error-robust fluorescence in situ hybridization (MERFISH) data of the entire LS | manual_final_filter_after_pdf_review |
| 10.7150/ijbs.43887 | 202 | Encoding method of single-cell spatial transcriptomics sequencing | AGREE | title: Encoding method of single-cell spatial transcriptomics sequencing; q0 notes: Pass-A abstract: OTHER → FILTERED; q1 panel-design fields are empty; q2 panel_scope=UNCLEAR; q2 spatial=UNCLEAR; q2 paper_type=OTHER. | accepted_agree |

| DOI | Year | Title | Audit decision | Evidence (verbatim quote) | Validation status / scope flag |
|---|---|---|---|---|---|
| 10.7150/thno.112339 | 2025 | Pharmacological and genetic inhibition of ARG2 in CXCR2<sup>Hi</sup> myeloid-derived suppressor cells combats sepsis-induced lymphopenia | AGREE | paper.title='Pharmacological and genetic inhibition of ARG2 in CXCR2<sup>Hi</sup> myeloid-derived suppressor cells combats sepsis-induced lymphopenia'; gemma_q2_paper_profile.spatial=NO; gemma_final=FILTERED. | accepted_agree |
| 10.7150/thno.99046 | 2024 | <i>DAB2</i><sup>+</sup> macrophages support <i>FAP</i><sup>+</sup> fibroblasts in shaping tumor barrier and inducing poor clinical outcomes in liver cancer | NEEDS_MANUAL | panel_scope=NOT_TARGETED with no target-gene panel source evidence; untargeted/no-panel records are out of scope. Original audit evidence: No panel-design evidence is present in the packet. | validated_high_risk_manual_filtered |
| 10.7185/gold2021.8278 | 2021 | Stable coexistence of aerobic and anaerobic transition | AGREE | title: Stable coexistence of aerobic and anaerobic transition; q0 notes: Q0 document type pre-filter marked NON_ARTICLE and FILTERED; q1 panel-design fields are empty; there is no panel-selection evidence. | accepted_agree |
| 10.7287/peerj.preprints.1388v2 | 2015 | Relationship between small-scale catch-per-unit-effort and abundance in New Zealand abalone (pāua, <i>Haliotis iris</i> ) fisheries | AGREE | title: Relationship between small-scale catch-per-unit-effort and abundance in New Zealand abalone (pāua, Haliotis iris) fisheries; q0 notes: Pass-A abstract: non-spatial → FILTERED; q1 panel-design fields are empty; q2 spatial=NO; q2 paper_type=EXP. | accepted_agree |
| 10.7302/25075 | 2024 | Computational Analysis and Statistical Methods in Spatial Transcriptomics and Neuroimaging | AGREE | title: Computational Analysis and Statistical Methods in Spatial Transcriptomics and Neuroimaging; q0 notes: Pass-A abstract: COMP → FILTERED; q1 panel-design fields are empty; q2 panel_scope=UNCLEAR; q2 spatial=YES; q2 paper_type=COMP. | accepted_agree |
| 10.7506/spkx1002-6630-20200419-250 | 2021 | Changes in the Quality and Protein Properties of Seasoned Fish during Repeated Freeze-Thaw Cycles | AGREE | title: Changes in the Quality and Protein Properties of Seasoned Fish during Repeated Freeze-Thaw Cycles; q0 notes: Pass-A abstract: non-spatial → FILTERED; q1 panel-design fields are empty; q2 spatial=NO; q2 paper_type=EXP. | accepted_agree |

| DOI | Year | Title | Audit decision | Evidence (verbatim quote) | Validation status / scope flag |
|---|---|---|---|---|---|
| 10.7554/elife.100796 | 2025 | Single-cell RNA sequencing of the holothurian regenerating intestine reveals the pluripotency of the coelomic epithelium. | AGREE | title: Single-cell RNA sequencing of the holothurian regenerating intestine reveals the pluripotency of the coelomic epithelium.; q0 notes: Pass-A abstract: non-spatial → FILTERED; q1 panel-design fields are empty; q2 spatial=NO; q2 paper_type=EXP. | accepted_agree |
| 10.7554/elife.105149 | 2025 | MerQuaCo: a computational tool for quality control in image-based spatial transcriptomics | NEEDS_MANUAL | Final manual decision: FILTERED because no local PDF was available and the packet did not provide usable final gene-panel source evidence. Original evidence: No gene-panel selection evidence found in the packet. | manual_final_filter_after_pdf_review |
| 10.7554/elife.105225 | 2025 | Modular DNA barcoding of nanobodies enables multiplexed in situ protein imaging and high-throughput biomolecule detection. | AGREE | title: Modular DNA barcoding of nanobodies enables multiplexed in situ protein imaging and high-throughput biomolecule detection.; q0 notes: Pass-A abstract: COMP → FILTERED; q1 panel-design fields are empty; q2 spatial=NO; q2 paper_type=COMP. | accepted_agree |
| 10.7554/elife.107531 | 2025 | CellCover Defines Marker Gene Panels Capturing Developmental Progression in Neocortical Neural Stem Cell Identity | AGREE | gemma_final: Pass-A abstract prefilter -> FILTERED; q1 panel-design fields are empty; q2 paper_type=COMP, spatial=NO, panel_scope=UNCLEAR. | accepted_agree |
| 10.7554/elife.43627 | 2019 | Flura-seq identifies organ-specific metabolic adaptations during early metastatic colonization. | AGREE | title: Flura-seq identifies organ-specific metabolic adaptations during early metastatic colonization.; q0 notes: Pass-A abstract: not targeted → FILTERED; q1 panel-design fields are empty; q2 spatial=YES; q2 paper_type=EXP. | accepted_agree |
| 10.7554/elife.55608 | 202 | <i>Fgf4</i> maintains <i>Hes7</i> levels critical for normal somite segmentation clock function. | NEEDS_MANUAL | Final manual decision: FILTERED because manual review found no usable evidence for final gene-panel source selection. Original evidence: We analyzed Fgf4 mutants by quantifying mRNAs fluorescently labeled by hybridization chain reaction | manual_final_filter_after_pdf_review |
| 10.7606/j.issn.1000-4025.2020.04.0588 | 202 | Genome Comparison between Elymus nutans Griseb. and E. | AGREE | title: Genome Comparison between Elymus nutans Griseb. | accepted_agree |

| DOI | Year | Title | Audit decision | Evidence (verbatim quote) | Validation status / scope flag |
|---|---|---|---|---|---|
| | | dahuricus Turcez. ex Griseb. with Cytogenetics | | and E. dahuricus Turcez. ex Griseb. with Cytogenetics; q0 notes: Pass-A abstract: non-spatial → FILTERED; q1 panel-design fields are empty; q2 spatial=NO; q2 paper_type=EXP. | |
| 10.7717/peerj.16192 | 2023 | SaPt-CNN-LSTM-AR-EA: a hybrid ensemble learning framework for time series-based multivariate DNA sequence prediction | AGREE | Title only; paper_type COMP; spatial blank; panel_scope blank; q1 panel-design fields are empty; gemma_final=FILTERED from citation-screening prefilter. | accepted_agree |
| 10.7759/cureus.86815 | 2025 | A Case of Pediatric Renal Cell Carcinoma With TFE3 Rearrangement Treated With Robot-Assisted Partial Nephrectomy | AGREE | title: A Case of Pediatric Renal Cell Carcinoma With TFE3 Rearrangement Treated With Robot-Assisted Partial Nephrectomy; q0 notes: Pass-A abstract: non-spatial → FILTERED; q1 panel-design fields are empty; q2 spatial=NO; q2 paper_type=EXP. | accepted_agree |
| 10.7860/njlm/2025/74501.2896 | 2025 | A Cross-sectional Study on Molecular Cartography: The Mapping of Down Syndrome with Cytogenetic Tools | AGREE | Paper title is about a cross-sectional study on molecular cartography with cytogenetic tools; q2 says spatial=NO and q1 panel-design fields are empty. | accepted_agree |
