## supplementary_methods_s1_doi_manifest_queries for "Lack of adoption of computational gene-panel-selection tools in targeted spatial transcriptomics"

### Supplementary Methods S1: Tool DOI manifest and citation-harvest query specifications

This document specifies (i) the canonical published and preprint DOIs for the 18 computational gene-panel selection tools reviewed in this manuscript, and (ii) the exact OpenAlex and Semantic Scholar queries used to harvest citing papers for each DOI. The intent is that any reader with the listed DOIs and the queries below can reproduce the citation harvest precisely (modulo any new citations accumulated after the harvest freeze date of 17 April 2026).

#### A. Canonical DOI manifest for the 18 tools

The manifest lists, per tool, the published DOI (when one exists at the time of the harvest), the preprint DOI (always present), and the preprint deposit date. Categories follow the manuscript: spatially-motivated tools (Spapros, gpsFISH, PERSIST, scGIST, ReconST, scGPD, panelScope, geneBasis, geneCover, scMAGS) and general marker-selection tools (the remaining 8). Where a published DOI is missing (ReconST, scGPD), only the preprint DOI is used for the citation query.

| Tool | Category | Published DOI | Preprint DOI | Preprint date |
|---|---|---|---|---|
| scGeneFit | general marker-selection | `10.1038/s41467-021-21453-4` | `10.1101/599654` | 2019-xx-xx |
| RankCorr | general marker-selection | `10.1186/s12859-020-03641-z` | `10.1101/679761` | 2019-xx-xx |
| SCMER | general marker-selection | `10.1038/s43588-021-00070-7` | `10.1101/2020.12.01.407262` | 2020-12-01 |
| scPNMF | general marker-selection | `10.1093/bioinformatics/btab273` | `10.1101/2021.02.09.430550` | 2021-02-09 |
| SMaSH | general marker-selection | `10.1186/s12859-022-04860-2` | `10.1101/2021.04.08.438978` | 2021-04-08 |
| COSG | general marker-selection | `10.1093/bib/bbab579` | `10.1101/2021.06.15.448484` | 2021-06-15 |
| geneBasis | spatially-motivated | `10.1186/s13059-021-02548-z` | `10.1101/2021.08.10.455720` | 2021-08-10 |
| scMAGS | spatially-motivated | `10.1016/j.compbiomed.2023.106634` | `10.1101/2022.03.22.485261` | 2022-03-22 |
| PERSIST | spatially-motivated | `10.1038/s41467-023-37392-1` | `10.1101/2022.05.13.491738` | 2022-05-13 |
| Spapros | spatially-motivated | `10.1038/s41592-024-02496-z` | `10.1101/2022.08.16.504115` | 2022-08-16 |
| scGIST | spatially-motivated | `10.1186/s13059-024-03185-y` | `10.1101/2022.09.25.509418` | 2022-09-25 |

| Tool | Category | Published DOI | Preprint DOI | Preprint date |
|---|---|---|---|---|
| gpsFISH | spatially-motivated | 10.1186/s13059-024-03174-1 | 10.1101/2023.02.03.527053 | 2023-02-03 |
| NS-Forest v2 | general marker-selection | 10.1101/gr.275569.121 | 10.1101/2020.09.23.308932 | 2020-09-23 |
| NS-Forest v4.0 | general marker-selection | 10.1186/s44330-024-00015-2 | 10.1101/2024.04.22.590194 | 2024-04-22 |
| geneCover | spatially-motivated | 10.1101/gr.280539.125 | 10.1101/2024.10.30.621151 | 2024-10-30 |
| panelScope | spatially-motivated | 10.1093/bib/bbaf478 | 10.1101/2025.02.19.639174 | 2025-02-19 |
| scGPD | spatially-motivated | – | 10.1101/2025.10.02.680117 | 2025-10-02 |
| ReconST | spatially-motivated | – | 10.1101/2025.10.08.681071 | 2025-10-08 |

The canonical machine-readable form of this table is manuscript/tables/ST_gene_selection_tools.csv.

#### B. DOI normalization

DOIs are normalised before all lookups and deduplication: convert to lowercase, strip surrounding whitespace, and remove any leading 'https://doi.org/' or 'doi:' prefix. A canonical DOI is the lowercase form without any URL prefix (e.g. '10.1186/s13059-021-02548-z').

#### C. OpenAlex citation query

##### C.1 DOI → OpenAlex short id

Resolve a tool's published or preprint DOI to its OpenAlex short work id (e.g. W1234567890). The short id is then used as the cited work in the citation query below. Both the published DOI and the preprint DOI are resolved when both exist.

**Endpoint:** GET https://api.openalex.org/works/doi:<normalized_doi>

**Params:**

```
api_key = <OA_API_KEY>
select = id
```

##### C.2 Citing-papers query

**Endpoint:** GET https://api.openalex.org/works

**Params:**

```
api_key = <OA_API_KEY>
filter = cites:<openalex_short_id>
select =
```

```
id,doi,title,publication_year,type,abstract_inverted_index,open_access,primary_location
per_page = 200
cursor = * (initial); next_cursor for each subsequent page
```

**Pagination.** Cursor-paginated. The first request uses cursor=*; subsequent requests use meta.next_cursor returned by the previous response. Iteration stops when results is empty or next_cursor is missing.

**Rate.** 100 ms sleep between pages.

**Fields used (per returned citing record):**

- id (OpenAlex short id, used as oa_id)
- doi (used as the citing-paper DOI after normalisation)
- title
- publication_year
- type (used as oa_type)
- abstract_inverted_index (reconstructed to plain text abstract)
- open_access.oa_url (used as oa_pdf_url)
- primary_location.source.display_name (used as oa_journal)

**DOI-less records.** Records returned without a DOI are kept under key ‘no_doi:’ so they remain identifiable for downstream review.

#### D. Semantic Scholar citation query

##### D.1 DOI → S2 paperId (with search fallback)

Resolve a tool’s published or preprint DOI to a Semantic Scholar paperId. The primary path uses the direct DOI: endpoint. If that returns 404 (some publisher and preprint DOIs are not indexed under the DOI key but are findable by full-text search), the fallback path searches /paper/search and accepts the first hit whose externalIds.DOI matches the normalised query DOI exactly. This dual lookup is important for PRISMA completeness because S2 indexing of biorxiv / chemrxiv / publisher DOIs is uneven.

**Primary endpoint:** `GET https://api.semanticscholar.org/graph/v1/paper/DOI:<normalized_doi>`

**Primary params:**

```
fields = paperId,externalIds
header = x-api-key: <S2_API_KEY>
```

**Fallback endpoint (on 404 or empty response):** `GET https://api.semanticscholar.org/graph/v1/paper/search`

**Fallback params:**

```
query = <normalized_doi>
fields = paperId,externalIds
limit = 5
header = x-api-key: <S2_API_KEY>
```

##### D.2 Citing-papers query

**Endpoint:** `GET https://api.semanticscholar.org/graph/v1/paper/<paperId>/citations`

**Params:**

```
fields = paperId,externalIds,title,year,publicationTypes,abstract,openAccessPdf,journal,venue
limit = 1000
offset = 0, 1000, 2000, ... (incremented per page)
header = x-api-key: <S2_API_KEY>
```

**Pagination.** Offset-paginated. Each page requests limit=1000 and the offset is incremented by the size of the previous page. Iteration stops when the returned data array is empty.

**Fields used (per returned citing record):**

- citingPaper.paperId (used as s2_id)
- citingPaper.externalIds (DOI extracted and normalised)
- citingPaper.title
- citingPaper.year
- citingPaper.journal / venue (combined into journal field)
- citingPaper.publicationTypes (used as s2_pub_types)
- citingPaper.abstract
- citingPaper.openAccessPdf.url (used as s2_pdf_url)

**DOI-less records.** Records returned without a DOI are kept under key ‘no_doi_s2:’.

#### E. Per-tool deduplication and version handling

After harvest, citing-paper records are deduplicated within each tool by normalised DOI. Same-tool preprint/published version pairs are then collapsed using a fuzzy-title similarity threshold of 0.80 (VERSION_DEDUP_SIMILARITY in citing_papers.py). Cross-tool duplicates are intentionally NOT deduplicated at this stage so that multi-tool citation records remain visible; the unique-paper count is computed by collapsing on normalised DOI across the full harvest.
