## supplementary_methods_s2_citation_prompts for "Lack of adoption of computational gene-panel-selection tools in targeted spatial transcriptomics"

### Supplementary Methods S2: Citation classification prompts

This document reproduces the deterministic pre-filter rules, the prompt templates, and the structured-output schema used to classify each tool-paper citation record. Classification ran in a deterministic pre-filter followed by two model stages:

1. A deterministic metadata and title pre-filter assigned a final label to records that could be typed unambiguously from structured metadata or the title alone, with no model call (Section S2.1).
2. A local Gemma4 model (served through Ollama) classified each remaining record's document type, panel-selection use of the cited tool, and primary paper profile, using only the extracted PDF text (Section S2.2).
3. An independent GPT-5.4 audit reviewed every model classification against a self-contained audit packet and either agreed, disagreed, or routed the record to manual review (Section S2.3).

Both model stages ran in JSON mode and required verbatim PDF quotes for every evidence field. The Gemma4 prompts are reproduced verbatim from the source classifier; the GPT-5.4 audit instructions are reproduced verbatim from the audit packets actually sent to the model (using the first packet as the canonical example).

#### S2.1 Deterministic metadata and title pre-filter

Before any model call, each tool-paper record was tested against a deterministic pre-filter that assigns a final label from structured metadata or the title alone. The pre-filter is conservative by design: it fires only on unambiguous non-primary article types and otherwise returns control to the model stage (Section S2.2). Two checks were applied in order; the first to match assigns the label.

**(1) Metadata check.** Using the OpenAlex work type (`oa_type`) and the Semantic Scholar publication types (`s2_pub_types`):

- → **REV** if the OpenAlex type is `review`, or the Semantic Scholar types contain `Review` but not `JournalArticle` (the second condition avoids reclassifying primary research that some sources dual-tag as a review).
- → **OTHER** if the OpenAlex type is `book-chapter` or `dissertation`, or the Semantic Scholar types contain `Book` or `ClinicalTrial`.

**(2) Title check.** Applied only when the metadata check does not fire, using case-insensitive regular expressions anchored to the start of the title. A match on the correction pattern assigns **OTHER**; a match on the editorial pattern assigns **REV**:

```
correction / retraction → OTHER:
^\s*(correction|erratum|corrigendum|retraction|withdrawn|expression of concern)\b

editorial / special-issue → REV:
^\s*(editorial|preface|foreword|in memoriam|introduction to the special issue)\b|special
issue\b
```

Pre-filter labels were emitted at high confidence and tagged with their basis (`metadata_prefilter` or `title_prefilter`) for traceability, and remained subject to the independent GPT-5.4 audit (Section S2.3), which could overturn an incorrect assignment (audit error code `metadata_prefilter_wrong`). Of 626 tool-paper records that entered citation screening, 86 were resolved before model classification — 77 by these deterministic rules (76 by metadata, 1 by title) and 9 by manual review — while the remaining 540 proceeded to Gemma4 classification (Section S2.2).

#### S2.2 Gemma4 citation-classification prompt

For each tool-paper record routed to full-text classification, the extracted PDF text was passed three times to Gemma4 (Ollama backend), once per sub-prompt below. Each sub-prompt requests a single JSON object matching the schema embedded in its header. ### S2.2.1 Document-type sub-prompt (`DOC_TYPE_PROMPT`)

```
You are classifying a scientific PDF using ONLY the provided full extracted PDF text.

Return one JSON object only with this exact schema:
{{
  "doc_type": "MAIN_ARTICLE|EDITORIAL|CORRECTION|SUPPLEMENT|UNCLEAR",
  "doc_confidence": "HIGH|LOW",
  "doc_evidence": "exact quote from the PDF text, or NOT FOUND"
}}

Decision rules:
- Use only the provided PDF text.
- doc_evidence must be a verbatim quote copied from the PDF text.
- If direct support is missing, use NOT FOUND and LOW confidence.

Document type:
- MAIN_ARTICLE: the primary research or review article itself
- EDITORIAL: editorial, preface, perspective, commentary, or special-issue introduction without
full research content
- CORRECTION: erratum, corrigendum, retraction, expression of concern, correction notice
- SUPPLEMENT: supplementary information, appendix, supplementary methods/tables/figures,
supporting information
- UNCLEAR: cannot determine reliably
```

##### S2.2.2 Tool-use sub-prompt (`TOOL_USE_PROMPT`)

```
You are judging whether a scientific PDF reports that the authors actually USED a specific
named tool for panel / marker / gene-set selection, using ONLY the provided full extracted PDF
text.

Return one JSON object only with this exact schema:
{{
  "tool_mention": "YES|NO",
  "panel_selection_use": "EXP_USE|BENCHMARK|NOT_USED|UNCERTAIN",
  "tool_confidence": "HIGH|LOW",
  "evidence_tool": "exact quote from the PDF text, or NOT FOUND",
  "evidence_use": "exact quote from the PDF text, or NOT FOUND"
}}

Target tool context:
{tool_context}

Decision rules:
- Use only the provided PDF text.
- All evidence fields must be verbatim quotes copied from the PDF text.
- tool_mention is YES only if the PDF text explicitly names the tool, an alias, or one of the
tool-paper DOIs.

panel_selection_use definitions (choose the single best fit):
- EXP_USE = the authors RAN the tool to design or select the gene/probe/marker panel that they
```

```
then USE in their own real spatial-transcriptomics (or adjacent wet-lab) experiments in this
paper. The resulting panel is applied to real biological or clinical samples.
  Example: "We selected 484 genes with gpsFISH and profiled them via DART-FISH on human kidney
sections."
  CRITICAL: Using the tool for routine scRNA-seq downstream analysis — such as finding top-N
DEGs per cluster for cell type annotation, or for any purpose that does NOT produce a
spatial/wet-lab gene panel — is NOT EXP_USE. Label those as NOT_USED.
  NOT EXP_USE examples: "we used COSG to find the top 50 marker genes of each cluster"; "marker
genes were identified with COSG for cell type annotation"; "we applied scGeneFit to identify
DEGs for clustering visualisation".
- BENCHMARK = the authors RAN the tool purely for EVALUATION / COMPARISON / CHARACTERIZATION
purposes — e.g. as a baseline competitor against their own new method, or to compare multiple
panel-selection tools against each other. The tool's output is not used to design a panel for a
real downstream wet-lab experiment in this paper.
  Examples: "we applied scGPD and three baseline gene selection methods: PERSIST, scGIST, and
geneBasis to identify gene panels...";
           "We compared the results of COSG, NSForest, scGeneFit, scMAGS, and our method to
evaluate their abilities to identify marker genes";
           "We conducted panel selection by following gpsFISH's vignette (n = 117) [to
characterize it with our panelScope framework]".
- NOT_USED = the tool is absent, cited-only, mentioned in passing, appears only in the
reference list / related-work / background, OR was run only for routine scRNA-seq analysis (DEG
finding, cluster annotation) with no spatial/wet-lab panel outcome.
- UNCERTAIN = the tool is mentioned and possibly run, but the text does not make clear whether
it was EXP_USE vs BENCHMARK, or whether it was run at all.

- Do not infer use from citation alone.
- If direct support is missing, use NOT FOUND and LOW confidence.
```

##### S2.2.3 Paper-profile sub-prompt (`PROFILE_PROMPT`)

```
You are classifying the primary type of a scientific PDF using ONLY the provided full extracted
PDF text.

Return one JSON object only with this exact schema:
{{
  "paper_type": "COMP|REV|EXP|OTHER|UNCLEAR",
  "spatial": "YES|NO|UNCLEAR",
  "profile_confidence": "HIGH|LOW",
  "evidence_type": "exact quote from the PDF text, or NOT FOUND",
  "evidence_spatial": "exact quote from the PDF text, or NOT FOUND"
}}

Decision rules:
- Use only the provided PDF text.
- All evidence fields must be verbatim quotes copied from the PDF text.

paper_type definitions (choose the single primary contribution):
- COMP = the paper's primary contribution is a new computational
method/algorithm/model/software library
  (no new wet-lab data generation; any data shown is illustrative/benchmarking reuse of public
datasets).
  Examples: a new deep-learning model for cell-type deconvolution; a new marker-gene selection
algorithm.
- EXP = the paper primarily reports **new wet-lab experimental work**. This includes BOTH:
    (i) new biological/clinical findings generated from fresh samples, AND
    (ii) new experimental assays, probe chemistries, imaging platforms, or physical
         instrumentation — even when the paper also describes novel computational analysis.
  Tie-breaker: if the paper introduces a new wet-lab assay / probe chemistry / imaging protocol
  (e.g. padlock probes, smFISH/MERFISH/seqFISH variants, microfluidics, Visium/Xenium-like
platforms),
  classify as EXP regardless of accompanying computational components.
  Examples: "we developed DART-FISH, a padlock-probe ST technology" → EXP;
           "we compared Xenium, MERSCOPE, Visium on tumor cryosections" → EXP;
```

```
          "we present a reinforcement-learning algorithm for FOV selection" → COMP.
- REV = reviews, perspectives, commentary, surveys, meta-analyses.
- OTHER = protocols, resources, datasets, book chapters, guidelines, and similar non-primary
paper types.

spatial:
- spatial=YES only if the PDF text explicitly reports a spatial transcriptomics or in situ
transcriptomics platform/experiment.
- spatial=NO if no spatial transcriptomics experiment is reported.

- Use UNCLEAR when the PDF text does not provide enough direct evidence.
- If direct support is missing, use NOT FOUND and LOW confidence.
```

#### S2.3 GPT-5.4 citation-audit prompt

Each Gemma4 citation classification was packaged into a self-contained JSON audit packet (one packet per tool-paper record). Each packet included the paper metadata, the Gemma4 outputs (label + verbatim evidence quotes + confidence), citation-source inventory, and the `audit_instructions` block reproduced below. GPT-5.4 returned one JSON object per packet, matching the output schema reproduced after the instruction block. ### S2.3.1 Audit instructions

```
{
  "classification_set": [
    "EXP-USED",
    "EXP-CITED",
    "COMP-BENCH",
    "COMP",
    "REV",
    "NON-ST",
    "OTHER"
  ],
  "key_rules": [
    "This is a tool-paper audit, not a paper-only audit: the question is how this paper relates
  to the named tool.",
    "Use EXP-USED only if the evidence explicitly shows that the named tool was used to choose,
  rank, optimize, or validate genes for the paper's own final targeted spatial panel, probe set,
  or target list.",
    "A mention that the tool selected genes, identified markers, ranked features, or reported
  DE genes is NOT enough unless the nearby text also links those genes to panel design or
  targeted spatial usage.",
    "Nearby text means the same sentence, an adjacent sentence, the same paragraph, a figure
  legend, or the same methods block. Do not infer the link from distant context elsewhere in the
  packet.",
    "If the tool is mentioned but there is no close-context statement connecting it to panel
  selection for a targeted spatial experiment, treat that as a negative signal, not a positive
  one.",
    "Use EXP-CITED when the paper is experimental/spatial-transcriptomics but only cites the
  tool, uses it for marker reporting without adopting it for panel selection, or cites it as
  background.",
    "Use COMP-BENCH only for computational or methods papers that explicitly benchmark,
  compare, characterize, or evaluate the named tool.",
    "Use COMP for computational or methods papers that cite the named tool without benchmarking
  or using it directly.",
    "Use REV for reviews, perspectives, tutorials, editorials, and similar synthesis
  articles.",
    "Use NON-ST for papers that are not spatial transcriptomics studies.",
    "Use OTHER for protocols, resources, corrections, clinical reports, or other out-of-scope
  paper types.",
    "Before assigning EXP-USED, confirm all three conditions: targeted spatial or in situ
  study, an actual panel/probe/target-gene selection event, and an explicit statement that the
  named tool drove that selection.",
    "If evidence is insufficient or contradictory, use audit_decision=NEEDS_MANUAL rather than
  inventing certainty."
```

```
  ],
  "task": "Audit Gemma's citation-screening label for this one tool-paper record. Use only this packet unless an external PDF snippet is explicitly provided."
}
```

##### S2.3.2 Audit output schema

```
{
  "type": "object",
  "required": [
    "audit_decision",
    "confidence",
    "Rec. classification",
    "tool_paper_key",
    "paper_key",
    "tool",
    "doi",
    "error_type",
    "reason",
    "evidence_used"
  ],
  "properties": {
    "audit_decision": {
      "enum": [
        "AGREE",
        "DISAGREE",
        "NEEDS_MANUAL"
      ]
    },
    "confidence": {
      "enum": [
        "high",
        "medium",
        "low"
      ]
    },
    "Rec. classification": {
      "enum": [
        "EXP-USED",
        "EXP-CITED",
        "COMP-BENCH",
        "COMP",
        "REV",
        "NON-ST",
        "OTHER"
      ]
    },
    "tool_paper_key": {
      "type": "string"
    },
    "paper_key": {
      "type": "string"
    },
    "tool": {
      "type": "string"
    },
    "doi": {
      "type": "string"
    },
    "error_type": {
      "enum": [
        "none",
        "tool_use_overcalled",
        "tool_use_missed",
```

```
        "benchmark_missed",
        "paper_type_misread",
        "spatial_scope_misread",
        "metadata_prefilter_wrong",
        "wrong_pdf",
        "insufficient_evidence",
        "other"
      ]
    },
    "reason": {
      "type": "string"
    },
    "evidence_used": {
      "type": "string"
    }
  }
}
```
