## supplementary_methods_s3_corpus_queries for "Lack of adoption of computational gene-panel-selection tools in targeted spatial transcriptomics"

### Supplementary Methods S3: Topic corpus query specifications

This document reproduces the full query strings and API call parameters used to harvest the targeted spatial transcriptomics topic corpus from four databases — Europe PMC, OpenAlex, Scopus, and Semantic Scholar. The same per-tool keyword/phrase set is used across all four databases; the surface syntax differs because each backend has its own query language, field-scoping rules, and pagination scheme. The intent is that any reader with the queries and parameters below can reproduce the topic-corpus harvest precisely (modulo any new records added after the publication cutoff of 31 March 2026). All four sources were queried over the same window: papers published from 2015-01-01 through 2026-03-31 (inclusive). Preprints are included where the source indexes them (Europe PMC: SRC:PPR; OpenAlex: type:preprint; Semantic Scholar: empty publicationTypes kept). Scopus does not index preprints, so its contribution is peer-reviewed records only.

#### S3.1 Europe PMC query strings and API parameters

**Endpoint:** `GET https://www.ebi.ac.uk/europepmc/webservices/rest/search`

**Request parameters (per query):**

```
query = <per-tool query string, see S3.1 list>
format = json
resultType = core (includes abstractText; default 'lite' omits it)
cursorMark = * (initial); nextCursorMark for subsequent pages
pageSize = 1000
```

**Pagination.** Cursor-paginated via cursorMark / nextCursorMark. Each query is paged to exhaustion.

**Field-scoping rule.** All searches are field-scoped to TITLE and ABSTRACT only. Europe PMC defaults to full-text search, which inflates results with papers that mention a platform only in passing. The helper _ta(…) emits TITLE:term OR ABSTRACT:term clauses for every primary search term. For AND-qualified queries (ExSeq, HCR-FISH, in situ sequencing, etc.) only the distinctive primary terms are restricted to title+abstract; the disambiguating qualifiers (e.g. ‘spatial’, ‘tissue’, ’transcriptom*’) are left unrestricted so they narrow a generic term rather than acting as identifiers.

**Shared clauses (composed into every per-tool query as `<query> AND _type _excl AND _date`):**

- `_type`:

```
(PUB_TYPE:"Journal Article" OR SRC:PPR)
```

- `_excl (non-research exclusions)`:

```
AND NOT PUB_TYPE:"review" AND NOT PUB_TYPE:"review-article" AND NOT PUB_TYPE:"systematic-
review" AND NOT PUB_TYPE:"meta-analysis" AND NOT PUB_TYPE:"editorial" AND NOT PUB_TYPE:"letter"
AND NOT PUB_TYPE:"comment" AND NOT PUB_TYPE:"article-commentary" AND NOT PUB_TYPE:"discussion"
AND NOT PUB_TYPE:"abstract" AND NOT PUB_TYPE:"case-report" AND NOT PUB_TYPE:"case reports" AND
NOT PUB_TYPE:"dataset" AND NOT PUB_TYPE:"news" AND NOT PUB_TYPE:"retraction of publication" AND
```

```
NOT PUB_TYPE:"erratum" AND NOT PUB_TYPE:"preprint-withdrawal" AND NOT PUB_TYPE:"preprint-
removal"
```

- `_date`:

```
FIRST_PDATE:[2015-01-01 TO 2026-03-31]
```

**Date semantics.** FIRST_PDATE records the date of first publication for journal articles and the server-posting date for preprints (SRC:PPR covers bioRxiv, medRxiv, ChemRxiv, F1000Research, and other preprint servers indexed by Europe PMC).

**Per-tool query strings:**

| # | Term label | Query string |
|---|---|---|
| 1 | MERFISH | `(TITLE:MERFISH OR ABSTRACT:MERFISH OR TITLE:"multiplexed error-robust FISH" OR ABSTRACT:"multiplexed error-robust FISH") AND (PUB_TYPE:"Journal Article" OR SRC:PPR) AND NOT PUB_TYPE:"review" AND NOT PUB_TYPE:"review-article" AND NOT PUB_TYPE:"systematic-review" AND NOT PUB_TYPE:"meta-analysis" AND NOT PUB_TYPE:"editorial" AND NOT PUB_TYPE:"letter" AND NOT PUB_TYPE:"comment" AND NOT PUB_TYPE:"article-commentary" AND NOT PUB_TYPE:"discussion" AND NOT PUB_TYPE:"abstract" AND NOT PUB_TYPE:"case-report" AND NOT PUB_TYPE:"case reports" AND NOT PUB_TYPE:"dataset" AND NOT PUB_TYPE:"news" AND NOT PUB_TYPE:"retraction of publication" AND NOT PUB_TYPE:"erratum" AND NOT PUB_TYPE:"preprint-withdrawal" AND NOT PUB_TYPE:"preprint-removal" AND FIRST_PDATE:[2015-01-01 TO 2026-03-31]` |
| 2 | seqFISH | `(TITLE:seqFISH OR ABSTRACT:seqFISH OR TITLE:"seqFISH+" OR ABSTRACT:"seqFISH+" OR TITLE:"sequential FISH" OR ABSTRACT:"sequential FISH") AND (PUB_TYPE:"Journal Article" OR SRC:PPR) AND NOT PUB_TYPE:"review" AND NOT` |

| # | Term label | Query string |
| --- | --- | --- |
| | | PUB_TYPE:"review-article" AND NOT PUB_TYPE:"systematic-review" AND NOT PUB_TYPE:"meta-analysis" AND NOT PUB_TYPE:"editorial" AND NOT PUB_TYPE:"letter" AND NOT PUB_TYPE:"comment" AND NOT PUB_TYPE:"article-commentary" AND NOT PUB_TYPE:"discussion" AND NOT PUB_TYPE:"abstract" AND NOT PUB_TYPE:"case-report" AND NOT PUB_TYPE:"case reports" AND NOT PUB_TYPE:"dataset" AND NOT PUB_TYPE:"news" AND NOT PUB_TYPE:"retraction of publication" AND NOT PUB_TYPE:"erratum" AND NOT PUB_TYPE:"preprint-withdrawal" AND NOT PUB_TYPE:"preprint-removal" AND FIRST_PDATE:[2015-01-01 TO 2026-03-31] |
| 3 | Xenium | (TITLE:Xenium OR ABSTRACT:Xenium OR TITLE:"10x Genomics Xenium" OR ABSTRACT:"10x Genomics Xenium" OR TITLE:"Xenium In Situ" OR ABSTRACT:"Xenium In Situ") AND (PUB_TYPE:"Journal Article" OR SRC:PPR) AND NOT PUB_TYPE:"review" AND NOT PUB_TYPE:"review-article" AND NOT PUB_TYPE:"systematic-review" AND NOT PUB_TYPE:"meta-analysis" AND NOT PUB_TYPE:"editorial" AND NOT PUB_TYPE:"letter" AND NOT PUB_TYPE:"comment" AND NOT PUB_TYPE:"article-commentary" AND NOT PUB_TYPE:"discussion" AND NOT PUB_TYPE:"abstract" AND NOT PUB_TYPE:"case-report" AND NOT PUB_TYPE:"case reports" AND NOT PUB_TYPE:"dataset" AND NOT PUB_TYPE:"news" AND NOT PUB_TYPE:"retraction of publication" AND NOT PUB_TYPE:"erratum" AND NOT PUB_TYPE:"preprint-withdrawal" AND NOT PUB_TYPE:"preprint-removal" AND FIRST_PDATE:[2015-01-01 TO 2026-03-31] |

| # | Term label | Query string |
|---|---|---|
| 4 | CARTANA | `((TITLE:CARTANA OR ABSTRACT:CARTANA OR TITLE:"direct RNA targeted in situ sequencing" OR ABSTRACT:"direct RNA targeted in situ sequencing" OR TITLE:"10x in situ sequencing" OR ABSTRACT:"10x in situ sequencing") AND (spatial OR transcriptom* OR tissue OR section*)) AND (PUB_TYPE:"Journal Article" OR SRC:PPR) AND NOT PUB_TYPE:"review" AND NOT PUB_TYPE:"review-article" AND NOT PUB_TYPE:"systematic-review" AND NOT PUB_TYPE:"meta-analysis" AND NOT PUB_TYPE:"editorial" AND NOT PUB_TYPE:"letter" AND NOT PUB_TYPE:"comment" AND NOT PUB_TYPE:"article-commentary" AND NOT PUB_TYPE:"discussion" AND NOT PUB_TYPE:"abstract" AND NOT PUB_TYPE:"case-report" AND NOT PUB_TYPE:"case reports" AND NOT PUB_TYPE:"dataset" AND NOT PUB_TYPE:"news" AND NOT PUB_TYPE:"retraction of publication" AND NOT PUB_TYPE:"erratum" AND NOT PUB_TYPE:"preprint-withdrawal" AND NOT PUB_TYPE:"preprint-removal" AND FIRST_PDATE:[2015-01-01 TO 2026-03-31]` |
| 5 | STARmap | `(TITLE:"STARmap" OR ABSTRACT:"STARmap" OR TITLE:"STARmap PLUS" OR ABSTRACT:"STARmap PLUS") AND (PUB_TYPE:"Journal Article" OR SRC:PPR) AND NOT PUB_TYPE:"review" AND NOT PUB_TYPE:"review-article" AND NOT PUB_TYPE:"systematic-review" AND NOT PUB_TYPE:"meta-analysis" AND NOT PUB_TYPE:"editorial" AND NOT PUB_TYPE:"letter" AND NOT PUB_TYPE:"comment" AND NOT PUB_TYPE:"article-commentary" AND NOT PUB_TYPE:"discussion" AND NOT PUB_TYPE:"abstract" AND NOT PUB_TYPE:"case-report" AND NOT` |

| # | Term label | Query string |
| --- | --- | --- |
| | | `PUB_TYPE:"case reports" AND NOT PUB_TYPE:"dataset" AND NOT PUB_TYPE:"news" AND NOT PUB_TYPE:"retraction of publication" AND NOT PUB_TYPE:"erratum" AND NOT PUB_TYPE:"preprint-withdrawal" AND NOT PUB_TYPE:"preprint-removal" AND FIRST_PDATE:[2015-01-01 TO 2026-03-31]` |
| 6 | osmFISH | `(TITLE:osmFISH OR ABSTRACT:osmFISH) AND (PUB_TYPE:"Journal Article" OR SRC:PPR) AND NOT PUB_TYPE:"review" AND NOT PUB_TYPE:"review-article" AND NOT PUB_TYPE:"systematic-review" AND NOT PUB_TYPE:"meta-analysis" AND NOT PUB_TYPE:"editorial" AND NOT PUB_TYPE:"letter" AND NOT PUB_TYPE:"comment" AND NOT PUB_TYPE:"article-commentary" AND NOT PUB_TYPE:"discussion" AND NOT PUB_TYPE:"abstract" AND NOT PUB_TYPE:"case-report" AND NOT PUB_TYPE:"case reports" AND NOT PUB_TYPE:"dataset" AND NOT PUB_TYPE:"news" AND NOT PUB_TYPE:"retraction of publication" AND NOT PUB_TYPE:"erratum" AND NOT PUB_TYPE:"preprint-withdrawal" AND NOT PUB_TYPE:"preprint-removal" AND FIRST_PDATE:[2015-01-01 TO 2026-03-31]` |
| 7 | EEL FISH | `((TITLE:"EEL FISH" OR ABSTRACT:"EEL FISH" OR TITLE:"EEL-FISH" OR ABSTRACT:"EEL-FISH") AND (transcriptom* OR "in situ" OR spatial OR probe*)) AND (PUB_TYPE:"Journal Article" OR SRC:PPR) AND NOT PUB_TYPE:"review" AND NOT PUB_TYPE:"review-article" AND NOT PUB_TYPE:"systematic-review" AND NOT PUB_TYPE:"meta-analysis" AND NOT PUB_TYPE:"editorial" AND NOT PUB_TYPE:"letter" AND NOT PUB_TYPE:"comment" AND NOT PUB_TYPE:"article-commentary"` |

| # | Term label | Query string |
| --- | --- | --- |
| | | AND NOT PUB_TYPE:"discussion" AND NOT PUB_TYPE:"abstract" AND NOT PUB_TYPE:"case-report" AND NOT PUB_TYPE:"case reports" AND NOT PUB_TYPE:"dataset" AND NOT PUB_TYPE:"news" AND NOT PUB_TYPE:"retraction of publication" AND NOT PUB_TYPE:"erratum" AND NOT PUB_TYPE:"preprint-withdrawal" AND NOT PUB_TYPE:"preprint-removal" AND FIRST_PDATE:[2015-01-01 TO 2026-03-31] |
| 8 | HybISS | (TITLE:HybISS OR ABSTRACT:HybISS) AND (PUB_TYPE:"Journal Article" OR SRC:PPR) AND NOT PUB_TYPE:"review" AND NOT PUB_TYPE:"review-article" AND NOT PUB_TYPE:"systematic-review" AND NOT PUB_TYPE:"meta-analysis" AND NOT PUB_TYPE:"editorial" AND NOT PUB_TYPE:"letter" AND NOT PUB_TYPE:"comment" AND NOT PUB_TYPE:"article-commentary" AND NOT PUB_TYPE:"discussion" AND NOT PUB_TYPE:"abstract" AND NOT PUB_TYPE:"case-report" AND NOT PUB_TYPE:"case reports" AND NOT PUB_TYPE:"dataset" AND NOT PUB_TYPE:"news" AND NOT PUB_TYPE:"retraction of publication" AND NOT PUB_TYPE:"erratum" AND NOT PUB_TYPE:"preprint-withdrawal" AND NOT PUB_TYPE:"preprint-removal" AND FIRST_PDATE:[2015-01-01 TO 2026-03-31] |
| 9 | BaristaSeq | (TITLE:BaristaSeq OR ABSTRACT:BaristaSeq) AND (PUB_TYPE:"Journal Article" OR SRC:PPR) AND NOT PUB_TYPE:"review" AND NOT PUB_TYPE:"review-article" AND NOT PUB_TYPE:"systematic-review" AND NOT PUB_TYPE:"meta-analysis" AND NOT PUB_TYPE:"editorial" AND NOT PUB_TYPE:"letter" AND NOT PUB_TYPE:"comment" AND NOT PUB_TYPE:"article-commentary" |

| # | Term label | Query string |
|---|---|---|
| | | AND NOT PUB_TYPE:"discussion" AND NOT PUB_TYPE:"abstract" AND NOT PUB_TYPE:"case-report" AND NOT PUB_TYPE:"case reports" AND NOT PUB_TYPE:"dataset" AND NOT PUB_TYPE:"news" AND NOT PUB_TYPE:"retraction of publication" AND NOT PUB_TYPE:"erratum" AND NOT PUB_TYPE:"preprint-withdrawal" AND NOT PUB_TYPE:"preprint-removal" AND FIRST_PDATE:[2015-01-01 TO 2026-03-31] |
| 10 | SCRINSHOT | (TITLE:SCRINSHOT OR ABSTRACT:SCRINSHOT) AND (PUB_TYPE:"Journal Article" OR SRC:PPR) AND NOT PUB_TYPE:"review" AND NOT PUB_TYPE:"review-article" AND NOT PUB_TYPE:"systematic-review" AND NOT PUB_TYPE:"meta-analysis" AND NOT PUB_TYPE:"editorial" AND NOT PUB_TYPE:"letter" AND NOT PUB_TYPE:"comment" AND NOT PUB_TYPE:"article-commentary" AND NOT PUB_TYPE:"discussion" AND NOT PUB_TYPE:"abstract" AND NOT PUB_TYPE:"case-report" AND NOT PUB_TYPE:"case reports" AND NOT PUB_TYPE:"dataset" AND NOT PUB_TYPE:"news" AND NOT PUB_TYPE:"retraction of publication" AND NOT PUB_TYPE:"erratum" AND NOT PUB_TYPE:"preprint-withdrawal" AND NOT PUB_TYPE:"preprint-removal" AND FIRST_PDATE:[2015-01-01 TO 2026-03-31] |
| 11 | EASI-FISH | (TITLE:"EASI-FISH" OR ABSTRACT:"EASI-FISH") AND (PUB_TYPE:"Journal Article" OR SRC:PPR) AND NOT PUB_TYPE:"review" AND NOT PUB_TYPE:"review-article" AND NOT PUB_TYPE:"systematic-review" AND NOT PUB_TYPE:"meta-analysis" AND NOT PUB_TYPE:"editorial" AND NOT PUB_TYPE:"letter" AND NOT PUB_TYPE:"comment" AND NOT PUB_TYPE:"article-commentary" |

| # | Term label | Query string |
| --- | --- | --- |
| | | AND NOT PUB_TYPE:"discussion" AND NOT PUB_TYPE:"abstract" AND NOT PUB_TYPE:"case-report" AND NOT PUB_TYPE:"case reports" AND NOT PUB_TYPE:"dataset" AND NOT PUB_TYPE:"news" AND NOT PUB_TYPE:"retraction of publication" AND NOT PUB_TYPE:"erratum" AND NOT PUB_TYPE:"preprint-withdrawal" AND NOT PUB_TYPE:"preprint-removal" AND FIRST_PDATE:[2015-01-01 TO 2026-03-31] |
| 12 | SPLIT-FISH | (TITLE:"SPLIT-FISH" OR ABSTRACT:"SPLIT-FISH") AND (PUB_TYPE:"Journal Article" OR SRC:PPR) AND NOT PUB_TYPE:"review" AND NOT PUB_TYPE:"review-article" AND NOT PUB_TYPE:"systematic-review" AND NOT PUB_TYPE:"meta-analysis" AND NOT PUB_TYPE:"editorial" AND NOT PUB_TYPE:"letter" AND NOT PUB_TYPE:"comment" AND NOT PUB_TYPE:"article-commentary" AND NOT PUB_TYPE:"discussion" AND NOT PUB_TYPE:"abstract" AND NOT PUB_TYPE:"case-report" AND NOT PUB_TYPE:"case reports" AND NOT PUB_TYPE:"dataset" AND NOT PUB_TYPE:"news" AND NOT PUB_TYPE:"retraction of publication" AND NOT PUB_TYPE:"erratum" AND NOT PUB_TYPE:"preprint-withdrawal" AND NOT PUB_TYPE:"preprint-removal" AND FIRST_PDATE:[2015-01-01 TO 2026-03-31] |
| 13 | targeted spatial | ((TITLE:"targeted spatial transcriptomics" OR ABSTRACT:"targeted spatial transcriptomics") AND (panel* OR probe* OR "gene panel" OR "probe design" OR imaging OR "in situ")) AND (PUB_TYPE:"Journal Article" OR SRC:PPR) AND NOT PUB_TYPE:"review" AND NOT PUB_TYPE:"review-article" AND NOT PUB_TYPE:"systematic-review" AND NOT |

| # | Term label | Query string |
|---|---|---|
| | | PUB_TYPE:"meta-analysis" AND NOT PUB_TYPE:"editorial" AND NOT PUB_TYPE:"letter" AND NOT PUB_TYPE:"comment" AND NOT PUB_TYPE:"article-commentary" AND NOT PUB_TYPE:"discussion" AND NOT PUB_TYPE:"abstract" AND NOT PUB_TYPE:"case-report" AND NOT PUB_TYPE:"case reports" AND NOT PUB_TYPE:"dataset" AND NOT PUB_TYPE:"news" AND NOT PUB_TYPE:"retraction of publication" AND NOT PUB_TYPE:"erratum" AND NOT PUB_TYPE:"preprint-withdrawal" AND NOT PUB_TYPE:"preprint-removal" AND FIRST_PDATE:[2015-01-01 TO 2026-03-31] |
| 14 | in situ transcriptomics | (TITLE:"in situ transcriptomics" OR ABSTRACT:"in situ transcriptomics") AND (PUB_TYPE:"Journal Article" OR SRC:PPR) AND NOT PUB_TYPE:"review" AND NOT PUB_TYPE:"review-article" AND NOT PUB_TYPE:"systematic-review" AND NOT PUB_TYPE:"meta-analysis" AND NOT PUB_TYPE:"editorial" AND NOT PUB_TYPE:"letter" AND NOT PUB_TYPE:"comment" AND NOT PUB_TYPE:"article-commentary" AND NOT PUB_TYPE:"discussion" AND NOT PUB_TYPE:"abstract" AND NOT PUB_TYPE:"case-report" AND NOT PUB_TYPE:"case reports" AND NOT PUB_TYPE:"dataset" AND NOT PUB_TYPE:"news" AND NOT PUB_TYPE:"retraction of publication" AND NOT PUB_TYPE:"erratum" AND NOT PUB_TYPE:"preprint-withdrawal" AND NOT PUB_TYPE:"preprint-removal" AND FIRST_PDATE:[2015-01-01 TO 2026-03-31] |
| 15 | MERSCOPE | ((TITLE:MERSCOPE OR ABSTRACT:MERSCOPE OR TITLE:"Vizgen MERSCOPE" OR ABSTRACT:"Vizgen MERSCOPE") OR ((TITLE:VIZGEN OR ABSTRACT:VIZGEN) AND (MERFISH |

| # | Term label | Query string |
|---|---|---|
| | | `OR spatial OR transcriptom*))) AND (PUB_TYPE:"Journal Article" OR SRC:PPR) AND NOT PUB_TYPE:"review" AND NOT PUB_TYPE:"review-article" AND NOT PUB_TYPE:"systematic-review" AND NOT PUB_TYPE:"meta-analysis" AND NOT PUB_TYPE:"editorial" AND NOT PUB_TYPE:"letter" AND NOT PUB_TYPE:"comment" AND NOT PUB_TYPE:"article-commentary" AND NOT PUB_TYPE:"discussion" AND NOT PUB_TYPE:"abstract" AND NOT PUB_TYPE:"case-report" AND NOT PUB_TYPE:"case reports" AND NOT PUB_TYPE:"dataset" AND NOT PUB_TYPE:"news" AND NOT PUB_TYPE:"retraction of publication" AND NOT PUB_TYPE:"erratum" AND NOT PUB_TYPE:"preprint-withdrawal" AND NOT PUB_TYPE:"preprint-removal" AND FIRST_PDATE:[2015-01-01 TO 2026-03-31]` |
| 16 | CosMx | `((TITLE:CosMx OR ABSTRACT:CosMx OR TITLE:"CosMx SMI" OR ABSTRACT:"CosMx SMI" OR TITLE:"NanoString CosMx" OR ABSTRACT:"NanoString CosMx") OR ((TITLE:"spatial molecular imager" OR ABSTRACT:"spatial molecular imager") AND NanoString)) AND (PUB_TYPE:"Journal Article" OR SRC:PPR) AND NOT PUB_TYPE:"review" AND NOT PUB_TYPE:"review-article" AND NOT PUB_TYPE:"systematic-review" AND NOT PUB_TYPE:"meta-analysis" AND NOT PUB_TYPE:"editorial" AND NOT PUB_TYPE:"letter" AND NOT PUB_TYPE:"comment" AND NOT PUB_TYPE:"article-commentary" AND NOT PUB_TYPE:"discussion" AND NOT PUB_TYPE:"abstract" AND NOT PUB_TYPE:"case-report" AND NOT PUB_TYPE:"case reports" AND NOT PUB_TYPE:"dataset" AND NOT PUB_TYPE:"news" AND NOT` |

| # | Term label | Query string |
|---|---|---|
| | | PUB_TYPE:"retraction of publication" AND NOT PUB_TYPE:"erratum" AND NOT PUB_TYPE:"preprint-withdrawal" AND NOT PUB_TYPE:"preprint-removal" AND FIRST_PDATE:[2015-01-01 TO 2026-03-31] |
| 17 | Molecular Cartography | ((TITLE:"Molecular Cartography" OR ABSTRACT:"Molecular Cartography") OR ((TITLE:"Resolve Biosciences" OR ABSTRACT:"Resolve Biosciences") AND spatial)) AND (PUB_TYPE:"Journal Article" OR SRC:PPR) AND NOT PUB_TYPE:"review" AND NOT PUB_TYPE:"review-article" AND NOT PUB_TYPE:"systematic-review" AND NOT PUB_TYPE:"meta-analysis" AND NOT PUB_TYPE:"editorial" AND NOT PUB_TYPE:"letter" AND NOT PUB_TYPE:"comment" AND NOT PUB_TYPE:"article-commentary" AND NOT PUB_TYPE:"discussion" AND NOT PUB_TYPE:"abstract" AND NOT PUB_TYPE:"case-report" AND NOT PUB_TYPE:"case reports" AND NOT PUB_TYPE:"dataset" AND NOT PUB_TYPE:"news" AND NOT PUB_TYPE:"retraction of publication" AND NOT PUB_TYPE:"erratum" AND NOT PUB_TYPE:"preprint-withdrawal" AND NOT PUB_TYPE:"preprint-removal" AND FIRST_PDATE:[2015-01-01 TO 2026-03-31] |
| 18 | ExSeq | ((TITLE:ExSeq OR ABSTRACT:ExSeq OR TITLE:"expansion sequencing" OR ABSTRACT:"expansion sequencing") AND (spatial OR transcriptom* OR "in situ")) AND (PUB_TYPE:"Journal Article" OR SRC:PPR) AND NOT PUB_TYPE:"review" AND NOT PUB_TYPE:"review-article" AND NOT PUB_TYPE:"systematic-review" AND NOT PUB_TYPE:"meta-analysis" AND NOT PUB_TYPE:"editorial" AND NOT PUB_TYPE:"letter" AND NOT |

| # | Term label | Query string |
|---|---|---|
| | | PUB_TYPE:"comment" AND NOT PUB_TYPE:"article-commentary" AND NOT PUB_TYPE:"discussion" AND NOT PUB_TYPE:"abstract" AND NOT PUB_TYPE:"case-report" AND NOT PUB_TYPE:"case reports" AND NOT PUB_TYPE:"dataset" AND NOT PUB_TYPE:"news" AND NOT PUB_TYPE:"retraction of publication" AND NOT PUB_TYPE:"erratum" AND NOT PUB_TYPE:"preprint-withdrawal" AND NOT PUB_TYPE:"preprint-removal" AND FIRST_PDATE:[2015-01-01 TO 2026-03-31] |
| 19 | SABER-FISH | ((TITLE:"SABER-FISH" OR ABSTRACT:"SABER-FISH" OR TITLE:"SABER FISH" OR ABSTRACT:"SABER FISH" OR TITLE:"signal amplification by exchange reaction" OR ABSTRACT:"signal amplification by exchange reaction") AND (multiplex* OR spatial)) AND (PUB_TYPE:"Journal Article" OR SRC:PPR) AND NOT PUB_TYPE:"review" AND NOT PUB_TYPE:"review-article" AND NOT PUB_TYPE:"systematic-review" AND NOT PUB_TYPE:"meta-analysis" AND NOT PUB_TYPE:"editorial" AND NOT PUB_TYPE:"letter" AND NOT PUB_TYPE:"comment" AND NOT PUB_TYPE:"article-commentary" AND NOT PUB_TYPE:"discussion" AND NOT PUB_TYPE:"abstract" AND NOT PUB_TYPE:"case-report" AND NOT PUB_TYPE:"case reports" AND NOT PUB_TYPE:"dataset" AND NOT PUB_TYPE:"news" AND NOT PUB_TYPE:"retraction of publication" AND NOT PUB_TYPE:"erratum" AND NOT PUB_TYPE:"preprint-withdrawal" AND NOT PUB_TYPE:"preprint-removal" AND FIRST_PDATE:[2015-01-01 TO 2026-03-31] |
| 20 | HCR-FISH | ((TITLE:"HCR-FISH" OR ABSTRACT:"HCR-FISH" OR TITLE:"hybridization chain |

| # | Term label | Query string |
|---|---|---|
| | | `reaction" OR ABSTRACT:"hybridization chain reaction") AND (multiplex* OR panel* OR "gene panel" OR "high-plex") AND (spatial OR "spatial transcriptom*" OR "in situ transcriptom*" OR transcriptom*)) AND (PUB_TYPE:"Journal Article" OR SRC:PPR) AND NOT PUB_TYPE:"review" AND NOT PUB_TYPE:"review-article" AND NOT PUB_TYPE:"systematic-review" AND NOT PUB_TYPE:"meta-analysis" AND NOT PUB_TYPE:"editorial" AND NOT PUB_TYPE:"letter" AND NOT PUB_TYPE:"comment" AND NOT PUB_TYPE:"article-commentary" AND NOT PUB_TYPE:"discussion" AND NOT PUB_TYPE:"abstract" AND NOT PUB_TYPE:"case-report" AND NOT PUB_TYPE:"case reports" AND NOT PUB_TYPE:"dataset" AND NOT PUB_TYPE:"news" AND NOT PUB_TYPE:"retraction of publication" AND NOT PUB_TYPE:"erratum" AND NOT PUB_TYPE:"preprint-withdrawal" AND NOT PUB_TYPE:"preprint-removal" AND FIRST_PDATE:[2015-01-01 TO 2026-03-31]` |
| 21 | in situ sequencing | `((TITLE:"in situ sequencing" OR ABSTRACT:"in situ sequencing" OR TITLE:"direct RNA targeted in situ sequencing" OR ABSTRACT:"direct RNA targeted in situ sequencing") AND ((spatial OR tissue OR section*) AND (transcriptom* OR target* OR panel* OR "cell type"))) AND (PUB_TYPE:"Journal Article" OR SRC:PPR) AND NOT PUB_TYPE:"review" AND NOT PUB_TYPE:"review-article" AND NOT PUB_TYPE:"systematic-review" AND NOT PUB_TYPE:"meta-analysis" AND NOT PUB_TYPE:"editorial" AND NOT PUB_TYPE:"letter" AND NOT PUB_TYPE:"comment" AND NOT PUB_TYPE:"article-commentary"` |

| # | Term label | Query string |
|---|---|---|
| | | AND NOT PUB_TYPE:"discussion" AND NOT PUB_TYPE:"abstract" AND NOT PUB_TYPE:"case-report" AND NOT PUB_TYPE:"case reports" AND NOT PUB_TYPE:"dataset" AND NOT PUB_TYPE:"news" AND NOT PUB_TYPE:"retraction of publication" AND NOT PUB_TYPE:"erratum" AND NOT PUB_TYPE:"preprint-withdrawal" AND NOT PUB_TYPE:"preprint-removal" AND FIRST_PDATE:[2015-01-01 TO 2026-03-31] |

#### S3.2 OpenAlex query strings and filter parameters

**Endpoint:** `GET https://api.openalex.org/works`

**Request parameters (per query):**

```
api_key = <OA_API_KEY>
filter = from_publication_date:2015-01-01,to_publication_date:2026-03-
31,type:article|preprint,title_and_abstract.search:<per-tool search string, see S3.2 list>
select =
id,doi,title,publication_year,publication_date,primary_location,open_access,authorships,abstrac
t_inverted_index,cited_by_count,type,language
per_page = 200
cursor = * (initial); meta.next_cursor for subsequent pages
```

**Pagination.** Cursor-paginated. Each query is paged to exhaustion. 100 ms sleep between pages. Per-query fetches run in parallel via a 4-worker thread pool; results are written in deterministic label order.

**Field-scoping rule.** title_and_abstract.search restricts matching to title and abstract text only (no full-text). type:article|preprint includes both peer-reviewed articles and preprint server records.

**Per-tool search strings (substituted into `title_and_abstract.search:` in the filter above):**

| # | Term label | Query string |
|---|---|---|
| 1 | MERFISH | MERFISH OR "multiplexed error-robust FISH" |
| 2 | seqFISH | seqFISH OR "seqFISH+" OR "sequential FISH" |
| 3 | Xenium | Xenium OR "10x Genomics Xenium" OR "Xenium In Situ" |
| 4 | CARTANA | CARTANA OR "direct RNA targeted in situ sequencing" OR "10x in situ sequencing" |
| 5 | MERSCOPE | MERSCOPE OR "Vizgen MERSCOPE" OR (VIZGEN AND |

| # | Term label | Query string |
|---|---|---|
| | | `(MERFISH OR spatial OR transcriptom*))` |
| 6 | CosMx | `CosMx OR "CosMx SMI" OR "NanoString CosMx" OR ("spatial molecular imager" AND NanoString)` |
| 7 | Molecular Cartography | `"Molecular Cartography" OR ("Resolve Biosciences" AND spatial)` |
| 8 | STARmap | `"STARmap" OR "STARmap PLUS"` |
| 9 | osmFISH | `osmFISH` |
| 10 | EEL FISH | `("EEL FISH" OR "EEL-FISH") AND (transcriptom* OR "in situ" OR spatial OR probe*)` |
| 11 | HybISS | `HybISS` |
| 12 | BaristaSeq | `BaristaSeq` |
| 13 | SCRINSHOT | `SCRINSHOT` |
| 14 | EASI-FISH | `"EASI-FISH"` |
| 15 | ExSeq | `(ExSeq OR "expansion sequencing") AND (spatial OR transcriptom* OR "in situ")` |
| 16 | SABER-FISH | `("SABER-FISH" OR "SABER FISH" OR "signal amplification by exchange reaction") AND (multiplex* OR spatial)` |
| 17 | HCR-FISH | `("HCR-FISH" OR "hybridization chain reaction") AND (multiplex* OR panel* OR "gene panel" OR "high-plex") AND (spatial OR "spatial transcriptom*" OR "in situ transcriptom*" OR transcriptom*)` |
| 18 | in situ sequencing | `("in situ sequencing" OR "direct RNA targeted in situ sequencing") AND ((spatial OR tissue OR section*) AND (transcriptom* OR target* OR panel* OR "cell type"))` |
| 19 | SPLIT-FISH | `"SPLIT-FISH"` |
| 20 | targeted spatial | `"targeted spatial transcriptomics" AND (panel* OR probe* OR "gene panel" OR "probe design" OR imaging OR "in situ")` |
| 21 | in situ transcriptomics | `"in situ transcriptomics"` |

#### S3.3 Scopus query strings

**Endpoint:** `GET https://api.elsevier.com/content/search/scopus`

**Headers:**

```
X-ELS-APIKey = <SCOPUS_API_KEY>
Accept = application/json
```

**Request parameters (per query):**

```
query = <per-tool query string, see S3.3 list>
start = 0, 25, 50, ... (incremented per page)
count = 25
view = COMPLETE (includes dc:description abstract; falls back to STANDARD when the API key
lacks COMPLETE entitlement)
```

**Pagination.** Offset-paginated via start + count. Each query is paged to exhaustion or to min(total, 5000) records (Scopus API hard cap). 200 ms sleep between pages.

**Scopus-specific syntax:**

- **TITLE-ABS-KEY scoping.** Every primary term (and any AND-qualifier subclause) is wrapped in TITLE-ABS-KEY(…). Scopus does not search full text by default, but this is explicit and avoids accidental MeSH or affiliation matches.
- **DOCTYPE / SRCTYPE filter.** DOCTYPE(ar) AND SRCTYPE(j) AND PUBYEAR > 2014 AND PUBYEAR < 2027

**Preprint coverage.** DOCTYPE(ar) restricts to journal articles. SRCTYPE(j) adds a second guard against conference proceedings and other non-journal content. Scopus does not index preprints, so the corpus contribution from this source is intentionally limited to peer-reviewed records.

**Per-tool query strings (already include the TITLE-ABS-KEY wrap and the date / doctype filter):**

| # | Term label | Query string |
|---|---|---|
| 1 | MERFISH | `TITLE-ABS-KEY(MERFISH OR "multiplexed error-robust FISH") AND DOCTYPE(ar) AND SRCTYPE(j) AND PUBYEAR > 2014 AND PUBYEAR < 2027` |
| 2 | seqFISH | `TITLE-ABS-KEY(seqFISH OR "seqFISH+" OR "sequential FISH") AND DOCTYPE(ar) AND SRCTYPE(j) AND PUBYEAR > 2014 AND PUBYEAR < 2027` |
| 3 | Xenium | `TITLE-ABS-KEY(Xenium OR "10x Genomics Xenium" OR "Xenium In Situ") AND DOCTYPE(ar) AND SRCTYPE(j) AND PUBYEAR > 2014 AND PUBYEAR < 2027` |
| 4 | CARTANA | `TITLE-ABS-KEY(CARTANA OR "direct RNA targeted in situ sequencing" OR "10x in situ` |

| # | Term label | Query string |
|---|---|---|
| | | sequencing") AND DOCTYPE(ar) AND SRCTYPE(j) AND PUBYEAR > 2014 AND PUBYEAR < 2027 |
| 5 | MERSCOPE | TITLE-ABS-KEY(MERSCOPE OR "Vizgen MERSCOPE" OR (VIZGEN AND (MERFISH OR spatial OR transcriptom*))) AND DOCTYPE(ar) AND SRCTYPE(j) AND PUBYEAR > 2014 AND PUBYEAR < 2027 |
| 6 | CosMx | TITLE-ABS-KEY(CosMx OR "CosMx SMI" OR "NanoString CosMx" OR ("spatial molecular imager" AND NanoString)) AND DOCTYPE(ar) AND SRCTYPE(j) AND PUBYEAR > 2014 AND PUBYEAR < 2027 |
| 7 | Molecular Cartography | TITLE-ABS-KEY("Molecular Cartography" OR ("Resolve Biosciences" AND spatial)) AND DOCTYPE(ar) AND SRCTYPE(j) AND PUBYEAR > 2014 AND PUBYEAR < 2027 |
| 8 | STARmap | TITLE-ABS-KEY("STARmap" OR "STARmap PLUS") AND DOCTYPE(ar) AND SRCTYPE(j) AND PUBYEAR > 2014 AND PUBYEAR < 2027 |
| 9 | osmFISH | TITLE-ABS-KEY(osmFISH) AND DOCTYPE(ar) AND SRCTYPE(j) AND PUBYEAR > 2014 AND PUBYEAR < 2027 |
| 10 | EEL FISH | TITLE-ABS-KEY(("EEL FISH" OR "EEL-FISH") AND (transcriptom* OR "in situ" OR spatial OR probe*)) AND DOCTYPE(ar) AND SRCTYPE(j) AND PUBYEAR > 2014 AND PUBYEAR < 2027 |
| 11 | HybISS | TITLE-ABS-KEY(HybISS) AND DOCTYPE(ar) AND SRCTYPE(j) AND PUBYEAR > 2014 AND PUBYEAR < 2027 |
| 12 | BaristaSeq | TITLE-ABS-KEY(BaristaSeq) AND DOCTYPE(ar) AND SRCTYPE(j) AND PUBYEAR > 2014 AND PUBYEAR < 2027 |
| 13 | SCRINSHOT | TITLE-ABS-KEY(SCRINSHOT) AND DOCTYPE(ar) AND SRCTYPE(j) AND PUBYEAR > 2014 AND PUBYEAR < 2027 |

| # | Term label | Query string |
|---|---|---|
| 14 | EASI-FISH | `TITLE-ABS-KEY("EASI-FISH") AND DOCTYPE(ar) AND SRCTYPE(j) AND PUBYEAR > 2014 AND PUBYEAR < 2027` |
| 15 | ExSeq | `TITLE-ABS-KEY((ExSeq OR "expansion sequencing") AND (spatial OR transcriptom* OR "in situ")) AND DOCTYPE(ar) AND SRCTYPE(j) AND PUBYEAR > 2014 AND PUBYEAR < 2027` |
| 16 | SABER-FISH | `TITLE-ABS-KEY(("SABER-FISH" OR "SABER FISH" OR "signal amplification by exchange reaction") AND (multiplex* OR spatial)) AND DOCTYPE(ar) AND SRCTYPE(j) AND PUBYEAR > 2014 AND PUBYEAR < 2027` |
| 17 | HCR-FISH | `TITLE-ABS-KEY(("HCR-FISH" OR "hybridization chain reaction") AND (multiplex* OR panel* OR "gene panel" OR "high-plex") AND (spatial OR "spatial transcriptom*" OR "in situ transcriptom*" OR transcriptom*)) AND DOCTYPE(ar) AND SRCTYPE(j) AND PUBYEAR > 2014 AND PUBYEAR < 2027` |
| 18 | in situ sequencing | `TITLE-ABS-KEY(("in situ sequencing" OR "direct RNA targeted in situ sequencing") AND ((spatial OR tissue OR section*) AND (transcriptom* OR target* OR panel* OR "cell type"))) AND DOCTYPE(ar) AND SRCTYPE(j) AND PUBYEAR > 2014 AND PUBYEAR < 2027` |
| 19 | SPLIT-FISH | `TITLE-ABS-KEY("SPLIT-FISH") AND DOCTYPE(ar) AND SRCTYPE(j) AND PUBYEAR > 2014 AND PUBYEAR < 2027` |
| 20 | targeted spatial | `TITLE-ABS-KEY("targeted spatial transcriptomics" AND (panel* OR probe* OR "gene panel" OR "probe design" OR imaging OR "in situ")) AND DOCTYPE(ar) AND SRCTYPE(j) AND PUBYEAR > 2014 AND PUBYEAR < 2027` |
| 21 | in situ transcriptomics | `TITLE-ABS-KEY("in situ transcriptomics") AND DOCTYPE(ar) AND SRCTYPE(j)` |

| # | Term label | Query string |
|---|---|---|
| | | AND PUBYEAR > 2014 AND PUBYEAR < 2027 |

#### S3.4 Semantic Scholar query strings

**Endpoint:** `GET https://api.semanticscholar.org/graph/v1/paper/search/bulk`

**Headers:**

```
x-api-key = <S2_API_KEY>
```

**Request parameters (per query):**

```
query = <per-tool query string, see S3.4 list — uses S2 bulk-search syntax>
publicationDateOrYear = 2015-01-01:2026-03-31
fields =
paperId,externalIds,title,abstract,year,publicationDate,journal,publicationTypes,authors,fields
OfStudy,openAccessPdf,citationCount
limit = 1000
token = <empty on first page; data.token on subsequent pages>
```

**Pagination.** Token-paginated. Each query is paged to exhaustion.

**Semantic Scholar bulk-search syntax:**

- `|` — OR
- `+` — AND
- `-` — NOT
- `*` — prefix wildcard
- `"phrase"` — exact phrase

**publicationTypes filter rationale.** The publicationTypes filter is intentionally NOT applied at the API level: Semantic Scholar has no ‘Preprint’ enum, and bioRxiv / medRxiv preprints have an empty publicationTypes array. Filtering on publicationTypes=JournalArticle would silently drop those preprints. Non-research types (Review, Conference, Editorial, etc.) are excluded downstream in deduplicate.py instead.

**Per-tool query strings:**

| # | Term label | Query string |
|---|---|---|
| 1 | MERFISH | MERFISH \| "multiplexed error-robust FISH" |
| 2 | seqFISH | seqFISH \| "seqFISH+" \| "sequential FISH" |
| 3 | Xenium | Xenium \| "10x Genomics Xenium" \| "Xenium In Situ" |
| 4 | CARTANA | CARTANA \| "direct RNA targeted in situ sequencing" \| "10x in situ sequencing" |
| 5 | MERSCOPE | MERSCOPE \| "Vizgen MERSCOPE" \| (VIZGEN + |

| # | Term label | Query string |
| --- | --- | --- |
| | | (MERFISH \| spatial \| transcriptom*)) |
| 6 | CosMx | CosMx \| "CosMx SMI" \| "NanoString CosMx" \| ("spatial molecular imager" + NanoString) |
| 7 | Molecular Cartography | "Molecular Cartography" \| ("Resolve Biosciences" + spatial) |
| 8 | STARmap | "STARmap" \| "STARmap PLUS" |
| 9 | osmFISH | osmFISH |
| 10 | EEL FISH | ("EEL FISH" \| "EEL-FISH") + (transcriptom* \| "in situ" \| spatial \| probe*) |
| 11 | HybISS | HybISS |
| 12 | BaristaSeq | BaristaSeq |
| 13 | SCRINSHOT | SCRINSHOT |
| 14 | EASI-FISH | "EASI-FISH" |
| 15 | ExSeq | (ExSeq \| "expansion sequencing") + (spatial \| transcriptom* \| "in situ") |
| 16 | SABER-FISH | ("SABER-FISH" \| "SABER FISH" \| "signal amplification by exchange reaction") + (multiplex* \| spatial) |
| 17 | HCR-FISH | ("HCR-FISH" \| "hybridization chain reaction") + (multiplex* \| panel* \| "gene panel" \| "high-plex") + (spatial \| "spatial transcriptom*" \| "in situ transcriptom*" \| transcriptom*) |
| 18 | in situ sequencing | ("in situ sequencing" \| "direct RNA targeted in situ sequencing") + ((spatial \| tissue \| section*) + (transcriptom* \| target* \| panel* \| "cell type")) |
| 19 | SPLIT-FISH | "SPLIT-FISH" |
| 20 | targeted spatial | "targeted spatial transcriptomics" + (panel* \| probe* \| "gene panel" \| "probe design" \| imaging \| "in situ") |
| 21 | in situ transcriptomics | "in situ transcriptomics" |
