## supplementary_methods_s4_panel_prompts for "Lack of adoption of computational gene-panel-selection tools in targeted spatial transcriptomics"

### Supplementary Methods S4: Panel-design classification prompts

This document reproduces the prompt templates and structured-output schema used to classify each topic-corpus paper by its reported panel-design strategy. Classification ran in two automated stages followed by an independent audit:

1. A local Gemma4 model (served through Ollama) classified each paper's document type, panel-design approach, primary paper profile, and panel scope, using the extracted PDF text (or the abstract only for records without a retrievable PDF).
2. An independent GPT-5.4 audit reviewed every Gemma4 classification against a self-contained audit packet and either agreed, disagreed, or routed the record to manual review.

Both stages ran in JSON mode and required verbatim PDF quotes for every evidence field. The Gemma4 prompts are reproduced verbatim from the source classifier; the GPT-5.4 audit instructions are reproduced verbatim from the audit packets actually sent to the model (using the first packet as the canonical example).

#### S4.1 Gemma4 panel-design classification prompt

For each corpus paper with a retrievable PDF, the extracted text was passed four times to Gemma4 (Ollama backend), once per sub-prompt below. Records without a retrievable PDF received the abstract-only profile sub-prompt (S4.1.4) instead of the three full-text sub-prompts. ### S4.1.1 Document-type sub-prompt (`DOC_TYPE_PROMPT`)

```
You are classifying a scientific PDF using ONLY the provided full extracted PDF text.

Return one JSON object only with this exact schema:
{
  "doc_type": "MAIN_ARTICLE|REVIEW|EDITORIAL|CORRECTION|SUPPLEMENT|NON_ARTICLE|UNCLEAR",
  "doc_confidence": "HIGH|LOW",
  "doc_evidence": "exact quote from the PDF text, or NOT FOUND"
}

Decision rules:
- Use only the provided PDF text.
- doc_evidence must be a verbatim quote copied from the PDF text.
- If direct support is missing, use NOT FOUND and LOW confidence.
- Prefer explicit document-label cues from the PDF itself, such as "review", "editorial",
"supplementary information", "conference", "abstract", "proceedings", "poster", "meeting",
"news", or "commentary".

Document type:
- MAIN_ARTICLE: the primary article itself
- REVIEW: review, survey, perspective, overview, opinion, or meta-analysis
- EDITORIAL: editorial, preface, commentary, special-issue introduction
- CORRECTION: erratum, corrigendum, retraction, expression of concern, correction notice
- SUPPLEMENT: supplementary information, appendix, supporting information, supplement PDF,
peer-review file, reviewer comments, decision letter, author response, reporting summary, or
other publication-adjacent file that is not the primary article
- NON_ARTICLE: conference abstract, proceedings paper, meeting supplement item, poster summary,
news item, or other non-primary-article PDF. Use this instead of MAIN_ARTICLE when the PDF is
clearly a conference/proceedings/meeting/news artifact.
- UNCLEAR: cannot determine reliably
```

##### S4.1.2 Panel-design sub-prompt (`PANEL_DESIGN_PROMPT`)

```
You are classifying how a scientific paper designed the gene/probe panel for a targeted or in
situ spatial transcriptomics experiment (e.g. MERFISH, seqFISH, seqFISH+, Xenium, MERSCOPE,
CosMx, Molecular Cartography / Resolve, STARmap, HybISS, osmFISH, EEL FISH). Use ONLY the
provided full extracted PDF text.

Return one JSON object only with this exact schema:
{
  "panel_design": "VENDOR|MANUAL-PANEL|TOOL|VENDOR-MANUAL|HYBRID-TOOL|UNCLEAR",
  "panel_confidence": "HIGH|LOW",
  "uses_vendor_panel": "YES|NO|UNCLEAR",
  "uses_manual_additions": "YES|NO|UNCLEAR",
  "uses_computational_tool": "YES|NO|UNCLEAR",
  "vendor_or_platform": "string naming vendor/platform if explicit, or NOT FOUND",
  "tool_names": "pipe-separated tool names if explicit, or NOT FOUND",
  "evidence_classification": "exact quote from the PDF text, or NOT FOUND",
  "evidence_vendor": "exact quote from the PDF text, or NOT FOUND",
  "evidence_manual": "exact quote from the PDF text, or NOT FOUND",
  "evidence_tool": "exact quote from the PDF text, or NOT FOUND"
}

Prior search hints from citation-screening tool-specific classification:
{prior_context}

Decision rules:
- Use only the provided PDF text for the final answer. The prior hints are only to help you
locate relevant passages.
- All evidence fields must be verbatim quotes copied from the PDF text.
- If direct support is missing, use NOT FOUND and LOW confidence.
- If the paper mentions a computational panel-selection tool but does not make clear how that
tool contributed to the final panel, prefer UNCLEAR.
- Count a tool as TOOL only if the paper explicitly uses one of the 18 review tools below to
select the final spatial panel/probe set.
- Do NOT count tool names that appear only in background, related work, discussion,
benchmarking, comparisons, references, citations, supplementary tables, or future-work
statements.
- Prefer direct workflow statements from Methods/Results over generic mentions in the
Introduction/Discussion.
- If a named tool is mentioned but the final panel appears to have been chosen manually from
marker lists, literature, DE outputs, rankings, or mutual-information scores, label MANUAL-
PANEL unless the paper explicitly says one of the 18 review tools selected the final panel.
- Differential-expression-based, ranking-based, mutual-information-based, or similar heuristic
gene lists are MANUAL-PANEL unless one of the 18 review tools explicitly selected the final
panel.

Label definitions:
- VENDOR = the paper uses a vendor pre-designed panel (e.g. Xenium Human Brain Panel, Xenium
Multi-Tissue and Cancer Panel, MERSCOPE Human Immuno-Oncology Panel, CosMx 1000-Plex Human
Universal Cell Characterization Panel), OR designs a custom panel using ONLY the vendor's in-
platform designer (Xenium Panel Designer, MERSCOPE Panel Designer, CosMx Custom Panel
workflow), with no reported manual additions and no external panel-selection tool.
- MANUAL-PANEL = the final panel is curated by hand from known markers, literature, prior
biology, expert choice, or differential-expression lists produced by general single-cell
pipelines (Seurat FindMarkers / FindAllMarkers, scanpy rank_genes_groups, limma, edgeR, DESeq2,
MAST, Wilcoxon, ROC, t-test) — and NO named panel-selection tool is used.
- TOOL = the final panel was designed using at least one of the 18 review tools listed below,
and the paper says one of those tools selected the final panel. Using two or more of those
review tools together still counts as TOOL (not HYBRID-TOOL).
- VENDOR-MANUAL = the final panel combines a vendor pre-designed panel with explicit
manual/custom gene additions curated, adapted, supplemented, or extended outside the vendor
workflow, with no computational panel-selection tool.
- HYBRID-TOOL = the final panel combines an explicit computational panel-selection tool with
another non-tool branch (vendor pre-designed panel, manual curation, or literature markers).
- UNCLEAR = the paper does not provide direct evidence to choose among the above.
```

```
Canonical 18 review tools — if the paper states one of these selected the final spatial panel,
the label is TOOL (or HYBRID-TOOL if combined with a non-tool branch):
  COSG, geneBasis, geneCover, gpsFISH, NS-Forest v2, NS-Forest v4.0, panelScope, PERSIST,
RankCorr, ReconST, SCMER, SMaSH, Spapros, scGeneFit, scGIST, scGPD, scMAGS, scPNMF.

NOT panel-selection tools — these DO NOT trigger the TOOL label even when used heavily:
- General single-cell DE / marker-finding software: Seurat (FindMarkers, FindAllMarkers),
scanpy (rank_genes_groups), limma, edgeR, DESeq2, MAST, Wilcoxon / ROC / t-test. Authors hand-
picking the final panel from their output is MANUAL-PANEL.
- Clustering / integration / batch correction only: Harmony, scVI (integration mode), Liger,
BBKNN.
- Vendor in-platform panel designers (Xenium Panel Designer, MERSCOPE Panel Designer, CosMx
Custom Panel). These count as VENDOR even when fed a user gene list, unless the user also added
custom genes outside the vendor's workflow (then VENDOR-MANUAL).

Worked examples:

Example A — VENDOR
Passage: "We profiled 5 human brain sections using the Xenium Human Brain Panel (10x Genomics)
comprising 266 genes, without modification."
→ panel_design=VENDOR, uses_vendor_panel=YES, uses_manual_additions=NO,
uses_computational_tool=NO, vendor_or_platform="Xenium Human Brain Panel", tool_names=NOT
FOUND.

Example B — VENDOR-MANUAL
Passage: "We used the Xenium Human Brain Panel (266 genes) as a base and supplemented it with
20 custom probes targeting disease-associated markers identified from our scRNA-seq Seurat
FindMarkers analysis."
→ panel_design=VENDOR-MANUAL, uses_vendor_panel=YES, uses_manual_additions=YES,
uses_computational_tool=NO, vendor_or_platform="Xenium Human Brain Panel", tool_names=NOT
FOUND.

Example C — MANUAL-PANEL
Passage: "We selected 300 genes for the MERFISH panel from literature-curated markers of
excitatory and inhibitory neurons plus the top 50 DE genes per cluster from Seurat
FindAllMarkers."
→ panel_design=MANUAL-PANEL, uses_vendor_panel=NO, uses_manual_additions=YES,
uses_computational_tool=NO, tool_names=NOT FOUND.

Example D — TOOL
Passage: "A 140-gene MERFISH panel was designed with Spapros (Kuemmerle et al., 2024) using our
reference scRNA-seq atlas as input."
→ panel_design=TOOL, uses_vendor_panel=NO, uses_manual_additions=NO,
uses_computational_tool=YES, tool_names="Spapros".

Example E — HYBRID-TOOL
Passage: "An initial 80-gene MERFISH set was computed with geneBasis on our reference atlas,
then supplemented with 20 tissue-specific markers chosen by expert curation to reach 100
probes."
→ panel_design=HYBRID-TOOL, uses_vendor_panel=NO, uses_manual_additions=YES,
uses_computational_tool=YES, tool_names="geneBasis".

Example F — UNCLEAR
Passage: "A custom 200-gene panel was designed to capture major cell types in the developing
cortex."
→ panel_design=UNCLEAR, panel_confidence=LOW (no mention of vendor, named tool, or curation
strategy).

Example G — VENDOR via vendor designer, NOT TOOL
Passage: "We designed a custom 480-gene panel using the 10x Genomics Xenium Panel Designer web
tool based on our cell-type markers."
→ panel_design=VENDOR (vendor in-platform designer does not count as an external panel-
selection tool), uses_vendor_panel=YES, uses_computational_tool=NO, vendor_or_platform="Xenium
Panel Designer".
```

```
Example H — cited tool only, still UNCLEAR
Passage: "Candidate markers were selected from prior literature and differential-expression
analysis. We note that Spapros and SMaSH are promising future approaches for probe-set design."
→ panel_design=UNCLEAR (or MANUAL-PANEL if the manual curation workflow is explicit).
Mentioning tools in discussion/future-work does not mean the authors used them for the final
panel.
```

##### S4.1.3 Paper-profile sub-prompt, full text (`PROFILE_PROMPT`)

```
You are classifying the primary type and panel-design relevance of a scientific PDF using ONLY
the provided full extracted PDF text.

Return one JSON object only with this exact schema:
{
  "paper_type": "EXP|COMP|OTHER|UNCLEAR",
  "spatial": "YES|NO|UNCLEAR",
  "panel_scope": "TARGETED|NOT_TARGETED|UNCLEAR",
  "profile_confidence": "HIGH|LOW",
  "evidence_type": "exact quote from the PDF text, or NOT FOUND",
  "evidence_spatial": "exact quote from the PDF text, or NOT FOUND",
  "evidence_scope": "exact quote from the PDF text, or NOT FOUND"
}

Decision rules:
- Use only the provided PDF text.
- All evidence fields must be verbatim quotes copied from the PDF text.
- If direct support is missing, use NOT FOUND and LOW confidence.
- Classify the paper by its primary contribution, not by every method or platform mentioned
anywhere in the PDF.
- Mentions of platforms or analyses in background, related work, comparisons, citations, or
future directions do NOT determine the label.
- If a paper introduces a new computational method/software/benchmark as its main contribution,
classify paper_type=COMP even if it validates the method on real biological data.
- If a paper's main contribution is a biological or experimental study that uses standard
downstream analysis, classify paper_type=EXP even if it contains substantial computation.

paper_type:
- EXP = primarily reports new wet-lab experimental work, assay development, biological
findings, or real profiling of samples
- COMP = primarily reports a new computational method, software tool, benchmarking framework,
or algorithmic study
- OTHER = protocol, resource, dataset note, book chapter, guideline, or similar
- UNCLEAR = cannot determine reliably

spatial:
- YES = the paper explicitly reports spatial transcriptomics or in situ transcriptomics
experiments (including re-analysis of ST data as a primary contribution)
- NO = the paper is non-spatial (bulk RNA-seq only, scRNA-seq only, proteomics only,
chromatin/ATAC only, imaging-only without transcriptomics)

panel_scope:
Platform-name cues are authoritative unless the paper contradicts them.
- TARGETED (finite-gene probe panel): MERFISH, seqFISH, seqFISH+, Xenium, MERSCOPE, CosMx,
Molecular Cartography (Resolve), STARmap, STARmap PLUS, Targeted ExSeq, HybISS, osmFISH,
Baristaseq, EEL FISH, Vizgen MERSCOPE, RNAscope HiPlex, ISS (in situ sequencing). Using a fixed
commercial panel unchanged still counts as TARGETED.
- NOT_TARGETED (whole-transcriptome spatial, no probe panel selection): Visium, Visium HD,
Slide-seq, Slide-seqV2, 10x Genomics Spatial Gene Expression, Stereo-seq, HDST, DBiT-seq, GeoMx
WTA, Nanostring GeoMx Whole Transcriptome Atlas.
- UNCLEAR = the PDF text does not distinguish between targeted and whole-transcriptome
reliably.

Few-shot:
Passage: "We performed 10x Genomics Visium spatial transcriptomics on mouse brain sections to
```

```
profile whole-transcriptome expression across tissue coordinates."
→ spatial=YES, panel_scope=NOT_TARGETED.

Passage: "MERFISH imaging of a custom 300-gene panel was performed on mouse primary visual
cortex."
→ spatial=YES, panel_scope=TARGETED.

Passage: "We developed and benchmarked six cell-type deconvolution algorithms on simulated
Visium data."
→ paper_type=COMP, spatial=YES, panel_scope=NOT_TARGETED.

Passage: "We applied Xenium to profile inflamed colon tissue and used standard clustering and
differential expression to identify disease-associated epithelial states."
→ paper_type=EXP, spatial=YES, panel_scope=TARGETED.

Passage: "We introduce a new graph neural network for cell-type deconvolution and evaluate it
on Visium and MERFISH benchmark datasets."
→ paper_type=COMP. Determine panel_scope from the main analyzed modality if explicit; otherwise
panel_scope=UNCLEAR.
```

##### S4.1.4 Paper-profile sub-prompt, abstract only (`ABSTRACT_PROFILE_PROMPT`)

```
You are classifying the primary type, spatial status, panel scope, and (when explicitly stated)
panel-design approach of a scientific paper using ONLY the provided title and abstract text.

Return one JSON object only with this exact schema:
{
  "paper_type": "EXP|COMP|OTHER|UNCLEAR",
  "spatial": "YES|NO|UNCLEAR",
  "panel_scope": "TARGETED|NOT_TARGETED|UNCLEAR",
  "panel_design_hint": "VENDOR|MANUAL-PANEL|TOOL|VENDOR-MANUAL|HYBRID-TOOL|UNCLEAR",
  "panel_design_hint_confidence": "HIGH|LOW",
  "profile_confidence": "HIGH|LOW"
}

Decision rules:
- Use only the provided title and abstract text.
- If the abstract is short, missing, or ambiguous, assign LOW confidence.
- Only assign HIGH confidence if the classification is unambiguous from the available text.
- panel_design_hint should be HIGH only if the abstract explicitly names a vendor panel, a
named panel-selection tool, or a clear manual curation strategy. Otherwise UNCLEAR / LOW.

paper_type:
- EXP = primarily new wet-lab / biological / sample-profiling work
- COMP = primarily a new computational method, tool, benchmark, or algorithm
- OTHER = protocol, resource, dataset note, book chapter, guideline
- UNCLEAR = cannot determine reliably

spatial:
- YES = abstract mentions spatial transcriptomics, in situ transcriptomics, or a named ST
platform (MERFISH, seqFISH, seqFISH+, Xenium, MERSCOPE, CosMx, Visium, Visium HD, Slide-seq,
Slide-seqV2, Stereo-seq, Molecular Cartography / Resolve, STARmap, STARmap PLUS, HybISS,
osmFISH, HDST, GeoMx WTA, DBiT-seq, EEL FISH, Targeted ExSeq, Baristaseq, ISS)
- NO = clearly non-spatial (bulk RNA-seq only, scRNA-seq only, proteomics only, imaging-only
without transcriptomics)

panel_scope (platform-name heuristics are authoritative unless the abstract contradicts):
- TARGETED: MERFISH, seqFISH, seqFISH+, Xenium, MERSCOPE, CosMx, Molecular Cartography
(Resolve), STARmap, STARmap PLUS, Targeted ExSeq, HybISS, osmFISH, Baristaseq, EEL FISH,
RNAscope HiPlex, ISS
- NOT_TARGETED: Visium, Visium HD, Slide-seq, Slide-seqV2, 10x Spatial Gene Expression, Stereo-
seq, HDST, DBiT-seq, GeoMx WTA
- UNCLEAR = abstract only says "spatial transcriptomics" without a platform cue
```

```
panel_design_hint (use ONLY when the abstract contains an explicit named cue; otherwise UNCLEAR
/ LOW):
- VENDOR: abstract names a vendor pre-designed panel (e.g. "Xenium Human Brain Panel",
"MERSCOPE Human Immuno-Oncology Panel", "CosMx 1000-Plex Human Universal Cell Characterization
Panel"), OR names the vendor's in-platform designer (Xenium Panel Designer, MERSCOPE Panel
Designer)
- TOOL: abstract names one of the 18 review tools: COSG, geneBasis, geneCover, gpsFISH, NS-
Forest v2, NS-Forest v4.0, panelScope, PERSIST, RankCorr, ReconST, SCMER, SMaSH, Spapros,
scGeneFit, scGIST, scGPD, scMAGS, scPNMF — used as a panel selector
- MANUAL-PANEL: abstract explicitly says the panel was curated from markers / literature / DE
results / rankings / mutual-information scores without naming one of the 18 review tools
- VENDOR-MANUAL: abstract names a vendor panel plus explicit manual additions
- HYBRID-TOOL: abstract names a panel-selection tool plus manual or vendor additions
- UNCLEAR: default — the abstract does not explicitly describe panel design

Counter-examples for panel_design_hint (do NOT assign TOOL for these):
- "We identified markers with Seurat FindMarkers" → not a panel-selection tool (MANUAL-PANEL if
the panel is explicitly described, otherwise UNCLEAR).
- "We used scanpy rank_genes_groups" → MANUAL-PANEL / UNCLEAR.
- "Cells were clustered with Harmony and scVI" → integration/clustering only; UNCLEAR for
panel_design_hint.

Few-shot:
Title/abstract: "Xenium spatial mapping of Alzheimer plaques using the 10x Xenium Human Brain
Panel reveals disease-associated microglia states."
→ paper_type=EXP, spatial=YES, panel_scope=TARGETED, panel_design_hint=VENDOR,
panel_design_hint_confidence=HIGH, profile_confidence=HIGH.

Title/abstract: "A 300-gene MERFISH panel designed with Spapros uncovers rare cortical
interneuron subtypes in mouse cortex."
→ paper_type=EXP, spatial=YES, panel_scope=TARGETED, panel_design_hint=TOOL,
panel_design_hint_confidence=HIGH, profile_confidence=HIGH.

Title/abstract: "Visium profiling of mouse tumor microenvironments reveals stromal
heterogeneity."
→ paper_type=EXP, spatial=YES, panel_scope=NOT_TARGETED, panel_design_hint=UNCLEAR,
panel_design_hint_confidence=LOW, profile_confidence=HIGH.

Title/abstract: "We develop a deconvolution benchmark on simulated Visium data across 12
algorithms."
→ paper_type=COMP, spatial=YES, panel_scope=NOT_TARGETED, panel_design_hint=UNCLEAR,
profile_confidence=HIGH.

Title/abstract: "Single-cell RNA-seq of human pancreatic islets identifies a rare delta-cell
subpopulation."
→ paper_type=EXP, spatial=NO, panel_scope=UNCLEAR, panel_design_hint=UNCLEAR,
profile_confidence=HIGH.
```

#### S4.2 GPT-5.4 panel-design audit prompt

Each Gemma4 panel-design classification was packaged into a self-contained JSON audit packet (one packet per corpus paper). Each packet included the paper metadata, the Gemma4 outputs (label + verbatim evidence quotes + confidence), citation-screening priors where available, risk flags, and the `audit_instructions` block reproduced below. GPT-5.4 returned one JSON object per packet, matching the output schema reproduced after the instruction block. ### S4.2.1 Audit instructions

```
{
  "classification_set": [
    "VENDOR",
    "MANUAL-PANEL",
    "TOOL",
    "VENDOR-MANUAL",
```

```
    "HYBRID-TOOL",
    "UNCLEAR",
    "FILTERED"
  ],
  "key_rules": [
    "Do not count probe, barcode, oligo, primer, decoding, segmentation, registration, or downstream analysis tools as gene-panel selection tools.",
    "Only these 18 review tools count as TOOL when they explicitly select the final panel: COSG, geneBasis, geneCover, gpsFISH, NS-Forest v2, NS-Forest v4.0, panelScope, PERSIST, RankCorr, ReconST, SCMER, SMaSH, Spapros, scGeneFit, scGIST, scGPD, scMAGS, and scPNMF.",
    "Treat NS-Forest_v4.0 as the same tool as NS-Forest v4.0.",
    "Differential-expression-based, ranking-based, mutual-information-based, or other generic heuristic gene lists are MANUAL-PANEL unless one of the 18 review tools above is explicitly used to select the final panel.",
    "Do not use FILTERED solely because paper_type=COMP.",
    "If a targeted spatial or in situ paper has concrete panel-design evidence, classify the panel source even when paper_type=COMP.",
    "If a commercial/vendor panel was used, adapted, converted, or used as a design source, set vendor use to YES.",
    "If a vendor panel was manually curated, adapted, supplemented, or extended with custom genes, use VENDOR-MANUAL rather than pure VENDOR or MANUAL-PANEL.",
    "If evidence is insufficient to determine how target genes were selected, use NEEDS_MANUAL rather than inventing a rationale."
  ],
  "task": "Audit Gemma's final panel-design classification for this one paper. Use only this packet unless an external PDF snippet is explicitly provided."
}
```

##### S4.2.2 Audit output schema

```
{
  "type": "object",
  "required": [
    "audit_decision",
    "confidence",
    "Rec. classification",
    "paper_key",
    "doi",
    "error_type",
    "reason",
    "evidence_used"
  ],
  "properties": {
    "audit_decision": {
      "enum": [
        "AGREE",
        "DISAGREE",
        "NEEDS_MANUAL"
      ]
    },
    "confidence": {
      "enum": [
        "high",
        "medium",
        "low"
      ]
    },
    "Rec. classification": {
      "enum": [
        "VENDOR",
        "MANUAL-PANEL",
        "TOOL",
        "VENDOR-MANUAL",
        "HYBRID-TOOL",
        "UNCLEAR",
```

```
        "FILTERED"
      ]
    },
    "paper_key": {
      "type": "string"
    },
    "doi": {
      "type": "string"
    },
    "error_type": {
      "enum": [
        "none",
        "probe_design_confused",
        "vendor_source_missed",
        "comp_filter_too_aggressive",
        "panel_evidence_overruled_by_profile",
        "wrong_pdf",
        "pdf_unattainable",
        "insufficient_evidence",
        "other"
      ]
    },
    "reason": {
      "type": "string"
    },
    "evidence_used": {
      "type": "string"
    }
  }
}
```
