## Supplementary figures and images for "Lack of adoption of computational gene-panel-selection tools in targeted spatial transcriptomics"

### Fig_SUPP 1

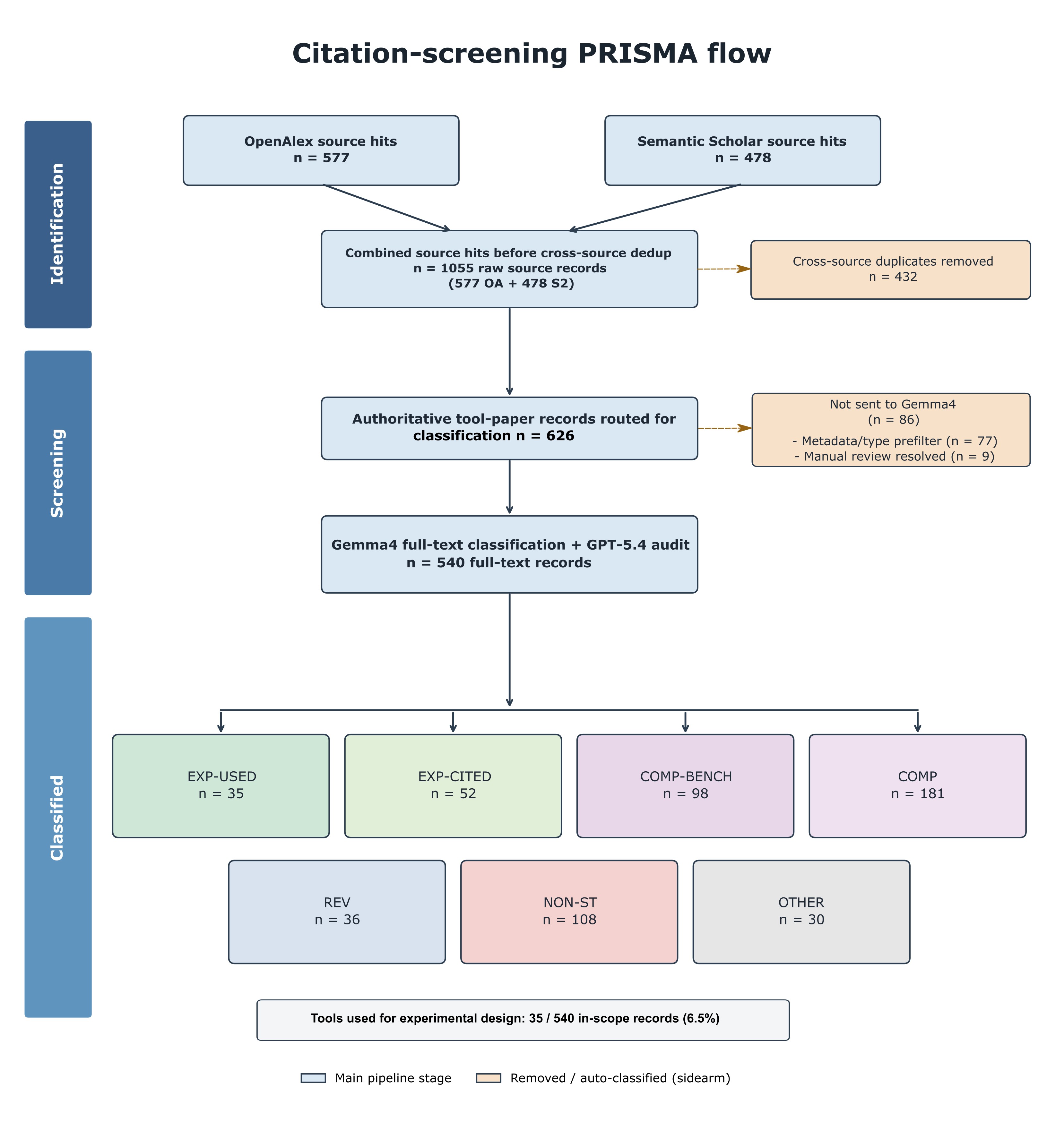

### Fig_SUPP 2

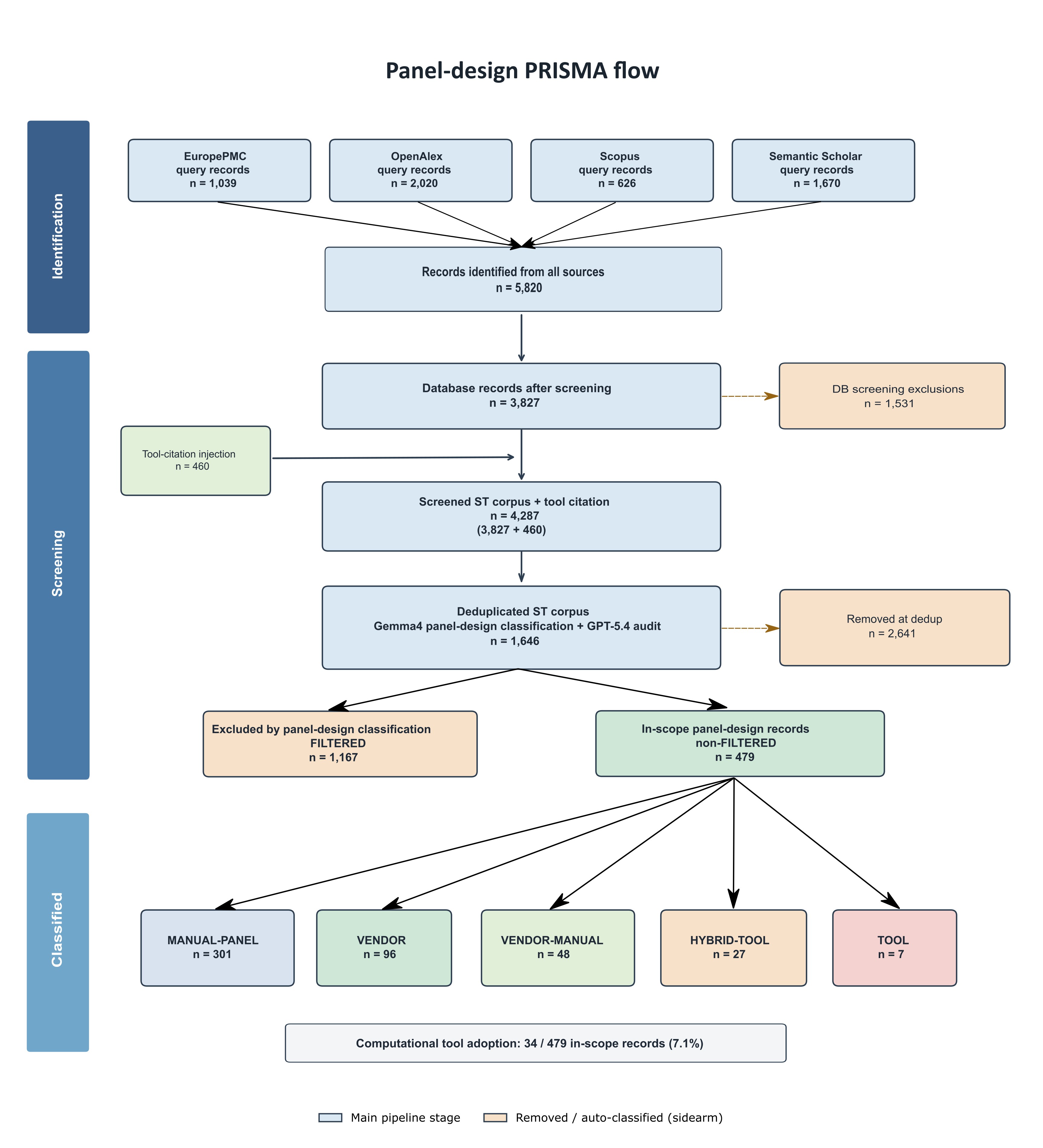
